# KIAA1841, a novel SANT and BTB domain-containing protein, inhibits class switch recombination

**DOI:** 10.1101/2020.08.12.247809

**Authors:** Simin Zheng, Allysia J. Matthews, Numa Rahman, Kayleigh Herrick-Reynolds, Jee Eun Choi, Emily Sible, Yan Kee Ng, Daniela Rhodes, Stephen J. Elledge, Bao Q. Vuong

## Abstract

Class switch recombination (CSR) enables B cells to produce different immunoglobulin isotypes and mount an effective immune response against pathogens. Timely resolution of CSR prevents damage due to an uncontrolled and prolonged immune response. While many positive regulators of CSR have been described, negative regulators of CSR are relatively unknown. Using a shRNA library screen in a mouse B cell line, we have identified the novel protein KIAA1841 (NM_027860) as a negative regulator of CSR. KIAA1841 is an uncharacterized protein of 82kD containing SANT and BTB domains. The BTB domain of KIAA1841 exhibited characteristic properties such as self-dimerization and interaction with co-repressor proteins. Overexpression of KIAA1841 inhibited CSR in primary mouse splenic B cells, and inhibition of CSR is dependent on the BTB domain while the SANT domain is largely dispensable. Thus, we have identified a new member of the BTB family that serves as a negative regulator of CSR.

## Introduction

Immunoglobulin (Ig) production by B cells is an essential component of the immune response against pathogens and malignancies. Upon encounter with antigen, mature B cells are activated and undergo class switch recombination (CSR)^1-3^. This process changes the isotype of the Ig produced from IgM to IgG, IgE or IgA, and thus couples Ig specificity, which is determined by the variable domains, to a broad spectrum of effector functions for complete humoral immunity. Through a DNA rearrangement reaction, CSR replaces the default constant region Cµ in the Ig heavy chain (IgH) locus with one of the downstream constant regions (Cγ, Cε or Cα) that codes for the new isotype. Following stimulation for CSR, B cells express the enzyme activation-induced cytidine deaminase (AID), which localizes to repetitive GC-rich switch (S) regions that precede each constant region^4,5^. AID deamination of the S regions initiates the formation of DNA double-strand breaks (DSBs) in Sµ and a downstream acceptor S region to delete the intervening DNA and rearrange the IgH locus to express a different Ig isotype.

Following AID deamination of S region DNA, two complementary pathways, base excision repair (BER) and mismatch repair (MMR), generate the DSBs that are necessary for CSR. In BER, uracil DNA glycosylase (UNG) removes the AID-generated uracil base to create an abasic site, which is processed into a DNA break by apurinic/apyrimidinic endonucleases (APE)^6,7^. During MMR, a complex of proteins, which includes PMS1 homolog 2 (PMS2) and Exonuclease I (EXO1), converts AID-induced U:G mismatches into DSBs^8^. Proteins of the non-homologous end-joining pathway, such as Ku70, Ku80, X-ray repair cross complementing 4 (XRCC4) and Ligase IV (Lig4), ligate DSBs in donor and acceptor S regions together to complete CSR^1^. Similarly, other factors positively regulate CSR by modulating AID expression, targeting AID to S regions, generating DSBs, and repairing DSBs^1,3,9-12^.

Chromatin architecture also influences CSR and proteins that regulate epigenetic states have been reported to promote CSR. For example, loss of the PTIP protein decreased histone methylation and chromatin accessibility, altered germline transcription, and resulted in defects in CSR^13,14^. Phosphorylation and acetylation of histone H3 modulates AID recruitment to S regions by 14-3-3 and conditional inactivation of methyltransferases (*Suv4-20h1* and *Suv4-20h2*), which modify histone H4, in B cells significantly impairs CSR^15,16^.

In contrast, few negative regulators of CSR have been described. Previously characterized negative regulators include the poly(ADP) ribose polymerase Parp3^17^ and the aryl hydrocarbon receptor AhR^18^, with many more yet to be uncovered. Parp3 prevents excessive accumulation of AID at S regions^17^, while AhR limits AID expression^18^. These negative regulators of CSR fine tune humoral immune responses and AID-mediated DNA damage, which if dysregulated can lead to autoimmunity, genomic instability and lymphomagenesis^19-23^. To identify novel negative regulators of CSR, we performed an shRNA library screen in the CH12 mouse B cell line. Here, we report the identification and characterization of a novel SANT- and BTB-domain containing protein (KIAA1841) as a negative regulator of CSR.

## Results

### Identification of KIAA1841 as a negative regulator of CSR

To identify novel negative regulators of CSR, we performed an shRNA screen using the CH12 mouse B lymphoma cell line (**Fig. 1A**). CH12 cells are a well-established model system for the study of CSR, as they can be induced to class switch from IgM to IgA in culture when stimulated with anti-CD40, interleukin 4 (IL-4) and transforming growth factor β (TGF-β), which is henceforth referred to as CIT^5,18,24-27^. CH12 cells were infected with a lentiviral shRNA library comprised of 67,676 shRNAs targeting 28,801 genes in the mouse genome^28,29^. Transduced cells were selected with puromycin and stimulated with CIT to undergo CSR to IgA. IgA- and IgA+ populations were sorted and harvested for genomic DNA. The genomic DNA was used as the template in PCRs to generate fluorescence-labeled half-hairpin amplicons; IgA-amplicons were labeled with Cy5 while IgA+ amplicons were labeled with Cy3. The relative abundance of individual shRNAs was determined by competitive hybridization to microarray (**Fig. 1A**). As depletion of negative regulators would likely promote CSR, we expected shRNAs that targeted them to be enriched in the IgA+ population as compared to the IgA-population. Genes with shRNAs showing at least 2-fold enrichment in IgA+ compared to IgA-samples, i.e. log^2^(Cy3/Cy5)>1, were identified as negative regulators (**Supplementary Table 1**). As controls in the screen, known positive regulators of CSR resulted in negative log^2^(Cy3/Cy5) values, indicating that shRNAs for these genes are enriched in the IgA-population and their knockdown prevented CSR (**Supplementary Table 2**). For instance, log^2^(Cy3/Cy5) of AID = −1.42, demonstrating the validity of the shRNA screen (**Supplementary Table 2**).

**Figure 1:**
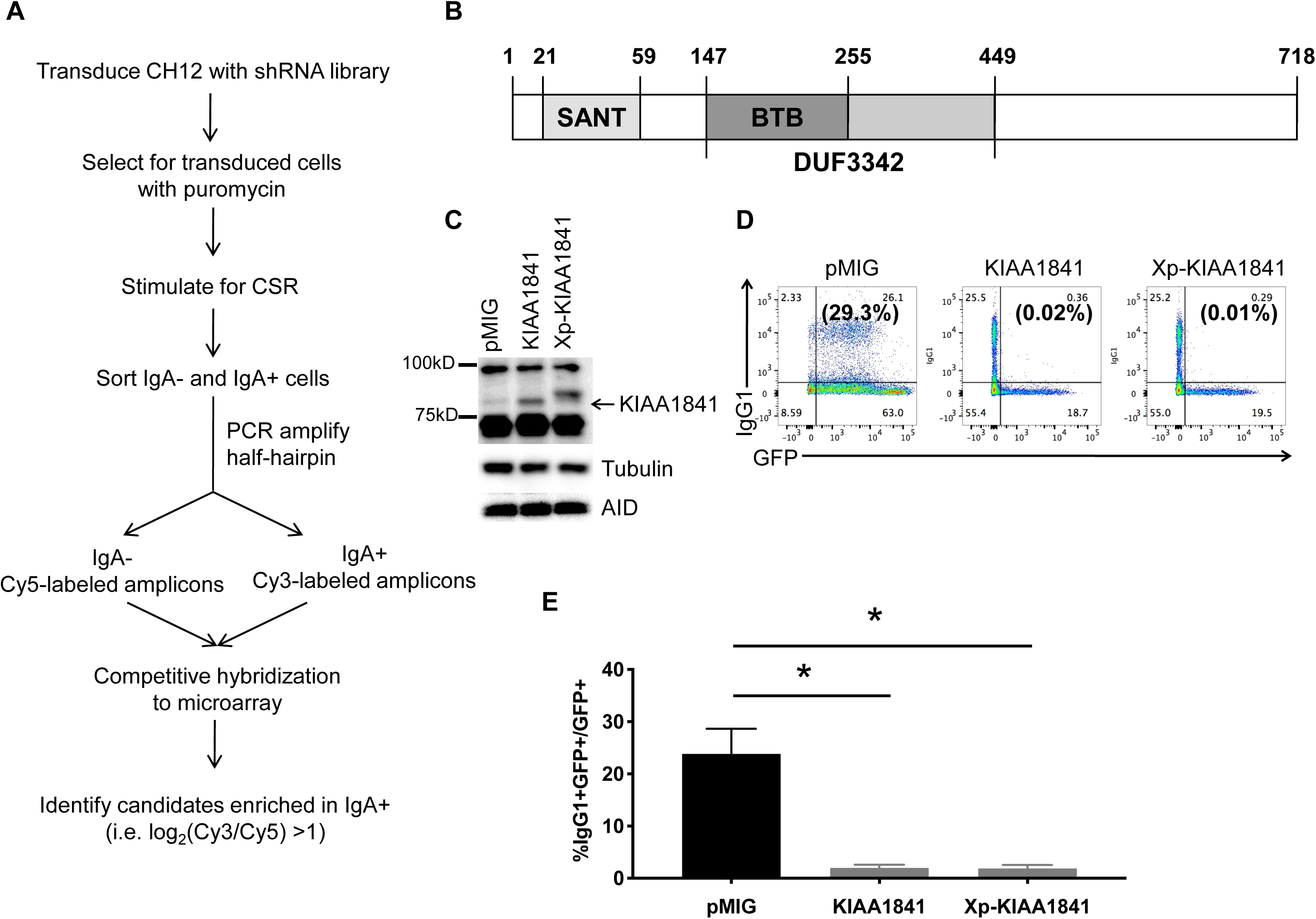
shRNA screen identifies KIAA1841 as a negative regulator of CSR. **(A)** Workflow for the shRNA screen. CH12 cells were infected with the lentiviral shRNA library, selected with puromycin and stimulated to undergo CSR to IgA. IgA- and IgA+ cells were sorted and differentially labeled half-hairpin amplicons were generated by PCR using genomic DNA (Cy5 or Cy3 respectively). Relative abundance of individual shRNAs were determined by competitive hybridization to microarray using labeled half-hairpin amplicons. Potential negative regulators of CSR were identified as genes with at least 2-fold enrichment of targeting shRNA in the IgA+ population as compared to the IgA-population (log2(Cy3/Cy5)>1). **(B)** Schematic representation of the domains of KIAA1841. **(C-E)** Expression of KIAA1841 inhibits CSR. Splenic B cells were isolated from wild-type mice, stimulated for CSR with LPS+IL4, and transduced with retroviral vector control (pMIG) or vectors expressing untagged or Xpress-tagged (Xp) KIAA1841. **(C)** Overexpression of KIAA1841 and expression of AID were determined by immunoblot. Tubulin was used as a loading control. **(D)** CSR to IgG1 among the transduced cell population was determined by flow cytometry. A representative experiment is shown. The numbers in the corners of each plot indicate the percentage of cells in each quadrant while the numbers in parentheses indicate the percentage of IgG1+ cells within the GFP+ gate. **(E)** The mean %IgG1+ within the GFP+ gate from three independent experiments +/-SD is shown. * p < 0.05, two-tailed paired student’s t-test.

The shRNA screen identified an uncharacterized mRNA, NM_027860 (RIKEN cDNA 0610010F05 gene, 0610010F05Rik), as a candidate negative regulator of CSR (log^2^(Cy3/Cy5) = 3.09). NM_027860 encodes for a well-conserved protein KIAA1841 (**Supplementary Fig. 1**) that is expressed broadly in many cell types^30,31^. A NCBI genome database search predicts that invertebrates (e.g. Drosophila melanogaster CG6761), vertebrates (e.g. Homo sapiens, Danio rerio), and single cell organisms (e.g. Trypanosoma grayi, Ruminococcus albus 7) express an ortholog of KIAA1841. The mouse KIAA1841 gene is located on chromosome 11 and consists of 28 exons. While a potential splice isoform missing exons 3, 4, and 5, which encodes the first 146 amino acids, has been reported^32,33^, we were unable to detect this isoform in mouse splenic B cells (data not shown). We cloned the full-length KIAA1841 cDNA from stimulated mouse splenic B cells. Full-length KIAA1841 comprises of 718 amino acids with a predicted molecular weight of 82 kD (**Fig. 1B**). Amino acid sequence alignment by BLAST and structure prediction by Phyre2^34^ further revealed that this protein has a putative SANT domain (’switching-defective protein 3 (Swi3), adaptor 2 (Ada2), nuclear receptor co-repressor (N-CoR), transcription factor (TF)IIIB’) encoded by amino acids 21-59, and a BTB domain (‘broad-complex, tramtrack and bric-a-brac’) encoded by amino acids 147-255. The KIAA1841 BTB domain overlaps a conserved domain of unknown function (DUF3342, pfam11822), which includes amino acids 147-449 (**Fig. 1B**).

To validate the identification of KIAA1841 as a negative regulator of CSR, we retrovirally overexpressed untagged and Xpress epitope-tagged KIAA1841 in LPS+IL4-stimulated mouse splenic B cells (**Fig. 1C**). Both the untagged and Xpress-tagged KIAA1841 significantly reduced CSR to IgG1 (**Fig. 1D, 1E**), reinforcing the role of KIAA1841 as a negative regulator of CSR. Overexpression of KIAA1841 did not affect germline transcription (**Supplementary Fig. 2**), AID mRNA and protein expression (**Supplementary Fig. 2**; **Fig. 1C**), or B cell proliferation (**Supplementary Fig. 3**). Thus, overexpression of KIAA1841 inhibits CSR without affecting fundamental processes that are required for CSR.

### Putative BTB domain of KIAA1841 mediates dimerization

We next investigated the function of the putative BTB domain of KIAA1841. Many cellular processes require BTB domain-containing proteins, including transcription and protein degradation^35^. In the immune system, several developmental pathways are controlled by BTB-containing proteins. For instance, promyelocytic leukemia zinc finger (PLZF) controls the development of NKT cells^36,37^ and B cell lymphoma 6 (BCL6) is essential for the formation and function of follicular helper T cells (T^FH^) and germinal center B cells38,39. The BTB domain is a well-characterized structural motif that mediates dimerization^35^. The putative BTB domain of KIAA1841 identified by Phyre2 is well-conserved with the BTB domains of PLZF and BCL6 (**Fig. 2A**). As full-length KIAA1841 is poorly expressed and aggregated in *E*.*coli* (data not shown), a fragment containing the putative BTB domain, KIAA1841(BTB), was expressed as a His^6^-tagged recombinant protein and purified (**Fig. 2B**) before analysis on a size-exclusion chromatography column (**Fig. 2C**). Comparison with protein molecular weight standards separated on the column under the same conditions indicated that KIAA1841(BTB) migrates as a dimer (**Fig. 2C**). To further verify that KIAA1841(BTB) dimerizes, we crosslinked purified KIAA1841(BTB) with glutaraldehyde before analysis on a denaturing SDS-PAGE (**Fig. 2D**). When crosslinked, the 15kD fragment of KIAA1841(BTB) formed a species approximately 30kD in size, corresponding to the expected molecular weight of a dimer (**Fig. 2D**). Thus, KIAA1841 contains a BTB domain that dimerizes *in vitro*.

**Figure 2:**
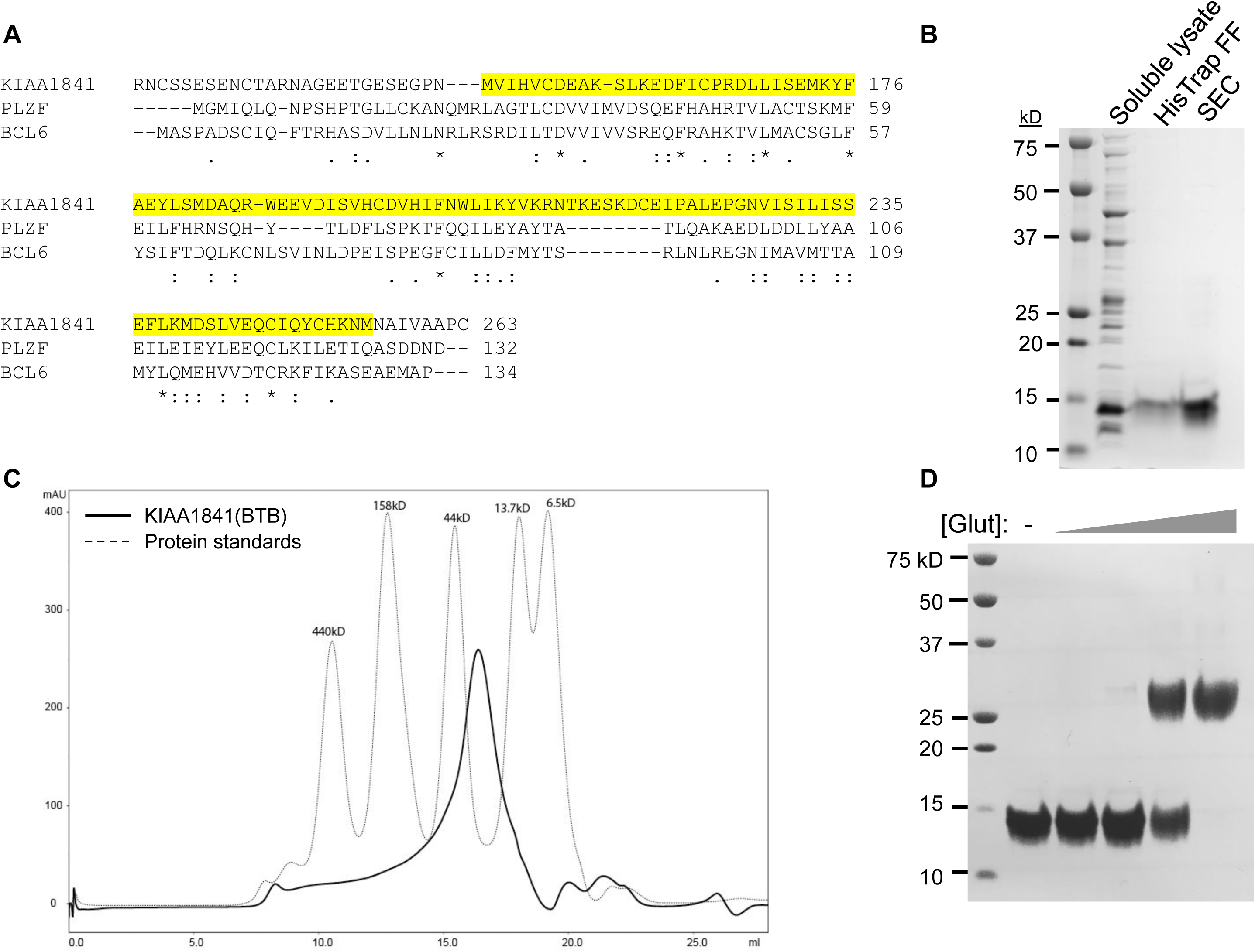
The BTB domain of KIAA1841 dimerizes *in vitro*. **(A)** Sequence alignment of the putative BTB domain of KIAA1841 with the BTB domains of PLZF and BCL6. Alignment was performed using ClustalOmega; *, conserved residue,:, strongly similar residues, ., weakly similar residues. The BTB domain of KIAA1841 (amino acids 147-255) that was predicted by Phyre2 protein fold recognition software is highlighted in yellow. Numbers indicate amino acid residues in each sequence. **(B**) Purification of recombinant the BTB domain of KIAA1841, KIAA1841(BTB). The predicted BTB domain of KIAA1841 (amino acids 144-261) was purified as a His6-tagged protein by HisTrap FF column and size exclusion chromatography (SEC). A Coomassie-stained gel is shown. The predicted size of His6-KIAA1841(BTB) is 15kD. **(C)** SEC analysis of purified KIAA1841(BTB). SEC was performed using Superdex-200 10/30 column. The chromatogram of molecular weight standards is indicated by the dotted line. **(D)** Glutaraldehyde cross-linking of purified KIAA1841(BTB). Purified KIAA1841(BTB) was treated with 5-fold increasing concentrations of glutaraldehyde (Glut), before analysis by SDS-PAGE and Coomassie stain.

To determine if the BTB domain of KIAA1841 promotes dimerization *in vivo*, dimerization assays were performed in 293T cells. Full-length flag-strepII-tagged wild-type (WT) or deletion mutants of KIAA1841, which lacked the BTB domain (▵BTB, amino acids 147-255), or helix 3 of the SANT domain^40^ (▵SANT, amino acids 48-59) (**Supplementary Figure 4**), were co-expressed with N-terminal GFP-tagged WT KIAA1841 (GFP-KIAA1841) in 293T cells (**Fig. 3**). Flag-Strep-KIAA1841(WT) and the KIAA1841 deletion mutant proteins were isolated with Strep-Tactin XT beads and examined for their ability to interact with GFP-KIAA1841 (**Fig. 3**). Unlike WT and ▵SANT, the ▵BTB mutant protein showed reduced binding to GFP-KIAA1841 (**Fig. 3**), demonstrating that the BTB domain mediates dimerization of KIAA1841 *in vivo*. The binding of the ▵SANT mutant protein to GFP-KIAA1841 indicates this domain is not required for dimerization of KIAA1841 (**Fig. 3**), and its function in KIAA1841 remains to be determined.

**Figure 3:**
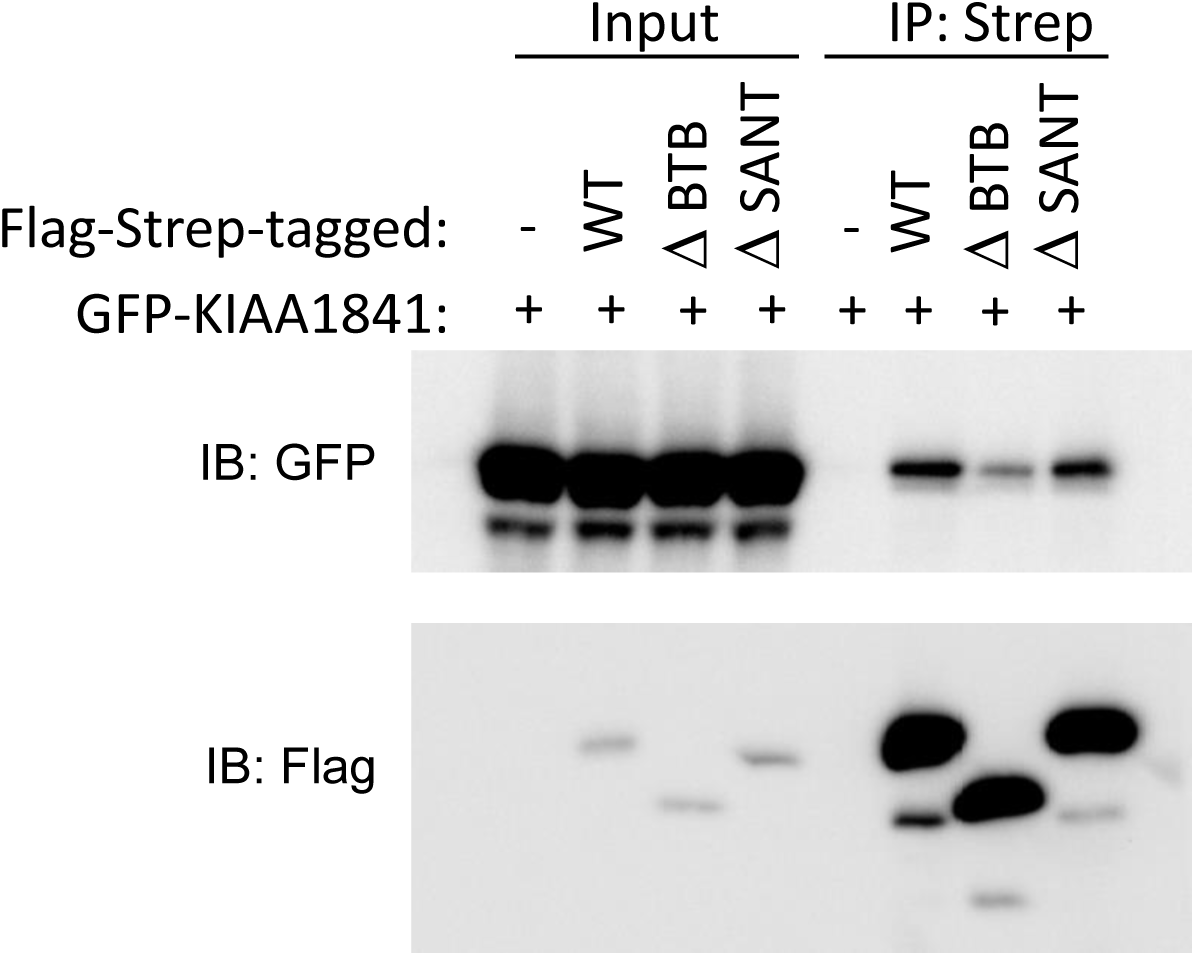
The BTB domain of KIAA1841 facilitates dimerization *in vivo*. 293T cells were co-transfected with GFP-tagged KIAA1841 and either Flag-Strep-tagged wild-type (WT), ▵BTB or ▵SANT KIAA1841. Flag-Strep-tagged proteins were pulled down using Strep-Tactin XT beads and bound proteins were analyzed by immunoblot using anti-GFP and anti-Flag antibodies. The results are representative of three independent pull-down experiments.

### KIAA1841 interacts with co-repressors through its putative BTB domain

BTB-containing proteins commonly function by interacting with co-repressors, such as histone deacetylases (HDACs), nuclear co-repressors (N-CoR) and silencing mediator of retinoic acid and thyroid hormone receptor (SMRT), via their BTB domain^35^. To determine if the BTB domain of KIAA1841 can bind to co-repressors, we purified recombinant glutathione-S-transferase (GST) GST-tagged HDAC1 and a fragment of SMRT previously reported to interact with the BTB domain of PLZF^41^ (**Fig. 4A**). These proteins were then used in binding assays with purified His^6^-tagged KIAA1841(BTB), where a His^6^-tagged BTB domain of PLZF was used as a positive control. Consistent with a previous report41, His^6^-PLZF(BTB) interacted with HDAC1 and the SMRT fragment (**Fig. 4B**). Similarly, His^6^-KIAA1841(BTB) also bound to these co-repressor proteins (**Fig. 4B**). Taken together with the earlier dimerization findings, these data suggest that KIAA1841 contains a functional BTB domain that mediates dimer formation and binding to co-repressors.

**Figure 4:**
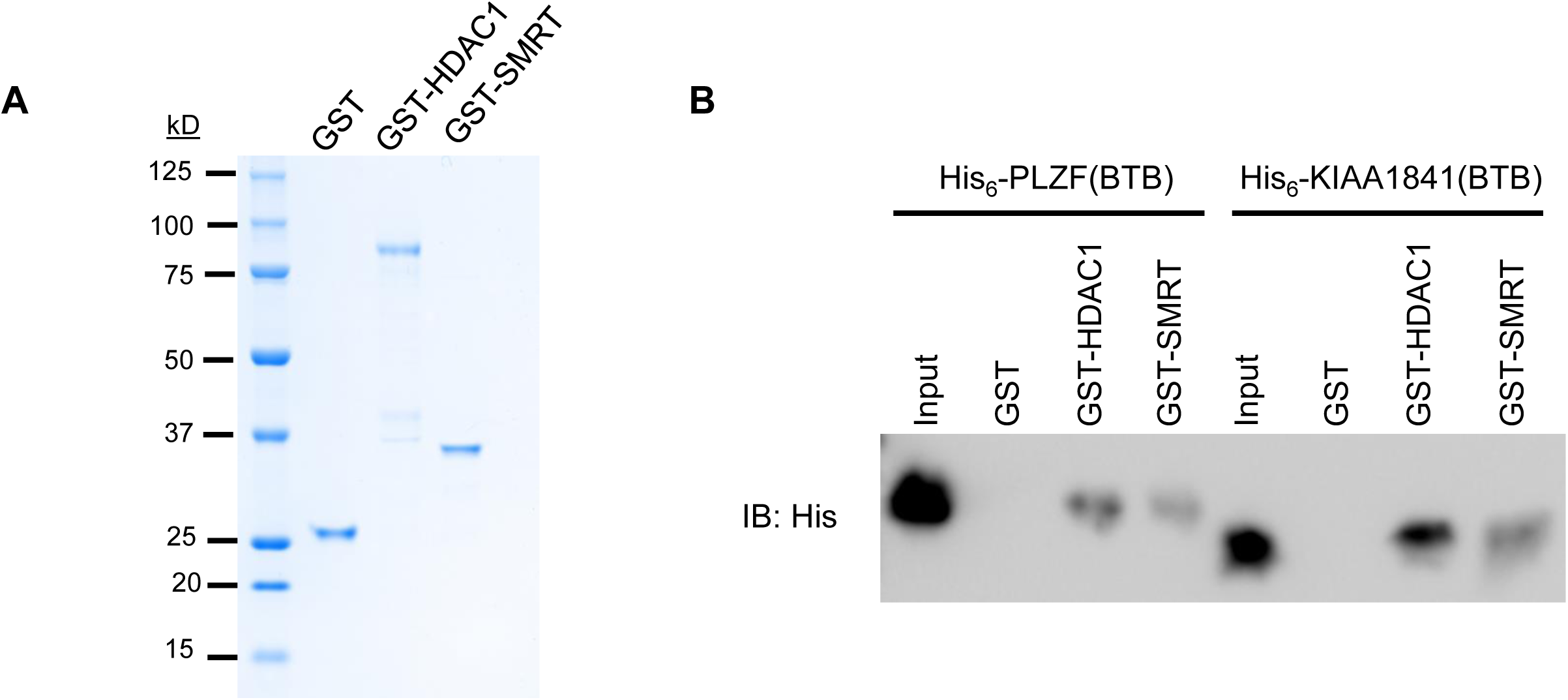
The BTB domain of KIAA1841 interacts with co-repressors. **(A)** Recombinant GST, GST-HDAC1 and GST-SMRT fragment were purified and analyzed by Coomassie stain. **(B)** Purified GST, GST-HDAC1 or GST-SMRT fragment were incubated with equimolar purified His-tagged PLZF(BTB) or KIAA1841(BTB), followed by GST-pull down and analysis by immunoblot (IB) using anti-His antibody. The results are representative of three independent pull-down experiments.

### The BTB domain of KIAA1841 is important for the inhibition of CSR

To evaluate the role of the KIAA1841 BTB and SANT domains in the regulation of CSR, Flag-Strep-tagged ▵BTB or ▵SANT deletion mutants were expressed in mouse splenic B cells that were stimulated for CSR to IgG1 by LPS+IL4. As shown earlier (**Fig. 1D, 1E**), retroviral expression of wild-type (WT) KIAA1841 protein (**Fig. 5A**) reduced CSR to IgG1 in splenic B cells (**Fig. 5B, 5C**). While the ▵SANT mutant inhibited CSR to similar levels as WT KIAA1841, the ▵BTB mutant only partially suppressed CSR (**Fig. 5B, 5C**). This suggests that the BTB domain of KIAA1841 plays an important role in the negative regulation of CSR, while the SANT domain is largely dispensable. As WT levels of CSR were not achieved with ▵BTB mutant expression, other regions of KIAA1841 likely function in tandem with the BTB domain to mediate the suppression of CSR.

**Figure 5:**
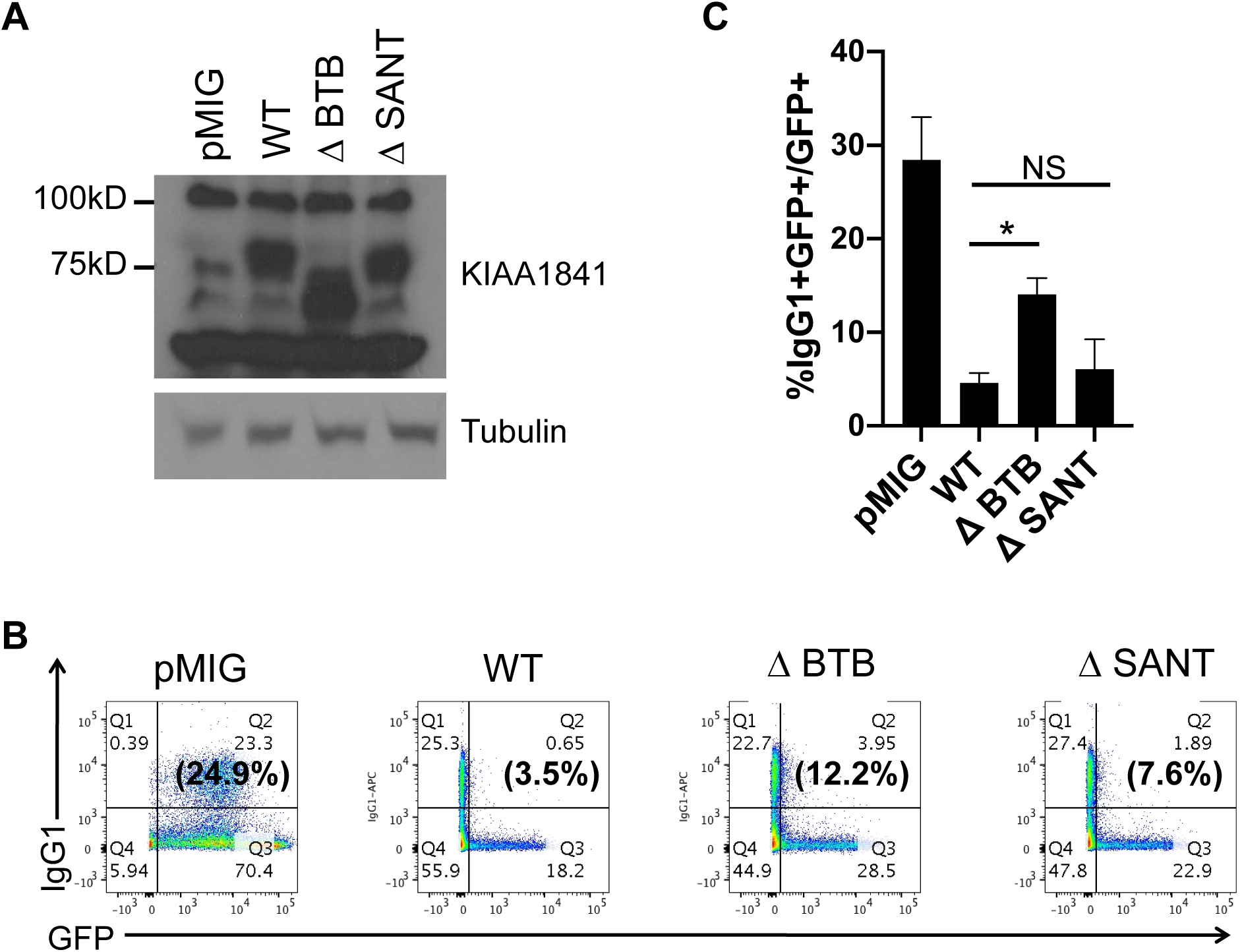
The BTB domain of KIAA1841 is necessary for inhibiting CSR. Splenic B cells were isolated from mice, stimulated with LPS+IL4, and transduced with retroviral vector control (pMIG) or vectors expressing wild-type (WT), ▵BTB or ▵SANT KIAA1841. **(A)** Expression of KIAA1841 and mutants were determined by immunoblot with KIAA1841 antibodies. Tubulin was used as a loading control. **(B)** CSR to IgG1 among the transduced GFP+ cells was determined by flow cytometry. A representative experiment is shown. The numbers in the corners of each plot indicate the percentage of cells in each quadrant while the numbers in parentheses indicate the percentage of IgG1+ cells within the GFP+ gate. **(C)** The mean %IgG1+ within the GFP+ gate from three independent experiments +/-SD is shown. * p < 0.05. NS, p = not significant, p ≥ 0.05, two-tailed paired student’s t-test.

## Discussion

Our study has identified a novel SANT- and BTB-domain containing protein, KIAA1841, that inhibits CSR. The predicted BTB domain of KIAA1841 displays characteristic properties of BTB domains, including homodimerization (**Fig. 2, Fig. 3**) as well as interaction with co-repressor proteins *in vitro* (**Fig. 4**). To the best of our knowledge, this represents the first report demonstrating this protein as a member of the BTB protein family.

BTB protein family members often serve as important transcriptional regulators that control many developmental processes^35^. For instance, PLZF interacts with N-CoR, SMRT and HDACs to mediate transcriptional repression^41^, and plays an important role in the control of the differentiation of NKT cells^36,37^, as well as the modulation of the inflammatory response in macrophages^42^. Similarly, BCL6 interacts with the corepressor BCoR to control the function of B cells and T follicular helper cells in germinal centers^43,44^. While the mechanisms by which KIAA1841 inhibits CSR are still unclear, its ability to interact with co-repressor proteins such as HDAC1 and SMRT via the BTB domain leads us to hypothesize that KIAA1841 acts as a transcriptional regulator in line with other BTB protein family members (**Fig. 4**). KIAA1841 may recruit these co-repressors to transcriptional targets to downregulate gene expression. We have shown that germline switch transcripts and AID mRNA levels are not directly regulated by KIAA1841 overexpression (**Supplementary Fig. 2**), thus the transcriptional targets of KIAA1841 that mediate its inhibitory effects on CSR await further investigation.

While the BTB domain alone is sufficient for homodimerization *in vitro* (**Fig. 2**), other regions of KIAA1841 contribute to dimer formation. Deletion of the BTB domain alone only partially impaired dimerization *in vivo* (**Fig. 3**) and resulted in a corresponding partial rescue of CSR inhibition when compared to the WT protein (**Fig. 5B, 5C**). BTB domains can also mediate heterodimerization^45^. These heterodimers may have distinct functions from homodimers, which adds another layer of regulation that can influence cellular fitness^46,47^. For instance, BCL6 has been shown to heterodimerize with Miz-1 via its BTB domain to inhibit expression of the Miz-1 target, cyclin-dependent kinase inhibitor p21, thereby allowing for proliferation of germinal center B cells^46^. Thus, KIAA1841 may similarly associate with BCL6, or other BTB protein family members, to inhibit CSR. Although the SANT domain is dispensable for dimerization (**Fig. 3**) and inhibition of CSR (**Fig. 5**), it may be involved in other non-CSR functions of KIAA1841. SANT domains are often found in chromatin remodeling proteins and function by interacting with histone tails^40,48,49^. In concert with co-repressors recruited via the BTB domain, the KIAA1841 SANT domain may modify chromatin, regulate gene transcription, and/or germinal center B cell development. These and other roles of KIAA1841 beyond inhibition of CSR await further investigation.

## Acknowledgements

This work was supported by The National Institute on Minority Health and Health Disparities (5G12MD007603), The National Cancer Institute (2U54CA132378), and The National Institute of General Medical Sciences (1SC1GM132035-01). A.J M. was supported by The American Association of Immunologists Careers in Immunology Fellowship. B.Q.V. was supported by The American Association of Immunologists Early Career Faculty Travel Grant and the PSC-CUNY Enhanced Research Award.

## Experimental Procedures

### Cell culture

Primary B cells were purified from spleens of wild-type male and female C57BL/6 mice (2-4 months old) and stimulated with LPS and IL-4 to induce CSR to IgG1 as described^50^. Retroviral infection of splenic B cells was performed as described^51^. CH12 cells were maintained and stimulated with anti-CD40, IL-4 and TGF-β to induce CSR to IgA as described^52^. C57BL/6 mice were purchased from The Jackson Laboratory. Experiments using mice were conducted according to protocols approved by The City College of New York Institutional Animal Care and Use Committee.

### Antibodies

The polyclonal KIAA1841 antibody was produced by immunizing rabbits with the C-terminal KIAA1841 peptide (RSKSRFGQGRPA) and isolating reactive antibodies from sera (Covance). Antibodies for flow cytometry include: anti-IgG1-APC (X56, BD Pharmingen), anti-IgA-FITC (C10-3, BD Pharmingen). Antibodies for immunoblot include: anti-AID^53^, anti-tubulin (DM1A, Sigma), anti-penta-His (34660, Qiagen), anti-GFP-HRP (B-2, Santa Cruz), anti-Flag (M2, Sigma).

### shRNA library screen

The lentiviral shRNA library targeting the mouse genome was a kind gift from S.J. Elledge^28,29^. Screening for negative regulators of CSR was adapted from previous protocols^28,29^. Briefly, CH12 cells were infected with the lentiviral shRNA library. Successfully transduced cells were selected with 3 μg/ml puromycin for 72 hours and subsequently stimulated to undergo CSR to IgA for 96 hours. Cells were sorted into IgA- and IgA+ populations and genomic DNA was prepared from the sorted cells. Differentially labeled (Cy5–IgA-; Cy3–IgA+) half hairpin amplicons were PCR amplified and allowed to undergo competitive hybridization onto microarrays (Agilent). Candidate negative regulators were identified as genes that are enriched in the IgA+ population at least 2-fold over the IgA-population; log^2^(Cy3/Cy5)>1.

### Protein purification

Residues 144-261 of KIAA1841 containing the BTB domain was expressed with an N-terminal His^6^-tag in BL21DE3(RIPL) *E*.*coli* and induced with 0.2mM IPTG at 16°C for 17-20 hours. Cells were lysed in lysis buffer (20mM Tris, pH 8.5; 0.5M NaCl, 50mM imidazole, 1mM TCEP, 0.5mg/ml benzonase, 1mM PMSF, cOmplete^™^ protease inhibitor, 100ug/ml lysozyme) using the LM20 Microfluidizer (Microfluidics). Lysates were clarified by centrifugation and His^6^-KIAA1841(BTB) was isolated on a HisTrap FF column (GE Healthcare). The column was washed with 10 column volumes (CVs) of lysis buffer and bound His^6^-KIAA1841(BTB) protein was eluted with 5 CVs of elution buffer (20mM Tris, pH8.5, 0.15M NaCl, 100mM imidazole, 1mM TCEP). Fractions containing His^6^-KIAA1841(BTB) were pooled, concentrated and further purified by size exclusion chromatography (SEC) using Superdex 200 10/300 GL(GE Healthcare) equilibrated and eluted with SEC buffer (10mM HEPES, pH8.0; 0.3M NaCl, 10% glycerol, 1mM TCEP). The BTB domain of PLZF (Ahman, 1998; Melnick, 2002), His^6^-PLZF(BTB), was expressed and purified as described above.

GST-tagged mouse HDAC1 and residues 1383-1467 of mouse SMRT were expressed in BL21DE3(RIPL) *E*.*coli* and induced with 0.2mM IPTG at 16°C for 17-20 hours. This region corresponds to region 1414-1498 of human SMRT, which was previously reported to interact with BTB domain of PLZF^41^. Cells were lysed in lysis buffer (20mM Tris, pH 7.5; 0.5M NaCl, 10% glycerol, 1mM TCEP, 0.5mg/ml benzonase, 1mM PMSF, cOmplete^™^ protease inhibitor, 100ug/ml lysozyme) using the LM20 Microfluidizer (Microfluidics). Lysates were clarified by centrifugation and GST-tagged protein were isolated on glutathione sepharose 4B resin (GE Healthcare) at 4°C for 2 hours. The resin was washed with 30 CVs of lysis buffer and bound proteins eluted with elution buffer (50mM Tris pH 7.5, 0.15M NaCl, 10% glycerol, 15mM glutathione, 1mM TCEP). Fractions containing GST-HDAC1 and GST-SMRT(1383-1467) were pooled, concentrated and further purified on S200 10/30 GL (GE Healthcare) using (10mM HEPES, pH7.5, 0.3M NaCl, 10% glycerol, 1mM TCEP).

### Glutaraldehyde crosslinking

Purified His^6^-KIAA1841(BTB) protein at 0.1mg/ml in sample buffer (10mM HEPES, pH8.0; 0.3M NaCl, 10% glycerol, 1mM TCEP) was crosslinked with 5-fold increasing concentration of glutaraldehyde (i.e. 0.0008%, 0.0004%, 0.02%, 0.1% (v/v)) in a total volume of 10μl at 37°C for 5 min. The reaction was quenched by adding 1μl of 1M Tris, pH 8.0 and boiled in 1X SDS loading buffer, before separation on a 12% SDS-PAGE denaturing gel and stained with coomassie blue to visualize the migration of the resultant protein species.

### In vitro co-repressor binding assay

Purified His^6^-KIAA1841(BTB) or His^6^-PLZF(BTB) at 200nM was mixed with equimolar amounts of purified GST, GST-HDAC1 or GST-SMRT(1383-1467) fragment in binding buffer (50mM HEPES pH 8.0, 150mM NaCl, 0.1% IGEPAL-CA630, 2mM EDTA), in a total volume of 400 μl, at 4°C for 2 hours. Glutathione sepharose 4B resin (GE Healthcare), which was pre-washed thrice with binding buffer, was added and mixed with the proteins at 4°C for an additional 2 hours, then washed thrice with binding buffer. Bound complexes were released by boiling in 2X SDS loading buffer and analyzed by immunoblot with anti-His antibody. 30 ng His^6^-KIAA1841(BTB) or 33 ng His^6^-PLZF(BTB) (i.e. 5% of that used in the assay) were loaded as inputs.

### In vivo dimerization assay

293T cells were co-transfected with GFP-tagged KIAA1841 and either Flag-Strep-tagged WT KIAA; KIAA1841▵BTB, which lacks amino acids 147-255; or KIAA1841▵SANT, which lacks amino acids 48-59. After transfection (48hr), cells from two confluent wells of a 6-well plate were washed with PBS and lysed with 200 μl lysis buffer (20mM Tris, pH 7.5, 150mM NaCl, 5% glycerol, 0.5% NP40) supplemented with 1mM PMSF, 1mM DTT, and cOmplete mini protease inhibitor cocktail (Roche). Lysates were clarified by centrifugation at 21,000g for 15min at 4°C, and incubated with 20μl MagStrep “type 3” XT beads (IBA) for 2 hours at 4°C to pull down Flag-Strep-tagged KIAA1841 and mutant proteins. The beads were washed thrice with lysis buffer and bound proteins were recovered by boiling in 1X SDS loading dye and analyzed by immunoblot.

