## Supplemental figures and tables for "KIAA1841, a novel SANT and BTB domain-containing protein, inhibits class switch recombination"

### Supplemental figure and table legends

**Supplementary Figure 1: KIAA1841 is well conserved among vertebrates.** NCBI HomoloGene pairwise alignment scores of human KIAA1841 sequence compared to other vertebrate orthologs.

**Supplementary Figure 2: μ-germline transcripts (GLT) and AID mRNA expression are not affected by overexpression of KIAA1841.**

Splenic B cells were isolated from wild-type mice, and transduced with retroviral vector control (pMIG) or a vector expressing KIAA1841. Levels of μ-GLT and AID mRNA were determined at 72h post-infection by RT-qPCR, and normalized to β-actin mRNA and the pMIG control. The mean of three independent experiments +/- SD is shown. NS, p = not significant, $p \geq 0.05$, two-tailed paired student's t-test.

**Supplementary Figure 3: Overexpression of untagged and Xpress-tagged KIAA1841 does not affect cell proliferation.**

Splenic B cells were isolated from wild-type mice, stimulated with LPS+IL4, and transduced with retroviral vector control (pMIG) or a vector expressing KIAA1841 or Xpress (Xp)-tagged KIAA1841. Cells are labeled with seminaphthorhodafluor (SNARF) cell tracking dye and SNARF intensity was measured at 0h, 24h, and 48h after retroviral infection. Histograms of transduced cells (GFP+) for pMIG control, KIAA1841 and Xp-KIAA1841 were superimposed at each time point. Three independent retroviral infection experiments are shown.

**Supplementary Figure 4: Sequence alignment of the putative SANT domain of KIAA1841 with the SANT domain of SWI3.** Alignment was performed using ClustalOmega; *, conserved residue, :, strongly similar residues, ., weakly similar residues. Numbers indicate amino acid residues in each sequence. Helix 3, which has been reported to be important for the chromatin remodeling function of SWI3 (Boyer et. al., 2002), is well conserved in KIAA1841.

**Supplementary Table 1: Negative regulators of CSR recovered by the shRNA screen.** Genes with shRNAs showing at least 2-fold enrichment in IgA+ compared to IgA- samples, i.e. $\log_2$(Cy3/Cy5)>1, and FDR less than 10 were identified as negative regulators. The novel gene of interest, NM_027860, which encodes for the protein KIAA1841, is highlighted in yellow.

**Supplementary Table 2: Positive regulators of CSR recovered by the shRNA screen.** Genes with shRNAs showing at least 2-fold enrichment in IgA- compared to IgA+ samples, i.e. $\log_2$(Cy3/Cy5)<-1, and FDR less than 10, were identified as positive regulators. Known positive regulator, activation-induced cytidine deaminase (*Aicda*), is highlighted in green.

### Pairwise Alignment Scores

| Gene | | Identity (%) | |
|---|---|---|---|
| Species | Symbol | Protein | DNA |
| **H.sapiens** | **KIAA1841** | | |
| vs. P.troglodytes | KIAA1841 | 99.2 | 99.5 |
| vs. M.mulatta | LOC717910 | 97.8 | 97.7 |
| vs. C.lupus | KIAA1841 | 92.3 | 92.2 |
| vs. M.musculus | 0610010F05Rik | 88.6 | 88.2 |
| vs. R.norvegicus | RGD1305110 | 88.4 | 87.2 |
| vs. G.gallus | KIAA1841 | 75.9 | 77.1 |
| vs. X.tropicalis | kiaa1841 | 70.9 | 70.4 |
| vs. D.rerio | LOC793832 | 61.7 | 62.8 |

**Supplementary Figure 1**

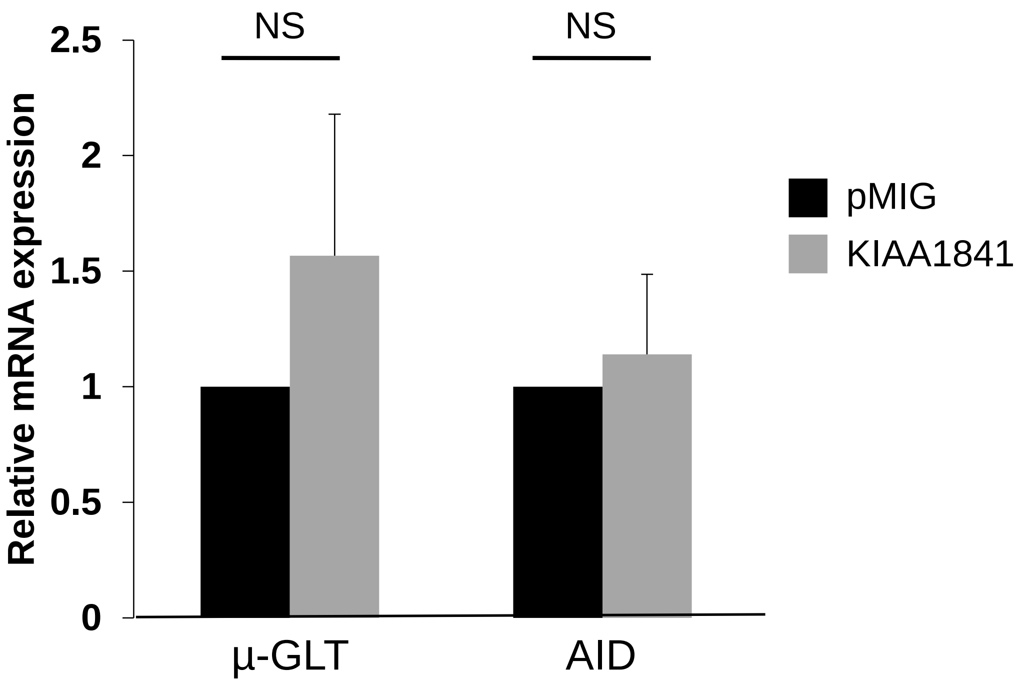

**Supplementary Figure 2**

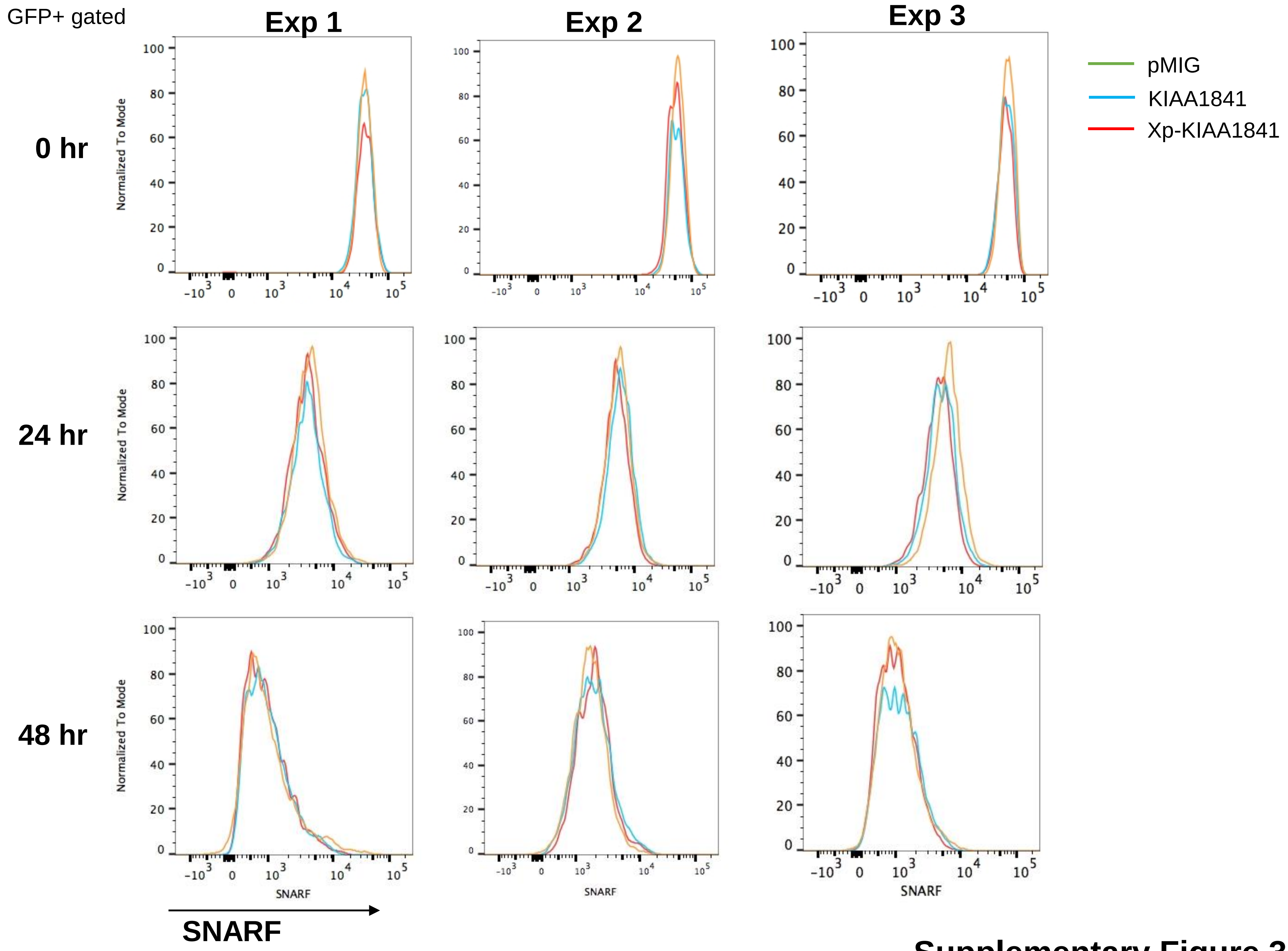

**Supplementary Figure 3**

```
KIAA1841  MVLDMILYPLIGI-PQTINWETVARLVPGLTPKECVKRFDELKS------ 60
SWI3      WSKEDLQKLLKGIQEFGADWYKVAKNVGNKSPEQCILRFLQLPIEDKFLY 576
             : :   * **     :* .**: * . :*::*: ** :*
```

Helix 3

**Supplementary Figure 4**

**Supplementary Table 1: Negative regulators of CSR recovered by the shRNA screen**

| Gene.Name | log2.ratio | q.value... | Hit? | Codex_hai | Codex_97 | Codex_19 | Codex_Accession | Codex_gene.name | Codex_gene.symbol | OB_Releas | OB_Plate | OB_Well_I | OB_Well_( |
|---|---|---|---|---|---|---|---|---|---|---|---|---|---|
| 092308m1_V2MM_141890 | 3.829646 | 3.004939 | TRUE | HP_36260 | TGCTGTTG | GCTTTATC | XM_157111 | Mus musculus LOC212310 (LOC212310), mR. | | 2.4 | SM2050 | h | 12 |
| mPool2_V2MM_133332 | 3.684535 | 3.004939 | TRUE | HP_35407 | TGCTGTTG | GATTCTCT | XM_149195 | RIKEN cDNA E230025N21 gene | E230025N21Rik | 2.11 | SM2373 | e | 7 |
| mPool2_V2MM_2305 | 3.67322 | 2.205692 | TRUE | HP_22592 | TGCTGTTG | CTCCTAAA | XM_132611 | similar to variable region of immunoglobulin kappa light chain | | 2.11 | SM2381 | h | 4 |
| mPool7_V2MM_141895 | 3.615101 | 3.004939 | TRUE | HP_36260 | TGCTGTTG | CGCTTTAT | XM_157111 | Mus musculus LOC212310 (LOC212310), mRNA. | | 2.1 | SM2329 | a | 8 |
| mPool7_V2MM_88676 | 3.592123 | 2.205692 | TRUE | HP_31011 | TGCTGTTG | GTACAGA | NM_175463 | RIKEN cDNA A930012O16 gene | A930012O16Rik | 2.9 | SM2257 | e | 8 |
| mPool7_V2MM_88676 | 3.592123 | 2.205692 | TRUE | HP_31011 | TGCTGTTG | GTACAGA | NM_175463 | RIKEN cD A930012O16 gene | A930012O16Rik | 2.9 | SM2257 | e | 8 |
| mPool2_V2MM_145554 | 3.486776 | 3.004939 | TRUE | HP_36626 | TGCTGTTG | CATTCTTA | XM_161053 | Mus musculus LOC230170 (LOC230170), mR. | | 2.4 | SM2040 | e | 9 |
| mPool2_V2MM_145554 | 3.486776 | 3.004939 | TRUE | HP_36626 | TGCTGTTG | CATTCTTA | XM_161053 | Mus musculus LOC230170 (LOC230170), mRNA. | | 2.4 | SM2040 | e | 9 |
| mPool2_V2MM_92494 | 3.390594 | 3.004939 | TRUE | HP_31384 | TGCTGTTG | GTTGGAA | NM_178720 | RIKEN cDNA 9430016A21 gene | 9430016A21Rik | 2.6 | SM2137 | a | 5 |
| mPool2_V2MM_138848 | 3.381957 | 2.205692 | TRUE | HP_35956 | TGCTGTTG | GACATTA | XM_154566 | Mus musculus LOC238925 (LOC238925), mR. | | 2.4 | SM2030 | d | 11 |
| mPool2_V2MM_138848 | 3.381957 | 2.205692 | TRUE | HP_35956 | TGCTGTTG | GACATTA | XM_154566 | Mus musculus LOC238925 (LOC238925), mRNA. | | 2.4 | SM2030 | d | 11 |
| mPool2_V2MM_134275 | 3.329965 | 2.205692 | TRUE | HP_35500 | TGCTGTTG | GGCTAGT | XM_357654 | hypothetical LOC384452 | | 2.11 | SM2368 | f | 8 |
| mPool7_V2MM_199459 | 3.328607 | 2.205692 | TRUE | HP_41750 | TGCTGTTG | CCATTTAA | XM_289192 | Mus musculus LOC332813 (LOC332813), mRNA. | | 2.9 | SM2276 | c | 3 |
| mPool2_V2MM_17530 | 3.318772 | 2.205692 | TRUE | HP_24080 | TGCTGTTG | GATACTTT | NM_134054 | RIKEN cDNA 1110002B05 gene | 1110002B05Rik | 2.11 | SM2378 | g | 1 |
| 092308m1_V2MM_171708 | 3.314424 | 1.959154 | TRUE | HP_39219 | TGCTGTTG | CATGTTAC | XM_286679 | Mus musculus hypothetical gene supported by AK039985 (LOC328540), mR. | | 2.5 | SM2074 | b | 9 |
| mPool2_V2MM_36734 | 3.311664 | 3.004939 | TRUE | HP_25950 | TGCTGTTG | CTTGTACA | NM_023816 | RIKEN cDNA 1700012M14 gene | 1700012M14Rik | 2.11 | SM2392 | a | 9 |
| mPool2_V2MM_84705 | 3.309396 | 3.004939 | TRUE | HP_14696 | TGCTGTTG | CTAATCAT | NM_028930 | transmembrane channel-like gene family 5 | Tmc5 | 2.6 | SM2137 | b | 10 |
| 092308m1_V2MM_121144 | 3.273188 | 3.004939 | TRUE | HP_34202 | TGCTGTTG | CGGCATC | XM_141454 | Mus musculus similar to lactate dehydrogese [Mus musculus] (LOC228430), mR. | | 2.6 | SM2106 | g | 6 |
| 092308m1_V2MM_124008 | 3.268085 | 2.205692 | TRUE | HP_34486 | TGCTGTTG | GTCTGGA | XM_143251 | Mus musculus similar to hypothetical protein [Macaca fascicularis] (LOC242043), | | 2.6 | SM2102 | b | 7 |
| mPool7_V2MM_113184 | 3.252594 | 2.205692 | TRUE | HP_33417 | TGCTGTTG | CTACATTC | XM_136853 | Mus musculus LOC241181 (LOC241181), mRNA. | | 2.1 | SM2304 | d | 4 |
| mPool2_V2MM_120733 | 3.242869 | 2.205692 | TRUE | HP_34162 | TGCTGTTG | GAAATTTC | XM_141156 | Mus musculus similar to olfactory receptor MOR264-19 [Mus musculus] (LOC228 | | 2.11 | SM2359 | f | 5 |
| mPool7_V2MM_79723 | 3.198836 | 3.004939 | TRUE | HP_30137 | TGCTGTTG | GCCAGGA | NM_009828 | cyclin A2 | Ccna2 | 2.8 | SM2227 | d | 7 |
| 092308m1_V2MM_159994 | 3.196066 | 2.205692 | TRUE | HP_38058 | TGCTGTTG | CATCTAAC | AK048916 | cD sequence BC052055 | BC052055 | 2.6 | SM2115 | f | 1 |
| mPool2_V2MM_14135 | 3.188079 | 2.205692 | TRUE | HP_23748 | TGCTGTTG | CCCTGAA | NM_007672 | cerebellar degeneration-related 2 | Cdr2 | 2.11 | SM2379 | d | 10 |
| mPool2_V2MM_212177 | 3.187273 | 1.959154 | TRUE | HP_42985 | TGCTGTTG | CTTCTATC | XM_148779 | RIKEN cDNA 4930541O19 gene | 4930541O19Rik | 2.11 | SM2365 | f | 7 |
| mPool2_V2MM_17214 | 3.164486 | 2.205692 | TRUE | HP_24049 | TGCTGTTG | GAGCTAA | NM_011623 | topoisomerase (DNA) II alpha | Top2a | 2.11 | SM2387 | h | 6 |
| mPool7_V2MM_67258 | 3.157407 | 2.205692 | TRUE | HP_28924 | TGCTGTTG | CCGTTTGC | NM_021546 | amyloid beta (A4) precursor protein-binding, family A, m | Apba2bp | 2.8 | SM2237 | d | 7 |
| mPool2_V2MM_208999 | 3.152098 | 3.004939 | TRUE | HP_42677 | TGCTGTTG | CTCAGGA | XM_157672 | similar to Suppressor of G2 allele of SKP1 homolog (Sgt1) (Putative 40-6-3 protei | | 2.1 | SM2338 | a | 5 |
| mPool2_V2MM_94467 | 3.148552 | 2.205692 | TRUE | HP_31575 | TGCTGTTG | CAGCATG | BC052532 | cDNA sequence BC006779 | BC006779 | 2.6 | SM2142 | a | 7 |
| mPool7_V2MM_202647 | 3.143102 | 2.205692 | TRUE | HP_42064 | TGCTGTTG | GTATGATT | AK045028 | histone deacetylase 10 | Hdac10 | 2.1 | SM2317 | h | 10 |
| mPool2_V2MM_1433 | 3.143072 | 2.205692 | TRUE | HP_22507 | TGCTGTTG | CCGCATAC | NM_007855 | twist homolog 2 (Drosophila) | Twist2 | 2.11 | SM2386 | g | 12 |
| mPool2_V2MM_134121 | 3.135473 | 3.004939 | TRUE | HP_35484 | TGCTGTTG | CTGTCTTT | NM_177582 | hypothetical protein B930074I24 | | 2.1 | SM2349 | h | 4 |
| 092308m1_V2MM_180277 | 3.099315 | 2.205692 | TRUE | HP_40075 | TGCTGTTG | CTTTAGAT | XM_288480 | Mus musculus LOC331828 (LOC331828), mRNA. | | 2.5 | SM2072 | h | 3 |
| 092308m1_V2MM_180277 | 3.099315 | 2.205692 | TRUE | HP_40075 | TGCTGTTG | CTTTAGAT | XM_288480 | Mus musculus LOC331828 (LOC331828), mR. | | 2.5 | SM2072 | h | 3 |
| 092308m1_V2MM_156420 | 3.091247 | 2.205692 | TRUE | HP_37705 | TGCTGTTG | CTGCTTTA | NM_027860 | RIKEN cD 0610010F05 gene | 0610010F05Rik | 2.6 | SM2115 | e | 4 |
| 092308m1_V2MM_70303 | 3.088266 | 3.004939 | TRUE | HP_29221 | TGCTGTTG | CAACCAA | NM_028911 | RIKEN cD 4933414I15 gene | 4933414I15Rik | 2.3 | SM2006 | e | 3 |
| mPool7_V2MM_197834 | 3.086251 | 2.205692 | TRUE | HP_41589 | TGCTGTTG | GATGATA | XM_111554 | similar to Aspartate aminotransferase, mitochondrial precursor (Transamise A) (( | | 2.9 | SM2266 | b | 12 |
| mPool7_V2MM_197834 | 3.086251 | 2.205692 | TRUE | HP_41589 | TGCTGTTG | GATGATA | XM_111554 | similar to Aspartate aminotransferase, mitochondrial precursor (Transaminase A | | 2.9 | SM2266 | b | 12 |
| mPool7_V2MM_205300 | 3.075308 | 2.205692 | TRUE | HP_42316 | TGCTGTTG | CAAAGTT | XM_138746 | Mus musculus similar to thymine D glycosylase [Mus musculus] (LOC238743), mR | | 2.1 | SM2306 | d | 1 |
| mPool7_V2MM_205300 | 3.075308 | 2.205692 | TRUE | HP_42316 | TGCTGTTG | CAAAGTT | XM_138746 | Mus musculus similar to thymine DNA glycosylase [Mus musculus] (LOC238743), | | 2.1 | SM2306 | d | 1 |
| 092308m1_V2MM_135215 | 3.072112 | 1.959154 | TRUE | HP_35593 | TGCTGTTG | CTTTATTT | XM_489372 | similar to RIKEN cD 6720401G13 gene | | 2.5 | SM2057 | a | 3 |
| mPool2_V2MM_212145 | 3.039733 | 3.004939 | TRUE | HP_42982 | TGCTGTTG | CCCTTTAA | AK017143 | RIKEN cDNA 5031425E22 gene | 5031425E22Rik | 2.11 | SM2362 | d | 9 |
| mPool7_V2MM_144853 | 3.031093 | 2.205692 | TRUE | HP_36556 | TGCTGTTG | GAGCTTA | XM_160377 | Mus musculus LOC229769 (LOC229769), mRNA. | | 2.1 | SM2331 | f | 2 |
| 092308m1_V2MM_155240 | 3.030417 | 2.205692 | TRUE | HP_37588 | TGCTGTTG | GCCTGAA | XM_197286 | Mus musculus LOC271407 (LOC271407), mR. | | 2.6 | SM2114 | g | 7 |
| mPool2_V2MM_106531 | 3.008607 | 3.004939 | TRUE | HP_32763 | TGCTGTTG | GAAACAT | NM_001002927 | preproenkephalin 1 | Penk1 | 2.1 | SM2316 | h | 4 |
| mPool2_V2MM_210729 | 3.001055 | 2.205692 | TRUE | HP_42846 | TGCTGTTG | CGCAGTT | XM_359191 | gene model 1773, (NCBI) | Gm1773 | 2.11 | SM2363 | b | 5 |
| mPool2_V2MM_123973 | 2.985185 | 2.205692 | TRUE | HP_34482 | TGCTGTTG | GACAGGA | XM_143235 | Mus musculus similar to embryonic blastocoelar extracellular matrix protein prec | | 2.11 | SM2351 | d | 7 |
| 092308m1_V2MM_171296 | 2.978236 | 2.205692 | TRUE | HP_39178 | TGCTGTTG | CCTAGAG | XM_286593 | Mus musculus hypothetical gene supported by AK051402 (LOC328380), mR. | | 2.5 | SM2080 | h | 5 |
| mPool2_V2MM_142344 | 2.975649 | 2.205692 | TRUE | HP_36305 | TGCTGTTG | CACGAAA | XM_157589 | Mus musculus LOC225224 (LOC225224), mRNA. | | 2.4 | SM2039 | c | 5 |
| mPool2_V2MM_142344 | 2.975649 | 2.205692 | TRUE | HP_36305 | TGCTGTTG | CACGAAA | XM_157589 | Mus musculus LOC225224 (LOC225224), mR. | | 2.4 | SM2039 | c | 5 |
| mPool2_V2MM_204673 | 2.974126 | 3.004939 | TRUE | HP_42256 | TGCTGTTG | GATGAAT | XM_129159 | RIKEN cDNA 1810073H04 gene | 1810073H04Rik | 2.1 | SM2318 | f | 8 |

| | | | | | | | | | | | | | |
|---|---|---|---|---|---|---|---|---|---|---|---|---|---|
| 092308m1_V2MM_172099 | 2.955371 | 3.004939 | TRUE | HP_39258 | TGCTGTTG | CATTCTGA | XM_286756 | Mus musculus hypothetical gene supported by AK048704 (LOC328685), mRNA. | | 2.5 | SM2090 | h | 5 |
| 092308m1_V2MM_172099 | 2.955371 | 3.004939 | TRUE | HP_39258 | TGCTGTTG | CATTCTGA | XM_286756 | Mus musculus hypothetical gene supported by AK048704 (LOC328685), mR. | | 2.5 | SM2090 | h | 5 |
| mPool2_V2MM_128454 | 2.921823 | 2.205692 | TRUE | HP_34926 | TGCTGTTG | CAGTAGA | XM_145689 | similar to glyceraldehyde-3-phosphate dehydrogenase | | 2.11 | SM2367 | f | 8 |
| mPool2_V2MM_136210 | 2.918504 | 3.004939 | TRUE | HP_35693 | TGCTGTTG | CCTACGAA | XM_151496 | Mus musculus LOC241166 (LOC241166), mRNA. | | 2.5 | SM2053 | h | 11 |
| mPool2_V2MM_136210 | 2.918504 | 3.004939 | TRUE | HP_35693 | TGCTGTTG | CCTACGAA | XM_151496 | Mus musculus LOC241166 (LOC241166), mR. | | 2.5 | SM2053 | h | 11 |
| 092308m1_V2MM_124037 | 2.908688 | 3.004939 | TRUE | HP_34489 | TGCTGTTG | CAATTGTC | XM_143269 | Mus musculus similar to Heat shock protein HSP 90-beta (HSP 84) (LOC229352), r | | 2.6 | SM2103 | g | 10 |
| mPool2_V2MM_126884 | 2.906645 | 3.004939 | TRUE | HP_34772 | TGCTGTTG | CTTGATTA | XM_144862 | Mus musculus similar to Gag [Ovis aries] (LOC232180), mRNA. | | 2.11 | SM2360 | d | 5 |
| mPool2_V2MM_10177 | 2.90313 | 3.004939 | TRUE | HP_23361 | TGCTGTTG | CTACTATT | NM_007785 | casein gamma | Csng | 2.11 | SM2374 | a | 10 |
| 092308m1_V2MM_186411 | 2.895897 | 3.004939 | TRUE | HP_40688 | TGCTGTTG | CCCTTCAA | XM_289534 | Mus musculus LOC333278 (LOC333278), mR. | | 2.5 | SM2084 | g | 3 |
| 092308m1_V2MM_186411 | 2.895897 | 3.004939 | TRUE | HP_40688 | TGCTGTTG | CCCTTCAA | XM_289534 | Mus musculus LOC333278 (LOC333278), mRNA. | | 2.5 | SM2084 | g | 3 |
| mPool7_V2MM_180521 | 2.887155 | 2.205692 | TRUE | HP_40099 | TGCTGTTG | CAGCCAGA | XM_288521 | Mus musculus LOC331896 (LOC331896), mRNA. | | 2.9 | SM2277 | c | 7 |
| mPool2_V2MM_139762 | 2.886327 | 3.004939 | TRUE | HP_36048 | TGCTGTTG | CCAATTAC | XM_155380 | similar to Rpl7a protein | | 2.5 | SM2055 | f | 2 |
| mPool2_V2MM_139762 | 2.886327 | 3.004939 | TRUE | HP_36048 | TGCTGTTG | CCAATTAC | XM_155380 | similar to Rpl7a protein | | 2.5 | SM2055 | f | 2 |
| mPool2_V2MM_86036 | 2.885936 | 2.205692 | TRUE | HP_30751 | TGCTGTTG | CATGTACA | NM_146605 | olfactory receptor 828 | Olfr828 | 2.6 | SM2145 | b | 6 |
| mPool2_V2MM_33233 | 2.885463 | 2.205692 | TRUE | HP_25609 | TGCTGTTG | CACTAAAT | AK044117 | RIKEN cDNA C430014H23 gene | C430014H23Rik | 2.11 | SM2391 | e | 11 |
| mPool7_V2MM_117268 | 2.884369 | 3.004939 | TRUE | HP_33821 | TGCTGTTG | CAGATGAT | XM_139295 | caspase recruitment domain family, member 6 | Card6 | 2.1 | SM2314 | a | 1 |
| 092308m1_V2MM_124502 | 2.881563 | 3.004939 | TRUE | HP_34535 | TGCTGTTG | GCCTTCTT | XM_146378 | similar to HMG-1 | | 2.6 | SM2103 | f | 7 |
| 092308m1_V2MM_136210 | 2.874993 | 3.004939 | TRUE | HP_35693 | TGCTGTTG | CCTACGAA | XM_151496 | Mus musculus LOC241166 (LOC241166), mR. | | 2.5 | SM2053 | h | 11 |
| 092308m1_V2MM_136210 | 2.874993 | 3.004939 | TRUE | HP_35693 | TGCTGTTG | CCTACGAA | XM_151496 | Mus musculus LOC241166 (LOC241166), mRNA. | | 2.5 | SM2053 | h | 11 |
| mPool7_V2MM_86351 | 2.86126 | 3.004939 | TRUE | HP_30782 | TGCTGTTG | CCTTTATA | NM_146691 | olfactory receptor 1467 | Olfr1467 | 2.9 | SM2256 | e | 8 |
| mPool4_V2MM_262088 | 2.857863 | 75.83547 | TRUE | HP_52497 | TGCTGTTG | CTTGTACA | NM_134206 | vomeronasal 1 receptor, G5 | V1rg5 | 2.15 | SM2582 | f | 8 |
| mPool2_V2MM_159248 | 2.854788 | 3.004939 | TRUE | HP_37984 | TGCTGTTG | GGGTTATC | XM_283022 | zinc finger protein, subfamily 1A, 3 (Aiolos) | Zfpn1a3 | 2.6 | SM2132 | c | 5 |
| 092308m1_V2MM_62859 | 2.854269 | 2.205692 | TRUE | HP_28495 | TGCTGTTG | CATTCTTC | NM_010672 | keratin associated protein 6-1 | Krtap6-1 | 2.4 | SM2020 | f | 4 |
| mPool7_V2MM_179575 | 2.838696 | 2.205692 | TRUE | HP_40004 | TGCTGTTG | CTTCAAAT | XM_288356 | Mus musculus LOC331675 (LOC331675), mRNA. | | 2.9 | SM2287 | g | 12 |
| 092308m1_V2MM_142344 | 2.837502 | 2.205692 | TRUE | HP_36305 | TGCTGTTG | CACGAAAC | XM_157589 | Mus musculus LOC225224 (LOC225224), mRNA. | | 2.4 | SM2039 | c | 5 |
| 092308m1_V2MM_142344 | 2.837502 | 2.205692 | TRUE | HP_36305 | TGCTGTTG | CACGAAAC | XM_157589 | Mus musculus LOC225224 (LOC225224), mR. | | 2.4 | SM2039 | c | 5 |
| mPool7_V2MM_180277 | 2.837056 | 2.205692 | TRUE | HP_40075 | TGCTGTTG | CTTTAGAT | XM_288480 | Mus musculus LOC331828 (LOC331828), mRNA. | | 2.5 | SM2072 | h | 3 |
| mPool7_V2MM_180277 | 2.837056 | 2.205692 | TRUE | HP_40075 | TGCTGTTG | CTTTAGAT | XM_288480 | Mus musculus LOC331828 (LOC331828), mR. | | 2.5 | SM2072 | h | 3 |
| 092308m1_V2MM_149019 | 2.83581 | 1.959154 | TRUE | HP_36973 | TGCTGTTG | CTGGGAAC | XM_164412 | RIKEN cD 1600027J07 gene | 1600027J07Rik | 2.5 | SM2058 | h | 4 |
| mPool2_V2MM_89163 | 2.828845 | 2.205692 | TRUE | HP_31059 | TGCTGTTG | CCTCCAAC | NM_178220 | arrestin, beta 1 | Arrb1 | 2.6 | SM2143 | b | 9 |
| 092308m1_V2MM_187186 | 2.825406 | 2.205692 | TRUE | HP_40765 | TGCTGTTG | CTCCTACT | XM_289663 | Mus musculus LOC333427 (LOC333427), mR. | | 2.5 | SM2069 | f | 3 |
| 092308m1_V2MM_67710 | 2.824968 | 2.205692 | TRUE | HP_28969 | TGCTGTTG | CTATTACG | AB086123 | calcium channel, voltage-dependent, L type, alpha 1D sul | Cac1d | 2.3 | SM2008 | e | 10 |
| 092308m1_V2MM_83700 | 2.824072 | 3.004939 | TRUE | HP_30522 | TGCTGTTG | GAGAATGC | NM_023750 | zinc finger protein 84 | Zfp84 | 2.6 | SM2112 | g | 4 |
| mPool7_V2MM_198010 | 2.823741 | 2.205692 | TRUE | HP_41607 | TGCTGTTG | CTATTTCT | XM_287838 | Mus musculus hypothetical gene supported by AK048651 (LOC330802), mRNA. | | 2.9 | SM2277 | b | 5 |
| 092308m1_V2MM_176389 | 2.82183 | 2.205692 | TRUE | HP_39687 | TGCTGTTG | CCTTAAAT | AK129335 | RIKEN cDNA C230080I20 gene | C230080I20Rik | 2.5 | SM2077 | f | 6 |
| 092308m1_V2MM_176389 | 2.82183 | 2.205692 | TRUE | HP_39687 | TGCTGTTG | CCTTAAAT | AK129335 | RIKEN cD C230080I20 gene | C230080I20Rik | 2.5 | SM2077 | f | 6 |
| mPool7_V2MM_171714 | 2.819084 | 2.205692 | TRUE | HP_39220 | TGCTGTTG | CAGTGTTC | XM_286680 | Mus musculus hypothetical gene supported by AK043364 (LOC328593), mRNA. | | 2.9 | SM2294 | g | 11 |
| mPool2_V2MM_97290 | 2.816496 | 2.205692 | TRUE | HP_31853 | TGCTGTTG | CACCATTT | XM_111418 | Mus musculus similar to ribosomal protein S9, cytosolic [validated] - rat (LOC218 | | 2.6 | SM2144 | e | 11 |
| 092308m1_V2MM_67000 | 2.815659 | 2.205692 | TRUE | HP_28900 | TGCTGTTG | CCCAGTCA | NM_028979 | cytochrome P450, family 2, subfamily j, polypeptide 9 | Cyp2j9 | 2.4 | SM2028 | b | 6 |
| mPool7_V2MM_177652 | 2.809365 | 3.004939 | TRUE | HP_39813 | TGCTGTTG | GAATTGGA | AK084093 | RIKEN cDNA D130086K05 gene | D130086K05Rik | 2.5 | SM2060 | f | 7 |
| mPool7_V2MM_177652 | 2.809365 | 3.004939 | TRUE | HP_39813 | TGCTGTTG | GAATTGGA | AK084093 | RIKEN cD D130086K05 gene | D130086K05Rik | 2.5 | SM2060 | f | 7 |
| mPool7_V2MM_102459 | 2.802223 | 3.004939 | TRUE | HP_32361 | TGCTGTTG | CTGAACTT | XM_484567 | RIKEN cDNA 2310061N02 gene | 2310061N02Rik | 2.1 | SM2302 | g | 2 |
| 092308m1_V2MM_174236 | 2.780406 | 3.004939 | TRUE | HP_39472 | TGCTGTTG | CTTCCCTT | AK087574 | RIKEN cD E230011G24 gene | E230011G24Rik | 2.5 | SM2082 | h | 8 |
| 092308m1_V2MM_142084 | 2.768636 | 2.205692 | TRUE | HP_36279 | TGCTGTTG | GACAGAAC | XM_157307 | Mus musculus LOC225477 (LOC225477), mR. | | 2.4 | SM2046 | e | 8 |
| 092308m1_V2MM_142084 | 2.768636 | 2.205692 | TRUE | HP_36279 | TGCTGTTG | GACAGAAC | XM_157307 | Mus musculus LOC225477 (LOC225477), mRNA. | | 2.4 | SM2046 | e | 8 |
| mPool2_V2MM_18361 | 2.753327 | 3.004939 | TRUE | HP_24160 | TGCTGTTG | GCCTGTTT | NM_022880 | solute carrier family 29 (nucleoside transporters), memb | Slc29a1 | 2.11 | SM2377 | a | 3 |
| 092308m1_V2MM_175268 | 2.751147 | 3.004939 | TRUE | HP_39575 | TGCTGTTG | GTCTTAAT | XM_287405 | Mus musculus hypothetical gene supported by AK037004 (LOC330002), mR. | | 2.5 | SM2086 | d | 10 |
| mPool2_V2MM_120290 | 2.748502 | 1.959154 | TRUE | HP_34118 | TGCTGTTG | CTTCATTG | XM_140842 | Mus musculus similar to glyceraldehyde-3-phosphate dehydrogenase [Mus musc | | 2.11 | SM2370 | e | 4 |
| 092308m1_V2MM_172614 | 2.744887 | 2.205692 | TRUE | HP_39310 | TGCTGTTG | CAGGAAAC | XM_286868 | hypothetical protein 9330117O12 | | 2.5 | SM2072 | f | 12 |
| mPool2_V2MM_91867 | 2.741015 | 1.959154 | TRUE | HP_12628 | TGCTGTTG | CCTACAAA | NM_178607 | ring finger protein 24 | Rnf24 | 2.6 | SM2109 | h | 2 |
| mPool2_V2MM_91867 | 2.741015 | 1.959154 | TRUE | HP_12628 | TGCTGTTG | CCTACAAA | NM_178607 | ring finger protein 24 | Rnf24 | 2.6 | SM2109 | h | 2 |
| mPool2_V2MM_148147 | 2.740015 | 3.004939 | TRUE | HP_36885 | TGCTGTTG | GAAGGTTC | XM_163530 | Mus musculus LOC244133 (LOC244133), mR. | | 2.4 | SM2040 | h | 1 |
| mPool2_V2MM_148147 | 2.740015 | 3.004939 | TRUE | HP_36885 | TGCTGTTG | GAAGGTTC | XM_163530 | Mus musculus LOC244133 (LOC244133), mRNA. | | 2.4 | SM2040 | h | 1 |

| | | | | | | | | | | | | | |
|---|---|---|---|---|---|---|---|---|---|---|---|---|---|
| mPool2_V2MM_16143 | 2.733176 | 2.205692 | TRUE | HP_23944 | TGCTGTTG | CTTCCAGT | NM_007520 | BTB and CNC homology 1 | Bach1 | 2.11 | SM2389 | e | 1 |
| mPool7_V2MM_196342 | 2.72588 | 2.205692 | TRUE | HP_41444 | TGCTGTTG | GGACTTTA | NM_177769 | hypothetical protein 4831417L10 | | 2.9 | SM2254 | h | 7 |
| 092308m1_V2MM_85041 | 2.725737 | 3.004939 | TRUE | HP_30653 | TGCTGTTG | GTGTTCTC | NM_030139 | zinc finger protein 449 | Zfp449 | 2.6 | SM2113 | f | 4 |
| 092308m1_V2MM_138848 | 2.720962 | 2.205692 | TRUE | HP_35956 | TGCTGTTG | GACATTAA | XM_154566 | Mus musculus LOC238925 (LOC238925), mR. | | 2.4 | SM2030 | d | 11 |
| 092308m1_V2MM_138848 | 2.720962 | 2.205692 | TRUE | HP_35956 | TGCTGTTG | GACATTAA | XM_154566 | Mus musculus LOC238925 (LOC238925), mRNA. | | 2.4 | SM2030 | d | 11 |
| mPool7_V2MM_100846 | 2.716422 | 2.205692 | TRUE | HP_32202 | TGCTGTTG | GACTGTAT | XM_127139 | echinoderm microtubule associated protein like 1 | Eml1 | 2.1 | SM2310 | c | 10 |
| mPool7_V2MM_80549 | 2.715909 | 2.205692 | TRUE | HP_30217 | TGCTGTTG | GGAAATGT | NM_011192 | proteaseome (prosome, macropain) 28 subunit, 3 | Psme3 | 2.8 | SM2227 | b | 6 |
| 092308m1_V2MM_83476 | 2.7159 | 3.004939 | TRUE | HP_30501 | TGCTGTTG | CACTTTAT | NM_017368 | CUG triplet repeat, R binding protein 1 | Cugbp1 | 2.6 | SM2112 | e | 2 |
| 092308m1_V2MM_159791 | 2.714818 | 2.205692 | TRUE | HP_38038 | TGCTGTTG | GTTTCTAT | AK081652 | pam, highwire, rpm 1 | Phr1 | 2.6 | SM2116 | d | 3 |
| 092308m1_V2MM_73747 | 2.710122 | 2.205692 | TRUE | HP_29557 | TGCTGTTG | GTCAGGTT | NM_146808 | olfactory receptor 1240 | Olfr1240 | 2.3 | SM2015 | b | 2 |
| 092308m1_V2MM_170566 | 2.708142 | 1.959154 | TRUE | HP_39105 | TGCTGTTG | GCGTTTGT | XM_286444 | Mus musculus hypothetical gene supported by AK028654 (LOC328131), mR. | | 2.5 | SM2083 | b | 7 |
| 092308m1_V2MM_177051 | 2.695729 | 3.004939 | TRUE | HP_39753 | TGCTGTTG | GAAGTATT | AK047086 | hypothetical protein B930018H19 | | 2.5 | SM2092 | f | 7 |
| 092308m1_V2MM_132337 | 2.694129 | 3.004939 | TRUE | HP_35309 | TGCTGTTG | CTCCTTCT | XM_148441 | RIKEN cD 5730478M09 gene | 5730478M09Rik | 2.6 | SM2103 | g | 11 |
| mPool2_V2MM_100946 | 2.69225 | 2.205692 | TRUE | HP_32212 | TGCTGTTG | CATTCTTA | XM_127279 | RIKEN cDNA 1110037N09 gene | 1110037N09Rik | 2.6 | SM2149 | b | 8 |
| mPool7_V2MM_74811 | 2.690379 | 2.205692 | TRUE | HP_29660 | TGCTGTTG | CAGGTTTG | BC005723 | RIKEN cDNA 1700029K01 gene | 1700029K01Rik | 2.8 | SM2246 | e | 8 |
| 092308m1_V2MM_178563 | 2.684133 | 1.959154 | TRUE | HP_39903 | TGCTGTTG | CAATTTAA | XM_288183 | Mus musculus LOC329171 (LOC329171), mR. | | 2.5 | SM2082 | d | 4 |
| mPool2_V2MM_153839 | 2.680466 | 3.004939 | TRUE | HP_37448 | TGCTGTTG | GCTTGGGA | BC030424 | A930003O13 | | 2.6 | SM2132 | h | 3 |
| mPool2_V2MM_213746 | 2.672153 | 3.004939 | TRUE | HP_43136 | TGCTGTTG | CTGGTTAA | XM_149205 | ring finger protein 24 | Rnf24 | 2.11 | SM2371 | a | 9 |
| 092308m1_V2MM_87899 | 2.671989 | 2.205692 | TRUE | HP_30934 | TGCTGTTG | GGCATTGC | NM_175270 | RIKEN cD 5730467H21 gene | 5730467H21Rik | 2.6 | SM2106 | c | 10 |
| mPool2_V2MM_145984 | 2.667191 | 2.205692 | TRUE | HP_36669 | TGCTGTTG | CCATGTCT | XM_161687 | Mus musculus LOC242888 (LOC242888), mRNA. | | 2.1 | SM2345 | h | 4 |
| mPool7_V2MM_96447 | 2.66463 | 2.205692 | TRUE | HP_31770 | TGCTGTTG | GATTTCTG | XM_110690 | RIKEN cDNA 1810033B17 gene | 1810033B17Rik | 2.9 | SM2253 | d | 11 |
| mPool7_V2MM_96447 | 2.66463 | 2.205692 | TRUE | HP_31770 | TGCTGTTG | GATTTCTG | XM_110690 | RIKEN cD 1810033B17 gene | 1810033B17Rik | 2.9 | SM2253 | d | 11 |
| mPool2_V2MM_119501 | 2.661508 | 1.959154 | TRUE | HP_34041 | TGCTGTTG | GTATTTCA | XM_140461 | RIKEN cDNA 2410072D24 gene | 2410072D24Rik | 2.11 | SM2361 | h | 1 |
| mPool2_V2MM_136543 | 2.661019 | 2.205692 | TRUE | HP_35726 | TGCTGTTG | GTTATCTT | XM_151819 | Mus musculus LOC213261 (LOC213261), mRNA. | | 2.4 | SM2034 | c | 9 |
| mPool2_V2MM_136543 | 2.661019 | 2.205692 | TRUE | HP_35726 | TGCTGTTG | GTTATCTT | XM_151819 | Mus musculus LOC213261 (LOC213261), mR. | | 2.4 | SM2034 | c | 9 |
| mPool7_V2MM_110292 | 2.660996 | 0 | TRUE | NA | NA | NA | NA | NA | NA | NA | NA | NA | NA |
| mPool2_V2MM_120180 | 2.659371 | 3.004939 | TRUE | HP_34108 | TGCTGTTG | CTCTCCTA | XM_140790 | similar to Rpl7a protein | | 2.6 | SM2110 | a | 12 |
| mPool2_V2MM_120180 | 2.659371 | 3.004939 | TRUE | HP_34108 | TGCTGTTG | CTCTCCTA | XM_140790 | similar to Rpl7a protein | | 2.6 | SM2110 | a | 12 |
| 092308m1_V2MM_142189 | 2.654747 | 2.205692 | TRUE | HP_36290 | TGCTGTTG | GCTAATTT | XM_157391 | Mus musculus LOC240317 (LOC240317), mR. | | 2.4 | SM2032 | g | 8 |
| 092308m1_V2MM_142189 | 2.654747 | 2.205692 | TRUE | HP_36290 | TGCTGTTG | GCTAATTT | XM_157391 | Mus musculus LOC240317 (LOC240317), mRNA. | | 2.4 | SM2032 | g | 8 |
| 092308m1_V2MM_141892 | 2.654311 | 2.205692 | TRUE | HP_36260 | TGCTGTTG | CTTTATCC | XM_157111 | Mus musculus LOC212310 (LOC212310), mR. | | 2.4 | SM2046 | b | 11 |
| mPool2_V2MM_113323 | 2.650507 | 3.004939 | TRUE | HP_33431 | TGCTGTTG | CCCAGAAC | XM_136918 | Mus musculus similar to glyceraldehyde-3-phosphate dehydrogenase [Mus musc | | 2.7 | SM2154 | e | 9 |
| mPool7_V2MM_196482 | 2.648947 | 3.004939 | TRUE | HP_41458 | TGCTGTTG | CTGTCTTT | XM_125673 | CXXC finger 6 | Cxxc6 | 2.9 | SM2266 | g | 2 |
| mPool7_V2MM_113323 | 2.64584 | 3.004939 | TRUE | HP_33431 | TGCTGTTG | CCCAGAAC | XM_136918 | Mus musculus similar to glyceraldehyde-3-phosphate dehydrogenase [Mus musc | | 2.7 | SM2154 | e | 9 |
| 092308m1_V2MM_176355 | 2.645547 | 2.205692 | TRUE | HP_39683 | TGCTGTTG | CTCTGTTT | XM_489154 | RIKEN cD 9330102E08 gene | 9330102E08Rik | 2.5 | SM2065 | b | 4 |
| mPool2_V2MM_144423 | 2.642873 | 3.004939 | TRUE | HP_36513 | TGCTGTTG | GAATTGGA | XM_159993 | Mus musculus LOC241988 (LOC241988), mRNA. | | 2.1 | SM2344 | h | 1 |
| mPool2_V2MM_124466 | 2.637419 | 3.004939 | TRUE | HP_34531 | TGCTGTTG | CGGCAATT | XM_143471 | potassium voltage-gated channel, shaker-related subfam | Kcna10 | 2.11 | SM2357 | c | 5 |
| mPool2_V2MM_13552 | 2.621147 | 3.004939 | TRUE | HP_23691 | TGCTGTTG | CATTTGTA | NM_026405 | RAB32, member RAS oncogene family | Rab32 | 2.11 | SM2380 | b | 3 |
| mPool2_V2MM_32012 | 2.620331 | 3.004939 | TRUE | HP_25490 | TGCTGTTG | GATGGAAC | NM_023500 | McLeod syndrome gene homolog | Xkh | 2.11 | SM2389 | a | 12 |
| mPool2_V2MM_124810 | 2.612072 | 3.004939 | TRUE | HP_34565 | TGCTGTTG | CTTTCAAA | XM_143641 | Mus musculus similar to TRAV19 [Mus musculus] (LOC230001), mRNA. | | 2.11 | SM2352 | d | 7 |
| 092308m1_V2MM_91867 | 2.607876 | 2.205692 | TRUE | HP_12628 | TGCTGTTG | CCTACAAA | NM_178607 | ring finger protein 24 | Rnf24 | 2.6 | SM2109 | h | 2 |
| 092308m1_V2MM_91867 | 2.607876 | 2.205692 | TRUE | HP_12628 | TGCTGTTG | CCTACAAA | NM_178607 | ring finger protein 24 | Rnf24 | 2.6 | SM2109 | h | 2 |
| 092308m1_V2MM_98561 | 2.596861 | 2.205692 | TRUE | HP_31978 | TGCTGTTG | CGTATTTG | XM_112650 | aquaporin 9 | Aqp9 | 2.6 | SM2112 | h | 8 |
| mPool7_V2MM_172099 | 2.592874 | 2.205692 | TRUE | HP_39258 | TGCTGTTG | CATTCTGA | XM_286756 | Mus musculus hypothetical gene supported by AK048704 (LOC328685), mRNA. | | 2.5 | SM2090 | h | 5 |
| mPool7_V2MM_172099 | 2.592874 | 2.205692 | TRUE | HP_39258 | TGCTGTTG | CATTCTGA | XM_286756 | Mus musculus hypothetical gene supported by AK048704 (LOC328685), mR. | | 2.5 | SM2090 | h | 5 |
| 092308m1_V2MM_150902 | 2.591938 | 2.205692 | TRUE | HP_37158 | TGCTGTTG | CTGAGTTA | BC059023 | expressed sequence AW060220 | AW060220 | 2.4 | SM2037 | b | 11 |
| 092308m1_V2MM_152512 | 2.591886 | 2.205692 | TRUE | HP_37317 | TGCTGTTG | GCCACTTC | XM_195409 | Mus musculus similar to protease [Trichosurus vulpecula] (LOC270243), mR. | | 2.4 | SM2030 | c | 4 |
| 092308m1_V2MM_78324 | 2.59181 | 2.205692 | TRUE | HP_30001 | TGCTGTTG | GACCCAAT | NM_021557 | retinol dehydrogese 11 | Rdh11 | 2.3 | SM2007 | g | 7 |
| mPool2_V2MM_208908 | 2.588866 | 3.004939 | TRUE | HP_42668 | TGCTGTTG | CATGTTAC | XM_163904 | Mus musculus LOC244280 (LOC244280), mRNA. | | 2.1 | SM2336 | f | 5 |
| mPool7_V2MM_184400 | 2.583425 | 1.959154 | TRUE | HP_40487 | TGCTGTTG | CCCATTTA | XM_289192 | Mus musculus LOC332813 (LOC332813), mRNA. | | 2.9 | SM2295 | c | 5 |
| 092308m1_V2MM_154074 | 2.581221 | 3.004939 | TRUE | HP_37471 | TGCTGTTG | CCTGTTTA | NM_175535 | Rho GTPase activating protein 20 | Arhgap20 | 2.6 | SM2114 | f | 6 |
| mPool7_V2MM_107136 | 2.566738 | 2.205692 | TRUE | HP_32822 | TGCTGTTG | CTGTTTAC | BC060239 | RIKEN cDNA 2610109H07 gene | 2610109H07Rik | 2.1 | SM2313 | c | 8 |
| 092308m3_V2MM_98132 | 2.560958 | 75.83547 | TRUE | HP_31936 | TGCTGTTG | CCCTTTGG | XM_194868 | Mus musculus similar to hypothetical protein [Plasmodium yoelii yoelii] (LOC268 | | 2.13 | SM2472 | g | 11 |

| | | | | | | | | | | | | | |
|---|---|---|---|---|---|---|---|---|---|---|---|---|---|
| 092308m1_V2MM_173028 | 2.560137 | 3.004939 | TRUE | HP_39351 | TGCTGTTG | GATCCAA | XM_286944 | Mus musculus hypothetical gene supported by AK089863 (LOC329049), mR. | | 2.5 | SM2067 | b | 6 |
| 092308m1_V2MM_63008 | 2.556561 | 2.205692 | TRUE | HP_28509 | TGCTGTTG | CCGATATT | NM_013818 | GTP binding protein 1 | Gtpbp1 | 2.4 | SM2025 | c | 6 |
| mPool7_V2MM_94358 | 2.544241 | 3.004939 | TRUE | HP_31565 | TGCTGTTG | GTGGTAA | NM_183137 | RIKEN cDNA 2410002I01 gene | 2410002I01Rik | 2.8 | SM2249 | g | 8 |
| mPool2_V2MM_21743 | 2.543583 | 3.004939 | TRUE | HP_24491 | TGCTGTTG | CATTTCTT | NM_145586 | RIKEN cDNA 8430420C20 gene | 8430420C20Rik | 2.11 | SM2389 | f | 9 |
| mPool2_V2MM_87747 | 2.543217 | 3.004939 | TRUE | HP_30919 | TGCTGTTG | CCAACTCA | NM_175236 | alcohol dehydrogese, iron containing, 1 | Adhfe1 | 2.6 | SM2108 | e | 12 |
| mPool2_V2MM_87747 | 2.543217 | 3.004939 | TRUE | HP_30919 | TGCTGTTG | CCAACTCA | NM_175236 | alcohol dehydrogenase, iron containing, 1 | Adhfe1 | 2.6 | SM2108 | e | 12 |
| mPool2_V2MM_5090 | 2.537657 | 1.959154 | TRUE | HP_22864 | TGCTGTTG | CATCTACA | NM_010114 | kallikrein 22 | Klk22 | 2.11 | SM2384 | d | 6 |
| mPool7_V2MM_171247 | 2.534961 | 2.205692 | TRUE | HP_39173 | TGCTGTTG | CCTTTATA | XM_286584 | hypothetical LOC328369 | | 2.5 | SM2080 | h | 2 |
| mPool7_V2MM_171247 | 2.534961 | 2.205692 | TRUE | HP_39173 | TGCTGTTG | CCTTTATA | XM_286584 | hypothetical LOC328369 | | 2.5 | SM2080 | h | 2 |
| mPool2_V2MM_8473 | 2.534002 | 2.205692 | TRUE | HP_23195 | TGCTGTTG | GTCTTTAT | NM_008148 | glycoprotein 5 (platelet) | Gp5 | 2.11 | SM2377 | d | 7 |
| 092308m1_V2MM_180103 | 2.533756 | 3.004939 | TRUE | HP_40057 | TGCTGTTG | GTTGATTT | XM_288450 | Mus musculus LOC331788 (LOC331788), mR. | | 2.5 | SM2082 | h | 7 |
| 092308m1_V2MM_145554 | 2.532402 | 2.205692 | TRUE | HP_36626 | TGCTGTTG | CATTCTTA | XM_161053 | Mus musculus LOC230170 (LOC230170), mRNA. | | 2.4 | SM2040 | e | 9 |
| 092308m1_V2MM_145554 | 2.532402 | 2.205692 | TRUE | HP_36626 | TGCTGTTG | CATTCTTA | XM_161053 | Mus musculus LOC230170 (LOC230170), mR. | | 2.4 | SM2040 | e | 9 |
| mPool7_V2MM_176389 | 2.532005 | 3.004939 | TRUE | HP_39687 | TGCTGTTG | CCTTAAAT | AK129335 | RIKEN cD C230080I20 gene | C230080I20Rik | 2.5 | SM2077 | f | 6 |
| mPool7_V2MM_176389 | 2.532005 | 3.004939 | TRUE | HP_39687 | TGCTGTTG | CCTTAAAT | AK129335 | RIKEN cDNA C230080I20 gene | C230080I20Rik | 2.5 | SM2077 | f | 6 |
| mPool7_V2MM_139283 | 2.530036 | 3.004939 | TRUE | HP_36000 | TGCTGTTG | CATTCTTT | XM_154903 | Mus musculus LOC210773 (LOC210773), mRNA. | | 2.1 | SM2327 | f | 8 |
| mPool2_V2MM_134455 | 2.529677 | 2.205692 | TRUE | HP_35518 | TGCTGTTG | GTACATCT | XM_149889 | Mus musculus LOC232903 (LOC232903), mRNA. | | 2.11 | SM2368 | h | 6 |
| mPool7_V2MM_114037 | 2.528273 | 2.205692 | TRUE | HP_33502 | TGCTGTTG | CCATTTGA | XM_137252 | Mus musculus similar to nuclear transport factor 2 [Rattus norvegicus] (LOC2375 | | 2.7 | SM2162 | b | 5 |
| mPool2_V2MM_12274 | 2.527311 | 3.004939 | TRUE | HP_23567 | TGCTGTTG | GTGGATG | NM_007658 | cell division cycle 25 homolog A (S. cerevisiae) | Cdc25a | 2.11 | SM2378 | c | 10 |
| mPool2_V2MM_8682 | 2.521539 | 3.004939 | TRUE | HP_23216 | TGCTGTTG | GTTAGCTT | NM_008258 | hematological and neurological expressed sequence 1 | Hn1 | 2.11 | SM2383 | a | 10 |
| mPool2_V2MM_91937 | 2.520823 | 2.205692 | TRUE | HP_31330 | TGCTGTTG | GTACTTGT | NM_178619 | RIKEN cDNA 1810026J23 gene | 1810026J23Rik | 2.6 | SM2138 | f | 1 |
| mPool2_V2MM_146410 | 2.520178 | 2.205692 | TRUE | HP_36712 | TGCTGTTG | CCTAAGTT | XM_161991 | Mus musculus LOC243104 (LOC243104), mRNA. | | 2.1 | SM2347 | d | 7 |
| mPool7_V2MM_89146 | 2.517894 | 3.004939 | TRUE | HP_31057 | TGCTGTTG | GCCCTAAC | NM_176942 | gamma-aminobutyric acid (GABA-A) receptor, subunit al | Gabra5 | 2.9 | SM2262 | b | 5 |
| 092308m1_V2MM_181334 | 2.512468 | 1.959154 | TRUE | HP_40180 | TGCTGTTG | CCTATTAC | XM_288658 | Mus musculus LOC328501 (LOC328501), mR. | | 2.5 | SM2085 | d | 11 |
| 092308m1_V2MM_181334 | 2.512468 | 1.959154 | TRUE | HP_40180 | TGCTGTTG | CCTATTAC | XM_288658 | Mus musculus LOC328501 (LOC328501), mRNA. | | 2.5 | SM2085 | d | 11 |
| mPool2_V2MM_90692 | 2.508323 | 3.004939 | TRUE | HP_31210 | TGCTGTTG | GGCACTA | AK033743 | lipocalin 6 | Lcn6 | 2.6 | SM2139 | h | 6 |
| mPool2_V2MM_114037 | 2.507135 | 2.205692 | TRUE | HP_33502 | TGCTGTTG | CCATTTGA | XM_137252 | Mus musculus similar to nuclear transport factor 2 [Rattus norvegicus] (LOC2375 | | 2.7 | SM2162 | b | 5 |
| mPool2_V2MM_127305 | 2.5064 | 2.205692 | TRUE | HP_34813 | TGCTGTTG | GTGAAGT | AK032959 | RIKEN cDNA A430107O13 gene | A430107O13Rik | 2.11 | SM2351 | f | 3 |
| 092308m1_V2MM_131144 | 2.505932 | 2.205692 | TRUE | HP_35191 | TGCTGTTG | CAGAAAT | XM_147444 | RIKEN cD 4930485B16 gene | 4930485B16Rik | 2.5 | SM2100 | a | 7 |
| 092308m1_V2MM_93146 | 2.505727 | 1.959154 | TRUE | HP_31447 | TGCTGTTG | GTGGATA | AK083705 | expressed sequence AI256775 | AI256775 | 2.6 | SM2110 | a | 2 |
| 092308m1_V2MM_93146 | 2.505727 | 1.959154 | TRUE | HP_31447 | TGCTGTTG | GTGGATA | AK083705 | expressed sequence AI256775 | AI256775 | 2.6 | SM2110 | a | 2 |
| 092308m1_V2MM_154975 | 2.502832 | 2.205692 | TRUE | HP_37561 | TGCTGTTG | GTGAACT | XM_197114 | hypothetical LOC380894 | | 2.6 | SM2122 | e | 1 |
| mPool7_V2MM_63851 | 2.498773 | 2.205692 | TRUE | HP_28591 | TGCTGTTG | CTCATTAT | NM_146868 | olfactory receptor 894 | Olfr894 | 2.8 | SM2239 | f | 10 |
| 092308m1_V2MM_184354 | 2.48845 | 2.205692 | TRUE | HP_40482 | TGCTGTTG | CAGCATTT | XM_289184 | Mus musculus LOC332804 (LOC332804), mRNA. | | 2.5 | SM2069 | f | 4 |
| 092308m1_V2MM_184354 | 2.48845 | 2.205692 | TRUE | HP_40482 | TGCTGTTG | CAGCATTT | XM_289184 | Mus musculus LOC332804 (LOC332804), mR. | | 2.5 | SM2069 | f | 4 |
| mPool7_V2MM_171879 | 2.483358 | 1.959154 | TRUE | HP_39236 | TGCTGTTG | CATTGCTC | AK078693 | RIKEN cDNA 7530414M10 gene | 7530414M10Rik | 2.9 | SM2272 | a | 11 |
| mPool2_V2MM_82528 | 2.478454 | 3.004939 | TRUE | HP_30410 | TGCTGTTG | GAGCTATT | NM_009700 | aquaporin 4 | Aqp4 | 2.7 | SM2164 | g | 12 |
| mPool2_V2MM_82528 | 2.478454 | 3.004939 | TRUE | HP_30410 | TGCTGTTG | GAGCTATT | NM_009700 | aquaporin 4 | Aqp4 | 2.7 | SM2164 | g | 12 |
| mPool2_V2MM_5145 | 2.478236 | 3.004939 | TRUE | HP_22870 | TGCTGTTG | CATTGTAA | NM_027398 | Kv channel-interacting protein 1 | Kcnip1 | 2.11 | SM2389 | c | 1 |
| mPool2_V2MM_120050 | 2.477423 | 3.004939 | TRUE | HP_34095 | TGCTGTTG | CTAAACAT | XM_140736 | Mus musculus similar to pol protein [Homo sapiens] (LOC226212), mRNA. | | 2.11 | SM2363 | h | 7 |
| 092308m1_V2MM_179548 | 2.465744 | 3.004939 | TRUE | HP_40002 | TGCTGTTG | CCTACAAT | XM_288352 | Mus musculus LOC331672 (LOC331672), mRNA. | | 2.5 | SM2091 | d | 5 |
| 092308m1_V2MM_179548 | 2.465744 | 3.004939 | TRUE | HP_40002 | TGCTGTTG | CCTACAAT | XM_288352 | Mus musculus LOC331672 (LOC331672), mR. | | 2.5 | SM2091 | d | 5 |
| mPool7_V2MM_205982 | 2.462338 | 0 | TRUE | HP_42382 | TGCTGTTG | GTTGAAA | BC054817 | RIKEN cDNA 1810011E08 gene | 1810011E08Rik | 2.9 | SM2299 | b | 1 |
| mPool7_V2MM_84945 | 2.461374 | 2.205692 | TRUE | HP_30644 | TGCTGTTG | CTAATTCC | NM_029891 | RIKEN cDNA 9430034D17 gene | 9430034D17Rik | 2.9 | SM2254 | d | 4 |
| 092308m1_V2MM_128357 | 2.455774 | 3.004939 | TRUE | HP_34917 | TGCTGTTG | CCCATTAC | XM_145657 | Mus musculus similar to membrane and microfilament-associated protein p58 [R | | 2.6 | SM2106 | c | 2 |
| 092308m1_V2MM_178888 | 2.451542 | 3.004939 | TRUE | HP_39936 | TGCTGTTG | CTGATTAT | XM_288237 | Mus musculus LOC332503 (LOC332503), mR. | | 2.5 | SM2086 | e | 9 |
| 092308m1_V2MM_167815 | 2.446867 | 2.205692 | TRUE | HP_38832 | TGCTGTTG | CCACTTCA | XM_285849 | Mus musculus similar to protease [Trichosurus vulpecula] (LOC329987), mR. | | 2.6 | SM2123 | d | 8 |
| 092308m1_V2MM_177650 | 2.445472 | 2.205692 | TRUE | HP_39813 | TGCTGTTG | GTCCTTGA | AK084093 | RIKEN cD D130086K05 gene | D130086K05Rik | 2.5 | SM2095 | a | 3 |
| 092308m1_V2MM_179034 | 2.443166 | 3.004939 | TRUE | HP_39950 | TGCTGTTG | CAGGATA | XM_288262 | Mus musculus LOC329282 (LOC329282), mRNA. | | 2.5 | SM2089 | b | 6 |
| 092308m1_V2MM_179034 | 2.443166 | 3.004939 | TRUE | HP_39950 | TGCTGTTG | CAGGATA | XM_288262 | Mus musculus LOC329282 (LOC329282), mR. | | 2.5 | SM2089 | b | 6 |
| 092308m1_V2MM_64431 | 2.439037 | 3.004939 | TRUE | HP_28648 | TGCTGTTG | CTGGGAG | XM_195585 | Mus musculus similar to MAP/microtubule affinity-regulating kise like 1; MARK4 | | 2.4 | SM2020 | c | 12 |
| mPool7_V2MM_184599 | 2.437695 | 2.205692 | TRUE | HP_40507 | TGCTGTTG | CACTTTCT | XM_289225 | Mus musculus LOC332850 (LOC332850), mRNA. | | 2.9 | SM2290 | e | 12 |
| mPool7_V2MM_184599 | 2.437695 | 2.205692 | TRUE | HP_40507 | TGCTGTTG | CACTTTCT | XM_289225 | Mus musculus LOC332850 (LOC332850), mR. | | 2.9 | SM2290 | e | 12 |

| | | | | | | | | | | | | | |
|---|---|---|---|---|---|---|---|---|---|---|---|---|---|
| 092308m1_V2MM_80443 | 2.43119 | 3.004939 | TRUE | HP_30207 | TGCTGTTG | GCTTAAAC | NM_009850 | CD3 antigen, gamma polypeptide | Cd3g | 2.4 | SM2024 | f | 9 |
| mPool2_V2MM_30861 | 2.430164 | 2.205692 | TRUE | HP_25377 | TGCTGTTG | CTCAGAAT | NM_015825 | SH3-binding domain glutamic acid-rich protein | Sh3bgr | 2.11 | SM2390 | f | 9 |
| mPool2_V2MM_123245 | 2.427093 | 2.205692 | TRUE | HP_34410 | TGCTGTTG | CTTATATT | XM_142826 | Mus musculus similar to major allergen 1-like; lacrimal gland protein [Mus muscu | | 2.11 | SM2353 | f | 2 |
| mPool7_V2MM_101648 | 2.426232 | 3.004939 | TRUE | HP_32281 | TGCTGTTG | CCTTTGAT | XM_127760 | Mus musculus neurofilament, light polypeptide (Nefl), mRNA. | | 2.1 | SM2301 | d | 3 |
| 092308m1_V2MM_167262 | 2.425224 | 3.004939 | TRUE | HP_38777 | TGCTGTTG | CACATTCC | XM_485247 | similar to arylacetamide deacetylase | | 2.6 | SM2124 | d | 6 |
| mPool7_V2MM_66116 | 2.422509 | 3.004939 | TRUE | HP_28813 | TGCTGTTG | GTGTGGA | NM_011210 | protein tyrosine phosphatase, receptor type, C | Ptprc | 2.8 | SM2242 | c | 4 |
| 092308m1_V2MM_153010 | 2.409313 | 3.004939 | TRUE | HP_37367 | TGCTGTTG | CTGCTTAA | XM_195653 | Mus musculus similar to Amphoterin (LOC272391), mR. | | 2.6 | SM2117 | c | 4 |
| mPool2_V2MM_3920 | 2.4076 | 2.205692 | TRUE | HP_22750 | TGCTGTTG | GGACTTGT | NM_010836 | homeo box, msh-like 3 | Msx3 | 2.11 | SM2377 | h | 12 |
| 092308m1_V2MM_136213 | 2.405075 | 2.205692 | TRUE | HP_35693 | TGCTGTTG | CTACGAAA | XM_151496 | Mus musculus LOC241166 (LOC241166), mR. | | 2.5 | SM2052 | b | 2 |
| mPool4_V2MM_105816 | 2.403439 | 75.83547 | TRUE | HP_32692 | TGCTGTTG | CCATTGGA | XM_130816 | similar to Retrovirus-related POL polyprotein | | 2.16 | SM2625 | e | 4 |
| mPool2_V2MM_84527 | 2.397941 | 3.004939 | TRUE | HP_30603 | TGCTGTTG | CATAGTAA | NM_028259 | ribosomal protein S6 kinase, polypeptide 1 | Rps6kb1 | 2.6 | SM2146 | g | 12 |
| 092308m1_V2MM_171406 | 2.396564 | 2.205692 | TRUE | HP_39189 | TGCTGTTG | CTCGTTGA | XM_286618 | hypothetical protein E030037F02 | | 2.5 | SM2073 | b | 12 |
| 092308m1_V2MM_66137 | 2.393384 | 3.004939 | TRUE | HP_28815 | TGCTGTTG | GTTAATAC | NM_025741 | RIKEN cD 4931412G03 gene | 4931412G03Rik | 2.4 | SM2027 | e | 10 |
| mPool7_V2MM_174664 | 2.390888 | 3.004939 | TRUE | HP_39514 | TGCTGTTG | CTGTATTC | AK028474 | hypothetical protein 4631419I20 | | 2.9 | SM2276 | f | 9 |
| mPool7_V2MM_201943 | 2.387435 | 2.205692 | TRUE | HP_41996 | TGCTGTTG | GGGTTTAC | AK052405 | RIKEN cDNA C130089K02 gene | C130089K02Rik | 2.9 | SM2291 | d | 8 |
| 092308m1_V2MM_171247 | 2.385208 | 2.205692 | TRUE | HP_39173 | TGCTGTTG | CCTTTATA | XM_286584 | hypothetical LOC328369 | | 2.5 | SM2080 | h | 2 |
| 092308m1_V2MM_171247 | 2.385208 | 2.205692 | TRUE | HP_39173 | TGCTGTTG | CCTTTATA | XM_286584 | hypothetical LOC328369 | | 2.5 | SM2080 | h | 2 |
| 092308m1_V2MM_185463 | 2.385014 | 3.004939 | TRUE | HP_40593 | TGCTGTTG | GTCTAAGT | XM_289375 | Mus musculus LOC333041 (LOC333041), mR. | | 2.5 | SM2062 | c | 12 |
| 092308m1_V2MM_185463 | 2.385014 | 3.004939 | TRUE | HP_40593 | TGCTGTTG | GTCTAAGT | XM_289375 | Mus musculus LOC333041 (LOC333041), mRNA. | | 2.5 | SM2062 | c | 12 |
| mPool7_V2MM_90186 | 2.381069 | 1.959154 | TRUE | HP_31160 | TGCTGTTG | CATCCAGA | NM_177750 | hypothetical protein B930041G04 | | 2.9 | SM2251 | e | 3 |
| mPool2_V2MM_129470 | 2.377483 | 3.004939 | TRUE | HP_35027 | TGCTGTTG | CTGAGTAT | XM_486101 | similar to hypothetical protein FLJ25801 | | 2.1 | SM2350 | g | 6 |
| mPool2_V2MM_210414 | 2.37623 | 3.004939 | TRUE | HP_42816 | TGCTGTTG | CTAAATAC | XM_143858 | similar to Elongation factor 1-alpha 1 (EF-1-alpha-1) (Elongation factor 1 A-1) (eEl | | 2.11 | SM2367 | h | 11 |
| 092308m1_V2MM_165904 | 2.373889 | 2.205692 | TRUE | HP_38643 | TGCTGTTG | CGCTATGT | XM_285354 | olfactory receptor 1065 | Olfr1065 | 2.6 | SM2123 | c | 6 |
| 092308m1_V2MM_62055 | 2.373677 | 3.004939 | TRUE | HP_28417 | TGCTGTTG | CACCGTCT | NM_011059 | peptidyl arginine deimise, type I | Padi1 | 2.3 | SM2018 | h | 4 |
| mPool2_V2MM_1570 | 2.373061 | 3.004939 | TRUE | HP_22520 | TGCTGTTG | CCTTAAAT | NM_025818 | RIKEN cDNA 1200014J11 gene | 1200014J11Rik | 2.11 | SM2384 | f | 9 |
| 092308m1_V2MM_131266 | 2.372112 | 3.004939 | TRUE | HP_35203 | TGCTGTTG | CAGCTAAC | NM_213729 | expressed sequence AI842396 | AI842396 | 2.5 | SM2097 | g | 3 |
| mPool2_V2MM_6688 | 2.371223 | 2.205692 | TRUE | HP_23021 | TGCTGTTG | CTTGATTT | NM_054094 | butyryl Coenzyme A synthetase 1 | Bucs1 | 2.11 | SM2389 | g | 2 |
| mPool7_V2MM_199554 | 2.366627 | 3.004939 | TRUE | HP_41759 | TGCTGTTG | CTCATTAT | XM_287452 | Mus musculus hypothetical gene supported by AK035988 (LOC330083), mRNA. | | 2.9 | SM2272 | b | 9 |
| mPool2_V2MM_157089 | 2.366555 | 3.004939 | TRUE | HP_37771 | TGCTGTTG | CTTGGCAT | XM_204855 | Mus musculus similar to hypothetical protein D930020E02 (LOC277170), mRNA. | | 2.6 | SM2127 | h | 10 |
| mPool7_V2MM_86833 | 2.364566 | 1.959154 | TRUE | HP_30830 | TGCTGTTG | GTTATTAA | NM_173783 | RIKEN cDNA B430216B18 gene | B430216B18Rik | 2.9 | SM2265 | f | 2 |
| mPool2_V2MM_212162 | 2.361153 | 2.205692 | TRUE | HP_42983 | TGCTGTTG | CCAGATTT | XM_141639 | Mus musculus similar to protease [Mus musculus] (LOC236704), mRNA. | | 2.11 | SM2362 | c | 2 |
| mPool7_V2MM_113519 | 2.360552 | 3.004939 | TRUE | HP_33450 | TGCTGTTG | CTTGCTCT | XM_137005 | Mus musculus similar to Hypothetical protein KIAA0144 (LOC212406), mRNA. | | 2.7 | SM2154 | f | 2 |
| 092308m1_V2MM_176364 | 2.353708 | 2.205692 | TRUE | HP_39684 | TGCTGTTG | CATTCATT | XM_287651 | Mus musculus hypothetical gene supported by AK079645 (LOC330422), mR. | | 2.5 | SM2079 | b | 1 |
| 092308m1_V2MM_176364 | 2.353708 | 2.205692 | TRUE | HP_39684 | TGCTGTTG | CATTCATT | XM_287651 | Mus musculus hypothetical gene supported by AK079645 (LOC330422), mRNA. | | 2.5 | SM2079 | b | 1 |
| mPool2_V2MM_162783 | 2.353143 | 1.959154 | TRUE | HP_38334 | TGCTGTTG | CTTTGGAA | XM_284493 | Mus musculus hypothetical gene supported by AK083277 (LOC330932), mRNA. | | 2.6 | SM2130 | e | 8 |
| mPool5_V2MM_14453 | 2.352201 | 70.87591 | TRUE | HP_23779 | TGCTGTTG | CCTCCCAC | NM_025875 | RNA binding motif protein 8a | Rbm8a | NA | | | NA |
| mPool2_V2MM_1149 | 2.350584 | 2.205692 | TRUE | HP_22479 | TGCTGTTG | CATCAACT | NM_011721 | Werner syndrome homolog (human) | Wrn | 2.11 | SM2385 | a | 3 |
| mPool7_V2MM_197705 | 2.344005 | 3.004939 | TRUE | HP_35074 | TGCTGTTG | GCGAGTAT | XM_356738 | similar to T-cell receptor alpha variable region family 13 subfamily 1 | | 2.9 | SM2263 | h | 6 |
| mPool2_V2MM_121994 | 2.343889 | 2.205692 | TRUE | HP_34286 | TGCTGTTG | CAGTTATA | NM_001003916 | cDNA sequence AK129302 | AK129302 | 2.1 | SM2350 | g | 2 |
| mPool2_V2MM_121891 | 2.34312 | 3.004939 | TRUE | HP_34276 | TGCTGTTG | CTGAGTCT | XM_141921 | gene model 370, (NCBI) | Gm370 | 2.1 | SM2350 | f | 6 |
| mPool2_V2MM_209788 | 2.342931 | 2.205692 | TRUE | HP_42755 | TGCTGTTG | CCAATTTC | XM_163354 | Mus musculus LOC215451 (LOC215451), mRNA. | | 2.1 | SM2336 | g | 12 |
| mPool2_V2MM_142189 | 2.340758 | 1.959154 | TRUE | HP_36290 | TGCTGTTG | GCTAATTT | XM_157391 | Mus musculus LOC240317 (LOC240317), mRNA. | | 2.4 | SM2032 | g | 8 |
| mPool2_V2MM_142189 | 2.340758 | 1.959154 | TRUE | HP_36290 | TGCTGTTG | GCTAATTT | XM_157391 | Mus musculus LOC240317 (LOC240317), mR. | | 2.4 | SM2032 | g | 8 |
| mPool2_V2MM_5233 | 2.336743 | 2.205692 | TRUE | HP_22879 | TGCTGTTG | CCACTGTT | NM_025806 | RIKEN cDNA 1100001H23 gene | 1100001H23Rik | 2.11 | SM2383 | d | 4 |
| mPool2_V2MM_141349 | 2.327546 | 3.004939 | TRUE | HP_36206 | TGCTGTTG | GAGAAATT | XM_156599 | Mus musculus LOC239923 (LOC239923), mRNA. | | 2.1 | SM2345 | h | 9 |
| 092308m1_V2MM_169742 | 2.326999 | 3.004939 | TRUE | HP_39023 | TGCTGTTG | CTCTCTCT | AK081287 | hypothetical protein C030046G05 | | 2.6 | SM2119 | g | 1 |
| mPool7_V2MM_178545 | 2.31982 | 2.205692 | TRUE | HP_39901 | TGCTGTTG | GCTTAGAT | XM_288180 | Mus musculus LOC332444 (LOC332444), mR. | | 2.5 | SM2092 | f | 11 |
| mPool7_V2MM_178545 | 2.31982 | 2.205692 | TRUE | HP_39901 | TGCTGTTG | GCTTAGAT | XM_288180 | Mus musculus LOC332444 (LOC332444), mRNA. | | 2.5 | SM2092 | f | 11 |
| mPool2_V2MM_127435 | 2.319779 | 3.004939 | TRUE | HP_34826 | TGCTGTTG | CTAGGAAT | XM_145191 | Mus musculus similar to fatty acid binding protein ( heart ) like [Bos taurus] (LOC | | 2.11 | SM2359 | f | 7 |
| mPool4_V2MM_205800 | 2.318956 | 62.64426 | TRUE | HP_42365 | TGCTGTTG | GCCTGTTT | AK010497 | RIKEN cDNA 2410015J15 gene | 2410015J15Rik | 2.16 | SM2614 | h | 2 |
| mPool2_V2MM_21542 | 2.318763 | 3.004939 | TRUE | HP_24471 | TGCTGTTG | CAGTTGAC | NM_023530 | phospholipase A2, group XIIB | Pla2g12b | 2.11 | SM2392 | c | 7 |
| mPool2_V2MM_134459 | 2.317929 | 2.205692 | TRUE | HP_35518 | TGCTGTTG | CTAGAACT | XM_149889 | Mus musculus LOC232903 (LOC232903), mRNA. | | 2.11 | SM2354 | d | 11 |
| 092308m1_V2MM_127416 | 2.3129 | 2.205692 | TRUE | HP_34824 | TGCTGTTG | CAGAGACC | XM_145185 | similar to glyceraldehyde-3-phosphate dehydrogese (phosphorylating) (EC 1.2.1. | | 2.6 | SM2102 | b | 4 |

| | | | | | | | | | | | | | |
|---|---|---|---|---|---|---|---|---|---|---|---|---|---|
| mPool2_V2MM_131495 | 2.312512 | 2.205692 | TRUE | HP_35225 | TGCTGTTG | GTTATATA | XM_147679 | Mus musculus LOC209126 (LOC209126), mRNA. | | 2.11 | SM2353 | d | 4 |
| mPool7_V2MM_184354 | 2.311222 | 2.205692 | TRUE | HP_40482 | TGCTGTTG | CAGCATTT | XM_289184 | Mus musculus LOC332804 (LOC332804), mR. | | 2.5 | SM2069 | f | 4 |
| mPool7_V2MM_184354 | 2.311222 | 2.205692 | TRUE | HP_40482 | TGCTGTTG | CAGCATTT | XM_289184 | Mus musculus LOC332804 (LOC332804), mRNA. | | 2.5 | SM2069 | f | 4 |
| mPool2_V2MM_115702 | 2.309701 | 3.004939 | TRUE | HP_33666 | TGCTGTTG | GCTGGAA( | XM_138606 | similar to glyceraldehyde-3-phosphate dehydrogenase | | 2.1 | SM2323 | c | 1 |
| mPool2_V2MM_131249 | 2.309261 | 3.004939 | TRUE | HP_35201 | TGCTGTTG | GTTGATTA | AK015910 | RIKEN cDNA 4930527B16 gene | 4930527B16Rik | 2.1 | SM2347 | b | 8 |
| mPool2_V2MM_85581 | 2.308452 | 2.205692 | TRUE | HP_30706 | TGCTGTTG | GCTGAACT | NM_146417 | olfactory receptor 877 | Olfr877 | 2.6 | SM2143 | h | 4 |
| mPool2_V2MM_132910 | 2.3083 | 3.004939 | TRUE | HP_35365 | TGCTGTTG | GGCATTCT | BC020411 | RIKEN cDNA 2700010L10 gene | 2700010L10Rik | 2.11 | SM2370 | d | 10 |
| mPool7_V2MM_171663 | 2.308269 | 2.205692 | TRUE | HP_39215 | TGCTGTTG | CATATTCT | AK078525 | activin A receptor, type 1B | Acvr1b | 2.5 | SM2062 | a | 5 |
| mPool7_V2MM_171663 | 2.308269 | 2.205692 | TRUE | HP_39215 | TGCTGTTG | CATATTCT | AK078525 | activin A receptor, type 1B | Acvr1b | 2.5 | SM2062 | a | 5 |
| mPool2_V2MM_213136 | 2.305802 | 2.205692 | TRUE | HP_43078 | TGCTGTTG | CCTTGAAT | AK029061 | RIKEN cDNA 1700020O03 gene | 1700020O03Rik | 2.11 | SM2366 | f | 5 |
| mPool2_V2MM_101248 | 2.302802 | 2.205692 | TRUE | HP_32242 | TGCTGTTG | CTGACTTC | AK083705 | expressed sequence AI256775 | AI256775 | 2.7 | SM2157 | c | 1 |
| mPool2_V2MM_5330 | 2.299528 | 3.004939 | TRUE | HP_22888 | TGCTGTTG | CCCAAGAT | NM_007560 | bone morphogenetic protein receptor, type 1B | Bmpr1b | 2.11 | SM2378 | e | 1 |
| mPool2_V2MM_164305 | 2.297209 | 3.004939 | TRUE | HP_38485 | TGCTGTTG | CTGGATTG | XM_284973 | Mus musculus similar to thyroid hormone receptor-associated protein, 150 kDa s | | 2.6 | SM2130 | f | 7 |
| 092308m1_V2MM_95502 | 2.297144 | 2.205692 | TRUE | HP_31677 | TGCTGTTG | GGCAATTT | NM_198017 | RIKEN cD C430003P19 gene | C430003P19Rik | 2.6 | SM2112 | h | 5 |
| 092308m1_V2MM_174732 | 2.294221 | 2.205692 | TRUE | HP_39521 | TGCTGTTG | GCAGTAA/ | AK034327 | hypothetical protein 9330178D15 | | 2.9 | SM2277 | f | 7 |
| 092308m1_V2MM_174732 | 2.294221 | 2.205692 | TRUE | HP_39521 | TGCTGTTG | GCAGTAA/ | AK034327 | hypothetical protein 9330178D15 | | 2.9 | SM2277 | f | 7 |
| 092308m1_V2MM_175229 | 2.294108 | 3.004939 | TRUE | HP_39571 | TGCTGTTG | CATCGAA( | XM_287395 | Mus musculus hypothetical gene supported by AK038877 (LOC329949), mR. | | 2.5 | SM2071 | b | 4 |
| 092308m1_V2MM_91379 | 2.291987 | 2.205692 | TRUE | HP_31275 | TGCTGTTG | CATACACT | BC052827 | Unknown (protein for IMAGE:30033377) | | 2.6 | SM2113 | a | 6 |
| mPool7_V2MM_78352 | 2.291468 | 3.004939 | TRUE | HP_30004 | TGCTGTTG | GACCTATG | NM_029624 | RIKEN cDNA 2400010G15 gene | 2400010G15Rik | 2.8 | SM2243 | h | 7 |
| mPool2_V2MM_132669 | 2.290049 | 2.205692 | TRUE | HP_35341 | TGCTGTTG | CACAGTTG | AK014914 | RIKEN cDNA 4921517O11 gene | 4921517O11Rik | 2.11 | SM2367 | e | 3 |
| mPool2_V2MM_133624 | 2.287794 | 2.205692 | TRUE | HP_35435 | TGCTGTTG | CTGACATG | AK014550 | RIKEN cDNA 4632404M16 gene | 4632404M16Rik | 2.11 | SM2358 | d | 4 |
| mPool7_V2MM_117028 | 2.286222 | 1.959154 | TRUE | HP_33797 | TGCTGTTG | CTGCTGCA | XM_139196 | Mus musculus similar to Hypothetical protein KIAA1002 (LOC219081), mRNA. | | 2.9 | SM2299 | e | 9 |
| mPool2_V2MM_109420 | 2.275413 | 2.205692 | TRUE | HP_33047 | TGCTGTTG | CTTCCAAG | XM_133877 | RIKEN cDNA 4933402N03 gene | 4933402N03Rik | 2.7 | SM2152 | f | 1 |
| mPool2_V2MM_30402 | 2.274178 | 2.205692 | TRUE | HP_25333 | TGCTGTTG | CGATGAAT | AK083102 | RIKEN cDNA C630015F21 gene | C630015F21Rik | 2.11 | SM2392 | g | 6 |
| mPool7_V2MM_179548 | 2.273032 | 2.205692 | TRUE | HP_40002 | TGCTGTTG | CCTACAAT | XM_288352 | Mus musculus LOC331672 (LOC331672), mRNA. | | 2.5 | SM2091 | d | 5 |
| mPool7_V2MM_179548 | 2.273032 | 2.205692 | TRUE | HP_40002 | TGCTGTTG | CCTACAAT | XM_288352 | Mus musculus LOC331672 (LOC331672), mR. | | 2.5 | SM2091 | d | 5 |
| mPool2_V2MM_169959 | 2.259909 | 3.004939 | TRUE | HP_39045 | TGCTGTTG | CAGGGTAT | XM_286366 | Mus musculus hypothetical gene supported by AK050116 (LOC327909), mRNA. | | 2.6 | SM2128 | e | 3 |
| 092308m1_V2MM_148639 | 2.256828 | 2.205692 | TRUE | HP_36935 | TGCTGTTG | CTCCCTGC | XM_163974 | Mus musculus LOC244370 (LOC244370), mR. | | 2.4 | SM2042 | b | 5 |
| mPool7_V2MM_171303 | 2.256793 | 2.205692 | TRUE | HP_39179 | TGCTGTTG | CTGCTCAC | AK084007 | RIKEN cDNA D130076A03 gene | D130076A03Rik | 2.9 | SM2272 | c | 3 |
| mPool2_V2MM_101731 | 2.251934 | 3.004939 | TRUE | HP_32289 | TGCTGTTG | CACTGTTA | XM_127850 | gene model 293, (NCBI) | Gm293 | 2.7 | SM2157 | c | 3 |
| mPool2_V2MM_136024 | 2.25192 | 2.205692 | TRUE | HP_35674 | TGCTGTTG | GGCTCTG/ | XM_151323 | Mus musculus LOC211516 (LOC211516), mRNA. | | 2.1 | SM2335 | h | 9 |
| mPool2_V2MM_6131 | 2.248339 | 2.205692 | TRUE | HP_22967 | TGCTGTTG | CTCAGTAC | NM_010789 | myeloid ecotropic viral integration site 1 | Meis1 | 2.11 | SM2388 | g | 5 |
| mPool2_V2MM_208005 | 2.245995 | 2.205692 | TRUE | HP_42580 | TGCTGTTG | GTACTATA | XM_489317 | LOC434549 | | 2.1 | SM2336 | g | 4 |
| 092308m1_V2MM_155480 | 2.244592 | 3.004939 | TRUE | HP_37611 | TGCTGTTG | CTCAGTAA | XM_197476 | Mus musculus LOC271802 (LOC271802), mR. | | 2.6 | SM2123 | d | 4 |
| mPool2_V2MM_3044 | 2.242759 | 2.205692 | TRUE | HP_22665 | TGCTGTTG | CTTATCTC | NM_007664 | cadherin 2 | Cdh2 | 2.7 | SM2166 | d | 11 |
| mPool2_V2MM_122581 | 2.240322 | 3.004939 | TRUE | HP_34344 | TGCTGTTG | GAGGTTAT | XM_488394 | similar to 40S ribosomal protein SA (P40) (34/67 kDa laminin receptor) | | 2.11 | SM2354 | f | 11 |
| 092308m1_V2MM_66431 | 2.237994 | 3.004939 | TRUE | HP_28844 | TGCTGTTG | CAGAGACT | NM_172991 | RIKEN cD C030048B08 gene | C030048B08Rik | 2.3 | SM2008 | e | 3 |
| mPool7_V2MM_100261 | 2.237624 | 3.004939 | TRUE | HP_32145 | TGCTGTTG | CAATTCAT | XM_126724 | fetal Alzheimer antigen | Falz | 2.7 | SM2152 | c | 9 |
| 092308m1_V2MM_171557 | 2.237398 | 2.205692 | TRUE | HP_39204 | TGCTGTTG | GAAATGTT | XM_286648 | Mus musculus hypothetical gene supported by AK037505 (LOC328492), mR. | | 2.5 | SM2078 | e | 1 |
| mPool7_V2MM_195100 | 2.236993 | 2.205692 | TRUE | HP_41324 | TGCTGTTG | CTGTATAT | NM_146632 | olfactory receptor 116 | Olfr116 | 2.9 | SM2254 | f | 8 |
| mPool7_V2MM_196394 | 2.234656 | 3.004939 | TRUE | HP_41449 | TGCTGTTG | CCGTAAAT | NM_026505 | BMP and activin membrane-bound inhibitor, homolog (X | Bambi | 2.9 | SM2266 | a | 7 |
| 092308m1_V2MM_142501 | 2.234342 | 3.004939 | TRUE | HP_36321 | TGCTGTTG | GACACTCA | XM_157672 | similar to Suppressor of G2 allele of SKP1 homolog (Sgt1) (Putative 40-6-3 protei | | 2.4 | SM2049 | d | 8 |
| mPool2_V2MM_101888 | 2.234022 | 2.205692 | TRUE | HP_32305 | TGCTGTTG | GCGCCTTT | NM_027707 | RIKEN cDNA 4933421G18 gene | 4933421G18Rik | 2.1 | SM2309 | h | 11 |
| mPool2_V2MM_16861 | 2.233186 | 2.205692 | TRUE | HP_24014 | TGCTGTTG | GACTGGTT | NM_008268 | homeo box B5 | Hoxb5 | 2.11 | SM2389 | f | 4 |
| mPool7_V2MM_179034 | 2.232598 | 3.004939 | TRUE | HP_39950 | TGCTGTTG | CAGGATAT | XM_288262 | Mus musculus LOC329282 (LOC329282), mR. | | 2.5 | SM2089 | b | 6 |
| mPool7_V2MM_179034 | 2.232598 | 3.004939 | TRUE | HP_39950 | TGCTGTTG | CAGGATAT | XM_288262 | Mus musculus LOC329282 (LOC329282), mRNA. | | 2.5 | SM2089 | b | 6 |
| mPool2_V2MM_159712 | 2.232264 | 2.205692 | TRUE | HP_38030 | TGCTGTTG | CAGGAAG | XM_283222 | Mus musculus hypothetical gene supported by AK034489 (LOC328439), mRNA. | | 2.6 | SM2126 | a | 7 |
| 092308m1_V2MM_158630 | 2.229339 | 3.004939 | TRUE | HP_37924 | TGCTGTTG | CAGCCTTA | XM_206878 | Mus musculus LOC279358 (LOC279358), mR. | | 2.6 | SM2116 | g | 9 |
| mPool2_V2MM_93146 | 2.226003 | 2.205692 | TRUE | HP_31447 | TGCTGTTG | GTGGATA/ | AK083705 | expressed sequence AI256775 | AI256775 | 2.6 | SM2110 | a | 2 |
| mPool2_V2MM_93146 | 2.226003 | 2.205692 | TRUE | HP_31447 | TGCTGTTG | GTGGATA/ | AK083705 | expressed sequence AI256775 | AI256775 | 2.6 | SM2110 | a | 2 |
| mPool2_V2MM_8740 | 2.223547 | 3.004939 | TRUE | HP_23222 | TGCTGTTG | CTTTGCAT | NM_027137 | small proline rich-like 7 | Sprrl7 | 2.11 | SM2384 | c | 2 |
| mPool2_V2MM_97667 | 2.223155 | 2.205692 | TRUE | HP_31890 | TGCTGTTG | GAAGGGA | XM_111809 | Mus musculus LOC241767 (LOC241767), mRNA. | | 2.6 | SM2144 | d | 3 |
| mPool2_V2MM_125139 | 2.221033 | 3.004939 | TRUE | HP_34598 | TGCTGTTG | GATCTTAC | XM_143811 | gene model 427, (NCBI) | Gm427 | 2.1 | SM2349 | b | 7 |

| | | | | | | | | | | | | | |
|---|---|---|---|---|---|---|---|---|---|---|---|---|---|
| mPool2_V2MM_113519 | 2.220567 | 3.004939 | TRUE | HP_33450 | TGCTGTTG | CTTGCTCT | XM_137005 | Mus musculus similar to Hypothetical protein KIAA0144 (LOC212406), mRNA. | | 2.7 | SM2154 | f | 2 |
| mPool7_V2MM_205096 | 2.220502 | 3.004939 | TRUE | HP_42297 | TGCTGTTG | CTCTGTAA | XM_136300 | Mus musculus similar to MRC OX-45 surface antigen precursor (BCM1 surface an | | 2.1 | SM2313 | h | 6 |
| mPool2_V2MM_15479 | 2.220285 | 3.004939 | TRUE | HP_23880 | TGCTGTTG | CTCTTATC | AK047140 | MAP/microtubule affinity-regulating kinase 1 | Mark1 | 2.11 | SM2383 | b | 1 |
| 092308m1_V2MM_70919 | 2.215476 | 3.004939 | TRUE | HP_29282 | TGCTGTTG | CAGCTTCA | NM_146796 | olfactory receptor 1499 | Olfr1499 | 2.4 | SM2022 | a | 8 |
| 092308m1_V2MM_144230 | 2.213874 | 3.004939 | TRUE | HP_36494 | TGCTGTTG | CATTGTAC | XM_159825 | Mus musculus LOC207745 (LOC207745), mRNA. | | 2.5 | SM2058 | a | 5 |
| 092308m1_V2MM_144230 | 2.213874 | 3.004939 | TRUE | HP_36494 | TGCTGTTG | CATTGTAC | XM_159825 | Mus musculus LOC207745 (LOC207745), mR. | | 2.5 | SM2058 | a | 5 |
| 092308m1_V2MM_184599 | 2.213551 | 3.004939 | TRUE | HP_40507 | TGCTGTTG | CACTTTCT | XM_289225 | Mus musculus LOC332850 (LOC332850), mRNA. | | 2.9 | SM2290 | e | 12 |
| 092308m1_V2MM_184599 | 2.213551 | 3.004939 | TRUE | HP_40507 | TGCTGTTG | CACTTTCT | XM_289225 | Mus musculus LOC332850 (LOC332850), mR. | | 2.9 | SM2290 | e | 12 |
| 092308m1_V2MM_148845 | 2.212773 | 2.205692 | TRUE | HP_36955 | TGCTGTTG | CACGTTTA | XM_164238 | Mus musculus LOC244499 (LOC244499), mR. | | 2.5 | SM2055 | b | 6 |
| mPool2_V2MM_214871 | 2.209404 | 2.205692 | TRUE | HP_43244 | TGCTGTTG | CCTGATTT | XM_143387 | Mus musculus similar to 60S ribosomal protein L21 (LOC242101), mRNA. | | 2.11 | SM2356 | e | 6 |
| mPool7_V2MM_199693 | 2.207545 | 3.004939 | TRUE | HP_41773 | TGCTGTTG | CAGGTTGA | AK086149 | RIKEN cDNA A730017D01 gene | A730017D01Rik | 2.9 | SM2291 | c | 11 |
| 092308m1_V2MM_173562 | 2.204047 | 2.205692 | TRUE | HP_39404 | TGCTGTTG | CACACAGT | AK048422 | hypothetical protein C130058N19 | | 2.5 | SM2086 | e | 11 |
| 092308m1_V2MM_170236 | 2.202563 | 2.205692 | TRUE | HP_39073 | TGCTGTTG | CGTTTATG | XM_286417 | Mus musculus hypothetical gene supported by AK036142 (LOC328071), mR. | | 2.6 | SM2116 | g | 8 |
| mPool2_V2MM_19590 | 2.201509 | 3.004939 | TRUE | HP_24280 | TGCTGTTG | GGTTGGA | NM_152229 | nuclear receptor subfamily 2, group E, member 1 | Nr2e1 | 2.7 | SM2164 | f | 3 |
| mPool2_V2MM_135781 | 2.201352 | 3.004939 | TRUE | HP_35650 | TGCTGTTG | CTGTTGTT | XM_151047 | Mus musculus LOC211853 (LOC211853), mRNA. | | 2.1 | SM2343 | a | 10 |
| 092308m1_V2MM_125703 | 2.198339 | 2.205692 | TRUE | HP_34654 | TGCTGTTG | GGCCTTAC | XM_144185 | Mus musculus similar to hypothetical protein C230069C04 (LOC242842), mR. | | 2.5 | SM2099 | c | 11 |
| mPool7_V2MM_174650 | 2.19784 | 3.004939 | TRUE | HP_39513 | TGCTGTTG | CTCTTCTT | AK082502 | RIKEN cDNA C230057H02 gene | C230057H02Rik | 2.5 | SM2089 | h | 10 |
| mPool7_V2MM_174650 | 2.19784 | 3.004939 | TRUE | HP_39513 | TGCTGTTG | CTCTTCTT | AK082502 | RIKEN cD C230057H02 gene | C230057H02Rik | 2.5 | SM2089 | h | 10 |
| mPool7_V2MM_174555 | 2.197047 | 2.205692 | TRUE | HP_39504 | TGCTGTTG | GTTTATAG | XM_287252 | Mus musculus hypothetical gene supported by AK084244 (LOC329673), mRNA. | | 2.9 | SM2291 | e | 4 |
| mPool7_V2MM_204410 | 2.197023 | 3.004939 | TRUE | HP_42231 | TGCTGTTG | GGTCTACC | XM_139317 | Mus musculus similar to Metal-response element-binding transcription factor 2 ( | | 2.9 | SM2298 | d | 8 |
| mPool2_V2MM_118884 | 2.196167 | 3.004939 | TRUE | HP_33981 | TGCTGTTG | GTACCTCA | XM_140107 | RIKEN cDNA 1700090G07 gene | 1700090G07Rik | 2.11 | SM2373 | b | 9 |
| mPool7_V2MM_198063 | 2.196087 | 2.205692 | TRUE | HP_41612 | TGCTGTTG | GCTGTAGC | XM_287341 | Mus musculus hypothetical gene supported by AK080995 (LOC329843), mRNA. | | 2.9 | SM2295 | d | 8 |
| mPool2_V2MM_169542 | 2.192794 | 3.004939 | TRUE | HP_39003 | TGCTGTTG | GCTCAGA | XM_286284 | Mus musculus hypothetical gene supported by AK049492 (LOC327825), mRNA. | | 2.6 | SM2126 | g | 3 |
| 092308m1_V2MM_76661 | 2.192463 | 2.205692 | TRUE | HP_29841 | TGCTGTTG | CTACTGTT | NM_146732 | olfactory receptor 488 | Olfr488 | 2.4 | SM2025 | c | 4 |
| 092308m1_V2MM_85838 | 2.188618 | 2.205692 | TRUE | HP_30731 | TGCTGTTG | CACAATCA | NM_146487 | olfactory receptor 130 | Olfr130 | 2.6 | SM2107 | f | 5 |
| mPool2_V2MM_133964 | 2.187805 | 3.004939 | TRUE | HP_35469 | TGCTGTTG | CTATGGAT | AK034038 | RIKEN cDNA 2310058O09 gene | 2310058O09Rik | 2.5 | SM2100 | a | 2 |
| mPool2_V2MM_133964 | 2.187805 | 3.004939 | TRUE | HP_35469 | TGCTGTTG | CTATGGAT | AK034038 | RIKEN cD 2310058O09 gene | 2310058O09Rik | 2.5 | SM2100 | a | 2 |
| mPool2_V2MM_159312 | 2.187191 | 2.205692 | TRUE | HP_37990 | TGCTGTTG | GTATGTTA | XM_283051 | Mus musculus hypothetical gene supported by AK048793 (LOC328053), mRNA. | | 2.6 | SM2129 | c | 8 |
| 092308m1_V2MM_77667 | 2.186486 | 1.959154 | TRUE | HP_29938 | TGCTGTTG | CTGTTTGT | NM_029478 | RIKEN cD 4930579A11 gene | 4930579A11Rik | 2.3 | SM2016 | d | 8 |
| mPool7_V2MM_67301 | 2.186257 | 3.004939 | TRUE | HP_28929 | TGCTGTTG | CCTCAAGA | NM_025555 | RIKEN cDNA 2410004B18 gene | 2410004B18Rik | 2.8 | SM2231 | b | 2 |
| mPool2_V2MM_157803 | 2.182956 | 3.004939 | TRUE | HP_37841 | TGCTGTTG | GGCAAATC | XM_205638 | Mus musculus similar to hypothetical protein FLJ38725 [Homo sapiens] (LOC277! | | 2.6 | SM2127 | g | 11 |
| mPool7_V2MM_139690 | 2.181734 | 2.205692 | TRUE | HP_36041 | TGCTGTTG | CAGACAAT | XM_155311 | Mus musculus LOC211071 (LOC211071), mRNA. | | 2.5 | SM2056 | f | 9 |
| mPool7_V2MM_139690 | 2.181734 | 2.205692 | TRUE | HP_36041 | TGCTGTTG | CAGACAAT | XM_155311 | Mus musculus LOC211071 (LOC211071), mR. | | 2.5 | SM2056 | f | 9 |
| 092308m1_V2MM_157112 | 2.181489 | 3.004939 | TRUE | HP_37773 | TGCTGTTG | CCGAAGTC | XM_204889 | Mus musculus similar to nuclear domain 10 protein [Homo sapiens] (LOC278732) | | 2.6 | SM2116 | e | 6 |
| 092308m1_V2MM_83357 | 2.18136 | 3.004939 | TRUE | HP_30490 | TGCTGTTG | CGCAGTA | NM_013511 | erythrocyte protein band 4.1-like 2 | Epb4.1l2 | 2.6 | SM2113 | a | 2 |
| mPool2_V2MM_31619 | 2.17997 | 3.004939 | TRUE | HP_25451 | TGCTGTTG | CTTATTGC | NM_019911 | tryptophan 2,3-dioxygenase | Tdo2 | 2.11 | SM2389 | e | 6 |
| mPool2_V2MM_206238 | 2.179866 | 2.205692 | TRUE | HP_42407 | TGCTGTTG | CTGGAACT | XM_138606 | similar to glyceraldehyde-3-phosphate dehydrogenase | | 2.1 | SM2318 | h | 12 |
| mPool4_V2MM_160042 | 2.179119 | 75.83547 | TRUE | NA | NA | NA | NA | NA | NA | NA | NA | NA | NA |
| 092308m1_V2MM_73701 | 2.178346 | 2.205692 | TRUE | HP_29552 | TGCTGTTG | GTAATCTA | NM_175034 | solute carrier family 24, member 5 | Slc24a5 | 2.4 | SM2022 | h | 3 |
| mPool2_V2MM_212273 | 2.176676 | 2.205692 | TRUE | HP_42994 | TGCTGTTG | GATAGAA | XM_141423 | Mus musculus LOC241709 (LOC241709), mRNA. | | 2.1 | SM2350 | b | 11 |
| mPool2_V2MM_1146 | 2.176472 | 2.205692 | TRUE | HP_22479 | TGCTGTTG | CATATTGA | NM_030611 | aldo-keto reductase family 1, member C6 | Akr1c6 | 2.11 | SM2377 | h | 7 |
| 092308m1_V2MM_132762 | 2.173996 | 3.004939 | TRUE | HP_35351 | TGCTGTTG | CTCCAGAC | XM_148876 | Mus musculus LOC225654 (LOC225654), mR. | | 2.6 | SM2101 | h | 7 |
| mPool2_V2MM_205508 | 2.173072 | 2.205692 | TRUE | NA | NA | NA | NA | NA | NA | NA | NA | NA | NA |
| 092308m1_V2MM_124416 | 2.172018 | 3.004939 | TRUE | HP_34526 | TGCTGTTG | GATCCAAT | XM_143434 | Mus musculus similar to ribosomal protein L34 [Rattus rattus] (LOC229740), mR. | | 2.6 | SM2114 | h | 3 |
| 092308m1_V2MM_161647 | 2.171355 | 2.205692 | TRUE | HP_38221 | TGCTGTTG | CATTCATA | XM_283996 | Mus musculus hypothetical gene supported by AK052563 (LOC329911), mR. | | 2.6 | SM2118 | b | 8 |
| mPool7_V2MM_73979 | 2.169804 | 2.205692 | TRUE | HP_29579 | TGCTGTTG | GTTGTAAC | NM_010721 | lamin B1 | Lmnb1 | 2.8 | SM2244 | d | 10 |
| 092308m1_V2MM_164671 | 2.169254 | 3.004939 | TRUE | HP_38521 | TGCTGTTG | GGCTTAAC | XM_487308 | similar to 40S ribosomal protein S2 | | 2.6 | SM2119 | b | 1 |
| 092308m1_V2MM_169628 | 2.168786 | 2.205692 | TRUE | HP_39012 | TGCTGTTG | GCAGTACT | AK081017 | hypothetical protein B930054O08 | | 2.6 | SM2119 | h | 9 |
| 092308m1_V2MM_162965 | 2.167877 | 3.004939 | TRUE | HP_38352 | TGCTGTTG | CTGCTCTG | XM_284565 | Mus musculus hypothetical gene supported by AK044672 (LOC331077), mR. | | 2.6 | SM2116 | b | 2 |
| mPool7_V2MM_88730 | 2.165839 | 1.959154 | TRUE | HP_31016 | TGCTGTTG | CTTATGAC | NM_175477 | zinc finger protein 574 | Zfp574 | 2.9 | SM2255 | a | 8 |
| mPool7_V2MM_103978 | 2.165484 | 2.205692 | TRUE | HP_32511 | TGCTGTTG | GTTGGAGT | AK011565 | RIKEN cDNA 2610027F03 gene | 2610027F03Rik | 2.1 | SM2304 | g | 5 |
| 092308m1_V2MM_120180 | 2.162264 | 2.205692 | TRUE | HP_34108 | TGCTGTTG | CTCTCCTA | XM_140790 | similar to Rpl7a protein | | 2.6 | SM2110 | a | 12 |
| 092308m1_V2MM_120180 | 2.162264 | 2.205692 | TRUE | HP_34108 | TGCTGTTG | CTCTCCTA | XM_140790 | similar to Rpl7a protein | | 2.6 | SM2110 | a | 12 |

| | | | | | | | | | | | | | |
|---|---|---|---|---|---|---|---|---|---|---|---|---|---|
| mPool7_V2MM_78895 | 2.16209 | 2.205692 | TRUE | HP_30057 | TGCTGTTG | GAGGATA | NM_026029 | RIKEN cDNA 2700085E05 gene | 2700085E05Rik | 2.8 | SM2244 | b | 10 |
| mPool7_V2MM_111145 | 2.161757 | 2.205692 | TRUE | HP_37701 | TGCTGTTG | CCCATTTA | XM_484460 | similar to 2610510D13Rik protein | | 2.6 | SM2114 | c | 6 |
| 092308m3_V2MM_55457 | 2.161418 | 75.83547 | TRUE | NA | NA | NA | NA | NA | NA | NA | NA | NA | NA |
| mPool2_V2MM_212428 | 2.152989 | 3.004939 | TRUE | HP_43009 | TGCTGTTG | GAGGAAA | AK082964 | cAMP responsive element binding protein 3-like 2 | Creb3l2 | 2.11 | SM2359 | d | 1 |
| mPool7_V2MM_95623 | 2.152689 | 1.959154 | TRUE | HP_31689 | TGCTGTTG | CAAGTAA | NM_198108 | cDNA sequence BC023055 | BC023055 | 2.8 | SM2250 | e | 12 |
| mPool7_V2MM_95623 | 2.152689 | 1.959154 | TRUE | HP_31689 | TGCTGTTG | CAAGTAA | NM_198108 | cD sequence BC023055 | BC023055 | 2.8 | SM2250 | e | 12 |
| mPool2_V2MM_10140 | 2.151917 | 1.959154 | TRUE | HP_23358 | TGCTGTTG | CTAATAAA | NM_007853 | degenerative spermatocyte homolog (Drosophila) | Degs | 2.11 | SM2386 | h | 11 |
| mPool7_V2MM_171488 | 2.150559 | 3.004939 | TRUE | HP_39197 | TGCTGTTG | CCAATAAT | XM_286635 | Mus musculus hypothetical gene supported by AK040469 (LOC328470), mRNA. | | 2.9 | SM2271 | c | 7 |
| mPool7_V2MM_176364 | 2.149908 | 1.959154 | TRUE | HP_39684 | TGCTGTTG | CATTCATT | XM_287651 | Mus musculus hypothetical gene supported by AK079645 (LOC330422), mRNA. | | 2.5 | SM2079 | b | 1 |
| mPool7_V2MM_176364 | 2.149908 | 1.959154 | TRUE | HP_39684 | TGCTGTTG | CATTCATT | XM_287651 | Mus musculus hypothetical gene supported by AK079645 (LOC330422), mR. | | 2.5 | SM2079 | b | 1 |
| mPool2_V2MM_132821 | 2.149411 | 2.205692 | TRUE | HP_35357 | TGCTGTTG | GGAGCTA | XM_148904 | oxysterol binding protein | Osbp | 2.11 | SM2368 | d | 10 |
| 092308m1_V2MM_154888 | 2.148857 | 2.205692 | TRUE | HP_37552 | TGCTGTTG | CACTTATT | XM_197073 | RIKEN cD 9330128J19 gene | 9330128J19Rik | 2.6 | SM2124 | a | 10 |
| 092308m1_V2MM_73153 | 2.147094 | 2.205692 | TRUE | HP_29499 | TGCTGTTG | GCATATCT | NM_026197 | RIKEN cD 2810013M15 gene | 2810013M15Rik | 2.3 | SM2012 | e | 10 |
| mPool7_V2MM_67969 | 2.146374 | 2.205692 | TRUE | HP_28994 | TGCTGTTG | CTGAATTT | NM_028815 | leucine-rich repeats and IQ motif containing 2 | Lrriq2 | 2.8 | SM2234 | d | 10 |
| mPool2_V2MM_112469 | 2.143769 | 3.004939 | TRUE | HP_33347 | TGCTGTTG | GGCAATC | XM_355265 | similar to usherin isoform B | | 2.7 | SM2151 | a | 12 |
| 092308m1_V2MM_175866 | 2.140792 | 2.205692 | TRUE | HP_39634 | TGCTGTTG | CAGCTAG | XM_287544 | Mus musculus hypothetical gene supported by AK035359 (LOC330238), mR. | | 2.5 | SM2060 | b | 4 |
| 092308m1_V2MM_174286 | 2.139789 | 1.959154 | TRUE | HP_39477 | TGCTGTTG | GTTATAAC | XM_287196 | Mus musculus hypothetical gene supported by AK034924 (LOC329550), mR. | | 2.5 | SM2076 | a | 10 |
| 092308m1_V2MM_171386 | 2.137622 | 2.205692 | TRUE | HP_39187 | TGCTGTTG | GGAGTTT | AK053660 | hypothetical protein E130119H08 | | 2.5 | SM2068 | c | 5 |
| mPool2_V2MM_124518 | 2.136591 | 3.004939 | TRUE | HP_34537 | TGCTGTTG | GGTACTA | XM_143495 | ribosomal protein S20 | Rps20 | 2.11 | SM2357 | g | 2 |
| 092308m1_V2MM_174338 | 2.136563 | 2.205692 | TRUE | HP_39482 | TGCTGTTG | CACACAG | AK083044 | RIKEN cD C530038A18 gene | C530038A18Rik | 2.5 | SM2087 | g | 8 |
| mPool7_V2MM_204042 | 2.134788 | 3.004939 | TRUE | HP_42196 | TGCTGTTG | GAACCTG | AK015711 | RIKEN cDNA 4930505O20 gene | 4930505O20Rik | 2.1 | SM2317 | d | 7 |
| 092308m1_V2MM_174678 | 2.132732 | 3.004939 | TRUE | HP_39516 | TGCTGTTG | CTAAGTTT | XM_287282 | Mus musculus hypothetical gene supported by AK041457 (LOC329726), mR. | | 2.5 | SM2062 | g | 2 |
| mPool7_V2MM_80803 | 2.129937 | 3.004939 | TRUE | HP_30242 | TGCTGTTG | GGCAAAT | NM_028796 | RIKEN cDNA 1700012B15 gene | 1700012B15Rik | 2.8 | SM2248 | h | 2 |
| mPool2_V2MM_106336 | 2.127353 | 2.205692 | TRUE | HP_32743 | TGCTGTTG | GTGATAA | AK004863 | alcohol dehydrogenase 6 (class V), pseudogene 1 | Adh6-ps1 | 2.7 | SM2163 | d | 6 |
| 092308m1_V2MM_126821 | 2.122218 | 3.004939 | TRUE | HP_34765 | TGCTGTTG | GAAATCA | XM_144829 | immunoglobulin kappa chain variable 28 (V28) | Igk-V28 | 2.6 | SM2103 | b | 5 |
| 092308m1_V2MM_126821 | 2.122218 | 3.004939 | TRUE | HP_34765 | TGCTGTTG | GAAATCA | XM_144829 | immunoglobulin kappa chain variable 28 (V28) | Igk-V28 | 2.6 | SM2103 | b | 5 |
| mPool2_V2MM_136270 | 2.119626 | 3.004939 | TRUE | HP_35699 | TGCTGTTG | GGAACAA | XM_151594 | Mus musculus LOC213763 (LOC213763), mR. | | 2.4 | SM2039 | h | 8 |
| mPool2_V2MM_136270 | 2.119626 | 3.004939 | TRUE | HP_35699 | TGCTGTTG | GGAACAA | XM_151594 | Mus musculus LOC213763 (LOC213763), mRNA. | | 2.4 | SM2039 | h | 8 |
| 092308m1_V2MM_156971 | 2.116529 | 1.959154 | TRUE | HP_37759 | TGCTGTTG | GCCATTAT | XM_204668 | similar to growth arrest-specific 2 like 2; GAS2-related protein | | 2.6 | SM2115 | f | 3 |
| mPool7_V2MM_80858 | 2.11584 | 3.004939 | TRUE | HP_30248 | TGCTGTTG | GGCAGAG | NM_025765 | RIKEN cDNA 4933437N03 gene | 4933437N03Rik | 2.8 | SM2240 | f | 9 |
| mPool2_V2MM_211508 | 2.115203 | 3.004939 | TRUE | HP_42920 | TGCTGTTG | GTTGGAT | XM_139696 | Mus musculus similar to 60S RIBOSOMAL PROTEIN L9 (LOC224311), mRNA. | | 2.1 | SM2349 | a | 6 |
| mPool7_V2MM_116075 | 2.114921 | 2.205692 | TRUE | HP_33703 | TGCTGTTG | GTGCTTAT | XM_138743 | Mus musculus similar to Gag [Ovis aries] (LOC218358), mRNA. | | 2.7 | SM2154 | e | 2 |
| mPool2_V2MM_160396 | 2.112521 | 2.205692 | TRUE | HP_38097 | TGCTGTTG | CTGGCTA | AK080941 | ankyrin repeat and KH domain containing 1 | Ankhd1 | 2.6 | SM2128 | b | 3 |
| mPool7_V2MM_204277 | 2.109901 | 2.205692 | TRUE | HP_42218 | TGCTGTTG | CTGCTTAT | AK019136 | RIKEN cDNA 2410150O07 gene | 2410150O07Rik | 2.9 | SM2299 | g | 1 |
| 092308m1_V2MM_121702 | 2.107362 | 2.205692 | TRUE | HP_34258 | TGCTGTTG | GCAAGAA | XM_141796 | Mus musculus similar to RIKEN cD 4933402E13 [Mus musculus] (LOC236842), mR | | 2.5 | SM2099 | d | 3 |
| 092308m1_V2MM_121702 | 2.107362 | 2.205692 | TRUE | HP_34258 | TGCTGTTG | GCAAGAA | XM_141796 | Mus musculus similar to RIKEN cDNA 4933402E13 [Mus musculus] (LOC236842), | | 2.5 | SM2099 | d | 3 |
| 092308m1_V2MM_141720 | 2.106854 | 3.004939 | TRUE | HP_36243 | TGCTGTTG | CCTGATTT | XM_156900 | Mus musculus LOC224946 (LOC224946), mR. | | 2.4 | SM2042 | f | 4 |
| 092308m1_V2MM_133964 | 2.105841 | 3.004939 | TRUE | HP_35469 | TGCTGTTG | CTATGGA | AK034038 | RIKEN cD 2310058O09 gene | 2310058O09Rik | 2.5 | SM2100 | a | 2 |
| 092308m1_V2MM_133964 | 2.105841 | 3.004939 | TRUE | HP_35469 | TGCTGTTG | CTATGGA | AK034038 | RIKEN cDNA 2310058O09 gene | 2310058O09Rik | 2.5 | SM2100 | a | 2 |
| mPool2_V2MM_91636 | 2.10493 | 2.205692 | TRUE | HP_31300 | TGCTGTTG | GGACTAA | AK046533 | zinc finger, DHHC domain containing 2 | Zdhhc2 | 2.6 | SM2144 | d | 6 |
| 092308m1_V2MM_117680 | 2.104055 | 2.205692 | TRUE | HP_33862 | TGCTGTTG | GGGATAA | XM_139486 | Mus musculus LOC239417 (LOC239417), mR. | | 2.5 | SM2097 | a | 1 |
| mPool2_V2MM_100261 | 2.103912 | 3.004939 | TRUE | HP_32145 | TGCTGTTG | CAATTCAT | XM_126724 | fetal Alzheimer antigen | Falz | 2.7 | SM2152 | c | 9 |
| mPool4_V2MM_22779 | 2.102706 | 75.83547 | TRUE | NA | NA | NA | NA | NA | NA | NA | NA | NA | NA |
| mPool7_V2MM_74961 | 2.102405 | 3.004939 | TRUE | HP_29675 | TGCTGTTG | CATAGAT | NM_175244 | RIKEN cDNA 1700064K09 gene | 1700064K09Rik | 2.8 | SM2245 | f | 3 |
| mPool7_V2MM_144466 | 2.102218 | 3.004939 | TRUE | HP_36517 | TGCTGTTG | CTACTTCT | XM_160031 | Mus musculus LOC241981 (LOC241981), mR. | | 2.4 | SM2042 | e | 6 |
| mPool7_V2MM_144466 | 2.102218 | 3.004939 | TRUE | HP_36517 | TGCTGTTG | CTACTTCT | XM_160031 | Mus musculus LOC241981 (LOC241981), mRNA. | | 2.4 | SM2042 | e | 6 |
| mPool2_V2MM_189909 | 2.098125 | 2.205692 | TRUE | HP_25841 | TGCTGTTG | CTATCAA | NM_177326 | p21 (CDKN1A)-activated kinase 2 | Pak2 | 2.11 | SM2391 | c | 12 |
| mPool7_V2MM_62468 | 2.097133 | 2.205692 | TRUE | HP_28457 | TGCTGTTG | CAGCTAG | NM_030218 | RIKEN cDNA 9130017N09 gene | 9130017N09Rik | 2.8 | SM2229 | d | 7 |
| mPool2_V2MM_91761 | 2.096039 | 2.205692 | TRUE | HP_31313 | TGCTGTTG | CTCATTGT | NM_178444 | EGF-like domain 7 | Egfl7 | 2.6 | SM2146 | f | 10 |
| 092308m1_V2MM_147365 | 2.095732 | 3.004939 | TRUE | HP_36807 | TGCTGTTG | GGTTTAG | XM_162901 | Mus musculus LOC208785 (LOC208785), mR. | | 2.4 | SM2033 | a | 6 |
| 092308m1_V2MM_165237 | 2.095658 | 3.004939 | TRUE | HP_38577 | TGCTGTTG | CAAGTAT | XM_285204 | Mus musculus similar to enolase 1, alpha non-neuron; alpha-enolase; 2-phospho | | 2.6 | SM2114 | g | 9 |
| mPool7_V2MM_200431 | 2.09333 | 2.205692 | TRUE | HP_41846 | TGCTGTTG | CATCTATA | XM_287522 | Mus musculus hypothetical gene supported by AK046921 (LOC330195), mRNA. | | 2.9 | SM2286 | f | 6 |
| 092308m1_V2MM_88186 | 2.093269 | 2.205692 | TRUE | HP_30963 | TGCTGTTG | CACTAAA | NM_153538 | zinc finger, CCHC domain containing 6 | Zcchc6 | 2.6 | SM2110 | d | 6 |

| | | | | | | | | | | | | | |
|---|---|---|---|---|---|---|---|---|---|---|---|---|---|
| mPool2_V2MM_83274 | 2.093266 | 3.004939 | TRUE | HP_30482 | TGCTGTTG | GGTGAAAT | NM_011815 | FYN binding protein | Fyb | 2.6 | SM2132 | g | 10 |
| 092308m1_V2MM_135644 | 2.090897 | 2.205692 | TRUE | HP_35636 | TGCTGTTG | CATCTATA | XM_150884 | Mus musculus LOC240965 (LOC240965), mR. | | 2.5 | SM2052 | e | 5 |
| mPool2_V2MM_19518 | 2.090357 | 3.004939 | TRUE | HP_24273 | TGCTGTTG | GGTGCAA | NM_134191 | vomeronasal 1 receptor, E2 | V1re2 | 2.11 | SM2384 | f | 3 |
| 092308m1_V2MM_71999 | 2.089876 | 3.004939 | TRUE | HP_29386 | TGCTGTTG | CTCGTAAT | NM_146813 | olfactory receptor 651 | Olfr651 | 2.4 | SM2028 | e | 6 |
| mPool2_V2MM_22093 | 2.088642 | 1.959154 | TRUE | HP_24525 | TGCTGTTG | CCTTTGTT | NM_024191 | ADP-ribosylation factor-like 2 binding protein | Arl2bp | 2.11 | SM2389 | f | 7 |
| mPool2_V2MM_96594 | 2.088408 | 3.004939 | TRUE | HP_31784 | TGCTGTTG | CTGAGGA | XM_110868 | similar to ORF1 | | 2.6 | SM2142 | d | 12 |
| mPool7_V2MM_197067 | 2.087911 | 2.205692 | TRUE | HP_41515 | TGCTGTTG | CAATCAGA | NM_173780 | Kruppel-like factor 8 | Klf8 | 2.9 | SM2258 | h | 9 |
| mPool7_V2MM_198688 | 2.087593 | 2.205692 | TRUE | HP_41674 | TGCTGTTG | CTTACTGA | AK048069 | hypothetical protein LOC328277 | | 2.9 | SM2296 | g | 3 |
| mPool7_V2MM_184921 | 2.087238 | 3.004939 | TRUE | HP_40539 | TGCTGTTG | GAGCTTCT | XM_289281 | Mus musculus LOC332932 (LOC332932), mRNA. | | 2.5 | SM2069 | a | 6 |
| mPool7_V2MM_184921 | 2.087238 | 3.004939 | TRUE | HP_40539 | TGCTGTTG | GAGCTTCT | XM_289281 | Mus musculus LOC332932 (LOC332932), mR. | | 2.5 | SM2069 | a | 6 |
| mPool2_V2MM_205219 | 2.085947 | 2.205692 | TRUE | HP_42309 | TGCTGTTG | CTCACATT | XM_138748 | Mus musculus similar to envelope protein [Ovis aries] (LOC218362), mRNA. | | 2.1 | SM2319 | a | 3 |
| mPool2_V2MM_143769 | 2.082516 | 2.205692 | TRUE | HP_36448 | TGCTGTTG | CTTGATAG | XM_159101 | hypothetical LOC208429 | | 2.1 | SM2342 | a | 9 |
| mPool2_V2MM_143769 | 2.082516 | 2.205692 | TRUE | HP_36448 | TGCTGTTG | CTTGATAG | XM_159101 | hypothetical LOC208429 | | 2.1 | SM2342 | a | 9 |
| 092308m1_V2MM_179819 | 2.08022 | 3.004939 | TRUE | HP_40029 | TGCTGTTG | CTACTTCA | XM_288402 | Mus musculus LOC327894 (LOC327894), mR. | | 2.5 | SM2090 | e | 11 |
| 092308m1_V2MM_174650 | 2.078315 | 3.004939 | TRUE | HP_39513 | TGCTGTTG | CTCTTCTT | AK082502 | RIKEN cD C230057H02 gene | C230057H02Rik | 2.5 | SM2089 | h | 10 |
| 092308m1_V2MM_174650 | 2.078315 | 3.004939 | TRUE | HP_39513 | TGCTGTTG | CTCTTCTT | AK082502 | RIKEN cDNA C230057H02 gene | C230057H02Rik | 2.5 | SM2089 | h | 10 |
| mPool7_V2MM_197174 | 2.077952 | 2.205692 | TRUE | HP_41525 | TGCTGTTG | GATCATAT | NM_146693 | olfactory receptor 1462 | Olfr1462 | 2.9 | SM2260 | g | 3 |
| 092308m1_V2MM_184921 | 2.077901 | 3.004939 | TRUE | HP_40539 | TGCTGTTG | GAGCTTCT | XM_289281 | Mus musculus LOC332932 (LOC332932), mR. | | 2.5 | SM2069 | a | 6 |
| 092308m1_V2MM_184921 | 2.077901 | 3.004939 | TRUE | HP_40539 | TGCTGTTG | GAGCTTCT | XM_289281 | Mus musculus LOC332932 (LOC332932), mRNA. | | 2.5 | SM2069 | a | 6 |
| mPool7_V2MM_201173 | 2.077697 | 2.205692 | TRUE | HP_41920 | TGCTGTTG | CCTTAAGC | XM_289648 | Mus musculus LOC333392 (LOC333392), mRNA. | | 2.9 | SM2294 | b | 12 |
| mPool2_V2MM_101746 | 2.076362 | 2.205692 | TRUE | HP_32291 | TGCTGTTG | GCTGAAGT | XM_127854 | A kinase (PRKA) anchor protein 11 | Akap11 | 2.7 | SM2152 | h | 5 |
| 092308m1_V2MM_175564 | 2.075031 | 2.205692 | TRUE | HP_39604 | TGCTGTTG | GCCATTGC | XM_489100 | hypothetical gene supported by AK031827 | | 2.5 | SM2078 | d | 9 |
| 092308m1_V2MM_175564 | 2.075031 | 2.205692 | TRUE | HP_39604 | TGCTGTTG | GCCATTGC | XM_489100 | hypothetical gene supported by AK031827 | | 2.5 | SM2078 | d | 9 |
| 092308m1_V2MM_171663 | 2.07375 | 2.205692 | TRUE | HP_39215 | TGCTGTTG | CATATTCT | AK078525 | activin A receptor, type 1B | Acvr1b | 2.5 | SM2062 | a | 5 |
| 092308m1_V2MM_171663 | 2.07375 | 2.205692 | TRUE | HP_39215 | TGCTGTTG | CATATTCT | AK078525 | activin A receptor, type 1B | Acvr1b | 2.5 | SM2062 | a | 5 |
| mPool2_V2MM_208454 | 2.073222 | 3.004939 | TRUE | HP_42624 | TGCTGTTG | GAGAAACT | XM_160842 | Mus musculus LOC242351 (LOC242351), mRNA. | | 2.1 | SM2336 | c | 5 |
| mPool7_V2MM_70814 | 2.07193 | 3.004939 | TRUE | HP_29271 | TGCTGTTG | CAGATAAT | NM_025711 | asporin | Aspn | 2.8 | SM2240 | e | 4 |
| 092308m1_V2MM_173334 | 2.071353 | 3.004939 | TRUE | HP_39382 | TGCTGTTG | CACAAATT | XM_287004 | Mus musculus hypothetical gene supported by AK081108 (LOC329138), mR. | | 2.5 | SM2067 | c | 12 |
| mPool2_V2MM_102674 | 2.07014 | 3.004939 | TRUE | HP_32382 | TGCTGTTG | CCCTTGGA | AK076291 | RIKEN cDNA 4632424B03 gene | | 2.7 | SM2153 | b | 6 |
| mPool2_V2MM_100923 | 2.069492 | 3.004939 | TRUE | HP_32209 | TGCTGTTG | CAGGATAT | AK005458 | prolactin-like protein N | Prlpn | 2.1 | SM2324 | g | 1 |
| mPool7_V2MM_179101 | 2.06843 | 3.004939 | TRUE | HP_39957 | TGCTGTTG | CCACATTC | XM_288274 | Mus musculus LOC332550 (LOC332550), mRNA. | | 2.5 | SM2092 | a | 9 |
| mPool7_V2MM_179101 | 2.06843 | 3.004939 | TRUE | HP_39957 | TGCTGTTG | CCACATTC | XM_288274 | Mus musculus LOC332550 (LOC332550), mR. | | 2.5 | SM2092 | a | 9 |
| 092308m1_V2MM_179268 | 2.068282 | 2.205692 | TRUE | HP_39974 | TGCTGTTG | CAGTCCTC | XM_288302 | Mus musculus LOC331620 (LOC331620), mR. | | 2.5 | SM2094 | b | 10 |
| 092308m1_V2MM_137537 | 2.067598 | 3.004939 | TRUE | HP_35825 | TGCTGTTG | CTTAAAGT | XM_153261 | similar to Rho GTPase activating protein 20 | | 2.4 | SM2046 | h | 8 |
| 092308m1_V2MM_137537 | 2.067598 | 3.004939 | TRUE | HP_35825 | TGCTGTTG | CTTAAAGT | XM_153261 | similar to Rho GTPase activating protein 20 | | 2.4 | SM2046 | h | 8 |
| mPool7_V2MM_182263 | 2.067132 | 2.205692 | TRUE | HP_40273 | TGCTGTTG | CTCATTTA | XM_288815 | Mus musculus LOC332291 (LOC332291), mRNA. | | 2.9 | SM2295 | a | 8 |
| mPool2_V2MM_3059 | 2.067122 | 3.004939 | TRUE | HP_22666 | TGCTGTTG | CTTCATTG | NM_026437 | RIKEN cDNA 1810055E12 gene | 1810055E12Rik | 2.11 | SM2376 | f | 6 |
| mPool7_V2MM_205775 | 2.063797 | 2.205692 | TRUE | HP_28627 | TGCTGTTG | CTGAGTTT | NM_030236 | F-box only protein 34 | Fbxo34 | 2.1 | SM2302 | g | 8 |
| 092308m1_V2MM_174942 | 2.063045 | 3.004939 | TRUE | HP_39542 | TGCTGTTG | CTGGAAAT | XM_287341 | Mus musculus hypothetical gene supported by AK080995 (LOC329843), mR. | | 2.5 | SM2077 | a | 7 |
| 092308m1_V2MM_184975 | 2.062451 | 2.205692 | TRUE | HP_40544 | TGCTGTTG | CTGCTTGA | XM_289290 | Mus musculus LOC332940 (LOC332940), mR. | | 2.5 | SM2095 | a | 4 |
| mPool7_V2MM_78316 | 2.061128 | 3.004939 | TRUE | HP_30000 | TGCTGTTG | GACCAGAT | NM_011192 | proteaseome (prosome, macropain) 28 subunit, 3 | Psme3 | 2.8 | SM2232 | a | 10 |
| mPool2_V2MM_104747 | 2.060998 | 2.205692 | TRUE | HP_32587 | TGCTGTTG | GAGCTTAT | AK173248 | zinc finger, CCHC domain containing 2 | Zcchc2 | 2.6 | SM2149 | c | 4 |
| mPool2_V2MM_144169 | 2.058879 | 2.205692 | TRUE | HP_36488 | TGCTGTTG | CATTTGTG | XM_159785 | Mus musculus LOC207783 (LOC207783), mRNA. | | 2.1 | SM2340 | h | 5 |
| 092308m1_V2MM_135868 | 2.057805 | 2.205692 | TRUE | HP_35658 | TGCTGTTG | CATGACAC | XM_151129 | Mus musculus LOC212730 (LOC212730), mR. | | 2.4 | SM2031 | d | 11 |
| 092308m1_V2MM_141175 | 2.055914 | 2.205692 | TRUE | HP_36188 | TGCTGTTG | CGGAAACA | XM_156493 | Mus musculus LOC239875 (LOC239875), mR. | | NA | | | NA |
| mPool2_V2MM_16625 | 2.05208 | 3.004939 | TRUE | HP_23991 | TGCTGTTG | GACAGAG | NM_134129 | PRP19/PSO4 homolog (S. cerevisiae) | Prp19 | 2.11 | SM2384 | d | 11 |
| mPool2_V2MM_18303 | 2.051061 | 2.205692 | TRUE | HP_24155 | TGCTGTTG | GCCTCTAC | NM_011607 | tenascin C | Tnc | 2.7 | SM2166 | c | 5 |
| mPool2_V2MM_7219 | 2.048636 | 3.004939 | TRUE | HP_23073 | TGCTGTTG | GAGCTACA | NM_007798 | cathepsin B | Ctsb | 2.11 | SM2383 | a | 7 |
| 092308m1_V2MM_160569 | 2.047524 | 3.004939 | TRUE | HP_38114 | TGCTGTTG | GTTTCTGA | XM_283567 | Mus musculus similar to hypothetical protein 7 - rat (LOC329062), mR. | | 2.6 | SM2118 | h | 6 |
| 092308m3_V2MM_41756 | 2.046211 | 60.13437 | TRUE | HP_26438 | TGCTGTTG | CAGATATT | NM_009451 | tubulin, beta 4 | Tubb4 | 2.12 | SM2416 | e | 11 |
| mPool7_V2MM_86068 | 2.045252 | 1.959154 | TRUE | HP_30754 | TGCTGTTG | CTATCATG | NM_146613 | olfactory receptor 973 | Olfr973 | 2.9 | SM2267 | g | 7 |
| 092308m1_V2MM_130662 | 2.044505 | 3.004939 | TRUE | HP_35144 | TGCTGTTG | CATCCTTT | XM_146968 | Mus musculus similar to hypothetical protein MGC18257 [Homo sapiens] (LOC21 | | 2.5 | SM2100 | c | 9 |
| 092308m1_V2MM_70007 | 2.042037 | 3.004939 | TRUE | HP_29192 | TGCTGTTG | GTGCAATC | NM_007380 | abl-interactor 1 | Abi1 | 2.4 | SM2022 | a | 6 |

| | | | | | | | | | | | | | |
|---|---|---|---|---|---|---|---|---|---|---|---|---|---|
| 092308m1_V2MM_151908 | 2.041136 | 3.004939 | TRUE | HP_37257 | TGCTGTTG | CGTTTATC | XM_194993 | olfactory receptor 171 | Olfr171 | 2.4 | SM2042 | e | 4 |
| mPool2_V2MM_166901 | 2.039306 | 3.004939 | TRUE | HP_38742 | TGCTGTTG | CTGATATT | XM_285628 | Mus musculus similar to zinc finger protein 93 homolog; zinc finger protein homo | | 2.6 | SM2128 | a | 6 |
| mPool7_V2MM_179484 | 2.039014 | 1.959154 | TRUE | HP_39995 | TGCTGTTG | CACTAATT | XM_288341 | Mus musculus LOC331656 (LOC331656), mRNA. | | 2.5 | SM2076 | b | 2 |
| mPool7_V2MM_179484 | 2.039014 | 1.959154 | TRUE | HP_39995 | TGCTGTTG | CACTAATT | XM_288341 | Mus musculus LOC331656 (LOC331656), mR. | | 2.5 | SM2076 | b | 2 |
| mPool2_V2MM_6274 | 2.038377 | 2.205692 | TRUE | HP_22981 | TGCTGTTG | CTCTTATA | NM_019586 | ubiquitin-conjugating enzyme E2, J1 | Ube2j1 | 2.11 | SM2381 | c | 2 |
| mPool7_V2MM_151720 | 2.03783 | 3.004939 | TRUE | HP_37239 | TGCTGTTG | CGGATTA/ | XM_489722 | similar to Rpl17 protein | | 2.4 | SM2035 | b | 4 |
| mPool7_V2MM_151720 | 2.03783 | 3.004939 | TRUE | HP_37239 | TGCTGTTG | CGGATTA/ | XM_489722 | similar to Rpl17 protein | | 2.4 | SM2035 | b | 4 |
| 092308m1_V2MM_136460 | 2.0376 | 3.004939 | TRUE | HP_35718 | TGCTGTTG | CTCTGTAT | XM_151736 | Mus musculus LOC212058 (LOC212058), mR. | | 2.5 | SM2057 | a | 5 |
| mPool2_V2MM_210700 | 2.035306 | 3.004939 | TRUE | HP_42843 | TGCTGTTG | CAGACCA( | XM_141415 | Mus musculus similar to Peptidyl-prolyl cis-trans isomerase A (PPIase) (Rotamase | | 2.1 | SM2350 | g | 8 |
| 092308m1_V2MM_142526 | 2.034763 | 2.205692 | TRUE | HP_36323 | TGCTGTTG | CTTAGGA/ | XM_157697 | Mus musculus LOC240481 (LOC240481), mR. | | 2.5 | SM2056 | c | 9 |
| 092308m1_V2MM_142526 | 2.034763 | 2.205692 | TRUE | HP_36323 | TGCTGTTG | CTTAGGA/ | XM_157697 | Mus musculus LOC240481 (LOC240481), mRNA. | | 2.5 | SM2056 | c | 9 |
| mPool2_V2MM_117317 | 2.034715 | 2.205692 | TRUE | HP_33826 | TGCTGTTG | CCTATGTC | XM_139321 | Mus musculus similar to protease [Mus musculus] (LOC223346), mRNA. | | 2.7 | SM2160 | f | 9 |
| mPool2_V2MM_85468 | 2.034221 | 3.004939 | TRUE | HP_30694 | TGCTGTTG | GCCATTG( | NM_146390 | olfactory receptor 1323 | Olfr1323 | 2.6 | SM2143 | e | 8 |
| mPool2_V2MM_147923 | 2.03199 | 2.205692 | TRUE | HP_36863 | TGCTGTTG | CAATTTCC | XM_163354 | Mus musculus LOC215451 (LOC215451), mRNA. | | 2.5 | SM2052 | e | 6 |
| mPool2_V2MM_147923 | 2.03199 | 2.205692 | TRUE | HP_36863 | TGCTGTTG | CAATTTCC | XM_163354 | Mus musculus LOC215451 (LOC215451), mR. | | 2.5 | SM2052 | e | 6 |
| mPool2_V2MM_9420 | 2.030441 | 3.004939 | TRUE | HP_23288 | TGCTGTTG | CAGCTGT/ | NM_019571 | transmembrane 4 superfamily member 9 | Tm4sf9 | 2.11 | SM2380 | f | 3 |
| 092308m1_V2MM_66196 | 2.028476 | 3.004939 | TRUE | HP_28821 | TGCTGTTG | CAACAAA1 | NM_133196 | cleavage stimulation factor, 3 pre-R subunit 2 | Cstf2 | 2.4 | SM2029 | e | 4 |
| mPool2_V2MM_39295 | 2.025544 | 3.004939 | TRUE | HP_26198 | TGCTGTTG | GCTTCAA/ | NM_130451 | solute carrier family 2 (facilitated glucose transporter), m | Slc2a10 | NA | | | NA |
| mPool2_V2MM_84154 | 2.02289 | 1.959154 | TRUE | HP_30567 | TGCTGTTG | CTATACCT | NM_026989 | RIKEN cDNA 2610019N13 gene | 2610019N13Rik | 2.6 | SM2133 | f | 1 |
| mPool2_V2MM_5641 | 2.022473 | 2.205692 | TRUE | HP_22918 | TGCTGTTG | CCTAGAA( | NM_007622 | chromobox homolog 1 (Drosophila HP1 beta) | Cbx1 | 2.11 | SM2382 | d | 6 |
| mPool7_V2MM_177038 | 2.021263 | 3.004939 | TRUE | HP_39751 | TGCTGTTG | CCTTAAAT | AK039009 | hypothetical protein A230084N10 | | 2.5 | SM2085 | e | 9 |
| mPool7_V2MM_177038 | 2.021263 | 3.004939 | TRUE | HP_39751 | TGCTGTTG | CCTTAAAT | AK039009 | hypothetical protein A230084N10 | | 2.5 | SM2085 | e | 9 |
| mPool2_V2MM_204607 | 2.018482 | 2.205692 | TRUE | HP_42250 | TGCTGTTG | GTAGAATT | AK050925 | RIKEN cDNA 4930513F16 gene | 4930513F16Rik | 2.1 | SM2319 | a | 6 |
| mPool7_V2MM_69056 | 2.01462 | 2.205692 | TRUE | HP_29099 | TGCTGTTG | GATGTGT/ | NM_008041 | formyl peptide receptor, related sequence 4 | Fpr-rs4 | 2.8 | SM2242 | e | 6 |
| mPool7_V2MM_70275 | 2.013936 | 3.004939 | TRUE | HP_29219 | TGCTGTTG | CAAATATA | NM_030150 | DNA segment, Chr 11, Lothar Hennighausen 2, expressed | D11Lgp2e | 2.8 | SM2245 | g | 7 |
| 092308m1_V2MM_143932 | 2.011254 | 3.004939 | TRUE | HP_36464 | TGCTGTTG | CAGAGAT/ | XM_159469 | Mus musculus LOC211711 (LOC211711), mRNA. | | 2.4 | SM2033 | d | 1 |
| 092308m1_V2MM_143932 | 2.011254 | 3.004939 | TRUE | HP_36464 | TGCTGTTG | CAGAGAT/ | XM_159469 | Mus musculus LOC211711 (LOC211711), mR. | | 2.4 | SM2033 | d | 1 |
| mPool7_V2MM_67685 | 2.009994 | 2.205692 | TRUE | HP_28966 | TGCTGTTG | CTATACAA | NM_011241 | RAN GTPase activating protein 1 | Rangap1 | 2.8 | SM2231 | e | 11 |
| mPool2_V2MM_118382 | 2.009705 | 3.004939 | TRUE | HP_33931 | TGCTGTTG | CTTGAGG/ | XM_139821 | similar to RIKEN cDNA 1700029I01 | | 2.11 | SM2368 | b | 6 |
| mPool7_V2MM_174732 | 2.008312 | 2.205692 | TRUE | HP_39521 | TGCTGTTG | GCAGTAA/ | AK034327 | hypothetical protein 9330178D15 | | 2.9 | SM2277 | f | 7 |
| mPool7_V2MM_174732 | 2.008312 | 2.205692 | TRUE | HP_39521 | TGCTGTTG | GCAGTAA/ | AK034327 | hypothetical protein 9330178D15 | | 2.9 | SM2277 | f | 7 |
| mPool7_V2MM_181893 | 2.007928 | 2.205692 | TRUE | HP_40236 | TGCTGTTG | GTTTATTG | XM_288752 | Mus musculus LOC332202 (LOC332202), mRNA. | | 2.5 | SM2089 | c | 5 |
| mPool7_V2MM_181893 | 2.007928 | 2.205692 | TRUE | HP_40236 | TGCTGTTG | GTTTATTG | XM_288752 | Mus musculus LOC332202 (LOC332202), mR. | | 2.5 | SM2089 | c | 5 |
| 092308m1_V2MM_91659 | 2.007065 | 3.004939 | TRUE | HP_31302 | TGCTGTTG | CGGAAGT( | NM_178399 | RIKEN cD 3110035E14 gene | 3110035E14Rik | 2.6 | SM2109 | c | 9 |
| 092308m1_V2MM_170685 | 2.00697 | 3.004939 | TRUE | HP_39117 | TGCTGTTG | GGTTTATC | XM_286470 | Mus musculus hypothetical gene supported by AK048337 (LOC328164), mR. | | 2.5 | SM2095 | g | 5 |
| 092308m1_V2MM_175033 | 2.005484 | 3.004939 | TRUE | HP_39551 | TGCTGTTG | GATAGCC/ | AK029093 | RIKEN cDNA E030045D18 gene | E030045D18Rik | 2.5 | SM2080 | c | 7 |
| 092308m1_V2MM_175033 | 2.005484 | 3.004939 | TRUE | HP_39551 | TGCTGTTG | GATAGCC/ | AK029093 | RIKEN cD E030045D18 gene | E030045D18Rik | 2.5 | SM2080 | c | 7 |
| mPool2_V2MM_108684 | 2.003483 | 2.205692 | TRUE | HP_32975 | TGCTGTTG | CTGAGTT( | XM_133185 | similar to hypothetical protein FLJ12895 | | 2.7 | SM2159 | h | 3 |
| mPool7_V2MM_93943 | 2.003002 | 3.004939 | TRUE | HP_31524 | TGCTGTTG | CTTGCCCA | NM_182840 | elastin microfibril interfacer 3 | Emilin3 | 2.8 | SM2249 | e | 9 |
| 092308m1_V2MM_165554 | 2.000053 | 3.004939 | TRUE | HP_38608 | TGCTGTTG | CTGGAAGT | XM_487540 | LOC435573 | | 2.6 | SM2120 | c | 10 |
| mPool2_V2MM_9341 | 1.998236 | 1.959154 | TRUE | HP_23280 | TGCTGTTG | CAGATTC( | NM_010903 | nuclear factor, erythroid derived 2, like 3 | Nfe2l3 | 2.11 | SM2386 | b | 9 |
| 092308m1_V2MM_137497 | 1.997 | 3.004939 | TRUE | HP_35821 | TGCTGTTG | GAAATTA1 | XM_153221 | RIKEN cDNA 1110038H03 gene | 1110038H03Rik | 2.4 | SM2042 | h | 10 |
| 092308m1_V2MM_137497 | 1.997 | 3.004939 | TRUE | HP_35821 | TGCTGTTG | GAAATTA1 | XM_153221 | RIKEN cD 1110038H03 gene | 1110038H03Rik | 2.4 | SM2042 | h | 10 |
| mPool2_V2MM_1383 | 1.995442 | 3.004939 | TRUE | HP_22502 | TGCTGTTG | CCATAGA( | NM_018748 | golgi autoantigen, golgin subfamily a, 4 | Golga4 | 2.11 | SM2386 | f | 11 |
| mPool2_V2MM_163514 | 1.993409 | 3.004939 | TRUE | HP_38406 | TGCTGTTG | CTGAATAT | XM_284801 | RIKEN cDNA 1700029B21 gene | 1700029B21Rik | 2.6 | SM2127 | a | 8 |
| mPool7_V2MM_204599 | 1.992271 | 3.004939 | TRUE | HP_42249 | TGCTGTTG | CGCCAAG( | XM_486831 | similar to RIKEN cDNA 4921509A18 gene | | 2.1 | SM2314 | a | 9 |
| 092308m1_V2MM_135381 | 1.991633 | 2.205692 | TRUE | HP_35610 | TGCTGTTG | CTGTAATC | XM_150596 | Mus musculus LOC240784 (LOC240784), mR. | | 2.5 | SM2058 | e | 12 |
| mPool2_V2MM_144938 | 1.991622 | 3.004939 | TRUE | HP_36564 | TGCTGTTG | CAATTGA( | XM_160439 | Mus musculus LOC242208 (LOC242208), mR. | | 2.4 | SM2045 | a | 10 |
| mPool2_V2MM_144938 | 1.991622 | 3.004939 | TRUE | HP_36564 | TGCTGTTG | CAATTGA( | XM_160439 | Mus musculus LOC242208 (LOC242208), mRNA. | | 2.4 | SM2045 | a | 10 |
| 092308m1_V2MM_144938 | 1.990391 | 3.004939 | TRUE | HP_36564 | TGCTGTTG | CAATTGA( | XM_160439 | Mus musculus LOC242208 (LOC242208), mRNA. | | 2.4 | SM2045 | a | 10 |
| 092308m1_V2MM_144938 | 1.990391 | 3.004939 | TRUE | HP_36564 | TGCTGTTG | CAATTGA( | XM_160439 | Mus musculus LOC242208 (LOC242208), mR. | | 2.4 | SM2045 | a | 10 |
| mPool2_V2MM_116119 | 1.986981 | 2.205692 | TRUE | HP_33707 | TGCTGTTG | CGTTTGAT | XM_138761 | Mus musculus similar to NADH-UBIQUINONE OXIDOREDUCTASE CHAIN 1 (LOC23 | | 2.6 | SM2149 | f | 4 |
| 092308m1_V2MM_75728 | 1.98636 | 2.205692 | TRUE | HP_29750 | TGCTGTTG | CCCTTATA | NM_146700 | olfactory receptor 1453 | Olfr1453 | 2.4 | SM2020 | c | 8 |

| | | | | | | | | | | | | | |
|---|---|---|---|---|---|---|---|---|---|---|---|---|---|
| mPool2_V2MM_104265 | 1.985448 | 3.004939 | TRUE | HP_32539 | TGCTGTTG | CATGCATC | XM_129647 | glutamyl-prolyl-tRNA synthetase | Eprs | 2.6 | SM2149 | f | 9 |
| 092308m1_V2MM_136201 | 1.985416 | 2.205692 | TRUE | HP_35692 | TGCTGTTG | CGGGAAT | NM_207031 | prostate cancer associated protein 5 | Pcap5 | 2.4 | SM2035 | d | 10 |
| mPool2_V2MM_134989 | 1.985106 | 1.959154 | TRUE | HP_35571 | TGCTGTTG | GAAATCA | AK015733 | RIKEN cDNA 4930509E16 gene | 4930509E16Rik | 2.11 | SM2363 | g | 12 |
| 092308m1_V2MM_135825 | 1.983802 | 3.004939 | TRUE | HP_35654 | TGCTGTTG | GAGTTAC | XM_151088 | Mus musculus LOC212291 (LOC212291), mR. | | 2.4 | SM2047 | h | 12 |
| mPool2_V2MM_208932 | 1.983489 | 2.205692 | TRUE | HP_42671 | TGCTGTTG | CCCAATTT | XM_163354 | Mus musculus LOC215451 (LOC215451), mRNA. | | 2.1 | SM2345 | d | 7 |
| 092308m1_V2MM_176881 | 1.979243 | 3.004939 | TRUE | HP_39736 | TGCTGTTG | GTGATATT | AK085344 | RIKEN cD D630014A15 gene | D630014A15Rik | 2.5 | SM2081 | e | 7 |
| mPool2_V2MM_92572 | 1.978452 | 3.004939 | TRUE | HP_31391 | TGCTGTTG | CCCAATAC | NM_178735 | RIKEN cDNA A730069N07 gene | A730069N07Rik | 2.6 | SM2143 | f | 5 |
| 092308m1_V2MM_91071 | 1.978003 | 3.004939 | TRUE | HP_31248 | TGCTGTTG | CTTCAAGA | NM_177915 | immunoglobulin superfamily, member 1 | Igsf1 | 2.6 | SM2113 | e | 12 |
| 092308m1_V2MM_176857 | 1.977693 | 3.004939 | TRUE | HP_39733 | TGCTGTTG | GTACCAAT | XM_287757 | non-coding transcript 1 | Nctc1 | 2.5 | SM2063 | h | 8 |
| mPool7_V2MM_108179 | 1.975357 | 3.004939 | TRUE | HP_32925 | TGCTGTTG | CCAGTTAT | BC058338 | RIKEN cDNA 2210404G23 gene | 2210404G23Rik | 2.1 | SM2315 | c | 8 |
| mPool7_V2MM_78418 | 1.973083 | 2.205692 | TRUE | HP_30010 | TGCTGTTG | GACTATCA | NM_022332 | Suppression of tumorigenicity 7 | St7 | 2.8 | SM2234 | e | 12 |
| mPool7_V2MM_201299 | 1.970236 | 2.205692 | TRUE | HP_41932 | TGCTGTTG | CAATCTAT | AK043696 | RIKEN cDNA A830021F12 gene | A830021F12Rik | 2.9 | SM2284 | d | 7 |
| mPool2_V2MM_139604 | 1.969442 | 3.004939 | TRUE | HP_36032 | TGCTGTTG | CAGAGAG | XM_155248 | Mus musculus LOC223326 (LOC223326), mRNA. | | 2.1 | SM2347 | e | 1 |
| 092308m1_V2MM_138094 | 1.967412 | 3.004939 | TRUE | HP_35881 | TGCTGTTG | GACAGTA | XM_153883 | Mus musculus LOC238525 (LOC238525), mR. | | 2.5 | SM2054 | a | 9 |
| 092308m1_V2MM_138094 | 1.967412 | 3.004939 | TRUE | HP_35881 | TGCTGTTG | GACAGTA | XM_153883 | Mus musculus LOC238525 (LOC238525), mRNA. | | 2.5 | SM2054 | a | 9 |
| 092308m3_V2MM_255457 | 1.966382 | 75.83547 | TRUE | HP_31891 | TGCTGTTG | GGCTTACA | NM_138596 | D segment, Chr 13, Wayne State University 50, expressed | D13Wsu50e | 2.13 | SM2477 | h | 9 |
| mPool7_V2MM_71610 | 1.96447 | 2.205692 | TRUE | HP_29348 | TGCTGTTG | CGTAATCT | XM_142620 | similar to hypothetical protein FLJ90396 | | 2.8 | SM2232 | b | 12 |
| mPool2_V2MM_170124 | 1.964206 | 2.205692 | TRUE | HP_39061 | TGCTGTTG | CAGGTAA | AK029830 | trans-acting transcription factor 6 | Sp6 | 2.6 | SM2128 | d | 6 |
| mPool2_V2MM_109155 | 1.963502 | 3.004939 | TRUE | HP_33022 | TGCTGTTG | GAGATCTT | BC057029 | phosphodiesterase 2A, cGMP-stimulated | Pde2a | 2.6 | SM2149 | c | 7 |
| 092308m1_V2MM_170459 | 1.963127 | 3.004939 | TRUE | HP_39095 | TGCTGTTG | GGCTTTAA | XM_283177 | Mus musculus similar to hypothetical protein 3 - rat (LOC328328), mRNA. | | 2.5 | SM2077 | e | 6 |
| 092308m1_V2MM_170459 | 1.963127 | 3.004939 | TRUE | HP_39095 | TGCTGTTG | GGCTTTAA | XM_283177 | Mus musculus similar to hypothetical protein 3 - rat (LOC328328), mR. | | 2.5 | SM2077 | e | 6 |
| mPool7_V2MM_87336 | 1.962984 | 2.205692 | TRUE | HP_30879 | TGCTGTTG | GCATGTAT | NM_175141 | RIKEN cDNA 1810030O07 gene | 1810030O07Rik | 2.9 | SM2269 | f | 11 |
| mPool2_V2MM_123668 | 1.962399 | 3.004939 | TRUE | HP_34452 | TGCTGTTG | GGTATATC | XM_143088 | Mus musculus similar to small nuclear ribonucleic protein (LOC229081), mRNA. | | 2.11 | SM2371 | g | 11 |
| 092308m1_V2MM_71710 | 1.961597 | 2.205692 | TRUE | HP_29358 | TGCTGTTG | CTACCTTG | NM_025695 | SMC6 structural maintence of chromosomes 6-like 1 (yea | Smc6l1 | 2.3 | SM2016 | e | 2 |
| mPool2_V2MM_100109 | 1.96141 | 3.004939 | TRUE | HP_32130 | TGCTGTTG | CATTTCTA | NM_009546 | tripartite motif protein 25 | Trim25 | 2.7 | SM2158 | g | 10 |
| 092308m1_V2MM_136270 | 1.960709 | 2.205692 | TRUE | HP_35699 | TGCTGTTG | GGAACAA | XM_151594 | Mus musculus LOC213763 (LOC213763), mR. | | 2.4 | SM2039 | h | 8 |
| 092308m1_V2MM_136270 | 1.960709 | 2.205692 | TRUE | HP_35699 | TGCTGTTG | GGAACAA | XM_151594 | Mus musculus LOC213763 (LOC213763), mRNA. | | 2.4 | SM2039 | h | 8 |
| 092308m1_V2MM_83740 | 1.960038 | 1.959154 | TRUE | HP_30526 | TGCTGTTG | CTGAGTGT | NM_024288 | RIKEN cD 1110007A06 gene | 1110007A06Rik | 2.6 | SM2113 | c | 8 |
| 092308m1_V2MM_170903 | 1.95963 | 3.004939 | TRUE | HP_39139 | TGCTGTTG | CTTCCTAA | AK046068 | RIKEN cD A330033J07 gene | A330033J07Rik | 2.5 | SM2091 | b | 9 |
| 092308m1_V2MM_132136 | 1.956419 | 2.205692 | TRUE | HP_35289 | TGCTGTTG | CTCTCCCG | XM_148108 | RIKEN cD 4930451G09 gene | 4930451G09Rik | 2.6 | SM2103 | e | 8 |
| mPool7_V2MM_203240 | 1.956169 | 3.004939 | TRUE | HP_42120 | TGCTGTTG | GGAAACG | XM_136539 | Mus musculus similar to HMG-box containing protein 1 - rat (LOC226885), mRNA | | 2.1 | SM2313 | a | 7 |
| mPool2_V2MM_139675 | 1.953193 | 2.205692 | TRUE | HP_36039 | TGCTGTTG | CTCTATTC | XM_155296 | Mus musculus LOC211057 (LOC211057), mR. | | 2.1 | SM2336 | c | 3 |
| mPool2_V2MM_139675 | 1.953193 | 2.205692 | TRUE | HP_36039 | TGCTGTTG | CTCTATTC | XM_155296 | Mus musculus LOC211057 (LOC211057), mRNA. | | 2.1 | SM2336 | c | 3 |
| mPool2_V2MM_168754 | 1.952362 | 2.205692 | TRUE | HP_38925 | TGCTGTTG | CTTAGCAT | XM_286058 | Mus musculus similar to putative pheromone receptor [Rattus norvegicus] (LOC3 | | 2.6 | SM2131 | c | 10 |
| 092308m1_V2MM_147048 | 1.952156 | 2.205692 | TRUE | HP_36775 | TGCTGTTG | CACAAAG | XM_162569 | Mus musculus LOC243509 (LOC243509), mRNA. | | 2.4 | SM2041 | e | 9 |
| 092308m1_V2MM_147048 | 1.952156 | 2.205692 | TRUE | HP_36775 | TGCTGTTG | CACAAAG | XM_162569 | Mus musculus LOC243509 (LOC243509), mR. | | 2.4 | SM2041 | e | 9 |
| mPool7_V2MM_73386 | 1.951822 | 3.004939 | TRUE | HP_29522 | TGCTGTTG | GGAAATGT | NM_025435 | RIKEN cDNA 1500006O09 gene | 1500006O09Rik | 2.8 | SM2234 | c | 12 |
| mPool7_V2MM_182325 | 1.950208 | 3.004939 | TRUE | HP_40279 | TGCTGTTG | CAGTCAAT | XM_288825 | Mus musculus LOC332298 (LOC332298), mR. | | 2.5 | SM2062 | c | 5 |
| mPool7_V2MM_182325 | 1.950208 | 3.004939 | TRUE | HP_40279 | TGCTGTTG | CAGTCAAT | XM_288825 | Mus musculus LOC332298 (LOC332298), mRNA. | | 2.5 | SM2062 | c | 5 |
| mPool7_V2MM_202377 | 1.946956 | 0 | TRUE | HP_92884 | TGCTGTTG | CTAAGAAT | XM_128090 | RIKEN cDNA D330037H05 gene | D330037H05Rik | 2.9 | SM2296 | d | 11 |
| 092308m3_V2MM_47630 | 1.945047 | 75.83547 | TRUE | HP_27010 | TGCTGTTG | CTGTGGCT | NM_138592 | ubiquitin specific protease 39 | Usp39 | 2.12 | SM2416 | b | 5 |
| 092308m1_V2MM_140408 | 1.944185 | 2.205692 | TRUE | HP_36112 | TGCTGTTG | CAGTTTAA | XM_155942 | Mus musculus similar to Voltage-dependent anion-selective channel protein 3 (V | | 2.4 | SM2038 | h | 11 |
| mPool2_V2MM_146584 | 1.943609 | 3.004939 | TRUE | HP_36729 | TGCTGTTG | CATAATTG | XM_162162 | Mus musculus LOC243184 (LOC243184), mR. | | 2.5 | SM2058 | a | 9 |
| mPool2_V2MM_146584 | 1.943609 | 3.004939 | TRUE | HP_36729 | TGCTGTTG | CATAATTG | XM_162162 | Mus musculus LOC243184 (LOC243184), mRNA. | | 2.5 | SM2058 | a | 9 |
| mPool7_V2MM_105217 | 1.942843 | 3.004939 | TRUE | HP_32633 | TGCTGTTG | CAAGTTAT | XM_130324 | RIKEN cDNA 2810410A08 gene | 2810410A08Rik | 2.1 | SM2306 | h | 4 |
| mPool7_V2MM_107148 | 1.94244 | 2.205692 | TRUE | HP_32823 | TGCTGTTG | CCCAGTGC | NM_021604 | agrin | Agrn | 2.1 | SM2301 | e | 5 |
| mPool2_V2MM_121702 | 1.940567 | 2.205692 | TRUE | HP_34258 | TGCTGTTG | GCAAGAAT | XM_141796 | Mus musculus similar to RIKEN cDNA 4933402E13 [Mus musculus] (LOC236842), | | 2.5 | SM2099 | d | 3 |
| mPool2_V2MM_121702 | 1.940567 | 2.205692 | TRUE | HP_34258 | TGCTGTTG | GCAAGAAT | XM_141796 | Mus musculus similar to RIKEN cD 4933402E13 [Mus musculus] (LOC236842), mR | | 2.5 | SM2099 | d | 3 |
| mPool7_V2MM_197536 | 1.940227 | 3.004939 | TRUE | HP_41560 | TGCTGTTG | GGTGTAAT | NM_011264 | REV3-like, catalytic subunit of DNA polymerase zeta RAD | Rev3l | 2.9 | SM2253 | a | 4 |
| 092308m1_V2MM_154123 | 1.939896 | 3.004939 | TRUE | HP_37476 | TGCTGTTG | GCTGTTAT | NM_201354 | gene model 672, (NCBI) | Gm672 | 2.5 | SM2091 | e | 12 |
| mPool2_V2MM_88570 | 1.939516 | 3.004939 | TRUE | HP_31000 | TGCTGTTG | CACTTAGT | NM_175436 | zinc finger protein 526 | Zfp526 | 2.6 | SM2147 | e | 3 |
| mPool2_V2MM_106861 | 1.938486 | 3.004939 | TRUE | HP_32795 | TGCTGTTG | GTTTATAT | AK053970 | RIKEN cDNA 4930538K18 gene | 4930538K18Rik | 2.6 | SM2149 | a | 11 |
| 092308m3_V2MM_113015 | 1.938188 | 75.83547 | TRUE | NA | NA | NA | NA | NA | NA | NA | NA | NA | NA |

| | | | | | | | | | | | | | |
|---|---|---|---|---|---|---|---|---|---|---|---|---|---|
| mPool7_V2MM_84065 | 1.937373 | 2.205692 | TRUE | HP_30558 | TGCTGTTG | GTCATCTA | NM_026808 | RIKEN cDNA 1110028A07 gene | 1110028A07Rik | 2.8 | SM2250 | e | 3 |
| mPool7_V2MM_88813 | 1.936547 | 3.004939 | TRUE | HP_31024 | TGCTGTTG | GACCTTTC | NM_175499 | SLIT and NTRK-like family, member 6 | Slitrk6 | 2.9 | SM2252 | e | 11 |
| mPool7_V2MM_88813 | 1.936547 | 3.004939 | TRUE | HP_31024 | TGCTGTTG | GACCTTTC | NM_175499 | SLIT and NTRK-like family, member 6 | Slitrk6 | 2.9 | SM2252 | e | 11 |
| 092308m1_V2MM_112685 | 1.936069 | 2.205692 | TRUE | HP_29847 | TGCTGTTG | CTATTAGC | XM_486245 | similar to Rpl7a protein | | 2.4 | SM2028 | b | 1 |
| 092308m1_V2MM_112685 | 1.936069 | 2.205692 | TRUE | HP_29847 | TGCTGTTG | CTATTAGC | XM_486245 | similar to Rpl7a protein | | 2.4 | SM2028 | b | 1 |
| 092308m1_V2MM_64665 | 1.934134 | 3.004939 | TRUE | HP_28671 | TGCTGTTG | CTTATTAC | NM_009863 | cell division cycle 7 (S. cerevisiae) | Cdc7 | 2.4 | SM2022 | e | 9 |
| mPool2_V2MM_215307 | 1.931471 | 1.959154 | TRUE | HP_43286 | TGCTGTTG | GATGTGA | XM_147068 | Mus musculus similar to glyceraldehyde-3-phosphate dehydrogenase [Mus musc | | 2.11 | SM2358 | e | 12 |
| 092308m1_V2MM_172849 | 1.930505 | 3.004939 | TRUE | HP_39333 | TGCTGTTG | CCATATAT | AK085159 | RIKEN cDNA 4933403F05 gene | 4933403F05Rik | 2.5 | SM2069 | f | 8 |
| 092308m1_V2MM_172849 | 1.930505 | 3.004939 | TRUE | HP_39333 | TGCTGTTG | CCATATAT | AK085159 | RIKEN cD 4933403F05 gene | 4933403F05Rik | 2.5 | SM2069 | f | 8 |
| mPool2_V2MM_109617 | 1.929899 | 2.205692 | TRUE | HP_33067 | TGCTGTTG | CTCACTCA | XM_486079 | RIKEN cDNA 4930453L07 gene | 4930453L07Rik | 2.7 | SM2153 | b | 3 |
| 092308m3_V2MM_33008 | 1.929111 | 75.83547 | TRUE | HP_25587 | TGCTGTTG | CAATGTGT | NM_008542 | MAD homolog 6 (Drosophila) | Smad6 | 2.12 | SM2429 | a | 10 |
| mPool2_V2MM_163177 | 1.928686 | 2.205692 | TRUE | HP_38373 | TGCTGTTG | GGCTTTAA | XM_284694 | Mus musculus hypothetical gene supported by AK039538 (LOC331396), mRNA. | | 2.6 | SM2131 | h | 8 |
| 092308m1_V2MM_136154 | 1.928177 | 2.205692 | TRUE | HP_35687 | TGCTGTTG | GTTTATAA | XM_151438 | Mus musculus LOC241149 (LOC241149), mR. | | 2.5 | SM2059 | g | 3 |
| mPool2_V2MM_212123 | 1.926596 | 3.004939 | TRUE | HP_42980 | TGCTGTTG | GGAGAGA | XM_145549 | similar to hypothetical protein FLJ38281 | | 2.11 | SM2364 | e | 10 |
| mPool2_V2MM_116075 | 1.923422 | 3.004939 | TRUE | HP_33703 | TGCTGTTG | GTGCTTAT | XM_138743 | Mus musculus similar to Gag [Ovis aries] (LOC218358), mRNA. | | 2.7 | SM2154 | e | 2 |
| mPool2_V2MM_92325 | 1.923364 | 3.004939 | TRUE | HP_31367 | TGCTGTTG | CAGCGAT( | NM_178689 | RIKEN cDNA 6430537H07 gene | 6430537H07Rik | 2.6 | SM2137 | d | 6 |
| mPool2_V2MM_108460 | 1.922955 | 1.959154 | TRUE | HP_32953 | TGCTGTTG | CTTATTTG | NM_019697 | potassium voltage-gated channel, Shal-related family, me | Kcnd2 | 2.1 | SM2309 | d | 9 |
| mPool2_V2MM_144897 | 1.9224 | 3.004939 | TRUE | HP_36560 | TGCTGTTG | CAACCTTT | XM_160409 | Mus musculus LOC242190 (LOC242190), mR. | | 2.1 | SM2334 | c | 11 |
| mPool2_V2MM_144897 | 1.9224 | 3.004939 | TRUE | HP_36560 | TGCTGTTG | CAACCTTT | XM_160409 | Mus musculus LOC242190 (LOC242190), mRNA. | | 2.1 | SM2334 | c | 11 |
| mPool2_V2MM_215055 | 1.921813 | 2.205692 | TRUE | HP_43262 | TGCTGTTG | CCTTCGTT | XM_141782 | Mus musculus similar to H3 histone, family 3B [Mus musculus] (LOC245434), mRI | | 2.11 | SM2355 | e | 1 |
| 092308m1_V2MM_185834 | 1.921408 | 3.004939 | TRUE | HP_40630 | TGCTGTTG | CCTGCATT | XM_289437 | Mus musculus LOC333103 (LOC333103), mR. | | 2.5 | SM2094 | g | 2 |
| 092308m1_V2MM_139690 | 1.921377 | 3.004939 | TRUE | HP_36041 | TGCTGTTG | CAGACAAT | XM_155311 | Mus musculus LOC211071 (LOC211071), mR. | | 2.5 | SM2056 | f | 9 |
| 092308m1_V2MM_139690 | 1.921377 | 3.004939 | TRUE | HP_36041 | TGCTGTTG | CAGACAAT | XM_155311 | Mus musculus LOC211071 (LOC211071), mRNA. | | 2.5 | SM2056 | f | 9 |
| mPool2_V2MM_126633 | 1.919759 | 3.004939 | TRUE | HP_34747 | TGCTGTTG | GCTGCATT | XM_144678 | Mus musculus similar to ribosomal protein L19 [Rattus norvegicus] (LOC213268), | | 2.11 | SM2360 | g | 12 |
| mPool2_V2MM_212264 | 1.918986 | 1.959154 | TRUE | HP_42993 | TGCTGTTG | CTCTTGGT | XM_143544 | Mus musculus similar to CL2BB protein - rat (LOC215282), mRNA. | | 2.11 | SM2352 | d | 1 |
| mPool7_V2MM_79053 | 1.918247 | 3.004939 | TRUE | HP_30072 | TGCTGTTG | GAGTGGT | NM_025453 | RIKEN cDNA 1810018L02 gene | 1810018L02Rik | 2.8 | SM2236 | e | 6 |
| 092308m1_V2MM_177264 | 1.91691 | 3.004939 | TRUE | HP_39774 | TGCTGTTG | GCATTTAC | AK045718 | RIKEN cD C030044C12 gene | C030044C12Rik | 2.5 | SM2078 | h | 8 |
| mPool2_V2MM_986 | 1.915567 | 2.205692 | TRUE | HP_22464 | TGCTGTTG | CAGTAAAT | XM_484397 | diacylglycerol kinase, eta | Dgkh | 2.11 | SM2379 | g | 6 |
| mPool7_V2MM_81233 | 1.911495 | 2.205692 | TRUE | HP_30284 | TGCTGTTG | GGTGTAGT | NM_172815 | RIKEN cDNA 2610028F08 gene | 2610028F08Rik | 2.8 | SM2248 | c | 4 |
| 092308m3_V2MM_96391 | 1.911202 | 75.83547 | TRUE | HP_31764 | TGCTGTTG | GAAATGTT | AK015302 | RIKEN cD 4933432K03 gene | 4933432K03Rik | 2.9 | SM2251 | d | 3 |
| 092308m3_V2MM_96391 | 1.911202 | 75.83547 | TRUE | HP_31764 | TGCTGTTG | GAAATGTT | AK015302 | RIKEN cDNA 4933432K03 gene | 4933432K03Rik | 2.9 | SM2251 | d | 3 |
| mPool7_V2MM_180607 | 1.910923 | 3.004939 | TRUE | HP_40108 | TGCTGTTG | CTATGAAT | XM_358439 | gene model 1573, (NCBI) | Gm1573 | 2.5 | SM2065 | h | 7 |
| mPool7_V2MM_180607 | 1.910923 | 3.004939 | TRUE | HP_40108 | TGCTGTTG | CTATGAAT | XM_358439 | gene model 1573, (NCBI) | Gm1573 | 2.5 | SM2065 | h | 7 |
| mPool2_V2MM_105912 | 1.910609 | 3.004939 | TRUE | HP_32701 | TGCTGTTG | CAGAGAC | AK015427 | RIKEN cDNA 4930449A18 gene | 4930449A18Rik | 2.7 | SM2158 | e | 11 |
| mPool6_V2MM_69203 | 1.910461 | 75.83547 | TRUE | HP_29114 | TGCTGTTG | GCATCTGC | NM_010319 | guanine nucleotide binding protein (G protein), gamma 7 | Gng7 | 2.7 | SM2175 | g | 5 |
| mPool7_V2MM_204746 | 1.909558 | 3.004939 | TRUE | HP_42263 | TGCTGTTG | CTGTCAAA | XM_129787 | similar to ALY | | 2.1 | SM2303 | e | 11 |
| 092308m1_V2MM_165290 | 1.908716 | 2.205692 | TRUE | HP_38582 | TGCTGTTG | CTGGTGAT | XM_484618 | similar to hypothetical protein FLJ38755 | | 2.6 | SM2115 | h | 6 |
| mPool6_V2MM_35599 | 1.907829 | 75.83547 | TRUE | HP_25839 | TGCTGTTG | CTAGGGA | NM_152804 | polo-like kinase 2 (Drosophila) | Plk2 | 2.7 | SM2183 | h | 9 |
| 092308m1_V2MM_136543 | 1.905996 | 2.205692 | TRUE | HP_35726 | TGCTGTTG | GTTATCTT | XM_151819 | Mus musculus LOC213261 (LOC213261), mRNA. | | 2.4 | SM2034 | c | 9 |
| 092308m1_V2MM_136543 | 1.905996 | 2.205692 | TRUE | HP_35726 | TGCTGTTG | GTTATCTT | XM_151819 | Mus musculus LOC213261 (LOC213261), mR. | | 2.4 | SM2034 | c | 9 |
| 092308m1_V2MM_142682 | 1.904666 | 3.004939 | TRUE | HP_36339 | TGCTGTTG | CTCTCTGT | XM_157813 | Mus musculus LOC240599 (LOC240599), mR. | | 2.4 | SM2049 | c | 10 |
| mPool7_V2MM_112685 | 1.901594 | 2.205692 | TRUE | HP_29847 | TGCTGTTG | CTATTAGC | XM_486245 | similar to Rpl7a protein | | 2.4 | SM2028 | b | 1 |
| mPool7_V2MM_112685 | 1.901594 | 2.205692 | TRUE | HP_29847 | TGCTGTTG | CTATTAGC | XM_486245 | similar to Rpl7a protein | | 2.4 | SM2028 | b | 1 |
| 092308m1_V2MM_161230 | 1.901168 | 2.205692 | TRUE | HP_38180 | TGCTGTTG | CCAGGTCT | XM_283818 | cadherin-like 26 | Cdh26 | 2.6 | SM2115 | h | 9 |
| mPool7_V2MM_81478 | 1.898915 | 3.004939 | TRUE | HP_30308 | TGCTGTTG | GTCAGTTT | NM_172815 | RIKEN cDNA 2610028F08 gene | 2610028F08Rik | 2.8 | SM2229 | g | 1 |
| mPool2_V2MM_210988 | 1.898246 | 3.004939 | TRUE | HP_37786 | TGCTGTTG | CCTATGTA | XM_205092 | Mus musculus similar to Zinc finger protein 180 (HHZ168) (LOC277541), mRNA. | | 2.11 | SM2364 | e | 9 |
| mPool2_V2MM_106763 | 1.896387 | 3.004939 | TRUE | HP_32785 | TGCTGTTG | CTTGCTTA | BC037594 | cDNA sequence BC037594 | BC037594 | 2.7 | SM2152 | h | 12 |
| mPool2_V2MM_87110 | 1.895813 | 3.004939 | TRUE | HP_30857 | TGCTGTTG | CAGATAAT | NM_175024 | RIKEN cDNA G630049C14 gene | G630049C14Rik | 2.6 | SM2149 | c | 8 |
| 092308m1_V2MM_123096 | 1.895295 | 3.004939 | TRUE | HP_34395 | TGCTGTTG | CACAGGT( | XM_142669 | similar to 60 kDa heat shock protein, mitochondrial precursor (Hsp60) (60 kDa ch | | 2.6 | SM2102 | b | 6 |
| 092308m1_V2MM_123096 | 1.895295 | 3.004939 | TRUE | HP_34395 | TGCTGTTG | CACAGGT( | XM_142669 | similar to 60 kDa heat shock protein, mitochondrial precursor (Hsp60) (60 kDa ch | | 2.6 | SM2102 | b | 6 |
| mPool2_V2MM_100653 | 1.895155 | 2.205692 | TRUE | HP_32183 | TGCTGTTG | CAATTGGA | AK032388 | serologically defined colon cancer antigen 1 | Sdccag1 | 2.7 | SM2154 | a | 2 |
| mPool7_V2MM_183702 | 1.894866 | 2.205692 | TRUE | HP_40417 | TGCTGTTG | CATCAAAC | AK018908 | RIKEN cDNA 1700085A12 gene | 1700085A12Rik | 2.9 | SM2275 | f | 7 |
| mPool2_V2MM_12204 | 1.893768 | 2.205692 | TRUE | HP_23560 | TGCTGTTG | GTGACAA | NM_134191 | vomeronasal 1 receptor, E2 | V1re2 | 2.11 | SM2377 | f | 10 |

| | | | | | | | | | | | | | |
|---|---|---|---|---|---|---|---|---|---|---|---|---|---|
| mPool2_V2MM_110673 | 1.893199 | 3.004939 | TRUE | HP_33170 | TGCTGTTG | GAGATTGT | BC051163 | mannose phosphate isomerase 1 | Mpi1 | 2.7 | SM2162 | b | 11 |
| mPool7_V2MM_101164 | 1.893125 | 3.004939 | TRUE | HP_32233 | TGCTGTTG | GACAGACA | AK046032 | RIKEN cDNA 2900024O10 gene | 2900024O10Rik | 2.6 | SM2150 | b | 10 |
| mPool7_V2MM_143114 | 1.892393 | 3.004939 | TRUE | HP_36382 | TGCTGTTG | GGATTACT | XM_158386 | Mus musculus LOC241409 (LOC241409), mRNA. | | 2.4 | SM2043 | a | 12 |
| mPool7_V2MM_143114 | 1.892393 | 3.004939 | TRUE | HP_36382 | TGCTGTTG | GGATTACT | XM_158386 | Mus musculus LOC241409 (LOC241409), mR. | | 2.4 | SM2043 | a | 12 |
| mPool7_V2MM_97784 | 1.89095 | 3.004939 | TRUE | HP_31902 | TGCTGTTG | CTATAGAA | XM_111892 | RIKEN cDNA 1700123J19 gene | 1700123J19Rik | 2.9 | SM2256 | d | 2 |
| mPool7_V2MM_97784 | 1.89095 | 3.004939 | TRUE | HP_31902 | TGCTGTTG | CTATAGAA | XM_111892 | RIKEN cD 1700123J19 gene | 1700123J19Rik | 2.9 | SM2256 | d | 2 |
| mPool2_V2MM_35528 | 1.890658 | 2.205692 | TRUE | HP_25832 | TGCTGTTG | CTAAGACA | NM_027457 | RIKEN cDNA 5730437N04 gene | 5730437N04Rik | 2.11 | SM2389 | a | 6 |
| mPool7_V2MM_200071 | 1.890457 | 3.004939 | TRUE | HP_41810 | TGCTGTTG | GGACTTTA | XM_286571 | Mus musculus hypothetical gene supported by AK051599 (LOC328315), mRNA. | | 2.9 | SM2289 | d | 5 |
| mPool2_V2MM_141195 | 1.889642 | 1.959154 | TRUE | HP_36190 | TGCTGTTG | GACAAATT | XM_156514 | Mus musculus LOC239894 (LOC239894), mRNA. | | 2.4 | SM2042 | d | 4 |
| mPool2_V2MM_141195 | 1.889642 | 1.959154 | TRUE | HP_36190 | TGCTGTTG | GACAAATT | XM_156514 | Mus musculus LOC239894 (LOC239894), mR. | | 2.4 | SM2042 | d | 4 |
| 092308m1_V2MM_153410 | 1.889635 | 3.004939 | TRUE | HP_37406 | TGCTGTTG | CTGTCCGT | XM_196098 | Mus musculus LOC268660 (LOC268660), mR. | | 2.6 | SM2121 | e | 12 |
| mPool7_V2MM_151904 | 1.888824 | 3.004939 | TRUE | HP_37257 | TGCTGTTG | GCGTTTAT | XM_194993 | olfactory receptor 171 | Olfr171 | 2.1 | SM2327 | c | 4 |
| mPool7_V2MM_176524 | 1.885764 | 1.959154 | TRUE | HP_39700 | TGCTGTTG | CTCCATTG | XM_489175 | hypothetical gene supported by AK049985 | | 2.5 | SM2089 | c | 4 |
| mPool7_V2MM_176524 | 1.885764 | 1.959154 | TRUE | HP_39700 | TGCTGTTG | CTCCATTG | XM_489175 | hypothetical gene supported by AK049985 | | 2.5 | SM2089 | c | 4 |
| mPool2_V2MM_87591 | 1.884702 | 3.004939 | TRUE | HP_30904 | TGCTGTTG | GAACGTCT | BC060730 | ring finger protein 38 | Rnf38 | 2.6 | SM2144 | b | 3 |
| mPool7_V2MM_176192 | 1.88464 | 2.205692 | TRUE | HP_39667 | TGCTGTTG | CTTATTCC | XM_287620 | Mus musculus hypothetical gene supported by AK036508 (LOC330383), mRNA. | | 2.9 | SM2292 | b | 9 |
| mPool7_V2MM_174946 | 1.884374 | 2.205692 | TRUE | HP_39543 | TGCTGTTG | CGCTGTAC | XM_287341 | Mus musculus hypothetical gene supported by AK080995 (LOC329843), mRNA. | | 2.9 | SM2284 | a | 3 |
| 092308m1_V2MM_179537 | 1.884121 | 3.004939 | TRUE | HP_40001 | TGCTGTTG | CAGTTTGC | XM_288350 | Mus musculus LOC331670 (LOC331670), mR. | | 2.5 | SM2094 | b | 6 |
| 092308m1_V2MM_69539 | 1.882686 | 2.205692 | TRUE | HP_29146 | TGCTGTTG | GCTATTTA | NM_009923 | cyclic nucleotide phosphodiesterase 1 | Cnp1 | 2.4 | SM2019 | a | 12 |
| 092308m3_V2MM_259618 | 1.881581 | 75.83547 | TRUE | HP_31891 | TGCTGTTG | GCTTACAC | NM_138596 | D segment, Chr 13, Wayne State University 50, expressec | D13Wsu50e | 2.13 | SM2453 | b | 6 |
| 092308m1_V2MM_173441 | 1.881264 | 1.959154 | TRUE | HP_39392 | TGCTGTTG | CAAACATA | AK035520 | hypothetical protein 9530060I07 | | 2.5 | SM2081 | c | 2 |
| 092308m1_V2MM_173441 | 1.881264 | 1.959154 | TRUE | HP_39392 | TGCTGTTG | CAAACATA | AK035520 | hypothetical protein 9530060I07 | | 2.5 | SM2081 | c | 2 |
| mPool7_V2MM_115972 | 1.880251 | 3.004939 | TRUE | HP_33692 | TGCTGTTG | CTCCTTCA | XM_138705 | Mus musculus similar to RNA polymerase II transcriptional coactivator [Mus mus | | 2.1 | SM2301 | h | 1 |
| mPool2_V2MM_3018 | 1.879415 | 3.004939 | TRUE | HP_86735 | TGCTGTTG | CTGTTTGC | NM_011448 | SRY-box containing gene 9 | Sox9 | 2.11 | SM2376 | g | 3 |
| 092308m1_V2MM_153317 | 1.877659 | 2.205692 | TRUE | HP_37397 | TGCTGTTG | GTCAGTAT | XM_484109 | RIKEN cD 4930417G10 gene | 4930417G10Rik | 2.6 | SM2118 | h | 5 |
| mPool7_V2MM_101389 | 1.876482 | 3.004939 | TRUE | HP_32256 | TGCTGTTG | GTGTTTAC | AK007938 | RIKEN cDNA 1810062O18 gene | 1810062O18Rik | 2.1 | SM2305 | h | 2 |
| mPool2_V2MM_12514 | 1.874201 | 3.004939 | TRUE | HP_23590 | TGCTGTTG | CAAGTTAA | AF463504 | metastasis associated 1 | Mta1 | 2.11 | SM2376 | c | 3 |
| mPool7_V2MM_198072 | 1.87376 | 2.205692 | TRUE | HP_41613 | TGCTGTTG | CTATCATT | XM_287636 | Mus musculus hypothetical gene supported by AK039232 (LOC330394), mRNA. | | 2.9 | SM2279 | d | 2 |
| 092308m3_V2MM_91799 | 1.873063 | 59.99801 | TRUE | HP_31316 | TGCTGTTG | CCTCTAAT | NM_178595 | RIKEN cD 2210013M04 gene | 2210013M04Rik | 2.13 | SM2479 | f | 5 |
| 092308m1_V2MM_150297 | 1.870964 | 1.959154 | TRUE | HP_37099 | TGCTGTTG | GAATTATT | BC023730 | cell division cycle 27 homolog (S. cerevisiae) | Cdc27 | 2.4 | SM2039 | e | 7 |
| mPool2_V2MM_211266 | 1.870358 | 3.004939 | TRUE | HP_34019 | TGCTGTTG | CTCTCAAT | XM_486101 | similar to hypothetical protein FLJ25801 | | 2.1 | SM2336 | e | 8 |
| mPool7_V2MM_108856 | 1.868995 | 3.004939 | TRUE | HP_32992 | TGCTGTTG | CTGAGATC | AK002462 | RIKEN cDNA 0610010E21 gene | 0610010E21Rik | 2.1 | SM2317 | f | 12 |
| mPool2_V2MM_90839 | 1.868748 | 2.205692 | TRUE | HP_31225 | TGCTGTTG | CTACAAAC | NM_177869 | expressed sequence AI847670 | AI847670 | 2.6 | SM2145 | g | 8 |
| mPool2_V2MM_141774 | 1.868049 | 2.205692 | TRUE | HP_36248 | TGCTGTTG | CTTCAATA | XM_156976 | Mus musculus LOC210827 (LOC210827), mRNA. | | 2.4 | SM2049 | b | 6 |
| mPool2_V2MM_141774 | 1.868049 | 2.205692 | TRUE | HP_36248 | TGCTGTTG | CTTCAATA | XM_156976 | Mus musculus LOC210827 (LOC210827), mR. | | 2.4 | SM2049 | b | 6 |
| 092308m1_V2MM_92527 | 1.867673 | 3.004939 | TRUE | HP_31387 | TGCTGTTG | CCAATTAA | NM_178727 | RIKEN cD D630039A03 gene | D630039A03Rik | 2.6 | SM2106 | c | 11 |
| 092308m1_V2MM_92527 | 1.867673 | 3.004939 | TRUE | HP_31387 | TGCTGTTG | CCAATTAA | NM_178727 | RIKEN cDNA D630039A03 gene | D630039A03Rik | 2.6 | SM2106 | c | 11 |
| mPool7_V2MM_79585 | 1.865885 | 3.004939 | TRUE | HP_30124 | TGCTGTTG | GCATGTTT | NM_011370 | cytoplasmic FMR1 interacting protein 1 | Cyfip1 | 2.8 | SM2243 | b | 2 |
| 092308m3_V2MM_196667 | 1.864094 | 75.83547 | TRUE | HP_41476 | TGCTGTTG | CCAGCCTC | XM_113133 | Mus musculus LOC195552 (LOC195552), mR. | | 2.13 | SM2476 | f | 9 |
| mPool2_V2MM_106855 | 1.86392 | 1.959154 | TRUE | HP_32794 | TGCTGTTG | CGAACACT | XM_131605 | RIKEN cDNA 4921539E11 gene | 4921539E11Rik | 2.7 | SM2155 | e | 7 |
| mPool7_V2MM_83283 | 1.86167 | 3.004939 | TRUE | HP_30483 | TGCTGTTG | CTGTTCAG | NM_011914 | Wolf-Hirschhorn syndrome candidate 2 (human) | Whsc2 | 2.9 | SM2266 | g | 6 |
| mPool7_V2MM_83283 | 1.86167 | 3.004939 | TRUE | HP_30483 | TGCTGTTG | CTGTTCAG | NM_011914 | Wolf-Hirschhorn syndrome candidate 2 (human) | Whsc2 | 2.9 | SM2266 | g | 6 |
| mPool7_V2MM_112410 | 1.86156 | 3.004939 | TRUE | HP_33341 | TGCTGTTG | CAGGAAAC | XM_136489 | similar to fatty acid desaturase | | 2.9 | SM2297 | d | 12 |
| mPool7_V2MM_112410 | 1.86156 | 3.004939 | TRUE | HP_33341 | TGCTGTTG | CAGGAAAC | XM_136489 | similar to fatty acid desaturase | | 2.9 | SM2297 | d | 12 |
| 092308m1_V2MM_179484 | 1.860226 | 2.205692 | TRUE | HP_39995 | TGCTGTTG | CACTAATT | XM_288341 | Mus musculus LOC331656 (LOC331656), mR. | | 2.5 | SM2076 | b | 2 |
| 092308m1_V2MM_179484 | 1.860226 | 2.205692 | TRUE | HP_39995 | TGCTGTTG | CACTAATT | XM_288341 | Mus musculus LOC331656 (LOC331656), mRNA. | | 2.5 | SM2076 | b | 2 |
| mPool7_V2MM_88202 | 1.858436 | 3.004939 | TRUE | HP_30964 | TGCTGTTG | CAATATTT | NM_175342 | RIKEN cD C330003B14 gene | C330003B14Rik | 2.9 | SM2267 | h | 7 |
| mPool7_V2MM_88202 | 1.858436 | 3.004939 | TRUE | HP_30964 | TGCTGTTG | CAATATTT | NM_175342 | RIKEN cDNA C330003B14 gene | C330003B14Rik | 2.9 | SM2267 | h | 7 |
| 092308m1_V2MM_152938 | 1.857777 | 3.004939 | TRUE | HP_37360 | TGCTGTTG | GAGTGAA | XM_195621 | Mus musculus similar to squamous cell carcinoma antigen 2 [Mus musculus] (LOC | | 2.6 | SM2117 | g | 7 |
| mPool7_V2MM_109155 | 1.857661 | 2.205692 | TRUE | HP_33022 | TGCTGTTG | GAGATCTT | BC057029 | phosphodiesterase 2A, cGMP-stimulated | Pde2a | 2.6 | SM2149 | c | 7 |
| 092308m1_V2MM_147923 | 1.857462 | 3.004939 | TRUE | HP_36863 | TGCTGTTG | CAATTTCC | XM_163354 | Mus musculus LOC215451 (LOC215451), mRNA. | | 2.5 | SM2052 | e | 6 |
| 092308m1_V2MM_147923 | 1.857462 | 3.004939 | TRUE | HP_36863 | TGCTGTTG | CAATTTCC | XM_163354 | Mus musculus LOC215451 (LOC215451), mR. | | 2.5 | SM2052 | e | 6 |
| 092308m1_V2MM_174387 | 1.856481 | 2.205692 | TRUE | HP_39487 | TGCTGTTG | GGTTAGTA | XM_287220 | Mus musculus hypothetical gene supported by AK045957 (LOC329604), mR. | | 2.5 | SM2095 | d | 12 |

| | | | | | | | | | | | | | |
|---|---|---|---|---|---|---|---|---|---|---|---|---|---|
| mPool2_V2MM_145646 | 1.855991 | 3.004939 | TRUE | HP_36635 | TGCTGTTG | CTGCAAGT | XM_161218 | hypothetical gene supported by AK039704 | | 2.1 | SM2341 | h | 12 |
| mPool7_V2MM_206248 | 1.855908 | 2.205692 | TRUE | HP_42408 | TGCTGTTG | CTCTGCTA | AK042897 | RIKEN cDNA A930021C24 gene | A930021C24Rik | 2.9 | SM2300 | d | 7 |
| 092308m1_V2MM_160405 | 1.855381 | 2.205692 | TRUE | HP_38098 | TGCTGTTG | CAGTCTAC | AK088434 | RIKEN cD E430016J11 gene | E430016J11Rik | 2.6 | SM2118 | b | 4 |
| mPool7_V2MM_102528 | 1.854369 | 3.004939 | TRUE | HP_32367 | TGCTGTTG | CTGCCATC | AK036186 | Wilms tumour 1-associating protein | Wtap | 2.1 | SM2303 | g | 5 |
| mPool4_V2MM_162346 | 1.854303 | 75.83547 | TRUE | NA | NA | NA | NA | NA | NA | NA | NA | NA | NA |
| mPool7_V2MM_94539 | 1.854125 | 2.205692 | TRUE | HP_31582 | TGCTGTTG | CAGGAGT | NM_183177 | cDNA sequence BC052046 | BC052046 | 2.9 | SM2253 | e | 1 |
| mPool2_V2MM_213976 | 1.852894 | 3.004939 | TRUE | HP_43159 | TGCTGTTG | CTCTAATG | BC034735 | RIKEN cDNA 2310047A01 gene | 2310047A01Rik | 2.11 | SM2370 | d | 7 |
| mPool2_V2MM_133635 | 1.8528 | 2.205692 | TRUE | HP_35436 | TGCTGTTG | CCTGGAGT | BC050760 | synaptotagmin 6 | Syt6 | 2.11 | SM2357 | h | 9 |
| mPool2_V2MM_127171 | 1.852547 | 2.205692 | TRUE | HP_34800 | TGCTGTTG | GATATACT | XM_145041 | Mus musculus similar to RNA-binding region (RNP1, RRM) containing 2; splicing f | | 2.11 | SM2369 | a | 1 |
| 092308m3_V2MM_222383 | 1.852379 | 75.83547 | TRUE | NA | NA | NA | NA | NA | NA | NA | NA | NA | NA |
| mPool2_V2MM_86707 | 1.852379 | 2.205692 | TRUE | HP_76909 | TGCTGTTG | GTACTGAT | NM_173760 | expressed sequence AW555814 | AW555814 | 2.6 | SM2147 | h | 6 |
| 092308m1_V2MM_177190 | 1.852005 | 2.205692 | TRUE | HP_39767 | TGCTGTTG | GTGCAAAT | XM_287819 | Mus musculus hypothetical gene supported by AK090231 (LOC330797), mR. | | 2.5 | SM2063 | c | 8 |
| 092308m1_V2MM_181893 | 1.848692 | 3.004939 | TRUE | HP_40236 | TGCTGTTG | GTTTATTG | XM_288752 | Mus musculus LOC332202 (LOC332202), mR. | | 2.5 | SM2089 | c | 5 |
| 092308m1_V2MM_181893 | 1.848692 | 3.004939 | TRUE | HP_40236 | TGCTGTTG | GTTTATTG | XM_288752 | Mus musculus LOC332202 (LOC332202), mRNA. | | 2.5 | SM2089 | c | 5 |
| mPool7_V2MM_106336 | 1.847994 | 2.205692 | TRUE | HP_32743 | TGCTGTTG | GTGATAAA | AK004863 | alcohol dehydrogenase 6 (class V), pseudogene 1 | Adh6-ps1 | 2.7 | SM2163 | d | 6 |
| 092308m3_V2MM_53137 | 1.847345 | 62.64426 | TRUE | HP_27550 | TGCTGTTG | GGGCAGA | NM_016658 | galactose-1-phosphate uridyl transferase | Galt | 2.12 | SM2416 | a | 7 |
| mPool7_V2MM_176089 | 1.846342 | 1.959154 | TRUE | HP_39657 | TGCTGTTG | CCAGGAA | XM_287594 | Mus musculus hypothetical gene supported by AK034606 (LOC330311), mRNA. | | 2.5 | SM2063 | h | 12 |
| mPool7_V2MM_176089 | 1.846342 | 1.959154 | TRUE | HP_39657 | TGCTGTTG | CCAGGAA | XM_287594 | Mus musculus hypothetical gene supported by AK034606 (LOC330311), mR. | | 2.5 | SM2063 | h | 12 |
| mPool7_V2MM_98374 | 1.844994 | 3.004939 | TRUE | HP_31960 | TGCTGTTG | CAGAAGA | XM_112433 | PREDICTED: Mus musculus RIKEN cDNA E130307J04 gene (E130307J04Rik), mRNA | | 2.9 | SM2267 | e | 7 |
| 092308m1_V2MM_62022 | 1.844357 | 3.004939 | TRUE | HP_28413 | TGCTGTTG | CACCATGT | BC060160 | transient receptor potential cation channel, subfamily C, | Trpc4 | 2.3 | SM2014 | b | 12 |
| mPool7_V2MM_197843 | 1.844133 | 2.205692 | TRUE | HP_41590 | TGCTGTTG | CAGATTTG | NM_029498 | zinc finger protein 198 | Zfp198 | 2.8 | SM2250 | d | 11 |
| 092308m3_V2MM_50774 | 1.843957 | 35.99881 | TRUE | HP_27318 | TGCTGTTG | CATTTAGT | D83204 | protein tyrosine phosphatase, receptor type, J | Ptprj | 2.12 | SM2418 | b | 4 |
| 092308m1_V2MM_75845 | 1.843484 | 3.004939 | TRUE | HP_29761 | TGCTGTTG | CCGGAAG | NM_010720 | lipase, endothelial | Lipg | 2.3 | SM2016 | g | 8 |
| mPool2_V2MM_139267 | 1.843441 | 2.205692 | TRUE | HP_35998 | TGCTGTTG | GTAGAAAT | XM_154892 | Mus musculus LOC214538 (LOC214538), mRNA. | | 2.1 | SM2342 | c | 2 |
| 092308m3_V2MM_141128 | 1.842728 | 59.99801 | TRUE | HP_36184 | TGCTGTTG | CGGATCCC | XM_156476 | Mus musculus LOC239968 (LOC239968), mR. | | 2.14 | SM2528 | c | 8 |
| mPool2_V2MM_105738 | 1.842222 | 3.004939 | TRUE | HP_32684 | TGCTGTTG | CAGCAATT | AK005136 | phosphatase and actin regulator 3 | Phactr3 | 2.1 | SM2323 | c | 11 |
| mPool2_V2MM_105738 | 1.842222 | 3.004939 | TRUE | HP_32684 | TGCTGTTG | CAGCAATT | AK005136 | phosphatase and actin regulator 3 | Phactr3 | 2.1 | SM2323 | c | 11 |
| 092308m3_V2MM_144135 | 1.841662 | 66.95036 | TRUE | HP_36484 | TGCTGTTG | GTAAGCGC | XM_159771 | Mus musculus LOC241861 (LOC241861), mR. | | 2.14 | SM2520 | d | 12 |
| 092308m1_V2MM_140674 | 1.841327 | 3.004939 | TRUE | HP_36139 | TGCTGTTG | CACTAGAA | XM_156197 | Mus musculus LOC208209 (LOC208209), mRNA. | | 2.5 | SM2051 | d | 12 |
| 092308m1_V2MM_140674 | 1.841327 | 3.004939 | TRUE | HP_36139 | TGCTGTTG | CACTAGAA | XM_156197 | Mus musculus LOC208209 (LOC208209), mR. | | 2.5 | SM2051 | d | 12 |
| mPool2_V2MM_102079 | 1.841301 | 3.004939 | TRUE | HP_32323 | TGCTGTTG | GAGAATAT | NM_198606 | gene model 83, (NCBI) | Gm83 | 2.7 | SM2162 | e | 2 |
| mPool7_V2MM_147049 | 1.837501 | 3.004939 | TRUE | HP_36776 | TGCTGTTG | CTCTTTGT | XM_162569 | Mus musculus LOC243509 (LOC243509), mR. | | 2.4 | SM2046 | d | 5 |
| mPool7_V2MM_147049 | 1.837501 | 3.004939 | TRUE | HP_36776 | TGCTGTTG | CTCTTTGT | XM_162569 | Mus musculus LOC243509 (LOC243509), mRNA. | | 2.4 | SM2046 | d | 5 |
| mPool7_V2MM_92818 | 1.836666 | 3.004939 | TRUE | HP_31416 | TGCTGTTG | CAGACACT | AK079097 | RIKEN cDNA 9330186A19 gene | 9330186A19Rik | 2.9 | SM2266 | e | 9 |
| 092308m1_V2MM_160106 | 1.836561 | 3.004939 | TRUE | HP_38069 | TGCTGTTG | CCATAAAT | XM_283375 | Mus musculus hypothetical gene supported by AK053631 (LOC328710), mR. | | 2.6 | SM2119 | g | 6 |
| 092308m1_V2MM_157052 | 1.835591 | 3.004939 | TRUE | HP_37767 | TGCTGTTG | GATGTTTA | XM_204790 | Mus musculus similar to vomerosal 1 receptor, I9 [Mus musculus] (LOC277108), r | | 2.6 | SM2124 | g | 1 |
| mPool2_V2MM_92506 | 1.834843 | 3.004939 | TRUE | HP_31385 | TGCTGTTG | GTAATGAT | NM_178723 | zinc finger protein 533 | Zfp533 | 2.6 | SM2139 | a | 5 |
| 092308m1_V2MM_140837 | 1.834628 | 3.004939 | TRUE | HP_36155 | TGCTGTTG | CTTTCTTT | XM_156295 | Mus musculus similar to pol protein [Phascolarctos cinereus] (LOC224231), mRNA | | 2.1 | SM2331 | f | 6 |
| 092308m1_V2MM_140837 | 1.834628 | 3.004939 | TRUE | HP_36155 | TGCTGTTG | CTTTCTTT | XM_156295 | Mus musculus similar to pol protein [Phascolarctos cinereus] (LOC224231), mR. | | 2.1 | SM2331 | f | 6 |
| 092308m1_V2MM_67924 | 1.834546 | 3.004939 | TRUE | HP_28989 | TGCTGTTG | CTCTGTGA | NM_172998 | RIKEN cD B830028P19 gene | B830028P19Rik | 2.3 | SM2011 | a | 4 |
| mPool7_V2MM_170459 | 1.833132 | 3.004939 | TRUE | HP_39095 | TGCTGTTG | GGCTTTAA | XM_283177 | Mus musculus similar to hypothetical protein 3 - rat (LOC328328), mRNA. | | 2.5 | SM2077 | e | 6 |
| mPool7_V2MM_170459 | 1.833132 | 3.004939 | TRUE | HP_39095 | TGCTGTTG | GGCTTTAA | XM_283177 | Mus musculus similar to hypothetical protein 3 - rat (LOC328328), mR. | | 2.5 | SM2077 | e | 6 |
| mPool2_V2MM_30536 | 1.830474 | 2.205692 | TRUE | HP_25346 | TGCTGTTG | CGGAATAT | NM_011941 | mitogen activated protein kinase binding protein 1 | Mapkbp1 | 2.11 | SM2392 | d | 6 |
| mPool7_V2MM_90885 | 1.829432 | 3.004939 | TRUE | HP_31230 | TGCTGTTG | CTTTGAGC | NM_177880 | hypothetical protein 4933411G11 | | 2.9 | SM2254 | g | 12 |
| mPool7_V2MM_90842 | 1.828726 | 3.004939 | TRUE | HP_31225 | TGCTGTTG | GGTTTACT | NM_177869 | expressed sequence AI847670 | AI847670 | 2.8 | SM2250 | a | 4 |
| mPool7_V2MM_90842 | 1.828726 | 3.004939 | TRUE | HP_31225 | TGCTGTTG | GGTTTACT | NM_177869 | expressed sequence AI847670 | AI847670 | 2.8 | SM2250 | a | 4 |
| 092308m1_V2MM_135185 | 1.825928 | 3.004939 | TRUE | HP_35590 | TGCTGTTG | GTCTAAAT | XM_150392 | protein phosphatase 1A, magnesium dependent, alpha is | Ppm1a | 2.4 | SM2045 | e | 12 |
| 092308m1_V2MM_135185 | 1.825928 | 3.004939 | TRUE | HP_35590 | TGCTGTTG | GTCTAAAT | XM_150392 | protein phosphatase 1A, magnesium dependent, alpha is | Ppm1a | 2.4 | SM2045 | e | 12 |
| mPool2_V2MM_1304 | 1.825761 | 2.205692 | TRUE | HP_22495 | TGCTGTTG | CCAATTCT | NM_008125 | gap junction membrane channel protein beta 2 | Gjb2 | 2.11 | SM2379 | h | 6 |
| mPool7_V2MM_195719 | 1.82571 | 2.205692 | TRUE | HP_41384 | TGCTGTTG | GATCATCT | NM_178376 | Ras-related GTP binding A | Rraga | 2.9 | SM2259 | e | 4 |
| mPool7_V2MM_63424 | 1.823698 | 2.205692 | TRUE | HP_28550 | TGCTGTTG | CGGATATC | NM_177383 | G protein-coupled receptor 21 | Gpr21 | 2.8 | SM2243 | c | 12 |
| mPool7_V2MM_103070 | 1.821865 | 2.205692 | TRUE | HP_32421 | TGCTGTTG | CTATTTCT | NM_199466 | echinoderm microtubule associated protein like 4 | Eml4 | 2.1 | SM2321 | c | 11 |
| mPool2_V2MM_127282 | 1.821688 | 2.205692 | TRUE | HP_34811 | TGCTGTTG | GAAAGGA | XM_145108 | Mus musculus LOC243714 (LOC243714), mRNA. | | 2.11 | SM2369 | g | 5 |

| | | | | | | | | | | | | | | | |
|---|---|---|---|---|---|---|---|---|---|---|---|---|---|---|---|
| 092308m1_V2MM_137091 | 1.821151 | 2.205692 | TRUE | HP_35781 | TGCTGTTG | GGCTCTAT | XM_152567 | Mus musculus LOC237736 (LOC237736), mR. | | | 2.5 | SM2052 | c | | 3 |
| mPool7_V2MM_103172 | 1.820546 | 3.004939 | TRUE | HP_32431 | TGCTGTTG | GAGACAT | XM_128858 | RIKEN cDNA 4833403I15 gene | 4833403I15Rik | | 2.1 | SM2315 | h | | 4 |
| 092308m1_V2MM_178263 | 1.820437 | 2.205692 | TRUE | HP_39873 | TGCTGTTG | CATCCTAA | XM_288120 | Mus musculus hypothetical gene supported by AK054126 (LOC331473), mR. | | | 2.5 | SM2079 | h | | 1 |
| mPool7_V2MM_89907 | 1.820017 | 3.004939 | TRUE | HP_31133 | TGCTGTTG | CATCTTCT | BC025913 | biogenesis of lysosome-related organelles complex-1, sul | Bloc1s3 | | 2.9 | SM2266 | f | | 11 |
| mPool7_V2MM_89907 | 1.820017 | 3.004939 | TRUE | HP_31133 | TGCTGTTG | CATCTTCT | BC025913 | biogenesis of lysosome-related organelles complex-1, sul | Bloc1s3 | | 2.9 | SM2266 | f | | 11 |
| mPool7_V2MM_175564 | 1.818907 | 3.004939 | TRUE | HP_39604 | TGCTGTTG | GCCATTG | XM_489100 | hypothetical gene supported by AK031827 | | | 2.5 | SM2078 | d | | 9 |
| mPool7_V2MM_175564 | 1.818907 | 3.004939 | TRUE | HP_39604 | TGCTGTTG | GCCATTG | XM_489100 | hypothetical gene supported by AK031827 | | | 2.5 | SM2078 | d | | 9 |
| 092308m1_V2MM_175874 | 1.817468 | 2.205692 | TRUE | HP_39635 | TGCTGTTG | CAAATCTT | XM_287545 | similar to PABP1-dependent poly A-specific ribonuclease subunit PAN3; PABP-de | | | 2.5 | SM2066 | a | | 6 |
| 092308m3_V2MM_139822 | 1.817454 | 61.29828 | TRUE | HP_36054 | TGCTGTTG | GGGCCAG | XM_155408 | Mus musculus LOC239495 (LOC239495), mR. | | | 2.14 | SM2514 | d | | 12 |
| mPool6_V2MM_77615 | 1.81742 | 75.83547 | TRUE | HP_29933 | TGCTGTTG | CTGTGTTT | NM_025784 | BCS1-like (yeast) | Bcs1l | | 2.7 | SM2175 | b | | 7 |
| 092308m1_V2MM_178019 | 1.814586 | 2.205692 | TRUE | HP_39849 | TGCTGTTG | CAGGGAA | XM_288010 | Mus musculus hypothetical gene supported by AK076622 (LOC331162), mR. | | | 2.5 | SM2094 | h | | 1 |
| 092308m1_V2MM_178353 | 1.813933 | 3.004939 | TRUE | HP_39882 | TGCTGTTG | CCAAGTA | XM_288143 | Mus musculus hypothetical gene supported by AK046508 (LOC331558), mRNA. | | | 2.5 | SM2069 | g | | 10 |
| 092308m1_V2MM_178353 | 1.813933 | 3.004939 | TRUE | HP_39882 | TGCTGTTG | CCAAGTA | XM_288143 | Mus musculus hypothetical gene supported by AK046508 (LOC331558), mR. | | | 2.5 | SM2069 | g | | 10 |
| mPool7_V2MM_150314 | 1.813855 | 3.004939 | TRUE | HP_37100 | TGCTGTTG | GTTTCCTA | NM_177171 | RIKEN cDNA D930036F22 gene | D930036F22Rik | | 2.1 | SM2328 | g | | 8 |
| mPool7_V2MM_92636 | 1.813438 | 3.004939 | TRUE | HP_31398 | TGCTGTTG | GTAGGAG | NM_178745 | RIKEN cDNA 6330442E10 gene | 6330442E10Rik | | 2.9 | SM2262 | c | | 6 |
| mPool2_V2MM_160247 | 1.813302 | 2.205692 | TRUE | HP_38083 | TGCTGTTG | CCAGGAA | XM_283435 | Mus musculus similar to RIKEN cDNA 2010003O02 [Mus musculus] (LOC328796), | | | 2.6 | SM2126 | d | | 8 |
| mPool7_V2MM_74933 | 1.812848 | 2.205692 | TRUE | HP_29672 | TGCTGTTG | CAGTTTAT | NM_028836 | chitobiase, di-N-acetyl- | Ctbs | | 2.8 | SM2236 | g | | 4 |
| mPool2_V2MM_145856 | 1.812021 | 3.004939 | TRUE | HP_36656 | TGCTGTTG | GTCAGAA | XM_161546 | Mus musculus LOC242761 (LOC242761), mRNA. | | | 2.1 | SM2345 | e | | 4 |
| mPool2_V2MM_140268 | 1.811281 | 3.004939 | TRUE | HP_36098 | TGCTGTTG | GCTATGT | XM_155816 | Mus musculus LOC239490 (LOC239490), mR. | | | 2.4 | SM2042 | g | | 7 |
| mPool2_V2MM_140268 | 1.811281 | 3.004939 | TRUE | HP_36098 | TGCTGTTG | GCTATGT | XM_155816 | Mus musculus LOC239490 (LOC239490), mRNA. | | | 2.4 | SM2042 | g | | 7 |
| 092308m3_V2MM_208411 | 1.811041 | 66.95036 | TRUE | HP_42620 | TGCTGTTG | GCACTTG | XM_159344 | Mus musculus LOC245565 (LOC245565), mR. | | | 2.14 | SM2520 | b | | 9 |
| mPool7_V2MM_84662 | 1.808608 | 2.205692 | TRUE | HP_30616 | TGCTGTTG | CGTTGATT | NM_028848 | RIKEN cDNA 4930513F16 gene | 4930513F16Rik | | 2.9 | SM2259 | b | | 8 |
| mPool7_V2MM_77897 | 1.808162 | 3.004939 | TRUE | HP_10504 | TGCTGTTG | GAAACCA | AB086123 | calcium channel, voltage-dependent, L type, alpha 1D sul | Cacna1d | | 2.8 | SM2236 | a | | 11 |
| 092308m1_V2MM_180607 | 1.807655 | 3.004939 | TRUE | HP_40108 | TGCTGTTG | CTATGAAT | XM_358439 | gene model 1573, (NCBI) | Gm1573 | | 2.5 | SM2065 | h | | 7 |
| 092308m1_V2MM_180607 | 1.807655 | 3.004939 | TRUE | HP_40108 | TGCTGTTG | CTATGAAT | XM_358439 | gene model 1573, (NCBI) | Gm1573 | | 2.5 | SM2065 | h | | 7 |
| 092308m1_V2MM_154308 | 1.806754 | 3.004939 | TRUE | HP_37494 | TGCTGTTG | CCCTTGGT | AK045988 | RIKEN cD 6530403G13 gene | 6530403G13Rik | | 2.6 | SM2115 | d | | 9 |
| mPool2_V2MM_143833 | 1.804551 | 2.205692 | TRUE | HP_36454 | TGCTGTTG | CATGAAC | XM_159183 | Mus musculus LOC245402 (LOC245402), mR. | | | 2.4 | SM2038 | f | | 10 |
| mPool2_V2MM_143833 | 1.804551 | 2.205692 | TRUE | HP_36454 | TGCTGTTG | CATGAAC | XM_159183 | Mus musculus LOC245402 (LOC245402), mRNA. | | | 2.4 | SM2038 | f | | 10 |
| 092308m1_V2MM_171110 | 1.801508 | 2.205692 | TRUE | HP_39159 | TGCTGTTG | CGTATTTA | XM_286558 | Mus musculus hypothetical gene supported by AK044419 (LOC328319), mR. | | | 2.5 | SM2072 | b | | 7 |
| 092308m1_V2MM_69402 | 1.801135 | 3.004939 | TRUE | HP_36745 | TGCTGTTG | GCCTGAC | BC082537 | RIKEN cD 2900042E01 gene | 2900042E01Rik | | 2.3 | SM2003 | f | | 3 |
| mPool6_V2MM_158860 | 1.800228 | 75.83547 | TRUE | HP_37946 | TGCTGTTG | GCTGCTG | XM_282894 | Mus musculus hypothetical gene supported by AK042566 (LOC327726), mRNA. | | | 2.8 | SM2208 | e | | 6 |
| mPool4_V2MM_15470 | 1.800172 | 75.83547 | TRUE | HP_23879 | TGCTGTTG | CTCTGTGT | NM_011407 | schlafen 1 | Slfn1 | | 2.15 | SM2573 | b | | 2 |
| mPool7_V2MM_84868 | 1.799685 | 2.205692 | TRUE | HP_30636 | TGCTGTTG | CTTCATCC | NM_029705 | Machado-Joseph disease (spinocerebellar ataxia 3, olivo | Mjd | | 2.9 | SM2261 | c | | 5 |
| mPool7_V2MM_196416 | 1.799305 | 3.004939 | TRUE | HP_41452 | TGCTGTTG | CTGTAAG | NM_175479 | RIKEN cDNA A330008L17 gene | A330008L17Rik | | 2.9 | SM2254 | a | | 4 |
| mPool2_V2MM_212683 | 1.798955 | 2.205692 | TRUE | HP_43034 | TGCTGTTG | CACATCAT | XM_142433 | similar to hypothetical protein | | | 2.11 | SM2370 | a | | 6 |
| mPool2_V2MM_110315 | 1.796038 | 3.004939 | TRUE | HP_33135 | TGCTGTTG | GACCCTA | XM_134580 | gene model 177, (NCBI) | Gm177 | | 2.6 | SM2149 | h | | 5 |
| 092308m1_V2MM_151863 | 1.795947 | 2.205692 | TRUE | HP_37253 | TGCTGTTG | CCCATGTT | XM_194974 | similar to ribosomal protein | | | 2.5 | SM2052 | h | | 3 |
| mPool2_V2MM_85938 | 1.7941 | 3.004939 | TRUE | HP_30741 | TGCTGTTG | CAGCTATT | NM_146524 | olfactory receptor 855 | Olfr855 | | 2.6 | SM2146 | c | | 10 |
| 092308m1_V2MM_146584 | 1.793485 | 3.004939 | TRUE | HP_36729 | TGCTGTTG | CATAATTG | XM_162162 | Mus musculus LOC243184 (LOC243184), mR. | | | 2.5 | SM2058 | a | | 9 |
| 092308m1_V2MM_146584 | 1.793485 | 3.004939 | TRUE | HP_36729 | TGCTGTTG | CATAATTG | XM_162162 | Mus musculus LOC243184 (LOC243184), mRNA. | | | 2.5 | SM2058 | a | | 9 |
| mPool7_V2MM_177261 | 1.792334 | 3.004939 | TRUE | NA | NA | NA | NA | NA | NA | NA | | NA | NA | NA | |
| 092308m1_V2MM_139509 | 1.790566 | 3.004939 | TRUE | HP_36022 | TGCTGTTG | CTACTATC | XM_155129 | Mus musculus LOC239255 (LOC239255), mRNA. | | | 2.4 | SM2040 | h | | 5 |
| 092308m1_V2MM_139509 | 1.790566 | 3.004939 | TRUE | HP_36022 | TGCTGTTG | CTACTATC | XM_155129 | Mus musculus LOC239255 (LOC239255), mR. | | | 2.4 | SM2040 | h | | 5 |
| 092308m1_V2MM_187096 | 1.788755 | 2.205692 | TRUE | HP_40756 | TGCTGTTG | CCCTTAAG | XM_289648 | Mus musculus LOC333392 (LOC333392), mR. | | | 2.5 | SM2072 | e | | 8 |
| mPool7_V2MM_68109 | 1.787386 | 1.959154 | TRUE | HP_10776 | TGCTGTTG | CTGGAAA | NM_026360 | DEAD (Asp-Glu-Ala-Asp) box polypeptide 47 | Ddx47 | | 2.8 | SM2243 | d | | 1 |
| mPool7_V2MM_173441 | 1.787223 | 3.004939 | TRUE | HP_39392 | TGCTGTTG | CAAACATA | AK035520 | hypothetical protein 9530060I07 | | | 2.5 | SM2081 | c | | 2 |
| mPool7_V2MM_173441 | 1.787223 | 3.004939 | TRUE | HP_39392 | TGCTGTTG | CAAACATA | AK035520 | hypothetical protein 9530060I07 | | | 2.5 | SM2081 | c | | 2 |
| 092308m1_V2MM_158543 | 1.785906 | 3.004939 | TRUE | HP_37915 | TGCTGTTG | CTCAAGCA | XM_206745 | Mus musculus LOC279156 (LOC279156), mR. | | | 2.6 | SM2115 | e | | 10 |
| mPool2_V2MM_133812 | 1.78557 | 3.004939 | TRUE | HP_35454 | TGCTGTTG | CTGCTTAT | NM_177669 | RIKEN cDNA A630098G03 gene | A630098G03Rik | | 2.11 | SM2364 | a | | 10 |
| mPool2_V2MM_214867 | 1.785552 | 3.004939 | TRUE | HP_43243 | TGCTGTTG | CATGATTA | XM_485031 | olfactory receptor 48 | Olfr48 | | 2.11 | SM2351 | f | | 12 |
| 092308m1_V2MM_158554 | 1.784753 | 3.004939 | TRUE | HP_37916 | TGCTGTTG | GCCCAAAT | XM_206756 | similar to F1N19.3 | | | 2.6 | SM2121 | f | | 9 |
| 092308m1_V2MM_74731 | 1.783928 | 3.004939 | TRUE | HP_29652 | TGCTGTTG | CAGGAAC | NM_173431 | RIKEN cD 1700047E16 gene | 1700047E16Rik | | 2.3 | SM2009 | e | | 9 |
| mPool2_V2MM_128028 | 1.783348 | 3.004939 | TRUE | HP_34884 | TGCTGTTG | GCCTTTAG | XM_145500 | RIKEN cDNA 1110003H10 gene | 1110003H10Rik | | 2.11 | SM2351 | h | | 6 |

| | | | | | | | | | | | | | |
|---|---|---|---|---|---|---|---|---|---|---|---|---|---|
| mPool2_V2MM_101379 | 1.782748 | 3.004939 | TRUE | NA | NA | NA | NA | NA | NA | NA | NA | NA | NA |
| 092308m1_V2MM_68358 | 1.782434 | 3.004939 | TRUE | HP_29032 | TGCTGTTG | CTTTCTTG | NM_146851 | olfactory receptor 295 | Olfr295 | 2.3 | SM2009 | f | 4 |
| mPool7_V2MM_202589 | 1.782339 | 3.004939 | TRUE | HP_42058 | TGCTGTTG | CCTGGAG | XM_127675 | Mus musculus similar to KIAA1607 protein [Homo sapiens] (LOC218897), mRNA. | | 2.1 | SM2306 | b | 1 |
| mPool2_V2MM_20404 | 1.781467 | 2.205692 | TRUE | HP_22249 | TGCTGTTG | GTTATATA | NM_010756 | v-maf musculoaponeurotic fibrosarcoma oncogene famil | Mafg | 2.11 | SM2386 | f | 10 |
| mPool2_V2MM_134442 | 1.781304 | 3.004939 | TRUE | HP_35516 | TGCTGTTG | CTGTTCTT | AK046418 | RIKEN cD 4930415J05 gene | 4930415J05Rik | 2.6 | SM2104 | c | 10 |
| mPool2_V2MM_134442 | 1.781304 | 3.004939 | TRUE | HP_35516 | TGCTGTTG | CTGTTCTT | AK046418 | RIKEN cDNA 4930415J05 gene | 4930415J05Rik | 2.6 | SM2104 | c | 10 |
| 092308m1_V2MM_183610 | 1.781 | 2.205692 | TRUE | HP_40408 | TGCTGTTG | CCGCTTAA | XM_289049 | Mus musculus LOC333489 (LOC333489), mR. | | 2.5 | SM2081 | b | 8 |
| 092308m3_V2MM_206507 | 1.780459 | 75.83547 | TRUE | HP_42432 | TGCTGTTG | CACACTTT | XM_128374 | zinc finger protein 294 | Zfp294 | 2.1 | SM2318 | g | 12 |
| 092308m3_V2MM_206507 | 1.780459 | 75.83547 | TRUE | HP_42432 | TGCTGTTG | CACACTTT | XM_128374 | zinc finger protein 294 | Zfp294 | 2.1 | SM2318 | g | 12 |
| 092308m1_V2MM_142730 | 1.779285 | 2.205692 | TRUE | HP_36344 | TGCTGTTG | GTGAATAT | XM_157867 | Mus musculus LOC240547 (LOC240547), mR. | | 2.5 | SM2059 | g | 4 |
| 092308m1_V2MM_142730 | 1.779285 | 2.205692 | TRUE | HP_36344 | TGCTGTTG | GTGAATAT | XM_157867 | Mus musculus LOC240547 (LOC240547), mRNA. | | 2.5 | SM2059 | g | 4 |
| mPool7_V2MM_171353 | 1.779272 | 1.959154 | TRUE | HP_39184 | TGCTGTTG | GATTTATT | XM_488707 | hypothetical gene supported by AK048353 | | 2.9 | SM2274 | b | 3 |
| 092308m1_V2MM_125180 | 1.777149 | 3.004939 | TRUE | HP_34602 | TGCTGTTG | CAGCAAG | BC059253 | CDK5 regulatory subunit associated protein 2 | Cdk5rap2 | 2.6 | SM2101 | a | 7 |
| 092308m1_V2MM_170810 | 1.775773 | 3.004939 | TRUE | HP_39130 | TGCTGTTG | CCCAGAA | AK045681 | hypothetical protein B230303A05 | | 2.5 | SM2080 | a | 8 |
| mPool2_V2MM_125198 | 1.775367 | 3.004939 | TRUE | HP_34604 | TGCTGTTG | CATGTTGA | XM_143850 | Mus musculus similar to pre-mRNA splicing factor Prp16; pre-mRNA splicing facto | | 2.11 | SM2366 | c | 8 |
| 092308m3_V2MM_59498 | 1.774868 | 62.64426 | TRUE | HP_28167 | TGCTGTTG | GCCTTTGC | NM_029295 | chemokine-like factor | Cklf | 2.12 | SM2407 | c | 9 |
| mPool7_V2MM_85962 | 1.774285 | 3.004939 | TRUE | HP_30744 | TGCTGTTG | CATGGAT | NM_146530 | olfactory receptor 1186 | Olfr1186 | 2.9 | SM2268 | e | 12 |
| mPool7_V2MM_179499 | 1.773159 | 3.004939 | TRUE | HP_39997 | TGCTGTTG | CAGAGTTT | XM_288344 | Mus musculus LOC327806 (LOC327806), mRNA. | | 2.5 | SM2062 | e | 2 |
| mPool7_V2MM_179499 | 1.773159 | 3.004939 | TRUE | HP_39997 | TGCTGTTG | CAGAGTTT | XM_288344 | Mus musculus LOC327806 (LOC327806), mR. | | 2.5 | SM2062 | e | 2 |
| mPool2_V2MM_212338 | 1.772796 | 2.205692 | TRUE | HP_43000 | TGCTGTTG | GACATAAT | AK048574 | RIKEN cDNA C130078N17 gene | C130078N17Rik | 2.1 | SM2350 | a | 3 |
| mPool7_V2MM_67315 | 1.772663 | 3.004939 | TRUE | HP_28930 | TGCTGTTG | CCTCCGCT | NM_021532 | dapper homolog 1, antagonist of beta-catenin (xenopus) | Dact1 | 2.8 | SM2230 | h | 1 |
| 092308m1_V2MM_62745 | 1.771299 | 3.004939 | TRUE | HP_28484 | TGCTGTTG | CATCCAAT | Z11981 | plasmacytoma variant translocation 1 | Pvt1 | 2.3 | SM2013 | h | 7 |
| mPool2_V2MM_133636 | 1.771069 | 3.004939 | TRUE | HP_35437 | TGCTGTTG | CTGGAGTT | BC050760 | synaptotagmin 6 | Syt6 | 2.11 | SM2374 | f | 3 |
| mPool7_V2MM_75235 | 1.770862 | 3.004939 | TRUE | HP_29701 | TGCTGTTG | CCACCAAT | NM_026391 | protein phosphatase 2, regulatory subunit B, delta isofor | Ppp2r2d | 2.8 | SM2240 | b | 4 |
| mPool7_V2MM_140837 | 1.77001 | 2.205692 | TRUE | HP_36155 | TGCTGTTG | CTTTCTTT | XM_156295 | Mus musculus similar to pol protein [Phascolarctos cinereus] (LOC224231), mRN | | 2.1 | SM2331 | f | 6 |
| mPool7_V2MM_140837 | 1.77001 | 2.205692 | TRUE | HP_36155 | TGCTGTTG | CTTTCTTT | XM_156295 | Mus musculus similar to pol protein [Phascolarctos cinereus] (LOC224231), mR. | | 2.1 | SM2331 | f | 6 |
| 092308m1_V2MM_128385 | 1.767592 | 3.004939 | TRUE | HP_34919 | TGCTGTTG | CTCATGTT | XM_145667 | transient receptor potential cation channel, subfamily M | Trpm1 | 2.5 | SM2100 | g | 12 |
| mPool7_V2MM_195901 | 1.765877 | 3.004939 | TRUE | HP_41402 | TGCTGTTG | CATGGTT | NM_178665 | LIM domain containing preferred translocation partner ir | Lpp | 2.8 | SM2250 | b | 1 |
| mPool2_V2MM_132692 | 1.764918 | 3.004939 | TRUE | HP_35344 | TGCTGTTG | GAGACATT | AK044430 | mutated in colorectal cancers | Mcc | 2.1 | SM2350 | a | 8 |
| mPool7_V2MM_151228 | 1.762266 | 2.205692 | TRUE | HP_37189 | TGCTGTTG | GTGAAAT | NM_001002272 | trophinin | Tro | 2.4 | SM2050 | c | 10 |
| mPool7_V2MM_151228 | 1.762266 | 2.205692 | TRUE | HP_37189 | TGCTGTTG | GTGAAAT | NM_001002272 | trophinin | Tro | 2.4 | SM2050 | c | 10 |
| mPool7_V2MM_179069 | 1.759908 | 3.004939 | TRUE | HP_39954 | TGCTGTTG | CTATTAAA | XM_288269 | Mus musculus LOC332542 (LOC332542), mR. | | 2.5 | SM2091 | c | 7 |
| mPool7_V2MM_179069 | 1.759908 | 3.004939 | TRUE | HP_39954 | TGCTGTTG | CTATTAAA | XM_288269 | Mus musculus LOC332542 (LOC332542), mRNA. | | 2.5 | SM2091 | c | 7 |
| mPool7_V2MM_196009 | 1.758391 | 3.004939 | TRUE | HP_41412 | TGCTGTTG | CTTGTATA | NM_173757 | mitochondrial ribosomal protein S27 | Mrps27 | 2.9 | SM2266 | e | 5 |
| mPool7_V2MM_86066 | 1.757119 | 3.004939 | TRUE | HP_30754 | TGCTGTTG | GAGTTTAC | NM_146613 | olfactory receptor 973 | Olfr973 | 2.9 | SM2270 | b | 6 |
| mPool7_V2MM_170871 | 1.757114 | 3.004939 | TRUE | HP_39136 | TGCTGTTG | CGTTAAAT | XM_286509 | Mus musculus hypothetical gene supported by AK037084 (LOC328225), mR. | | 2.5 | SM2072 | g | 5 |
| mPool7_V2MM_170871 | 1.757114 | 3.004939 | TRUE | HP_39136 | TGCTGTTG | CGTTAAAT | XM_286509 | Mus musculus hypothetical gene supported by AK037084 (LOC328225), mRNA. | | 2.5 | SM2072 | g | 5 |
| mPool2_V2MM_192335 | 1.756741 | 1.959154 | TRUE | HP_33789 | TGCTGTTG | GTAACATA | XM_354831 | olfactomedin 4 | Olfm4 | 2.1 | SM2308 | h | 6 |
| mPool2_V2MM_126250 | 1.756123 | 2.205692 | TRUE | HP_34709 | TGCTGTTG | CGTACAAC | XM_144444 | Mus musculus similar to 60S ACIDIC RIBOSOMAL PROTEIN P1 (LOC243162), mRN | | 2.11 | SM2355 | e | 7 |
| mPool2_V2MM_210602 | 1.755649 | 3.004939 | TRUE | HP_42834 | TGCTGTTG | CAACCACA | XM_143694 | Mus musculus similar to glyceraldehyde-3-phosphate dehydrogenase [Mus musc | | 2.11 | SM2362 | e | 12 |
| mPool2_V2MM_211682 | 1.755483 | 3.004939 | TRUE | HP_42937 | TGCTGTTG | GGCAATA | XM_143642 | Mus musculus similar to eukaryotic translation initiation factor 4A1; initiation fac | | 2.11 | SM2352 | f | 11 |
| mPool7_V2MM_172300 | 1.754816 | 2.205692 | TRUE | HP_39278 | TGCTGTTG | GTTTGTAA | XM_286796 | Mus musculus hypothetical gene supported by AK042793 (LOC328772), mR. | | 2.5 | SM2079 | c | 5 |
| mPool7_V2MM_172300 | 1.754816 | 2.205692 | TRUE | HP_39278 | TGCTGTTG | GTTTGTAA | XM_286796 | Mus musculus hypothetical gene supported by AK042793 (LOC328772), mRNA. | | 2.5 | SM2079 | c | 5 |
| mPool2_V2MM_4516 | 1.754703 | 3.004939 | TRUE | HP_22808 | TGCTGTTG | GTTATAAA | NM_027374 | peptidylprolyl isomerase (cyclophilin)-like 3 | Ppil3 | 2.11 | SM2375 | b | 1 |
| mPool7_V2MM_195279 | 1.754677 | 3.004939 | TRUE | HP_41342 | TGCTGTTG | CCATATGT | NM_146436 | olfactory receptor 998 | Olfr998 | 2.9 | SM2257 | d | 12 |
| mPool7_V2MM_195279 | 1.754677 | 3.004939 | TRUE | HP_41342 | TGCTGTTG | CCATATGT | NM_146436 | olfactory receptor 998 | Olfr998 | 2.9 | SM2257 | d | 12 |
| mPool2_V2MM_151870 | 1.752762 | 3.004939 | TRUE | HP_37253 | TGCTGTTG | CTCTGTGT | XM_194977 | similar to Cytochrome P450, family 2, subfamily d, polypeptide 13 | | 2.1 | SM2345 | g | 5 |
| 092308m1_V2MM_176983 | 1.750992 | 2.205692 | TRUE | HP_39746 | TGCTGTTG | CTACTTAC | AK081575 | expressed sequence W91709 | W91709 | 2.5 | SM2069 | b | 3 |
| mPool2_V2MM_6424 | 1.750325 | 2.205692 | TRUE | HP_22996 | TGCTGTTG | CTGCTAAA | NM_177992 | guanosine monophosphate reductase 2 | Gmpr2 | 2.11 | SM2377 | f | 4 |
| 092308m1_V2MM_170621 | 1.749821 | 2.205692 | TRUE | HP_39111 | TGCTGTTG | CTGTCAGT | XM_286455 | Mus musculus hypothetical gene supported by AK039743 (LOC328163), mR. | | 2.5 | SM2060 | f | 3 |
| mPool2_V2MM_210737 | 1.7487 | 2.205692 | TRUE | HP_42847 | TGCTGTTG | CTAAGTAT | XM_144834 | gene model 1409, (NCBI) | Gm1409 | 2.11 | SM2371 | e | 3 |
| mPool2_V2MM_128586 | 1.748574 | 3.004939 | TRUE | HP_34939 | TGCTGTTG | CTACTGAC | XM_145773 | Mus musculus similar to olfactory receptor MOR10-1 [Mus musculus] (LOC23359 | | 2.11 | SM2364 | g | 2 |
| mPool2_V2MM_170258 | 1.748132 | 2.205692 | TRUE | HP_39075 | TGCTGTTG | CACACACT | AK083360 | hypothetical protein C920021A13 | | 2.6 | SM2132 | b | 2 |

| | | | | | | | | | | | | | |
|---|---|---|---|---|---|---|---|---|---|---|---|---|---|
| mPool7_V2MM_116096 | 1.747098 | 2.205692 | TRUE | HP_33705 | TGCTGTTG | CTGATATT | XM_138748 | Mus musculus similar to envelope protein [Ovis aries] (LOC218362), mRNA. | | 2.6 | SM2150 | c | 2 |
| mPool2_V2MM_10590 | 1.746522 | 3.004939 | TRUE | HP_23401 | TGCTGTTG | CTGAAACT | NM_007687 | cofilin 1, non-muscle | Cfl1 | 2.11 | SM2386 | b | 3 |
| 092308m1_V2MM_167494 | 1.746275 | 3.004939 | TRUE | HP_38800 | TGCTGTTG | CACAGAG | XM_285777 | Mus musculus similar to Involved in chitin synthase III activity, also required for h | | 2.6 | SM2117 | g | 10 |
| mPool7_V2MM_103760 | 1.746163 | 1.959154 | TRUE | HP_32489 | TGCTGTTG | GCTTATTT | AK008974 | RIKEN cD 2210418G03 gene | 2210418G03Rik | 2.9 | SM2297 | a | 4 |
| mPool7_V2MM_103760 | 1.746163 | 1.959154 | TRUE | HP_32489 | TGCTGTTG | GCTTATTT | AK008974 | RIKEN cDNA 2210418G03 gene | 2210418G03Rik | 2.9 | SM2297 | a | 4 |
| mPool7_V2MM_175995 | 1.745676 | 3.004939 | TRUE | HP_39647 | TGCTGTTG | CACACCCT | AK033923 | hypothetical protein 9330117B14 | | 2.9 | SM2274 | f | 4 |
| mPool2_V2MM_121810 | 1.745608 | 2.205692 | TRUE | HP_34268 | TGCTGTTG | CTGTCTTA | XM_141857 | Mus musculus similar to RIKEN cDNA 1700042B14 [Mus musculus] (LOC236872), | | 2.11 | SM2364 | h | 7 |
| 092308m1_V2MM_161736 | 1.742692 | 3.004939 | TRUE | HP_38230 | TGCTGTTG | CTCTCGGA | XM_284026 | Mus musculus hypothetical gene supported by AK030056 (LOC329969), mR. | | 2.6 | SM2115 | d | 10 |
| mPool2_V2MM_32277 | 1.741137 | 3.004939 | TRUE | HP_25516 | TGCTGTTG | GGAAATTA | NM_023587 | protein tyrosine phosphatase-like (proline instead of cata | Ptplb | 2.11 | SM2392 | b | 11 |
| mPool7_V2MM_89806 | 1.741072 | 3.004939 | TRUE | HP_31123 | TGCTGTTG | GTGAGCG | NM_177674 | hypothetical protein A530045M11 | | 2.9 | SM2269 | b | 6 |
| 092308m1_V2MM_175197 | 1.740383 | 2.205692 | TRUE | HP_39568 | TGCTGTTG | CTGTTTCT | XM_287390 | Mus musculus hypothetical gene supported by AK089399 (LOC329956), mR. | | 2.5 | SM2095 | c | 11 |
| mPool7_V2MM_199289 | 1.738678 | 3.004939 | TRUE | HP_41733 | TGCTGTTG | CATTGTTT | XM_286732 | expressed sequence AU022320 | AU022320 | 2.9 | SM2288 | h | 11 |
| mPool2_V2MM_130962 | 1.738175 | 3.004939 | TRUE | HP_35173 | TGCTGTTG | GTCTGTAC | XM_147124 | RIKEN cDNA 5830454E08 gene | 5830454E08Rik | 2.11 | SM2360 | d | 8 |
| mPool7_V2MM_85524 | 1.738001 | 0 | TRUE | HP_30700 | TGCTGTTG | GTCCTACT | NM_146402 | olfactory receptor 1303 | Olfr1303 | 2.9 | SM2254 | h | 9 |
| 092308m3_V2MM_196361 | 1.737549 | 75.83547 | TRUE | HP_41446 | TGCTGTTG | CCTGGAA | NM_010165 | eyes absent 2 homolog (Drosophila) | Eya2 | 2.12 | SM2450 | h | 9 |
| mPool7_V2MM_90042 | 1.737532 | 2.205692 | TRUE | HP_31146 | TGCTGTTG | GTGAGTG | NM_177721 | RAN binding protein 6 | Ranbp6 | 2.9 | SM2254 | a | 3 |
| mPool7_V2MM_90042 | 1.737532 | 2.205692 | TRUE | HP_31146 | TGCTGTTG | GTGAGTG | NM_177721 | RAN binding protein 6 | Ranbp6 | 2.9 | SM2254 | a | 3 |
| 092308m1_V2MM_144171 | 1.736964 | 3.004939 | TRUE | HP_36488 | TGCTGTTG | CCATTTGT | XM_159785 | Mus musculus LOC207783 (LOC207783), mR. | | 2.5 | SM2056 | a | 3 |
| mPool2_V2MM_4227 | 1.735435 | 2.205692 | TRUE | HP_22780 | TGCTGTTG | GGGTTTAT | NM_007568 | betacellulin, epidermal growth factor family member | Btc | 2.11 | SM2385 | b | 2 |
| 092308m1_V2MM_161793 | 1.734441 | 3.004939 | TRUE | HP_38235 | TGCTGTTG | CAGAGTA | XM_284053 | six transmembrane epithelial antigen of prostate 2 | Steap2 | 2.6 | SM2118 | b | 7 |
| mPool2_V2MM_2066 | 1.733813 | 3.004939 | TRUE | HP_22569 | TGCTGTTG | CTACTACT | NM_008288 | hydroxysteroid 11-beta dehydrogenase 1 | Hsd11b1 | 2.11 | SM2385 | g | 3 |
| 092308m1_V2MM_127129 | 1.733755 | 3.004939 | TRUE | HP_34796 | TGCTGTTG | CCACCTTA | XM_145005 | IQ motif and Sec7 domain 3 | Iqsec3 | 2.6 | SM2104 | a | 9 |
| mPool2_V2MM_107909 | 1.732894 | 2.205692 | TRUE | HP_32898 | TGCTGTTG | CTGATCAT | AK014627 | RIKEN cDNA 4731417B20 gene | 4731417B20Rik | 2.6 | SM2148 | c | 11 |
| mPool7_V2MM_83844 | 1.732078 | 2.205692 | TRUE | HP_30536 | TGCTGTTG | GATTAAAT | NM_025700 | phosphoglucomutase 1 | Pgm1 | 2.9 | SM2270 | c | 9 |
| mPool7_V2MM_83844 | 1.732078 | 2.205692 | TRUE | HP_30536 | TGCTGTTG | GATTAAAT | NM_025700 | phosphoglucomutase 1 | Pgm1 | 2.9 | SM2270 | c | 9 |
| mPool2_V2MM_210885 | 1.731604 | 2.205692 | TRUE | HP_42862 | TGCTGTTG | GCCAGATT | XM_143332 | RIKEN cDNA 2010204N08 gene | 2010204N08Rik | 2.11 | SM2368 | g | 5 |
| mPool2_V2MM_132088 | 1.731498 | 2.205692 | TRUE | HP_35284 | TGCTGTTG | GTTATAGA | XM_489508 | RIKEN cDNA 1810013L24 gene | 1810013L24Rik | 2.11 | SM2359 | c | 8 |
| 092308m3_V2MM_32715 | 1.731085 | 66.95036 | TRUE | HP_25559 | TGCTGTTG | GTGCTTCC | NM_145602 | N-myc downstream regulated gene 4 | Ndrg4 | 2.11 | SM2400 | h | 9 |
| mPool2_V2MM_142730 | 1.730535 | 2.205692 | TRUE | HP_36344 | TGCTGTTG | GTGAATAT | XM_157867 | Mus musculus LOC240547 (LOC240547), mR. | | 2.5 | SM2059 | g | 4 |
| mPool2_V2MM_142730 | 1.730535 | 2.205692 | TRUE | HP_36344 | TGCTGTTG | GTGAATAT | XM_157867 | Mus musculus LOC240547 (LOC240547), mRNA. | | 2.5 | SM2059 | g | 4 |
| mPool2_V2MM_122425 | 1.729666 | 3.004939 | TRUE | HP_34329 | TGCTGTTG | GGATAAGT | XM_142191 | gene model 384, (NCBI) | Gm384 | 2.1 | SM2349 | a | 10 |
| 092308m1_V2MM_136395 | 1.729153 | 2.205692 | TRUE | HP_35711 | TGCTGTTG | CTGCAGA | XM_151691 | Mus musculus LOC237325 (LOC237325), mR. | | 2.4 | SM2046 | d | 9 |
| mPool2_V2MM_129626 | 1.728163 | 2.205692 | TRUE | HP_35042 | TGCTGTTG | CTAATACT | XM_146376 | Mus musculus similar to Ddx10 protein (LOC234281), mRNA. | | 2.11 | SM2373 | f | 1 |
| mPool7_V2MM_207918 | 1.727276 | 1.959154 | TRUE | HP_42571 | TGCTGTTG | CAGATAG | AY462058 | kininogen 1 | Kng1 | 2.1 | SM2332 | g | 6 |
| mPool5_V2MM_59506 | 1.727234 | 66.95036 | TRUE | HP_28168 | TGCTGTTG | GCGAAGA | NM_010111 | ephrin B2 | Efnb2 | NA | | | NA |
| mPool7_V2MM_116116 | 1.727206 | 1.959154 | TRUE | HP_33707 | TGCTGTTG | CTACAATA | XM_138757 | Mus musculus similar to ribosomal protein L7a, cytosolic [validated] - rat (LOC23 | | 2.9 | SM2300 | c | 1 |
| mPool2_V2MM_142526 | 1.726834 | 0 | TRUE | HP_36323 | TGCTGTTG | CTTAGGA | XM_157697 | Mus musculus LOC240481 (LOC240481), mRNA. | | 2.5 | SM2056 | c | 9 |
| mPool2_V2MM_142526 | 1.726834 | 0 | TRUE | HP_36323 | TGCTGTTG | CTTAGGA | XM_157697 | Mus musculus LOC240481 (LOC240481), mR. | | 2.5 | SM2056 | c | 9 |
| mPool4_V2MM_206874 | 1.726041 | 59.99801 | TRUE | HP_42469 | TGCTGTTG | CGCACCA | XM_155817 | Mus musculus LOC239491 (LOC239491), mRNA. | | 2.16 | SM2628 | h | 10 |
| 092308m1_V2MM_70012 | 1.725899 | 3.004939 | TRUE | HP_29193 | TGCTGTTG | GTGCCATT | XM_358362 | protein tyrosine phosphatase, receptor type Z, polypepti | Ptprz1 | 2.3 | SM2018 | g | 1 |
| mPool2_V2MM_121852 | 1.725074 | 3.004939 | TRUE | HP_34272 | TGCTGTTG | GACACCTT | XM_141896 | Mus musculus similar to actin, beta [Bos taurus] (LOC212460), mRNA. | | 2.11 | SM2372 | h | 6 |
| mPool7_V2MM_179672 | 1.724805 | 2.205692 | TRUE | HP_40014 | TGCTGTTG | AAATCTCA | XM_288374 | Mus musculus LOC331689 (LOC331689), mRNA. | | 2.9 | SM2274 | a | 6 |
| mPool2_V2MM_137497 | 1.722985 | 3.004939 | TRUE | HP_35821 | TGCTGTTG | GAAATTAT | XM_153221 | RIKEN cD 1110038H03 gene | 1110038H03Rik | 2.4 | SM2042 | h | 10 |
| mPool2_V2MM_137497 | 1.722985 | 3.004939 | TRUE | HP_35821 | TGCTGTTG | GAAATTAT | XM_153221 | RIKEN cDNA 1110038H03 gene | 1110038H03Rik | 2.4 | SM2042 | h | 10 |
| 092308m3_V2MM_107546 | 1.722655 | 62.64426 | TRUE | HP_32862 | TGCTGTTG | GTAGCAA | XM_284166 | RIKEN cD 5430427O21 gene | 5430427O21Rik | 2.1 | SM2320 | c | 8 |
| 092308m3_V2MM_107546 | 1.722655 | 62.64426 | TRUE | HP_32862 | TGCTGTTG | GTAGCAA | XM_284166 | RIKEN cDNA 5430427O21 gene | 5430427O21Rik | 2.1 | SM2320 | c | 8 |
| mPool2_V2MM_130006 | 1.722553 | 2.205692 | TRUE | HP_35080 | TGCTGTTG | GAGCTAAT | XM_146551 | Mus musculus LOC209349 (LOC209349), mR. | | 2.6 | SM2102 | a | 1 |
| mPool2_V2MM_130006 | 1.722553 | 2.205692 | TRUE | HP_35080 | TGCTGTTG | GAGCTAAT | XM_146551 | Mus musculus LOC209349 (LOC209349), mRNA. | | 2.6 | SM2102 | a | 1 |
| 092308m1_V2MM_177760 | 1.721438 | 3.004939 | TRUE | HP_39824 | TGCTGTTG | GAGAATG. | XM_489292 | hypothetical protein 4933400C23 | | 2.5 | SM2087 | h | 7 |
| 092308m1_V2MM_71136 | 1.720414 | 2.205692 | TRUE | HP_29303 | TGCTGTTG | CATATATA | NM_010652 | killer cell lectin-like receptor subfamily C, member 1 | Klrc1 | 2.4 | SM2021 | c | 5 |
| mPool2_V2MM_2250 | 1.720029 | 3.004939 | TRUE | HP_22587 | TGCTGTTG | CTCATTGC | NM_025972 | N-acylsphingosine amidohydrolase (acid ceramidase)-like | Asahl | 2.11 | SM2383 | b | 7 |
| 092308m1_V2MM_90381 | 1.719739 | 3.004939 | TRUE | HP_31180 | TGCTGTTG | CACTACAT | BC072604 | DEP domain containing 5 | Depdc5 | 2.6 | SM2113 | a | 3 |
| mPool6_V2MM_159578 | 1.719659 | 75.83547 | TRUE | NA | NA | NA | NA | NA | NA | NA | NA | NA | NA |

| | | | | | | | | | | | | | |
|---|---|---|---|---|---|---|---|---|---|---|---|---|---|
| mPool7_V2MM_78398 | 1.719277 | 3.004939 | TRUE | HP_30008 | TGCTGTTG | GACGGAT | NM_011240 | RAN binding protein 2 | Ranbp2 | 2.8 | SM2240 | h | 6 |
| mPool4_V2MM_21305 | 1.716725 | 60.13437 | TRUE | HP_24448 | TGCTGTTG | CAGCGAG | NM_173070 | small proline-rich protein 4 | Sprr4 | 2.16 | SM2619 | a | 2 |
| 092308m1_V2MM_179320 | 1.716666 | 3.004939 | TRUE | HP_39979 | TGCTGTTG | GAATTTAT | XM_288311 | gene model 1550, (NCBI) | Gm1550 | 2.5 | SM2080 | e | 5 |
| 092308m1_V2MM_179320 | 1.716666 | 3.004939 | TRUE | HP_39979 | TGCTGTTG | GAATTTAT | XM_288311 | gene model 1550, (NCBI) | Gm1550 | 2.5 | SM2080 | e | 5 |
| mPool2_V2MM_214253 | 1.716632 | 3.004939 | TRUE | HP_43185 | TGCTGTTG | CAATTATA | XM_140452 | Mus musculus LOC240195 (LOC240195), mRNA. | | 2.11 | SM2369 | a | 7 |
| mPool7_V2MM_104337 | 1.714827 | 3.004939 | TRUE | HP_32547 | TGCTGTTG | GAGAGAA | XM_129704 | potassium voltage-gated channel, subfamily Q, member | Kcnq5 | 2.7 | SM2153 | h | 9 |
| 092308m1_V2MM_77656 | 1.714723 | 3.004939 | TRUE | HP_29937 | TGCTGTTG | CTGTTTAA | AK004695 | vacuolar protein sorting 11 (yeast) | Vps11 | 2.3 | SM2001 | f | 9 |
| mPool7_V2MM_199967 | 1.714685 | 2.205692 | TRUE | HP_41800 | TGCTGTTG | CTGAGGA | XM_289070 | Mus musculus LOC333520 (LOC333520), mRNA. | | 2.9 | SM2289 | h | 4 |
| 092308m1_V2MM_177436 | 1.714304 | 2.205692 | TRUE | HP_39791 | TGCTGTTG | CATGTATT | XM_287874 | Mus musculus hypothetical gene supported by AK084195 (LOC330879), mR. | | 2.5 | SM2071 | h | 8 |
| mPool7_V2MM_136545 | 1.71387 | 3.004939 | TRUE | HP_35726 | TGCTGTTG | GAGTTAT | XM_151819 | Mus musculus LOC213261 (LOC213261), mRNA. | | 2.1 | SM2332 | a | 10 |
| mPool2_V2MM_111896 | 1.712256 | 2.205692 | TRUE | HP_33290 | TGCTGTTG | CGAGGTT | XM_136234 | Mus musculus similar to Rbm6 protein (LOC240782), mRNA. | | 2.7 | SM2154 | h | 10 |
| mPool7_V2MM_72556 | 1.711047 | 2.205692 | TRUE | HP_29441 | TGCTGTTG | CTGTTATA | XM_283218 | kinesin family member 13B | Kif13b | 2.8 | SM2229 | h | 7 |
| mPool2_V2MM_1143 | 1.710174 | 2.205692 | TRUE | HP_22479 | TGCTGTTG | CATATTCA | NM_027216 | solute carrier family 39 (metal ion transporter), member | Slc39a11 | 2.11 | SM2380 | d | 6 |
| 092308m1_V2MM_176265 | 1.710005 | 1.959154 | TRUE | HP_39674 | TGCTGTTG | GAGCTAT | AK028992 | RIKEN cD 6230400G14 gene | 6230400G14Rik | 2.5 | SM2076 | c | 11 |
| 092308m1_V2MM_179579 | 1.709722 | 2.205692 | TRUE | HP_40005 | TGCTGTTG | GAGTATC | XM_288357 | Mus musculus LOC331677 (LOC331677), mRNA. | | 2.5 | SM2087 | g | 7 |
| 092308m1_V2MM_179579 | 1.709722 | 2.205692 | TRUE | HP_40005 | TGCTGTTG | GAGTATC | XM_288357 | Mus musculus LOC331677 (LOC331677), mR. | | 2.5 | SM2087 | g | 7 |
| mPool2_V2MM_110362 | 1.709357 | 2.205692 | TRUE | HP_33139 | TGCTGTTG | CTGAGTTT | XM_134619 | matrix metalloproteinase 27 | Mmp27 | 2.1 | SM2323 | d | 9 |
| mPool2_V2MM_39693 | 1.70718 | 3.004939 | TRUE | HP_26237 | TGCTGTTG | GGCAGAA | NM_028139 | ataxin 7-like 4 | Atxn7l4 | 2.11 | SM2390 | c | 12 |
| mPool2_V2MM_21683 | 1.705785 | 2.205692 | TRUE | HP_24485 | TGCTGTTG | CATGTTAA | NM_030712 | chemokine (C-X-C motif) receptor 6 | Cxcr6 | 2.11 | SM2391 | a | 8 |
| 092308m1_V2MM_160811 | 1.704706 | 1.959154 | TRUE | HP_38138 | TGCTGTTG | CATATTAT | AK040883 | leucine-rich repeat-containing G protein-coupled recepto | Lgr6 | 2.6 | SM2125 | e | 10 |
| mPool2_V2MM_126032 | 1.703954 | 3.004939 | TRUE | HP_34687 | TGCTGTTG | CAGCCAC | XM_144331 | Mus musculus similar to Adaptor protein complex AP-2, alpha 2 subunit (LOC242 | | 2.1 | SM2349 | c | 9 |
| 092308m1_V2MM_172832 | 1.703612 | 3.004939 | TRUE | HP_39331 | TGCTGTTG | GATTAAA | XM_286907 | Mus musculus hypothetical gene supported by AK048737 (LOC328947), mRNA. | | 2.5 | SM2089 | f | 8 |
| 092308m1_V2MM_172832 | 1.703612 | 3.004939 | TRUE | HP_39331 | TGCTGTTG | GATTAAA | XM_286907 | Mus musculus hypothetical gene supported by AK048737 (LOC328947), mR. | | 2.5 | SM2089 | f | 8 |
| mPool2_V2MM_131554 | 1.703386 | 3.004939 | TRUE | HP_35231 | TGCTGTTG | CCCATGA | BC013539 | immunoglobulin heavy chain (J558 family) | Igh-VJ558 | 2.11 | SM2360 | f | 7 |
| mPool2_V2MM_136416 | 1.702517 | 2.205692 | TRUE | HP_35713 | TGCTGTTG | CTGATAC | XM_151707 | Mus musculus LOC237268 (LOC237268), mRNA. | | 2.5 | SM2054 | d | 3 |
| mPool2_V2MM_136416 | 1.702517 | 2.205692 | TRUE | HP_35713 | TGCTGTTG | CTGATAC | XM_151707 | Mus musculus LOC237268 (LOC237268), mR. | | 2.5 | SM2054 | d | 3 |
| 092308m1_V2MM_141774 | 1.701996 | 3.004939 | TRUE | HP_36248 | TGCTGTTG | CTTCAATA | XM_156976 | Mus musculus LOC210827 (LOC210827), mR. | | 2.4 | SM2049 | b | 6 |
| 092308m1_V2MM_141774 | 1.701996 | 3.004939 | TRUE | HP_36248 | TGCTGTTG | CTTCAATA | XM_156976 | Mus musculus LOC210827 (LOC210827), mRNA. | | 2.4 | SM2049 | b | 6 |
| mPool2_V2MM_128057 | 1.700216 | 3.004939 | TRUE | HP_34887 | TGCTGTTG | CACTAGT | XM_145511 | heat shock protein, alpha-crystallin-related, B6 | Hspb6 | 2.11 | SM2373 | a | 3 |
| mPool7_V2MM_78595 | 1.699674 | 3.004939 | TRUE | HP_30027 | TGCTGTTG | GAGACAG | NM_013709 | Sh3 domain YSC-like 1 | Sh3yl1 | 2.8 | SM2233 | c | 4 |
| 092308m3_V2MM_59009 | 1.69903 | 75.83547 | TRUE | HP_28122 | TGCTGTTG | GCAGTCA | NM_028561 | spermatogenesis associated glutamate (E)-rich protein 4l | Speer4b | 2.12 | SM2436 | c | 10 |
| mPool2_V2MM_214957 | 1.698929 | 3.004939 | TRUE | HP_43252 | TGCTGTTG | CCAAGAA | XM_141921 | gene model 370, (NCBI) | Gm370 | 2.11 | SM2359 | c | 4 |
| mPool2_V2MM_193324 | 1.697744 | 3.004939 | TRUE | HP_32914 | TGCTGTTG | GGACCAA | XM_132579 | PEST-containing nuclear protein | | 2.1 | SM2318 | c | 10 |
| mPool2_V2MM_193324 | 1.697744 | 3.004939 | TRUE | HP_32914 | TGCTGTTG | GGACCAA | XM_132579 | PEST-containing nuclear protein | | 2.1 | SM2318 | c | 10 |
| 092308m3_V2MM_48710 | 1.697484 | 59.99801 | TRUE | HP_27116 | TGCTGTTG | GCCAGTT | NM_177337 | ADP-ribosylation factor-like 11 | Arl11 | 2.12 | SM2446 | g | 2 |
| mPool7_V2MM_71767 | 1.69744 | 2.205692 | TRUE | HP_29363 | TGCTGTTG | CTATTTAT | XM_139060 | similar to TRAV3-1 | | 2.8 | SM2230 | d | 5 |
| mPool2_V2MM_125590 | 1.697176 | 2.205692 | TRUE | HP_34643 | TGCTGTTG | GGGACCA | XM_144109 | gene model 435, (NCBI) | Gm435 | 2.11 | SM2369 | c | 3 |
| mPool7_V2MM_108133 | 1.697088 | 3.004939 | TRUE | HP_32920 | TGCTGTTG | GCTGGAA | AK036732 | hypothetical protein 9830168K20 | | 2.1 | SM2306 | a | 11 |
| mPool6_V2MM_159050 | 1.694999 | 69.72441 | TRUE | HP_37965 | TGCTGTTG | CCAGCAG | XM_282962 | Mus musculus hypothetical gene supported by AK029003 (LOC327858), mRNA. | | 2.7 | SM2195 | h | 9 |
| mPool2_V2MM_113276 | 1.694849 | 2.205692 | TRUE | HP_33426 | TGCTGTTG | CACAGTA | XM_136902 | Mus musculus similar to hypothetical protein A830058L05 [Mus musculus] (LOC2 | | 2.7 | SM2152 | f | 4 |
| mPool2_V2MM_13348 | 1.694825 | 2.205692 | TRUE | HP_23671 | TGCTGTTG | CAGTTATT | NM_010738 | lymphocyte antigen 6 complex, locus A | Ly6a | 2.11 | SM2379 | b | 5 |
| mPool4_V2MM_220434 | 1.69329 | 75.83547 | TRUE | HP_45339 | TGCTGTTG | CTGTGAG | NM_028737 | RIKEN cDNA 4931406B18 gene | 4931406B18Rik | 2.15 | SM2590 | f | 10 |
| mPool2_V2MM_230 | 1.692944 | 2.205692 | TRUE | HP_22390 | TGCTGTTG | CAATTAAT | NM_010823 | myeloproliferative leukemia virus oncogene | Mpl | 2.11 | SM2383 | h | 3 |
| 092308m1_V2MM_138667 | 1.692732 | 3.004939 | TRUE | HP_35938 | TGCTGTTG | GAGCACT | XM_154403 | Mus musculus LOC238812 (LOC238812), mR. | | 2.5 | SM2056 | b | 6 |
| 092308m1_V2MM_138667 | 1.692732 | 3.004939 | TRUE | HP_35938 | TGCTGTTG | GAGCACT | XM_154403 | Mus musculus LOC238812 (LOC238812), mRNA. | | 2.5 | SM2056 | b | 6 |
| mPool7_V2MM_181334 | 1.692111 | 3.004939 | TRUE | HP_40180 | TGCTGTTG | CCTATTAC | XM_288658 | Mus musculus LOC328501 (LOC328501), mR. | | 2.5 | SM2085 | d | 11 |
| mPool7_V2MM_181334 | 1.692111 | 3.004939 | TRUE | HP_40180 | TGCTGTTG | CCTATTAC | XM_288658 | Mus musculus LOC328501 (LOC328501), mRNA. | | 2.5 | SM2085 | d | 11 |
| mPool6_V2MM_169583 | 1.691789 | 75.83547 | TRUE | HP_39007 | TGCTGTTG | CAGGGAA | AK081028 | hypothetical protein B930071A02 | | 2.8 | SM2213 | a | 8 |
| mPool7_V2MM_179579 | 1.691606 | 2.205692 | TRUE | HP_40005 | TGCTGTTG | GAGTATC | XM_288357 | Mus musculus LOC331677 (LOC331677), mRNA. | | 2.5 | SM2087 | g | 7 |
| mPool7_V2MM_179579 | 1.691606 | 2.205692 | TRUE | HP_40005 | TGCTGTTG | GAGTATC | XM_288357 | Mus musculus LOC331677 (LOC331677), mR. | | 2.5 | SM2087 | g | 7 |
| mPool2_V2MM_107243 | 1.6913 | 1.959154 | TRUE | HP_32833 | TGCTGTTG | CCTGTAAT | AK122544 | DNA segment, Chr 5, Wayne State University 178, expres | D5Wsu178e | 2.1 | SM2319 | f | 1 |
| mPool2_V2MM_111848 | 1.690387 | 2.205692 | TRUE | HP_33285 | TGCTGTTG | CAGCTAT | XM_136205 | G protein-coupled receptor 39 | Gpr39 | 2.1 | SM2320 | d | 7 |
| mPool2_V2MM_123198 | 1.690271 | 3.004939 | TRUE | HP_34405 | TGCTGTTG | CAGAATC | XM_142755 | similar to hypothetical protein FLJ90396 | | 2.11 | SM2355 | a | 8 |

| | | | | | | | | | | | | | |
|---|---|---|---|---|---|---|---|---|---|---|---|---|---|
| mPool7_V2MM_172684 | 1.687238 | 2.205692 | TRUE | HP_39317 | TGCTGTTG | CTAGAGA( | XM_286882 | Mus musculus hypothetical gene supported by AK085983 (LOC328888), mRNA. | | 2.9 | SM2282 | b | 5 |
| mPool7_V2MM_173731 | 1.686504 | 3.004939 | TRUE | HP_39421 | TGCTGTTG | CTTAATTG | XM_488936 | LOC433405 | | 2.9 | SM2273 | c | 8 |
| mPool2_V2MM_212465 | 1.686325 | 3.004939 | TRUE | HP_43013 | TGCTGTTG | CAGCTTTG | NM_207632 | olfactory receptor 1118 | Olfr1118 | 2.11 | SM2355 | a | 5 |
| mPool7_V2MM_89292 | 1.683966 | 3.004939 | TRUE | HP_31072 | TGCTGTTG | GTGGATA | NM_177564 | cDNA sequence BC022224 | BC022224 | 2.9 | SM2259 | c | 2 |
| 092308m1_V2MM_136416 | 1.683669 | 3.004939 | TRUE | HP_35713 | TGCTGTTG | CTGATACC | XM_151707 | Mus musculus LOC237268 (LOC237268), mR. | | 2.5 | SM2054 | d | 3 |
| 092308m1_V2MM_136416 | 1.683669 | 3.004939 | TRUE | HP_35713 | TGCTGTTG | CTGATACC | XM_151707 | Mus musculus LOC237268 (LOC237268), mRNA. | | 2.5 | SM2054 | d | 3 |
| mPool2_V2MM_207912 | 1.683554 | 3.004939 | TRUE | HP_42570 | TGCTGTTG | CATCTTCC | XM_195479 | Mus musculus similar to argonaute 5 protein [Mus musculus] (LOC269496), mRN | | 2.1 | SM2335 | b | 8 |
| mPool5_V2MM_160937 | 1.680462 | 75.83547 | TRUE | HP_38151 | TGCTGTTG | GCCGCAA( | XM_486014 | similar to zinc finger and BTB domain containing 2 | | NA | | | NA |
| mPool4_V2MM_114976 | 1.680189 | 75.83547 | TRUE | HP_33594 | TGCTGTTG | CTTTGGAT | XM_138051 | Mus musculus similar to Heterogeneous nuclear ribonucleoprotein A1 (Helix-des | | 2.16 | SM2626 | d | 6 |
| 092308m1_V2MM_164058 | 1.67844 | 3.004939 | TRUE | HP_38460 | TGCTGTTG | GCATGTA | XM_486406 | similar to godotropin inducible ovarian transcription factor 1 | | 2.6 | SM2120 | e | 9 |
| mPool2_V2MM_144690 | 1.676626 | 3.004939 | TRUE | HP_36540 | TGCTGTTG | GCTATGG | XM_160200 | Mus musculus LOC213229 (LOC213229), mRNA. | | 2.1 | SM2334 | g | 2 |
| mPool2_V2MM_125824 | 1.676015 | 3.004939 | TRUE | HP_34666 | TGCTGTTG | CAAATAG | XM_144242 | heterogeneous nuclear ribonucleoprotein A3 | Hnrpa3 | 2.11 | SM2354 | e | 8 |
| mPool7_V2MM_152662 | 1.675901 | 3.004939 | TRUE | HP_37332 | TGCTGTTG | CCAAGAA( | XM_195495 | Mus musculus similar to ribosomal protein S17 [Mus musculus] (LOC269534), mF | | 2.1 | SM2329 | c | 3 |
| mPool2_V2MM_213282 | 1.675459 | 3.004939 | TRUE | HP_43092 | TGCTGTTG | GGATTATT | XM_141458 | Mus musculus similar to olfactory receptor MOR245-10 [Mus musculus] (LOC228 | | 2.11 | SM2362 | a | 4 |
| mPool2_V2MM_211088 | 1.674746 | 3.004939 | TRUE | HP_42881 | TGCTGTTG | GACTGAA( | XM_147017 | similar to Chromosome 6 open reading frame 117 | | 2.1 | SM2348 | f | 1 |
| 092308m1_V2MM_149887 | 1.67447 | 3.004939 | TRUE | HP_37059 | TGCTGTTG | GTTTAGTT | XM_165303 | Mus musculus LOC208460 (LOC208460), mR. | | 2.4 | SM2049 | a | 6 |
| mPool2_V2MM_103007 | 1.673055 | 1.959154 | TRUE | HP_32415 | TGCTGTTG | GAACTGA | XM_128751 | cysteine-rich motor neuron 1 | Crim1 | 2.1 | SM2311 | c | 3 |
| mPool7_V2MM_141500 | 1.672061 | 3.004939 | TRUE | HP_36221 | TGCTGTTG | ATCCCAAC | XM_156723 | Mus musculus LOC210877 (LOC210877), mR. | | 2.5 | SM2057 | d | 7 |
| mPool7_V2MM_141500 | 1.672061 | 3.004939 | TRUE | HP_36221 | TGCTGTTG | ATCCCAAC | XM_156723 | Mus musculus LOC210877 (LOC210877), mRNA. | | 2.5 | SM2057 | d | 7 |
| 092308m1_V2MM_132242 | 1.671695 | 2.205692 | TRUE | HP_35299 | TGCTGTTG | CCTCCTAA | AK080331 | RIKEN cD A630055G03 gene | A630055G03Rik | 2.6 | SM2111 | b | 9 |
| 092308m1_V2MM_63607 | 1.670955 | 3.004939 | TRUE | HP_28568 | TGCTGTTG | CTAAGCA | NM_029682 | RIKEN cD 1700095N21 gene | 1700095N21Rik | 2.3 | SM2013 | c | 5 |
| 092308m1_V2MM_123913 | 1.669969 | 3.004939 | TRUE | HP_34476 | TGCTGTTG | GACTACA( | XM_143206 | Mus musculus similar to zo pellucida protein 1 [Gallus gallus] (LOC215202), mR. | | 2.5 | SM2099 | c | 1 |
| mPool7_V2MM_81409 | 1.669353 | 3.004939 | TRUE | HP_30301 | TGCTGTTG | GTATTTAT | NM_172859 | RIKEN cDNA 2810039F03 gene | 2810039F03Rik | 2.8 | SM2231 | a | 4 |
| 092308m1_V2MM_171812 | 1.66744 | 2.205692 | TRUE | HP_39229 | TGCTGTTG | CTGCTTAT | XM_286699 | Mus musculus hypothetical gene supported by AK029886 (LOC328542), mR. | | 2.5 | SM2073 | h | 7 |
| mPool2_V2MM_95470 | 1.667251 | 3.004939 | TRUE | HP_31674 | TGCTGTTG | CCGTCTGT | NM_198010 | ankyrin repeat domain 17 | Ankrd17 | 2.6 | SM2134 | g | 5 |
| mPool7_V2MM_84826 | 1.665726 | 3.004939 | TRUE | HP_30632 | TGCTGTTG | CTGGGAC | NM_029545 | RIKEN cDNA 6530401N04 gene | 6530401N04Rik | 2.9 | SM2258 | d | 12 |
| mPool7_V2MM_195814 | 1.665548 | 1.959154 | TRUE | HP_41393 | TGCTGTTG | GGGCATT( | NM_175270 | RIKEN cDNA 5730467H21 gene | 5730467H21Rik | 2.9 | SM2251 | f | 3 |
| 092308m1_V2MM_181041 | 1.665303 | 2.205692 | TRUE | HP_40151 | TGCTGTTG | GACATTAT | XM_288609 | Mus musculus LOC332013 (LOC332013), mR. | | 2.5 | SM2062 | c | 10 |
| 092308m1_V2MM_181041 | 1.665303 | 2.205692 | TRUE | HP_40151 | TGCTGTTG | GACATTAT | XM_288609 | Mus musculus LOC332013 (LOC332013), mRNA. | | 2.5 | SM2062 | c | 10 |
| 092308m1_V2MM_150691 | 1.664663 | 2.205692 | TRUE | HP_37137 | TGCTGTTG | GTAGGAC | NM_177857 | RIKEN cD A930010I20 gene | A930010I20Rik | 2.4 | SM2041 | g | 4 |
| mPool2_V2MM_112258 | 1.664356 | 2.205692 | TRUE | HP_33326 | TGCTGTTG | GGCCATTT | XM_488159 | similar to protease | | 2.7 | SM2157 | b | 10 |
| 092308m3_V2MM_83872 | 1.662958 | 35.99881 | TRUE | HP_30539 | TGCTGTTG | CTGAGTA | NM_025890 | RIKEN cD 2410004A20 gene | 2410004A20Rik | 2.13 | SM2458 | g | 4 |
| 092308m3_V2MM_25446 | 1.662316 | 59.99801 | TRUE | HP_24851 | TGCTGTTG | CAGCCAG | NM_172411 | RIKEN cD 2310007B03 gene | 2310007B03Rik | 2.12 | SM2427 | g | 12 |
| mPool7_V2MM_95902 | 1.660192 | 1.959154 | TRUE | NA | NA | NA | NA | NA | NA | NA | NA | NA | NA |
| 092308m1_V2MM_154894 | 1.65988 | 3.004939 | TRUE | HP_37553 | TGCTGTTG | CCATGACA | XM_197074 | Mus musculus LOC271145 (LOC271145), mR. | | 2.6 | SM2122 | h | 2 |
| mPool2_V2MM_214331 | 1.659023 | 2.205692 | TRUE | HP_43192 | TGCTGTTG | GAAAGTT( | XM_147679 | Mus musculus LOC209126 (LOC209126), mRNA. | | 2.11 | SM2365 | c | 8 |
| 092308m3_V2MM_59041 | 1.657501 | 61.29828 | TRUE | HP_28125 | TGCTGTTG | GCATCCA( | NM_177030 | RIKEN cD C330023D02 gene | C330023D02Rik | 2.12 | SM2416 | g | 6 |
| mPool2_V2MM_118588 | 1.657123 | 3.004939 | TRUE | HP_33952 | TGCTGTTG | GATGAGG | XM_139952 | Mus musculus similar to olfactory receptor MOR121-1 [Mus musculus] (LOC2097 | | 2.11 | SM2355 | a | 3 |
| mPool2_V2MM_97315 | 1.65693 | 3.004939 | TRUE | HP_31855 | TGCTGTTG | GATCAAG | XM_111430 | Mus musculus similar to serine protease inhibitor 10 [Mus musculus] (LOC19343 | | 2.6 | SM2107 | d | 4 |
| mPool2_V2MM_97315 | 1.65693 | 3.004939 | TRUE | HP_31855 | TGCTGTTG | GATCAAG | XM_111430 | Mus musculus similar to serine protease inhibitor 10 [Mus musculus] (LOC19343 | | 2.6 | SM2107 | d | 4 |
| mPool2_V2MM_101471 | 1.65571 | 3.004939 | TRUE | HP_32264 | TGCTGTTG | GCTCTGTT | AK044563 | DNA segment, Chr 14, Abbott 1 expressed | D14Abb1e | 2.7 | SM2153 | b | 4 |
| mPool2_V2MM_115134 | 1.654911 | 3.004939 | TRUE | HP_33610 | TGCTGTTG | CATGAGTT | XM_138128 | Mus musculus similar to Embryonic growth-associated (LOC217630), mRNA. | | 2.6 | SM2149 | c | 2 |
| 092308m3_V2MM_49151 | 1.65458 | 62.64426 | TRUE | HP_27159 | TGCTGTTG | GGCAAAG | NM_172524 | RIKEN cD 9530066K23 gene | 9530066K23Rik | 2.12 | SM2416 | a | 4 |
| mPool2_V2MM_104352 | 1.654459 | 3.004939 | TRUE | HP_32548 | TGCTGTTG | CCTATTGC | AK077997 | RIKEN cDNA 6130401J04 gene | 6130401J04Rik | 2.7 | SM2157 | e | 2 |
| mPool7_V2MM_64588 | 1.654208 | 2.205692 | TRUE | HP_28663 | TGCTGTTG | CTGTGTTT | NM_026367 | G patch domain containing 2 | Gpatc2 | 2.8 | SM2232 | e | 10 |
| mPool6_V2MM_161630 | 1.653826 | 60.13437 | TRUE | HP_38219 | TGCTGTTG | CCCTGATT | XM_283985 | Mus musculus hypothetical gene supported by AK089333 (LOC329899), mRNA. | | 2.8 | SM2209 | d | 9 |
| 092308m1_V2MM_62958 | 1.653809 | 3.004939 | TRUE | HP_28505 | TGCTGTTG | CCAGTAA | NM_011316 | serum amyloid A 4 | Saa4 | 2.4 | SM2020 | g | 8 |
| 092308m1_V2MM_87165 | 1.65377 | 2.205692 | TRUE | HP_30862 | TGCTGTTG | CTGGTTAC | NM_175099 | replication initiator 1 | Repin1 | 2.6 | SM2108 | f | 11 |
| mPool7_V2MM_187884 | 1.653066 | 1.959154 | TRUE | HP_40834 | TGCTGTTG | CATGGAC | XM_489130 | hypothetical gene supported by AK018238; AK046289 | | 2.5 | SM2076 | h | 2 |
| mPool7_V2MM_187884 | 1.653066 | 1.959154 | TRUE | HP_40834 | TGCTGTTG | CATGGAC | XM_489130 | hypothetical gene supported by AK018238; AK046289 | | 2.5 | SM2076 | h | 2 |
| mPool2_V2MM_149064 | 1.652872 | 2.205692 | TRUE | HP_36977 | TGCTGTTG | CCATGATA | XM_164446 | Mus musculus LOC244637 (LOC244637), mRNA. | | 2.1 | SM2344 | c | 9 |
| 092308m3_V2MM_237848 | 1.652659 | 60.13437 | TRUE | HP_48969 | TGCTGTTG | CTATGAG( | XM_154800 | Mus musculus LOC239116 (LOC239116), mR. | | 2.14 | SM2528 | d | 5 |
| mPool7_V2MM_84788 | 1.652556 | 2.205692 | TRUE | HP_30629 | TGCTGTTG | CCTCCATT | NM_029282 | RIKEN cDNA 2610036D13 gene | 2610036D13Rik | 2.9 | SM2256 | g | 1 |

| | | | | | | | | | | | | | |
|---|---|---|---|---|---|---|---|---|---|---|---|---|---|
| 092308m1_V2MM_141969 | 1.651983 | 3.004939 | TRUE | HP_36268 | TGCTGTTG | CCTGGAAT | XM_157241 | Mus musculus LOC240232 (LOC240232), mR. | | 2.5 | SM2052 | h | 11 |
| mPool2_V2MM_19547 | 1.651104 | 3.004939 | TRUE | HP_24276 | TGCTGTTG | GGTGTAAA | XM_133269 | Mus musculus similar to Glycogen synthase kinase-3 alpha (GSK-3 alpha) (Factor | | 2.11 | SM2376 | g | 4 |
| mPool7_V2MM_92122 | 1.650871 | 3.004939 | TRUE | HP_31348 | TGCTGTTG | CATTGAAT | NM_178653 | RIKEN cD C330023F11 gene | C330023F11Rik | 2.9 | SM2263 | g | 9 |
| mPool7_V2MM_92122 | 1.650871 | 3.004939 | TRUE | HP_31348 | TGCTGTTG | CATTGAAT | NM_178653 | RIKEN cDNA C330023F11 gene | C330023F11Rik | 2.9 | SM2263 | g | 9 |
| mPool7_V2MM_62581 | 1.650682 | 3.004939 | TRUE | HP_28468 | TGCTGTTG | CAGGTAGT | NM_146936 | olfactory receptor 1417 | Olfr1417 | 2.8 | SM2242 | c | 7 |
| 092308m1_V2MM_123721 | 1.650348 | 3.004939 | TRUE | HP_34457 | TGCTGTTG | CTTTATCG | XM_143117 | Mus musculus similar to peptidylprolyl isomerase D (cyclophilin D); hCyP40 [Hom | | 2.6 | SM2114 | g | 3 |
| mPool7_V2MM_81341 | 1.649455 | 2.205692 | TRUE | HP_30294 | TGCTGTTG | GTACGATA | NM_021704 | chemokine (C-X-C motif) ligand 12 | Cxcl12 | 2.8 | SM2243 | h | 11 |
| mPool7_V2MM_184802 | 1.648989 | 3.004939 | TRUE | HP_40527 | TGCTGTTG | CGGTTAAT | XM_289261 | Mus musculus LOC332907 (LOC332907), mRNA. | | 2.5 | SM2086 | c | 5 |
| mPool7_V2MM_184802 | 1.648989 | 3.004939 | TRUE | HP_40527 | TGCTGTTG | CGGTTAAT | XM_289261 | Mus musculus LOC332907 (LOC332907), mR. | | 2.5 | SM2086 | c | 5 |
| mPool7_V2MM_116331 | 1.648744 | 2.205692 | TRUE | HP_33728 | TGCTGTTG | GAGTCTAC | XM_138876 | gene model 280, (NCBI) | Gm280 | 2.1 | SM2302 | e | 3 |
| 092308m1_V2MM_165864 | 1.647286 | 3.004939 | TRUE | HP_38639 | TGCTGTTG | GTCAATTC | XM_285342 | cardiomyopathy associated 3 | Cmya3 | 2.6 | SM2125 | a | 8 |
| mPool7_V2MM_172357 | 1.644231 | 2.205692 | TRUE | HP_39284 | TGCTGTTG | CCATGGTC | AK051520 | hypothetical protein D130054H01 | | 2.5 | SM2095 | c | 6 |
| mPool7_V2MM_172357 | 1.644231 | 2.205692 | TRUE | HP_39284 | TGCTGTTG | CCATGGTC | AK051520 | hypothetical protein D130054H01 | | 2.5 | SM2095 | c | 6 |
| 092308m3_V2MM_238015 | 1.643452 | 75.83547 | TRUE | HP_48985 | TGCTGTTG | GCCTCTGC | XM_158288 | Mus musculus LOC241355 (LOC241355), mR. | | 2.14 | SM2517 | g | 5 |
| 092308m3_V2MM_112866 | 1.643403 | 66.95036 | TRUE | HP_33385 | TGCTGTTG | GCTGTCGA | AK122502 | cD sequence BC056494 | BC056494 | 2.13 | SM2486 | h | 2 |
| mPool6_V2MM_68227 | 1.642929 | 75.83547 | TRUE | HP_29019 | TGCTGTTG | CTGTGTGT | NM_026100 | RIKEN cDNA 1700055O19 gene | 1700055O19Rik | 2.7 | SM2177 | b | 11 |
| 092308m1_V2MM_187364 | 1.642089 | 3.004939 | TRUE | HP_40783 | TGCTGTTG | GGGAAAT | XM_289743 | Mus musculus LOC333651 (LOC333651), mR. | | 2.5 | SM2080 | b | 9 |
| mPool2_V2MM_213434 | 1.640589 | 2.205692 | TRUE | HP_43106 | TGCTGTTG | CGGGACCA | XM_144109 | gene model 435, (NCBI) | Gm435 | 2.11 | SM2362 | e | 1 |
| mPool7_V2MM_67030 | 1.640248 | 3.004939 | TRUE | HP_28903 | TGCTGTTG | CCCATTAC | NM_010484 | solute carrier family 6 (neurotransmitter transporter, ser | Slc6a4 | 2.8 | SM2233 | g | 1 |
| mPool6_V2MM_160680 | 1.638854 | 69.72441 | TRUE | HP_38125 | TGCTGTTG | GAGCTCAT | NM_010678 | lymphoid nuclear protein related to AF4 | Laf4 | 2.8 | SM2207 | c | 9 |
| 092308m1_V2MM_186219 | 1.638758 | 3.004939 | TRUE | HP_40669 | TGCTGTTG | CAGAGAAT | XM_289502 | Mus musculus LOC333219 (LOC333219), mRNA. | | 2.5 | SM2066 | d | 12 |
| 092308m1_V2MM_186219 | 1.638758 | 3.004939 | TRUE | HP_40669 | TGCTGTTG | CAGAGAAT | XM_289502 | Mus musculus LOC333219 (LOC333219), mR. | | 2.5 | SM2066 | d | 12 |
| mPool2_V2MM_214049 | 1.635909 | 3.004939 | TRUE | HP_43166 | TGCTGTTG | GAATTTAT | XM_143381 | Mus musculus similar to 60S acidic ribosomal protein P1 (LOC229548), mRNA. | | 2.11 | SM2354 | g | 3 |
| 092308m1_V2MM_146848 | 1.635419 | 3.004939 | TRUE | HP_36755 | TGCTGTTG | CGGGTAAC | XM_162361 | Mus musculus LOC210681 (LOC210681), mR. | | 2.4 | SM2045 | b | 3 |
| 092308m1_V2MM_146848 | 1.635419 | 3.004939 | TRUE | HP_36755 | TGCTGTTG | CGGGTAAC | XM_162361 | Mus musculus LOC210681 (LOC210681), mRNA. | | 2.4 | SM2045 | b | 3 |
| mPool2_V2MM_128478 | 1.635243 | 2.205692 | TRUE | HP_34929 | TGCTGTTG | GTCATAGA | XM_145695 | Mus musculus similar to glyceraldehyde-3-phosphate dehydrogenase [Mus musc | | 2.11 | SM2364 | a | 5 |
| mPool2_V2MM_30073 | 1.634736 | 3.004939 | TRUE | HP_25301 | TGCTGTTG | CAGTACAT | NM_148948 | Dicer1, Dcr-1 homolog (Drosophila) | Dicer1 | 2.11 | SM2391 | b | 5 |
| 092308m3_V2MM_48109 | 1.633919 | 75.83547 | TRUE | HP_27057 | TGCTGTTG | GACTGGAC | NM_016784 | pleiotropic regulator 1, PRL1 homolog (Arabidopsis) | Plrg1 | 2.12 | SM2437 | d | 2 |
| mPool2_V2MM_12429 | 1.631928 | 2.205692 | TRUE | HP_23582 | TGCTGTTG | CAAATATT | NM_007550 | Bloom syndrome homolog (human) | Blm | 2.11 | SM2383 | c | 5 |
| 092308m3_V2MM_104592 | 1.631834 | 73.80895 | TRUE | HP_32572 | TGCTGTTG | CTGACAGA | NM_198899 | UDP-glucose ceramide glucosyltransferase-like 1 | Ugcgl1 | 2.9 | SM2296 | d | 12 |
| 092308m3_V2MM_104592 | 1.631834 | 73.80895 | TRUE | HP_32572 | TGCTGTTG | CTGACAGA | NM_198899 | UDP-glucose ceramide glucosyltransferase-like 1 | Ugcgl1 | 2.9 | SM2296 | d | 12 |
| mPool2_V2MM_123892 | 1.631352 | 3.004939 | TRUE | HP_34474 | TGCTGTTG | CGAGTTCA | XM_143201 | similar to Polyadenylate-binding protein 4 (Poly(A)-binding protein 4) (PABP 4) (I | | 2.11 | SM2361 | g | 7 |
| 092308m1_V2MM_64758 | 1.629823 | 3.004939 | TRUE | HP_28680 | TGCTGTTG | CTTGTAAC | NM_026202 | RIKEN cD 2610529H08 gene | 2610529H08Rik | 2.3 | SM2007 | h | 1 |
| 092308m1_V2MM_185449 | 1.62962 | 2.205692 | TRUE | HP_40592 | TGCTGTTG | GACCTTTC | XM_289372 | SEC31-like 1 (S. cerevisiae) | Sec31l1 | 2.5 | SM2094 | b | 3 |
| 092308m1_V2MM_170776 | 1.629174 | 3.004939 | TRUE | HP_39126 | TGCTGTTG | CAATCTTA | AK049051 | engulfment and cell motility 1, ced-12 homolog (C. elega | Elmo1 | 2.5 | SM2069 | c | 4 |
| 092308m1_V2MM_170776 | 1.629174 | 3.004939 | TRUE | HP_39126 | TGCTGTTG | CAATCTTA | AK049051 | engulfment and cell motility 1, ced-12 homolog (C. elega | Elmo1 | 2.5 | SM2069 | c | 4 |
| 092308m1_V2MM_182147 | 1.62846 | 3.004939 | TRUE | HP_40262 | TGCTGTTG | GGTGTACA | XM_288795 | Mus musculus LOC332264 (LOC332264), mRNA. | | 2.5 | SM2095 | a | 12 |
| 092308m1_V2MM_182147 | 1.62846 | 3.004939 | TRUE | HP_40262 | TGCTGTTG | GGTGTACA | XM_288795 | Mus musculus LOC332264 (LOC332264), mR. | | 2.5 | SM2095 | a | 12 |
| mPool7_V2MM_64772 | 1.627981 | 2.205692 | TRUE | HP_28682 | TGCTGTTG | CTTGTTAT | NM_007484 | ras homolog gene family, member C | Rhoc | 2.8 | SM2234 | a | 12 |
| mPool7_V2MM_88549 | 1.627651 | 2.205692 | TRUE | HP_30998 | TGCTGTTG | GTCTGTAA | NM_175431 | RIKEN cDNA A830093I24 gene | A830093I24Rik | 2.9 | SM2262 | c | 4 |
| mPool7_V2MM_90928 | 1.626334 | 3.004939 | TRUE | HP_31234 | TGCTGTTG | CTGCTAAC | NM_177888 | zinc finger protein 78 | Zfp78 | 2.9 | SM2255 | g | 8 |
| mPool7_V2MM_90928 | 1.626334 | 3.004939 | TRUE | HP_31234 | TGCTGTTG | CTGCTAAC | NM_177888 | zinc finger protein 78 | Zfp78 | 2.9 | SM2255 | g | 8 |
| mPool2_V2MM_117781 | 1.626197 | 3.004939 | TRUE | HP_33872 | TGCTGTTG | CAGAGGT | XM_139533 | RIKEN cDNA A630029G22 gene | A630029G22Rik | 2.11 | SM2369 | f | 8 |
| mPool2_V2MM_202297 | 1.625141 | 3.004939 | TRUE | HP_42030 | TGCTGTTG | CACCTTCT | XM_136883 | Mus musculus similar to KIAA1007 protein; adrenal gland protein AD-005 [Homo | | 2.1 | SM2333 | c | 4 |
| 092308m3_V2MM_233281 | 1.623726 | 59.99801 | TRUE | HP_48519 | TGCTGTTG | CACAGCAA | NM_199147 | cD sequence BC062185 | BC062185 | 2.14 | SM2503 | f | 12 |
| 092308m3_V2MM_208742 | 1.621922 | 75.83547 | TRUE | HP_42652 | TGCTGTTG | CCAACTTA | XM_154583 | Mus musculus LOC238932 (LOC238932), mR. | | 2.14 | SM2512 | g | 8 |
| 092308m1_V2MM_95911 | 1.62175 | 2.205692 | TRUE | HP_31717 | TGCTGTTG | CTGGTGAC | XM_109763 | Mus musculus LOC192909 (LOC192909), mR. | | 2.6 | SM2110 | a | 11 |
| 092308m1_V2MM_147408 | 1.618192 | 2.205692 | TRUE | HP_36811 | TGCTGTTG | GAAACAGT | XM_162954 | Mus musculus LOC243724 (LOC243724), mR. | | 2.5 | SM2058 | d | 5 |
| 092308m1_V2MM_147408 | 1.618192 | 2.205692 | TRUE | HP_36811 | TGCTGTTG | GAAACAGT | XM_162954 | Mus musculus LOC243724 (LOC243724), mRNA. | | 2.5 | SM2058 | d | 5 |
| 092308m1_V2MM_153474 | 1.617355 | 3.004939 | TRUE | HP_37412 | TGCTGTTG | CCTTGGTA | AK013526 | RIKEN cD 2900011G08 gene | 2900011G08Rik | 2.6 | SM2115 | b | 12 |
| mPool7_V2MM_62079 | 1.616822 | 2.205692 | TRUE | HP_28419 | TGCTGTTG | CACCTTAA | NM_146864 | olfactory receptor 768 | Olfr768 | 2.8 | SM2236 | b | 8 |
| mPool2_V2MM_142836 | 1.615709 | 2.205692 | TRUE | HP_36354 | TGCTGTTG | GGTCCTAT | XM_157999 | Mus musculus LOC213992 (LOC213992), mRNA. | | 2.1 | SM2333 | f | 1 |
| mPool2_V2MM_142836 | 1.615709 | 2.205692 | TRUE | HP_36354 | TGCTGTTG | GGTCCTAT | XM_157999 | Mus musculus LOC213992 (LOC213992), mR. | | 2.1 | SM2333 | f | 1 |

| | | | | | | | | | | | | | |
|---|---|---|---|---|---|---|---|---|---|---|---|---|---|
| 092308m1_V2MM_184054 | 1.615425 | 3.004939 | TRUE | HP_40452 | TGCTGTTG | GCCTAGAT | XM_289132 | Mus musculus LOC331157 (LOC331157), mR. | | 2.5 | SM2081 | b | 7 |
| mPool2_V2MM_121026 | 1.615336 | 3.004939 | TRUE | HP_34191 | TGCTGTTG | CATTGGCC | XM_141393 | Mus musculus LOC241683 (LOC241683), mRNA. | | 2.11 | SM2361 | d | 2 |
| mPool2_V2MM_137537 | 1.613534 | 2.205692 | TRUE | HP_35825 | TGCTGTTG | CTTAAAGT | XM_153261 | similar to Rho GTPase activating protein 20 | | 2.4 | SM2046 | h | 8 |
| mPool2_V2MM_137537 | 1.613534 | 2.205692 | TRUE | HP_35825 | TGCTGTTG | CTTAAAGT | XM_153261 | similar to Rho GTPase activating protein 20 | | 2.4 | SM2046 | h | 8 |
| mPool2_V2MM_116048 | 1.613069 | 3.004939 | TRUE | HP_33700 | TGCTGTTG | GTCTGGAT | XM_488036 | similar to MrgA4 RF-amide G protein-coupled receptor | | 2.1 | SM2319 | g | 10 |
| mPool7_V2MM_170515 | 1.612718 | 2.205692 | TRUE | HP_39100 | TGCTGTTG | GCATATTA | AK087847 | RIKEN cDNA E330029E12 gene | E330029E12Rik | 2.5 | SM2068 | f | 4 |
| mPool7_V2MM_170515 | 1.612718 | 2.205692 | TRUE | HP_39100 | TGCTGTTG | GCATATTA | AK087847 | RIKEN cD E330029E12 gene | E330029E12Rik | 2.5 | SM2068 | f | 4 |
| mPool2_V2MM_91548 | 1.612562 | 3.004939 | TRUE | NA | NA | NA | NA | NA | NA | NA | NA | NA | NA |
| mPool7_V2MM_186370 | 1.612237 | 3.004939 | TRUE | HP_40684 | TGCTGTTG | GTGAAGG | XM_289527 | RIKEN cDNA A030007E19 gene | A030007E19Rik | 2.9 | SM2282 | c | 12 |
| mPool2_V2MM_113507 | 1.611841 | 3.004939 | TRUE | HP_33449 | TGCTGTTG | GTCTTTAT | XM_136994 | gene model 229, (NCBI) | Gm229 | 2.7 | SM2157 | h | 3 |
| mPool2_V2MM_115503 | 1.611368 | 3.004939 | TRUE | HP_33646 | TGCTGTTG | CCCTTCAA | XM_138425 | Mus musculus similar to general transcription factor IIa, 2 (12kD subunit) [Rattus | | 2.7 | SM2159 | b | 11 |
| 092308m1_V2MM_136568 | 1.611111 | 3.004939 | TRUE | HP_35728 | TGCTGTTG | CCTCATCC | XM_151865 | Mus musculus LOC215140 (LOC215140), mRNA. | | 2.4 | SM2050 | b | 7 |
| 092308m1_V2MM_136568 | 1.611111 | 3.004939 | TRUE | HP_35728 | TGCTGTTG | CCTCATCC | XM_151865 | Mus musculus LOC215140 (LOC215140), mR. | | 2.4 | SM2050 | b | 7 |
| mPool2_V2MM_25233 | 1.610976 | 1.959154 | TRUE | HP_24830 | TGCTGTTG | CACATCAA | NM_133680 | RIKEN cDNA 5730414C17 gene | 5730414C17Rik | 2.11 | SM2391 | f | 4 |
| mPool2_V2MM_122861 | 1.609653 | 3.004939 | TRUE | HP_34371 | TGCTGTTG | CATAAAGC | XM_142466 | Mus musculus similar to GLE1-like, RNA export mediator; GLE1 (yeast homolog)-l | | 2.11 | SM2358 | a | 9 |
| 092308m1_V2MM_75421 | 1.609311 | 3.004939 | TRUE | HP_29720 | TGCTGTTG | CCATTGGA | NM_010424 | hemochromatosis | Hfe | 2.3 | SM2002 | h | 8 |
| mPool7_V2MM_181041 | 1.609267 | 1.959154 | TRUE | HP_40151 | TGCTGTTG | GACATTAT | XM_288609 | Mus musculus LOC332013 (LOC332013), mR. | | 2.5 | SM2062 | c | 10 |
| mPool7_V2MM_181041 | 1.609267 | 1.959154 | TRUE | HP_40151 | TGCTGTTG | GACATTAT | XM_288609 | Mus musculus LOC332013 (LOC332013), mRNA. | | 2.5 | SM2062 | c | 10 |
| 092308m1_V2MM_179098 | 1.609079 | 3.004939 | TRUE | HP_39957 | TGCTGTTG | CACATTCA | XM_288274 | Mus musculus LOC332550 (LOC332550), mR. | | 2.5 | SM2086 | d | 11 |
| mPool7_V2MM_206355 | 1.609063 | 3.004939 | TRUE | HP_42418 | TGCTGTTG | CTGTTAAA | AK032192 | potassium inwardly-rectifying channel, subfamily J, mem | Kcnj16 | 2.9 | SM2298 | b | 7 |
| mPool2_V2MM_123965 | 1.608449 | 2.205692 | TRUE | HP_34481 | TGCTGTTG | CAGGAATC | XM_143234 | Mus musculus similar to embryonic blastocoelar extracellular matrix protein prec | | 2.11 | SM2367 | f | 4 |
| 092308m1_V2MM_173192 | 1.608417 | 2.205692 | TRUE | HP_39367 | TGCTGTTG | CAGAGTA | XM_286976 | Mus musculus hypothetical gene supported by AK013638; AK030685; AK049099; | | 2.5 | SM2084 | d | 9 |
| 092308m3_V2MM_202996 | 1.607836 | 75.83547 | TRUE | HP_42097 | TGCTGTTG | CATCAGA | NM_133738 | anthrax toxin receptor 2 | Antxr2 | 2.13 | SM2497 | e | 7 |
| mPool2_V2MM_88659 | 1.60698 | 2.205692 | TRUE | HP_31009 | TGCTGTTG | CCATTTGC | NM_175460 | nicotinamide nucleotide adenylyltransferase 2 | Nmnat2 | 2.6 | SM2134 | c | 3 |
| mPool4_V2MM_163217 | 1.602686 | 61.29828 | TRUE | HP_38377 | TGCTGTTG | GTGTTCAA | XM_284721 | gene model 774, (NCBI) | Gm774 | 2.16 | SM2649 | a | 8 |
| mPool7_V2MM_147048 | 1.602061 | 2.205692 | TRUE | HP_36775 | TGCTGTTG | CACAAAG | XM_162569 | Mus musculus LOC243509 (LOC243509), mRNA. | | 2.4 | SM2041 | e | 9 |
| mPool7_V2MM_147048 | 1.602061 | 2.205692 | TRUE | HP_36775 | TGCTGTTG | CACAAAG | XM_162569 | Mus musculus LOC243509 (LOC243509), mR. | | 2.4 | SM2041 | e | 9 |
| 092308m1_V2MM_151228 | 1.601286 | 2.205692 | TRUE | HP_37189 | TGCTGTTG | GTGAAAT | NM_001002272 | trophinin | Tro | 2.4 | SM2050 | c | 10 |
| 092308m1_V2MM_151228 | 1.601286 | 2.205692 | TRUE | HP_37189 | TGCTGTTG | GTGAAAT | NM_001002272 | trophinin | Tro | 2.4 | SM2050 | c | 10 |
| 092308m1_V2MM_130006 | 1.59975 | 3.004939 | TRUE | HP_35080 | TGCTGTTG | GAGCTAAT | XM_146551 | Mus musculus LOC209349 (LOC209349), mR. | | 2.6 | SM2102 | a | 1 |
| 092308m1_V2MM_130006 | 1.59975 | 3.004939 | TRUE | HP_35080 | TGCTGTTG | GAGCTAAT | XM_146551 | Mus musculus LOC209349 (LOC209349), mRNA. | | 2.6 | SM2102 | a | 1 |
| 092308m1_V2MM_84816 | 1.599032 | 3.004939 | TRUE | HP_30631 | TGCTGTTG | CAGTAACT | NM_029498 | zinc finger protein 198 | Zfp198 | 2.6 | SM2109 | d | 8 |
| mPool2_V2MM_125294 | 1.598047 | 2.205692 | TRUE | HP_34613 | TGCTGTTG | CACCTCCT | XM_143897 | Mus musculus similar to zinc finger protein 352 (LOC242525), mRNA. | | 2.1 | SM2349 | b | 5 |
| mPool7_V2MM_179143 | 1.597158 | 2.205692 | TRUE | HP_39961 | TGCTGTTG | GGGAGAC | XM_288281 | Mus musculus LOC332557 (LOC332557), mRNA. | | 2.9 | SM2296 | e | 5 |
| mPool7_V2MM_186219 | 1.597042 | 3.004939 | TRUE | HP_40669 | TGCTGTTG | CAGAGAAT | XM_289502 | Mus musculus LOC333219 (LOC333219), mR. | | 2.5 | SM2066 | d | 12 |
| mPool7_V2MM_186219 | 1.597042 | 3.004939 | TRUE | HP_40669 | TGCTGTTG | CAGAGAAT | XM_289502 | Mus musculus LOC333219 (LOC333219), mRNA. | | 2.5 | SM2066 | d | 12 |
| mPool6_V2MM_66304 | 1.596576 | 61.29828 | TRUE | HP_28832 | TGCTGTTG | CACCAAG | NM_011156 | prolyl endopeptidase | Prep | 2.7 | SM2173 | g | 2 |
| 092308m1_V2MM_132069 | 1.596359 | 3.004939 | TRUE | HP_35282 | TGCTGTTG | CTGTTTGC | BC028850 | RIKEN cD 1500002B03 gene | 1500002B03Rik | 2.6 | SM2104 | h | 9 |
| mPool6_V2MM_159548 | 1.596288 | 59.99801 | TRUE | HP_38014 | TGCTGTTG | GCCTAGGT | AK079330 | RIKEN cDNA 9630025I21 gene | 9630025I21Rik | 2.7 | SM2200 | e | 1 |
| mPool2_V2MM_33465 | 1.595741 | 3.004939 | TRUE | HP_25631 | TGCTGTTG | CAGATGAC | NM_144895 | spastic paraplegia 20, spartin (Troyer syndrome) homolo | Spg20 | 2.11 | SM2389 | h | 12 |
| 092308m1_V2MM_148147 | 1.595412 | 3.004939 | TRUE | HP_36885 | TGCTGTTG | GAAGGTTC | XM_163530 | Mus musculus LOC244133 (LOC244133), mRNA. | | 2.4 | SM2040 | h | 1 |
| 092308m1_V2MM_148147 | 1.595412 | 3.004939 | TRUE | HP_36885 | TGCTGTTG | GAAGGTTC | XM_163530 | Mus musculus LOC244133 (LOC244133), mR. | | 2.4 | SM2040 | h | 1 |
| mPool7_V2MM_68331 | 1.595013 | 2.205692 | TRUE | HP_29029 | TGCTGTTG | CTTTAAGT | NM_030081 | zinc finger, FYVE domain containing 20 | Zfyve20 | 2.8 | SM2247 | a | 2 |
| mPool2_V2MM_159305 | 1.594981 | 2.205692 | TRUE | HP_37990 | TGCTGTTG | GAGGTAG | XM_283042 | Mus musculus hypothetical gene supported by AK051127 (LOC328052), mRNA. | | 2.6 | SM2128 | h | 1 |
| mPool5_V2MM_45643 | 1.594664 | 69.72441 | TRUE | HP_26817 | TGCTGTTG | CAAGGACT | NM_010171 | coagulation factor III | F3 | NA | | | NA |
| mPool7_V2MM_64143 | 1.594046 | 3.004939 | TRUE | HP_28620 | TGCTGTTG | CTGACACC | NM_025781 | RIKEN cDNA 9030409E16 gene | 9030409E16Rik | 2.8 | SM2242 | f | 5 |
| 092308m3_V2MM_227395 | 1.594037 | 75.83547 | TRUE | HP_47939 | TGCTGTTG | GCCTTTGA | NM_183217 | cD sequence BC042698 | BC042698 | 2.13 | SM2469 | c | 11 |
| 092308m3_V2MM_30001 | 1.592893 | 35.99881 | TRUE | HP_25294 | TGCTGTTG | CAGGCTGC | NM_015740 | biogenesis of lysosome-related organelles complex-1, sul | Bloc1s1 | 2.12 | SM2431 | f | 4 |
| mPool2_V2MM_17541 | 1.591709 | 3.004939 | TRUE | HP_24081 | TGCTGTTG | GATAGGTT | NM_133206 | zinc and ring finger 1 | Znrf1 | 2.11 | SM2381 | c | 12 |
| mPool7_V2MM_202608 | 1.590471 | 2.205692 | TRUE | HP_42060 | TGCTGTTG | CCAAGTTT | AK015845 | RIKEN cDNA 4930519L02 gene | 4930519L02Rik | 2.9 | SM2299 | h | 12 |
| mPool7_V2MM_113916 | 1.590406 | 1.959154 | TRUE | HP_33490 | TGCTGTTG | GATCTATT | XM_137192 | similar to EC1-V2R pheromone receptor protein | | 2.1 | SM2306 | f | 11 |
| 092308m3_V2MM_194750 | 1.589511 | 61.29828 | TRUE | HP_41290 | TGCTGTTG | GATGCTCC | NM_178645 | bleomycin hydrolase | Blmh | 2.12 | SM2450 | h | 4 |
| 092308m3_V2MM_112923 | 1.588719 | 69.72441 | TRUE | HP_33391 | TGCTGTTG | GGTAAGCT | XM_136719 | Mus musculus LOC213721 (LOC213721), mR. | | 2.13 | SM2494 | c | 1 |

| | | | | | | | | | | | | | |
|---|---|---|---|---|---|---|---|---|---|---|---|---|---|
| mPool2_V2MM_21587 | 1.587724 | 3.004939 | TRUE | HP_24475 | TGCTGTTG | CATATCAT | NM_011991 | COP9 (constitutive photomorphogenic) homolog, subuni | Cops3 | 2.7 | SM2165 | b | 11 |
| 092308m1_V2MM_173404 | 1.587555 | 3.004939 | TRUE | HP_39389 | TGCTGTTG | GTAATAA( | NM_172463 | sushi, nidogen and EGF-like domains 1 | Sned1 | 2.5 | SM2079 | a | 8 |
| 092308m3_V2MM_206242 | 1.586854 | 75.83547 | TRUE | HP_42407 | TGCTGTTG | GCAACGC | XM_137051 | Mus musculus similar to Gag [Ovis aries] (LOC212923), mRNA. | | 2.9 | SM2300 | d | 5 |
| 092308m3_V2MM_206242 | 1.586854 | 75.83547 | TRUE | HP_42407 | TGCTGTTG | GCAACGC | XM_137051 | Mus musculus similar to Gag [Ovis aries] (LOC212923), mR. | | 2.9 | SM2300 | d | 5 |
| mPool7_V2MM_173267 | 1.58674 | 3.004939 | TRUE | HP_39375 | TGCTGTTG | GTAATTAT | XM_286992 | Mus musculus hypothetical gene supported by AK031258 (LOC329151), mRNA. | | 2.9 | SM2284 | e | 8 |
| mPool2_V2MM_135185 | 1.586555 | 3.004939 | TRUE | HP_35590 | TGCTGTTG | GTCTAAAT | XM_150392 | protein phosphatase 1A, magnesium dependent, alpha is | Ppm1a | 2.4 | SM2045 | e | 12 |
| mPool2_V2MM_135185 | 1.586555 | 3.004939 | TRUE | HP_35590 | TGCTGTTG | GTCTAAAT | XM_150392 | protein phosphatase 1A, magnesium dependent, alpha is | Ppm1a | 2.4 | SM2045 | e | 12 |
| 092308m1_V2MM_63506 | 1.586532 | 3.004939 | TRUE | HP_28558 | TGCTGTTG | CGGTGCT( | NM_029597 | RIKEN cD 1700018B08 gene | 1700018B08Rik | 2.4 | SM2025 | g | 4 |
| mPool7_V2MM_96391 | 1.585834 | 3.004939 | TRUE | HP_31764 | TGCTGTTG | GAAATGT | AK015302 | RIKEN cDNA 4933432K03 gene | 4933432K03Rik | 2.9 | SM2251 | d | 3 |
| mPool7_V2MM_96391 | 1.585834 | 3.004939 | TRUE | HP_31764 | TGCTGTTG | GAAATGT | AK015302 | RIKEN cD 4933432K03 gene | 4933432K03Rik | 2.9 | SM2251 | d | 3 |
| mPool2_V2MM_112209 | 1.585509 | 2.205692 | TRUE | HP_33321 | TGCTGTTG | CTGCATA( | XM_136373 | RIKEN cDNA 4930413P14 gene | 4930413P14Rik | 2.6 | SM2148 | b | 11 |
| mPool6_V2MM_5275 | 1.584245 | 59.99801 | TRUE | HP_22882 | TGCTGTTG | CCAGTGT( | NM_011629 | nuclear receptor subfamily 2, group C, member 1 | Nr2c1 | 2.7 | SM2172 | a | 6 |
| mPool7_V2MM_87647 | 1.583858 | 2.205692 | TRUE | HP_30909 | TGCTGTTG | CCACTATA | NM_175212 | RIKEN cDNA 4930438D12 gene | 4930438D12Rik | 2.9 | SM2263 | f | 5 |
| mPool7_V2MM_87647 | 1.583858 | 2.205692 | TRUE | HP_30909 | TGCTGTTG | CCACTATA | NM_175212 | RIKEN cD 4930438D12 gene | 4930438D12Rik | 2.9 | SM2263 | f | 5 |
| mPool2_V2MM_146549 | 1.583004 | 2.205692 | TRUE | HP_36726 | TGCTGTTG | CCTGCTCT | XM_162150 | Mus musculus LOC243169 (LOC243169), mRNA. | | 2.1 | SM2337 | c | 10 |
| mPool7_V2MM_205921 | 1.582907 | 3.004939 | TRUE | HP_42376 | TGCTGTTG | CTGGGAA( | AK122392 | expressed sequence AI447928 | AI447928 | 2.1 | SM2307 | b | 5 |
| 092308m3_V2MM_85571 | 1.582464 | 61.29828 | TRUE | HP_30705 | TGCTGTTG | CCATGAA( | NM_146414 | olfactory receptor 1431 | Olfr1431 | 2.13 | SM2476 | g | 11 |
| 092308m3_V2MM_114909 | 1.582009 | 75.83547 | TRUE | HP_33588 | TGCTGTTG | GCACTGA | XM_138011 | RIKEN cDNA E030025L21 gene | E030025L21Rik | 2.1 | SM2306 | b | 6 |
| 092308m3_V2MM_114909 | 1.582009 | 75.83547 | TRUE | HP_33588 | TGCTGTTG | GCACTGA | XM_138011 | RIKEN cD E030025L21 gene | E030025L21Rik | 2.1 | SM2306 | b | 6 |
| mPool2_V2MM_4273 | 1.581661 | 3.004939 | TRUE | HP_22784 | TGCTGTTG | GTACTTCA | NM_019580 | membrane interacting protein of RGS16 | | 2.11 | SM2379 | b | 7 |
| mPool5_V2MM_171369 | 1.580833 | 75.83547 | TRUE | HP_39185 | TGCTGTTG | GACCCAC | XM_286611 | Mus musculus hypothetical gene supported by AK040135 (LOC328411), mRNA. | | NA | | | NA |
| mPool2_V2MM_104822 | 1.580602 | 2.205692 | TRUE | HP_32595 | TGCTGTTG | GCTGTACA | AK015207 | RIKEN cDNA 4930426L09 gene | 4930426L09Rik | 2.7 | SM2153 | d | 8 |
| mPool2_V2MM_211363 | 1.578616 | 2.205692 | TRUE | HP_42906 | TGCTGTTG | CAACTGA( | XM_489556 | similar to hypothetical protein | | 2.11 | SM2361 | h | 10 |
| mPool2_V2MM_105368 | 1.578457 | 3.004939 | TRUE | HP_32648 | TGCTGTTG | CCCATGA | AK016695 | RIKEN cDNA 4933406J08 gene | 4933406J08Rik | 2.1 | SM2319 | b | 11 |
| 092308m1_V2MM_178182 | 1.577267 | 2.205692 | TRUE | HP_39865 | TGCTGTTG | CATATATC | AK035100 | RIKEN cD 9430088F21 gene | 9430088F21Rik | 2.5 | SM2072 | a | 3 |
| 092308m3_V2MM_33913 | 1.576187 | 75.83547 | TRUE | HP_25674 | TGCTGTTG | CATGATGT | NM_024435 | neurotensin | Nts | 2.11 | SM2393 | h | 2 |
| 092308m1_V2MM_174234 | 1.575705 | 3.004939 | TRUE | HP_39472 | TGCTGTTG | CTTGCTGT | AK087574 | RIKEN cD E230011G24 gene | E230011G24Rik | 2.5 | SM2060 | h | 8 |
| mPool2_V2MM_210407 | 1.575421 | 3.004939 | TRUE | HP_42816 | TGCTGTTG | GGTGCAG | XM_142191 | gene model 384, (NCBI) | Gm384 | 2.11 | SM2373 | f | 12 |
| mPool2_V2MM_101245 | 1.574392 | 3.004939 | TRUE | HP_32241 | TGCTGTTG | CTAACCAA | AK083705 | expressed sequence AI256775 | AI256775 | 2.1 | SM2324 | e | 1 |
| mPool2_V2MM_101245 | 1.574392 | 3.004939 | TRUE | HP_32241 | TGCTGTTG | CTAACCAA | AK083705 | expressed sequence AI256775 | AI256775 | 2.1 | SM2324 | e | 1 |
| mPool2_V2MM_97289 | 1.573556 | 2.205692 | TRUE | HP_31853 | TGCTGTTG | CTCTGAAA | XM_111418 | Mus musculus similar to ribosomal protein S9, cytosolic [validated] - rat (LOC218 | | 2.1 | SM2318 | c | 12 |
| 092308m1_V2MM_178624 | 1.572382 | 3.004939 | TRUE | HP_39909 | TGCTGTTG | GTGATAA( | XM_288193 | Mus musculus LOC332456 (LOC332456), mR. | | 2.5 | SM2088 | b | 1 |
| 092308m1_V2MM_178624 | 1.572382 | 3.004939 | TRUE | HP_39909 | TGCTGTTG | GTGATAA( | XM_288193 | Mus musculus LOC332456 (LOC332456), mRNA. | | 2.5 | SM2088 | b | 1 |
| 092308m1_V2MM_119523 | 1.572155 | 3.004939 | TRUE | HP_34044 | TGCTGTTG | CTTCCACA | XM_140470 | Mus musculus similar to ribosomal protein L10 [Rattus norvegicus] (LOC225234), | | 2.5 | SM2097 | d | 1 |
| 092308m1_V2MM_119523 | 1.572155 | 3.004939 | TRUE | HP_34044 | TGCTGTTG | CTTCCACA | XM_140470 | Mus musculus similar to ribosomal protein L10 [Rattus norvegicus] (LOC225234), | | 2.5 | SM2097 | d | 1 |
| mPool2_V2MM_88856 | 1.572011 | 2.205692 | TRUE | HP_31029 | TGCTGTTG | CTCCTTTG | NM_175510 | RIKEN cDNA C030018G13 gene | C030018G13Rik | 2.6 | SM2134 | e | 1 |
| 092308m1_V2MM_141195 | 1.57155 | 2.205692 | TRUE | HP_36190 | TGCTGTTG | GACAAAT | XM_156514 | Mus musculus LOC239894 (LOC239894), mRNA. | | 2.4 | SM2042 | d | 4 |
| 092308m1_V2MM_141195 | 1.57155 | 2.205692 | TRUE | HP_36190 | TGCTGTTG | GACAAAT | XM_156514 | Mus musculus LOC239894 (LOC239894), mR. | | 2.4 | SM2042 | d | 4 |
| 092308m1_V2MM_144387 | 1.571385 | 3.004939 | TRUE | HP_36509 | TGCTGTTG | CTGGTCA( | XM_159943 | Mus musculus LOC241958 (LOC241958), mR. | | 2.4 | SM2038 | d | 2 |
| 092308m1_V2MM_78999 | 1.571372 | 2.205692 | TRUE | HP_30067 | TGCTGTTG | GAGTATA( | NM_146775 | olfactory receptor 473 | Olfr473 | 2.4 | SM2026 | a | 9 |
| 092308m1_V2MM_70732 | 1.57053 | 3.004939 | TRUE | HP_29263 | TGCTGTTG | CAGAATG | NM_053090 | down-regulated by Ctnnb1, a | | 2.4 | SM2021 | e | 5 |
| mPool2_V2MM_212032 | 1.570461 | 2.205692 | TRUE | HP_42971 | TGCTGTTG | CACTTATT | XM_141313 | Mus musculus LOC214223 (LOC214223), mRNA. | | 2.11 | SM2367 | e | 1 |
| 092308m3_V2MM_197834 | 1.5698 | 75.83547 | TRUE | HP_41589 | TGCTGTTG | GATGATA | XM_111554 | similar to Aspartate aminotransferase, mitochondrial precursor (Transaminase A | | 2.9 | SM2266 | b | 12 |
| 092308m3_V2MM_197834 | 1.5698 | 75.83547 | TRUE | HP_41589 | TGCTGTTG | GATGATA | XM_111554 | similar to Aspartate aminotransferase, mitochondrial precursor (Transamise A) ( | | 2.9 | SM2266 | b | 12 |
| mPool2_V2MM_112686 | 1.56727 | 3.004939 | TRUE | HP_18032 | TGCTGTTG | GAAGCTA | XM_486245 | similar to Rpl7a protein | | 2.6 | SM2150 | f | 9 |
| 092308m1_V2MM_123687 | 1.566683 | 3.004939 | TRUE | HP_34454 | TGCTGTTG | CAATGAA | XM_143099 | Mus musculus similar to glyceraldehyde-3-phosphate dehydrogese [Mus muscul | | 2.5 | SM2097 | d | 9 |
| mPool2_V2MM_107197 | 1.56653 | 3.004939 | TRUE | HP_32828 | TGCTGTTG | CAGCCAA( | AK017309 | peroxisome biogenesis factor 1 | Pex1 | 2.7 | SM2154 | g | 3 |
| mPool2_V2MM_118608 | 1.565198 | 3.004939 | TRUE | HP_33954 | TGCTGTTG | CTCTAAAT | XM_139997 | Mus musculus similar to glyceraldehyde-3-phosphate dehydrogenase [Mus musc | | 2.11 | SM2372 | d | 1 |
| mPool2_V2MM_143932 | 1.564727 | 3.004939 | TRUE | HP_36464 | TGCTGTTG | CAGAGAT | XM_159469 | Mus musculus LOC211711 (LOC211711), mR. | | 2.4 | SM2033 | d | 1 |
| mPool2_V2MM_143932 | 1.564727 | 3.004939 | TRUE | HP_36464 | TGCTGTTG | CAGAGAT | XM_159469 | Mus musculus LOC211711 (LOC211711), mRNA. | | 2.4 | SM2033 | d | 1 |
| 092308m1_V2MM_179050 | 1.563952 | 3.004939 | TRUE | HP_39952 | TGCTGTTG | CCAGTTTG | XM_288266 | Mus musculus LOC332539 (LOC332539), mRNA. | | 2.5 | SM2070 | b | 6 |
| 092308m1_V2MM_179050 | 1.563952 | 3.004939 | TRUE | HP_39952 | TGCTGTTG | CCAGTTTG | XM_288266 | Mus musculus LOC332539 (LOC332539), mR. | | 2.5 | SM2070 | b | 6 |
| mPool5_V2MM_19921 | 1.56273 | 66.95036 | TRUE | HP_24312 | TGCTGTTG | GTCTCGG | NM_025807 | solute carrier family 16 (monocarboxylic acid transporter | Slc16a9 | NA | | | NA |

| | | | | | | | | | | | | | | | |
|---|---|---|---|---|---|---|---|---|---|---|---|---|---|---|---|
| mPool4_V2MM_214759 | 1.561206 | 75.83547 | TRUE | HP_43233 | TGCTGTTG | CAACCATC | XM_149204 | RIKEN cDNA 2610507N02 gene | 2610507N02Rik | | 2.16 | SM2639 | c | | 11 |
| 092308m1_V2MM_147040 | 1.55998 | 3.004939 | TRUE | HP_36775 | TGCTGTTG | CAGTTATC | XM_162552 | Mus musculus LOC243500 (LOC243500), mR. | | | 2.4 | SM2045 | a | | 5 |
| mPool2_V2MM_123084 | 1.559656 | 2.205692 | TRUE | HP_34394 | TGCTGTTG | GGTGAAAT | XM_142640 | serine (or cysteine) proteinase inhibitor, clade B (ovalbur | Serpinb3b | | 2.11 | SM2368 | e | | 1 |
| 092308m1_V2MM_141042 | 1.559204 | 3.004939 | TRUE | HP_36175 | TGCTGTTG | GCCTATCC | XM_156433 | Mus musculus LOC239944 (LOC239944), mR. | | | 2.4 | SM2045 | c | | 6 |
| 092308m3_V2MM_91895 | 1.557893 | 35.99881 | TRUE | HP_31326 | TGCTGTTG | GCCATTGC | NM_178612 | RIKEN cD 2610019P18 gene | 2610019P18Rik | | 2.13 | SM2485 | h | | 9 |
| mPool2_V2MM_211803 | 1.557815 | 3.004939 | TRUE | HP_42948 | TGCTGTTG | GCACCTAT | XM_143256 | similar to hypothetical protein C130079G13 | | | 2.11 | SM2353 | e | | 3 |
| mPool7_V2MM_175565 | 1.557728 | 3.004939 | TRUE | HP_39604 | TGCTGTTG | CTGATAGC | XM_489100 | hypothetical gene supported by AK031827 | | | 2.9 | SM2287 | b | | 2 |
| mPool2_V2MM_125842 | 1.557424 | 3.004939 | TRUE | HP_34668 | TGCTGTTG | GAGCCATT | XM_144249 | Mus musculus similar to vacuole membrane protein 1 [Rattus norvegicus] (LOC23 | | | 2.11 | SM2365 | a | | 4 |
| mPool7_V2MM_202467 | 1.557327 | 2.205692 | TRUE | HP_42047 | TGCTGTTG | CTGGTGAC | AK075654 | RIKEN cDNA 3830408G10 gene | 3830408G10Rik | | 2.1 | SM2313 | c | | 12 |
| mPool2_V2MM_208600 | 1.55657 | 2.205692 | TRUE | HP_32966 | TGCTGTTG | GAGAAGA | XM_289903 | RIKEN cDNA 2500002G23 gene | 2500002G23Rik | | 2.1 | SM2334 | g | | 6 |
| mPool4_V2MM_235772 | 1.555647 | 75.83547 | TRUE | HP_48763 | TGCTGTTG | GTGTACAT | XM_155527 | Mus musculus LOC207548 (LOC207548), mRNA. | | | 2.16 | SM2644 | e | | 11 |
| 092308m3_V2MM_226600 | 1.554549 | 60.13437 | TRUE | HP_47861 | TGCTGTTG | CTGTCTCT | NM_178690 | RIKEN cD 4732493F09 gene | 4732493F09Rik | | 2.13 | SM2482 | d | | 6 |
| mPool7_V2MM_196687 | 1.554047 | 3.004939 | TRUE | HP_41478 | TGCTGTTG | CTGTGAAA | NM_175519 | potassium channel tetramerisation domain containing 8 | Kctd8 | | 2.9 | SM2265 | g | | 5 |
| 092308m1_V2MM_169472 | 1.553705 | 3.004939 | TRUE | HP_38996 | TGCTGTTG | CCTCCGTA | XM_286267 | Mus musculus hypothetical gene supported by AK038448 (LOC327785), mR. | | | 2.6 | SM2122 | b | | 10 |
| 092308m3_V2MM_103006 | 1.553484 | 75.83547 | TRUE | NA | NA | NA | NA | NA | NA | NA | | NA | NA | NA | |
| mPool7_V2MM_91650 | 1.552805 | 3.004939 | TRUE | HP_31301 | TGCTGTTG | GTTATTAC | NM_178397 | RIKEN cDNA 2210404D11 gene | 2210404D11Rik | | 2.9 | SM2255 | g | | 12 |
| mPool2_V2MM_128961 | 1.551638 | 2.205692 | TRUE | HP_34977 | TGCTGTTG | GATTATCA | XM_146032 | Mus musculus similar to ORC1-related protein (LOC233711), mRNA. | | | 2.11 | SM2365 | d | | 9 |
| 092308m3_V2MM_226910 | 1.551575 | 75.83547 | TRUE | HP_47891 | TGCTGTTG | GTGTTTAT | NM_178613 | RIKEN cD 4933433P14 gene | 4933433P14Rik | | 2.13 | SM2474 | d | | 11 |
| mPool6_V2MM_81066 | 1.550831 | 75.83547 | TRUE | HP_30268 | TGCTGTTG | GGCTGCTT | NM_008071 | gamma-aminobutyric acid (GABA-A) receptor, subunit be | Gabrb3 | | 2.7 | SM2181 | h | | 11 |
| mPool7_V2MM_84322 | 1.549913 | 3.004939 | TRUE | HP_30583 | TGCTGTTG | CTGTATAA | XM_484255 | nucleolar protein 8 | Nol8 | | 2.9 | SM2268 | h | | 7 |
| mPool7_V2MM_82060 | 1.549911 | 3.004939 | TRUE | HP_30364 | TGCTGTTG | GTTCAAGT | NM_146966 | olfactory receptor 1247 | Olfr1247 | | 2.8 | SM2229 | f | | 4 |
| 092308m1_V2MM_145690 | 1.549745 | 2.205692 | TRUE | HP_36640 | TGCTGTTG | GAGGAAA | XM_161242 | Mus musculus LOC242544 (LOC242544), mR. | | NA | | | | NA | |
| mPool2_V2MM_144083 | 1.549507 | 1.959154 | TRUE | HP_36479 | TGCTGTTG | CCAGATAC | AY462058 | kininogen 1 | Kng1 | | 2.1 | SM2335 | e | | 2 |
| mPool2_V2MM_209186 | 1.548832 | 3.004939 | TRUE | HP_42696 | TGCTGTTG | CTGGAAAT | XM_151696 | Mus musculus LOC237328 (LOC237328), mRNA. | | | 2.1 | SM2336 | h | | 2 |
| mPool7_V2MM_183932 | 1.548526 | 3.004939 | TRUE | HP_40440 | TGCTGTTG | CACCTCCT | XM_289106 | Mus musculus LOC331565 (LOC331565), mRNA. | | | 2.9 | SM2277 | g | | 2 |
| mPool7_V2MM_174757 | 1.548195 | 3.004939 | TRUE | HP_39524 | TGCTGTTG | CAATCCGC | XM_287297 | Mus musculus hypothetical gene supported by AK080820 (LOC329769), mRNA. | | | 2.9 | SM2288 | e | | 12 |
| mPool2_V2MM_98015 | 1.547167 | 1.959154 | TRUE | HP_31925 | TGCTGTTG | GGATATTC | XM_112038 | Mus musculus similar to putative pheromone receptor (LOC195691), mRNA. | | | 2.6 | SM2145 | g | | 3 |
| 092308m3_V2MM_194721 | 1.546031 | 75.83547 | TRUE | HP_41287 | TGCTGTTG | CACCCGGT | NM_027008 | potassium channel tetramerisation domain containing 5 | Kctd5 | | 2.8 | SM2249 | a | | 6 |
| 092308m3_V2MM_194721 | 1.546031 | 75.83547 | TRUE | HP_41287 | TGCTGTTG | CACCCGGT | NM_027008 | potassium channel tetramerisation domain containing 5 | Kctd5 | | 2.8 | SM2249 | a | | 6 |
| mPool2_V2MM_105383 | 1.545677 | 3.004939 | TRUE | HP_32649 | TGCTGTTG | GACTATAC | BC024760 | cDNA sequence BC024760 | BC024760 | | 2.7 | SM2154 | e | | 7 |
| mPool2_V2MM_18640 | 1.544649 | 3.004939 | TRUE | HP_24188 | TGCTGTTG | GCTCCTAT | NM_018864 | inositol (myo)-1(or 4)-monophosphatase 1 | Impa1 | | 2.11 | SM2381 | g | | 3 |
| 092308m3_V2MM_103386 | 1.543647 | 60.13437 | TRUE | HP_32452 | TGCTGTTG | CAGCTGTT | AK020725 | RIKEN cD 2700046A07 gene | 2700046A07Rik | | 2.14 | SM2506 | c | | 6 |
| mPool7_V2MM_173876 | 1.54347 | 2.205692 | TRUE | HP_39436 | TGCTGTTG | CAGTGTCA | AK038492 | UDP-N-acetyl-alpha-D-galactosamine:polypeptide N-acet | Galnt13 | | 2.9 | SM2271 | b | | 1 |
| mPool2_V2MM_13272 | 1.543097 | 2.205692 | TRUE | HP_23664 | TGCTGTTG | CAGTCATT | NM_009062 | regulator of G-protein signaling 4 | Rgs4 | | 2.11 | SM2375 | h | | 8 |
| mPool2_V2MM_213248 | 1.542836 | 3.004939 | TRUE | HP_43088 | TGCTGTTG | CGTATTAT | XM_141639 | Mus musculus similar to protease [Mus musculus] (LOC236704), mRNA. | | | 2.11 | SM2372 | d | | 11 |
| mPool6_V2MM_160575 | 1.541416 | 69.72441 | TRUE | HP_38115 | TGCTGTTG | CATCGCGT | XM_283569 | Mus musculus hypothetical gene supported by AK081501 (LOC329063), mRNA. | | | 2.8 | SM2206 | a | | 4 |
| mPool2_V2MM_124711 | 1.538964 | 3.004939 | TRUE | HP_34556 | TGCTGTTG | GTCAGTGT | XM_143599 | Mus musculus similar to RNA polymerase II transcriptional coactivator [Mus mus | | | 2.11 | SM2367 | c | | 2 |
| 092308m1_V2MM_174386 | 1.536601 | 3.004939 | TRUE | HP_39487 | TGCTGTTG | GGGTTAGT | XM_287220 | Mus musculus hypothetical gene supported by AK045957 (LOC329604), mR. | | | 2.5 | SM2079 | g | | 10 |
| 092308m1_V2MM_144635 | 1.536364 | 3.004939 | TRUE | HP_36534 | TGCTGTTG | CATCCCAC | XM_160151 | Mus musculus LOC212834 (LOC212834), mR. | | | 2.5 | SM2057 | f | | 9 |
| mPool7_V2MM_197062 | 1.536062 | 2.205692 | TRUE | HP_41514 | TGCTGTTG | GTCTATTC | NM_146660 | olfactory receptor 1135 | Olfr1135 | | 2.9 | SM2255 | g | | 6 |
| mPool2_V2MM_4364 | 1.535818 | 2.205692 | TRUE | HP_22793 | TGCTGTTG | GTCTTTGA | NM_027356 | RIKEN cDNA 2700038N03 gene | 2700038N03Rik | | 2.11 | SM2383 | e | | 11 |
| mPool2_V2MM_111805 | 1.535686 | 3.004939 | TRUE | HP_33281 | TGCTGTTG | GACAAATC | XM_136178 | expressed sequence AI132431 | AI132431 | | 2.7 | SM2158 | b | | 10 |
| 092308m1_V2MM_187047 | 1.534193 | 3.004939 | TRUE | HP_40751 | TGCTGTTG | CAAAGAAC | XM_289640 | Mus musculus LOC333402 (LOC333402), mR. | | | 2.5 | SM2072 | f | | 3 |
| 092308m1_V2MM_187047 | 1.534193 | 3.004939 | TRUE | HP_40751 | TGCTGTTG | CAAAGAAC | XM_289640 | Mus musculus LOC333402 (LOC333402), mRNA. | | | 2.5 | SM2072 | f | | 3 |
| mPool6_V2MM_155861 | 1.534173 | 69.72441 | TRUE | HP_37650 | TGCTGTTG | CATATTTG | XM_197863 | Mus musculus similar to hypothetical protein MGC19022 [Mus musculus] (LOC27 | | | 2.7 | SM2199 | c | | 7 |
| mPool4_V2MM_215022 | 1.534074 | 62.64426 | TRUE | HP_43259 | TGCTGTTG | CAGCCACT | XM_141013 | Mus musculus similar to mitochondrial ribosomal protein S36; similar to RIKEN cI | | | 2.1 | SM2349 | d | | 8 |
| mPool2_V2MM_213314 | 1.533775 | 3.004939 | TRUE | HP_43095 | TGCTGTTG | CCCTGTGA | XM_149823 | Mus musculus LOC232625 (LOC232625), mRNA. | | | 2.11 | SM2361 | b | | 3 |
| mPool5_V2MM_160442 | 1.533276 | 70.87591 | TRUE | HP_38102 | TGCTGTTG | GCGTCTAT | AK033711 | hypothetical protein 9130229N11 | | NA | | | | NA | |
| 092308m1_V2MM_172048 | 1.531848 | 2.205692 | TRUE | HP_39253 | TGCTGTTG | CAACAGTC | XM_286748 | Mus musculus hypothetical gene supported by AK079475 (LOC328668), mR. | | | 2.5 | SM2081 | a | | 11 |
| 092308m3_V2MM_88257 | 1.531505 | 66.95036 | TRUE | HP_30969 | TGCTGTTG | GTTAACAT | NM_175355 | RIKEN cD E330013P04 gene | E330013P04Rik | | 2.9 | SM2254 | e | | 10 |
| 092308m3_V2MM_88257 | 1.531505 | 66.95036 | TRUE | HP_30969 | TGCTGTTG | GTTAACAT | NM_175355 | RIKEN cDNA E330013P04 gene | E330013P04Rik | | 2.9 | SM2254 | e | | 10 |
| 092308m1_V2MM_167173 | 1.531002 | 3.004939 | TRUE | HP_38768 | TGCTGTTG | CAGCCTAT | XM_285713 | Mus musculus similar to variant-specific surface protein S7 [Giardia intestilis] (LO | | | 2.6 | SM2124 | d | | 10 |
| 092308m1_V2MM_70292 | 1.530813 | 3.004939 | TRUE | HP_29220 | TGCTGTTG | CAAATTTG | NM_010514 | insulin-like growth factor 2 | Igf2 | | 2.3 | SM2009 | g | | 10 |

| | | | | | | | | | | | | | |
|---|---|---|---|---|---|---|---|---|---|---|---|---|---|
| mPool2_V2MM_118341 | 1.530787 | 2.205692 | TRUE | HP_33927 | TGCTGTTG | CCTGTATA | XM_139796 | Mus musculus similar to putative pheromone receptor [Mus musculus] (LOC2245 | | 2.11 | SM2354 | a | 12 |
| mPool2_V2MM_96187 | 1.530785 | 3.004939 | TRUE | HP_31744 | TGCTGTTG | CTTCCTAA | XM_489704 | similar to G protein coupled receptor | | 2.6 | SM2138 | a | 1 |
| mPool7_V2MM_176671 | 1.529466 | 3.004939 | TRUE | HP_39715 | TGCTGTTG | CTCAGGA | XM_287716 | Mus musculus hypothetical gene supported by AK085843 (LOC330610), mRNA. | | 2.9 | SM2273 | f | 1 |
| mPool7_V2MM_173142 | 1.529013 | 3.004939 | TRUE | HP_39362 | TGCTGTTG | GGAGTAAT | XM_286966 | Mus musculus hypothetical gene supported by AK051123 (LOC329083), mR. | | 2.5 | SM2089 | f | 4 |
| mPool7_V2MM_173142 | 1.529013 | 3.004939 | TRUE | HP_39362 | TGCTGTTG | GGAGTAAT | XM_286966 | Mus musculus hypothetical gene supported by AK051123 (LOC329083), mRNA. | | 2.5 | SM2089 | f | 4 |
| 092308m1_V2MM_179101 | 1.528739 | 3.004939 | TRUE | HP_39957 | TGCTGTTG | CCACATTC | XM_288274 | Mus musculus LOC332550 (LOC332550), mR. | | 2.5 | SM2092 | a | 9 |
| 092308m1_V2MM_179101 | 1.528739 | 3.004939 | TRUE | HP_39957 | TGCTGTTG | CCACATTC | XM_288274 | Mus musculus LOC332550 (LOC332550), mRNA. | | 2.5 | SM2092 | a | 9 |
| mPool4_V2MM_221042 | 1.528693 | 75.83547 | TRUE | HP_45398 | TGCTGTTG | GTTCAAGT | NM_146828 | olfactory receptor 975 | Olfr975 | 2.15 | SM2598 | c | 10 |
| mPool2_V2MM_120787 | 1.527815 | 3.004939 | TRUE | HP_34167 | TGCTGTTG | CCAGTAAT | XM_485031 | olfactory receptor 48 | Olfr48 | 2.11 | SM2353 | a | 3 |
| 092308m1_V2MM_184437 | 1.527239 | 3.004939 | TRUE | HP_40490 | TGCTGTTG | CTGTCTAT | XM_289198 | Mus musculus LOC332831 (LOC332831), mR. | | 2.5 | SM2067 | b | 3 |
| mPool2_V2MM_169682 | 1.526893 | 3.004939 | TRUE | HP_39017 | TGCTGTTG | CAACCAAT | XM_286315 | Mus musculus hypothetical gene supported by AK049252 (LOC327897), mRNA. | | 2.6 | SM2128 | c | 4 |
| 092308m1_V2MM_72432 | 1.526423 | 2.205692 | TRUE | HP_29429 | TGCTGTTG | CTGGTCAC | NM_173433 | hypothetical protein 4932416A15 | | 2.3 | SM2012 | a | 5 |
| mPool7_V2MM_186962 | 1.525513 | 3.004939 | TRUE | HP_40743 | TGCTGTTG | CTATTTGG | XM_289626 | Mus musculus LOC333388 (LOC333388), mRNA. | | 2.9 | SM2292 | h | 6 |
| mPool2_V2MM_105540 | 1.525092 | 3.004939 | TRUE | HP_32665 | TGCTGTTG | GGCTAACT | NM_177652 | ryanodine receptor 3 | Ryr3 | 2.7 | SM2159 | c | 7 |
| mPool2_V2MM_98480 | 1.524275 | 3.004939 | TRUE | HP_31970 | TGCTGTTG | CCGTGTTG | XM_112566 | Mus musculus LOC193371 (LOC193371), mRNA. | | 2.6 | SM2138 | d | 4 |
| mPool4_V2MM_236964 | 1.523556 | 75.83547 | TRUE | NA | NA | NA | NA | NA | NA | NA | NA | NA | NA |
| mPool2_V2MM_87388 | 1.521684 | 2.205692 | TRUE | HP_30884 | TGCTGTTG | GACTCTTC | NM_175153 | RIKEN cDNA 2010321M09 gene | 2010321M09Rik | 2.6 | SM2141 | b | 2 |
| mPool2_V2MM_150955 | 1.520861 | 1.959154 | TRUE | HP_37163 | TGCTGTTG | CTCTTGAA | XM_194334 | RIKEN cDNA 0710005I19 gene | 0710005I19Rik | 2.1 | SM2345 | f | 5 |
| 092308m1_V2MM_82790 | 1.519708 | 2.205692 | TRUE | HP_30435 | TGCTGTTG | CCAGGAGT | NM_010470 | heterochromatin protein 1, binding protein 3 | Hp1bp3 | 2.6 | SM2109 | b | 5 |
| mPool7_V2MM_194397 | 1.519333 | 3.004939 | TRUE | HP_41256 | TGCTGTTG | GAAATAAT | NM_175009 | RIKEN cDNA 1810057B09 gene | 1810057B09Rik | 2.9 | SM2265 | b | 2 |
| mPool7_V2MM_194397 | 1.519333 | 3.004939 | TRUE | HP_41256 | TGCTGTTG | GAAATAAT | NM_175009 | RIKEN cD 1810057B09 gene | 1810057B09Rik | 2.9 | SM2265 | b | 2 |
| mPool4_V2MM_203900 | 1.518837 | 35.99881 | TRUE | HP_42182 | TGCTGTTG | CACCAGTC | XM_138918 | Mus musculus similar to erythrocyte membrane protein 3 [Plasmodium falciparu | | 2.16 | SM2621 | d | 5 |
| 092308m3_V2MM_86162 | 1.517734 | 75.83547 | TRUE | HP_30763 | TGCTGTTG | CTGCTCAT | NM_146635 | olfactory receptor 1489 | Olfr1489 | 2.13 | SM2458 | d | 10 |
| 092308m3_V2MM_93062 | 1.517347 | 59.99801 | TRUE | HP_31439 | TGCTGTTG | CAGTGAG | NM_178897 | expressed sequence AW125391 | AW125391 | 2.9 | SM2262 | h | 7 |
| 092308m3_V2MM_93062 | 1.517347 | 59.99801 | TRUE | HP_31439 | TGCTGTTG | CAGTGAG | NM_178897 | expressed sequence AW125391 | AW125391 | 2.9 | SM2262 | h | 7 |
| mPool2_V2MM_14561 | 1.516491 | 2.205692 | TRUE | HP_23790 | TGCTGTTG | CCTGTAAA | NM_008328 | interferon activated gene 203 | Ifi203 | 2.11 | SM2388 | h | 11 |
| mPool2_V2MM_83438 | 1.515601 | 3.004939 | TRUE | HP_30498 | TGCTGTTG | GATCCTAT | NM_015767 | tocopherol (alpha) transfer protein | Ttpa | 2.6 | SM2135 | f | 4 |
| 092308m1_V2MM_144680 | 1.515213 | 2.205692 | TRUE | HP_36539 | TGCTGTTG | CATATTAT | XM_160198 | Mus musculus LOC213218 (LOC213218), mR. | | 2.4 | SM2047 | c | 6 |
| 092308m1_V2MM_172940 | 1.514759 | 3.004939 | TRUE | HP_39342 | TGCTGTTG | CAGCCCAT | XM_286926 | gene model 815, (NCBI) | Gm815 | 2.5 | SM2060 | d | 2 |
| 092308m1_V2MM_143833 | 1.514155 | 3.004939 | TRUE | HP_36454 | TGCTGTTG | CATGAACA | XM_159183 | Mus musculus LOC245402 (LOC245402), mR. | | 2.4 | SM2038 | f | 10 |
| 092308m1_V2MM_143833 | 1.514155 | 3.004939 | TRUE | HP_36454 | TGCTGTTG | CATGAACA | XM_159183 | Mus musculus LOC245402 (LOC245402), mRNA. | | 2.4 | SM2038 | f | 10 |
| 092308m1_V2MM_182065 | 1.513401 | 3.004939 | TRUE | HP_40253 | TGCTGTTG | CTCATCCG | XM_288781 | Mus musculus LOC332243 (LOC332243), mR. | | 2.5 | SM2068 | h | 6 |
| mPool2_V2MM_31752 | 1.512977 | 3.004939 | TRUE | HP_25464 | TGCTGTTG | GAAAGTA | NM_019708 | short coiled-coil protein | Scoc | 2.11 | SM2392 | d | 2 |
| mPool2_V2MM_131650 | 1.512862 | 2.205692 | TRUE | HP_35241 | TGCTGTTG | CATGTGTT | BC023150 | heterogeneous nuclear ribonucleoprotein A0 | Hnrpa0 | 2.11 | SM2373 | c | 7 |
| mPool2_V2MM_25174 | 1.512408 | 2.205692 | TRUE | HP_24824 | TGCTGTTG | CAATGTTA | NM_009194 | solute carrier family 12, member 2 | Slc12a2 | 2.7 | SM2168 | c | 1 |
| 092308m1_V2MM_155615 | 1.511965 | 2.205692 | TRUE | HP_37625 | TGCTGTTG | CAAGATC | XM_197644 | Mus musculus LOC271898 (LOC271898), mR. | | 2.6 | SM2115 | b | 3 |
| mPool7_V2MM_206173 | 1.511791 | 3.004939 | TRUE | HP_42401 | TGCTGTTG | GAATGTA | XM_128618 | similar to zinc finger protein 97 | | 2.1 | SM2317 | h | 4 |
| 092308m1_V2MM_172295 | 1.510556 | 2.205692 | TRUE | HP_39278 | TGCTGTTG | CATACTGC | XM_286795 | Mus musculus hypothetical gene supported by AK035334 (LOC328762), mR. | | 2.5 | SM2076 | b | 11 |
| mPool2_V2MM_5054 | 1.510332 | 2.205692 | TRUE | HP_22861 | TGCTGTTG | CAGTTAG | NM_008260 | forkhead box A3 | Foxa3 | 2.11 | SM2387 | a | 6 |
| mPool7_V2MM_176264 | 1.50943 | 3.004939 | TRUE | HP_39674 | TGCTGTTG | CATGCTAT | AK028992 | RIKEN cDNA 6230400G14 gene | 6230400G14Rik | 2.9 | SM2273 | g | 3 |
| 092308m3_V2MM_96516 | 1.508449 | 35.99881 | TRUE | HP_31776 | TGCTGTTG | CAGAGCAT | XM_110787 | UPF3 regulator of nonsense transcripts homolog B (yeast | Upf3b | 2.12 | SM2450 | g | 3 |
| 092308m3_V2MM_202732 | 1.50821 | 75.83547 | TRUE | HP_42072 | TGCTGTTG | CCATTATT | XM_138204 | Mus musculus similar to RIKEN cDNA 2810037C14 [Mus musculus] (LOC217805), | | 2.1 | SM2319 | b | 7 |
| 092308m3_V2MM_202732 | 1.50821 | 75.83547 | TRUE | HP_42072 | TGCTGTTG | CCATTATT | XM_138204 | Mus musculus similar to RIKEN cD 2810037C14 [Mus musculus] (LOC217805), mR | | 2.1 | SM2319 | b | 7 |
| mPool7_V2MM_203995 | 1.50764 | 3.004939 | TRUE | HP_42191 | TGCTGTTG | GCTCTAAC | BC050039 | RIKEN cDNA D330013L20 gene | D330013L20Rik | 2.1 | SM2303 | g | 8 |
| mPool2_V2MM_83891 | 1.50646 | 2.205692 | TRUE | HP_30541 | TGCTGTTG | GAGCTGT | NM_026112 | zinc finger protein 606 | Zfp606 | 2.6 | SM2134 | e | 5 |
| mPool6_V2MM_27137 | 1.505746 | 75.83547 | TRUE | HP_25016 | TGCTGTTG | CTGTGTAC | AK041928 | RIKEN cDNA A630047E20 gene | A630047E20Rik | 2.7 | SM2182 | a | 3 |
| mPool7_V2MM_214362 | 1.505284 | 2.205692 | TRUE | HP_30416 | TGCTGTTG | GACTTTAT | X60958 | CD80 antigen | Cd80 | 2.9 | SM2270 | a | 8 |
| 092308m3_V2MM_126924 | 1.505177 | 75.83547 | TRUE | HP_28195 | TGCTGTTG | GCTGCCCT | NM_011653 | tubulin, alpha 1 | Tuba1 | 2.6 | SM2111 | c | 1 |
| mPool7_V2MM_183717 | 1.50354 | 3.004939 | TRUE | HP_40419 | TGCTGTTG | CAGAGACT | XM_289068 | Mus musculus LOC333518 (LOC333518), mRNA. | | 2.5 | SM2070 | d | 9 |
| mPool7_V2MM_183717 | 1.50354 | 3.004939 | TRUE | HP_40419 | TGCTGTTG | CAGAGACT | XM_289068 | Mus musculus LOC333518 (LOC333518), mR. | | 2.5 | SM2070 | d | 9 |
| 092308m3_V2MM_101003 | 1.503375 | 75.83547 | TRUE | HP_32217 | TGCTGTTG | GGTGTAT | XM_488906 | LOC433341 | | 2.13 | SM2490 | c | 8 |
| mPool7_V2MM_85669 | 1.503228 | 2.205692 | TRUE | HP_30715 | TGCTGTTG | CTGTATAT | NM_146438 | olfactory receptor 938 | Olfr938 | 2.9 | SM2267 | h | 4 |
| mPool7_V2MM_85669 | 1.503228 | 2.205692 | TRUE | HP_30715 | TGCTGTTG | CTGTATAT | NM_146438 | olfactory receptor 938 | Olfr938 | 2.9 | SM2267 | h | 4 |

| | | | | | | | | | | | | | | | |
|---|---|---|---|---|---|---|---|---|---|---|---|---|---|---|---|
| mPool2_V2MM_132452 | 1.501643 | 3.004939 | TRUE | HP_35320 | TGCTGTTG | CCAACTGA | XM_489556 | similar to hypothetical protein | | | 2.11 | SM2362 | b | | 4 |
| mPool2_V2MM_107065 | 1.501547 | 2.205692 | TRUE | HP_32815 | TGCTGTTG | CGCCCATG | NM_198610 | cDNA sequence BC055811 | BC055811 | | 2.1 | SM2323 | a | | 9 |
| mPool2_V2MM_34221 | 1.501173 | 2.205692 | TRUE | HP_25704 | TGCTGTTG | CCATACAG | NM_019720 | cytochrome b-561 domain containing 2 | Cyb561d2 | | 2.11 | SM2390 | h | | 5 |
| 092308m1_V2MM_127942 | 1.501066 | 3.004939 | TRUE | HP_34876 | TGCTGTTG | CAGTGATT | XM_145466 | similar to cytochrome P450 CYP2B21 | | | 2.6 | SM2104 | g | | 2 |
| mPool6_V2MM_79262 | 1.500817 | 62.64426 | TRUE | HP_30092 | TGCTGTTG | GATGTGAG | AF285588 | sal-like 4 (Drosophila) | Sall4 | | 2.8 | SM2218 | d | | 9 |
| mPool7_V2MM_104355 | 1.499271 | 3.004939 | TRUE | HP_32548 | TGCTGTTG | CCCTATTG | AK077997 | RIKEN cDNA 6130401J04 gene | 6130401J04Rik | | 2.6 | SM2149 | e | | 1 |
| mPool2_V2MM_131836 | 1.498981 | 3.004939 | TRUE | HP_35259 | TGCTGTTG | GAAGAAG | AK018473 | RIKEN cDNA 8430437N05 gene | 8430437N05Rik | | 2.11 | SM2357 | c | | 2 |
| mPool2_V2MM_89062 | 1.497919 | 3.004939 | TRUE | HP_31049 | TGCTGTTG | GGTATTGC | AK031726 | RIKEN cDNA C130073D16 gene | C130073D16Rik | | 2.6 | SM2139 | b | | 4 |
| mPool7_V2MM_106117 | 1.497455 | 2.205692 | TRUE | HP_32722 | TGCTGTTG | CTTTATGA | BC064474 | cingulin | Cgn | | 2.1 | SM2315 | g | | 5 |
| mPool7_V2MM_109915 | 1.497161 | 2.205692 | TRUE | HP_33096 | TGCTGTTG | CTGGATCA | AK016466 | RIKEN cDNA 4931420C21 gene | 4931420C21Rik | | 2.9 | SM2299 | g | | 7 |
| mPool2_V2MM_92734 | 1.496458 | 3.004939 | TRUE | HP_31407 | TGCTGTTG | GCTCCTTA | NM_178764 | RIKEN cDNA B930006L02 gene | B930006L02Rik | | 2.6 | SM2147 | g | | 3 |
| mPool7_V2MM_70831 | 1.495737 | 3.004939 | TRUE | HP_29273 | TGCTGTTG | CAGATGGA | NM_026055 | ribosomal protein L39 | Rpl39 | | 2.8 | SM2237 | c | | 6 |
| mPool2_V2MM_127131 | 1.495649 | 2.205692 | TRUE | HP_34796 | TGCTGTTG | GCCTCAAC | XM_145005 | IQ motif and Sec7 domain 3 | Iqsec3 | | 2.11 | SM2361 | c | | 5 |
| mPool4_V2MM_16927 | 1.494767 | 61.29828 | TRUE | HP_24021 | TGCTGTTG | GACTTTGT | NM_030597 | LSM2 homolog, U6 small nuclear RNA associated (S. cere | Lsm2 | | 2.15 | SM2577 | a | | 2 |
| mPool2_V2MM_19545 | 1.494639 | 2.205692 | TRUE | HP_24276 | TGCTGTTG | GGTGTAAA | NM_134190 | vomeronasal 1 receptor, E1 | V1re1 | | 2.11 | SM2374 | f | | 11 |
| mPool7_V2MM_182988 | 1.494619 | 3.004939 | TRUE | HP_40346 | TGCTGTTG | CAGTCTGT | XM_288940 | Mus musculus LOC332580 (LOC332580), mRNA. | | | 2.9 | SM2285 | f | | 5 |
| 092308m1_V2MM_169424 | 1.494514 | 3.004939 | TRUE | HP_38991 | TGCTGTTG | CCCTGAAC | AK039487 | RIKEN cD A330049N07 gene | A330049N07Rik | | 2.6 | SM2122 | e | | 3 |
| mPool2_V2MM_100395 | 1.494109 | 2.205692 | TRUE | HP_32158 | TGCTGTTG | CTATTTCT | XM_126847 | kelch repeat and BTB (POZ) domain containing 9 | Kbtbd9 | | 2.7 | SM2163 | e | | 1 |
| mPool2_V2MM_115091 | 1.493748 | 3.004939 | TRUE | NA | NA | NA | NA | NA | NA | NA | | NA | NA | NA | |
| 092308m1_V2MM_68318 | 1.493576 | 2.205692 | TRUE | HP_29028 | TGCTGTTG | CTTGGCAT | NM_013737 | phospholipase A2, group VII (platelet-activating factor ac | Pla2g7 | | 2.4 | SM2020 | h | | 2 |
| mPool2_V2MM_8298 | 1.493277 | 2.205692 | TRUE | HP_23178 | TGCTGTTG | GTATAAAC | XM_122407 | gene model 1818, (NCBI) | Gm1818 | | 2.11 | SM2383 | e | | 12 |
| mPool6_V2MM_154711 | 1.493077 | 35.99881 | TRUE | HP_37535 | TGCTGTTG | CAACGAGA | XM_196979 | Mus musculus LOC270985 (LOC270985), mRNA. | | | 2.7 | SM2197 | d | | 10 |
| mPool5_V2MM_182203 | 1.492862 | 62.64426 | TRUE | HP_40267 | TGCTGTTG | CTGCTGGC | XM_288804 | Mus musculus LOC332255 (LOC332255), mRNA. | | NA | | | | NA | |
| mPool2_V2MM_143911 | 1.492824 | 3.004939 | TRUE | HP_36462 | TGCTGTTG | CAGCTATA | XM_159415 | Mus musculus LOC245652 (LOC245652), mRNA. | | | 2.5 | SM2054 | f | | 5 |
| mPool2_V2MM_143911 | 1.492824 | 3.004939 | TRUE | HP_36462 | TGCTGTTG | CAGCTATA | XM_159415 | Mus musculus LOC245652 (LOC245652), mR. | | | 2.5 | SM2054 | f | | 5 |
| mPool2_V2MM_123086 | 1.492113 | 2.205692 | TRUE | HP_34394 | TGCTGTTG | GGGTGAA | XM_142640 | serine (or cysteine) proteinase inhibitor, clade B (ovalbur | Serpinb3b | | 2.6 | SM2104 | a | | 5 |
| mPool2_V2MM_123086 | 1.492113 | 2.205692 | TRUE | HP_34394 | TGCTGTTG | GGGTGAA | XM_142640 | serine (or cysteine) proteise inhibitor, clade B (ovalbumir | Serpinb3b | | 2.6 | SM2104 | a | | 5 |
| mPool7_V2MM_203153 | 1.491948 | 3.004939 | TRUE | HP_42111 | TGCTGTTG | GCTCTTAA | AK021149 | potassium voltage-gated channel, subfamily H (eag-relat | Kcnh3 | | 2.9 | SM2297 | d | | 9 |
| mPool2_V2MM_134899 | 1.491897 | 3.004939 | TRUE | HP_35562 | TGCTGTTG | GATACCTC | XM_486209 | similar to sc328 | | | 2.11 | SM2374 | g | | 1 |
| mPool7_V2MM_68429 | 1.490868 | 3.004939 | TRUE | HP_29038 | TGCTGTTG | GAAATTCT | AK046834 | RIKEN cDNA A230065J02 gene | A230065J02Rik | | 2.8 | SM2244 | b | | 6 |
| mPool2_V2MM_166417 | 1.490852 | 3.004939 | TRUE | HP_38694 | TGCTGTTG | CCCAACAT | XM_285495 | Mus musculus similar to TRAF6-inhibitory zinc finger protein; TRAF6-binding zinc | | | 2.6 | SM2132 | e | | 2 |
| mPool2_V2MM_104372 | 1.490102 | 3.004939 | TRUE | HP_32550 | TGCTGTTG | CTCCGATC | XM_129726 | RIKEN cDNA 2410118I19 gene | 2410118I19Rik | | 2.7 | SM2163 | g | | 2 |
| mPool4_V2MM_110485 | 1.488575 | 60.13437 | TRUE | HP_33151 | TGCTGTTG | CCCATTAT | AK014852 | RIKEN cDNA 4921509F24 gene | 4921509F24Rik | | 2.16 | SM2612 | h | | 2 |
| 092308m1_V2MM_78092 | 1.488531 | 3.004939 | TRUE | HP_29979 | TGCTGTTG | GAAGTTAA | NM_172817 | zinc finger protein 647 | Zfp647 | | 2.4 | SM2022 | c | | 5 |
| 092308m3_V2MM_46522 | 1.487615 | 75.83547 | TRUE | HP_10524 | TGCTGTTG | CCCTGTGT | XM_484805 | similar to Cytochrome P450, family 2, subfamily c, polypeptide 40 | | | 2.12 | SM2416 | e | | 7 |
| 092308m1_V2MM_67026 | 1.487005 | 3.004939 | TRUE | HP_28902 | TGCTGTTG | CCCATGCC | NM_053085 | transcription factor 23 | Tcf23 | | 2.3 | SM2016 | h | | 7 |
| 092308m3_V2MM_33765 | 1.485245 | 75.83547 | TRUE | HP_25660 | TGCTGTTG | CAGTGGA | NM_133658 | excision repair cross-complementing rodent repair defici | Ercc3 | | 2.12 | SM2432 | d | | 6 |
| mPool2_V2MM_118795 | 1.484785 | 3.004939 | TRUE | HP_33972 | TGCTGTTG | CTAATGAA | XM_140071 | Mus musculus similar to protease [Mus musculus] (LOC215435), mRNA. | | | 2.11 | SM2368 | b | | 8 |
| mPool4_V2MM_124214 | 1.484565 | 75.83547 | TRUE | HP_34506 | TGCTGTTG | CTCGTGGT | XM_143348 | Mus musculus similar to protease [Mus musculus] (LOC213288), mRNA. | | | 2.14 | SM2532 | c | | 9 |
| 092308m3_V2MM_34522 | 1.484526 | 59.99801 | TRUE | HP_25734 | TGCTGTTG | CCCGGAGC | NM_153407 | cD sequence BC035295 | BC035295 | | 2.12 | SM2427 | e | | 10 |
| mPool2_V2MM_109764 | 1.484005 | 2.205692 | TRUE | HP_33081 | TGCTGTTG | CTGATTGA | AK020544 | RIKEN cDNA 9530004P13 gene | 9530004P13Rik | | 2.6 | SM2148 | g | | 12 |
| 092308m3_V2MM_23256 | 1.483568 | 75.83547 | TRUE | HP_24639 | TGCTGTTG | CTGCTGCT | NM_009295 | syntaxin binding protein 1 | Stxbp1 | | 2.12 | SM2426 | b | | 5 |
| mPool2_V2MM_165536 | 1.483557 | 3.004939 | TRUE | HP_38607 | TGCTGTTG | CTGAGGA | XM_285270 | Mus musculus similar to RIKEN cDNA 4932409F11 [Mus musculus] (LOC328896), | | | 2.6 | SM2130 | b | | 9 |
| mPool2_V2MM_105600 | 1.482355 | 2.205692 | TRUE | HP_32671 | TGCTGTTG | GGGATGA | XM_130642 | RIKEN cDNA 5430405G05 gene | 5430405G05Rik | | 2.6 | SM2147 | a | | 9 |
| mPool7_V2MM_93732 | 1.482234 | 3.004939 | TRUE | HP_31503 | TGCTGTTG | CGAAATGA | NM_181748 | G protein-coupled receptor 120 | Gpr120 | | 2.9 | SM2263 | e | | 11 |
| mPool7_V2MM_93732 | 1.482234 | 3.004939 | TRUE | HP_31503 | TGCTGTTG | CGAAATGA | NM_181748 | G protein-coupled receptor 120 | Gpr120 | | 2.9 | SM2263 | e | | 11 |
| mPool7_V2MM_177109 | 1.482223 | 3.004939 | TRUE | HP_39758 | TGCTGTTG | GTGTAAAT | XM_287805 | Mus musculus hypothetical gene supported by AK032925 (LOC330762), mRNA. | | | 2.5 | SM2063 | h | | 6 |
| mPool7_V2MM_177109 | 1.482223 | 3.004939 | TRUE | HP_39758 | TGCTGTTG | GTGTAAAT | XM_287805 | Mus musculus hypothetical gene supported by AK032925 (LOC330762), mR. | | | 2.5 | SM2063 | h | | 6 |
| 092308m3_V2MM_29314 | 1.482012 | 75.83547 | TRUE | HP_25227 | TGCTGTTG | CAAAGTTC | NM_146253 | zinc finger protein 482 | Zfp482 | | 2.12 | SM2425 | c | | 1 |
| mPool2_V2MM_118493 | 1.481935 | 3.004939 | TRUE | HP_33942 | TGCTGTTG | GATATGAA | XM_139876 | cDNA sequence D10628 | D10628 | | 2.11 | SM2359 | d | | 4 |
| 092308m1_V2MM_123086 | 1.481921 | 2.205692 | TRUE | HP_34394 | TGCTGTTG | GGGTGAA | XM_142640 | serine (or cysteine) proteinase inhibitor, clade B (ovalbur | Serpinb3b | | 2.6 | SM2104 | a | | 5 |
| 092308m1_V2MM_123086 | 1.481921 | 2.205692 | TRUE | HP_34394 | TGCTGTTG | GGGTGAA | XM_142640 | serine (or cysteine) proteise inhibitor, clade B (ovalbumir | Serpinb3b | | 2.6 | SM2104 | a | | 5 |
| mPool2_V2MM_109641 | 1.48147 | 3.004939 | TRUE | HP_33069 | TGCTGTTG | GACCTTGT | XM_134026 | RIKEN cDNA 2310051N18 gene | 2310051N18Rik | | 2.6 | SM2150 | a | | 12 |

| | | | | | | | | | | | | | | | |
|---|---|---|---|---|---|---|---|---|---|---|---|---|---|---|---|
| mPool7_V2MM_202438 | 1.481425 | 3.004939 | TRUE | HP_42044 | TGCTGTTG | CTGCTTAA | XM_485379 | similar to 2610030H06Rik protein | | | 2.1 | SM2306 | h | | 2 |
| 092308m1_V2MM_177029 | 1.481365 | 2.205692 | TRUE | HP_39750 | TGCTGTTG | CCATTGGA | XM_287788 | Mus musculus hypothetical gene supported by AK036103 (LOC330739), mRNA. | | | 2.5 | SM2081 | a | | 1 |
| 092308m1_V2MM_177029 | 1.481365 | 2.205692 | TRUE | HP_39750 | TGCTGTTG | CCATTGGA | XM_287788 | Mus musculus hypothetical gene supported by AK036103 (LOC330739), mR. | | | 2.5 | SM2081 | a | | 1 |
| mPool2_V2MM_111245 | 1.481108 | 3.004939 | TRUE | HP_33226 | TGCTGTTG | CTGATGAT | AK015052 | RIKEN cDNA 4930402K13 gene | 4930402K13Rik | | 2.7 | SM2160 | h | | 10 |
| mPool7_V2MM_207952 | 1.481019 | 3.004939 | TRUE | HP_42574 | TGCTGTTG | GTCCAAGT | XM_162924 | Mus musculus LOC243703 (LOC243703), mRNA. | | | 2.1 | SM2331 | c | | 3 |
| mPool7_V2MM_116587 | 1.480241 | 3.004939 | TRUE | HP_33753 | TGCTGTTG | GATGCCTT | XM_138996 | Mus musculus LOC239041 (LOC239041), mRNA. | | | 2.1 | SM2303 | h | | 8 |
| 092308m1_V2MM_153117 | 1.479187 | 3.004939 | TRUE | HP_37377 | TGCTGTTG | GGTGGTAT | XM_195745 | Mus musculus similar to Rse H, putative [Plasmodium yoelii yoelii] (LOC270119), | | | 2.6 | SM2125 | a | | 2 |
| mPool2_V2MM_107262 | 1.479027 | 3.004939 | TRUE | HP_32835 | TGCTGTTG | CCAATGAC | NM_027855 | RIKEN cDNA 0610007C21 gene | 0610007C21Rik | | 2.1 | SM2323 | d | | 10 |
| 092308m1_V2MM_136532 | 1.478567 | 3.004939 | TRUE | HP_35725 | TGCTGTTG | CAAATGAA | XM_151813 | Mus musculus LOC213257 (LOC213257), mR. | | | 2.4 | SM2039 | d | | 6 |
| 092308m1_V2MM_176912 | 1.477147 | 3.004939 | TRUE | HP_39739 | TGCTGTTG | CTGCTTAT | XM_287767 | Mus musculus hypothetical gene supported by AK038473 (LOC330714), mR. | | | 2.5 | SM2066 | f | | 1 |
| 092308m1_V2MM_80619 | 1.476967 | 3.004939 | TRUE | HP_30224 | TGCTGTTG | GGAATTGT | NM_025692 | ubiquitin-activating enzyme E1-domain containing 1 | Ube1dc1 | | 2.4 | SM2027 | h | | 3 |
| mPool7_V2MM_68711 | 1.476635 | 2.205692 | TRUE | HP_29066 | TGCTGTTG | GACTTTAC | NM_026173 | RIKEN cDNA 1200014M14 gene | 1200014M14Rik | | 2.8 | SM2227 | b | | 11 |
| 092308m3_V2MM_27443 | 1.476623 | 35.99881 | TRUE | HP_25046 | TGCTGTTG | GACACAAC | NM_030718 | ABO blood group (transferase A, alpha 1-3-N-acetylgalact | Abo | | 2.12 | SM2421 | a | | 4 |
| mPool2_V2MM_27473 | 1.476308 | 1.959154 | TRUE | HP_25049 | TGCTGTTG | GACAGATC | NM_133767 | mitochondrial translational initiation factor 2 | Mtif2 | | 2.11 | SM2389 | c | | 11 |
| mPool2_V2MM_215158 | 1.476302 | 3.004939 | TRUE | HP_43271 | TGCTGTTG | CTGGAATA | XM_146473 | Mus musculus similar to nuclear autoantigenic sperm protein [Mus musculus] (LC | | | 2.11 | SM2366 | f | | 11 |
| 092308m1_V2MM_168196 | 1.476272 | 3.004939 | TRUE | HP_38870 | TGCTGTTG | CCTCCTGT | XM_485762 | similar to hypothetical protein C230069C04 | | | 2.6 | SM2125 | a | | 11 |
| mPool4_V2MM_139312 | 1.47614 | 62.64426 | TRUE | HP_36003 | TGCTGTTG | GACTTTGA | XM_154921 | Mus musculus LOC219168 (LOC219168), mRNA. | | | 2.16 | SM2635 | a | | 3 |
| mPool7_V2MM_98281 | 1.475935 | 3.004939 | TRUE | HP_31951 | TGCTGTTG | CAGGGAA | XM_112336 | Mus musculus LOC195594 (LOC195594), mRNA. | | | 2.9 | SM2270 | g | | 1 |
| mPool2_V2MM_117768 | 1.475755 | 2.205692 | TRUE | NA | NA | NA | NA | NA | NA | NA | | NA | NA | NA | |
| 092308m1_V2MM_166043 | 1.47574 | 2.205692 | TRUE | HP_38657 | TGCTGTTG | CCTCGGAC | XM_285397 | Mus musculus similar to KIAA1856 protein [Homo sapiens] (LOC329487), mR. | | | 2.6 | SM2115 | b | | 8 |
| mPool7_V2MM_65246 | 1.47525 | 2.205692 | TRUE | HP_28728 | TGCTGTTG | GCATATAT | AK085850 | RIKEN cDNA D830016O14 gene | D830016O14Rik | | 2.8 | SM2242 | h | | 7 |
| mPool2_V2MM_212572 | 1.475089 | 3.004939 | TRUE | HP_43023 | TGCTGTTG | CTAACTAA | XM_140908 | ribosomal protein S27a | Rps27a | | 2.11 | SM2369 | d | | 5 |
| mPool2_V2MM_127574 | 1.474393 | 2.205692 | TRUE | HP_34839 | TGCTGTTG | CTATTATG | XM_145254 | RIKEN cDNA A630082K20 gene | A630082K20Rik | | 2.11 | SM2354 | b | | 3 |
| mPool7_V2MM_68759 | 1.474098 | 3.004939 | TRUE | HP_29070 | TGCTGTTG | GAGAGAA | NM_053117 | par-6 partitioning defective 6 homolog gamma (C. elegar | Pard6g | | 2.8 | SM2231 | c | | 6 |
| mPool2_V2MM_189240 | 1.472754 | 2.205692 | TRUE | HP_25600 | TGCTGTTG | CACATTTC | NM_146062 | periphilin 1 | Pphln1 | | 2.11 | SM2389 | h | | 8 |
| mPool2_V2MM_158160 | 1.47241 | 2.205692 | TRUE | HP_37877 | TGCTGTTG | GTGGTTAA | XM_206274 | Mus musculus LOC278520 (LOC278520), mRNA. | | | 2.6 | SM2132 | d | | 8 |
| 092308m3_V2MM_225744 | 1.471041 | 75.83547 | TRUE | HP_47778 | TGCTGTTG | GTGACTGT | NM_181753 | opsin 5 | Opn5 | | 2.13 | SM2468 | b | | 11 |
| 092308m3_V2MM_233230 | 1.470003 | 75.83547 | TRUE | HP_48514 | TGCTGTTG | GCTTGTAA | XM_484981 | D segment, Chr 2, Brigham & Womens Genetics 1335 exp | D2Bwg1335e | | 2.13 | SM2486 | e | | 8 |
| 092308m1_V2MM_154518 | 1.469995 | 3.004939 | TRUE | HP_37515 | TGCTGTTG | CTCGTATT | XM_196806 | Mus musculus LOC270769 (LOC270769), mR. | | | 2.6 | SM2116 | b | | 11 |
| mPool7_V2MM_147548 | 1.469821 | 3.004939 | TRUE | HP_36825 | TGCTGTTG | CCACGTCA | XM_163078 | Mus musculus LOC243797 (LOC243797), mRNA. | | | 2.5 | SM2052 | c | | 6 |
| mPool7_V2MM_147548 | 1.469821 | 3.004939 | TRUE | HP_36825 | TGCTGTTG | CCACGTCA | XM_163078 | Mus musculus LOC243797 (LOC243797), mR. | | | 2.5 | SM2052 | c | | 6 |
| 092308m3_V2MM_205360 | 1.469072 | 75.83547 | TRUE | HP_42322 | TGCTGTTG | CCGAGAAC | XM_132788 | similar to KIAA0218 | | | 2.13 | SM2489 | c | | 11 |
| 092308m3_V2MM_100381 | 1.467128 | 62.64426 | TRUE | HP_32157 | TGCTGTTG | GCCTTCCC | AK006597 | RIKEN cD 1700034J04 gene | 1700034J04Rik | | 2.14 | SM2501 | a | | 1 |
| 092308m3_V2MM_98617 | 1.466784 | 75.83547 | TRUE | HP_31984 | TGCTGTTG | CAGATGCC | XM_112797 | Mus musculus LOC215587 (LOC215587), mR. | | | 2.9 | SM2270 | d | | 3 |
| 092308m3_V2MM_98617 | 1.466784 | 75.83547 | TRUE | HP_31984 | TGCTGTTG | CAGATGCC | XM_112797 | Mus musculus LOC215587 (LOC215587), mRNA. | | | 2.9 | SM2270 | d | | 3 |
| mPool7_V2MM_103607 | 1.466715 | 2.205692 | TRUE | HP_32474 | TGCTGTTG | CCTGTATT | XM_129159 | RIKEN cDNA 1810073H04 gene | 1810073H04Rik | | 2.1 | SM2303 | e | | 4 |
| mPool2_V2MM_22454 | 1.466102 | 3.004939 | TRUE | HP_24560 | TGCTGTTG | CGTATTAT | NM_009285 | stanniocalcin 1 | Stc1 | | 2.7 | SM2165 | g | | 9 |
| mPool2_V2MM_115703 | 1.466051 | 3.004939 | TRUE | HP_33666 | TGCTGTTG | GACAGACA | XM_138607 | similar to hypothetical protein FLJ30829; RNA-binding region (RNP1, RRM) contai | | | 2.7 | SM2155 | h | | 10 |
| mPool2_V2MM_109890 | 1.465927 | 3.004939 | TRUE | HP_33093 | TGCTGTTG | GAAACATC | XM_486116 | cDNA sequence D10627 | D10627 | | 2.7 | SM2163 | h | | 11 |
| mPool7_V2MM_100109 | 1.464743 | 3.004939 | TRUE | HP_32130 | TGCTGTTG | CATTTCTA | NM_009546 | tripartite motif protein 25 | Trim25 | | 2.7 | SM2158 | g | | 10 |
| mPool7_V2MM_85492 | 1.46439 | 3.004939 | TRUE | HP_30697 | TGCTGTTG | CAATACTC | NM_146395 | olfactory receptor 1276 | Olfr1276 | | 2.8 | SM2249 | a | | 10 |
| mPool7_V2MM_177132 | 1.464243 | 2.205692 | TRUE | HP_39761 | TGCTGTTG | CGAATTTC | XM_287810 | Mus musculus hypothetical gene supported by AK049065 (LOC330779), mR. | | | 2.5 | SM2087 | d | | 7 |
| mPool7_V2MM_177132 | 1.464243 | 2.205692 | TRUE | HP_39761 | TGCTGTTG | CGAATTTC | XM_287810 | Mus musculus hypothetical gene supported by AK049065 (LOC330779), mRNA. | | | 2.5 | SM2087 | d | | 7 |
| 092308m1_V2MM_126100 | 1.463406 | 3.004939 | TRUE | HP_34694 | TGCTGTTG | GTGGCAA | XM_144377 | Mus musculus similar to crumbs homolog 1 [Mus musculus] (LOC231351), mR. | | | 2.6 | SM2103 | a | | 4 |
| mPool2_V2MM_116096 | 1.463322 | 2.205692 | TRUE | HP_33705 | TGCTGTTG | CTGATATT | XM_138748 | Mus musculus similar to envelope protein [Ovis aries] (LOC218362), mRNA. | | | 2.6 | SM2150 | c | | 2 |
| mPool4_V2MM_21536 | 1.462926 | 61.29828 | TRUE | HP_24470 | TGCTGTTG | CAGTTCTG | NM_148931 | solute carrier family 6 (neurotransmitter transporter, gly | Slc6a5 | | 2.15 | SM2599 | e | | 2 |
| 092308m1_V2MM_181542 | 1.462457 | 3.004939 | TRUE | HP_40201 | TGCTGTTG | CGAAGGA | XM_288693 | Mus musculus LOC332121 (LOC332121), mR. | | | 2.5 | SM2060 | d | | 4 |
| 092308m1_V2MM_183622 | 1.460482 | 3.004939 | TRUE | HP_40409 | TGCTGTTG | CTTTAAGA | XM_289051 | similar to bM64F17.3 (putative novel protein) | | | 2.5 | SM2069 | e | | 2 |
| mPool4_V2MM_213671 | 1.459972 | 60.13437 | TRUE | HP_43129 | TGCTGTTG | CAATAGTC | XM_140140 | Mus musculus similar to uracil DNA glycosylase [Homo sapiens] (LOC225099), mF | | | 2.14 | SM2544 | f | | 2 |
| 092308m1_V2MM_149043 | 1.459215 | 3.004939 | TRUE | HP_36975 | TGCTGTTG | CTGCCTGT | XM_164432 | Mus musculus LOC244522 (LOC244522), mR. | | | 2.5 | SM2057 | f | | 12 |
| mPool2_V2MM_123300 | 1.459045 | 2.205692 | TRUE | HP_34415 | TGCTGTTG | GATAAGAC | XM_142871 | Mus musculus similar to hypothetical protein FLJ90396 [Homo sapiens] (LOC2363 | | | 2.11 | SM2352 | f | | 6 |
| 092308m1_V2MM_177871 | 1.456781 | 2.205692 | TRUE | NA | NA | NA | NA | NA | NA | NA | | NA | NA | NA | |
| 092308m1_V2MM_170974 | 1.456765 | 2.205692 | TRUE | HP_39146 | TGCTGTTG | CCTTTACT | XM_286529 | gene model 1576, (NCBI) | Gm1576 | | 2.5 | SM2093 | g | | 11 |

| | | | | | | | | | | | | | | |
|---|---|---|---|---|---|---|---|---|---|---|---|---|---|---|
| 092308m1_V2MM_170974 | 1.456765 | 2.205692 | TRUE | HP_39146 | TGCTGTTG | CCTTTACT | XM_286529 | gene model 1576, (NCBI) | Gm1576 | 2.5 | SM2093 | g | | 11 |
| mPool7_V2MM_76262 | 1.456687 | 3.004939 | TRUE | HP_29802 | TGCTGTTG | CGATCATT | NM_025422 | RIKEN cDNA 1110055L24 gene | 1110055L24Rik | 2.8 | SM2227 | g | | 1 |
| mPool2_V2MM_107397 | 1.45601 | 3.004939 | TRUE | HP_32848 | TGCTGTTG | GCTCTAGT | AK075654 | RIKEN cDNA 3830408G10 gene | 3830408G10Rik | 2.7 | SM2156 | e | | 7 |
| mPool2_V2MM_145046 | 1.455564 | 3.004939 | TRUE | HP_36575 | TGCTGTTG | CTGGTATT | XM_160513 | Mus musculus LOC215240 (LOC215240), mRNA. | | 2.1 | SM2340 | e | | 8 |
| 092308m1_V2MM_92238 | 1.455531 | 3.004939 | TRUE | HP_31359 | TGCTGTTG | CTAACTAC | NM_178674 | F-box and leucine-rich repeat protein 21 | Fbxl21 | 2.6 | SM2112 | b | | 12 |
| 092308m1_V2MM_131488 | 1.455522 | 2.205692 | TRUE | HP_35225 | TGCTGTTG | CTAGGAG | AK020496 | RIKEN cD 9430078K24 gene | 9430078K24Rik | 2.5 | SM2097 | d | | 5 |
| 092308m1_V2MM_176622 | 1.454717 | 2.205692 | TRUE | HP_39710 | TGCTGTTG | CCTGTATT | XM_287707 | Mus musculus hypothetical gene supported by AK045013 (LOC330567), mR. | | 2.5 | SM2065 | a | | 6 |
| 092308m3_V2MM_202739 | 1.452713 | 62.64426 | TRUE | HP_42072 | TGCTGTTG | CCACAGA | XM_132795 | gene model 154, (NCBI) | Gm154 | 2.13 | SM2494 | e | | 1 |
| mPool2_V2MM_83636 | 1.452372 | 3.004939 | TRUE | HP_30516 | TGCTGTTG | CTCATTGA | NM_023127 | polymerase (RNA) II (DNA directed) polypeptide K | Polr2k | 2.6 | SM2140 | e | | 7 |
| mPool7_V2MM_173969 | 1.451997 | 3.004939 | TRUE | HP_39445 | TGCTGTTG | CCTAGTGT | XM_287126 | Mus musculus hypothetical gene supported by AK082531 (LOC329411), mRNA. | | 2.9 | SM2278 | f | | 1 |
| mPool2_V2MM_42767 | 1.451741 | 3.004939 | TRUE | HP_26537 | TGCTGTTG | CGCATTGC | NM_009692 | apolipoprotein A-I | Apoa1 | 2.7 | SM2165 | g | | 12 |
| mPool2_V2MM_14313 | 1.451211 | 3.004939 | TRUE | HP_23766 | TGCTGTTG | CCGCTTCT | NM_019394 | melanoma inhibitory activity 1 | Mia1 | 2.11 | SM2381 | h | | 12 |
| 092308m1_V2MM_173934 | 1.450953 | 2.205692 | TRUE | HP_39442 | TGCTGTTG | CTATATCA | XM_287120 | Mus musculus hypothetical gene supported by AK034841 (LOC329386), mR. | | 2.5 | SM2069 | e | | 7 |
| 092308m1_V2MM_173934 | 1.450953 | 2.205692 | TRUE | HP_39442 | TGCTGTTG | CTATATCA | XM_287120 | Mus musculus hypothetical gene supported by AK034841 (LOC329386), mRNA. | | 2.5 | SM2069 | e | | 7 |
| mPool6_V2MM_73675 | 1.450365 | 75.83547 | TRUE | HP_12575 | TGCTGTTG | GGTGTTCA | NM_025739 | RIKEN cDNA 4931406I20 gene | 4931406I20Rik | 2.7 | SM2173 | b | | 4 |
| mPool4_V2MM_37040 | 1.449677 | 75.83547 | TRUE | HP_25981 | TGCTGTTG | GAATCTGT | NM_031385 | testis expressed gene 18 | Tex18 | 2.15 | SM2579 | g | | 8 |
| 092308m1_V2MM_63029 | 1.448431 | 3.004939 | TRUE | HP_28511 | TGCTGTTG | CCGGATGT | NM_030237 | spermatogenic Zip 1 | | 2.3 | SM2015 | h | | 6 |
| 092308m1_V2MM_151907 | 1.448138 | 3.004939 | TRUE | HP_37257 | TGCTGTTG | GGCGTTTA | XM_194993 | olfactory receptor 171 | Olfr171 | 2.4 | SM2040 | b | | 5 |
| mPool7_V2MM_95356 | 1.447855 | 3.004939 | TRUE | HP_31663 | TGCTGTTG | CTGTTTGT | NM_197985 | adiponectin receptor 2 | Adipor2 | 2.9 | SM2255 | a | | 10 |
| mPool5_V2MM_7657 | 1.447542 | 75.83547 | TRUE | HP_23116 | TGCTGTTG | GCCACAGT | NM_008170 | glutamate receptor, ionotropic, NMDA2A (epsilon 1) | Grin2a | NA | | | NA | |
| mPool2_V2MM_21718 | 1.447502 | 2.205692 | TRUE | HP_24488 | TGCTGTTG | CATTGAGC | NM_027564 | RIKEN cDNA 4921507P07 gene | 4921507P07Rik | 2.11 | SM2387 | g | | 9 |
| 092308m1_V2MM_125765 | 1.447302 | 3.004939 | TRUE | HP_34660 | TGCTGTTG | CCAGTCTA | XM_144221 | similar to hypothetical protein MGC19022 | | 2.6 | SM2111 | g | | 1 |
| 092308m1_V2MM_125765 | 1.447302 | 3.004939 | TRUE | HP_34660 | TGCTGTTG | CCAGTCTA | XM_144221 | similar to hypothetical protein MGC19022 | | 2.6 | SM2111 | g | | 1 |
| mPool6_V2MM_77610 | 1.44722 | 60.13437 | TRUE | HP_29932 | TGCTGTTG | CTGTGTGA | NM_025482 | tumor protein D52-like 2 | Tpd52l2 | 2.8 | SM2222 | a | | 4 |
| 092308m3_V2MM_112366 | 1.447196 | 69.72441 | TRUE | HP_33337 | TGCTGTTG | CCTTTGAT | XM_136450 | Mus musculus similar to glyceraldehyde-3-phosphate dehydrogese [Rattus norve | | 2.13 | SM2497 | h | | 10 |
| mPool7_V2MM_104352 | 1.446502 | 3.004939 | TRUE | HP_32548 | TGCTGTTG | CCTATTGC | AK077997 | RIKEN cDNA 6130401J04 gene | 6130401J04Rik | 2.7 | SM2157 | e | | 2 |
| 092308m1_V2MM_138711 | 1.446331 | 3.004939 | TRUE | HP_35943 | TGCTGTTG | CGAAATAT | XM_154438 | Mus musculus LOC238835 (LOC238835), mRNA. | | 2.4 | SM2045 | d | | 6 |
| 092308m1_V2MM_138711 | 1.446331 | 3.004939 | TRUE | HP_35943 | TGCTGTTG | CGAAATAT | XM_154438 | Mus musculus LOC238835 (LOC238835), mR. | | 2.4 | SM2045 | d | | 6 |
| mPool7_V2MM_179050 | 1.446286 | 3.004939 | TRUE | HP_39952 | TGCTGTTG | CCAGTTTG | XM_288266 | Mus musculus LOC332539 (LOC332539), mRNA. | | 2.5 | SM2070 | b | | 6 |
| mPool7_V2MM_179050 | 1.446286 | 3.004939 | TRUE | HP_39952 | TGCTGTTG | CCAGTTTG | XM_288266 | Mus musculus LOC332539 (LOC332539), mR. | | 2.5 | SM2070 | b | | 6 |
| 092308m3_V2MM_149943 | 1.445488 | 75.83547 | TRUE | HP_37065 | TGCTGTTG | CCACGAAT | XM_165354 | Mus musculus LOC214929 (LOC214929), mR. | | 2.14 | SM2528 | h | | 11 |
| mPool2_V2MM_9944 | 1.445152 | 3.004939 | TRUE | HP_23338 | TGCTGTTG | CGCTTTAT | NM_011677 | uracil-DNA glycosylase | Ung | 2.11 | SM2379 | h | | 4 |
| mPool4_V2MM_262310 | 1.444862 | 75.83547 | TRUE | HP_52140 | TGCTGTTG | GTGTCTGT | NM_203507 | cDNA sequence BC016198 | BC016198 | 2.16 | SM2617 | h | | 3 |
| mPool2_V2MM_9907 | 1.443454 | 3.004939 | TRUE | HP_23335 | TGCTGTTG | CGCCGTAA | NM_011643 | transient receptor potential cation channel, subfamily C, | Trpc1 | 2.11 | SM2385 | a | | 4 |
| mPool4_V2MM_16028 | 1.443318 | 75.83547 | TRUE | HP_23933 | TGCTGTTG | CTGTGTGT | NM_198412 | DnaJ (Hsp40) homolog, subfamily C, member 6 | Dnajc6 | NA | | | NA | |
| mPool2_V2MM_106877 | 1.442866 | 3.004939 | TRUE | HP_32797 | TGCTGTTG | CATACATC | XM_131619 | cDNA sequence BC022150 | BC022150 | 2.7 | SM2159 | c | | 12 |
| 092308m1_V2MM_77283 | 1.442456 | 3.004939 | TRUE | HP_29900 | TGCTGTTG | CTGCCATC | NM_026015 | RIKEN cD 2610510L01 gene | 2610510L01Rik | 2.3 | SM2016 | b | | 8 |
| mPool2_V2MM_211496 | 1.441848 | 3.004939 | TRUE | HP_42919 | TGCTGTTG | GATGAGCT | XM_156070 | similar to stefin A2 | | 2.11 | SM2372 | b | | 7 |
| 092308m3_V2MM_226323 | 1.440703 | 75.83547 | TRUE | HP_47835 | TGCTGTTG | GGAGAAA | NM_175029 | APG4 (ATG4) autophagy-related homolog C (S. cerevisiae | Apg4c | 2.13 | SM2473 | g | | 2 |
| mPool5_V2MM_173749 | 1.440344 | 70.87591 | TRUE | HP_39423 | TGCTGTTG | CAGGCTGC | XM_287087 | Mus musculus hypothetical gene supported by AK083495 (LOC329349), mRNA. | | NA | | | NA | |
| 092308m3_V2MM_188570 | 1.440253 | 75.83547 | TRUE | HP_32192 | TGCTGTTG | CAAATTAC | NM_011344 | Sel1 (suppressor of lin-12) 1 homolog (C. elegans) | Sel1h | 2.7 | SM2168 | h | | 10 |
| 092308m3_V2MM_188570 | 1.440253 | 75.83547 | TRUE | HP_32192 | TGCTGTTG | CAAATTAC | NM_011344 | Sel1 (suppressor of lin-12) 1 homolog (C. elegans) | Sel1h | 2.7 | SM2168 | h | | 10 |
| mPool4_V2MM_239018 | 1.440179 | 60.13437 | TRUE | HP_49083 | TGCTGTTG | GTATATTT | XM_143027 | Mus musculus similar to envelope protein [Trichosurus vulpecula] (LOC236517), | | 2.14 | SM2548 | h | | 2 |
| 092308m1_V2MM_156660 | 1.439884 | 2.205692 | TRUE | HP_37728 | TGCTGTTG | CCACTTTC | AK013636 | RIKEN cD 2900042B11 gene | 2900042B11Rik | 2.6 | SM2117 | c | | 7 |
| mPool6_V2MM_76955 | 1.438621 | 75.83547 | TRUE | HP_29868 | TGCTGTTG | CTCGTGGC | NM_021791 | double C2, gamma | Doc2g | 2.7 | SM2172 | f | | 11 |
| mPool7_V2MM_177678 | 1.437435 | 2.205692 | TRUE | HP_39815 | TGCTGTTG | GGATTATT | XM_287927 | Mus musculus hypothetical gene supported by AK035601 (LOC330954), mRNA. | | 2.5 | SM2067 | h | | 11 |
| mPool7_V2MM_177678 | 1.437435 | 2.205692 | TRUE | HP_39815 | TGCTGTTG | GGATTATT | XM_287927 | Mus musculus hypothetical gene supported by AK035601 (LOC330954), mR. | | 2.5 | SM2067 | h | | 11 |
| mPool7_V2MM_171932 | 1.437425 | 3.004939 | TRUE | HP_39241 | TGCTGTTG | CCCTTAAC | XM_286725 | Mus musculus hypothetical gene supported by AK036009 (LOC328675), mR. | | 2.5 | SM2088 | f | | 9 |
| mPool7_V2MM_171932 | 1.437425 | 3.004939 | TRUE | HP_39241 | TGCTGTTG | CCCTTAAC | XM_286725 | Mus musculus hypothetical gene supported by AK036009 (LOC328675), mRNA. | | 2.5 | SM2088 | f | | 9 |
| mPool2_V2MM_8277 | 1.436967 | 3.004939 | TRUE | HP_23176 | TGCTGTTG | GTAAGTGT | NM_028060 | solute carrier family 35, member F2 | Slc35f2 | 2.11 | SM2386 | b | | 7 |
| 092308m3_V2MM_144130 | 1.436502 | 75.83547 | TRUE | HP_36484 | TGCTGTTG | CTTGGCCA | XM_159764 | Mus musculus LOC241881 (LOC241881), mR. | | 2.14 | SM2526 | e | | 3 |
| mPool4_V2MM_78137 | 1.436109 | 60.13437 | TRUE | HP_29983 | TGCTGTTG | GAATGGAT | NM_025504 | RIKEN cDNA 2310004L02 gene | 2310004L02Rik | 2.15 | SM2588 | f | | 11 |
| 092308m1_V2MM_117818 | 1.436038 | 3.004939 | TRUE | HP_33875 | TGCTGTTG | CAGAAAGT | XM_139540 | PDZ domain containing RING finger 4 | | 2.6 | SM2104 | d | | 1 |

| | | | | | | | | | | | | | |
|---|---|---|---|---|---|---|---|---|---|---|---|---|---|
| mPool2_V2MM_22530 | 1.435996 | 3.004939 | TRUE | HP_24568 | TGCTGTTG | CTAACCAT | NM_030705 | mesoderm development candidate 1 | Mesdc1 | 2.11 | SM2390 | c | 8 |
| 092308m1_V2MM_187836 | 1.43561 | 2.205692 | TRUE | HP_40830 | TGCTGTTG | CTGAAATT | XM_290161 | Mus musculus LOC333779 (LOC333779), mRNA. | | 2.5 | SM2094 | h | 2 |
| 092308m1_V2MM_187836 | 1.43561 | 2.205692 | TRUE | HP_40830 | TGCTGTTG | CTGAAATT | XM_290161 | Mus musculus LOC333779 (LOC333779), mR. | | 2.5 | SM2094 | h | 2 |
| mPool6_V2MM_193327 | 1.435352 | 75.83547 | TRUE | HP_41179 | TGCTGTTG | CCGTTCTG | NM_001006680 | potassium voltage-gated channel, subfamily Q, member | Kcnq2 | 2.7 | SM2180 | f | 3 |
| 092308m3_V2MM_55020 | 1.435206 | 66.95036 | TRUE | HP_27732 | TGCTGTTG | CCCAGGA | AK129138 | zinc finger protein 96 | Zfp96 | 2.12 | SM2446 | h | 4 |
| 092308m1_V2MM_176179 | 1.435037 | 2.205692 | TRUE | HP_39666 | TGCTGTTG | GGAAAGT | AK046665 | RIKEN cD 9530026P05 gene | 9530026P05Rik | 2.5 | SM2072 | f | 9 |
| mPool2_V2MM_3528 | 1.434641 | 3.004939 | TRUE | HP_22712 | TGCTGTTG | GATAGGA | NM_026742 | RIKEN cDNA 1110007M04 gene | 1110007M04Rik | 2.11 | SM2375 | f | 9 |
| mPool4_V2MM_213659 | 1.434521 | 66.95036 | TRUE | HP_43128 | TGCTGTTG | GGAGCCT | XM_147019 | Mus musculus similar to WD40-repeat type I transmembrane protein A72.5 [Mus | | 2.16 | SM2631 | d | 12 |
| 092308m3_V2MM_40374 | 1.43424 | 60.13437 | TRUE | HP_26304 | TGCTGTTG | GTCCCTGC | NM_009289 | STE20-like kise (yeast) | Slk | 2.12 | SM2424 | c | 10 |
| 092308m1_V2MM_137283 | 1.433549 | 3.004939 | TRUE | HP_35800 | TGCTGTTG | CAACCAAC | XM_152972 | Mus musculus LOC237973 (LOC237973), mRNA. | | 2.5 | SM2053 | f | 4 |
| 092308m1_V2MM_137283 | 1.433549 | 3.004939 | TRUE | HP_35800 | TGCTGTTG | CAACCAAC | XM_152972 | Mus musculus LOC237973 (LOC237973), mR. | | 2.5 | SM2053 | f | 4 |
| 092308m3_V2MM_104323 | 1.433405 | 75.83547 | TRUE | HP_32545 | TGCTGTTG | CTTACAAA | AK036315 | RIKEN cDNA 9630055N22 gene | 9630055N22Rik | 2.1 | SM2304 | a | 12 |
| 092308m3_V2MM_104323 | 1.433405 | 75.83547 | TRUE | HP_32545 | TGCTGTTG | CTTACAAA | AK036315 | RIKEN cD 9630055N22 gene | 9630055N22Rik | 2.1 | SM2304 | a | 12 |
| 092308m1_V2MM_147733 | 1.430171 | 3.004939 | TRUE | HP_36844 | TGCTGTTG | CAATGGAT | XM_163222 | similar to PIN4 protein | | 2.4 | SM2044 | f | 3 |
| mPool4_V2MM_105972 | 1.42983 | 75.83547 | TRUE | HP_32707 | TGCTGTTG | CTGCTGCT | BC025640 | pygopus 2 | Pygo2 | 2.16 | SM2610 | h | 2 |
| 092308m1_V2MM_131449 | 1.429086 | 3.004939 | TRUE | HP_35221 | TGCTGTTG | CCCAGTCT | XM_488636 | hypothetical gene supported by AK048895 | | 2.5 | SM2099 | e | 9 |
| mPool4_V2MM_45737 | 1.427752 | 70.87591 | TRUE | HP_26826 | TGCTGTTG | CACCCTGC | NM_173398 | G protein-coupled receptor 171 | Gpr171 | 2.16 | SM2607 | c | 7 |
| mPool2_V2MM_96077 | 1.427742 | 3.004939 | TRUE | HP_31734 | TGCTGTTG | GCCCTGA | BC011440 | histone 1, H2bc | Hist1h2bc | 2.6 | SM2137 | h | 6 |
| mPool7_V2MM_90036 | 1.427581 | 3.004939 | TRUE | HP_31146 | TGCTGTTG | CTAGATG | NM_177718 | hypothetical protein A430031N04 | | 2.9 | SM2253 | c | 3 |
| mPool2_V2MM_150932 | 1.427536 | 3.004939 | TRUE | HP_37160 | TGCTGTTG | CAGGATG | NM_011090 | paired-Ig-like receptor A3 | Pira3 | 2.4 | SM2039 | h | 1 |
| mPool2_V2MM_150932 | 1.427536 | 3.004939 | TRUE | HP_37160 | TGCTGTTG | CAGGATG | NM_011090 | paired-Ig-like receptor A3 | Pira3 | 2.4 | SM2039 | h | 1 |
| 092308m1_V2MM_159051 | 1.42726 | 3.004939 | TRUE | HP_37965 | TGCTGTTG | CAGCAGA | XM_282962 | Mus musculus hypothetical gene supported by AK029003 (LOC327858), mR. | | 2.6 | SM2119 | h | 7 |
| 092308m1_V2MM_172934 | 1.426825 | 3.004939 | TRUE | HP_39342 | TGCTGTTG | CAGGGTC | XM_286925 | Mus musculus hypothetical gene supported by AK083639 (LOC329014), mRNA. | | 2.5 | SM2069 | d | 8 |
| 092308m1_V2MM_172934 | 1.426825 | 3.004939 | TRUE | HP_39342 | TGCTGTTG | CAGGGTC | XM_286925 | Mus musculus hypothetical gene supported by AK083639 (LOC329014), mR. | | 2.5 | SM2069 | d | 8 |
| mPool2_V2MM_210580 | 1.426637 | 3.004939 | TRUE | HP_42832 | TGCTGTTG | CCATTTCC | AK080244 | RIKEN cDNA A530088I07 gene | A530088I07Rik | 2.1 | SM2348 | h | 8 |
| mPool4_V2MM_4871 | 1.426268 | 75.83547 | TRUE | HP_22843 | TGCTGTTG | CAGATACT | NM_053271 | regulating synaptic membrane exocytosis 2 | Rims2 | 2.15 | SM2598 | b | 5 |
| 092308m1_V2MM_171932 | 1.425525 | 1.959154 | TRUE | HP_39241 | TGCTGTTG | CCCTTAAC | XM_286725 | Mus musculus hypothetical gene supported by AK036009 (LOC328675), mR. | | 2.5 | SM2088 | f | 9 |
| 092308m1_V2MM_171932 | 1.425525 | 1.959154 | TRUE | HP_39241 | TGCTGTTG | CCCTTAAC | XM_286725 | Mus musculus hypothetical gene supported by AK036009 (LOC328675), mRNA. | | 2.5 | SM2088 | f | 9 |
| 092308m1_V2MM_159279 | 1.42511 | 3.004939 | TRUE | HP_37987 | TGCTGTTG | GGTTAAG | AK036657 | RIKEN cD 5830411O07 gene | 5830411O07Rik | 2.6 | SM2114 | h | 12 |
| mPool7_V2MM_103050 | 1.424667 | 3.004939 | TRUE | HP_32419 | TGCTGTTG | CTATTGAA | XM_128767 | RIKEN cDNA 4921513D11 gene | 4921513D11Rik | 2.9 | SM2300 | f | 9 |
| mPool2_V2MM_85970 | 1.424203 | 2.205692 | TRUE | HP_30745 | TGCTGTTG | CTCAAATG | NM_146533 | olfactory receptor 1367 | Olfr1367 | 2.6 | SM2144 | d | 4 |
| mPool7_V2MM_101698 | 1.423955 | 2.205692 | TRUE | HP_32286 | TGCTGTTG | GTGGAGA | XM_127811 | RIKEN cDNA 1700020M16 gene | 1700020M16Rik | 2.9 | SM2299 | f | 12 |
| 092308m3_V2MM_224880 | 1.422472 | 75.83547 | TRUE | HP_47694 | TGCTGTTG | GCCGAAAT | NM_053157 | beta-amyloid binding protein precursor | Bbp | 2.12 | SM2450 | b | 12 |
| 092308m3_V2MM_101053 | 1.420914 | 75.83547 | TRUE | HP_32222 | TGCTGTTG | GACCTCAT | NM_175150 | RIKEN cD 2310047H23 gene | 2310047H23Rik | 2.1 | SM2308 | c | 1 |
| 092308m3_V2MM_101053 | 1.420914 | 75.83547 | TRUE | HP_32222 | TGCTGTTG | GACCTCAT | NM_175150 | RIKEN cDNA 2310047H23 gene | 2310047H23Rik | 2.1 | SM2308 | c | 1 |
| 092308m1_V2MM_69950 | 1.420911 | 2.205692 | TRUE | HP_29187 | TGCTGTTG | GTGAAATT | NM_025626 | RIKEN cD 3110001A13 gene | 3110001A13Rik | 2.4 | SM2020 | d | 7 |
| 092308m3_V2MM_209057 | 1.420835 | 61.29828 | TRUE | HP_42683 | TGCTGTTG | CTGCTCAC | XM_156248 | Mus musculus LOC224196 (LOC224196), mRNA. | | 2.1 | SM2332 | d | 9 |
| 092308m3_V2MM_209057 | 1.420835 | 61.29828 | TRUE | HP_42683 | TGCTGTTG | CTGCTCAC | XM_156248 | Mus musculus LOC224196 (LOC224196), mR. | | 2.1 | SM2332 | d | 9 |
| mPool4_V2MM_90391 | 1.420092 | 69.72441 | TRUE | HP_31181 | TGCTGTTG | GCTTCATG | NM_177787 | hypothetical protein 9830102E05 | | 2.16 | SM2626 | d | 8 |
| mPool2_V2MM_7999 | 1.419749 | 2.205692 | TRUE | HP_59095 | TGCTGTTG | GCTATAGT | NM_030675 | cerebral cavernous malformations 1 | Ccm1 | 2.11 | SM2381 | a | 2 |
| 092308m1_V2MM_156212 | 1.419701 | 3.004939 | TRUE | HP_37685 | TGCTGTTG | CCTTGAAC | XM_198207 | Mus musculus LOC272573 (LOC272573), mR. | | 2.6 | SM2116 | g | 1 |
| 092308m1_V2MM_169663 | 1.419463 | 3.004939 | TRUE | HP_39015 | TGCTGTTG | CTACTGAT | AK039689 | hypothetical protein A330087I24 | | 2.6 | SM2116 | a | 7 |
| mPool5_V2MM_8062 | 1.419216 | 75.83547 | TRUE | HP_23155 | TGCTGTTG | GCTGGATC | L42339 | sodium channel, voltage-gated, type I, alpha polypeptide | Scn1a | NA | | | NA |
| mPool7_V2MM_81537 | 1.419145 | 2.205692 | TRUE | HP_30314 | TGCTGTTG | GTCCTAAA | NM_176967 | RIKEN cDNA A430072C10 gene | A430072C10Rik | 2.8 | SM2243 | f | 8 |
| mPool2_V2MM_208283 | 1.419128 | 2.205692 | TRUE | HP_42607 | TGCTGTTG | CTACTAGC | XM_154578 | Mus musculus LOC218719 (LOC218719), mRNA. | | 2.1 | SM2342 | h | 2 |
| 092308m1_V2MM_141358 | 1.418852 | 2.205692 | TRUE | HP_36207 | TGCTGTTG | GTGGGTAC | XM_156600 | Mus musculus LOC239924 (LOC239924), mR. | | 2.4 | SM2046 | a | 7 |
| mPool2_V2MM_9686 | 1.418838 | 3.004939 | TRUE | HP_23314 | TGCTGTTG | CATCTATT | NM_007856 | 7-dehydrocholesterol reductase | Dhcr7 | 2.11 | SM2382 | h | 2 |
| mPool4_V2MM_50742 | 1.418562 | 75.83547 | TRUE | HP_27315 | TGCTGTTG | CATTCAAT | NM_177593 | hypothetical protein MGC48079 | | 2.16 | SM2630 | a | 10 |
| mPool7_V2MM_174322 | 1.41812 | 3.004939 | TRUE | HP_39480 | TGCTGTTG | GCATCCA | XM_287205 | Mus musculus hypothetical gene supported by AK037602 (LOC329578), mR. | | 2.5 | SM2063 | h | 9 |
| mPool7_V2MM_174322 | 1.41812 | 3.004939 | TRUE | HP_39480 | TGCTGTTG | GCATCCA | XM_287205 | Mus musculus hypothetical gene supported by AK037602 (LOC329578), mRNA. | | 2.5 | SM2063 | h | 9 |
| 092308m1_V2MM_159103 | 1.418021 | 2.205692 | TRUE | HP_37970 | TGCTGTTG | CTGCTAAC | XM_282974 | gene model 739, (NCBI) | Gm739 | 2.6 | SM2115 | c | 5 |
| mPool4_V2MM_103 | 1.417965 | 73.80895 | TRUE | HP_22378 | TGCTGTTG | CAACCAG | NM_030558 | carbonic anhydrase 15 | Car15 | 2.15 | SM2580 | e | 7 |
| 092308m1_V2MM_144203 | 1.416695 | 3.004939 | TRUE | HP_36491 | TGCTGTTG | GTCCCTTC | XM_159803 | Mus musculus LOC207831 (LOC207831), mR. | | 2.5 | SM2057 | g | 5 |

| | | | | | | | | | | | | | |
|---|---|---|---|---|---|---|---|---|---|---|---|---|---|
| mPool4_V2MM_217576 | 1.416598 | 75.83547 | TRUE | HP_45064 | TGCTGTTG | GTGTTTCT | NM_011966 | proteasome (prosome, macropain) subunit, alpha type 4 | Psma4 | 2.15 | SM2574 | e | 6 |
| mPool4_V2MM_188490 | 1.416037 | 59.99801 | TRUE | HP_29850 | TGCTGTTG | CTCAAGTT | NM_010550 | interleukin 11 receptor, alpha chain 2 | Il11ra2 | 2.15 | SM2571 | a | 12 |
| mPool2_V2MM_202759 | 1.415503 | 2.205692 | TRUE | HP_42074 | TGCTGTTG | CCAGAAC | XM_137990 | Mus musculus similar to glyceraldehyde-3-phosphate dehydrogenase (phosphory | | 2.1 | SM2324 | b | 5 |
| 092308m3_V2MM_93509 | 1.415109 | 66.95036 | TRUE | HP_31482 | TGCTGTTG | GCTGCTAC | NM_181413 | ankyrin repeat and SAM domain containing 1 | Anks1 | 2.13 | SM2458 | f | 10 |
| mPool7_V2MM_172037 | 1.413912 | 2.205692 | TRUE | HP_39252 | TGCTGTTG | GGAAGAG | XM_286742 | gene model 813, (NCBI) | Gm813 | 2.9 | SM2273 | d | 12 |
| 092308m3_V2MM_58014 | 1.41357 | 75.83547 | TRUE | HP_28025 | TGCTGTTG | GACTGGT | NM_008866 | lysophospholipase 1 | Lypla1 | 2.12 | SM2445 | a | 8 |
| mPool7_V2MM_174595 | 1.413392 | 3.004939 | TRUE | HP_39508 | TGCTGTTG | CTCCTTAA | XM_287259 | Mus musculus hypothetical gene supported by AK053945 (LOC329683), mR. | | 2.5 | SM2068 | f | 8 |
| mPool7_V2MM_174595 | 1.413392 | 3.004939 | TRUE | HP_39508 | TGCTGTTG | CTCCTTAA | XM_287259 | Mus musculus hypothetical gene supported by AK053945 (LOC329683), mRNA. | | 2.5 | SM2068 | f | 8 |
| 092308m3_V2MM_86500 | 1.412265 | 60.13437 | TRUE | HP_30797 | TGCTGTTG | CAGTCCAT | NM_172675 | syntaxin 16 | Stx16 | 2.13 | SM2460 | d | 4 |
| mPool2_V2MM_149801 | 1.411882 | 3.004939 | TRUE | HP_37051 | TGCTGTTG | CATTGATG | XM_165199 | Mus musculus LOC209819 (LOC209819), mRNA. | | 2.1 | SM2344 | f | 1 |
| mPool2_V2MM_128720 | 1.411333 | 3.004939 | TRUE | HP_34953 | TGCTGTTG | GTGTTCTT | XM_145886 | gene model 491, (NCBI) | Gm491 | 2.11 | SM2370 | h | 7 |
| 092308m3_V2MM_102242 | 1.411324 | 75.83547 | TRUE | HP_32339 | TGCTGTTG | GCCGTGGT | XM_128190 | DEAD (Asp-Glu-Ala-Asp) box polypeptide 23 | Ddx23 | 2.13 | SM2500 | d | 2 |
| mPool2_V2MM_13293 | 1.411054 | 3.004939 | TRUE | HP_23666 | TGCTGTTG | CAGTGAAC | NM_053229 | vomeronasal 1 receptor, B8 | V1rb8 | 2.11 | SM2375 | e | 11 |
| mPool2_V2MM_83233 | 1.410856 | 3.004939 | TRUE | HP_30478 | TGCTGTTG | CTGTAAGT | AK041435 | utrophin | Utrn | 2.6 | SM2144 | f | 2 |
| mPool7_V2MM_68027 | 1.41072 | 3.004939 | TRUE | HP_28999 | TGCTGTTG | CTGATTCT | AK031552 | RIKEN cDNA 4732447D17 gene | 4732447D17Rik | 2.8 | SM2235 | e | 6 |
| 092308m3_V2MM_54987 | 1.410596 | 75.83547 | TRUE | HP_27729 | TGCTGTTG | CCCACTTT | NM_153540 | expressed sequence C85492 | C85492 | 2.12 | SM2414 | c | 12 |
| 092308m1_V2MM_146127 | 1.410463 | 2.205692 | TRUE | HP_36683 | TGCTGTTG | CAGCTTAT | XM_161773 | hypothetical gene supported by AK036869 | | 2.4 | SM2050 | a | 1 |
| 092308m1_V2MM_146127 | 1.410463 | 2.205692 | TRUE | HP_36683 | TGCTGTTG | CAGCTTAT | XM_161773 | hypothetical gene supported by AK036869 | | 2.4 | SM2050 | a | 1 |
| 092308m3_V2MM_56950 | 1.410207 | 75.83547 | TRUE | HP_27921 | TGCTGTTG | CTGGCTCT | NM_017392 | cadherin EGF LAG seven-pass G-type receptor 2 | Celsr2 | 2.12 | SM2439 | c | 8 |
| 092308m1_V2MM_166469 | 1.408364 | 3.004939 | TRUE | HP_38699 | TGCTGTTG | CAGGATAT | XM_285507 | Mus musculus similar to nucleosome assembly protein 1-like 2; brain specific ger | | 2.6 | SM2116 | f | 10 |
| mPool7_V2MM_66479 | 1.408283 | 2.205692 | TRUE | HP_28849 | TGCTGTTG | CAGCAAAC | NM_009836 | chaperonin subunit 3 (gamma) | Cct3 | 2.8 | SM2228 | a | 9 |
| mPool2_V2MM_109720 | 1.407546 | 3.004939 | TRUE | HP_33077 | TGCTGTTG | GACATGGC | XM_134085 | RIKEN cDNA 1700083M11 gene | 1700083M11Rik | 2.1 | SM2312 | g | 5 |
| mPool2_V2MM_109720 | 1.407546 | 3.004939 | TRUE | HP_33077 | TGCTGTTG | GACATGGC | XM_134085 | RIKEN cD 1700083M11 gene | 1700083M11Rik | 2.1 | SM2312 | g | 5 |
| mPool7_V2MM_79534 | 1.406818 | 3.004939 | TRUE | HP_30119 | TGCTGTTG | GCATATTC | NM_029460 | RIKEN cDNA 1700108E19 gene | 1700108E19Rik | 2.8 | SM2248 | b | 7 |
| 092308m3_V2MM_194961 | 1.406358 | 75.83547 | TRUE | HP_41311 | TGCTGTTG | GGCCGTA | XM_111160 | Mus musculus similar to eukaryotic translation initiation factor 4A1; initiation fac | | 2.9 | SM2252 | b | 3 |
| 092308m3_V2MM_194961 | 1.406358 | 75.83547 | TRUE | HP_41311 | TGCTGTTG | GGCCGTA | XM_111160 | Mus musculus similar to eukaryotic translation initiation factor 4A1; initiation fac | | 2.9 | SM2252 | b | 3 |
| mPool7_V2MM_170974 | 1.405913 | 2.205692 | TRUE | HP_39146 | TGCTGTTG | CCTTTACT | XM_286529 | gene model 1576, (NCBI) | Gm1576 | 2.5 | SM2093 | g | 11 |
| mPool7_V2MM_170974 | 1.405913 | 2.205692 | TRUE | HP_39146 | TGCTGTTG | CCTTTACT | XM_286529 | gene model 1576, (NCBI) | Gm1576 | 2.5 | SM2093 | g | 11 |
| mPool2_V2MM_83061 | 1.404691 | 3.004939 | TRUE | HP_30461 | TGCTGTTG | CACAGAAC | NM_011278 | ring finger protein 4 | Rnf4 | 2.6 | SM2133 | a | 8 |
| mPool7_V2MM_200241 | 1.40469 | 3.004939 | TRUE | HP_41827 | TGCTGTTG | GGCATATT | XM_288127 | Mus musculus hypothetical gene supported by AK053873 (LOC331495), mRNA. | | 2.9 | SM2276 | h | 9 |
| mPool2_V2MM_144230 | 1.404636 | 3.004939 | TRUE | HP_36494 | TGCTGTTG | CATTGTAC | XM_159825 | Mus musculus LOC207745 (LOC207745), mRNA. | | 2.5 | SM2058 | a | 5 |
| mPool2_V2MM_144230 | 1.404636 | 3.004939 | TRUE | HP_36494 | TGCTGTTG | CATTGTAC | XM_159825 | Mus musculus LOC207745 (LOC207745), mR. | | 2.5 | SM2058 | a | 5 |
| mPool2_V2MM_12938 | 1.404025 | 3.004939 | TRUE | HP_23632 | TGCTGTTG | CAGAGATT | NM_018818 | choroidermia | Chm | 2.11 | SM2389 | a | 2 |
| 092308m3_V2MM_44128 | 1.403683 | 62.64426 | TRUE | HP_26669 | TGCTGTTG | CTGTTTGT | NM_172588 | RIKEN cD A130038L21 gene | A130038L21Rik | 2.12 | SM2411 | h | 3 |
| 092308m1_V2MM_61794 | 1.403167 | 3.004939 | TRUE | HP_28391 | TGCTGTTG | CAAGGTG | NM_011084 | phosphatidylinositol 3-kise, C2 domain containing, gamm | Pik3c2g | 2.4 | SM2019 | h | 12 |
| 092308m1_V2MM_146706 | 1.402544 | 3.004939 | TRUE | HP_36741 | TGCTGTTG | CAATGGTC | XM_162225 | Mus musculus LOC243248 (LOC243248), mR. | | 2.5 | SM2056 | c | 11 |
| mPool2_V2MM_157721 | 1.402144 | 3.004939 | TRUE | HP_37833 | TGCTGTTG | CCACTGTA | NM_024477 | expressed sequence AI428795 | AI428795 | 2.6 | SM2132 | e | 1 |
| mPool2_V2MM_985 | 1.402002 | 3.004939 | TRUE | HP_22464 | TGCTGTTG | CAGTAAAC | NM_012013 | factor in the germline alpha | Figla | 2.11 | SM2386 | c | 12 |
| 092308m3_V2MM_203319 | 1.401706 | 75.83547 | TRUE | HP_30983 | TGCTGTTG | CTCTTCTC | NM_175397 | RIKEN cD 5830484A20 gene | 5830484A20Rik | 2.9 | SM2269 | b | 7 |
| 092308m3_V2MM_203319 | 1.401706 | 75.83547 | TRUE | HP_30983 | TGCTGTTG | CTCTTCTC | NM_175397 | RIKEN cDNA 5830484A20 gene | 5830484A20Rik | 2.9 | SM2269 | b | 7 |
| 092308m3_V2MM_25958 | 1.401588 | 75.83547 | TRUE | HP_24901 | TGCTGTTG | CCCATCGC | NM_030739 | vomerosal 1 receptor, D4 | V1rd4 | 2.12 | SM2433 | a | 11 |
| 092308m1_V2MM_139518 | 1.401294 | 2.205692 | TRUE | HP_36023 | TGCTGTTG | CAGCATAT | XM_155135 | Mus musculus LOC239258 (LOC239258), mR. | | 2.5 | SM2052 | b | 10 |
| mPool4_V2MM_50539 | 1.400632 | 75.83547 | TRUE | HP_27295 | TGCTGTTG | CAGTGCA | AK029735 | RIKEN cDNA 4930518F03 gene | 4930518F03Rik | NA | | | NA |
| mPool2_V2MM_101669 | 1.399984 | 3.004939 | TRUE | HP_13937 | TGCTGTTG | CAGTAAAT | AK088299 | regulator of chromosome condensation (RCC1) and BTB ( | Rcbtb1 | 2.1 | SM2318 | f | 10 |
| 092308m1_V2MM_173349 | 1.399787 | 2.205692 | TRUE | HP_39383 | TGCTGTTG | CACCCTTC | XM_287009 | Mus musculus hypothetical gene supported by AK036841 (LOC329196), mR. | | 2.5 | SM2077 | a | 3 |
| 092308m3_V2MM_91083 | 1.399619 | 60.13437 | TRUE | HP_31249 | TGCTGTTG | CACTTAGC | AK034060 | hypothetical protein 9330152L17 | | 2.13 | SM2463 | g | 2 |
| mPool2_V2MM_188431 | 1.398882 | 3.004939 | TRUE | HP_33206 | TGCTGTTG | GAATTTAT | NM_009917 | chemokine (C-C motif) receptor 5 | Ccr5 | 2.7 | SM2165 | f | 8 |
| mPool2_V2MM_127737 | 1.396732 | 2.205692 | TRUE | HP_34855 | TGCTGTTG | GTGACTA | XM_145359 | Mus musculus similar to vomerosal 1 receptor, G1 [Mus musculus] (LOC232868), | | 2.6 | SM2114 | c | 4 |
| mPool2_V2MM_127737 | 1.396732 | 2.205692 | TRUE | HP_34855 | TGCTGTTG | GTGACTA | XM_145359 | Mus musculus similar to vomeronasal 1 receptor, G1 [Mus musculus] (LOC23286 | | 2.6 | SM2114 | c | 4 |
| 092308m3_V2MM_37115 | 1.396609 | 62.64426 | TRUE | HP_25988 | TGCTGTTG | GACACAGC | NM_146252 | TBC1 domain family, member 13 | Tbc1d13 | 2.12 | SM2431 | g | 6 |
| mPool2_V2MM_213927 | 1.396378 | 2.205692 | TRUE | HP_43154 | TGCTGTTG | CGGCAAC | XM_140048 | Mus musculus similar to erythrocyte protein band 4.1-like 4b; expressed in high-r | | 2.11 | SM2351 | a | 6 |
| mPool2_V2MM_159763 | 1.395906 | 3.004939 | TRUE | HP_38035 | TGCTGTTG | CAGATAAC | XM_283239 | Mus musculus hypothetical gene supported by AK048863 (LOC328455), mRNA. | | 2.6 | SM2128 | d | 1 |
| 092308m3_V2MM_141606 | 1.395755 | 69.72441 | TRUE | HP_36231 | TGCTGTTG | CTGTTTCA | XM_156811 | Mus musculus LOC211465 (LOC211465), mR. | | 2.14 | SM2518 | c | 11 |

| | | | | | | | | | | | | | |
|---|---|---|---|---|---|---|---|---|---|---|---|---|---|
| mPool2_V2MM_20622 | 1.395571 | 3.004939 | TRUE | HP_24381 | TGCTGTTG | CAAATTAC | NM_008707 | N-myristoyltransferase 1 | Nmt1 | 2.11 | SM2392 | c | 10 |
| mPool2_V2MM_213769 | 1.395301 | 2.205692 | TRUE | HP_43139 | TGCTGTTG | GTTGTTCC | XM_143184 | similar to ribosomal protein L21 | | 2.11 | SM2366 | d | 5 |
| mPool2_V2MM_169067 | 1.395029 | 3.004939 | TRUE | HP_38956 | TGCTGTTG | GGTATAA | XM_286130 | Mus musculus similar to ribosomal protein S6 (LOC330873), mRNA. | | 2.6 | SM2128 | a | 1 |
| mPool4_V2MM_128857 | 1.394013 | 61.29828 | TRUE | HP_34967 | TGCTGTTG | GAGAAGA | XM_145946 | Mus musculus similar to REN [Mus musculus] (LOC244145), mRNA. | | 2.14 | SM2531 | e | 5 |
| mPool4_V2MM_34724 | 1.393375 | 75.83547 | TRUE | HP_25753 | TGCTGTTG | CCGATACC | NM_019763 | Msx2 interacting nuclear target protein | | 2.15 | SM2574 | c | 12 |
| mPool7_V2MM_68362 | 1.39232 | 3.004939 | TRUE | HP_29032 | TGCTGTTG | CTTTGCAA | NM_176972 | ubiquitin specific protease 37 | Usp37 | 2.8 | SM2245 | e | 6 |
| 092308m1_V2MM_156354 | 1.391838 | 2.205692 | TRUE | HP_37699 | TGCTGTTG | CATTTAAA | NM_212484 | RIKEN cD A230103N10 gene | A230103N10Rik | 2.6 | SM2122 | d | 7 |
| 092308m1_V2MM_182797 | 1.391389 | 3.004939 | TRUE | HP_40327 | TGCTGTTG | GAGAGTC | XM_288908 | Mus musculus LOC332375 (LOC332375), mR. | | 2.5 | SM2095 | h | 7 |
| mPool2_V2MM_117469 | 1.391119 | 3.004939 | TRUE | HP_33841 | TGCTGTTG | GGGATTCT | XM_139384 | Mus musculus similar to DKFZP434I092 protein (LOC239370), mRNA. | | 2.1 | SM2348 | a | 1 |
| 092308m1_V2MM_184558 | 1.390746 | 2.205692 | TRUE | HP_40503 | TGCTGTTG | CCAATTAT | XM_289218 | Mus musculus LOC332821 (LOC332821), mR. | | 2.5 | SM2081 | f | 12 |
| mPool2_V2MM_13630 | 1.390481 | 2.205692 | TRUE | HP_23699 | TGCTGTTG | CCAATAAA | NM_026435 | RIKEN cDNA 1810045K17 gene | 1810045K17Rik | 2.11 | SM2379 | f | 4 |
| 092308m1_V2MM_86116 | 1.39026 | 3.004939 | TRUE | HP_30759 | TGCTGTTG | CAGTGCTT | NM_146622 | olfactory receptor 360 | Olfr360 | 2.6 | SM2113 | a | 1 |
| mPool2_V2MM_169930 | 1.389017 | 3.004939 | TRUE | HP_39042 | TGCTGTTG | CTGAATTT | XM_488597 | hypothetical protein A730013G04 | | 2.6 | SM2131 | g | 1 |
| 092308m1_V2MM_141250 | 1.388988 | 3.004939 | TRUE | HP_36196 | TGCTGTTG | CTATGGG | XM_156547 | Mus musculus LOC239907 (LOC239907), mR. | | 2.4 | SM2035 | e | 4 |
| 092308m1_V2MM_170863 | 1.388959 | 3.004939 | TRUE | HP_39135 | TGCTGTTG | CCGTTGA | AK078039 | hypothetical protein 6330400D04 | | 2.5 | SM2092 | a | 11 |
| mPool2_V2MM_103237 | 1.388613 | 3.004939 | TRUE | HP_32437 | TGCTGTTG | CTGAGAG | XM_128893 | Nedd4 family interacting protein 1 | Ndfip1 | 2.6 | SM2148 | h | 9 |
| 092308m1_V2MM_181803 | 1.388076 | 3.004939 | TRUE | HP_40227 | TGCTGTTG | GAGACGA | XM_288737 | Mus musculus LOC332180 (LOC332180), mRNA. | | 2.5 | SM2065 | h | 3 |
| 092308m1_V2MM_181803 | 1.388076 | 3.004939 | TRUE | HP_40227 | TGCTGTTG | GAGACGA | XM_288737 | Mus musculus LOC332180 (LOC332180), mR. | | 2.5 | SM2065 | h | 3 |
| mPool2_V2MM_134829 | 1.387898 | 3.004939 | TRUE | HP_35555 | TGCTGTTG | GTCTGGA | XM_489254 | RIKEN cDNA 6030466F02 gene | 6030466F02Rik | 2.11 | SM2371 | f | 5 |
| 092308m1_V2MM_66786 | 1.386769 | 3.004939 | TRUE | HP_28879 | TGCTGTTG | CCAAGGT | NM_007465 | baculoviral IAP repeat-containing 2 | Birc2 | 2.3 | SM2009 | f | 9 |
| mPool6_V2MM_82018 | 1.386704 | 75.83547 | TRUE | HP_30360 | TGCTGTTG | GTGTTTAT | NM_173434 | RIKEN cDNA 9930111J21 gene | 9930111J21Rik | 2.8 | SM2222 | d | 11 |
| mPool6_V2MM_75286 | 1.386459 | 60.13437 | TRUE | HP_29706 | TGCTGTTG | CCAGCATC | NM_026319 | coiled-coil domain containing 2 | Ccdc2 | 2.8 | SM2220 | f | 2 |
| 092308m1_V2MM_175917 | 1.385919 | 2.205692 | TRUE | HP_39639 | TGCTGTTG | CCATTCAG | AK029922 | RIKEN cDNA 4930528J11 gene | 4930528J11Rik | 2.5 | SM2083 | d | 4 |
| 092308m1_V2MM_175917 | 1.385919 | 2.205692 | TRUE | HP_39639 | TGCTGTTG | CCATTCAG | AK029922 | RIKEN cD 4930528J11 gene | 4930528J11Rik | 2.5 | SM2083 | d | 4 |
| mPool2_V2MM_107081 | 1.385118 | 3.004939 | TRUE | HP_32817 | TGCTGTTG | GAAGAGT | NM_133788 | isoprenylcysteine carboxyl methyltransferase | Icmt | 2.1 | SM2318 | d | 5 |
| 092308m3_V2MM_119927 | 1.38479 | 75.83547 | TRUE | HP_27731 | TGCTGTTG | CCCAGATC | NM_146696 | olfactory receptor 1477 | Olfr1477 | 2.12 | SM2410 | c | 1 |
| mPool7_V2MM_72120 | 1.384529 | 3.004939 | TRUE | HP_29398 | TGCTGTTG | CTCTTTAA | NM_146786 | olfactory receptor 914 | Olfr914 | 2.8 | SM2241 | g | 12 |
| mPool7_V2MM_78317 | 1.383992 | 3.004939 | TRUE | HP_30000 | TGCTGTTG | GACCAGG | NM_009945 | cytochrome c oxidase, subunit VIIa 2 | Cox7a2 | 2.8 | SM2248 | b | 2 |
| mPool2_V2MM_128145 | 1.383839 | 3.004939 | TRUE | HP_34895 | TGCTGTTG | CAGAAACT | XM_145551 | Mus musculus LOC233181 (LOC233181), mRNA. | | 2.1 | SM2350 | a | 5 |
| mPool2_V2MM_148872 | 1.383768 | 3.004939 | TRUE | HP_36958 | TGCTGTTG | CATCAACC | XM_164276 | Mus musculus LOC244567 (LOC244567), mR. | | 2.1 | SM2333 | b | 6 |
| mPool2_V2MM_148872 | 1.383768 | 3.004939 | TRUE | HP_36958 | TGCTGTTG | CATCAACC | XM_164276 | Mus musculus LOC244567 (LOC244567), mRNA. | | 2.1 | SM2333 | b | 6 |
| 092308m1_V2MM_160643 | 1.383334 | 3.004939 | TRUE | HP_38122 | TGCTGTTG | GGTATCA | XM_283590 | Mus musculus similar to hypothetical protein 7 - rat (LOC329168), mR. | | 2.6 | SM2119 | h | 6 |
| 092308m3_V2MM_96506 | 1.383121 | 75.83547 | TRUE | HP_31775 | TGCTGTTG | CTGTGTTT | XM_110772 | Mus musculus similar to proteasome (prosome, macropain) subunit, beta type, 7 | | 2.13 | SM2470 | a | 10 |
| 092308m1_V2MM_124311 | 1.383058 | 1.959154 | TRUE | HP_34516 | TGCTGTTG | CCTTTACC | XM_143381 | Mus musculus similar to 60S acidic ribosomal protein P1 (LOC229548), mRNA. | | 2.5 | SM2097 | h | 1 |
| 092308m1_V2MM_124311 | 1.383058 | 1.959154 | TRUE | HP_34516 | TGCTGTTG | CCTTTACC | XM_143381 | Mus musculus similar to 60S acidic ribosomal protein P1 (LOC229548), mR. | | 2.5 | SM2097 | h | 1 |
| 092308m1_V2MM_172357 | 1.383023 | 3.004939 | TRUE | HP_39284 | TGCTGTTG | CCATGGTC | AK051520 | hypothetical protein D130054H01 | | 2.5 | SM2095 | c | 6 |
| 092308m1_V2MM_172357 | 1.383023 | 3.004939 | TRUE | HP_39284 | TGCTGTTG | CCATGGTC | AK051520 | hypothetical protein D130054H01 | | 2.5 | SM2095 | c | 6 |
| mPool6_V2MM_168245 | 1.382849 | 59.99801 | TRUE | HP_38875 | TGCTGTTG | GACTTAGT | XM_285938 | Mus musculus similar to amiloride binding protein 1 precursor; Amiloride-binding | | 2.8 | SM2205 | f | 2 |
| 092308m1_V2MM_128297 | 1.382158 | 3.004939 | TRUE | HP_34911 | TGCTGTTG | CTTAGAAT | XM_145626 | Mus musculus similar to envelope protein [Ovis aries] (LOC233260), mR. | | 2.5 | SM2097 | h | 9 |
| mPool7_V2MM_63244 | 1.3817 | 3.004939 | TRUE | HP_28533 | TGCTGTTG | CGAGTTAC | AK080900 | cholinergic receptor, nicotinic, alpha polypeptide 5 | Chrna5 | 2.8 | SM2235 | c | 4 |
| mPool2_V2MM_15141 | 1.381519 | 2.205692 | TRUE | HP_23847 | TGCTGTTG | CTATTCAA | NM_018755 | plasma glutamate carboxypeptidase | | 2.11 | SM2377 | c | 12 |
| 092308m1_V2MM_92679 | 1.380676 | 1.959154 | TRUE | HP_31402 | TGCTGTTG | GACTTGA | AK030182 | RIKEN cD A430081C19 gene | A430081C19Rik | 2.6 | SM2110 | c | 7 |
| mPool7_V2MM_135449 | 1.379237 | 3.004939 | TRUE | HP_35617 | TGCTGTTG | CTTGTCTA | XM_150641 | Mus musculus LOC240827 (LOC240827), mR. | | 2.4 | SM2035 | a | 11 |
| mPool7_V2MM_135449 | 1.379237 | 3.004939 | TRUE | HP_35617 | TGCTGTTG | CTTGTCTA | XM_150641 | Mus musculus LOC240827 (LOC240827), mRNA. | | 2.4 | SM2035 | a | 11 |
| 092308m3_V2MM_29803 | 1.379024 | 61.29828 | TRUE | HP_25275 | TGCTGTTG | CAGAGTG | NM_011923 | angiopoietin-like 2 | Angptl2 | 2.12 | SM2426 | a | 5 |
| mPool2_V2MM_114333 | 1.378143 | 2.205692 | TRUE | HP_33531 | TGCTGTTG | GCCCTTAC | XM_137447 | Mus musculus similar to DNA-directed RNA polymerase beta subunit 2 [Leptocor | | 2.7 | SM2157 | a | 2 |
| 092308m1_V2MM_143911 | 1.37805 | 3.004939 | TRUE | HP_36462 | TGCTGTTG | CAGCTATA | XM_159415 | Mus musculus LOC245652 (LOC245652), mR. | | 2.5 | SM2054 | f | 5 |
| 092308m1_V2MM_143911 | 1.37805 | 3.004939 | TRUE | HP_36462 | TGCTGTTG | CAGCTATA | XM_159415 | Mus musculus LOC245652 (LOC245652), mRNA. | | 2.5 | SM2054 | f | 5 |
| mPool7_V2MM_62878 | 1.377679 | 2.205692 | TRUE | HP_28497 | TGCTGTTG | CATTTATA | NM_146885 | olfactory receptor 1294 | Olfr1294 | 2.8 | SM2233 | e | 8 |
| mPool2_V2MM_141179 | 1.377065 | 3.004939 | TRUE | HP_36189 | TGCTGTTG | GAAATTTA | XM_156499 | Mus musculus LOC239882 (LOC239882), mRNA. | | 2.4 | SM2036 | d | 2 |
| mPool2_V2MM_141179 | 1.377065 | 3.004939 | TRUE | HP_36189 | TGCTGTTG | GAAATTTA | XM_156499 | Mus musculus LOC239882 (LOC239882), mR. | | 2.4 | SM2036 | d | 2 |
| mPool7_V2MM_201386 | 1.377022 | 1.959154 | TRUE | HP_41941 | TGCTGTTG | CCTCCTCT | XM_287151 | Mus musculus hypothetical gene supported by AK039753 (LOC329508), mRNA. | | 2.9 | SM2285 | b | 9 |
| 092308m1_V2MM_179499 | 1.376769 | 3.004939 | TRUE | HP_39997 | TGCTGTTG | CAGAGTTT | XM_288344 | Mus musculus LOC327806 (LOC327806), mRNA. | | 2.5 | SM2062 | e | 2 |

| | | | | | | | | | | | | | |
|---|---|---|---|---|---|---|---|---|---|---|---|---|---|
| 092308m1_V2MM_179499 | 1.376769 | 3.004939 | TRUE | HP_39997 | TGCTGTTG | CAGAGTTT | XM_288344 | Mus musculus LOC327806 (LOC327806), mR. | | 2.5 | SM2062 | e | 2 |
| mPool4_V2MM_241261 | 1.37659 | 75.83547 | TRUE | HP_49302 | TGCTGTTG | GGCATTGA | XM_147063 | gene model 518, (NCBI) | Gm518 | 2.14 | SM2540 | d | 6 |
| mPool2_V2MM_214087 | 1.376574 | 3.004939 | TRUE | HP_43169 | TGCTGTTG | CCATTGAA | XM_141846 | Mus musculus similar to olfactory receptor MOR264-5 (LOC236866), mRNA. | | 2.11 | SM2357 | g | 8 |
| mPool2_V2MM_1363 | 1.375258 | 3.004939 | TRUE | HP_22500 | TGCTGTTG | CCAGGATT | NM_026793 | myc target 1 | Myct1 | 2.11 | SM2386 | h | 8 |
| mPool2_V2MM_10867 | 1.375201 | 3.004939 | TRUE | HP_23428 | TGCTGTTG | CTGGGATA | NM_010867 | myomesin 1 | Myom1 | 2.11 | SM2377 | f | 6 |
| 092308m3_V2MM_143329 | 1.373046 | 75.83547 | TRUE | HP_36404 | TGCTGTTG | CAGAGCAT | XM_158642 | Mus musculus LOC214960 (LOC214960), mR. | | 2.14 | SM2520 | f | 4 |
| 092308m1_V2MM_145363 | 1.372971 | 0 | TRUE | HP_36607 | TGCTGTTG | GAGCTAGT | XM_487784 | similar to hypothetical protein FLJ35802 | | 2.4 | SM2049 | g | 8 |
| mPool2_V2MM_202732 | 1.372744 | 3.004939 | TRUE | HP_42072 | TGCTGTTG | CCATTATT | XM_138204 | Mus musculus similar to RIKEN cDNA 2810037C14 [Mus musculus] (LOC217805), | | 2.1 | SM2319 | b | 7 |
| mPool2_V2MM_202732 | 1.372744 | 3.004939 | TRUE | HP_42072 | TGCTGTTG | CCATTATT | XM_138204 | Mus musculus similar to RIKEN cD 2810037C14 [Mus musculus] (LOC217805), mR | | 2.1 | SM2319 | b | 7 |
| 092308m3_V2MM_20653 | 1.372689 | 69.72441 | TRUE | HP_24384 | TGCTGTTG | CAACGTGT | NM_030724 | uridine monophosphate kise | Umpk | 2.12 | SM2422 | d | 9 |
| mPool7_V2MM_180746 | 1.372306 | 3.004939 | TRUE | HP_40122 | TGCTGTTG | CCCATATT | XM_288558 | Mus musculus LOC331956 (LOC331956), mRNA. | | 2.5 | SM2088 | h | 1 |
| mPool7_V2MM_180746 | 1.372306 | 3.004939 | TRUE | HP_40122 | TGCTGTTG | CCCATATT | XM_288558 | Mus musculus LOC331956 (LOC331956), mR. | | 2.5 | SM2088 | h | 1 |
| 092308m1_V2MM_181014 | 1.372215 | 3.004939 | TRUE | HP_40148 | TGCTGTTG | GAGATGTT | XM_288604 | LOC433061 | | 2.5 | SM2081 | f | 5 |
| 092308m3_V2MM_87414 | 1.371194 | 75.83547 | TRUE | HP_30887 | TGCTGTTG | CTGTCGAT | NM_175161 | RIKEN cD 4931422A03 gene | 4931422A03Rik | 2.13 | SM2469 | b | 12 |
| mPool2_V2MM_19053 | 1.371023 | 3.004939 | TRUE | HP_24228 | TGCTGTTG | GGAGTTAT | NM_007641 | membrane-spanning 4-domains, subfamily A, member 1 | Ms4a1 | 2.11 | SM2381 | b | 11 |
| mPool2_V2MM_34625 | 1.370983 | 3.004939 | TRUE | HP_25743 | TGCTGTTG | CCCTGGAC | NM_172310 | RIKEN cDNA A530046H20 gene | A530046H20Rik | 2.11 | SM2391 | h | 11 |
| mPool2_V2MM_129787 | 1.370312 | 2.205692 | TRUE | HP_35058 | TGCTGTTG | GAGGATAT | XM_146456 | Mus musculus similar to cytochrome c (LOC234608), mRNA. | | 2.11 | SM2357 | h | 8 |
| mPool2_V2MM_116258 | 1.369758 | 3.004939 | TRUE | HP_33721 | TGCTGTTG | CTAAGTGT | XM_138836 | Mus musculus similar to RIKEN cDNA 2610018L09; expressed sequence AV08361 | | 2.1 | SM2312 | f | 7 |
| mPool2_V2MM_4470 | 1.36973 | 3.004939 | TRUE | HP_22804 | TGCTGTTG | GTGTAACT | L42338 | sodium channel, voltage-gated, type IX, alpha polypeptid | Scn9a | 2.11 | SM2384 | a | 1 |
| 092308m3_V2MM_203923 | 1.369369 | 75.83547 | TRUE | HP_42184 | TGCTGTTG | GGCAACGC | XM_137051 | Mus musculus similar to Gag [Ovis aries] (LOC212923), mR. | | 2.13 | SM2487 | b | 9 |
| mPool2_V2MM_213066 | 1.369171 | 3.004939 | TRUE | HP_37786 | TGCTGTTG | CTATGTAT | XM_205092 | Mus musculus similar to Zinc finger protein 180 (HHZ168) (LOC277541), mRNA. | | 2.11 | SM2355 | h | 8 |
| 092308m1_V2MM_173142 | 1.368296 | 3.004939 | TRUE | HP_39362 | TGCTGTTG | GGAGTAAT | XM_286966 | Mus musculus hypothetical gene supported by AK051123 (LOC329083), mRNA. | | 2.5 | SM2089 | f | 4 |
| 092308m1_V2MM_173142 | 1.368296 | 3.004939 | TRUE | HP_39362 | TGCTGTTG | GGAGTAAT | XM_286966 | Mus musculus hypothetical gene supported by AK051123 (LOC329083), mR. | | 2.5 | SM2089 | f | 4 |
| mPool7_V2MM_90210 | 1.367455 | 3.004939 | TRUE | HP_31163 | TGCTGTTG | CTCATATT | NM_177756 | glycosyltransferase 25 domain containing 2 | Glt25d2 | 2.9 | SM2255 | f | 1 |
| mPool2_V2MM_7117 | 1.367284 | 2.205692 | TRUE | HP_23063 | TGCTGTTG | GAGACATT | NM_018827 | cytokine receptor-like factor 1 | Crlf1 | 2.11 | SM2384 | e | 2 |
| mPool4_V2MM_9537 | 1.367204 | 35.99881 | TRUE | HP_23299 | TGCTGTTG | CAGGTTGC | NM_028034 | RIKEN cDNA 2410004F06 gene | 2410004F06Rik | 2.16 | SM2603 | h | 4 |
| mPool2_V2MM_138233 | 1.36587 | 2.205692 | TRUE | HP_35895 | TGCTGTTG | GTGTTAAC | XM_154067 | Mus musculus LOC238611 (LOC238611), mRNA. | | 2.1 | SM2343 | d | 3 |
| mPool6_V2MM_156063 | 1.365658 | 75.83547 | TRUE | HP_37670 | TGCTGTTG | CAAGTGTC | XM_198081 | Mus musculus LOC272439 (LOC272439), mRNA. | | 2.8 | SM2207 | b | 6 |
| 092308m3_V2MM_44054 | 1.365549 | 75.83547 | TRUE | HP_26662 | TGCTGTTG | CTGTGCGT | NM_021421 | D segment, Chr 1, ERATO Doi 396, expressed | D1Ertd396e | 2.12 | SM2418 | c | 4 |
| mPool4_V2MM_154881 | 1.365505 | 75.83547 | TRUE | HP_37552 | TGCTGTTG | GAAATCAC | XM_197072 | Mus musculus LOC271143 (LOC271143), mRNA. | | NA | | | NA |
| mPool7_V2MM_197384 | 1.36546 | 3.004939 | TRUE | HP_41545 | TGCTGTTG | CTATTCTC | NM_178921 | expressed sequence AI987712 | AI987712 | 2.9 | SM2254 | f | 7 |
| 092308m1_V2MM_73666 | 1.365188 | 2.205692 | TRUE | HP_29549 | TGCTGTTG | GGTCTTTG | NM_029569 | ankyrin repeat and SOCs box-containing protein 5 | Asb5 | 2.3 | SM2004 | b | 11 |
| 092308m3_V2MM_233179 | 1.365051 | 75.83547 | TRUE | HP_40072 | TGCTGTTG | CAATATCA | XM_288475 | Mus musculus LOC331813 (LOC331813), mR. | | 2.13 | SM2486 | a | 5 |
| mPool2_V2MM_169309 | 1.364618 | 3.004939 | TRUE | HP_38980 | TGCTGTTG | CTGCTTCT | XM_286231 | Mus musculus hypothetical gene supported by AK028286 (LOC327723), mRNA. | | 2.6 | SM2128 | h | 7 |
| mPool5_V2MM_377 | 1.364452 | 66.95036 | TRUE | HP_22404 | TGCTGTTG | CACCCTAA | NM_007752 | ceruloplasmin | Cp | NA | | | NA |
| mPool2_V2MM_9971 | 1.364444 | 2.205692 | TRUE | HP_23341 | TGCTGTTG | CGGAGAG | NM_133208 | zinc finger protein 287 | Zfp287 | 2.11 | SM2378 | b | 12 |
| mPool4_V2MM_220522 | 1.362919 | 75.83547 | TRUE | HP_45348 | TGCTGTTG | GGTATAGT | NM_053096 | camello-like 2 | Cml2 | 2.15 | SM2592 | b | 2 |
| mPool2_V2MM_8819 | 1.362743 | 2.205692 | TRUE | HP_23229 | TGCTGTTG | CAAATATC | NM_177719 | RIKEN cDNA 4932411A10 gene | 4932411A10Rik | 2.11 | SM2378 | d | 7 |
| mPool7_V2MM_205643 | 1.36191 | 3.004939 | TRUE | HP_42349 | TGCTGTTG | CTTTGATG | XM_127760 | Mus musculus neurofilament, light polypeptide (Nefl), mRNA. | | 2.1 | SM2317 | g | 11 |
| 092308m1_V2MM_140937 | 1.361806 | 3.004939 | TRUE | HP_36165 | TGCTGTTG | GGTGCTAT | XM_156372 | similar to RIKEN cD 1810036I24 | | 2.4 | SM2042 | a | 2 |
| 092308m1_V2MM_117571 | 1.361793 | 2.205692 | TRUE | HP_33851 | TGCTGTTG | CATAATCA | XM_139413 | Mus musculus similar to env protein [Sus scrofa] (LOC223510), mR. | | 2.5 | SM2098 | e | 12 |
| 092308m1_V2MM_129870 | 1.361555 | 3.004939 | TRUE | HP_35067 | TGCTGTTG | CTCAGGGA | XM_146492 | RIKEN cDNA 1700031F13 gene | 1700031F13Rik | 2.6 | SM2105 | d | 1 |
| 092308m1_V2MM_129870 | 1.361555 | 3.004939 | TRUE | HP_35067 | TGCTGTTG | CTCAGGGA | XM_146492 | RIKEN cD 1700031F13 gene | 1700031F13Rik | 2.6 | SM2105 | d | 1 |
| mPool2_V2MM_25870 | 1.361468 | 3.004939 | TRUE | HP_24892 | TGCTGTTG | CCATTCAA | NM_145542 | S-adenosylhomocysteine hydrolase-like 1 | Ahcyl1 | 2.11 | SM2389 | d | 8 |
| 092308m1_V2MM_172020 | 1.360958 | 3.004939 | TRUE | HP_39250 | TGCTGTTG | CAGTTTAA | XM_286739 | Mus musculus hypothetical gene supported by AK086345 (LOC328673), mRNA. | | 2.5 | SM2071 | f | 10 |
| 092308m1_V2MM_172020 | 1.360958 | 3.004939 | TRUE | HP_39250 | TGCTGTTG | CAGTTTAA | XM_286739 | Mus musculus hypothetical gene supported by AK086345 (LOC328673), mR. | | 2.5 | SM2071 | f | 10 |
| mPool2_V2MM_88194 | 1.360565 | 3.004939 | TRUE | HP_30963 | TGCTGTTG | CATTCAGT | NM_175340 | NHL repeat containing 1 | Nhlrc1 | 2.6 | SM2144 | h | 6 |
| mPool7_V2MM_84338 | 1.35974 | 3.004939 | TRUE | HP_30585 | TGCTGTTG | CAAGGGA | NM_027642 | PHD finger protein 6 | Phf6 | 2.9 | SM2269 | g | 8 |
| 092308m3_V2MM_43881 | 1.359124 | 75.83547 | TRUE | HP_26645 | TGCTGTTG | CTGCTTTC | NM_177069 | RIKEN cD E330009P21 gene | E330009P21Rik | 2.12 | SM2413 | h | 8 |
| mPool7_V2MM_189131 | 1.358544 | 3.004939 | TRUE | HP_90702 | TGCTGTTG | GAGATATA | NM_025513 | exosome component 3 | Exosc3 | 2.1 | SM2302 | h | 7 |
| mPool7_V2MM_189131 | 1.358544 | 3.004939 | TRUE | HP_90702 | TGCTGTTG | GAGATATA | NM_025513 | exosome component 3 | Exosc3 | 2.1 | SM2302 | h | 7 |
| mPool7_V2MM_108516 | 1.358177 | 2.205692 | TRUE | HP_32958 | TGCTGTTG | GATATTTG | BC060132 | RIKEN cDNA 2410127E18 gene | 2410127E18Rik | 2.1 | SM2314 | h | 4 |
| mPool6_V2MM_58613 | 1.357369 | 75.83547 | TRUE | HP_28083 | TGCTGTTG | GAGTTCAT | NM_010206 | fibroblast growth factor receptor 1 | Fgfr1 | 2.7 | SM2185 | c | 2 |

| | | | | | | | | | | | | | |
|---|---|---|---|---|---|---|---|---|---|---|---|---|---|
| 092308m3_V2MM_210133 | 1.357159 | 75.83547 | TRUE | HP_42789 | TGCTGTTG | GAGAACA | XM_195264 | similar to 60S ribosomal protein L23a | | 2.14 | SM2523 | f | 10 |
| 092308m1_V2MM_164762 | 1.357066 | 3.004939 | TRUE | HP_38530 | TGCTGTTG | GATCCCTT | XM_285090 | RIKEN cD 4930563I02 gene | 4930563I02Rik | 2.6 | SM2119 | f | 12 |
| mPool6_V2MM_190132 | 1.356935 | 75.83547 | TRUE | HP_29412 | TGCTGTTG | CTGATTGT | XM_487040 | similar to proliferation associated nuclear element 1 isoform 1; proliferation assc | | 2.7 | SM2174 | h | 9 |
| mPool2_V2MM_214912 | 1.356424 | 3.004939 | TRUE | HP_43248 | TGCTGTTG | CTATGATT | XM_149213 | RIKEN cDNA 2610318C08 gene | 2610318C08Rik | 2.11 | SM2363 | b | 7 |
| mPool7_V2MM_68410 | 1.356248 | 3.004939 | TRUE | HP_29037 | TGCTGTTG | GAAATAA | XM_484640 | similar to Ab2-450 | | 2.8 | SM2245 | h | 3 |
| mPool4_V2MM_36459 | 1.35572 | 61.29828 | TRUE | HP_25923 | TGCTGTTG | CTGTAAG | NM_008618 | malate dehydrogenase 1, NAD (soluble) | Mdh1 | 2.15 | SM2575 | e | 8 |
| mPool2_V2MM_131867 | 1.355553 | 3.004939 | TRUE | HP_35262 | TGCTGTTG | CATAGTTT | XM_147865 | Mus musculus LOC219109 (LOC219109), mRNA. | | 2.11 | SM2353 | b | 8 |
| 092308m1_V2MM_124793 | 1.35512 | 2.205692 | TRUE | HP_34564 | TGCTGTTG | CTGACCTA | XM_143634 | Mus musculus similar to hypothetical protein [Dictyostelium discoideum] (LOC24 | | 2.5 | SM2097 | b | 2 |
| mPool7_V2MM_105148 | 1.355073 | 3.004939 | TRUE | HP_32627 | TGCTGTTG | GGCTTAG | AK079931 | peptidyl-prolyl isomerase G (cyclophilin G) | Ppig | 2.9 | SM2300 | a | 9 |
| 092308m1_V2MM_141179 | 1.35469 | 3.004939 | TRUE | HP_36189 | TGCTGTTG | GAAATTTA | XM_156499 | Mus musculus LOC239882 (LOC239882), mRNA. | | 2.4 | SM2036 | d | 2 |
| 092308m1_V2MM_141179 | 1.35469 | 3.004939 | TRUE | HP_36189 | TGCTGTTG | GAAATTTA | XM_156499 | Mus musculus LOC239882 (LOC239882), mR. | | 2.4 | SM2036 | d | 2 |
| mPool7_V2MM_203507 | 1.354421 | 3.004939 | TRUE | HP_42145 | TGCTGTTG | GTGCTCTA | XM_128618 | similar to zinc finger protein 97 | | 2.9 | SM2299 | a | 12 |
| mPool6_V2MM_154578 | 1.354106 | 35.99881 | TRUE | HP_37521 | TGCTGTTG | CATGGAA | XM_196854 | Mus musculus LOC270856 (LOC270856), mRNA. | | 2.7 | SM2191 | c | 8 |
| mPool2_V2MM_145920 | 1.353848 | 3.004939 | TRUE | HP_36663 | TGCTGTTG | CTGATTCC | XM_161614 | Mus musculus LOC242812 (LOC242812), mRNA. | | 2.1 | SM2337 | b | 7 |
| 092308m3_V2MM_94622 | 1.353759 | 61.29828 | TRUE | HP_31590 | TGCTGTTG | GATTTAAT | NM_183192 | cD sequence BC027582 | BC027582 | 2.13 | SM2483 | c | 7 |
| mPool4_V2MM_146407 | 1.353673 | 75.83547 | TRUE | HP_36711 | TGCTGTTG | CTAAGTTC | XM_161991 | Mus musculus LOC243104 (LOC243104), mRNA. | | 2.16 | SM2644 | h | 12 |
| 092308m1_V2MM_118369 | 1.353132 | 3.004939 | TRUE | HP_33930 | TGCTGTTG | CTTACAAA | XM_139818 | Mus musculus similar to hypothetical protein FLJ32191 [Homo sapiens] (LOC224 | | 2.6 | SM2101 | a | 6 |
| 092308m1_V2MM_118369 | 1.353132 | 3.004939 | TRUE | HP_33930 | TGCTGTTG | CTTACAAA | XM_139818 | Mus musculus similar to hypothetical protein FLJ32191 [Homo sapiens] (LOC224 | | 2.6 | SM2101 | a | 6 |
| 092308m3_V2MM_51219 | 1.352802 | 75.83547 | TRUE | HP_27362 | TGCTGTTG | CTACTATG | NM_010174 | fatty acid binding protein 3, muscle and heart | Fabp3 | 2.12 | SM2445 | f | 11 |
| mPool2_V2MM_23246 | 1.352559 | 2.205692 | TRUE | HP_24638 | TGCTGTTG | CTGCTGA | NM_009373 | transglutaminase 2, C polypeptide | Tgm2 | 2.7 | SM2166 | b | 9 |
| 092308m1_V2MM_179849 | 1.351574 | 3.004939 | TRUE | HP_40032 | TGCTGTTG | CAGCAAA | XM_288408 | Mus musculus LOC331731 (LOC331731), mR. | | 2.5 | SM2096 | b | 2 |
| 092308m3_V2MM_103016 | 1.351204 | 62.64426 | TRUE | HP_32416 | TGCTGTTG | CTGGGTT | XM_128755 | Mus musculus spastic paraplegia 4 homolog (human) (Spg4), mR. | | 2.13 | SM2494 | g | 10 |
| mPool7_V2MM_203752 | 1.350019 | 3.004939 | TRUE | HP_42168 | TGCTGTTG | GAAGCTA | XM_285947 | Mus musculus similar to ribosomal protein L7a; surfeit 3 [Mus musculus] (LOC33 | | 2.1 | SM2322 | e | 1 |
| mPool7_V2MM_196913 | 1.349989 | 3.004939 | TRUE | HP_41500 | TGCTGTTG | CCAATACA | NM_178735 | RIKEN cDNA A730069N07 gene | A730069N07Rik | 2.9 | SM2259 | c | 12 |
| mPool7_V2MM_67681 | 1.349842 | 2.205692 | TRUE | HP_28966 | TGCTGTTG | CTAGTATT | NM_172931 | RIKEN cDNA B230358A15 gene | B230358A15Rik | 2.8 | SM2228 | h | 6 |
| mPool4_V2MM_138982 | 1.349787 | 75.83547 | TRUE | HP_35970 | TGCTGTTG | GTCACTGA | XM_154625 | Mus musculus LOC218785 (LOC218785), mRNA. | | 2.16 | SM2646 | e | 4 |
| 092308m1_V2MM_155277 | 1.34867 | 3.004939 | TRUE | HP_37591 | TGCTGTTG | GCTATTGT | XM_197334 | Mus musculus LOC271472 (LOC271472), mR. | | 2.6 | SM2122 | a | 11 |
| 092308m1_V2MM_159110 | 1.347827 | 3.004939 | TRUE | HP_37971 | TGCTGTTG | CTATGGA | XM_282976 | similar to hypothetical protein FLJ20345 | | 2.6 | SM2121 | e | 3 |
| 092308m3_V2MM_37804 | 1.347705 | 35.99881 | TRUE | HP_26054 | TGCTGTTG | GAGCTGC | AK032985 | RIKEN cD 2210018M11 gene | 2210018M11Rik | 2.12 | SM2421 | a | 5 |
| mPool7_V2MM_62510 | 1.347177 | 3.004939 | TRUE | HP_28461 | TGCTGTTG | CAGGACTT | NM_009897 | creatine kinase, mitochondrial 1, ubiquitous | Ckmt1 | 2.8 | SM2244 | a | 2 |
| 092308m1_V2MM_182121 | 1.345644 | 3.004939 | TRUE | HP_40259 | TGCTGTTG | GTTAGAG | XM_288790 | Mus musculus LOC332251 (LOC332251), mR. | | 2.5 | SM2085 | c | 7 |
| 092308m3_V2MM_49039 | 1.344435 | 75.83547 | TRUE | HP_27148 | TGCTGTTG | GCTGCTTT | NM_177175 | RIKEN cD A930001M12 gene | A930001M12Rik | 2.12 | SM2403 | e | 10 |
| 092308m3_V2MM_26629 | 1.34437 | 60.13437 | TRUE | HP_24966 | TGCTGTTG | CTCACATT | NM_133865 | D cross-link repair 1B, PSO2 homolog (S. cerevisiae) | Dclre1b | 2.12 | SM2429 | e | 4 |
| mPool7_V2MM_187401 | 1.343034 | 3.004939 | TRUE | HP_28498 | TGCTGTTG | CATTTCAT | AK087793 | importin 11 | Ipo11 | 2.9 | SM2287 | f | 8 |
| mPool6_V2MM_4129 | 1.342903 | 61.29828 | TRUE | HP_22770 | TGCTGTTG | GGGACTG | NM_008193 | guanylate kinase 1 | Guk1 | 2.7 | SM2182 | h | 4 |
| mPool7_V2MM_197099 | 1.342702 | 1.959154 | TRUE | HP_41518 | TGCTGTTG | CTATTTCT | NM_173783 | RIKEN cDNA B430216B18 gene | B430216B18Rik | 2.9 | SM2254 | a | 11 |
| mPool7_V2MM_71118 | 1.342637 | 3.004939 | TRUE | HP_29301 | TGCTGTTG | CATAAGA | XM_128178 | splicing factor, arginine/serine-rich 2, interacting protein | Sfrs2ip | 2.8 | SM2228 | d | 6 |
| mPool2_V2MM_131509 | 1.341959 | 3.004939 | TRUE | HP_35227 | TGCTGTTG | CACCCAA | XM_147687 | cDNA sequence BC042761 | BC042761 | 2.7 | SM2160 | h | 7 |
| mPool4_V2MM_208232 | 1.341815 | 75.83547 | TRUE | HP_42602 | TGCTGTTG | CTGTGTTG | XM_162903 | Mus musculus LOC208790 (LOC208790), mRNA. | | 2.16 | SM2634 | f | 9 |
| mPool4_V2MM_107294 | 1.340547 | 75.83547 | TRUE | HP_32838 | TGCTGTTG | CTGGGCA | AK006670 | RIKEN cDNA 1700041E20 gene | 1700041E20Rik | 2.16 | SM2615 | b | 11 |
| mPool7_V2MM_181741 | 1.340467 | 3.004939 | TRUE | HP_40221 | TGCTGTTG | CAAGTTAC | XM_288727 | Mus musculus LOC332169 (LOC332169), mRNA. | | 2.5 | SM2082 | d | 7 |
| mPool7_V2MM_181741 | 1.340467 | 3.004939 | TRUE | HP_40221 | TGCTGTTG | CAAGTTAC | XM_288727 | Mus musculus LOC332169 (LOC332169), mR. | | 2.5 | SM2082 | d | 7 |
| 092308m3_V2MM_102585 | 1.340278 | 60.13437 | TRUE | HP_32373 | TGCTGTTG | CCAAGGA | BC019219 | RIKEN cD 2610036F08 gene | 2610036F08Rik | 2.13 | SM2489 | f | 2 |
| mPool7_V2MM_66554 | 1.340191 | 3.004939 | TRUE | HP_28856 | TGCTGTTG | CAGGGAT | NM_172914 | hypothetical protein 4933409I22 | | 2.8 | SM2240 | b | 6 |
| 092308m3_V2MM_28083 | 1.339447 | 61.29828 | TRUE | HP_25107 | TGCTGTTG | GCACCAG | M24780 | phosphoglycerate kise 1, pseudogene 1 | Pgk1-ps1 | 2.12 | SM2428 | c | 2 |
| 092308m3_V2MM_235610 | 1.339331 | 75.83547 | TRUE | HP_48747 | TGCTGTTG | GGAACTT | XM_157106 | Mus musculus LOC211989 (LOC211989), mR. | NA | | | NA | |
| 092308m3_V2MM_86934 | 1.338899 | 75.83547 | TRUE | HP_33284 | TGCTGTTG | GGTGGAC | NM_174852 | PHD finger protein 12 | Phf12 | 2.13 | SM2469 | g | 11 |
| 092308m1_V2MM_168117 | 1.33885 | 1.959154 | TRUE | HP_38862 | TGCTGTTG | GAAATTGT | XM_285912 | Mus musculus similar to RE12057p (LOC330264), mR. | | 2.6 | SM2117 | f | 8 |
| mPool2_V2MM_162105 | 1.338726 | 3.004939 | TRUE | HP_38266 | TGCTGTTG | GAAAGTA | XM_284203 | Mus musculus hypothetical gene supported by AK030693 (LOC330316), mRNA. | | 2.6 | SM2127 | f | 10 |
| mPool6_V2MM_155837 | 1.338314 | 69.72441 | TRUE | HP_37647 | TGCTGTTG | GTTGCTCT | XM_197851 | Mus musculus LOC269619 (LOC269619), mRNA. | | 2.8 | SM2212 | f | 9 |
| 092308m3_V2MM_48464 | 1.338009 | 75.83547 | TRUE | HP_27092 | TGCTGTTG | GATTCAA | AK044730 | RIKEN cD A930037J23 gene | A930037J23Rik | 2.12 | SM2445 | h | 6 |
| 092308m1_V2MM_94159 | 1.337499 | 3.004939 | TRUE | HP_31545 | TGCTGTTG | CCTTAACT | AK013327 | RIKEN cD 9130017K11 gene | 9130017K11Rik | 2.6 | SM2108 | e | 10 |
| 092308m1_V2MM_123221 | 1.33728 | 3.004939 | TRUE | HP_34407 | TGCTGTTG | CATGTATG | XM_142785 | RIKEN cD C030035D04 gene | C030035D04Rik | 2.6 | SM2103 | d | 7 |

| | | | | | | | | | | | | | |
|---|---|---|---|---|---|---|---|---|---|---|---|---|---|
| 092308m3_V2MM_93385 | 1.336599 | 75.83547 | TRUE | HP_31470 | TGCTGTTG | CACCCTAG | NM_181345 | nucleophosmin/nucleoplasmin 2 | Npm2 | 2.9 | SM2265 | e | 1 |
| 092308m3_V2MM_93385 | 1.336599 | 75.83547 | TRUE | HP_31470 | TGCTGTTG | CACCCTAG | NM_181345 | nucleophosmin/nucleoplasmin 2 | Npm2 | 2.9 | SM2265 | e | 1 |
| 092308m1_V2MM_181932 | 1.336473 | 3.004939 | TRUE | HP_40240 | TGCTGTTG | CACCAAG( | XM_288759 | Mus musculus LOC332197 (LOC332197), mR. | | 2.5 | SM2094 | d | 3 |
| mPool2_V2MM_100868 | 1.336464 | 3.004939 | TRUE | HP_32204 | TGCTGTTG | GACCATAT | XM_127169 | Mus musculus similar to periaxin (LOC217886), mRNA. | | 2.1 | SM2312 | c | 3 |
| mPool2_V2MM_114036 | 1.336367 | 2.205692 | TRUE | HP_33502 | TGCTGTTG | GTGAGACT | NM_001003717 | oxysterol binding protein-like 8 | Osbpl8 | 2.7 | SM2157 | d | 3 |
| 092308m1_V2MM_152690 | 1.335699 | 3.004939 | TRUE | HP_37335 | TGCTGTTG | CTGAATTT | XM_195505 | Mus musculus similar to hypothetical protein MGC34837 [Homo sapiens] (LOC26 | | 2.6 | SM2114 | c | 10 |
| mPool2_V2MM_210411 | 1.335659 | 2.205692 | TRUE | HP_42816 | TGCTGTTG | CAACATA/ | XM_145002 | Mus musculus similar to hypothetical protein MGC34648 [Homo sapiens] (LOC21 | | 2.11 | SM2354 | b | 10 |
| mPool7_V2MM_90177 | 1.335654 | 3.004939 | TRUE | HP_31160 | TGCTGTTG | GTTATCCT | NM_177748 | killer immunoglobulin-like receptor-like 2 | | 2.9 | SM2259 | d | 7 |
| mPool7_V2MM_90177 | 1.335654 | 3.004939 | TRUE | HP_31160 | TGCTGTTG | GTTATCCT | NM_177748 | killer immunoglobulin-like receptor-like 2 | | 2.9 | SM2259 | d | 7 |
| 092308m3_V2MM_85206 | 1.335163 | 61.29828 | TRUE | HP_30669 | TGCTGTTG | CAGTATAT | NM_133225 | acyl-Coenzyme A binding domain containing 3 | Acbd3 | 2.8 | SM2250 | c | 11 |
| 092308m3_V2MM_85206 | 1.335163 | 61.29828 | TRUE | HP_30669 | TGCTGTTG | CAGTATAT | NM_133225 | acyl-Coenzyme A binding domain containing 3 | Acbd3 | 2.8 | SM2250 | c | 11 |
| mPool2_V2MM_16553 | 1.334731 | 2.205692 | TRUE | HP_23984 | TGCTGTTG | GACAAAT( | NM_134199 | vomeronasal 1 receptor, F2 | V1rf2 | 2.11 | SM2381 | f | 10 |
| mPool4_V2MM_262223 | 1.334692 | 75.83547 | TRUE | HP_52297 | TGCTGTTG | CTATTCCT | XM_195633 | Mus musculus similar to vomeronasal receptor V1RD12 [Mus musculus] (LOC272 | | 2.16 | SM2622 | a | 8 |
| 092308m1_V2MM_147832 | 1.334451 | 3.004939 | TRUE | HP_36854 | TGCTGTTG | GAAATCT/ | XM_163306 | Mus musculus LOC215016 (LOC215016), mR. | | 2.4 | SM2049 | a | 9 |
| 092308m3_V2MM_45464 | 1.334246 | 66.95036 | TRUE | HP_26799 | TGCTGTTG | GTGCATAT | AK038854 | neurexin I | Nrxn1 | 2.12 | SM2407 | c | 4 |
| mPool6_V2MM_74693 | 1.334019 | 75.83547 | TRUE | HP_29649 | TGCTGTTG | CAGCTAAT | NM_007381 | acetyl-Coenzyme A dehydrogenase, long-chain | Acadl | 2.8 | SM2218 | c | 11 |
| 092308m1_V2MM_178348 | 1.3338 | 3.004939 | TRUE | HP_39882 | TGCTGTTG | GTCCTAAC | AK081752 | hypothetical protein C130073P14 | | 2.5 | SM2092 | a | 3 |
| mPool4_V2MM_210787 | 1.333753 | 66.95036 | TRUE | HP_42852 | TGCTGTTG | GTGAGCA, | XM_142291 | Mus musculus similar to differentiation inhibitor protein Id2A - human (LOC2372 | | 2.14 | SM2541 | f | 1 |
| 092308m3_V2MM_142011 | 1.332867 | 60.13437 | TRUE | HP_36272 | TGCTGTTG | CACATCAT | XM_157271 | Mus musculus LOC240249 (LOC240249), mR. | | 2.14 | SM2512 | g | 6 |
| mPool7_V2MM_185131 | 1.332667 | 3.004939 | TRUE | HP_40560 | TGCTGTTG | CTACTTGA | XM_289316 | Mus musculus LOC332973 (LOC332973), mR. | | 2.5 | SM2066 | f | 10 |
| mPool7_V2MM_185131 | 1.332667 | 3.004939 | TRUE | HP_40560 | TGCTGTTG | CTACTTGA | XM_289316 | Mus musculus LOC332973 (LOC332973), mRNA. | | 2.5 | SM2066 | f | 10 |
| 092308m1_V2MM_141085 | 1.331848 | 2.205692 | TRUE | HP_36179 | TGCTGTTG | CTATTGAG | XM_156459 | Mus musculus LOC239959 (LOC239959), mR. | | 2.4 | SM2036 | g | 12 |
| 092308m1_V2MM_141085 | 1.331848 | 2.205692 | TRUE | HP_36179 | TGCTGTTG | CTATTGAG | XM_156459 | Mus musculus LOC239959 (LOC239959), mRNA. | | 2.4 | SM2036 | g | 12 |
| 092308m3_V2MM_234348 | 1.331842 | 75.83547 | TRUE | HP_48622 | TGCTGTTG | GTTATCCT | XM_137038 | Mus musculus similar to protease [Mus musculus] (LOC215893), mR. | | 2.13 | SM2491 | a | 3 |
| 092308m3_V2MM_231569 | 1.331187 | 61.29828 | TRUE | HP_48353 | TGCTGTTG | CACATGG/ | XM_131217 | cD sequence BC007180 | BC007180 | 2.13 | SM2495 | c | 9 |
| mPool6_V2MM_166047 | 1.331078 | 75.83547 | TRUE | HP_38657 | TGCTGTTG | GTCACGG/ | XM_285398 | similar to High mobility group protein 1-like 10 (HMG-1L10) | | 2.8 | SM2206 | h | 4 |
| mPool2_V2MM_1144 | 1.33081 | 3.004939 | TRUE | HP_22479 | TGCTGTTG | CATATTCA | XM_136490 | Mus musculus similar to degenerative spermatocyte-like protein RDES [Rattus nc | | 2.11 | SM2377 | b | 8 |
| 092308m3_V2MM_189641 | 1.329225 | 69.72441 | TRUE | HP_32713 | TGCTGTTG | GCTTATGT | XM_355444 | RIKEN cD 1700021C14 gene | 1700021C14Rik | 2.14 | SM2507 | h | 10 |
| mPool7_V2MM_209654 | 1.328044 | 3.004939 | TRUE | HP_42742 | TGCTGTTG | CCCAAGG/ | XM_151310 | Mus musculus LOC211412 (LOC211412), mRNA. | | 2.1 | SM2329 | h | 11 |
| mPool7_V2MM_209654 | 1.328044 | 3.004939 | TRUE | HP_42742 | TGCTGTTG | CCCAAGG/ | XM_151310 | Mus musculus LOC211412 (LOC211412), mR. | | 2.1 | SM2329 | h | 11 |
| 092308m1_V2MM_64478 | 1.32689 | 3.004939 | TRUE | HP_28653 | TGCTGTTG | CTGGTGAT | NM_146836 | olfactory receptor 1132 | Olfr1132 | 2.3 | SM2010 | e | 3 |
| mPool2_V2MM_123173 | 1.326661 | 2.205692 | TRUE | HP_34402 | TGCTGTTG | CAATTTGA | XM_142727 | nuclear antigen Sp100 | Sp100 | 2.11 | SM2369 | d | 9 |
| mPool4_V2MM_140191 | 1.326409 | 70.87591 | TRUE | HP_36091 | TGCTGTTG | GTGATCAC | XM_155791 | Mus musculus LOC239475 (LOC239475), mRNA. | | 2.16 | SM2637 | g | 5 |
| mPool4_V2MM_110591 | 1.325895 | 75.83547 | TRUE | HP_33162 | TGCTGTTG | CAAGATT( | XM_489276 | RIKEN cDNA 1700036A12 gene | 1700036A12Rik | 2.16 | SM2608 | c | 8 |
| 092308m1_V2MM_158721 | 1.325738 | 3.004939 | TRUE | HP_37933 | TGCTGTTG | GACCTTTA | XM_207141 | similar to SmcY protein (Histocompatibility Y antigen) (H-Y) | | 2.6 | SM2121 | d | 6 |
| mPool7_V2MM_80618 | 1.324783 | 3.004939 | TRUE | HP_30224 | TGCTGTTG | GGAATTA( | NM_025702 | RIKEN cDNA 3526402H21 gene | 3526402H21Rik | 2.8 | SM2243 | c | 11 |
| 092308m1_V2MM_173571 | 1.324646 | 3.004939 | TRUE | HP_39405 | TGCTGTTG | CAGACAT( | XM_287051 | Mus musculus hypothetical gene supported by AK042187 (LOC329287), mR. | | 2.5 | SM2072 | e | 2 |
| 092308m3_V2MM_88748 | 1.324347 | 62.64426 | TRUE | HP_31018 | TGCTGTTG | GTTCTTAT | NM_175481 | glutamate receptor, ionotropic, kaite 4 | Grik4 | 2.13 | SM2480 | h | 12 |
| mPool7_V2MM_92571 | 1.323838 | 2.205692 | TRUE | HP_31391 | TGCTGTTG | GACCTTA/ | NM_178735 | RIKEN cDNA A730069N07 gene | A730069N07Rik | 2.9 | SM2253 | d | 7 |
| mPool2_V2MM_143843 | 1.323587 | 3.004939 | TRUE | HP_36455 | TGCTGTTG | GCTCACAC | XM_159211 | Mus musculus LOC245417 (LOC245417), mRNA. | | 2.1 | SM2324 | g | 11 |
| mPool7_V2MM_200046 | 1.323157 | 2.205692 | TRUE | HP_41808 | TGCTGTTG | CATCTACT | AK080130 | hypothetical protein A530065N20 | | 2.9 | SM2284 | c | 3 |
| mPool7_V2MM_195927 | 1.322947 | 3.004939 | TRUE | HP_41404 | TGCTGTTG | CAATGTTC | NM_176837 | Rho GTPase activating protein 18 | Arhgap18 | 2.9 | SM2264 | h | 1 |
| mPool2_V2MM_2099 | 1.322328 | 0 | TRUE | HP_22572 | TGCTGTTG | CTATACCT | NM_011406 | solute carrier family 8 (sodium/calcium exchanger), mem | Slc8a1 | 2.11 | SM2387 | d | 4 |
| 092308m3_V2MM_51919 | 1.322024 | 75.83547 | TRUE | HP_27430 | TGCTGTTG | CTGGAAT/ | NM_028534 | stromal membrane-associated protein 1 | Smap1 | 2.12 | SM2405 | a | 3 |
| mPool7_V2MM_195087 | 1.321884 | 3.004939 | TRUE | HP_41323 | TGCTGTTG | GGCTCAG | XM_110936 | similar to peptidyl-dipeptidase A (EC 3.4.15.1), testis - mouse | | 2.9 | SM2253 | a | 3 |
| mPool2_V2MM_108097 | 1.32131 | 3.004939 | TRUE | HP_32917 | TGCTGTTG | GTCACTTC | AK002701 | RIKEN cDNA 0610030E20 gene | 0610030E20Rik | 2.7 | SM2151 | g | 12 |
| 092308m1_V2MM_165846 | 1.321286 | 3.004939 | TRUE | HP_38638 | TGCTGTTG | CCCGAGA/ | NM_026361 | plakophilin 4 | Pkp4 | 2.6 | SM2118 | a | 10 |
| mPool2_V2MM_24795 | 1.320845 | 2.205692 | TRUE | HP_24788 | TGCTGTTG | GGTGTTGT | NM_019702 | Hbs1-like (S. cerevisiae) | Hbs1l | 2.11 | SM2390 | a | 3 |
| 092308m1_V2MM_181407 | 1.319822 | 3.004939 | TRUE | HP_40188 | TGCTGTTG | CAGACCA/ | XM_288670 | Mus musculus LOC332096 (LOC332096), mR. | | 2.5 | SM2060 | e | 4 |
| 092308m1_V2MM_177974 | 1.31922 | 2.205692 | TRUE | HP_39845 | TGCTGTTG | GACCTGAT | XM_287992 | Mus musculus hypothetical gene supported by AK078598 (LOC331116), mR. | | 2.5 | SM2091 | h | 9 |
| 092308m3_V2MM_197217 | 1.318688 | 75.83547 | TRUE | HP_41529 | TGCTGTTG | CAGCTCAC | XM_487805 | similar to limitin | | 2.9 | SM2255 | c | 7 |
| 092308m3_V2MM_197217 | 1.318688 | 75.83547 | TRUE | HP_41529 | TGCTGTTG | CAGCTCAC | XM_487805 | similar to limitin | | 2.9 | SM2255 | c | 7 |
| mPool2_V2MM_86235 | 1.318485 | 3.004939 | TRUE | HP_30771 | TGCTGTTG | CTGGCACT | NM_146658 | olfactory receptor 1131 | Olfr1131 | 2.6 | SM2141 | d | 8 |

| | | | | | | | | | | | | | |
|---|---|---|---|---|---|---|---|---|---|---|---|---|---|
| 092308m3_V2MM_43419 | 1.3182 | 75.83547 | TRUE | HP_26600 | TGCTGTTG | CTCCTCTA | NM_016667 | syntrophin, basic 1 | Sntb1 | 2.12 | SM2403 | b | 8 |
| mPool2_V2MM_115572 | 1.317861 | 3.004939 | TRUE | HP_33653 | TGCTGTTG | CCTGTTGC | XM_138503 | Mus musculus similar to spermine synthase; spermidine aminopropyltransferase | | 2.7 | SM2157 | h | 1 |
| 092308m1_V2MM_182565 | 1.317846 | 3.004939 | TRUE | HP_40303 | TGCTGTTG | GACTAAAT | XM_288868 | RIKEN cDNA 4930546C10 gene | 4930546C10Rik | 2.5 | SM2071 | d | 10 |
| 092308m1_V2MM_182565 | 1.317846 | 3.004939 | TRUE | HP_40303 | TGCTGTTG | GACTAAAT | XM_288868 | RIKEN cD 4930546C10 gene | 4930546C10Rik | 2.5 | SM2071 | d | 10 |
| mPool7_V2MM_85297 | 1.317825 | 3.004939 | TRUE | HP_30678 | TGCTGTTG | GCTTTATC | NM_133907 | ubiquitin protein ligase E3C | Ube3c | 2.9 | SM2267 | e | 12 |
| mPool7_V2MM_198272 | 1.317695 | 3.004939 | TRUE | HP_41632 | TGCTGTTG | CAGGGAT | XM_287379 | Mus musculus hypothetical gene supported by AK039622 (LOC329946), mRNA. | | 2.9 | SM2276 | h | 1 |
| mPool7_V2MM_65152 | 1.317428 | 3.004939 | TRUE | HP_28719 | TGCTGTTG | GATATGAC | NM_025687 | testis expressed gene 12 | Tex12 | 2.8 | SM2231 | f | 2 |
| 092308m1_V2MM_84959 | 1.317312 | 2.205692 | TRUE | HP_30645 | TGCTGTTG | CATAATTA | NM_029947 | PR domain containing 8 | Prdm8 | 2.6 | SM2110 | g | 2 |
| mPool2_V2MM_101887 | 1.31709 | 3.004939 | TRUE | HP_32305 | TGCTGTTG | GGCTAAAT | NM_027707 | RIKEN cDNA 4933421G18 gene | 4933421G18Rik | 2.7 | SM2157 | c | 8 |
| mPool2_V2MM_210897 | 1.316985 | 3.004939 | TRUE | HP_42863 | TGCTGTTG | GAGATTAC | AK021091 | tryptophan rich basic protein | Wrb | 2.11 | SM2353 | e | 5 |
| mPool4_V2MM_197645 | 1.316257 | 75.83547 | TRUE | HP_41571 | TGCTGTTG | CGTGATGC | XM_125547 | solute carrier family 22 (organic cation transporter), men | Slc22a16 | 2.16 | SM2624 | g | 4 |
| mPool2_V2MM_147313 | 1.31617 | 2.205692 | TRUE | HP_36802 | TGCTGTTG | GGAAACA | XM_162860 | Mus musculus LOC232484 (LOC232484), mR. | | 2.5 | SM2052 | f | 9 |
| mPool2_V2MM_147313 | 1.31617 | 2.205692 | TRUE | HP_36802 | TGCTGTTG | GGAAACA | XM_162860 | Mus musculus LOC232484 (LOC232484), mRNA. | | 2.5 | SM2052 | f | 9 |
| 092308m1_V2MM_125029 | 1.31501 | 3.004939 | TRUE | HP_34587 | TGCTGTTG | CATCCTGC | XM_143748 | forkhead box E1 (thyroid transcription factor 2) | Foxe1 | 2.6 | SM2111 | e | 8 |
| mPool2_V2MM_3511 | 1.314849 | 2.205692 | TRUE | HP_22710 | TGCTGTTG | GAGTTACA | NM_011562 | teratocarcinoma-derived growth factor | Tdgf1 | 2.11 | SM2380 | e | 10 |
| mPool6_V2MM_160019 | 1.314336 | 75.83547 | TRUE | HP_38061 | TGCTGTTG | CCAACTGT | AK084162 | Casitas B-lineage lymphoma b | Cblb | 2.7 | SM2196 | d | 11 |
| mPool2_V2MM_153912 | 1.314013 | 3.004939 | TRUE | HP_37455 | TGCTGTTG | CAGAGATT | BC068148 | RIKEN cDNA 4732460K03 gene | 4732460K03Rik | 2.6 | SM2128 | b | 4 |
| mPool4_V2MM_212376 | 1.31331 | 61.29828 | TRUE | HP_43004 | TGCTGTTG | GCACATGA | XM_358748 | zinc finger protein 597 | Zfp597 | 2.14 | SM2535 | a | 6 |
| mPool6_V2MM_223790 | 1.313082 | 61.29828 | TRUE | HP_47591 | TGCTGTTG | CCTAGGAA | XM_284817 | similar to bM64F17.3 (putative novel protein) | | 2.7 | SM2197 | h | 4 |
| 092308m1_V2MM_181239 | 1.313008 | 3.004939 | TRUE | HP_40171 | TGCTGTTG | CGGGTAG | XM_288642 | Mus musculus LOC332046 (LOC332046), mR. | | 2.5 | SM2070 | d | 11 |
| mPool6_V2MM_159820 | 1.312976 | 66.95036 | TRUE | HP_38041 | TGCTGTTG | GCATTCGT | AK048688 | F-box and leucine-rich repeat protein 7 | Fbxl7 | 2.8 | SM2206 | d | 1 |
| mPool7_V2MM_113276 | 1.312816 | 3.004939 | TRUE | HP_33426 | TGCTGTTG | CACAGTAT | XM_136902 | Mus musculus similar to hypothetical protein A830058L05 [Mus musculus] (LOC2 | | 2.7 | SM2152 | f | 4 |
| mPool4_V2MM_130642 | 1.3127 | 75.83547 | TRUE | HP_35142 | TGCTGTTG | CAGCCCAA | XM_146963 | RIKEN cDNA C530008M07 gene | C530008M07Rik | 2.11 | SM2372 | c | 6 |
| mPool2_V2MM_6731 | 1.311953 | 3.004939 | TRUE | HP_23025 | TGCTGTTG | CTTTGATT | NM_144500 | oxysterol binding protein-like 2 | Osbpl2 | 2.11 | SM2387 | h | 11 |
| 092308m1_V2MM_82112 | 1.310989 | 2.205692 | TRUE | HP_30369 | TGCTGTTG | GTTGCTTT | NM_176998 | RIKEN cD A830027B17 gene | A830027B17Rik | 2.3 | SM2016 | b | 9 |
| 092308m3_V2MM_58280 | 1.310958 | 75.83547 | TRUE | HP_28051 | TGCTGTTG | GAGATTAT | AK083747 | RIKEN cD D030074E01 gene | D030074E01Rik | 2.12 | SM2411 | d | 6 |
| 092308m3_V2MM_147598 | 1.310865 | 59.99801 | TRUE | HP_36830 | TGCTGTTG | GCAACGCA | XM_163110 | Mus musculus LOC243757 (LOC243757), mR. | | 2.4 | SM2044 | e | 9 |
| 092308m3_V2MM_36096 | 1.310844 | 75.83547 | TRUE | HP_25888 | TGCTGTTG | CTGAGAAA | NM_023595 | deoxyuridine triphosphatase | Dutp | 2.11 | SM2396 | c | 12 |
| mPool6_V2MM_162517 | 1.310776 | 75.83547 | TRUE | HP_38307 | TGCTGTTG | CACCCTAT | XM_284382 | Mus musculus hypothetical gene supported by AK032428 (LOC330700), mRNA. | | 2.8 | SM2208 | g | 10 |
| mPool4_V2MM_46831 | 1.310754 | 75.83547 | TRUE | HP_26932 | TGCTGTTG | CGACGTTC | NM_139228 | rhomboid, veinlet-like 4 (Drosophila) | Rhbdl4 | 2.15 | SM2600 | c | 1 |
| mPool2_V2MM_214537 | 1.310336 | 2.205692 | TRUE | HP_43212 | TGCTGTTG | GAAGTTAT | XM_141933 | similar to Smage-1 protein | | 2.11 | SM2368 | d | 1 |
| mPool6_V2MM_37864 | 1.30933 | 75.83547 | TRUE | HP_26060 | TGCTGTTG | GAGGCCA | NM_009247 | serine (or cysteine) proteinase inhibitor, clade A, membe | Serpina1e | 2.7 | SM2170 | g | 6 |
| 092308m1_V2MM_119905 | 1.308989 | 3.004939 | TRUE | HP_34081 | TGCTGTTG | CAACTAGT | XM_488868 | hypothetical gene supported by AK041445 | | 2.5 | SM2099 | b | 3 |
| 092308m1_V2MM_157072 | 1.308938 | 2.205692 | TRUE | HP_37769 | TGCTGTTG | GCTCCTGT | XM_204841 | gene model 703, (NCBI) | Gm703 | 2.6 | SM2116 | g | 7 |
| 092308m1_V2MM_131242 | 1.308465 | 3.004939 | TRUE | HP_35200 | TGCTGTTG | CTCCCAAC | AB093288 | cytoplasmic FMR1 interacting protein 2 | Cyfip2 | 2.5 | SM2100 | d | 9 |
| mPool2_V2MM_137283 | 1.308407 | 2.205692 | TRUE | HP_35800 | TGCTGTTG | CAACCAAC | XM_152972 | Mus musculus LOC237973 (LOC237973), mRNA. | | 2.5 | SM2053 | f | 4 |
| mPool2_V2MM_137283 | 1.308407 | 2.205692 | TRUE | HP_35800 | TGCTGTTG | CAACCAAC | XM_152972 | Mus musculus LOC237973 (LOC237973), mR. | | 2.5 | SM2053 | f | 4 |
| 092308m3_V2MM_23028 | 1.307768 | 61.29828 | TRUE | HP_24616 | TGCTGTTG | CTGAAATA | NM_145578 | ubiquitin-conjugating enzyme E2M (UBC12 homolog, yea | Ube2m | 2.12 | SM2430 | c | 7 |
| mPool6_V2MM_79122 | 1.307434 | 75.83547 | TRUE | HP_30079 | TGCTGTTG | GATACCTT | NM_010695 | lipocalin 4 | Lcn4 | 2.8 | SM2223 | g | 4 |
| mPool2_V2MM_125948 | 1.305386 | 3.004939 | TRUE | HP_34679 | TGCTGTTG | CCCAAGAT | XM_144299 | similar to CG31559-PA | | 2.11 | SM2362 | f | 2 |
| mPool7_V2MM_95102 | 1.304662 | 3.004939 | TRUE | HP_31638 | TGCTGTTG | GATTCAGA | NM_183390 | kelch-like 6 (Drosophila) | Klhl6 | 2.9 | SM2253 | e | 12 |
| mPool7_V2MM_95102 | 1.304662 | 3.004939 | TRUE | HP_31638 | TGCTGTTG | GATTCAGA | NM_183390 | kelch-like 6 (Drosophila) | Klhl6 | 2.9 | SM2253 | e | 12 |
| mPool4_V2MM_157080 | 1.30386 | 60.13437 | TRUE | HP_37770 | TGCTGTTG | GAATGGA | XM_204847 | Mus musculus similar to protease [Trichosurus vulpecula] (LOC277161), mRNA. | | NA | | | NA |
| mPool7_V2MM_186224 | 1.303803 | 3.004939 | TRUE | HP_40669 | TGCTGTTG | CAAGTAAA | XM_289503 | Mus musculus LOC333220 (LOC333220), mRNA. | | 2.5 | SM2060 | c | 4 |
| mPool7_V2MM_186224 | 1.303803 | 3.004939 | TRUE | HP_40669 | TGCTGTTG | CAAGTAAA | XM_289503 | Mus musculus LOC333220 (LOC333220), mR. | | 2.5 | SM2060 | c | 4 |
| 092308m3_V2MM_92129 | 1.302436 | 59.99801 | TRUE | HP_31349 | TGCTGTTG | GCAAGACI | NM_178655 | ankyrin 2, brain | Ank2 | 2.13 | SM2464 | g | 11 |
| 092308m1_V2MM_176254 | 1.301912 | 2.205692 | TRUE | HP_39673 | TGCTGTTG | CCAAATAT | XM_287631 | Mus musculus RIKEN cD 3110052D19 gene (3110052D19Rik), mR. | | 2.5 | SM2085 | f | 3 |
| 092308m1_V2MM_63085 | 1.301278 | 3.004939 | TRUE | HP_28517 | TGCTGTTG | CCTCAGTT | U47328 | histocompatibility 2, K1, K region | H2-K1 | 2.4 | SM2019 | d | 8 |
| 092308m3_V2MM_93253 | 1.30127 | 75.83547 | TRUE | HP_31458 | TGCTGTTG | GATGGAA | NM_178936 | RIKEN cD 4930577M16 gene | 4930577M16Rik | 2.9 | SM2265 | h | 5 |
| 092308m3_V2MM_93253 | 1.30127 | 75.83547 | TRUE | HP_31458 | TGCTGTTG | GATGGAA | NM_178936 | RIKEN cDNA 4930577M16 gene | 4930577M16Rik | 2.9 | SM2265 | h | 5 |
| mPool7_V2MM_171499 | 1.301051 | 3.004939 | TRUE | HP_39198 | TGCTGTTG | CTTGGATA | AK043702 | RIKEN cDNA A830021K08 gene | A830021K08Rik | 2.9 | SM2281 | b | 9 |
| mPool2_V2MM_104605 | 1.301025 | 3.004939 | TRUE | HP_32573 | TGCTGTTG | CACTGAGC | XM_129858 | similar to General transcription factor 3C polypeptide 3 (Transcription factor IIIC- | | 2.7 | SM2162 | d | 7 |
| mPool4_V2MM_92064 | 1.300716 | 75.83547 | TRUE | HP_31342 | TGCTGTTG | CCAAGGG | NM_178642 | expressed sequence AU040576 | AU040576 | 2.16 | SM2608 | f | 5 |

| | | | | | | | | | | | | | |
|---|---|---|---|---|---|---|---|---|---|---|---|---|---|
| mPool7_V2MM_170548 | 1.300373 | 3.004939 | TRUE | HP_39104 | TGCTGTTG | CTTGGTTA | XM_286438 | Mus musculus hypothetical gene supported by AK090201 (LOC328098), mRNA. | | 2.9 | SM2291 | f | 11 |
| 092308m1_V2MM_166416 | 1.299703 | 2.205692 | TRUE | HP_38694 | TGCTGTTG | CTGCAAAT | XM_285495 | Mus musculus similar to TRAF6-inhibitory zinc finger protein; TRAF6-binding zinc | | 2.6 | SM2122 | f | 1 |
| mPool4_V2MM_217595 | 1.299404 | 69.72441 | TRUE | HP_45066 | TGCTGTTG | CTGGTGGT | NM_011227 | RAB20, member RAS oncogene family | Rab20 | 2.15 | SM2570 | h | 4 |
| mPool4_V2MM_195467 | 1.299216 | 75.83547 | TRUE | HP_41360 | TGCTGTTG | CACGGAAT | NM_146994 | olfactory receptor 201 | Olfr201 | 2.15 | SM2593 | c | 3 |
| mPool7_V2MM_171062 | 1.296879 | 3.004939 | TRUE | HP_39155 | TGCTGTTG | GGTTTACT | AK052551 | hypothetical protein D430050G20 | | 2.5 | SM2075 | d | 1 |
| mPool4_V2MM_205752 | 1.296597 | 75.83547 | TRUE | HP_42360 | TGCTGTTG | CCAAGCA( | AK081462 | neuropilin (NRP) and tolloid (TLL)-like 2 | Neto2 | 2.16 | SM2618 | h | 2 |
| mPool2_V2MM_165073 | 1.296259 | 3.004939 | TRUE | HP_38561 | TGCTGTTG | CTTGACAA | XM_285162 | Mus musculus similar to Gag [Ovis aries] (LOC332147), mRNA. | | 2.6 | SM2128 | e | 6 |
| mPool2_V2MM_108926 | 1.296093 | 2.205692 | TRUE | HP_32999 | TGCTGTTG | CAAGCATT | XM_133390 | similar to low density lipoprotein receptor-related protein 3 | | 2.7 | SM2158 | d | 4 |
| mPool2_V2MM_95105 | 1.29546 | 2.205692 | TRUE | HP_31638 | TGCTGTTG | CTAAGCTA | NM_183391 | tumor necrosis factor (ligand) superfamily, member 18 | Tnfsf18 | 2.6 | SM2106 | f | 11 |
| mPool2_V2MM_95105 | 1.29546 | 2.205692 | TRUE | HP_31638 | TGCTGTTG | CTAAGCTA | NM_183391 | tumor necrosis factor (ligand) superfamily, member 18 | Tnfsf18 | 2.6 | SM2106 | f | 11 |
| mPool2_V2MM_132410 | 1.295242 | 2.205692 | TRUE | HP_35316 | TGCTGTTG | GAGTTTAT | XM_358797 | similar to Stefin homolog | | 2.11 | SM2362 | e | 10 |
| mPool6_V2MM_160009 | 1.295129 | 75.83547 | TRUE | HP_38060 | TGCTGTTG | CGCTTGCT | XM_489526 | hypothetical gene supported by AK080987 | | 2.8 | SM2202 | e | 11 |
| mPool7_V2MM_172897 | 1.294525 | 3.004939 | TRUE | HP_39338 | TGCTGTTG | CTCATTAT | XM_286918 | Mus musculus hypothetical gene supported by AK084135 (LOC328996), mRNA. | | 2.9 | SM2276 | b | 4 |
| 092308m3_V2MM_204747 | 1.29448 | 60.13437 | TRUE | HP_42263 | TGCTGTTG | GCCACAG/ | XM_132795 | gene model 154, (NCBI) | Gm154 | 2.1 | SM2322 | e | 4 |
| 092308m3_V2MM_204747 | 1.29448 | 60.13437 | TRUE | HP_42263 | TGCTGTTG | GCCACAG/ | XM_132795 | gene model 154, (NCBI) | Gm154 | 2.1 | SM2322 | e | 4 |
| 092308m1_V2MM_148152 | 1.294315 | 3.004939 | TRUE | HP_36886 | TGCTGTTG | GGTAAAT/ | XM_163535 | Mus musculus LOC244134 (LOC244134), mRNA. | | 2.5 | SM2052 | f | 11 |
| 092308m1_V2MM_148152 | 1.294315 | 3.004939 | TRUE | HP_36886 | TGCTGTTG | GGTAAAT/ | XM_163535 | Mus musculus LOC244134 (LOC244134), mR. | | 2.5 | SM2052 | f | 11 |
| 092308m1_V2MM_120042 | 1.294306 | 3.004939 | TRUE | HP_34095 | TGCTGTTG | CATGTCTT | XM_140732 | Mus musculus similar to siah1A protein [Rattus norvegicus] (LOC226200), mR. | | 2.6 | SM2101 | b | 11 |
| mPool7_V2MM_78169 | 1.293934 | 3.004939 | TRUE | HP_29986 | TGCTGTTG | GACAAAG. | NM_023051 | calsyntenin 1 | Clstn1 | 2.8 | SM2233 | e | 11 |
| mPool4_V2MM_4912 | 1.293914 | 70.87591 | TRUE | HP_22847 | TGCTGTTG | CAGCCAAT | BC034893 | RIKEN cDNA 3110037K17 gene | 3110037K17Rik | 2.15 | SM2597 | e | 3 |
| mPool4_V2MM_227 | 1.293673 | 75.83547 | TRUE | HP_22390 | TGCTGTTG | CAATGTGT | NM_027188 | SET and MYND domain containing 3 | Smyd3 | 2.15 | SM2582 | f | 1 |
| mPool2_V2MM_7217 | 1.293628 | 3.004939 | TRUE | HP_23073 | TGCTGTTG | GAGCTAAT | NM_026627 | RIKEN cDNA 1700113O17 gene | 1700113O17Rik | 2.11 | SM2376 | e | 6 |
| mPool7_V2MM_136065 | 1.29294 | 1.959154 | TRUE | HP_35678 | TGCTGTTG | GCTGAAT( | XM_151339 | Mus musculus LOC211802 (LOC211802), mRNA. | | 2.4 | SM2040 | e | 8 |
| mPool7_V2MM_136065 | 1.29294 | 1.959154 | TRUE | HP_35678 | TGCTGTTG | GCTGAAT( | XM_151339 | Mus musculus LOC211802 (LOC211802), mR. | | 2.4 | SM2040 | e | 8 |
| 092308m3_V2MM_141975 | 1.290409 | 59.99801 | TRUE | HP_36268 | TGCTGTTG | GATTATGT | NM_157245 | Mus musculus LOC240234 (LOC240234), mR. | | 2.14 | SM2525 | e | 3 |
| 092308m3_V2MM_82338 | 1.289881 | 75.83547 | TRUE | HP_30391 | TGCTGTTG | CCTACATG | NM_008400 | integrin alpha L | Itgal | 2.13 | SM2468 | e | 3 |
| mPool2_V2MM_118274 | 1.289156 | 3.004939 | TRUE | HP_33921 | TGCTGTTG | CAGTCATA | XM_139758 | Mus musculus similar to formyl peptide receptor, related sequence 3 [Mus musc | | 2.11 | SM2358 | a | 12 |
| 092308m1_V2MM_68971 | 1.28892 | 3.004939 | TRUE | HP_29091 | TGCTGTTG | GAGTTATA | NM_025510 | RIKEN cD 2310004I24 gene | 2310004I24Rik | 2.4 | SM2021 | b | 4 |
| mPool4_V2MM_20367 | 1.288546 | 61.29828 | TRUE | HP_24356 | TGCTGTTG | GTGTTGT( | NM_153080 | target of myb1-like 2 (chicken) | Tom1l2 | 2.16 | SM2602 | b | 12 |
| 092308m3_V2MM_25306 | 1.288441 | 35.99881 | TRUE | HP_24837 | TGCTGTTG | CACTGACT | NM_007908 | eukaryotic elongation factor-2 kise | Eef2k | 2.12 | SM2401 | g | 3 |
| mPool2_V2MM_18541 | 1.288124 | 2.205692 | TRUE | HP_24178 | TGCTGTTG | GCTAAAT( | NM_025820 | Crn, crooked neck-like 1 (Drosophila) | Crnkl1 | 2.11 | SM2377 | g | 7 |
| mPool2_V2MM_160947 | 1.287844 | 2.205692 | TRUE | HP_38152 | TGCTGTTG | GTCCTGCT | XM_484975 | hypothetical gene supported by AK048278 | | 2.6 | SM2131 | a | 9 |
| mPool2_V2MM_136212 | 1.287828 | 3.004939 | TRUE | HP_35693 | TGCTGTTG | GTCGAAAT | XM_151496 | Mus musculus LOC241166 (LOC241166), mR. | | 2.4 | SM2042 | c | 3 |
| mPool2_V2MM_136212 | 1.287828 | 3.004939 | TRUE | HP_35693 | TGCTGTTG | GTCGAAAT | XM_151496 | Mus musculus LOC241166 (LOC241166), mRNA. | | 2.4 | SM2042 | c | 3 |
| mPool7_V2MM_104283 | 1.287633 | 3.004939 | TRUE | HP_32541 | TGCTGTTG | GAACCAAT | XM_129655 | cDNA sequence BC013667 | BC013667 | 2.1 | SM2304 | h | 9 |
| mPool7_V2MM_104283 | 1.287633 | 3.004939 | TRUE | HP_32541 | TGCTGTTG | GAACCAAT | XM_129655 | cD sequence BC013667 | BC013667 | 2.1 | SM2304 | h | 9 |
| mPool7_V2MM_194879 | 1.28745 | 2.205692 | TRUE | HP_41303 | TGCTGTTG | CCCTTTCT | NM_183126 | RIKEN cDNA 6030498E09 gene | 6030498E09Rik | 2.9 | SM2267 | h | 8 |
| mPool7_V2MM_194879 | 1.28745 | 2.205692 | TRUE | HP_41303 | TGCTGTTG | CCCTTTCT | NM_183126 | RIKEN cD 6030498E09 gene | 6030498E09Rik | 2.9 | SM2267 | h | 8 |
| mPool6_V2MM_224007 | 1.285019 | 75.83547 | TRUE | HP_47610 | TGCTGTTG | GCTGTATT | XM_484861 | similar to axonemal dynein heavy chain 7 | | 2.7 | SM2198 | a | 9 |
| 092308m1_V2MM_155374 | 1.284987 | 3.004939 | TRUE | HP_37601 | TGCTGTTG | GAGATAA/ | XM_197402 | Mus musculus LOC271565 (LOC271565), mR. | | 2.6 | SM2122 | g | 12 |
| 092308m1_V2MM_174345 | 1.284919 | 3.004939 | TRUE | HP_39483 | TGCTGTTG | GATTTAAT | XM_287210 | Mus musculus LOC329595 (LOC329595), mR. | | 2.5 | SM2062 | g | 3 |
| mPool5_V2MM_66560 | 1.284912 | 61.29828 | TRUE | HP_28857 | TGCTGTTG | CAGGGTTT | NM_010323 | gonadotropin releasing hormone receptor | Gnrhr | NA | | NA | |
| 092308m3_V2MM_237569 | 1.28491 | 66.95036 | TRUE | HP_48941 | TGCTGTTG | GAGCTTCT | XM_155441 | Mus musculus LOC239511 (LOC239511), mR. | | 2.14 | SM2524 | a | 3 |
| 092308m1_V2MM_187569 | 1.284424 | 2.205692 | TRUE | HP_40803 | TGCTGTTG | CATTTAAT | XM_290006 | Mus musculus similar to pol polyprotein [Trichosurus vulpecula] (LOC333873), m | | 2.5 | SM2065 | f | 3 |
| mPool2_V2MM_28070 | 1.282798 | 2.205692 | TRUE | HP_25106 | TGCTGTTG | GCAATAA( | NM_030730 | steroid receptor-interacting SNF2 domain protein | | 2.11 | SM2391 | f | 10 |
| mPool6_V2MM_82021 | 1.282521 | 59.99801 | TRUE | HP_30360 | TGCTGTTG | GTGTTTCT | NM_146752 | olfactory receptor 1106 | Olfr1106 | 2.7 | SM2179 | a | 6 |
| 092308m1_V2MM_161814 | 1.282315 | 3.004939 | TRUE | HP_38237 | TGCTGTTG | CACCTGGT | AK122544 | D segment, Chr 5, Wayne State University 178, expressec | D5Wsu178e | 2.6 | SM2115 | h | 11 |
| 092308m1_V2MM_176186 | 1.282285 | 3.004939 | TRUE | HP_39666 | TGCTGTTG | GAAGCTAT | XM_287619 | Mus musculus hypothetical gene supported by AK035337 (LOC330373), mR. | | 2.5 | SM2094 | a | 8 |
| 092308m1_V2MM_66050 | 1.281985 | 3.004939 | TRUE | HP_80906 | TGCTGTTG | GTGCTTTA | NM_010446 | forkhead box A2 | Foxa2 | 2.4 | SM2027 | f | 10 |
| 092308m1_V2MM_163750 | 1.281981 | 2.205692 | TRUE | HP_38430 | TGCTGTTG | CCATATAA | XM_284847 | Mus musculus similar to KIAA1466 protein [Homo sapiens] (LOC329320), mR. | | 2.6 | SM2116 | f | 2 |
| mPool5_V2MM_912 | 1.281942 | 66.95036 | TRUE | HP_22456 | TGCTGTTG | CAGGCTG( | NM_007808 | cytochrome c, somatic | Cycs | NA | | NA | |
| mPool2_V2MM_101104 | 1.281923 | 2.205692 | TRUE | HP_32227 | TGCTGTTG | CTTACTTG | AK006112 | RIKEN cDNA 0610007P08 gene | 0610007P08Rik | 2.6 | SM2150 | c | 4 |
| mPool7_V2MM_187236 | 1.281511 | 2.205692 | TRUE | HP_40770 | TGCTGTTG | CATAGTTA | XM_289671 | Mus musculus LOC333434 (LOC333434), mRNA. | | 2.9 | SM2286 | g | 7 |

| | | | | | | | | | | | | | | | |
|---|---|---|---|---|---|---|---|---|---|---|---|---|---|---|---|
| 092308m3_V2MM_116020 | 1.280604 | 75.83547 | TRUE | HP_33697 | TGCTGTTG | CATCCAAC | XM_138719 | Mus musculus similar to endogenous retroviral family W, env(C7), member 1 (syr | | | 2.14 | SM2508 | f | | 12 |
| mPool4_V2MM_131730 | 1.280494 | 75.83547 | TRUE | HP_35249 | TGCTGTTG | GTCCGTGT | AK014636 | RIKEN cDNA 4733401I05 gene | 4733401I05Rik | | 2.16 | SM2631 | c | | 4 |
| mPool6_V2MM_238650 | 1.279919 | 75.83547 | TRUE | HP_47486 | TGCTGTTG | CCCACTGC | XM_141448 | Mus musculus similar to protease [Trichosurus vulpecula] (LOC212819), mRNA. | | | 2.8 | SM2212 | e | | 8 |
| mPool2_V2MM_30527 | 1.279875 | 3.004939 | TRUE | HP_25345 | TGCTGTTG | CGGAAACT | NM_145492 | zinc finger protein 521 | Zfp521 | | 2.11 | SM2389 | g | | 10 |
| 092308m3_V2MM_111611 | 1.279309 | 59.99801 | TRUE | HP_33262 | TGCTGTTG | CGCTATCT | XM_135982 | RIKEN cD 6530406P05 gene | 6530406P05Rik | | 2.13 | SM2491 | h | | 12 |
| 092308m1_V2MM_148248 | 1.279012 | 3.004939 | TRUE | HP_36895 | TGCTGTTG | CACAGGA( | XM_163622 | Mus musculus LOC207376 (LOC207376), mRNA. | | | 2.5 | SM2056 | f | | 2 |
| 092308m1_V2MM_148248 | 1.279012 | 3.004939 | TRUE | HP_36895 | TGCTGTTG | CACAGGA( | XM_163622 | Mus musculus LOC207376 (LOC207376), mR. | | | 2.5 | SM2056 | f | | 2 |
| mPool2_V2MM_212991 | 1.278948 | 3.004939 | TRUE | HP_43064 | TGCTGTTG | CAGCTTTG | XM_488592 | RIKEN cDNA 5133400J02 gene | 5133400J02Rik | | 2.11 | SM2352 | e | | 9 |
| 092308m3_V2MM_47599 | 1.278706 | 75.83547 | TRUE | HP_27007 | TGCTGTTG | CTGTCTGC | NM_080846 | RIKEN cD 2310056K19 gene | 2310056K19Rik | | 2.12 | SM2405 | d | | 12 |
| 092308m1_V2MM_137567 | 1.278446 | 3.004939 | TRUE | HP_35828 | TGCTGTTG | CCAGGAC/ | XM_153307 | Mus musculus LOC208340 (LOC208340), mR. | | | 2.5 | SM2053 | g | | 9 |
| 092308m1_V2MM_178239 | 1.277698 | 3.004939 | TRUE | HP_39871 | TGCTGTTG | CACCAAG( | AK032250 | hypothetical protein 6430511F03 | | | 2.5 | SM2086 | f | | 5 |
| 092308m3_V2MM_95294 | 1.276198 | 61.29828 | TRUE | HP_31657 | TGCTGTTG | CAGCCTAT | NM_197943 | RUN and TBC1 domain containing 1 | Rutbc1 | | 2.9 | SM2254 | d | | 3 |
| 092308m3_V2MM_95294 | 1.276198 | 61.29828 | TRUE | HP_31657 | TGCTGTTG | CAGCCTAT | NM_197943 | RUN and TBC1 domain containing 1 | Rutbc1 | | 2.9 | SM2254 | d | | 3 |
| 092308m3_V2MM_93163 | 1.275389 | 75.83547 | TRUE | HP_31449 | TGCTGTTG | GTCTGGAT | NM_178919 | expressed sequence AI451006 | AI451006 | | 2.13 | SM2473 | e | | 2 |
| mPool4_V2MM_227676 | 1.275148 | 75.83547 | TRUE | HP_47966 | TGCTGTTG | GTATTAAT | XM_286653 | Mus musculus hypothetical gene supported by AK038948 (LOC328512), mRNA. | | NA | | | | NA | |
| 092308m3_V2MM_95830 | 1.274739 | 75.83547 | TRUE | HP_31709 | TGCTGTTG | CTTGTGTG | AK047814 | RIKEN cD A730024A03 gene | A730024A03Rik | | 2.12 | SM2449 | d | | 5 |
| mPool2_V2MM_121209 | 1.274586 | 2.205692 | TRUE | HP_34209 | TGCTGTTG | CACAAAT/ | XM_141541 | similar to 40S ribosomal protein S16 | | | 2.11 | SM2369 | h | | 5 |
| 092308m3_V2MM_224463 | 1.274567 | 59.99801 | TRUE | HP_47653 | TGCTGTTG | GCATCGA( | AK032385 | D segment, Chr 14, ERATO Doi 171, expressed | D14Ertd171e | | 2.13 | SM2482 | h | | 2 |
| mPool4_V2MM_214845 | 1.274325 | 75.83547 | TRUE | HP_18032 | TGCTGTTG | GAAGCTAT | XM_486245 | similar to Rpl7a protein | | | 2.1 | SM2308 | h | | 9 |
| 092308m1_V2MM_169374 | 1.274293 | 2.205692 | TRUE | HP_38986 | TGCTGTTG | CAAATATG | AK045844 | hypothetical protein B230314J19 | | | 2.6 | SM2123 | d | | 10 |
| mPool7_V2MM_185237 | 1.273848 | 3.004939 | TRUE | HP_40570 | TGCTGTTG | GAGCTAA/ | XM_289336 | Mus musculus LOC333017 (LOC333017), mRNA. | | | 2.9 | SM2272 | h | | 8 |
| mPool5_V2MM_7199 | 1.273055 | 75.83547 | TRUE | HP_23071 | TGCTGTTG | GAGCCAGT | NM_054085 | expressed sequence AW319487 | AW319487 | NA | | | | NA | |
| 092308m3_V2MM_110527 | 1.273049 | 75.83547 | TRUE | HP_33156 | TGCTGTTG | CAGTTGA/ | XM_134736 | zinc finger protein 26 | Zfp26 | | 2.13 | SM2500 | c | | 10 |
| mPool7_V2MM_152047 | 1.271704 | 3.004939 | TRUE | HP_37271 | TGCTGTTG | GCAAATG/ | XM_195063 | Mus musculus similar to potassium channel regulatory factor (LOC271501), mRN. | | | 2.1 | SM2325 | e | | 6 |
| 092308m3_V2MM_225789 | 1.271662 | 75.83547 | TRUE | HP_47782 | TGCTGTTG | CACGAGG( | XM_125996 | similar to SEC14-like protein 3 (45 kDa secretory protein) (rsec45) | | | 2.13 | SM2479 | c | | 6 |
| mPool2_V2MM_125119 | 1.271155 | 3.004939 | TRUE | HP_34596 | TGCTGTTG | GAAGAAG | XM_143807 | Mus musculus similar to alpha-2u-globulin IV precursor - mouse (LOC209155), ml | | | 2.11 | SM2368 | h | | 9 |
| mPool7_V2MM_110117 | 1.271136 | 3.004939 | TRUE | HP_33115 | TGCTGTTG | GTGATATT | XM_134391 | Mus musculus nuclear factor of activated T-cells, cytoplasmic 3 (Nfatc3), mRNA. | | | 2.9 | SM2299 | d | | 4 |
| 092308m3_V2MM_86482 | 1.270881 | 66.95036 | TRUE | HP_30795 | TGCTGTTG | CCCTCAG( | NM_172601 | RAB2B, member RAS oncogene family | Rab2b | | 2.13 | SM2471 | a | | 10 |
| mPool4_V2MM_209512 | 1.270453 | 75.83547 | TRUE | HP_42728 | TGCTGTTG | CTCCACAT | XM_195276 | Mus musculus similar to hypothetical protein [Plasmodium falciparum 3D7] (LOC | | NA | | | | NA | |
| 092308m1_V2MM_163858 | 1.270306 | 3.004939 | TRUE | HP_38441 | TGCTGTTG | CAGTTGGT | XM_284870 | Mus musculus similar to hypothetical protein D930020E02 [Mus musculus] (LOC3 | | | 2.6 | SM2123 | c | | 9 |
| mPool2_V2MM_107492 | 1.270014 | 2.205692 | TRUE | HP_32857 | TGCTGTTG | GAATCGA/ | XM_132134 | RIKEN cDNA 1110019K23 gene | 1110019K23Rik | | 2.6 | SM2149 | f | | 7 |
| mPool2_V2MM_44691 | 1.270002 | 3.004939 | TRUE | HP_26724 | TGCTGTTG | GCATTGG( | NM_009692 | apolipoprotein A-I | Apoa1 | | 2.7 | SM2165 | b | | 6 |
| mPool2_V2MM_21454 | 1.269953 | 2.205692 | TRUE | HP_24462 | TGCTGTTG | CAGTAGT/ | NM_027722 | nudix (nucleoside diphosphate linked moiety X)-type mot | Nudt4 | | 2.11 | SM2390 | d | | 10 |
| 092308m3_V2MM_51736 | 1.269804 | 75.83547 | TRUE | HP_27412 | TGCTGTTG | CTGACTTG | NM_138683 | thrombospondin type 1 domain containing gene | | | 2.12 | SM2409 | c | | 3 |
| 092308m3_V2MM_111320 | 1.26953 | 75.83547 | TRUE | HP_33233 | TGCTGTTG | CATAGGA( | XM_135683 | similar to tripartite motif-containing 43 | | | 2.1 | SM2302 | e | | 11 |
| 092308m3_V2MM_111320 | 1.26953 | 75.83547 | TRUE | HP_33233 | TGCTGTTG | CATAGGA( | XM_135683 | similar to tripartite motif-containing 43 | | | 2.1 | SM2302 | e | | 11 |
| 092308m3_V2MM_20561 | 1.269264 | 75.83547 | TRUE | HP_24375 | TGCTGTTG | GAATGGC/ | NM_030734 | defensin beta 5 | Defb5 | | 2.12 | SM2434 | e | | 11 |
| 092308m1_V2MM_136398 | 1.268689 | 2.205692 | TRUE | HP_35711 | TGCTGTTG | GTTCTTCA | XM_151696 | Mus musculus LOC237328 (LOC237328), mR. | | | 2.4 | SM2043 | e | | 12 |
| 092308m1_V2MM_161461 | 1.268399 | 3.004939 | TRUE | HP_38202 | TGCTGTTG | GCCAGAAT | XM_283928 | Mus musculus hypothetical gene supported by AK042285 (LOC329760), mR. | | | 2.6 | SM2120 | b | | 12 |
| mPool7_V2MM_66155 | 1.268317 | 3.004939 | TRUE | HP_28817 | TGCTGTTG | GTTGAAG/ | NM_025630 | RIKEN cDNA 2010009L17 gene | 2010009L17Rik | | 2.8 | SM2231 | h | | 1 |
| mPool5_V2MM_230540 | 1.268193 | 75.83547 | TRUE | HP_48252 | TGCTGTTG | GATGTGT( | AK079812 | RIKEN cDNA 9130227L01 gene | 9130227L01Rik | NA | | | | NA | |
| 092308m1_V2MM_127737 | 1.266666 | 3.004939 | TRUE | HP_34855 | TGCTGTTG | GTGACTA/ | XM_145359 | Mus musculus similar to vomeronasal 1 receptor, G1 [Mus musculus] (LOC23286 | | | 2.6 | SM2114 | c | | 4 |
| 092308m1_V2MM_127737 | 1.266666 | 3.004939 | TRUE | HP_34855 | TGCTGTTG | GTGACTA/ | XM_145359 | Mus musculus similar to vomerosal 1 receptor, G1 [Mus musculus] (LOC232868), | | | 2.6 | SM2114 | c | | 4 |
| mPool4_V2MM_125960 | 1.266542 | 75.83547 | TRUE | HP_34680 | TGCTGTTG | CTACTGAT | XM_144307 | Mus musculus similar to 40S ribosomal protein S2 (LOC231294), mRNA. | | | 2.14 | SM2538 | b | | 11 |
| mPool2_V2MM_124154 | 1.266422 | 3.004939 | TRUE | HP_34500 | TGCTGTTG | GAAAGTA/ | XM_143326 | Mus musculus LOC229387 (LOC229387), mRNA. | | | 2.11 | SM2371 | b | | 5 |
| mPool7_V2MM_179995 | 1.265819 | 3.004939 | TRUE | HP_40046 | TGCTGTTG | CATATTTA | XM_288432 | Mus musculus LOC331767 (LOC331767), mR. | | | 2.5 | SM2069 | a | | 2 |
| mPool7_V2MM_179995 | 1.265819 | 3.004939 | TRUE | HP_40046 | TGCTGTTG | CATATTTA | XM_288432 | Mus musculus LOC331767 (LOC331767), mRNA. | | | 2.5 | SM2069 | a | | 2 |
| mPool5_V2MM_52592 | 1.265314 | 62.64426 | TRUE | HP_27496 | TGCTGTTG | GAGTATGT | NM_008970 | parathyroid hormone-like peptide | Pthlh | NA | | | | NA | |
| mPool2_V2MM_99002 | 1.264925 | 2.205692 | TRUE | HP_32021 | TGCTGTTG | GTCCAATT | AK013759 | RIKEN cDNA 2900069M18 gene | 2900069M18Rik | | 2.6 | SM2147 | a | | 7 |
| 092308m1_V2MM_181021 | 1.264763 | 3.004939 | TRUE | HP_40149 | TGCTGTTG | CAGATTAC | XM_288605 | Mus musculus LOC332003 (LOC332003), mRNA. | | | 2.5 | SM2095 | d | | 3 |
| 092308m1_V2MM_181021 | 1.264763 | 3.004939 | TRUE | HP_40149 | TGCTGTTG | CAGATTAC | XM_288605 | Mus musculus LOC332003 (LOC332003), mR. | | | 2.5 | SM2095 | d | | 3 |
| mPool5_V2MM_211 | 1.26442 | 75.83547 | TRUE | HP_22388 | TGCTGTTG | CAATGAA( | NM_008211 | H3 histone, family 3B | H3f3b | NA | | | | NA | |
| mPool6_V2MM_65669 | 1.264243 | 75.83547 | TRUE | HP_28770 | TGCTGTTG | GGCTGCA( | NM_172908 | RIKEN cDNA 9230106D23 gene | 9230106D23Rik | | 2.7 | SM2181 | g | | 3 |

| | | | | | | | | | | | | | |
|---|---|---|---|---|---|---|---|---|---|---|---|---|---|
| mPool5_V2MM_159885 | 1.264116 | 75.83547 | TRUE | HP_38047 | TGCTGTTG | CTGGCCTC | XM_283291 | Mus musculus hypothetical gene supported by AK027976 (LOC328554), mRNA. | | NA | | | NA |
| 092308m3_V2MM_103042 | 1.26304 | 75.83547 | TRUE | HP_32418 | TGCTGTTG | GATACCGT | XM_484662 | WD repeat domain 43 | Wrd43 | 2.13 | SM2490 | f | 6 |
| mPool7_V2MM_76348 | 1.263026 | 2.205692 | TRUE | HP_29810 | TGCTGTTG | CGCCTTCT | NM_011351 | sema domain, transmembrane domain (TM), and cytopla | Sema6c | 2.8 | SM2234 | h | 3 |
| mPool2_V2MM_163010 | 1.262763 | 3.004939 | TRUE | HP_38356 | TGCTGTTG | GTAGAAAC | XM_284594 | Mus musculus similar to RIKEN cDNA D130040H23; hypothetical protein D130040 | | 2.6 | SM2130 | g | 3 |
| mPool2_V2MM_118336 | 1.262731 | 3.004939 | TRUE | HP_33927 | TGCTGTTG | GATATACC | XM_139794 | Mus musculus similar to calcium-sensing receptor related protein 3 (LOC224578) | | 2.1 | SM2350 | b | 5 |
| mPool7_V2MM_203135 | 1.262663 | 3.004939 | TRUE | HP_42110 | TGCTGTTG | CCTGATGC | XM_131689 | RIKEN cDNA 4930429A08 gene | 4930429A08Rik | 2.1 | SM2315 | f | 10 |
| 092308m1_V2MM_179069 | 1.261419 | 3.004939 | TRUE | HP_39954 | TGCTGTTG | CTATTAAA | XM_288269 | Mus musculus LOC332542 (LOC332542), mRNA. | | 2.5 | SM2091 | c | 7 |
| 092308m1_V2MM_179069 | 1.261419 | 3.004939 | TRUE | HP_39954 | TGCTGTTG | CTATTAAA | XM_288269 | Mus musculus LOC332542 (LOC332542), mR. | | 2.5 | SM2091 | c | 7 |
| 092308m3_V2MM_91853 | 1.260797 | 75.83547 | TRUE | HP_31322 | TGCTGTTG | CTGCAATT | NM_178605 | D segment, Chr 13, Wayne State University 177, expresse | D13Wsu177e | 2.13 | SM2458 | f | 8 |
| 092308m3_V2MM_93732 | 1.260601 | 75.83547 | TRUE | HP_31503 | TGCTGTTG | CGAAATGA | NM_181748 | G protein-coupled receptor 120 | Gpr120 | 2.9 | SM2263 | e | 11 |
| 092308m3_V2MM_93732 | 1.260601 | 75.83547 | TRUE | HP_31503 | TGCTGTTG | CGAAATGA | NM_181748 | G protein-coupled receptor 120 | Gpr120 | 2.9 | SM2263 | e | 11 |
| 092308m3_V2MM_38908 | 1.26049 | 62.64426 | TRUE | HP_26161 | TGCTGTTG | GCCTTGGA | NM_146242 | leucine rich repeat containing 10 | Lrrc10 | 2.12 | SM2421 | a | 6 |
| 092308m3_V2MM_194705 | 1.260414 | 75.83547 | TRUE | HP_41285 | TGCTGTTG | GAAATCAC | NM_178119 | centaurin, gamma 2 | Centg2 | 2.13 | SM2478 | b | 9 |
| mPool2_V2MM_215236 | 1.259445 | 2.205692 | TRUE | HP_43279 | TGCTGTTG | CCTGTTAC | XM_145781 | Mus musculus similar to olfactory receptor MOR23-1 [Mus musculus] (LOC23360 | | 2.11 | SM2366 | b | 1 |
| 092308m3_V2MM_148738 | 1.259292 | 75.83547 | TRUE | HP_36944 | TGCTGTTG | CCAAGTGT | XM_164093 | Mus musculus LOC244445 (LOC244445), mR. | | 2.14 | SM2517 | d | 4 |
| mPool7_V2MM_77900 | 1.259168 | 2.205692 | TRUE | HP_29960 | TGCTGTTG | GAAACGTC | NM_029362 | RIKEN cDNA 2010012F05 gene | 2010012F05Rik | 2.8 | SM2248 | c | 5 |
| mPool7_V2MM_93273 | 1.258875 | 3.004939 | TRUE | HP_31460 | TGCTGTTG | GTCCTTTG | NM_180958 | RIKEN cDNA 4930532D21 gene | 4930532D21Rik | 2.9 | SM2269 | a | 12 |
| 092308m1_V2MM_95054 | 1.258243 | 3.004939 | TRUE | HP_31633 | TGCTGTTG | GCAGTTAA | NM_183313 | hypothetical protein D930018N13 | | 2.6 | SM2113 | h | 7 |
| 092308m1_V2MM_171454 | 1.25819 | 3.004939 | TRUE | HP_39194 | TGCTGTTG | CCAAGTAC | AK048966 | RIKEN cD C230086J09 gene | C230086J09Rik | 2.5 | SM2086 | f | 2 |
| 092308m1_V2MM_122466 | 1.257098 | 3.004939 | TRUE | HP_34333 | TGCTGTTG | GAGAGCT | BC053004 | glycosyltransferase 28 domain containing 1 | Glt28d1 | 2.6 | SM2101 | f | 11 |
| mPool2_V2MM_83049 | 1.257038 | 2.205692 | TRUE | HP_30460 | TGCTGTTG | CTGGGAAT | NM_011267 | regulator of G-protein signaling 16 | Rgs16 | 2.6 | SM2138 | a | 4 |
| 092308m1_V2MM_167663 | 1.256768 | 3.004939 | TRUE | HP_38817 | TGCTGTTG | GGCTAAGT | XM_285812 | Mus musculus similar to protease [Mus musculus] (LOC329865), mR. | | 2.6 | SM2124 | g | 11 |
| mPool7_V2MM_170526 | 1.256519 | 2.205692 | TRUE | HP_39101 | TGCTGTTG | GTGTCTAC | AK034287 | leucine rich repeat and fibronectin type III domain contai | Lrfn5 | 2.9 | SM2276 | a | 8 |
| 092308m3_V2MM_60972 | 1.255802 | 75.83547 | TRUE | HP_28311 | TGCTGTTG | GTCCTGAA | NM_009503 | valosin containing protein | Vcp | 2.12 | SM2412 | b | 2 |
| mPool2_V2MM_107743 | 1.255407 | 2.205692 | TRUE | HP_32882 | TGCTGTTG | CACGGAAA | AK016965 | RIKEN cDNA 4933428F06 gene | 4933428F06Rik | 2.7 | SM2158 | f | 1 |
| 092308m3_V2MM_105467 | 1.255256 | 75.83547 | TRUE | HP_32658 | TGCTGTTG | GTGGTCCA | XM_130523 | D segment, Chr 2, Brigham & Womens Genetics 1356 exp | D2Bwg1356e | 2.14 | SM2508 | e | 11 |
| mPool2_V2MM_157758 | 1.254475 | 3.004939 | TRUE | HP_37837 | TGCTGTTG | CTATAAAT | XM_205565 | RIKEN cDNA 4930526H21 gene | 4930526H21Rik | 2.6 | SM2126 | b | 2 |
| mPool2_V2MM_129889 | 1.254189 | 3.004939 | TRUE | HP_35069 | TGCTGTTG | GAAGAAAT | BC041776 | RIKEN cDNA D630040G17 gene | D630040G17Rik | 2.11 | SM2371 | e | 8 |
| 092308m3_V2MM_234254 | 1.25414 | 75.83547 | TRUE | HP_48613 | TGCTGTTG | CCACTTCT | XM_136417 | hypothetical LOC268730 | | 2.13 | SM2499 | h | 1 |
| mPool6_V2MM_155870 | 1.254106 | 75.83547 | TRUE | HP_37650 | TGCTGTTG | CAGGAGA | NM_181857 | DNA polymerase N | Poln | 2.7 | SM2190 | h | 9 |
| mPool2_V2MM_62998 | 1.25371 | 2.205692 | TRUE | HP_28508 | TGCTGTTG | CCGAGACA | NM_008087 | growth arrest specific 2 | Gas2 | 2.7 | SM2166 | d | 5 |
| 092308m3_V2MM_38939 | 1.253641 | 35.99881 | TRUE | HP_26164 | TGCTGTTG | GCGACTCT | NM_007996 | ferredoxin 1 | Fdx1 | 2.11 | SM2394 | a | 10 |
| 092308m1_V2MM_187100 | 1.253581 | 3.004939 | TRUE | HP_40756 | TGCTGTTG | CTGAGACA | XM_289649 | Mus musculus LOC333411 (LOC333411), mR. | | 2.5 | SM2070 | d | 2 |
| mPool7_V2MM_79004 | 1.253445 | 3.004939 | TRUE | HP_30067 | TGCTGTTG | GAGTATTC | NM_025479 | RIKEN cDNA 2810021B07 gene | 2810021B07Rik | 2.8 | SM2246 | h | 7 |
| mPool2_V2MM_120546 | 1.253206 | 3.004939 | TRUE | HP_34144 | TGCTGTTG | GTGAAATT | XM_140991 | Mus musculus similar to ribosomal protein S6 [Gallus gallus] (LOC241487), mRNA | | 2.11 | SM2354 | c | 10 |
| mPool7_V2MM_109298 | 1.252512 | 3.004939 | TRUE | HP_33036 | TGCTGTTG | GAAGAGA | XM_355946 | RIKEN cDNA C030048B12 gene | C030048B12Rik | 2.1 | SM2310 | b | 8 |
| mPool2_V2MM_212633 | 1.252406 | 3.004939 | TRUE | HP_43029 | TGCTGTTG | GGAACGTA | XM_140116 | similar to ribosomal protein S24 | | 2.11 | SM2352 | h | 9 |
| mPool7_V2MM_75847 | 1.252022 | 3.004939 | TRUE | HP_29761 | TGCTGTTG | CCGGATAT | NM_033560 | DNA segment, Chr 8, ERATO Doi 531, expressed | D8Ertd531e | 2.8 | SM2236 | e | 9 |
| mPool7_V2MM_98250 | 1.251865 | 3.004939 | TRUE | HP_31948 | TGCTGTTG | GGACAAG | XM_112298 | similar to RIKEN cDNA 1700029I01 | | 2.9 | SM2258 | f | 5 |
| 092308m3_V2MM_45265 | 1.2518 | 59.99801 | TRUE | HP_26780 | TGCTGTTG | GGTCAAGC | NM_010155 | Ets2 repressor factor | Erf | 2.12 | SM2437 | a | 7 |
| 092308m1_V2MM_74261 | 1.251215 | 2.205692 | TRUE | HP_29606 | TGCTGTTG | CACAGTAC | NM_183120 | RIKEN cD 1110048D14 gene | 1110048D14Rik | 2.4 | SM2024 | h | 2 |
| 092308m1_V2MM_62780 | 1.251181 | 3.004939 | TRUE | HP_28487 | TGCTGTTG | CATCTTAT | NM_146748 | olfactory receptor 661 | Olfr661 | 2.4 | SM2026 | f | 8 |
| mPool4_V2MM_225529 | 1.250858 | 75.83547 | TRUE | HP_47757 | TGCTGTTG | CTTTGGTC | NM_178386 | RIKEN cDNA 1700034J06 gene | 1700034J06Rik | 2.16 | SM2605 | b | 12 |
| 092308m3_V2MM_236006 | 1.250853 | 62.64426 | TRUE | HP_48786 | TGCTGTTG | CTTCCCAG | XM_150399 | Mus musculus LOC208932 (LOC208932), mR. | | 2.14 | SM2519 | a | 6 |
| 092308m1_V2MM_61824 | 1.250278 | 3.004939 | TRUE | HP_28394 | TGCTGTTG | CAATCAGA | NM_176900 | RIKEN cD C730034F03 gene | C730034F03Rik | 2.3 | SM2009 | f | 11 |
| mPool2_V2MM_9768 | 1.250272 | 3.004939 | TRUE | HP_23321 | TGCTGTTG | CATTGGAA | NM_008444 | kinesin family member 3B | Kif3b | 2.11 | SM2385 | g | 1 |
| mPool2_V2MM_9754 | 1.25027 | 3.004939 | TRUE | HP_23320 | TGCTGTTG | CATTCCCT | NM_011660 | thioredoxin 1 | Txn1 | 2.11 | SM2387 | f | 1 |
| mPool2_V2MM_111510 | 1.249649 | 3.004939 | TRUE | HP_33252 | TGCTGTTG | CGCTGAAC | AK018320 | RIKEN cDNA 6530403M18 gene | 6530403M18Rik | 2.7 | SM2153 | d | 10 |
| mPool2_V2MM_18604 | 1.249593 | 3.004939 | TRUE | HP_24185 | TGCTGTTG | GCTATTGC | NM_144559 | Fc receptor-like 3 | Fcrl3 | 2.11 | SM2385 | f | 5 |
| 092308m3_V2MM_143023 | 1.249406 | 75.83547 | TRUE | HP_36373 | TGCTGTTG | GTGAAGAC | XM_158268 | Mus musculus LOC241344 (LOC241344), mR. | | 2.14 | SM2526 | f | 9 |
| mPool7_V2MM_96180 | 1.249143 | 3.004939 | TRUE | HP_31744 | TGCTGTTG | CCTATTAT | XM_110248 | F-box protein 11 | Fbxo11 | 2.9 | SM2268 | a | 8 |
| 092308m3_V2MM_196430 | 1.247642 | 75.83547 | TRUE | HP_41453 | TGCTGTTG | CACTGTTT | NM_183311 | hypothetical protein B930076A02 | | 2.13 | SM2477 | f | 4 |
| 092308m1_V2MM_125452 | 1.247425 | 2.205692 | TRUE | HP_34629 | TGCTGTTG | CTGTCTAA | XM_144012 | Mus musculus similar to beta-adaptin - rat (LOC230638), mRNA. | | 2.6 | SM2102 | h | 5 |

| | | | | | | | | | | | | | |
|---|---|---|---|---|---|---|---|---|---|---|---|---|---|
| 092308m1_V2MM_125452 | 1.247425 | 2.205692 | TRUE | HP_34629 | TGCTGTTG | CTGTCTAA | XM_144012 | Mus musculus similar to beta-adaptin - rat (LOC230638), mR. | | 2.6 | SM2102 | h | 5 |
| mPool2_V2MM_213180 | 1.247231 | 3.004939 | TRUE | HP_43082 | TGCTGTTG | CTACTTAA | XM_142166 | Mus musculus similar to ribosomal protein S7 (LOC245623), mRNA. | | 2.11 | SM2373 | a | 9 |
| 092308m1_V2MM_170513 | 1.247013 | 1.959154 | TRUE | HP_39100 | TGCTGTTG | GCATAGTT | XM_488637 | hypothetical gene supported by AK087847 | | 2.5 | SM2062 | d | 7 |
| mPool4_V2MM_30524 | 1.24668 | 75.83547 | TRUE | HP_25344 | TGCTGTTG | CGCTTTGC | NM_024177 | mitochondrial ribosomal protein L38 | Mrpl38 | 2.15 | SM2572 | f | 6 |
| 092308m3_V2MM_232423 | 1.246578 | 75.83547 | TRUE | HP_48436 | TGCTGTTG | CATATACT | XM_111313 | similar to small nuclear R | | 2.13 | SM2500 | h | 12 |
| mPool2_V2MM_169983 | 1.246037 | 3.004939 | TRUE | HP_39047 | TGCTGTTG | CTCATAAT | XM_286370 | Mus musculus hypothetical gene supported by AK086479 (LOC327911), mRNA. | | 2.6 | SM2128 | g | 5 |
| 092308m3_V2MM_89430 | 1.245931 | 62.64426 | TRUE | HP_16585 | TGCTGTTG | CTTGTTGA | NM_177595 | RIKEN cD 9430023B20 gene | 9430023B20Rik | 2.13 | SM2456 | h | 12 |
| 092308m3_V2MM_233661 | 1.245839 | 70.87591 | TRUE | HP_48556 | TGCTGTTG | CCGAAGCA | XM_133622 | hepatitis B virus x associated protein | Hbxap | 2.13 | SM2498 | h | 4 |
| mPool2_V2MM_12638 | 1.245531 | 3.004939 | TRUE | HP_14376 | TGCTGTTG | CACAGTGT | NM_011593 | tissue inhibitor of metalloproteinase 1 | Timp1 | 2.11 | SM2380 | g | 11 |
| 092308m3_V2MM_30237 | 1.244755 | 75.83547 | TRUE | HP_25317 | TGCTGTTG | CATCTGGA | BC065171 | CCR4-NOT transcription complex, subunit 2 | Cnot2 | 2.12 | SM2424 | c | 5 |
| 092308m1_V2MM_63866 | 1.24412 | 3.004939 | TRUE | HP_28593 | TGCTGTTG | CTCCACCT | NM_025786 | RIKEN cD 9130020G10 gene | 9130020G10Rik | 2.3 | SM2016 | e | 7 |
| mPool2_V2MM_15761 | 1.243947 | 2.205692 | TRUE | HP_23907 | TGCTGTTG | CTGCTAAT | NM_008372 | interleukin 7 receptor | Il7r | 2.11 | SM2379 | f | 6 |
| mPool2_V2MM_132868 | 1.243817 | 3.004939 | TRUE | HP_35361 | TGCTGTTG | GGAAGGA | AK015981 | RIKEN cDNA 4930535F04 gene | 4930535F04Rik | 2.11 | SM2371 | c | 6 |
| mPool4_V2MM_226810 | 1.243374 | 61.29828 | TRUE | HP_47881 | TGCTGTTG | CCTAAGCA | NM_177827 | hypothetical protein 9530053H22 | | 2.16 | SM2624 | h | 4 |
| mPool7_V2MM_93408 | 1.243225 | 3.004939 | TRUE | HP_31472 | TGCTGTTG | CTGCAATT | NM_181392 | DNA segment, Chr 17, Wayne State University 155, expre | D17Wsu155e | 2.9 | SM2257 | e | 2 |
| 092308m3_V2MM_234885 | 1.24206 | 75.83547 | TRUE | HP_48675 | TGCTGTTG | CTATTGTG | XM_153848 | Mus musculus LOC238501 (LOC238501), mR. | | 2.14 | SM2514 | c | 1 |
| 092308m1_V2MM_177109 | 1.241297 | 3.004939 | TRUE | HP_39758 | TGCTGTTG | GTGTAAAT | XM_287805 | Mus musculus hypothetical gene supported by AK032925 (LOC330762), mR. | | 2.5 | SM2063 | h | 6 |
| 092308m1_V2MM_177109 | 1.241297 | 3.004939 | TRUE | HP_39758 | TGCTGTTG | GTGTAAAT | XM_287805 | Mus musculus hypothetical gene supported by AK032925 (LOC330762), mRNA. | | 2.5 | SM2063 | h | 6 |
| mPool4_V2MM_47449 | 1.241049 | 75.83547 | TRUE | HP_26993 | TGCTGTTG | CTGCCATT | NM_175441 | RIKEN cDNA D830007F02 gene | D830007F02Rik | 2.16 | SM2622 | d | 6 |
| mPool7_V2MM_204911 | 1.240922 | 3.004939 | TRUE | HP_42279 | TGCTGTTG | CCAAGAAC | XM_138950 | Mus musculus LOC239003 (LOC239003), mRNA. | | 2.1 | SM2306 | b | 9 |
| 092308m3_V2MM_48991 | 1.240044 | 75.83547 | TRUE | HP_27144 | TGCTGTTG | GCTCCAGT | NM_145927 | farnesyltransferase, CAAX box, beta | Fntb | 2.12 | SM2446 | f | 2 |
| mPool6_V2MM_46500 | 1.240012 | 59.99801 | TRUE | HP_26900 | TGCTGTTG | CCCTGCTC | NM_153805 | protein kinase N3 | Pkn3 | 2.7 | SM2186 | a | 1 |
| mPool6_V2MM_166521 | 1.239925 | 73.80895 | TRUE | HP_38704 | TGCTGTTG | CATAAGCA | XM_285524 | gene model 1539, (NCBI) | Gm1539 | 2.7 | SM2192 | d | 12 |
| mPool2_V2MM_100265 | 1.239674 | 3.004939 | TRUE | HP_32145 | TGCTGTTG | CCGCCATC | BC025575 | cDNA sequence BC025575 | BC025575 | 2.7 | SM2160 | h | 8 |
| 092308m1_V2MM_142060 | 1.239671 | 3.004939 | TRUE | HP_36277 | TGCTGTTG | GATGTTCA | XM_157294 | Mus musculus LOC240262 (LOC240262), mR. | | 2.4 | SM2032 | d | 8 |
| mPool2_V2MM_139300 | 1.239618 | 3.004939 | TRUE | HP_36002 | TGCTGTTG | GACAGCAT | XM_154905 | Mus musculus LOC210787 (LOC210787), mRNA. | | 2.1 | SM2334 | h | 12 |
| mPool7_V2MM_205425 | 1.238526 | 3.004939 | TRUE | HP_42328 | TGCTGTTG | CCGAGAGT | XM_128377 | RIKEN cDNA 4921511H13 gene | 4921511H13Rik | 2.1 | SM2303 | b | 3 |
| mPool5_V2MM_180385 | 1.238283 | 66.95036 | TRUE | HP_40085 | TGCTGTTG | CACAGTCC | XM_288498 | Mus musculus LOC328105 (LOC328105), mRNA. | | NA | | | NA |
| 092308m1_V2MM_69811 | 1.238273 | 3.004939 | TRUE | HP_29173 | TGCTGTTG | GTAGGGA | NM_010459 | homeo box B4 | Hoxb4 | 2.3 | SM2005 | g | 7 |
| mPool2_V2MM_151163 | 1.236129 | 3.004939 | TRUE | HP_37183 | TGCTGTTG | GGGATGA | NM_001004364 | gene model 592, (NCBI) | Gm592 | 2.1 | SM2343 | a | 12 |
| 092308m3_V2MM_203136 | 1.236094 | 75.83547 | TRUE | HP_42110 | TGCTGTTG | GCCTCATT | XM_129935 | Mus musculus similar to nuclear pore-targeting complex component of 58 kDa (L | | 2.9 | SM2299 | c | 4 |
| 092308m3_V2MM_203136 | 1.236094 | 75.83547 | TRUE | HP_42110 | TGCTGTTG | GCCTCATT | XM_129935 | Mus musculus similar to nuclear pore-targeting complex component of 58 kDa (L | | 2.9 | SM2299 | c | 4 |
| mPool6_V2MM_167391 | 1.235898 | 75.83547 | TRUE | HP_38790 | TGCTGTTG | GACTACAT | XM_285759 | Mus musculus similar to tumor protein, translationally-controlled 1 [Rattus norve | | 2.7 | SM2194 | b | 5 |
| mPool2_V2MM_94842 | 1.235321 | 3.004939 | TRUE | HP_31612 | TGCTGTTG | CCCAGTTG | NM_183269 | RIKEN cDNA 4930511N19 gene | 4930511N19Rik | 2.6 | SM2136 | b | 9 |
| 092308m1_V2MM_151402 | 1.234206 | 2.205692 | TRUE | HP_37207 | TGCTGTTG | CCATAGCT | XM_194738 | Mus musculus similar to ATP synthase, H+ transporting, mitochondrial F0 comple | | 2.5 | SM2059 | e | 5 |
| 092308m3_V2MM_233077 | 1.234182 | 75.83547 | TRUE | HP_48500 | TGCTGTTG | GTCTGTCT | AK021026 | RIKEN cD B430319H21 gene | B430319H21Rik | 2.13 | SM2500 | d | 1 |
| 092308m1_V2MM_183706 | 1.233168 | 2.205692 | TRUE | HP_40418 | TGCTGTTG | CATGAAAT | XM_289066 | RIKEN cD 1700085A12 gene | 1700085A12Rik | 2.5 | SM2072 | e | 4 |
| mPool6_V2MM_33930 | 1.233143 | 59.99801 | TRUE | HP_25676 | TGCTGTTG | CATGTCAT | NM_011849 | NIMA (never in mitosis gene a)-related expressed kinase | Nek4 | 2.7 | SM2182 | g | 2 |
| 092308m1_V2MM_176986 | 1.233134 | 2.205692 | TRUE | HP_39746 | TGCTGTTG | CATATATA | AK081575 | expressed sequence W91709 | W91709 | 2.5 | SM2083 | b | 1 |
| 092308m1_V2MM_176986 | 1.233134 | 2.205692 | TRUE | HP_39746 | TGCTGTTG | CATATATA | AK081575 | expressed sequence W91709 | W91709 | 2.5 | SM2083 | b | 1 |
| mPool7_V2MM_205147 | 1.233094 | 3.004939 | TRUE | HP_42302 | TGCTGTTG | GAGGATG | XM_127013 | RIKEN cDNA 2600001A11 gene | 2600001A11Rik | 2.9 | SM2300 | d | 4 |
| mPool7_V2MM_79645 | 1.23278 | 3.004939 | TRUE | HP_30130 | TGCTGTTG | GCCACAAC | NM_021887 | interleukin 21 receptor | Il21r | 2.8 | SM2238 | a | 6 |
| mPool7_V2MM_201416 | 1.232656 | 3.004939 | TRUE | HP_41944 | TGCTGTTG | CTGCCAAA | XM_288575 | Mus musculus LOC331967 (LOC331967), mRNA. | | 2.9 | SM2288 | a | 5 |
| mPool4_V2MM_159534 | 1.231956 | 66.95036 | TRUE | HP_38012 | TGCTGTTG | CTGGAGCT | XM_283144 | similar to regulator of sex-limitation candidate 24 | | 2.16 | SM2650 | c | 11 |
| mPool4_V2MM_41824 | 1.231886 | 62.64426 | TRUE | HP_26444 | TGCTGTTG | CAGCCAGC | NM_017400 | SH3-domain GRB2-like 3 | Sh3gl3 | 2.15 | SM2564 | b | 12 |
| mPool2_V2MM_3169 | 1.231545 | 3.004939 | TRUE | HP_22677 | TGCTGTTG | CTTGTTAT | XM_486385 | similar to RIKEN cDNA 2610042L04 | | 2.11 | SM2387 | a | 3 |
| mPool7_V2MM_145089 | 1.231136 | 3.004939 | TRUE | HP_36580 | TGCTGTTG | CTATTCTA | XM_160546 | Mus musculus LOC215301 (LOC215301), mR. | | 2.4 | SM2049 | a | 4 |
| mPool7_V2MM_145089 | 1.231136 | 3.004939 | TRUE | HP_36580 | TGCTGTTG | CTATTCTA | XM_160546 | Mus musculus LOC215301 (LOC215301), mRNA. | | 2.4 | SM2049 | a | 4 |
| 092308m1_V2MM_152312 | 1.231104 | 3.004939 | TRUE | HP_37297 | TGCTGTTG | GTAAGTCA | XM_195257 | Mus musculus similar to Major uriry protein 5 precursor (MUP 5) (LOC272721), m | | 2.4 | SM2031 | e | 5 |
| mPool4_V2MM_47689 | 1.231064 | 75.83547 | TRUE | HP_27016 | TGCTGTTG | CTTATTGG | NM_023868 | ryanodine receptor 2, cardiac | Ryr2 | 2.15 | SM2568 | h | 7 |
| 092308m1_V2MM_176141 | 1.231009 | 3.004939 | TRUE | HP_39662 | TGCTGTTG | CAGTGAAT | XM_485788 | hypothetical protein 9130221F21 | | 2.5 | SM2070 | a | 8 |
| mPool7_V2MM_144281 | 1.230849 | 2.205692 | TRUE | HP_36499 | TGCTGTTG | GCAAAGA | XM_159840 | Mus musculus LOC208254 (LOC208254), mRNA. | | 2.1 | SM2332 | c | 10 |
| mPool2_V2MM_189948 | 1.230282 | 3.004939 | TRUE | HP_34777 | TGCTGTTG | GTCGCCTC | XM_355798 | similar to carbohydrate (chondroitin 4) sulfotransferase 13; chondroitin 4-O-sulfo | | 2.11 | SM2355 | g | 6 |

| | | | | | | | | | | | | | |
|---|---|---|---|---|---|---|---|---|---|---|---|---|---|
| 092308m1_V2MM_160694 | 1.23006 | 2.205692 | TRUE | HP_38127 | TGCTGTTG | CTCCTTTA | XM_283612 | Mus musculus similar to pol polyprotein [Trichosurus vulpecula] (LOC329162), m | | 2.4 | SM2045 | f | 2 |
| mPool2_V2MM_123920 | 1.229957 | 3.004939 | TRUE | HP_34477 | TGCTGTTG | CTCCAGG | XM_143210 | Mus musculus similar to bendless protein [Rattus norvegicus] (LOC241994), mRN | | 2.11 | SM2359 | a | 3 |
| 092308m3_V2MM_225633 | 1.229808 | 59.99801 | TRUE | HP_47767 | TGCTGTTG | CAGAGTT | NM_011030 | procollagen-proline, 2-oxoglutarate 4-dioxygese (proline | P4ha1 | 2.13 | SM2454 | h | 10 |
| 092308m3_V2MM_100826 | 1.229612 | 75.83547 | TRUE | HP_32200 | TGCTGTTG | CATAGTCA | BC055027 | expressed sequence AI132487 | AI132487 | 2.1 | SM2310 | h | 1 |
| 092308m3_V2MM_100826 | 1.229612 | 75.83547 | TRUE | HP_32200 | TGCTGTTG | CATAGTCA | BC055027 | expressed sequence AI132487 | AI132487 | 2.1 | SM2310 | h | 1 |
| mPool2_V2MM_136163 | 1.229573 | 3.004939 | TRUE | HP_35688 | TGCTGTTG | CAGCTTAC | XM_151461 | Mus musculus LOC208337 (LOC208337), mRNA. | | 2.1 | SM2336 | d | 2 |
| mPool2_V2MM_127709 | 1.228634 | 2.205692 | TRUE | HP_34853 | TGCTGTTG | CGAAAGG | XM_145347 | Mus musculus similar to Zinc finger protein 134 (LOC209850), mRNA. | | 2.11 | SM2352 | g | 7 |
| 092308m3_V2MM_54981 | 1.228278 | 75.83547 | TRUE | HP_27729 | TGCTGTTG | CCCACTGC | NM_177027 | RIKEN cD 4930572I07 gene | 4930572I07Rik | 2.12 | SM2442 | b | 7 |
| mPool2_V2MM_37833 | 1.227882 | 2.205692 | TRUE | HP_26057 | TGCTGTTG | GAGGAAA | NM_031374 | testis expressed gene 15 | Tex15 | 2.11 | SM2392 | g | 1 |
| mPool4_V2MM_101480 | 1.227184 | 75.83547 | TRUE | HP_32265 | TGCTGTTG | CTGTGTG | XM_127657 | RIKEN cDNA 2310032M22 gene | | 2.16 | SM2606 | f | 12 |
| 092308m3_V2MM_90023 | 1.227109 | 75.83547 | TRUE | HP_31144 | TGCTGTTG | GACCATTC | AK030707 | expressed sequence AI836003 | AI836003 | 2.13 | SM2470 | c | 10 |
| mPool5_V2MM_11148 | 1.22701 | 61.29828 | TRUE | HP_23456 | TGCTGTTG | CTTGATGA | NM_007655 | CD79A antigen (immunoglobulin-associated alpha) | Cd79a | NA | | | NA |
| 092308m3_V2MM_145141 | 1.226871 | 75.83547 | TRUE | HP_36585 | TGCTGTTG | CTACTCTG | XM_160598 | Mus musculus LOC242260 (LOC242260), mR. | | 2.14 | SM2510 | b | 5 |
| mPool2_V2MM_8808 | 1.226809 | 3.004939 | TRUE | HP_23228 | TGCTGTTG | CAAAGTAT | NM_007646 | CD38 antigen | Cd38 | 2.11 | SM2375 | h | 2 |
| mPool2_V2MM_120784 | 1.225891 | 3.004939 | TRUE | HP_34167 | TGCTGTTG | CCCAGTAA | XM_485031 | olfactory receptor 48 | Olfr48 | 2.11 | SM2359 | b | 1 |
| 092308m3_V2MM_84260 | 1.225693 | 75.83547 | TRUE | HP_30577 | TGCTGTTG | CTTTCTAT | NM_027405 | RIKEN cD 1700020O03 gene | 1700020O03Rik | 2.13 | SM2473 | h | 2 |
| 092308m3_V2MM_232758 | 1.225394 | 35.99881 | TRUE | HP_48469 | TGCTGTTG | CACCTTTA | AK017151 | RIKEN cD 5033404E19 gene | 5033404E19Rik | 2.13 | SM2490 | g | 2 |
| mPool7_V2MM_186924 | 1.22426 | 3.004939 | TRUE | HP_40739 | TGCTGTTG | CTGCAAAT | XM_289619 | Mus musculus LOC333363 (LOC333363), mR. | | 2.5 | SM2068 | c | 10 |
| mPool7_V2MM_186924 | 1.22426 | 3.004939 | TRUE | HP_40739 | TGCTGTTG | CTGCAAAT | XM_289619 | Mus musculus LOC333363 (LOC333363), mRNA. | | 2.5 | SM2068 | c | 10 |
| mPool7_V2MM_199863 | 1.224071 | 3.004939 | TRUE | HP_41790 | TGCTGTTG | CTGTCATT | AK048402 | RIKEN cDNA C130057M05 gene | C130057M05Rik | 2.9 | SM2291 | b | 11 |
| 092308m1_V2MM_174282 | 1.223776 | 2.205692 | TRUE | HP_39476 | TGCTGTTG | CACCAGG | XM_287196 | Mus musculus hypothetical gene supported by AK034924 (LOC329550), mR. | | 2.5 | SM2084 | c | 1 |
| 092308m1_V2MM_174282 | 1.223776 | 2.205692 | TRUE | HP_39476 | TGCTGTTG | CACCAGG | XM_287196 | Mus musculus hypothetical gene supported by AK034924 (LOC329550), mRNA. | | 2.5 | SM2084 | c | 1 |
| 092308m1_V2MM_62732 | 1.22372 | 2.205692 | TRUE | HP_28483 | TGCTGTTG | CATATTTC | XM_143364 | Mus musculus similar to pyridoxine kise [Plasmodium falciparum 3D7] (LOC2135 | | 2.3 | SM2014 | c | 5 |
| 092308m1_V2MM_63001 | 1.223661 | 1.959154 | TRUE | HP_28509 | TGCTGTTG | CCGAGGA | NM_172920 | RIKEN cD 1100001I19 gene | 1100001I19Rik | 2.3 | SM2005 | b | 8 |
| 092308m1_V2MM_171804 | 1.222807 | 3.004939 | TRUE | HP_39229 | TGCTGTTG | GCACAAT | XM_286696 | Mus musculus hypothetical LOC328615 (LOC328615), mR. | | 2.5 | SM2090 | a | 10 |
| mPool4_V2MM_194251 | 1.222462 | 70.87591 | TRUE | HP_41242 | TGCTGTTG | CTCACTTG | XM_487805 | similar to limitin | | 2.16 | SM2607 | b | 4 |
| mPool2_V2MM_10286 | 1.22242 | 3.004939 | TRUE | HP_23372 | TGCTGTTG | CTCACCTA | NM_053224 | vomeronasal 1 receptor, A9 | V1ra9 | 2.11 | SM2381 | d | 7 |
| mPool2_V2MM_165780 | 1.221891 | 3.004939 | TRUE | HP_38631 | TGCTGTTG | GACTTAG | XM_285325 | Mus musculus similar to protease [Mus musculus] (LOC329363), mRNA. | | 2.6 | SM2127 | c | 2 |
| 092308m1_V2MM_175083 | 1.221261 | 3.004939 | TRUE | HP_39556 | TGCTGTTG | GACTGAA | AK003729 | RIKEN cD 1110015M06 gene | 1110015M06Rik | 2.5 | SM2060 | d | 10 |
| 092308m3_V2MM_233770 | 1.220894 | 75.83547 | TRUE | HP_48567 | TGCTGTTG | GTCCAGG | XM_132152 | RIKEN cD 3110007P09 gene | 3110007P09Rik | 2.14 | SM2505 | d | 2 |
| mPool7_V2MM_63636 | 1.220771 | 1.959154 | TRUE | HP_28571 | TGCTGTTG | CTACTACA | NM_146756 | olfactory receptor 608 | Olfr608 | 2.8 | SM2240 | e | 1 |
| 092308m3_V2MM_101123 | 1.22061 | 66.95036 | TRUE | HP_32229 | TGCTGTTG | CGCGGCA | NM_001001321 | solute carrier family 35, member D2 | Slc35d2 | 2.13 | SM2490 | f | 5 |
| mPool2_V2MM_212105 | 1.220542 | 3.004939 | TRUE | HP_42978 | TGCTGTTG | GGCATTG | XM_146652 | Mus musculus similar to protease [Mus musculus] (LOC234930), mRNA. | | 2.11 | SM2356 | g | 2 |
| 092308m3_V2MM_52597 | 1.219206 | 66.95036 | TRUE | HP_27497 | TGCTGTTG | GAGTGAA | NM_008966 | prostaglandin F receptor | Ptgfr | 2.12 | SM2446 | f | 10 |
| mPool7_V2MM_107197 | 1.217956 | 3.004939 | TRUE | HP_32828 | TGCTGTTG | CAGCCAA | AK017309 | peroxisome biogenesis factor 1 | Pex1 | 2.7 | SM2154 | g | 3 |
| mPool7_V2MM_178288 | 1.217727 | 3.004939 | TRUE | HP_39876 | TGCTGTTG | CAGAATT | XM_288127 | Mus musculus hypothetical gene supported by AK053873 (LOC331495), mRNA. | | 2.9 | SM2272 | e | 12 |
| mPool4_V2MM_121736 | 1.217604 | 75.83547 | TRUE | HP_34261 | TGCTGTTG | CATGTCTC | XM_141818 | Mus musculus similar to 60S RIBOSOMAL PROTEIN L18 (LOC236846), mRNA. | | 2.15 | SM2551 | g | 5 |
| mPool7_V2MM_78128 | 1.217157 | 3.004939 | TRUE | HP_29982 | TGCTGTTG | GAATGAA | NM_025418 | RIKEN cDNA 1110059P08 gene | 1110059P08Rik | 2.8 | SM2245 | a | 5 |
| 092308m3_V2MM_236535 | 1.217052 | 75.83547 | TRUE | HP_48839 | TGCTGTTG | CTGACATT | XM_157329 | Mus musculus LOC225501 (LOC225501), mR. | | 2.14 | SM2512 | d | 11 |
| mPool5_V2MM_34126 | 1.216719 | 59.99801 | TRUE | HP_25695 | TGCTGTTG | CCACTGCT | NM_009321 | tubulin cofactor a | Tbca | NA | | | NA |
| 092308m3_V2MM_42114 | 1.216047 | 75.83547 | TRUE | HP_26473 | TGCTGTTG | CAGTGTGT | NM_009541 | zinc finger and BTB domain containing 17 | Zbtb17 | 2.12 | SM2412 | b | 1 |
| 092308m1_V2MM_141173 | 1.215995 | 3.004939 | TRUE | HP_36188 | TGCTGTTG | GGAAACA | XM_156493 | Mus musculus LOC239875 (LOC239875), mR. | | 2.4 | SM2033 | h | 3 |
| mPool7_V2MM_178556 | 1.21597 | 3.004939 | TRUE | HP_39903 | TGCTGTTG | GTGTCATC | XM_288181 | Mus musculus LOC332445 (LOC332445), mRNA. | | 2.5 | SM2065 | e | 4 |
| mPool7_V2MM_178556 | 1.21597 | 3.004939 | TRUE | HP_39903 | TGCTGTTG | GTGTCATC | XM_288181 | Mus musculus LOC332445 (LOC332445), mR. | | 2.5 | SM2065 | e | 4 |
| mPool4_V2MM_129839 | 1.215958 | 69.72441 | TRUE | HP_35064 | TGCTGTTG | GAGCTCCT | XM_146477 | gene model 1705, (NCBI) | Gm1705 | 2.14 | SM2549 | a | 7 |
| 092308m3_V2MM_57096 | 1.215591 | 75.83547 | TRUE | HP_27936 | TGCTGTTG | CTGTCTGC | NM_153585 | CCR4-NOT transcription complex, subunit 10 | Cnot10 | 2.12 | SM2416 | f | 11 |
| 092308m1_V2MM_86452 | 1.215223 | 3.004939 | TRUE | HP_30792 | TGCTGTTG | CTCTTAAA | NM_172468 | RIKEN cDNA 4732481H14 gene | 4732481H14Rik | 2.6 | SM2113 | b | 1 |
| 092308m1_V2MM_86452 | 1.215223 | 3.004939 | TRUE | HP_30792 | TGCTGTTG | CTCTTAAA | NM_172468 | RIKEN cD 4732481H14 gene | 4732481H14Rik | 2.6 | SM2113 | b | 1 |
| 092308m3_V2MM_115897 | 1.215145 | 70.87591 | TRUE | HP_33685 | TGCTGTTG | GACCAAG | XM_138679 | Mus musculus similar to zinc finger protein 91 (HPF7, HTF10) [Homo sapiens] (LO | | 2.1 | SM2312 | e | 2 |
| 092308m3_V2MM_115897 | 1.215145 | 70.87591 | TRUE | HP_33685 | TGCTGTTG | GACCAAG | XM_138679 | Mus musculus similar to zinc finger protein 91 (HPF7, HTF10) [Homo sapiens] (LO | | 2.1 | SM2312 | e | 2 |
| mPool4_V2MM_214734 | 1.215122 | 75.83547 | TRUE | HP_43230 | TGCTGTTG | CTGGCACT | XM_145127 | Mus musculus similar to set gene (LOC232637), mRNA. | | 2.16 | SM2646 | b | 9 |
| 092308m3_V2MM_103162 | 1.215093 | 75.83547 | TRUE | HP_32430 | TGCTGTTG | GTGCAAA | XM_128854 | RIKEN cD 2900055J20 gene | 2900055J20Rik | 2.14 | SM2506 | h | 12 |
| 092308m1_V2MM_173900 | 1.214873 | 3.004939 | TRUE | HP_39438 | TGCTGTTG | GTAGGAA | XM_287114 | Mus musculus hypothetical gene supported by AK052519 (LOC329402), mR. | | 2.5 | SM2079 | h | 2 |

| | | | | | | | | | | | | | |
|---|---|---|---|---|---|---|---|---|---|---|---|---|---|
| 092308m1_V2MM_173900 | 1.214873 | 3.004939 | TRUE | HP_39438 | TGCTGTTG | GTAGGAA( | XM_287114 | Mus musculus hypothetical gene supported by AK052519 (LOC329402), mRNA. | | 2.5 | SM2079 | h | 2 |
| mPool2_V2MM_109640 | 1.214814 | 1.959154 | TRUE | HP_33069 | TGCTGTTG | GGATTTA( | XM_134026 | RIKEN cDNA 2310051N18 gene | 2310051N18Rik | 2.7 | SM2163 | f | 7 |
| mPool2_V2MM_131800 | 1.213732 | 3.004939 | TRUE | HP_35256 | TGCTGTTG | GGTTAAA( | XM_147845 | Mus musculus LOC219184 (LOC219184), mRNA. | | 2.11 | SM2367 | h | 2 |
| mPool4_V2MM_151462 | 1.213468 | 75.83547 | TRUE | HP_37213 | TGCTGTTG | GTCTCAAT | XM_194774 | Mus musculus similar to RIKEN cDNA 1110001A07 [Mus musculus] (LOC270833), | | 2.16 | SM2650 | b | 3 |
| mPool4_V2MM_186539 | 1.213016 | 61.29828 | TRUE | HP_40701 | TGCTGTTG | GTCATTAT | XM_289555 | Mus musculus LOC333293 (LOC333293), mRNA. | | NA | | | NA |
| 092308m1_V2MM_92813 | 1.212434 | 2.205692 | TRUE | HP_31415 | TGCTGTTG | CTGGCCT( | NM_178780 | RIKEN cD E130307J04 gene | E130307J04Rik | 2.6 | SM2108 | d | 6 |
| mPool7_V2MM_63423 | 1.212083 | 2.205692 | TRUE | HP_28550 | TGCTGTTG | CGGATAG, | BC043139 | RIKEN cDNA 2700082D03 gene | 2700082D03Rik | 2.8 | SM2244 | c | 8 |
| mPool7_V2MM_140674 | 1.211769 | 3.004939 | TRUE | HP_36139 | TGCTGTTG | CACTAGA/ | XM_156197 | Mus musculus LOC208209 (LOC208209), mRNA. | | 2.5 | SM2051 | d | 12 |
| mPool7_V2MM_140674 | 1.211769 | 3.004939 | TRUE | HP_36139 | TGCTGTTG | CACTAGA/ | XM_156197 | Mus musculus LOC208209 (LOC208209), mR. | | 2.5 | SM2051 | d | 12 |
| mPool2_V2MM_210622 | 1.21095 | 3.004939 | TRUE | HP_42836 | TGCTGTTG | GAAGTTA1 | AK015148 | RIKEN cDNA 4930415H17 gene | 4930415H17Rik | 2.11 | SM2356 | b | 12 |
| mPool2_V2MM_21024 | 1.210728 | 3.004939 | TRUE | HP_24420 | TGCTGTTG | CACTGATT | NM_146145 | Janus kinase 1 | Jak1 | 2.11 | SM2392 | e | 12 |
| mPool2_V2MM_109775 | 1.209404 | 3.004939 | TRUE | HP_33082 | TGCTGTTG | GATGTTAT | NM_027973 | myeloid leukemia factor 1 interacting protein | Mlf1ip | 2.7 | SM2155 | g | 9 |
| mPool4_V2MM_120134 | 1.209255 | 75.83547 | TRUE | HP_10688 | TGCTGTTG | CAGTGTG/ | XM_140772 | gene model 340, (NCBI) | Gm340 | 2.14 | SM2550 | b | 3 |
| 092308m1_V2MM_149037 | 1.208607 | 2.205692 | TRUE | HP_36974 | TGCTGTTG | GAGACTT/ | XM_164431 | Mus musculus LOC244521 (LOC244521), mR. | | 2.5 | SM2058 | c | 8 |
| mPool7_V2MM_83050 | 1.208544 | 3.004939 | TRUE | HP_30460 | TGCTGTTG | GATTTGCT | NM_011267 | regulator of G-protein signaling 16 | Rgs16 | 2.8 | SM2250 | h | 7 |
| mPool2_V2MM_132535 | 1.207848 | 2.205692 | TRUE | HP_35328 | TGCTGTTG | GTCTTAA/ | AF032130 | diabetic embryopathy 1 | Dep1 | 2.11 | SM2362 | h | 9 |
| mPool5_V2MM_6244 | 1.205951 | 60.13437 | TRUE | HP_22978 | TGCTGTTG | CTCTGAG1 | NM_007557 | bone morphogenetic protein 7 | Bmp7 | NA | | | NA |
| mPool2_V2MM_30550 | 1.205938 | 3.004939 | TRUE | HP_25347 | TGCTGTTG | CGGAGTA( | AK087545 | heterogeneous nuclear ribonucleoprotein U-like 1 | Hnrpul1 | 2.11 | SM2390 | g | 11 |
| mPool5_V2MM_158870 | 1.204813 | 61.29828 | TRUE | HP_37947 | TGCTGTTG | GATTGTG1 | XM_282897 | Mus musculus hypothetical gene supported by AK032987 (LOC327724), mRNA. | | NA | | | NA |
| 092308m1_V2MM_97001 | 1.204625 | 2.205692 | TRUE | HP_31824 | TGCTGTTG | GCCGTAA1 | XM_111160 | Mus musculus similar to eukaryotic translation initiation factor 4A1; initiation fac | | 2.6 | SM2109 | a | 11 |
| mPool2_V2MM_212790 | 1.204301 | 3.004939 | TRUE | HP_43044 | TGCTGTTG | GAAAGAG | XM_149459 | hypothetical LOC230628 | | 2.11 | SM2371 | f | 12 |
| 092308m1_V2MM_170774 | 1.203418 | 3.004939 | TRUE | HP_39126 | TGCTGTTG | CACTGAT( | AK049051 | engulfment and cell motility 1, ced-12 homolog (C. elega | Elmo1 | 2.5 | SM2089 | a | 4 |
| mPool2_V2MM_696 | 1.203314 | 2.205692 | TRUE | HP_22435 | TGCTGTTG | CAGAGTT1 | NM_144552 | syntaxin binding protein 6 (amisyn) | Stxbp6 | 2.11 | SM2380 | g | 6 |
| mPool4_V2MM_192385 | 1.203062 | 75.83547 | TRUE | HP_41118 | TGCTGTTG | CCCGGAG( | NM_198959 | hypocretin (orexin) receptor 1 | Hcrtr1 | 2.16 | SM2626 | h | 8 |
| mPool2_V2MM_2758 | 1.2025 | 3.004939 | TRUE | HP_22637 | TGCTGTTG | CTGGAGA1 | NM_172120 | vacuolar protein sorting 41 (yeast) | Vps41 | 2.11 | SM2385 | b | 3 |
| mPool2_V2MM_104021 | 1.20204 | 3.004939 | TRUE | HP_32515 | TGCTGTTG | CACTTTAA | XM_129443 | protein tyrosine phosphatase, receptor type, f polypepti( | Ppfia4 | 2.7 | SM2159 | f | 4 |
| mPool2_V2MM_94190 | 1.202007 | 3.004939 | TRUE | HP_31548 | TGCTGTTG | GTATTTGT | NM_183100 | RIKEN cDNA 5430414B19 gene | 5430414B19Rik | 2.6 | SM2146 | h | 6 |
| 092308m3_V2MM_54088 | 1.201759 | 60.13437 | TRUE | HP_27642 | TGCTGTTG | CAGATGA/ | XM_486633 | similar to Xlr-related, meiosis regulated | | 2.12 | SM2441 | c | 1 |
| 092308m1_V2MM_142075 | 1.200922 | 3.004939 | TRUE | HP_36278 | TGCTGTTG | GTACAAC1 | XM_157299 | Mus musculus LOC240266 (LOC240266), mRNA. | | 2.4 | SM2046 | a | 9 |
| 092308m1_V2MM_142075 | 1.200922 | 3.004939 | TRUE | HP_36278 | TGCTGTTG | GTACAAC1 | XM_157299 | Mus musculus LOC240266 (LOC240266), mR. | | 2.4 | SM2046 | a | 9 |
| 092308m1_V2MM_168480 | 1.200464 | 3.004939 | TRUE | HP_38898 | TGCTGTTG | CCTATACC | XM_286000 | Mus musculus similar to zinc finger protein 93 homolog; zinc finger protein homo | | 2.6 | SM2114 | h | 7 |
| 092308m1_V2MM_88753 | 1.200049 | 3.004939 | TRUE | HP_31018 | TGCTGTTG | CGGATAG, | AK028335 | ubiquitin specific protease 28 | Usp28 | 2.6 | SM2112 | g | 9 |
| 092308m1_V2MM_88753 | 1.200049 | 3.004939 | TRUE | HP_31018 | TGCTGTTG | CGGATAG, | AK028335 | ubiquitin specific protease 28 | Usp28 | 2.6 | SM2112 | g | 9 |
| 092308m1_V2MM_86955 | 1.200042 | 3.004939 | TRUE | HP_30842 | TGCTGTTG | CATTGCA/ | AK083909 | RIKEN cD C030011O14 gene | C030011O14Rik | 2.6 | SM2112 | f | 11 |
| mPool5_V2MM_199116 | 1.199548 | 75.83547 | TRUE | HP_41716 | TGCTGTTG | CTGTGAC1 | XM_289259 | Mus musculus LOC332905 (LOC332905), mRNA. | | NA | | | NA |
| mPool6_V2MM_154058 | 1.199513 | 35.99881 | TRUE | HP_37470 | TGCTGTTG | CCTGTAA( | BC056393 | RIKEN cDNA 2600009P04 gene | 2600009P04Rik | 2.7 | SM2191 | h | 6 |
| mPool2_V2MM_89236 | 1.19948 | 2.205692 | TRUE | HP_31066 | TGCTGTTG | GGATGAA( | BC056493 | G protein-coupled receptor 123 | Gpr123 | 2.6 | SM2146 | h | 7 |
| mPool6_V2MM_163145 | 1.19934 | 61.28368 | TRUE | HP_38370 | TGCTGTTG | CAGACAA/ | AK054302 | expressed sequence C87414 | C87414 | 2.8 | SM2202 | e | 6 |
| 092308m3_V2MM_27305 | 1.198866 | 75.83547 | TRUE | HP_25032 | TGCTGTTG | GAAACTA( | NM_028193 | BRF1 homolog, subunit of R polymerase III transcription i | Brf1 | 2.11 | SM2399 | h | 1 |
| mPool2_V2MM_107820 | 1.198858 | 2.205692 | TRUE | HP_32890 | TGCTGTTG | CACTTACT | AK006923 | RIKEN cDNA 1700069L16 gene | 1700069L16Rik | 2.1 | SM2323 | a | 6 |
| mPool2_V2MM_155880 | 1.198419 | 3.004939 | TRUE | HP_37651 | TGCTGTTG | GGCTCTCT | XM_197869 | Mus musculus LOC272173 (LOC272173), mRNA. | | 2.6 | SM2129 | b | 5 |
| mPool2_V2MM_106552 | 1.19833 | 2.205692 | TRUE | HP_32765 | TGCTGTTG | GTCACAA/ | NM_199467 | RIKEN cDNA F730047E07 gene | F730047E07Rik | 2.7 | SM2155 | h | 6 |
| mPool4_V2MM_131911 | 1.198155 | 75.83547 | TRUE | HP_35267 | TGCTGTTG | CTGTGTG( | XM_488731 | hypothetical protein A530026G17 | | 2.16 | SM2637 | d | 9 |
| mPool4_V2MM_37052 | 1.198027 | 69.72441 | TRUE | HP_25982 | TGCTGTTG | GAATGCC1 | NM_144859 | praja 2, RING-H2 motif containing | Pja2 | 2.15 | SM2583 | b | 5 |
| 092308m1_V2MM_68315 | 1.19782 | 3.004939 | TRUE | HP_29027 | TGCTGTTG | CTTGGAA1 | NM_030261 | sestrin 3 | Sesn3 | 2.4 | SM2024 | g | 4 |
| mPool4_V2MM_132131 | 1.197516 | 75.83547 | TRUE | HP_35288 | TGCTGTTG | CCCTTTGG | AK014907 | RIKEN cDNA 4921517D16 gene | 4921517D16Rik | 2.14 | SM2535 | h | 11 |
| mPool4_V2MM_44096 | 1.197252 | 75.83547 | TRUE | HP_26666 | TGCTGTTG | CTGTTCAC | NM_147218 | ATP-binding cassette, sub-family A (ABC1), member 6 | Abca6 | 2.15 | SM2599 | a | 12 |
| 092308m1_V2MM_144898 | 1.196292 | 2.205692 | TRUE | HP_36560 | TGCTGTTG | CCAACCTT | XM_160409 | Mus musculus LOC242190 (LOC242190), mR. | | 2.5 | SM2051 | e | 11 |
| 092308m3_V2MM_232072 | 1.196219 | 60.13437 | TRUE | HP_48402 | TGCTGTTG | GCGCTTCT | NM_022999 | proline-rich Gla (G-carboxyglutamic acid) polypeptide 2 | Prrg2 | 2.14 | SM2504 | d | 1 |
| mPool5_V2MM_12733 | 1.196184 | 75.83547 | TRUE | HP_23611 | TGCTGTTG | CACCTTTG | NM_010893 | neuraminidase 1 | Neu1 | NA | | | NA |
| 092308m1_V2MM_184101 | 1.195897 | 3.004939 | TRUE | HP_40457 | TGCTGTTG | CAGGCTA1 | XM_289141 | Mus musculus LOC332728 (LOC332728), mR. | | 2.5 | SM2084 | c | 3 |
| 092308m1_V2MM_184101 | 1.195897 | 3.004939 | TRUE | HP_40457 | TGCTGTTG | CAGGCTA1 | XM_289141 | Mus musculus LOC332728 (LOC332728), mRNA. | | 2.5 | SM2084 | c | 3 |
| mPool6_V2MM_169845 | 1.195666 | 75.83547 | TRUE | HP_39034 | TGCTGTTG | GAAGGGA | XM_286346 | Mus musculus hypothetical gene supported by AK076588 (LOC327906), mRNA. | | 2.7 | SM2192 | a | 4 |

| | | | | | | | | | | | | | | |
|---|---|---|---|---|---|---|---|---|---|---|---|---|---|---|
| mPool7_V2MM_65931 | 1.195444 | 3.004939 | TRUE | HP_28796 | TGCTGTTG | GTCCCACC | NM_025629 | thrombospondin, type I domain containing 6 | Thsd6 | | 2.8 | SM2245 | c | 5 |
| mPool6_V2MM_81411 | 1.195153 | 75.83547 | TRUE | HP_30301 | TGCTGTTG | GTATTTGT | NM_013663 | splicing factor, arginine/serine-rich 3 (SRp20) | Sfrs3 | | 2.8 | SM2226 | g | 12 |
| mPool6_V2MM_61849 | 1.1947 | 75.83547 | TRUE | HP_28397 | TGCTGTTG | CAATTACA | NM_025705 | discoidin, CUB and LCCL domain containing 1 | Dcbld1 | | 2.7 | SM2176 | a | 5 |
| mPool7_V2MM_96426 | 1.194547 | 3.004939 | TRUE | HP_31767 | TGCTGTTG | CACCAGGT | L17069 | myeloid/lymphoid or mixed-lineage leukemia | Mll | | 2.9 | SM2260 | d | 1 |
| 092308m3_V2MM_87630 | 1.194491 | 75.83547 | TRUE | HP_30908 | TGCTGTTG | CTGCTTGC | NM_175208 | RIKEN cD 4931431C16 gene | 4931431C16Rik | | 2.13 | SM2466 | e | 12 |
| 092308m3_V2MM_94116 | 1.194032 | 75.83547 | TRUE | HP_31541 | TGCTGTTG | CCCATGGT | NM_183036 | defensin beta 38 | Defb38 | | 2.9 | SM2256 | a | 6 |
| 092308m3_V2MM_94116 | 1.194032 | 75.83547 | TRUE | HP_31541 | TGCTGTTG | CCCATGGT | NM_183036 | defensin beta 38 | Defb38 | | 2.9 | SM2256 | a | 6 |
| mPool2_V2MM_165085 | 1.193766 | 3.004939 | TRUE | HP_38562 | TGCTGTTG | CCCAAAGA | XM_285164 | Mus musculus similar to protease [Mus musculus] (LOC332160), mRNA. | | | 2.6 | SM2130 | f | 3 |
| mPool4_V2MM_122191 | 1.19353 | 59.99801 | TRUE | HP_34306 | TGCTGTTG | CTTCAGCC | XM_484474 | similar to High mobility group protein 1 (HMG-1) (Amphoterin) (Heparin-binding | | | 2.14 | SM2534 | h | 12 |
| mPool2_V2MM_202256 | 1.193343 | 2.205692 | TRUE | HP_42026 | TGCTGTTG | GAGCCTAT | AK033513 | RIKEN cDNA 6030408C04 gene | 6030408C04Rik | | 2.1 | SM2323 | b | 12 |
| mPool6_V2MM_2520 | 1.193163 | 66.95036 | TRUE | HP_22613 | TGCTGTTG | CTGAAGAC | NM_007841 | DEAD (Asp-Glu-Ala-Asp) box polypeptide 6 | Ddx6 | | 2.7 | SM2170 | f | 2 |
| mPool7_V2MM_102530 | 1.192928 | 3.004939 | TRUE | HP_32367 | TGCTGTTG | GCTCCAAT | AK036186 | Wilms tumour 1-associating protein | Wtap | | 2.1 | SM2310 | e | 2 |
| mPool4_V2MM_45263 | 1.192751 | 75.83547 | TRUE | HP_26780 | TGCTGTTG | GGTATTCT | NM_177102 | RIKEN cDNA A830041P22 gene | A830041P22Rik | | 2.16 | SM2620 | d | 3 |
| 092308m3_V2MM_209703 | 1.192534 | 75.83547 | TRUE | HP_42746 | TGCTGTTG | GACCAGGC | XM_156218 | similar to plakoglobin | | | 2.1 | SM2327 | f | 9 |
| 092308m3_V2MM_209703 | 1.192534 | 75.83547 | TRUE | HP_42746 | TGCTGTTG | GACCAGGC | XM_156218 | similar to plakoglobin | | | 2.1 | SM2327 | f | 9 |
| 092308m3_V2MM_206814 | 1.19196 | 66.95036 | TRUE | HP_42463 | TGCTGTTG | GAAGTCTA | XM_158055 | Mus musculus LOC227534 (LOC227534), mR. | | | 2.14 | SM2520 | a | 12 |
| 092308m3_V2MM_109571 | 1.191822 | 75.83547 | TRUE | HP_33062 | TGCTGTTG | GCAAGCTA | AY660739 | RIKEN cDNA 4833424K13 gene | 4833424K13Rik | | 2.1 | SM2318 | f | 11 |
| 092308m3_V2MM_109571 | 1.191822 | 75.83547 | TRUE | HP_33062 | TGCTGTTG | GCAAGCTA | AY660739 | RIKEN cD 4833424K13 gene | 4833424K13Rik | | 2.1 | SM2318 | f | 11 |
| mPool5_V2MM_7534 | 1.191779 | 75.83547 | TRUE | HP_23103 | TGCTGTTG | GCAATGTC | NM_011697 | vascular endothelial growth factor B | Vegfb | NA | | | NA | |
| 092308m3_V2MM_52863 | 1.191195 | 69.72441 | TRUE | HP_27523 | TGCTGTTG | GCTGCAGA | NM_177078 | adrenergic receptor kise, beta 2 | Adrbk2 | | 2.12 | SM2439 | f | 7 |
| 092308m1_V2MM_73288 | 1.190996 | 2.205692 | TRUE | HP_29512 | TGCTGTTG | GCCTTTGT | AK080731 | RIKEN cD A930024F17 gene | A930024F17Rik | | 2.3 | SM2012 | f | 11 |
| 092308m1_V2MM_66531 | 1.19087 | 3.004939 | TRUE | HP_28854 | TGCTGTTG | CAGCTTGT | NM_053070 | carbonic anhydrase 7 | Car7 | | 2.4 | SM2026 | a | 2 |
| 092308m1_V2MM_179498 | 1.190267 | 3.004939 | TRUE | HP_39997 | TGCTGTTG | GCCAGAGT | XM_288344 | Mus musculus LOC327806 (LOC327806), mR. | | | 2.5 | SM2087 | c | 7 |
| 092308m1_V2MM_155307 | 1.189461 | 3.004939 | TRUE | HP_37594 | TGCTGTTG | GAACTAAC | XM_197341 | Mus musculus LOC271503 (LOC271503), mR. | | | 2.6 | SM2123 | b | 12 |
| 092308m3_V2MM_61595 | 1.18944 | 75.83547 | TRUE | HP_28372 | TGCTGTTG | GTTTATTG | NM_010203 | fibroblast growth factor 5 | Fgf5 | | 2.12 | SM2411 | d | 10 |
| mPool4_V2MM_242072 | 1.18903 | 75.83547 | TRUE | HP_49382 | TGCTGTTG | CTGTGAAT | XM_488025 | similar to hypothetical protein FLJ38281 | | | 2.14 | SM2545 | f | 11 |
| mPool5_V2MM_5818 | 1.188487 | 75.83547 | TRUE | HP_22936 | TGCTGTTG | CGAGGAA | NM_175493 | G protein-coupled receptor 68 | Gpr68 | NA | | | NA | |
| mPool2_V2MM_9386 | 1.188204 | 3.004939 | TRUE | HP_23284 | TGCTGTTG | CAGCTACA | NM_145141 | expressed sequence BB219290 | BB219290 | | 2.11 | SM2389 | b | 3 |
| mPool6_V2MM_70998 | 1.18798 | 73.80895 | TRUE | HP_29289 | TGCTGTTG | CAGGTCAC | NM_013848 | erythroblast membrane-associated protein | Ermap | | 2.8 | SM2226 | h | 9 |
| 092308m1_V2MM_177421 | 1.187738 | 2.205692 | TRUE | HP_39790 | TGCTGTTG | CATTTACA | XM_287871 | Mus musculus hypothetical gene supported by AK086986 (LOC330881), mR. | | | 2.5 | SM2076 | e | 9 |
| 092308m1_V2MM_160259 | 1.187652 | 3.004939 | TRUE | HP_38084 | TGCTGTTG | GTTTCCAA | XM_283439 | expressed sequence AW049306 | AW049306 | | 2.6 | SM2121 | c | 8 |
| 092308m3_V2MM_44914 | 1.187644 | 59.99801 | TRUE | HP_26746 | TGCTGTTG | GCTGTGAC | NM_010181 | fibrillin 2 | Fbn2 | | 2.12 | SM2446 | a | 1 |
| mPool2_V2MM_53930 | 1.187433 | 3.004939 | TRUE | HP_16090 | TGCTGTTG | CACTTAAC | NM_010167 | eyes absent 4 homolog (Drosophila) | Eya4 | | 2.7 | SM2166 | d | 8 |
| mPool2_V2MM_5320 | 1.18668 | 3.004939 | TRUE | HP_22887 | TGCTGTTG | CCATTATT | NM_025957 | CEA-related cell adhesion molecule 14 | Ceacam14 | | 2.11 | SM2386 | a | 4 |
| 092308m3_V2MM_44284 | 1.186638 | 75.83547 | TRUE | HP_26684 | TGCTGTTG | CTTGTTGA | NM_172618 | BTB (POZ) domain containing 9 | Btbd9 | | 2.12 | SM2418 | e | 2 |
| 092308m1_V2MM_129465 | 1.186589 | 3.004939 | TRUE | HP_35027 | TGCTGTTG | GGTTGTAT | XM_146310 | Mus musculus LOC244446 (LOC244446), mR. | | | 2.5 | SM2099 | e | 4 |
| mPool2_V2MM_162396 | 1.186203 | 2.205692 | TRUE | HP_38295 | TGCTGTTG | CTTGCTTA | XM_284334 | Mus musculus similar to hypothetical protein FLJ13373 [Homo sapiens] (LOC330 | | | 2.6 | SM2131 | b | 11 |
| 092308m3_V2MM_85465 | 1.186023 | 75.83547 | TRUE | HP_30694 | TGCTGTTG | GCCGCCAT | NM_146389 | olfactory receptor 1350 | Olfr1350 | | 2.13 | SM2453 | d | 9 |
| mPool2_V2MM_90015 | 1.185477 | 3.004939 | TRUE | HP_31144 | TGCTGTTG | CTGTGAAC | NM_177715 | potassium channel tetramerisation domain containing 12 | Kctd12 | | 2.6 | SM2133 | d | 4 |
| mPool4_V2MM_115429 | 1.18546 | 75.83547 | TRUE | HP_33639 | TGCTGTTG | CTGCAAGT | XM_487246 | similar to Ig heavy chain precursor V region (VAR100) - mouse | | | 2.16 | SM2620 | g | 6 |
| mPool4_V2MM_111822 | 1.185376 | 75.83547 | TRUE | HP_33282 | TGCTGTTG | CTGGCAAC | XM_355234 | kinesin family member 14 | Kif14 | | 2.16 | SM2613 | d | 6 |
| 092308m1_V2MM_66945 | 1.185253 | 3.004939 | TRUE | HP_28894 | TGCTGTTG | CCCACAGT | NM_011059 | peptidyl arginine deimise, type I | Padi1 | | 2.3 | SM2010 | g | 11 |
| mPool2_V2MM_10496 | 1.184955 | 3.004939 | TRUE | HP_23392 | TGCTGTTG | CTCTACGT | NM_027275 | RIKEN cDNA 2810422B04 gene | 2810422B04Rik | | 2.11 | SM2382 | a | 6 |
| mPool7_V2MM_178353 | 1.184255 | 3.004939 | TRUE | HP_39882 | TGCTGTTG | CCAAGTAC | XM_288143 | Mus musculus hypothetical gene supported by AK046508 (LOC331558), mR. | | | 2.5 | SM2069 | g | 10 |
| mPool7_V2MM_178353 | 1.184255 | 3.004939 | TRUE | HP_39882 | TGCTGTTG | CCAAGTAC | XM_288143 | Mus musculus hypothetical gene supported by AK046508 (LOC331558), mRNA. | | | 2.5 | SM2069 | g | 10 |
| mPool7_V2MM_102051 | 1.184227 | 3.004939 | TRUE | HP_32321 | TGCTGTTG | CCTTAAAT | NM_026813 | RIKEN cDNA 8430411H09 gene | 8430411H09Rik | | 2.9 | SM2299 | d | 7 |
| 092308m3_V2MM_38616 | 1.184218 | 75.83547 | TRUE | HP_26133 | TGCTGTTG | GCCATAAA | NM_144828 | protein phosphatase 1, regulatory (inhibitor) subunit 1B | Ppp1r1b | | 2.11 | SM2399 | f | 1 |
| 092308m3_V2MM_227514 | 1.184077 | 75.83547 | TRUE | HP_47951 | TGCTGTTG | CTACAGCT | XM_124482 | ATP synthase, H+ transporting, mitochondrial F0 complex | Atp5l | | 2.13 | SM2456 | d | 12 |
| mPool4_V2MM_158175 | 1.183038 | 60.13437 | TRUE | HP_37878 | TGCTGTTG | CTTGGCAC | XM_206288 | Mus musculus LOC237944 (LOC237944), mRNA. | | NA | | | NA | |
| mPool2_V2MM_134397 | 1.182529 | 3.004939 | TRUE | HP_35512 | TGCTGTTG | CTATAACA | XM_149838 | Mus musculus LOC232644 (LOC232644), mRNA. | | | 2.11 | SM2363 | d | 2 |
| mPool2_V2MM_2496 | 1.182396 | 3.004939 | TRUE | HP_22611 | TGCTGTTG | CTCTTGTA | NM_018866 | chemokine (C-X-C motif) ligand 13 | Cxcl13 | | 2.11 | SM2386 | h | 12 |
| mPool4_V2MM_87936 | 1.182141 | 59.99801 | TRUE | HP_30938 | TGCTGTTG | GACTTGCT | NM_175276 | formin-family protein FHOS2 | | | 2.16 | SM2624 | e | 10 |
| 092308m3_V2MM_56691 | 1.181326 | 75.83547 | TRUE | HP_27896 | TGCTGTTG | CTGACTGT | BC055876 | RIKEN cD 1110007C02 gene | 1110007C02Rik | | 2.12 | SM2443 | c | 7 |

| | | | | | | | | | | | | | | |
|---|---|---|---|---|---|---|---|---|---|---|---|---|---|---|
| mPool2_V2MM_118660 | 1.181325 | 2.205692 | TRUE | HP_33959 | TGCTGTTG | GTTCAAAC | AK053237 | myosin head domain containing 1 | Myohd1 | | 2.11 | SM2368 | e | 5 |
| 092308m3_V2MM_139311 | 1.181177 | 60.13437 | TRUE | HP_36003 | TGCTGTTG | CGACTTTG | XM_154921 | Mus musculus LOC219168 (LOC219168), mR. | | | 2.14 | SM2516 | c | 8 |
| 092308m3_V2MM_105808 | 1.180869 | 75.83547 | TRUE | HP_32691 | TGCTGTTG | CTGATTGC | NM_008875 | phospholipase D1 | Pld1 | | 2.13 | SM2490 | b | 8 |
| mPool7_V2MM_185502 | 1.180513 | 3.004939 | TRUE | HP_40597 | TGCTGTTG | CCTTCTCA | XM_289381 | Mus musculus LOC333054 (LOC333054), mRNA. | | | 2.5 | SM2071 | d | 5 |
| mPool7_V2MM_185502 | 1.180513 | 3.004939 | TRUE | HP_40597 | TGCTGTTG | CCTTCTCA | XM_289381 | Mus musculus LOC333054 (LOC333054), mR. | | | 2.5 | SM2071 | d | 5 |
| 092308m1_V2MM_158758 | 1.18036 | 3.004939 | TRUE | HP_37936 | TGCTGTTG | CACAAGA( | XM_207576 | Mus musculus similar to envelope protein [Sus scrofa] (LOC280316), mR. | | | 2.6 | SM2117 | d | 5 |
| mPool2_V2MM_164934 | 1.180347 | 2.205692 | TRUE | HP_38547 | TGCTGTTG | CAAGTTGC | XM_285134 | Mus musculus similar to pol protein [Sus scrofa] (LOC328552), mRNA. | | | 2.6 | SM2127 | b | 2 |
| mPool6_V2MM_68559 | 1.179988 | 75.83547 | TRUE | HP_29051 | TGCTGTTG | GACAGAA | NM_025670 | RIKEN cDNA 5730403B10 gene | 5730403B10Rik | | 2.8 | SM2215 | d | 12 |
| mPool4_V2MM_189499 | 1.179796 | 75.83547 | TRUE | HP_32730 | TGCTGTTG | GGCTCAG( | NM_194343 | tripartite motif-containing 45 | Trim45 | | 2.16 | SM2623 | d | 2 |
| 092308m1_V2MM_131052 | 1.179749 | 2.205692 | TRUE | HP_35182 | TGCTGTTG | CTTCCTTA | AK006428 | RIKEN cD 1700027J07 gene | 1700027J07Rik | | 2.5 | SM2097 | h | 5 |
| 092308m1_V2MM_67801 | 1.179586 | 3.004939 | TRUE | HP_28977 | TGCTGTTG | CTCCAAAC | NM_009803 | nuclear receptor subfamily 1, group I, member 3 | Nr1i3 | | 2.3 | SM2001 | f | 4 |
| mPool4_V2MM_34025 | 1.179514 | 75.83547 | TRUE | HP_25685 | TGCTGTTG | CCAAGAG | NM_028228 | PIN2/TRF1-interacting protein | | | 2.15 | SM2597 | g | 5 |
| mPool6_V2MM_156589 | 1.177245 | 75.83547 | TRUE | HP_37722 | TGCTGTTG | CTGTGCCT | XM_203991 | similar to SHPS-1 | | | 2.7 | SM2194 | b | 4 |
| 092308m1_V2MM_133518 | 1.177072 | 3.004939 | TRUE | HP_35425 | TGCTGTTG | CCTTCAAT | XM_149291 | hypothetical gene supported by AK012899 | | | 2.5 | SM2097 | g | 10 |
| mPool2_V2MM_87228 | 1.176901 | 3.004939 | TRUE | HP_30868 | TGCTGTTG | CGAAGAA | NM_175111 | Hspb associated protein 1 | Hspbap1 | | 2.6 | SM2141 | c | 3 |
| mPool2_V2MM_114347 | 1.175589 | 3.004939 | TRUE | HP_33532 | TGCTGTTG | GCAGTATT | XM_137453 | Mus musculus similar to protease [Ovis aries] (LOC216550), mRNA. | | | 2.1 | SM2323 | e | 8 |
| mPool7_V2MM_200840 | 1.17557 | 2.205692 | TRUE | HP_41887 | TGCTGTTG | CTTTAGTC | AK042686 | hypothetical protein A730015I17 | | | 2.9 | SM2283 | h | 9 |
| mPool7_V2MM_178028 | 1.175099 | 2.205692 | TRUE | HP_39850 | TGCTGTTG | GGTACAG( | XM_288012 | similar to p47 protein isoform a | | | 2.9 | SM2284 | c | 5 |
| mPool7_V2MM_65519 | 1.174923 | 3.004939 | TRUE | HP_28755 | TGCTGTTG | GGACTAG | NM_025769 | RIKEN cDNA 5430404L10 gene | 5430404L10Rik | | 2.8 | SM2239 | b | 6 |
| mPool4_V2MM_20096 | 1.174707 | 75.83547 | TRUE | HP_24330 | TGCTGTTG | GTGCCAG | NM_026449 | UDP-N-acetyl-alpha-D-galactosamine:polypeptide N-acet | Galnt15 | | 2.15 | SM2597 | b | 3 |
| mPool2_V2MM_142227 | 1.174661 | 2.205692 | TRUE | HP_36294 | TGCTGTTG | GTCATTAC | XM_157427 | Mus musculus LOC240337 (LOC240337), mRNA. | | | 2.1 | SM2334 | h | 1 |
| 092308m1_V2MM_165913 | 1.173436 | 3.004939 | TRUE | HP_38644 | TGCTGTTG | CAGATTTA | XM_146592 | Mus musculus similar to protease [Mus musculus] (LOC210103), mR. | | | 2.6 | SM2118 | c | 9 |
| mPool7_V2MM_147294 | 1.173302 | 3.004939 | TRUE | HP_36800 | TGCTGTTG | CAGTTAAC | XM_162853 | Mus musculus LOC243690 (LOC243690), mRNA. | | | 2.1 | SM2329 | e | 3 |
| mPool4_V2MM_44518 | 1.171515 | 61.29828 | TRUE | HP_26707 | TGCTGTTG | GAGCTGA | NM_010088 | decidual/trophoblast prolactin-related protein | Dtprp | | 2.15 | SM2575 | c | 9 |
| 092308m3_V2MM_37343 | 1.17137 | 60.13437 | TRUE | HP_26010 | TGCTGTTG | GACGATG | NM_007928 | MAP/microtubule affinity-regulating kise 2 | Mark2 | | 2.11 | SM2400 | h | 12 |
| mPool6_V2MM_33614 | 1.17129 | 75.83547 | TRUE | HP_25646 | TGCTGTTG | CAGCTTGT | NM_007962 | epithelial V-like antigen 1 | Eva1 | | 2.7 | SM2170 | a | 7 |
| mPool7_V2MM_62388 | 1.17114 | 3.004939 | TRUE | HP_28449 | TGCTGTTG | CAGATTAT | NM_007392 | actin, alpha 2, smooth muscle, aorta | Acta2 | | 2.8 | SM2246 | b | 6 |
| mPool2_V2MM_57065 | 1.170621 | 3.004939 | TRUE | HP_27932 | TGCTGTTG | CTGTATCT | NM_010107 | ephrin A1 | Efna1 | | 2.7 | SM2165 | c | 8 |
| 092308m3_V2MM_41743 | 1.170269 | 61.28368 | TRUE | HP_26436 | TGCTGTTG | CAGAGGT | NM_020565 | sulfotransferase family 3A, member 1 | Sult3a1 | | 2.12 | SM2417 | h | 6 |
| mPool2_V2MM_120226 | 1.170055 | 3.004939 | TRUE | HP_34112 | TGCTGTTG | CACAGAA | XM_140812 | gene model 343, (NCBI) | Gm343 | | 2.11 | SM2368 | h | 2 |
| mPool2_V2MM_120226 | 1.170055 | 3.004939 | TRUE | HP_34112 | TGCTGTTG | CACAGAA | XM_140812 | gene model 343, (NCBI) | Gm343 | | 2.11 | SM2368 | h | 2 |
| mPool2_V2MM_114615 | 1.169766 | 3.004939 | TRUE | HP_33559 | TGCTGTTG | GTGAGCT( | XM_137762 | similar to PROHIBITIN (B-CELL RECEPTOR ASSOCIATED PROTEIN 32) (BAP 32) | | | 2.7 | SM2161 | b | 2 |
| mPool2_V2MM_84461 | 1.169437 | 3.004939 | TRUE | HP_30597 | TGCTGTTG | CCCTTAAG | NM_028173 | translocating chain-associating membrane protein 1 | Tram1 | | 2.6 | SM2141 | h | 6 |
| mPool2_V2MM_166076 | 1.169275 | 3.004939 | TRUE | HP_38660 | TGCTGTTG | CAGCTAA | XM_285404 | Mus musculus similar to ATP synthase, H+ transporting, mitochondrial F0 comple | | | 2.6 | SM2130 | h | 5 |
| mPool4_V2MM_241187 | 1.168214 | 60.13437 | TRUE | HP_49295 | TGCTGTTG | CTCTAAAT | XM_357569 | gene model 1396, (NCBI) | Gm1396 | | 2.14 | SM2543 | g | 2 |
| mPool7_V2MM_114177 | 1.167586 | 2.205692 | TRUE | HP_33516 | TGCTGTTG | CTATCGAC | XM_137307 | Mus musculus similar to 60S RIBOSOMAL PROTEIN L29 (P23) (LOC216263), mRN | | | 2.7 | SM2158 | h | 1 |
| mPool6_V2MM_72850 | 1.167445 | 75.83547 | TRUE | HP_29469 | TGCTGTTG | GAAGCCT( | NM_010405 | hemoglobin X, alpha-like embryonic chain in Hba comple | Hba-x | | 2.7 | SM2180 | e | 5 |
| 092308m3_V2MM_117273 | 1.167386 | 75.83547 | TRUE | HP_33821 | TGCTGTTG | GATGTGT( | XM_139295 | caspase recruitment domain family, member 6 | Card6 | | 2.14 | SM2507 | e | 7 |
| 092308m1_V2MM_187780 | 1.167259 | 2.205692 | TRUE | HP_40824 | TGCTGTTG | CGAAATAT | XM_290140 | Mus musculus LOC333674 (LOC333674), mR. | | | 2.5 | SM2076 | h | 11 |
| mPool7_V2MM_70593 | 1.167152 | 3.004939 | TRUE | HP_29250 | TGCTGTTG | CACCTTAA | NM_025734 | potassium voltage-gated channel, subfamily G, member | Kcng4 | | 2.8 | SM2229 | g | 8 |
| mPool5_V2MM_178000 | 1.166189 | 75.83547 | TRUE | HP_39847 | TGCTGTTG | GCCGGTG | XM_287998 | RIKEN cDNA 1700008B15 gene | 1700008B15Rik | NA | | | NA | |
| mPool6_V2MM_224067 | 1.166102 | 70.87591 | TRUE | HP_47615 | TGCTGTTG | CAAAGGA | XM_484949 | hypothetical gene supported by AK040286 | | | 2.8 | SM2212 | c | 8 |
| mPool5_V2MM_7716 | 1.164838 | 66.95036 | TRUE | HP_23121 | TGCTGTTG | GCCATTGT | XM_136256 | Mus musculus similar to protease (LOC240779), mRNA. | | NA | | | NA | |
| mPool4_V2MM_118347 | 1.16421 | 61.29828 | TRUE | HP_33928 | TGCTGTTG | CACATGG | XM_139800 | similar to actin related protein 2/3 complex subunit 2; ARP2/3 protein complex s | | | 2.16 | SM2629 | a | 11 |
| mPool2_V2MM_146021 | 1.164176 | 3.004939 | TRUE | HP_36673 | TGCTGTTG | CTCTGAG | XM_161699 | Mus musculus LOC242896 (LOC242896), mRNA. | | | 2.1 | SM2336 | c | 8 |
| 092308m3_V2MM_96297 | 1.163738 | 75.83547 | TRUE | HP_25474 | TGCTGTTG | GACTGAT( | XM_284619 | similar to Cdv3 protein | | | 2.12 | SM2436 | c | 2 |
| mPool5_V2MM_164644 | 1.163568 | 75.83547 | TRUE | HP_38518 | TGCTGTTG | GACATATT | XM_285057 | Mus musculus similar to ubiquitin specific protease 10 [Homo sapiens] (LOC3283 | | NA | | | NA | |
| mPool2_V2MM_102905 | 1.163516 | 3.004939 | TRUE | HP_32405 | TGCTGTTG | GACATAAT | XM_128692 | dishevelled associated activator of morphogenesis 2 | Daam2 | | 2.7 | SM2157 | f | 6 |
| mPool7_V2MM_173934 | 1.163251 | 3.004939 | TRUE | HP_39442 | TGCTGTTG | CTATATCA | XM_287120 | Mus musculus hypothetical gene supported by AK034841 (LOC329386), mRNA. | | | 2.5 | SM2069 | e | 7 |
| mPool7_V2MM_173934 | 1.163251 | 3.004939 | TRUE | HP_39442 | TGCTGTTG | CTATATCA | XM_287120 | Mus musculus hypothetical gene supported by AK034841 (LOC329386), mR. | | | 2.5 | SM2069 | e | 7 |
| mPool2_V2MM_148209 | 1.163156 | 2.205692 | TRUE | HP_36892 | TGCTGTTG | GAAGTAG | XM_163597 | Mus musculus LOC210179 (LOC210179), mR. | | | 2.4 | SM2030 | e | 5 |
| mPool2_V2MM_148209 | 1.163156 | 2.205692 | TRUE | HP_36892 | TGCTGTTG | GAAGTAG | XM_163597 | Mus musculus LOC210179 (LOC210179), mRNA. | | | 2.4 | SM2030 | e | 5 |
| mPool2_V2MM_20 | 1.162433 | 3.004939 | TRUE | HP_22370 | TGCTGTTG | GTCTCCTA | D29668 | immunoglobulin kappa chain variable 8 (V8) | Igk-V8 | | 2.11 | SM2384 | h | 4 |

| | | | | | | | | | | | | | | | |
|---|---|---|---|---|---|---|---|---|---|---|---|---|---|---|---|
| 092308m3_V2MM_58849 | 1.161888 | 59.99801 | TRUE | HP_28106 | TGCTGTTG | GCAACAT | NM_147014 | olfactory receptor 1048 | Olfr1048 | | 2.12 | SM2411 | b | | 5 |
| mPool4_V2MM_11016 | 1.161439 | 75.83547 | TRUE | HP_23443 | TGCTGTTG | CTGTGTG | NM_175284 | frizzled homolog 10 (Drosophila) | Fzd10 | | 2.16 | SM2618 | f | | 12 |
| mPool6_V2MM_158511 | 1.161054 | 75.83547 | TRUE | HP_37912 | TGCTGTTG | GCTTTCAA | XM_206707 | Mus musculus LOC279102 (LOC279102), mRNA. | | | 2.7 | SM2189 | a | | 4 |
| mPool2_V2MM_132331 | 1.160861 | 3.004939 | TRUE | HP_35308 | TGCTGTTG | CCAGTTCT | XM_358823 | leucine rich repeat containing 15 | Lrrc15 | | 2.5 | SM2099 | f | | 9 |
| mPool2_V2MM_132331 | 1.160861 | 3.004939 | TRUE | HP_35308 | TGCTGTTG | CCAGTTCT | XM_358823 | leucine rich repeat containing 15 | Lrrc15 | | 2.5 | SM2099 | f | | 9 |
| 092308m3_V2MM_36537 | 1.160679 | 61.29828 | TRUE | HP_25931 | TGCTGTTG | CTGTGATT | NM_011985 | matrix metalloproteise 23 | Mmp23 | | 2.11 | SM2396 | c | | 5 |
| mPool7_V2MM_74105 | 1.160661 | 3.004939 | TRUE | HP_29591 | TGCTGTTG | CAAGAAT | NM_009888 | complement component factor h | Cfh | | 2.8 | SM2232 | a | | 11 |
| mPool2_V2MM_141085 | 1.160474 | 2.205692 | TRUE | HP_36179 | TGCTGTTG | CTATTGA | XM_156459 | Mus musculus LOC239959 (LOC239959), mR. | | | 2.4 | SM2036 | g | | 12 |
| mPool2_V2MM_141085 | 1.160474 | 2.205692 | TRUE | HP_36179 | TGCTGTTG | CTATTGA | XM_156459 | Mus musculus LOC239959 (LOC239959), mRNA. | | | 2.4 | SM2036 | g | | 12 |
| 092308m1_V2MM_145508 | 1.160388 | 3.004939 | TRUE | HP_36621 | TGCTGTTG | CTTGGTCT | XM_161006 | Mus musculus LOC242463 (LOC242463), mR. | | | 2.4 | SM2043 | c | | 12 |
| 092308m1_V2MM_145508 | 1.160388 | 3.004939 | TRUE | HP_36621 | TGCTGTTG | CTTGGTCT | XM_161006 | Mus musculus LOC242463 (LOC242463), mRNA. | | | 2.4 | SM2043 | c | | 12 |
| 092308m3_V2MM_151658 | 1.160368 | 35.99881 | TRUE | HP_37232 | TGCTGTTG | CTCCTGTA | XM_194896 | Mus musculus similar to MRDS1 protein [Homo sapiens] (LOC268648), mR. | | | 2.14 | SM2513 | g | | 7 |
| mPool2_V2MM_11352 | 1.160003 | 3.004939 | TRUE | HP_23476 | TGCTGTTG | GACTGGA | NM_008217 | hyaluronan synthase 3 | Has3 | | 2.7 | SM2165 | f | | 7 |
| mPool2_V2MM_97978 | 1.159849 | 3.004939 | TRUE | HP_31921 | TGCTGTTG | CCACTGT | XM_112001 | Mus musculus LOC194863 (LOC194863), mRNA. | | | 2.6 | SM2133 | b | | 10 |
| 092308m1_V2MM_95567 | 1.159462 | 3.004939 | TRUE | HP_19009 | TGCTGTTG | GGGATAA | NM_198033 | expressed sequence AW060766 | AW060766 | | 2.6 | SM2107 | h | | 3 |
| 092308m3_V2MM_109745 | 1.159295 | 75.83547 | TRUE | HP_33079 | TGCTGTTG | GAAACAA | AK019126 | LSM6 homolog, U6 small nuclear R associated (S. cerevisi | Lsm6 | | 2.1 | SM2311 | e | | 9 |
| 092308m3_V2MM_109745 | 1.159295 | 75.83547 | TRUE | HP_33079 | TGCTGTTG | GAAACAA | AK019126 | LSM6 homolog, U6 small nuclear RNA associated (S. cere | Lsm6 | | 2.1 | SM2311 | e | | 9 |
| mPool2_V2MM_82342 | 1.158811 | 3.004939 | TRUE | HP_30392 | TGCTGTTG | GTCACCAT | NM_008441 | kinesin family member 1B | Kif1b | | 2.6 | SM2144 | h | | 4 |
| 092308m3_V2MM_106033 | 1.15869 | 66.95036 | TRUE | HP_32713 | TGCTGTTG | GTGATAC | XM_130987 | retinoic acid receptor responder (tazarotene induced) 1 | Rarres1 | | 2.1 | SM2315 | a | | 5 |
| 092308m3_V2MM_106033 | 1.15869 | 66.95036 | TRUE | HP_32713 | TGCTGTTG | GTGATAC | XM_130987 | retinoic acid receptor responder (tazarotene induced) 1 | Rarres1 | | 2.1 | SM2315 | a | | 5 |
| mPool7_V2MM_146127 | 1.158583 | 3.004939 | TRUE | HP_36683 | TGCTGTTG | CAGCTTAT | XM_161773 | hypothetical gene supported by AK036869 | | | 2.4 | SM2050 | a | | 1 |
| mPool7_V2MM_146127 | 1.158583 | 3.004939 | TRUE | HP_36683 | TGCTGTTG | CAGCTTAT | XM_161773 | hypothetical gene supported by AK036869 | | | 2.4 | SM2050 | a | | 1 |
| 092308m3_V2MM_24874 | 1.158542 | 75.83547 | TRUE | HP_24796 | TGCTGTTG | GTCCCAG | NM_178775 | ribosomal protein S6 kise, 52kDa, polypeptide 1 | Rps6kc1 | | 2.12 | SM2426 | g | | 9 |
| 092308m1_V2MM_94697 | 1.158208 | 3.004939 | TRUE | HP_31598 | TGCTGTTG | CTACTAGT | NM_183214 | RIKEN cD A830019P07 gene | A830019P07Rik | | 2.6 | SM2109 | f | | 7 |
| mPool7_V2MM_108418 | 1.158064 | 3.004939 | TRUE | HP_32948 | TGCTGTTG | GGAATCA | BC035209 | PTPRF interacting protein, binding protein 1 (liprin beta 1 | Ppfibp1 | | 2.7 | SM2153 | c | | 8 |
| mPool5_V2MM_83212 | 1.157719 | 75.83547 | TRUE | HP_30476 | TGCTGTTG | GCTTGAG | NM_011591 | translocator of inner mitochondrial membrane 17b | Timm17b | NA | | | | NA | |
| mPool7_V2MM_110094 | 1.157703 | 2.205692 | TRUE | HP_33113 | TGCTGTTG | CTCAAGT | BC059844 | RIKEN cDNA 4930566A11 gene | 4930566A11Rik | | 2.1 | SM2303 | d | | 6 |
| mPool6_V2MM_168312 | 1.156912 | 75.83547 | TRUE | HP_38881 | TGCTGTTG | GGCATCC | XM_285956 | Mus musculus similar to glyceraldehyde-3-phosphate dehydrogenase [Rattus nor | | | 2.8 | SM2211 | f | | 5 |
| mPool2_V2MM_136965 | 1.155604 | 3.004939 | TRUE | HP_35768 | TGCTGTTG | GTTAGTA | XM_488581 | LOC432511 | | | 2.1 | SM2334 | d | | 12 |
| 092308m3_V2MM_206971 | 1.155346 | 75.83547 | TRUE | HP_42478 | TGCTGTTG | GTAGCAG | XM_162125 | Mus musculus LOC243151 (LOC243151), mR. | | | 2.1 | SM2325 | e | | 1 |
| 092308m3_V2MM_206971 | 1.155346 | 75.83547 | TRUE | HP_42478 | TGCTGTTG | GTAGCAG | XM_162125 | Mus musculus LOC243151 (LOC243151), mRNA. | | | 2.1 | SM2325 | e | | 1 |
| 092308m1_V2MM_144190 | 1.153888 | 2.205692 | TRUE | HP_36490 | TGCTGTTG | CCATGGA | XM_159798 | Mus musculus LOC207822 (LOC207822), mR. | | | 2.4 | SM2045 | h | | 7 |
| 092308m3_V2MM_33201 | 1.152973 | 75.83547 | TRUE | HP_25605 | TGCTGTTG | CACCTTGT | NM_133774 | StAR-related lipid transfer (START) domain containing 4 | Stard4 | | 2.12 | SM2435 | f | | 1 |
| mPool2_V2MM_124311 | 1.152429 | 2.205692 | TRUE | HP_34516 | TGCTGTTG | CCTTTACC | XM_143381 | Mus musculus similar to 60S acidic ribosomal protein P1 (LOC229548), mRNA. | | | 2.5 | SM2097 | h | | 1 |
| mPool2_V2MM_124311 | 1.152429 | 2.205692 | TRUE | HP_34516 | TGCTGTTG | CCTTTACC | XM_143381 | Mus musculus similar to 60S acidic ribosomal protein P1 (LOC229548), mR. | | | 2.5 | SM2097 | h | | 1 |
| mPool6_V2MM_79209 | 1.152404 | 75.83547 | TRUE | HP_30087 | TGCTGTTG | GATGATA | NM_026197 | RIKEN cDNA 2810013M15 gene | 2810013M15Rik | | 2.8 | SM2223 | g | | 2 |
| 092308m3_V2MM_22487 | 1.152294 | 62.64426 | TRUE | HP_24563 | TGCTGTTG | CGTGGTA | NM_178446 | cD sequence BC013481 | BC013481 | | 2.11 | SM2395 | a | | 2 |
| 092308m3_V2MM_55747 | 1.152215 | 75.83547 | TRUE | HP_27804 | TGCTGTTG | CGAACTA | NM_029292 | RIKEN cD 1700008F21 gene | 1700008F21Rik | | 2.12 | SM2441 | d | | 10 |
| 092308m1_V2MM_175542 | 1.151973 | 3.004939 | TRUE | HP_39602 | TGCTGTTG | GCTTCTGA | XM_287472 | gene model 834, (NCBI) | Gm834 | | 2.5 | SM2093 | f | | 12 |
| 092308m3_V2MM_126795 | 1.151875 | 60.13437 | TRUE | HP_27669 | TGCTGTTG | CAGTGCC | XM_144814 | gene model 459, (NCBI) | Gm459 | | 2.12 | SM2437 | a | | 3 |
| mPool4_V2MM_126431 | 1.151527 | 70.87591 | TRUE | HP_34727 | TGCTGTTG | GTTGTTCA | XM_144553 | Mus musculus similar to 25 oligoadenylate synthetase-2 [Rattus norvegicus] (LOC | | | 2.16 | SM2638 | a | | 7 |
| mPool7_V2MM_105520 | 1.151276 | 3.004939 | TRUE | HP_32663 | TGCTGTTG | GGCAGTT | XM_130565 | RIKEN cDNA 4732486I23 gene | 4732486I23Rik | | 2.1 | SM2313 | e | | 4 |
| mPool2_V2MM_96051 | 1.150916 | 3.004939 | TRUE | HP_31731 | TGCTGTTG | GAATTGG | XM_110002 | similar to oogenesin | | | 2.6 | SM2139 | a | | 2 |
| mPool2_V2MM_109275 | 1.150877 | 3.004939 | TRUE | HP_33033 | TGCTGTTG | CTCTCAGA | XM_133717 | RIKEN cDNA 4933427G17 gene | 4933427G17Rik | | 2.1 | SM2324 | a | | 10 |
| mPool2_V2MM_141475 | 1.150611 | 2.205692 | TRUE | HP_36218 | TGCTGTTG | CCCTCTCT | XM_156701 | Mus musculus LOC213062 (LOC213062), mRNA. | | | 2.1 | SM2342 | h | | 7 |
| mPool7_V2MM_148663 | 1.149645 | 3.004939 | TRUE | HP_36937 | TGCTGTTG | CTTTGATA | XM_164008 | Mus musculus LOC210862 (LOC210862), mR. | | | 2.1 | SM2328 | e | | 1 |
| mPool7_V2MM_148663 | 1.149645 | 3.004939 | TRUE | HP_36937 | TGCTGTTG | CTTTGATA | XM_164008 | Mus musculus LOC210862 (LOC210862), mRNA. | | | 2.1 | SM2328 | e | | 1 |
| mPool5_V2MM_4059 | 1.149432 | 75.83547 | TRUE | HP_22763 | TGCTGTTG | GGCTACC | NM_026582 | RIKEN cDNA 5031439A09 gene | 5031439A09Rik | NA | | | | NA | |
| mPool6_V2MM_70121 | 1.149132 | 75.83547 | TRUE | HP_29203 | TGCTGTTG | GTGTGTG | NM_028726 | RIKEN cDNA 4632404H12 gene | 4632404H12Rik | | 2.7 | SM2173 | d | | 7 |
| mPool7_V2MM_198418 | 1.14851 | 2.205692 | TRUE | HP_41647 | TGCTGTTG | GAGTTCTT | XM_287701 | Mus musculus hypothetical gene supported by AK081791 (LOC330572), mRNA. | | | 2.9 | SM2271 | c | | 12 |
| mPool7_V2MM_143579 | 1.148032 | 3.004939 | TRUE | HP_36429 | TGCTGTTG | CAGAGGA | XM_158861 | Mus musculus LOC241609 (LOC241609), mRNA. | | | 2.1 | SM2329 | c | | 2 |
| mPool2_V2MM_103648 | 1.147853 | 3.004939 | TRUE | HP_32478 | TGCTGTTG | CTCAGAA | XM_129179 | Mus musculus similar to Elongation factor 1-alpha 1 (EF-1-alpha-1) (Elongation fa | | | 2.7 | SM2151 | e | | 10 |
| mPool7_V2MM_115572 | 1.147766 | 3.004939 | TRUE | HP_33653 | TGCTGTTG | CCTGTTGC | XM_138503 | Mus musculus similar to spermine synthase; spermidine aminopropyltransferase | | | 2.7 | SM2157 | h | | 1 |

| | | | | | | | | | | | | | | |
|---|---|---|---|---|---|---|---|---|---|---|---|---|---|---|
| 092308m1_V2MM_158501 | 1.147538 | 2.205692 | TRUE | HP_37911 | TGCTGTTG | CTCACCAC | XM_206687 | Mus musculus LOC279081 (LOC279081), mR. | | 2.6 | SM2115 | d | | 5 |
| mPool7_V2MM_203726 | 1.147261 | 1.959154 | TRUE | HP_40923 | TGCTGTTG | CAAGAAAT | NM_199055 | carbohydrate (N-acetylgalactosamine 4-0) sulfotransfera | Chst9 | 2.1 | SM2301 | d | | 2 |
| mPool2_V2MM_96152 | 1.147171 | 3.004939 | TRUE | HP_31741 | TGCTGTTG | CTCGAATC | NM_010479 | heat shock protein 1A | Hspa1a | 2.6 | SM2140 | g | | 4 |
| mPool4_V2MM_108910 | 1.146959 | 75.83547 | TRUE | HP_32997 | TGCTGTTG | GCCTAGA | AK028022 | RIKEN cDNA 1190020J12 gene | 1190020J12Rik | 2.16 | SM2605 | c | | 12 |
| mPool4_V2MM_229820 | 1.146797 | 75.83547 | TRUE | HP_48180 | TGCTGTTG | GGCTCTTG | XM_488696 | hypothetical protein B930071L05 | | NA | | | NA | |
| 092308m3_V2MM_233858 | 1.146316 | 75.83547 | TRUE | HP_48575 | TGCTGTTG | CATTTGAC | XM_137252 | Mus musculus similar to nuclear transport factor 2 [Rattus norvegicus] (LOC2375 | | 2.14 | SM2503 | e | | 9 |
| mPool4_V2MM_33162 | 1.146267 | 75.83547 | TRUE | HP_25602 | TGCTGTTG | CACCCACA | NM_080451 | synaptopodin 2 | Synpo2 | 2.16 | SM2603 | h | | 1 |
| mPool7_V2MM_94142 | 1.146061 | 3.004939 | TRUE | HP_31543 | TGCTGTTG | GTGCGTTC | NM_183089 | RIKEN cDNA 2600005O03 gene | 2600005O03Rik | 2.9 | SM2256 | a | | 11 |
| mPool2_V2MM_97302 | 1.145999 | 3.004939 | TRUE | HP_31854 | TGCTGTTG | CAATCAGA | XM_111423 | Mus musculus similar to protease [Mus musculus] (LOC195221), mRNA. | | 2.6 | SM2143 | c | | 9 |
| 092308m1_V2MM_118470 | 1.145543 | 3.004939 | TRUE | HP_33940 | TGCTGTTG | CGAATGA | XM_139866 | similar to cytochrome P450 CYP4F16 | | 2.5 | SM2097 | b | | 11 |
| 092308m3_V2MM_231561 | 1.145432 | 75.83547 | TRUE | HP_48352 | TGCTGTTG | CAGAGGC | AK087368 | myeloid/lymphoid or mixed-lineage leukemia 3 | Mll3 | 2.14 | SM2502 | e | | 1 |
| 092308m1_V2MM_152581 | 1.145242 | 3.004939 | TRUE | HP_37324 | TGCTGTTG | GTGGACAT | XM_195448 | Mus musculus similar to DJ (Hsp40) homolog, subfamily B, member 6 isoform a; I | | 2.4 | SM2035 | g | | 3 |
| 092308m1_V2MM_152581 | 1.145242 | 3.004939 | TRUE | HP_37324 | TGCTGTTG | GTGGACAT | XM_195448 | Mus musculus similar to DnaJ (Hsp40) homolog, subfamily B, member 6 isoform a | | 2.4 | SM2035 | g | | 3 |
| mPool7_V2MM_171834 | 1.144883 | 3.004939 | TRUE | HP_39232 | TGCTGTTG | GGAAGAA | XM_286706 | hypothetical gene supported by AK040104 | | 2.5 | SM2090 | e | | 2 |
| mPool7_V2MM_171834 | 1.144883 | 3.004939 | TRUE | HP_39232 | TGCTGTTG | GGAAGAA | XM_286706 | hypothetical gene supported by AK040104 | | 2.5 | SM2090 | e | | 2 |
| 092308m1_V2MM_173927 | 1.144809 | 2.205692 | TRUE | HP_39441 | TGCTGTTG | CAGAAATC | AK033904 | hypothetical protein 9330112M16 | | 2.5 | SM2088 | a | | 1 |
| 092308m1_V2MM_173927 | 1.144809 | 2.205692 | TRUE | HP_39441 | TGCTGTTG | CAGAAATC | AK033904 | hypothetical protein 9330112M16 | | 2.5 | SM2088 | a | | 1 |
| mPool4_V2MM_124074 | 1.14462 | 75.83547 | TRUE | HP_34492 | TGCTGTTG | CAGATTTC | XM_143290 | Mus musculus similar to endothelial-monocyte activating polypeptide II (LOC229 | | 2.14 | SM2543 | b | | 9 |
| 092308m1_V2MM_67649 | 1.144481 | 3.004939 | TRUE | HP_28963 | TGCTGTTG | CTAAGTTA | AK051378 | mitogen-activated protein kise kise kise 13 LocusID: 9175 | Map3k13 | 2.4 | SM2024 | g | | 3 |
| 092308m3_V2MM_233523 | 1.14435 | 75.83547 | TRUE | HP_48543 | TGCTGTTG | CCGTCAAT | AK020441 | RIKEN cD 9430027B09 gene | 9430027B09Rik | 2.13 | SM2487 | c | | 10 |
| 092308m3_V2MM_151297 | 1.143842 | 75.83547 | TRUE | HP_37196 | TGCTGTTG | GTCTCTAT | XM_194682 | similar to L-lactate dehydrogese A chain (LDH-A) (LDH muscle subunit) (LDH-M) | | 2.14 | SM2523 | g | | 1 |
| mPool4_V2MM_181343 | 1.14296 | 62.64426 | TRUE | HP_40181 | TGCTGTTG | GTCTCCGA | XM_288660 | Mus musculus LOC332070 (LOC332070), mRNA. | | NA | | | NA | |
| mPool7_V2MM_173900 | 1.142758 | 3.004939 | TRUE | HP_39438 | TGCTGTTG | GTAGGAA | XM_287114 | Mus musculus hypothetical gene supported by AK052519 (LOC329402), mR. | | 2.5 | SM2079 | h | | 2 |
| mPool7_V2MM_173900 | 1.142758 | 3.004939 | TRUE | HP_39438 | TGCTGTTG | GTAGGAA | XM_287114 | Mus musculus hypothetical gene supported by AK052519 (LOC329402), mRNA. | | 2.5 | SM2079 | h | | 2 |
| 092308m3_V2MM_115106 | 1.142545 | 59.99801 | TRUE | HP_33607 | TGCTGTTG | CTATTGCT | XM_138114 | Mus musculus similar to glyceraldehyde-3-phosphate dehydrogese (LOC217616), | | 2.1 | SM2307 | b | | 9 |
| 092308m3_V2MM_115106 | 1.142545 | 59.99801 | TRUE | HP_33607 | TGCTGTTG | CTATTGCT | XM_138114 | Mus musculus similar to glyceraldehyde-3-phosphate dehydrogenase (LOC21761 | | 2.1 | SM2307 | b | | 9 |
| 092308m1_V2MM_120199 | 1.142293 | 2.205692 | TRUE | HP_34110 | TGCTGTTG | CGGTATA | XM_140800 | dehydrogenase E1 and transketolase domain containing | Dhtkd1 | 2.11 | SM2368 | b | | 3 |
| 092308m1_V2MM_120199 | 1.142293 | 2.205692 | TRUE | HP_34110 | TGCTGTTG | CGGTATA | XM_140800 | dehydrogese E1 and transketolase domain containing 1 | Dhtkd1 | 2.11 | SM2368 | b | | 3 |
| 092308m1_V2MM_174038 | 1.142277 | 3.004939 | TRUE | HP_39452 | TGCTGTTG | CATAGTCT | XM_287146 | Mus musculus hypothetical gene supported by AK080575 (LOC329468), mR. | | 2.5 | SM2069 | e | | 8 |
| mPool5_V2MM_12865 | 1.141786 | 35.99881 | TRUE | HP_23624 | TGCTGTTG | CAGAAGC | NM_007557 | bone morphogenetic protein 7 | Bmp7 | NA | | | NA | |
| mPool7_V2MM_177805 | 1.141473 | 2.205692 | TRUE | HP_39828 | TGCTGTTG | CAGATAAT | AK076537 | RIKEN cDNA 4833445I07 gene | 4833445I07Rik | 2.5 | SM2067 | h | | 6 |
| mPool7_V2MM_177805 | 1.141473 | 2.205692 | TRUE | HP_39828 | TGCTGTTG | CAGATAAT | AK076537 | RIKEN cD 4833445I07 gene | 4833445I07Rik | 2.5 | SM2067 | h | | 6 |
| 092308m3_V2MM_52677 | 1.141256 | 75.83547 | TRUE | HP_27505 | TGCTGTTG | GCAGCAG | XM_126673 | SMAD specific E3 ubiquitin protein ligase 2 | Smurf2 | 2.12 | SM2440 | e | | 4 |
| mPool2_V2MM_141569 | 1.141107 | 3.004939 | TRUE | HP_36228 | TGCTGTTG | CCTAGGA | XM_156781 | Mus musculus LOC210475 (LOC210475), mRNA. | | 2.1 | SM2335 | d | | 6 |
| mPool7_V2MM_171571 | 1.140959 | 3.004939 | TRUE | HP_39206 | TGCTGTTG | GACTCAA | AK030258 | RIKEN cDNA 4930556M19 gene | 4930556M19Rik | 2.9 | SM2295 | d | | 2 |
| 092308m3_V2MM_106878 | 1.138996 | 75.83547 | TRUE | HP_32797 | TGCTGTTG | GAGACCTA | XM_131619 | cD sequence BC022150 | BC022150 | 2.14 | SM2506 | h | | 5 |
| mPool7_V2MM_196450 | 1.138675 | 3.004939 | TRUE | HP_41455 | TGCTGTTG | CGTTGTTT | NM_172015 | isoleucine-tRNA synthetase | Iars | 2.9 | SM2254 | d | | 11 |
| mPool7_V2MM_196450 | 1.138675 | 3.004939 | TRUE | HP_41455 | TGCTGTTG | CGTTGTTT | NM_172015 | isoleucine-tR synthetase | Iars | 2.9 | SM2254 | d | | 11 |
| mPool4_V2MM_136264 | 1.13815 | 61.29828 | TRUE | HP_35698 | TGCTGTTG | CAAATATG | XM_151591 | Mus musculus LOC213852 (LOC213852), mRNA. | | 2.14 | SM2529 | e | | 5 |
| mPool7_V2MM_187047 | 1.138097 | 3.004939 | TRUE | HP_40751 | TGCTGTTG | CAAAGAA | XM_289640 | Mus musculus LOC333402 (LOC333402), mRNA. | | 2.5 | SM2072 | f | | 3 |
| mPool7_V2MM_187047 | 1.138097 | 3.004939 | TRUE | HP_40751 | TGCTGTTG | CAAAGAA | XM_289640 | Mus musculus LOC333402 (LOC333402), mR. | | 2.5 | SM2072 | f | | 3 |
| mPool4_V2MM_198581 | 1.137742 | 75.83547 | TRUE | HP_41663 | TGCTGTTG | GGACTGCA | XM_287402 | gene model 833, (NCBI) | Gm833 | NA | | | NA | |
| 092308m3_V2MM_204989 | 1.137483 | 75.83547 | TRUE | HP_42287 | TGCTGTTG | CCAATGGA | XM_486261 | glutaminyl-tR synthetase | Qars | 2.13 | SM2499 | a | | 9 |
| mPool2_V2MM_126885 | 1.137264 | 3.004939 | TRUE | HP_34772 | TGCTGTTG | CTGAGGCT | XM_144862 | Mus musculus similar to Gag [Ovis aries] (LOC232180), mRNA. | | 2.11 | SM2358 | a | | 8 |
| mPool2_V2MM_120199 | 1.137135 | 2.205692 | TRUE | HP_34110 | TGCTGTTG | CGGTATA | XM_140800 | dehydrogenase E1 and transketolase domain containing | Dhtkd1 | 2.11 | SM2368 | b | | 3 |
| mPool2_V2MM_120199 | 1.137135 | 2.205692 | TRUE | HP_34110 | TGCTGTTG | CGGTATA | XM_140800 | dehydrogese E1 and transketolase domain containing 1 | Dhtkd1 | 2.11 | SM2368 | b | | 3 |
| 092308m3_V2MM_112859 | 1.136834 | 66.95036 | TRUE | HP_33385 | TGCTGTTG | CAATGGAT | XM_136697 | cDNA sequence BC056494 | BC056494 | 2.1 | SM2321 | g | | 2 |
| 092308m3_V2MM_112859 | 1.136834 | 66.95036 | TRUE | HP_33385 | TGCTGTTG | CAATGGAT | XM_136697 | cD sequence BC056494 | BC056494 | 2.1 | SM2321 | g | | 2 |
| mPool7_V2MM_112723 | 1.136815 | 3.004939 | TRUE | HP_33371 | TGCTGTTG | CCGAGTA | XM_136631 | Mus musculus similar to ribosomal protein L31 [Rattus norvegicus] (LOC227150), | | 2.1 | SM2305 | d | | 10 |
| mPool7_V2MM_62723 | 1.13665 | 3.004939 | TRUE | HP_28482 | TGCTGTTG | CATATATA | NM_009988 | coxsackievirus and adenovirus receptor | Cxadr | 2.8 | SM2240 | h | | 7 |
| mPool4_V2MM_122872 | 1.136649 | 75.83547 | TRUE | HP_34373 | TGCTGTTG | GAGATAGT | XM_142472 | similar to putative pheromone receptor (Go-VN4) | | 2.14 | SM2531 | c | | 12 |
| mPool2_V2MM_121432 | 1.136327 | 3.004939 | TRUE | HP_34231 | TGCTGTTG | GAGGTCAT | XM_488332 | similar to LRRG00119 | | 2.11 | SM2368 | b | | 12 |
| 092308m1_V2MM_179440 | 1.135939 | 3.004939 | TRUE | HP_39991 | TGCTGTTG | CATAATCA | XM_288334 | Mus musculus LOC331652 (LOC331652), mR. | | 2.5 | SM2085 | g | | 2 |

| | | | | | | | | | | | | | |
|---|---|---|---|---|---|---|---|---|---|---|---|---|---|
| mPool2_V2MM_141176 | 1.135592 | 2.205692 | TRUE | HP_36188 | TGCTGTTG | GAAACAA( | XM_156493 | Mus musculus LOC239875 (LOC239875), mR. | | 2.4 | SM2050 | e | 3 |
| mPool2_V2MM_141176 | 1.135592 | 2.205692 | TRUE | HP_36188 | TGCTGTTG | GAAACAA( | XM_156493 | Mus musculus LOC239875 (LOC239875), mRNA. | | 2.4 | SM2050 | e | 3 |
| mPool2_V2MM_8880 | 1.135526 | 3.004939 | TRUE | HP_23236 | TGCTGTTG | CAAGCGG. | NM_026480 | RIKEN cDNA 2410146L05 gene | 2410146L05Rik | 2.11 | SM2377 | e | 1 |
| 092308m1_V2MM_178954 | 1.135254 | 3.004939 | TRUE | HP_39942 | TGCTGTTG | GGAAGCA | XM_288248 | Mus musculus LOC332522 (LOC332522), mRNA. | | 2.5 | SM2081 | b | 1 |
| 092308m1_V2MM_178954 | 1.135254 | 3.004939 | TRUE | HP_39942 | TGCTGTTG | GGAAGCA | XM_288248 | Mus musculus LOC332522 (LOC332522), mR. | | 2.5 | SM2081 | b | 1 |
| 092308m3_V2MM_113039 | 1.135088 | 75.83547 | TRUE | HP_33403 | TGCTGTTG | CTGGGAG | XM_136780 | Mus musculus similar to translation elongation factor eEF-1 alpha chain - rat (LO( | | 2.13 | SM2490 | f | 2 |
| mPool2_V2MM_158991 | 1.134922 | 2.205692 | TRUE | HP_37959 | TGCTGTTG | GTAGTTTA | XM_483938 | RIKEN cDNA 1500026H17 gene | 1500026H17Rik | 2.6 | SM2131 | a | 8 |
| mPool5_V2MM_169518 | 1.13484 | 75.83547 | TRUE | HP_39001 | TGCTGTTG | CTGTTTCT | AK037372 | RIKEN cDNA A130012E19 gene | A130012E19Rik | NA | | | NA |
| 092308m3_V2MM_194363 | 1.134718 | 75.83547 | TRUE | HP_41253 | TGCTGTTG | CGGTGTTT | NM_178613 | RIKEN cD 4933433P14 gene | 4933433P14Rik | 2.9 | SM2268 | g | 2 |
| 092308m3_V2MM_194363 | 1.134718 | 75.83547 | TRUE | HP_41253 | TGCTGTTG | CGGTGTTT | NM_178613 | RIKEN cDNA 4933433P14 gene | 4933433P14Rik | 2.9 | SM2268 | g | 2 |
| mPool6_V2MM_68187 | 1.134322 | 75.83547 | TRUE | HP_29015 | TGCTGTTG | CTGTCCCT | NM_009774 | budding uninhibited by benzimidazoles 3 homolog (S. cer | Bub3 | 2.8 | SM2225 | a | 11 |
| 092308m1_V2MM_146447 | 1.134263 | 3.004939 | TRUE | HP_36715 | TGCTGTTG | CGGATTA/ | XM_162010 | Mus musculus LOC243064 (LOC243064), mR. | | 2.4 | SM2031 | a | 4 |
| mPool2_V2MM_144266 | 1.132677 | 3.004939 | TRUE | HP_36497 | TGCTGTTG | GACAATG | XM_159833 | Mus musculus LOC208237 (LOC208237), mRNA. | | 2.1 | SM2335 | h | 8 |
| 092308m1_V2MM_178872 | 1.132057 | 3.004939 | TRUE | HP_39934 | TGCTGTTG | CCCATCAT | XM_288234 | Mus musculus LOC332510 (LOC332510), mR. | | 2.5 | SM2092 | e | 4 |
| 092308m3_V2MM_113763 | 1.131893 | 75.83547 | TRUE | HP_33475 | TGCTGTTG | CATGACA( | XM_137098 | Mus musculus similar to Nucleolar protein Nop56 (Nucleolar protein 5A) (LOC21! | | 2.13 | SM2489 | g | 7 |
| mPool4_V2MM_201687 | 1.131885 | 75.83547 | TRUE | HP_41971 | TGCTGTTG | CAAAGCA | XM_288529 | Mus musculus LOC331914 (LOC331914), mRNA. | | NA | | | NA |
| 092308m1_V2MM_160758 | 1.131563 | 3.004939 | TRUE | HP_38133 | TGCTGTTG | CATCGTTA | XM_283630 | Mus musculus hypothetical gene supported by AK077805 (LOC329212), mR. | | 2.6 | SM2123 | b | 7 |
| mPool2_V2MM_165660 | 1.130546 | 3.004939 | TRUE | HP_38619 | TGCTGTTG | CATGAAC/ | XM_484793 | similar to Peptidyl-prolyl cis-trans isomerase A (PPIase) (Rotamase) (Cyclophilin A | | 2.6 | SM2130 | a | 6 |
| mPool5_V2MM_3632 | 1.130373 | 75.83547 | TRUE | HP_22722 | TGCTGTTG | GCATTGA( | XM_356218 | mannosidase 2, alpha B1 | Man2b1 | NA | | | NA |
| mPool2_V2MM_111512 | 1.130306 | 2.205692 | TRUE | HP_33252 | TGCTGTTG | GGTAAAC | XM_135893 | coagulation factor IX | F9 | 2.7 | SM2164 | b | 1 |
| mPool2_V2MM_3115 | 1.130195 | 3.004939 | TRUE | HP_22672 | TGCTGTTG | CTTGACAA | XM_136256 | Mus musculus similar to protease (LOC240779), mRNA. | | 2.11 | SM2387 | d | 12 |
| mPool2_V2MM_120783 | 1.129895 | 2.205692 | TRUE | HP_34167 | TGCTGTTG | CAGTAATT | XM_485031 | olfactory receptor 48 | Olfr48 | 2.11 | SM2362 | a | 2 |
| mPool4_V2MM_49706 | 1.12972 | 75.83547 | TRUE | HP_27214 | TGCTGTTG | GTCTGCAT | XM_357629 | similar to immunoglobulin light chain variable region | | 2.16 | SM2633 | c | 4 |
| mPool2_V2MM_135742 | 1.129156 | 3.004939 | TRUE | HP_35646 | TGCTGTTG | CAAACTAT | XM_151017 | Mus musculus LOC241032 (LOC241032), mR. | | 2.4 | SM2034 | h | 3 |
| mPool2_V2MM_135742 | 1.129156 | 3.004939 | TRUE | HP_35646 | TGCTGTTG | CAAACTAT | XM_151017 | Mus musculus LOC241032 (LOC241032), mRNA. | | 2.4 | SM2034 | h | 3 |
| 092308m3_V2MM_100181 | 1.128946 | 75.83547 | TRUE | HP_32137 | TGCTGTTG | GGCTAAG. | BC048675 | RIKEN cDNA 1700012B07 gene | 1700012B07Rik | 2.1 | SM2309 | e | 1 |
| 092308m3_V2MM_100181 | 1.128946 | 75.83547 | TRUE | HP_32137 | TGCTGTTG | GGCTAAG. | BC048675 | RIKEN cD 1700012B07 gene | 1700012B07Rik | 2.1 | SM2309 | e | 1 |
| mPool5_V2MM_20464 | 1.12862 | 75.83547 | TRUE | HP_24365 | TGCTGTTG | GTTGATG( | NM_007831 | deleted in colorectal carcinoma | Dcc | NA | | | NA |
| mPool7_V2MM_107239 | 1.127445 | 3.004939 | TRUE | HP_32832 | TGCTGTTG | GCACACTT | XM_131953 | similar to CG9004-PA | | 2.1 | SM2315 | c | 4 |
| mPool5_V2MM_139958 | 1.127332 | 35.99881 | TRUE | HP_36067 | TGCTGTTG | CGCCTAG/ | XM_155564 | similar to CUB and Sushi multiple domains 3 isoform 1 | | NA | | | NA |
| mPool4_V2MM_41437 | 1.127305 | 75.83547 | TRUE | HP_26407 | TGCTGTTG | CACATGCT | NM_153601 | glutamate-ammonia ligase (glutamine synthase) domain | Gluld1 | 2.16 | SM2601 | e | 11 |
| mPool4_V2MM_212650 | 1.127075 | 61.29828 | TRUE | HP_43031 | TGCTGTTG | GAGATATT | XM_140011 | Mus musculus similar to probable cell division control protein p55CDC - rat (LOC2 | | 2.16 | SM2648 | b | 10 |
| mPool7_V2MM_70926 | 1.126741 | 3.004939 | TRUE | HP_29282 | TGCTGTTG | CAGCTTGT | NM_009875 | cyclin-dependent kinase inhibitor 1B (P27) | Cdkn1b | 2.8 | SM2229 | b | 3 |
| 092308m3_V2MM_94440 | 1.126522 | 75.83547 | TRUE | HP_31572 | TGCTGTTG | CTCTTCAA | XM_487324 | polymerase (R) III (D directed) polypeptide A | Polr3a | 2.13 | SM2453 | e | 12 |
| mPool7_V2MM_201249 | 1.126381 | 3.004939 | TRUE | HP_41927 | TGCTGTTG | CATAGTTG | XM_289314 | Mus musculus LOC332963 (LOC332963), mRNA. | | 2.9 | SM2289 | f | 12 |
| mPool2_V2MM_758 | 1.126109 | 3.004939 | TRUE | HP_22441 | TGCTGTTG | CAGCATTA | NM_026577 | RIKEN cDNA C530009C10 gene | C530009C10Rik | 2.11 | SM2385 | h | 4 |
| mPool5_V2MM_18015 | 1.126075 | 75.83547 | TRUE | HP_24127 | TGCTGTTG | GCCACAG | NM_008289 | hydroxysteroid 11-beta dehydrogenase 2 | Hsd11b2 | NA | | | NA |
| mPool2_V2MM_121742 | 1.125719 | 2.205692 | TRUE | HP_34262 | TGCTGTTG | CATGGGC | XM_141826 | Mus musculus similar to RIKEN cDNA 2610301B20; EST AI428449 [Mus musculus] | | 2.11 | SM2351 | b | 4 |
| mPool7_V2MM_196145 | 1.125673 | 3.004939 | TRUE | HP_41425 | TGCTGTTG | GGCAAAT | NM_183219 | hypothetical protein B230218O03 | | 2.9 | SM2268 | h | 6 |
| mPool7_V2MM_196145 | 1.125673 | 3.004939 | TRUE | HP_41425 | TGCTGTTG | GGCAAAT | NM_183219 | hypothetical protein B230218O03 | | 2.9 | SM2268 | h | 6 |
| mPool2_V2MM_214369 | 1.125612 | 3.004939 | TRUE | HP_43196 | TGCTGTTG | GGAGGAT | XM_139471 | Mus musculus similar to envelope protein [Ovis aries] (LOC239411), mRNA. | | 2.11 | SM2358 | e | 7 |
| 092308m3_V2MM_54599 | 1.125167 | 75.83547 | TRUE | HP_27692 | TGCTGTTG | CATTTGG( | NM_025329 | RIKEN cD 0610012D17 gene | 0610012D17Rik | 2.12 | SM2407 | e | 3 |
| mPool4_V2MM_132731 | 1.124736 | 66.95036 | TRUE | HP_35348 | TGCTGTTG | CGTGCAC/ | XM_148860 | armadillo repeat containing 4 | Armc4 | 2.14 | SM2541 | e | 7 |
| 092308m3_V2MM_146128 | 1.124712 | 75.83547 | TRUE | HP_36683 | TGCTGTTG | CAGTCAA( | XM_161773 | hypothetical gene supported by AK036869 | | 2.14 | SM2519 | a | 11 |
| 092308m3_V2MM_105815 | 1.124543 | 75.83547 | TRUE | NA | NA | NA | NA | NA | NA | NA | NA | NA | NA |
| 092308m3_V2MM_203704 | 1.124481 | 75.83547 | TRUE | HP_42163 | TGCTGTTG | CCTACAAA | XM_139220 | similar to ribosomal protein S15; rat insulinoma gene | | 2.13 | SM2498 | b | 5 |
| 092308m3_V2MM_143822 | 1.124345 | 59.99801 | TRUE | HP_36453 | TGCTGTTG | GAAATGG. | XM_159171 | Mus musculus LOC245389 (LOC245389), mR. | | 2.14 | SM2526 | e | 4 |
| 092308m3_V2MM_87599 | 1.124285 | 75.83547 | TRUE | HP_30904 | TGCTGTTG | CCTGTGT( | NM_175203 | RIKEN cD 4833408A19 gene | 4833408A19Rik | 2.13 | SM2453 | g | 2 |
| 092308m3_V2MM_35636 | 1.124259 | 75.83547 | TRUE | HP_25842 | TGCTGTTG | CTATGTGA | XM_111356 | immunoglobulin heavy chain (V7183 family) | Igh-V7183 | 2.11 | SM2397 | h | 10 |
| 092308m3_V2MM_226127 | 1.123799 | 75.83547 | TRUE | HP_47815 | TGCTGTTG | CGAGCATT | XM_483874 | RIKEN cD 1700021A07 gene | 1700021A07Rik | 2.13 | SM2453 | g | 5 |
| 092308m3_V2MM_25093 | 1.122007 | 60.13437 | TRUE | HP_24817 | TGCTGTTG | AGAGTCC/ | AF190666 | limb region 1 | Lmbr1 | 2.12 | SM2433 | g | 7 |
| mPool2_V2MM_102558 | 1.121838 | 3.004939 | TRUE | HP_32370 | TGCTGTTG | GTGCATAT | NM_009558 | zinc finger protein 51 | Zfp51 | 2.7 | SM2160 | b | 5 |
| 092308m1_V2MM_131538 | 1.121298 | 3.004939 | TRUE | HP_35230 | TGCTGTTG | CCACCTCT | AK005401 | RIKEN cD 1600002O04 gene | 1600002O04Rik | 2.6 | SM2111 | a | 5 |

| | | | | | | | | | | | | | | | |
|---|---|---|---|---|---|---|---|---|---|---|---|---|---|---|---|
| mPool4_V2MM_213633 | 1.121283 | 75.83547 | TRUE | HP_43125 | TGCTGTTG | GTCATCCT | NM_198673 | cDNA sequence BC049697 | BC049697 | | 2.14 | SM2543 | a | | 8 |
| 092308m1_V2MM_159773 | 1.121147 | 3.004939 | TRUE | HP_38036 | TGCTGTTG | CAACCAAT | XM_283241 | Mus musculus hypothetical gene supported by AK034333 (LOC328474), mR. | | | 2.6 | SM2117 | h | | 9 |
| mPool4_V2MM_25400 | 1.120282 | 75.83547 | TRUE | HP_24846 | TGCTGTTG | CAGAGGA | NM_145601 | cDNA sequence BC016201 | BC016201 | | 2.15 | SM2589 | b | | 6 |
| 092308m3_V2MM_38398 | 1.120219 | 59.99801 | TRUE | HP_26112 | TGCTGTTG | GCAGCCAT | AK044117 | RIKEN cD C430014H23 gene | C430014H23Rik | | 2.12 | SM2418 | h | | 7 |
| mPool2_V2MM_213506 | 1.12014 | 2.205692 | TRUE | HP_43113 | TGCTGTTG | GTCAAATT | XM_139486 | Mus musculus LOC239417 (LOC239417), mRNA. | | | 2.11 | SM2357 | b | | 2 |
| mPool2_V2MM_84812 | 1.120085 | 3.004939 | TRUE | HP_30631 | TGCTGTTG | CAGTAAAC | NM_029498 | zinc finger protein 198 | Zfp198 | | 2.6 | SM2141 | f | | 10 |
| 092308m3_V2MM_22500 | 1.119573 | 75.83547 | TRUE | HP_24565 | TGCTGTTG | CGTGTGTC | NM_009120 | SAR1a gene homolog 1 (S. cerevisiae) | Sara1 | | 2.11 | SM2400 | a | | 7 |
| 092308m3_V2MM_28986 | 1.119553 | 75.83547 | TRUE | HP_14238 | TGCTGTTG | GTGAAGC | NM_146150 | rdilysin, N-arginine dibasic convertase, NRD convertase 1 | Nrd1 | | 2.11 | SM2398 | d | | 1 |
| mPool2_V2MM_86459 | 1.119209 | 3.004939 | TRUE | HP_30793 | TGCTGTTG | CTGTGTCT | NM_172474 | RIKEN cDNA 5230400J09 gene | 5230400J09Rik | | 2.6 | SM2134 | g | | 1 |
| mPool4_V2MM_57320 | 1.118912 | 69.72441 | TRUE | HP_27957 | TGCTGTTG | CTTGAAGA | NM_172762 | DNA segment, Chr 8, ERATO Doi 233, expressed | D8Ertd233e | | 2.15 | SM2585 | d | | 9 |
| mPool2_V2MM_132235 | 1.118679 | 3.004939 | TRUE | HP_35298 | TGCTGTTG | CAGGAAA | AK045005 | Casitas B-lineage lymphoma b | Cblb | | 2.11 | SM2351 | a | | 3 |
| mPool5_V2MM_5387 | 1.11808 | 75.83547 | TRUE | HP_22893 | TGCTGTTG | CCCAGGG | NM_019466 | Down syndrome critical region homolog 1 (human) | Dscr1 | NA | | | | NA | |
| mPool6_V2MM_63892 | 1.118066 | 75.83547 | TRUE | HP_28595 | TGCTGTTG | CTCCCAGT | NM_030150 | DNA segment, Chr 11, Lothar Hennighausen 2, expressed | D11Lgp2e | | 2.8 | SM2219 | h | | 6 |
| mPool5_V2MM_7596 | 1.117929 | 73.80895 | TRUE | HP_23110 | TGCTGTTG | GCAGTTGA | NM_007674 | caudal type homeo box 4 | Cdx4 | NA | | | | NA | |
| mPool7_V2MM_203147 | 1.117295 | 3.004939 | TRUE | HP_42111 | TGCTGTTG | CAGCTGA | XM_130845 | RIKEN cDNA E430012K20 gene | E430012K20Rik | | 2.9 | SM2297 | h | | 12 |
| 092308m3_V2MM_83794 | 1.117049 | 75.83547 | TRUE | HP_30531 | TGCTGTTG | CCAATTCT | NM_025499 | RIKEN cD 1700027M21 gene | 1700027M21Rik | | 2.13 | SM2468 | g | | 11 |
| 092308m1_V2MM_173519 | 1.116759 | 1.959154 | TRUE | HP_39400 | TGCTGTTG | CAAGAAAT | XM_287042 | gene model 819, (NCBI) | Gm819 | | 2.5 | SM2082 | e | | 10 |
| mPool5_V2MM_166083 | 1.116696 | 75.83547 | TRUE | HP_38661 | TGCTGTTG | GCTGATGT | XM_487675 | similar to neonatal submandibular gland protein B | | NA | | | | NA | |
| mPool5_V2MM_13730 | 1.116324 | 59.99801 | TRUE | NA | NA | NA | NA | NA | NA | NA | | NA | NA | NA | |
| mPool2_V2MM_28792 | 1.116099 | 2.205692 | TRUE | HP_25177 | TGCTGTTG | GTAAGAA | NM_011762 | zinc finger protein 59 | Zfp59 | | 2.11 | SM2389 | e | | 11 |
| mPool5_V2MM_6492 | 1.116082 | 75.83547 | TRUE | HP_23002 | TGCTGTTG | CTGGCTTT | NM_011756 | zinc finger protein 36 | Zfp36 | NA | | | | NA | |
| 092308m3_V2MM_116394 | 1.115923 | 73.80895 | TRUE | HP_33734 | TGCTGTTG | CAAAGTTC | XM_138909 | Mus musculus similar to endogenous retroviral family W, env(C7), member 1 (syr | | | 2.13 | SM2487 | c | | 3 |
| mPool7_V2MM_149886 | 1.115651 | 3.004939 | TRUE | HP_37059 | TGCTGTTG | CAGGGTTT | XM_165303 | Mus musculus LOC208460 (LOC208460), mR. | | | 2.1 | SM2331 | a | | 8 |
| mPool7_V2MM_149886 | 1.115651 | 3.004939 | TRUE | HP_37059 | TGCTGTTG | CAGGGTTT | XM_165303 | Mus musculus LOC208460 (LOC208460), mRNA. | | | 2.1 | SM2331 | a | | 8 |
| mPool2_V2MM_86642 | 1.115368 | 2.205692 | TRUE | HP_30811 | TGCTGTTG | CATAGGA | NM_173746 | hypothetical protein C130086A10 | | | 2.6 | SM2141 | c | | 5 |
| mPool7_V2MM_67874 | 1.11536 | 1.959154 | TRUE | HP_28984 | TGCTGTTG | CTCTAAGT | NM_030014 | hook homolog 1 (Drosophila) | Hook1 | | 2.8 | SM2241 | c | | 5 |
| 092308m1_V2MM_70780 | 1.115202 | 3.004939 | TRUE | HP_29268 | TGCTGTTG | CAGAGCA | XM_195466 | gene model 659, (NCBI) | Gm659 | | 2.3 | SM2004 | g | | 8 |
| mPool6_V2MM_64852 | 1.115182 | 75.83547 | TRUE | HP_28690 | TGCTGTTG | GAACTTTC | NM_007435 | ATP-binding cassette, sub-family D (ALD), member 1 | Abcd1 | | 2.7 | SM2181 | c | | 1 |
| 092308m1_V2MM_134577 | 1.115112 | 3.004939 | TRUE | HP_35530 | TGCTGTTG | CCTCAAAT | AK017247 | RIKEN cD 5430400N05 gene | 5430400N05Rik | | 2.6 | SM2113 | e | | 11 |
| mPool2_V2MM_120124 | 1.114877 | 3.004939 | TRUE | HP_34103 | TGCTGTTG | GAGAGAA | XM_484819 | cDNA sequence BC029127 | BC029127 | | 2.11 | SM2367 | f | | 10 |
| mPool4_V2MM_179286 | 1.114359 | 75.83547 | TRUE | HP_39976 | TGCTGTTG | CTGTCTGT | XM_288305 | Mus musculus LOC331613 (LOC331613), mRNA. | | NA | | | | NA | |
| mPool7_V2MM_93767 | 1.113544 | 3.004939 | TRUE | HP_31506 | TGCTGTTG | CAGTTCTT | NM_181754 | G protein-coupled receptor 141 | Gpr141 | | 2.9 | SM2270 | f | | 7 |
| 092308m3_V2MM_43600 | 1.113469 | 75.83547 | TRUE | HP_26617 | TGCTGTTG | CTCTTTGT | NM_016804 | metaxin 2 | Mtx2 | | 2.12 | SM2417 | b | | 8 |
| 092308m3_V2MM_29298 | 1.113445 | 75.83547 | TRUE | HP_25226 | TGCTGTTG | CAAAGCA | NM_146145 | Janus kise 1 | Jak1 | | 2.12 | SM2423 | c | | 8 |
| 092308m1_V2MM_96559 | 1.113225 | 2.205692 | TRUE | HP_31780 | TGCTGTTG | CACCTATA | XM_110852 | amyotrophic lateral sclerosis 2 (juvenile) chromosome re | Als2cr19 | | 2.6 | SM2110 | c | | 11 |
| mPool2_V2MM_130410 | 1.113144 | 3.004939 | TRUE | HP_35119 | TGCTGTTG | CATAACAC | XM_146855 | similar to BCL2/adenovirus E1B 19kDa-interacting protein 1, NIP2; BCL2/adenovi | | | 2.11 | SM2365 | h | | 6 |
| mPool2_V2MM_2146 | 1.113022 | 2.205692 | TRUE | HP_22577 | TGCTGTTG | CTCACACA | NM_010743 | interleukin 1 receptor-like 1 | Il1rl1 | | 2.11 | SM2387 | f | | 10 |
| mPool7_V2MM_197100 | 1.112647 | 2.205692 | TRUE | HP_41518 | TGCTGTTG | CTCTTACC | AK083918 | spastic paraplegia 3A homolog (human) | Spg3a | | 2.9 | SM2254 | g | | 9 |
| mPool7_V2MM_197100 | 1.112647 | 2.205692 | TRUE | HP_41518 | TGCTGTTG | CTCTTACC | AK083918 | spastic paraplegia 3A homolog (human) | Spg3a | | 2.9 | SM2254 | g | | 9 |
| 092308m3_V2MM_35016 | 1.112546 | 59.99801 | TRUE | HP_25782 | TGCTGTTG | CCTGTGAA | NM_009322 | T-box brain gene 1 | Tbr1 | | 2.11 | SM2394 | c | | 12 |
| 092308m3_V2MM_210322 | 1.112281 | 59.99801 | TRUE | HP_42807 | TGCTGTTG | GAGAACTC | XM_162001 | Mus musculus LOC243061 (LOC243061), mR. | | | 2.14 | SM2519 | b | | 8 |
| mPool2_V2MM_125363 | 1.11227 | 2.205692 | TRUE | HP_34620 | TGCTGTTG | GACATTAC | NM_001004178 | hypothetical gene supported by BC064115 | | | 2.11 | SM2361 | a | | 11 |
| 092308m3_V2MM_47433 | 1.111944 | 61.29828 | TRUE | HP_26991 | TGCTGTTG | CTGCATTG | NM_145842 | vomerosal 1 receptor, E9 | V1re9 | | 2.12 | SM2445 | h | | 12 |
| mPool5_V2MM_8676 | 1.111489 | 75.83547 | TRUE | HP_23215 | TGCTGTTG | GTTAAGCA | NM_153091 | suppression of tumorigenicity 7-like | St7l | NA | | | | NA | |
| mPool6_V2MM_164021 | 1.111095 | 75.83547 | TRUE | HP_38457 | TGCTGTTG | CATCTTGG | XM_284905 | Mus musculus similar to pORF1 [Plasmodium yoelii yoelii] (LOC331722), mRNA. | | | 2.7 | SM2192 | e | | 2 |
| mPool7_V2MM_203319 | 1.109669 | 3.004939 | TRUE | HP_30983 | TGCTGTTG | CTCTTCTC | NM_175397 | RIKEN cD 5830484A20 gene | 5830484A20Rik | | 2.9 | SM2269 | b | | 7 |
| mPool7_V2MM_203319 | 1.109669 | 3.004939 | TRUE | HP_30983 | TGCTGTTG | CTCTTCTC | NM_175397 | RIKEN cDNA 5830484A20 gene | 5830484A20Rik | | 2.9 | SM2269 | b | | 7 |
| mPool6_V2MM_193872 | 1.109488 | 75.83547 | TRUE | HP_37932 | TGCTGTTG | CTATGAAC | XM_289920 | filamin, alpha | Flna | | 2.8 | SM2205 | g | | 10 |
| mPool6_V2MM_64404 | 1.109358 | 75.83547 | TRUE | HP_28645 | TGCTGTTG | CTGGCCCT | NM_009954 | breast cancer anti-estrogen resistance 1 | Bcar1 | | 2.7 | SM2178 | b | | 5 |
| mPool2_V2MM_113470 | 1.109327 | 1.959154 | TRUE | HP_33445 | TGCTGTTG | CTGACATC | XM_136985 | Mus musculus similar to triosephosphate isomerase 1 (LOC215969), mRNA. | | | 2.7 | SM2157 | b | | 11 |
| 092308m1_V2MM_156055 | 1.108968 | 3.004939 | TRUE | HP_37669 | TGCTGTTG | GAAGATCC | XM_198073 | Mus musculus LOC272426 (LOC272426), mR. | | | 2.6 | SM2122 | f | | 3 |
| mPool5_V2MM_154160 | 1.108963 | 75.83547 | TRUE | HP_37480 | TGCTGTTG | CACAGTGT | XM_196624 | Mus musculus LOC271637 (LOC271637), mRNA. | | NA | | | | NA | |
| mPool4_V2MM_262148 | 1.108854 | 75.83547 | TRUE | HP_52291 | TGCTGTTG | GCCTGCTT | NM_138658 | cDNA sequence AF085738 | AF085738 | | 2.15 | SM2576 | e | | 8 |

| | | | | | | | | | | | | | |
|---|---|---|---|---|---|---|---|---|---|---|---|---|---|
| mPool2_V2MM_11459 | 1.108714 | 3.004939 | TRUE | HP_23486 | TGCTGTTG | GAGTCAAT | NM_028018 | RIKEN cDNA 2400003C14 gene | 2400003C14Rik | 2.11 | SM2374 | f | 10 |
| mPool2_V2MM_86999 | 1.108382 | 3.004939 | TRUE | HP_10675 | TGCTGTTG | CAGATAAA | NM_174996 | DNA segment, Chr 4, ERATO Doi 22, expressed | D4Ertd22e | 2.6 | SM2107 | c | 11 |
| mPool2_V2MM_86999 | 1.108382 | 3.004939 | TRUE | HP_10675 | TGCTGTTG | CAGATAAA | NM_174996 | D segment, Chr 4, ERATO Doi 22, expressed | D4Ertd22e | 2.6 | SM2107 | c | 11 |
| 092308m1_V2MM_183421 | 1.108179 | 2.205692 | TRUE | HP_40389 | TGCTGTTG | GAACCTAC | XM_289016 | Mus musculus LOC332676 (LOC332676), mR. | | 2.5 | SM2094 | d | 7 |
| 092308m1_V2MM_183443 | 1.107854 | 3.004939 | TRUE | HP_40391 | TGCTGTTG | CACTGAGC | XM_289020 | Mus musculus LOC332650 (LOC332650), mR. | | 2.5 | SM2078 | b | 3 |
| mPool2_V2MM_6003 | 1.107806 | 2.205692 | TRUE | HP_22954 | TGCTGTTG | CGTCTTTG | NM_008410 | integral membrane protein 2B | Itm2b | 2.11 | SM2382 | g | 3 |
| 092308m1_V2MM_139106 | 1.107258 | 3.004939 | TRUE | HP_35982 | TGCTGTTG | CATCTTGC | XM_154777 | Mus musculus LOC239075 (LOC239075), mRNA. | | 2.4 | SM2043 | c | 6 |
| 092308m1_V2MM_139106 | 1.107258 | 3.004939 | TRUE | HP_35982 | TGCTGTTG | CATCTTGC | XM_154777 | Mus musculus LOC239075 (LOC239075), mR. | | 2.4 | SM2043 | c | 6 |
| 092308m3_V2MM_147852 | 1.107204 | 75.83547 | TRUE | HP_36856 | TGCTGTTG | CCAAGAAA | XM_163317 | Mus musculus LOC215331 (LOC215331), mR. | | 2.14 | SM2517 | c | 10 |
| 092308m3_V2MM_41221 | 1.106744 | 75.83547 | TRUE | HP_26386 | TGCTGTTG | CAAGATGT | NM_016774 | ATP synthase, H+ transporting mitochondrial F1 complex | Atp5b | 2.12 | SM2447 | d | 1 |
| mPool2_V2MM_203365 | 1.106155 | 3.004939 | TRUE | HP_42131 | TGCTGTTG | GAAATTTA | XM_137252 | Mus musculus similar to nuclear transport factor 2 [Rattus norvegicus] (LOC2375 | | 2.1 | SM2318 | b | 12 |
| mPool6_V2MM_160401 | 1.106057 | 75.83547 | TRUE | HP_38098 | TGCTGTTG | CATCTAAT | NM_175506 | a disintegrin-like and metalloprotease (reprolysin type) w | Adamts19 | 2.8 | SM2210 | g | 6 |
| 092308m3_V2MM_58035 | 1.105898 | 75.83547 | TRUE | HP_30978 | TGCTGTTG | GACTTATT | NM_016716 | cullin 3 | Cul3 | 2.12 | SM2413 | a | 8 |
| mPool2_V2MM_101136 | 1.105848 | 1.959154 | TRUE | HP_32231 | TGCTGTTG | GCATTTAT | AK019536 | RIKEN cDNA 4921504N20 gene | 4921504N20Rik | 2.7 | SM2159 | a | 1 |
| 092308m3_V2MM_85318 | 1.105816 | 59.99801 | TRUE | HP_30680 | TGCTGTTG | CTGGGAAC | NM_133979 | expressed sequence AI604832 | AI604832 | 2.13 | SM2465 | e | 8 |
| 092308m1_V2MM_132170 | 1.105423 | 3.004939 | TRUE | HP_35292 | TGCTGTTG | GGTAGCGT | XM_148178 | RIKEN cD D930038M13 gene | D930038M13Rik | 2.6 | SM2106 | a | 9 |
| mPool2_V2MM_126677 | 1.105078 | 3.004939 | TRUE | HP_34751 | TGCTGTTG | CAAGTCAA | XM_144734 | Mus musculus similar to RNA helicase TNZ2 - mouse (LOC245881), mRNA. | | 2.11 | SM2352 | a | 8 |
| mPool7_V2MM_204465 | 1.104199 | 2.205692 | TRUE | HP_42236 | TGCTGTTG | GCTGTTAA | AK032192 | potassium inwardly-rectifying channel, subfamily J, mem | Kcnj16 | 2.9 | SM2297 | g | 9 |
| 092308m1_V2MM_76998 | 1.103889 | 3.004939 | TRUE | HP_29873 | TGCTGTTG | CTCTCTTA | NM_146641 | olfactory receptor 1164 | Olfr1164 | 2.3 | SM2011 | c | 4 |
| mPool2_V2MM_206908 | 1.103539 | 2.205692 | TRUE | HP_42472 | TGCTGTTG | GAAATGAT | XM_157374 | Mus musculus LOC240302 (LOC240302), mRNA. | | 2.1 | SM2341 | a | 2 |
| mPool4_V2MM_43506 | 1.103405 | 75.83547 | TRUE | HP_26608 | TGCTGTTG | CTCTCTTG | NM_010178 | FUS interacting protein (serine-arginine rich) 1 | Fusip1 | 2.15 | SM2570 | g | 6 |
| 092308m1_V2MM_185131 | 1.102773 | 3.004939 | TRUE | HP_40560 | TGCTGTTG | CTACTTGA | XM_289316 | Mus musculus LOC332973 (LOC332973), mR. | | 2.5 | SM2066 | f | 10 |
| 092308m1_V2MM_185131 | 1.102773 | 3.004939 | TRUE | HP_40560 | TGCTGTTG | CTACTTGA | XM_289316 | Mus musculus LOC332973 (LOC332973), mRNA. | | 2.5 | SM2066 | f | 10 |
| mPool2_V2MM_213256 | 1.102731 | 3.004939 | TRUE | HP_43089 | TGCTGTTG | GAAATACA | BC066073 | EST AA407452 | AA407452 | 2.1 | SM2350 | c | 6 |
| mPool2_V2MM_114532 | 1.101883 | 3.004939 | TRUE | HP_33550 | TGCTGTTG | CTGGAAAC | XM_137579 | Mus musculus similar to olfactory receptor 30; olfactory receptor MOR281-1 [Mu | | 2.1 | SM2318 | g | 10 |
| mPool4_V2MM_28237 | 1.101222 | 75.83547 | TRUE | HP_25123 | TGCTGTTG | GCCAGCAT | NM_172262 | amine oxidase, flavin containing 1 | Aof1 | 2.15 | SM2594 | e | 7 |
| 092308m3_V2MM_93505 | 1.100611 | 75.83547 | TRUE | HP_31481 | TGCTGTTG | CACTTGAC | NM_181412 | zinc finger, BED domain containing 4 | Zbed4 | 2.13 | SM2455 | e | 10 |
| mPool7_V2MM_89836 | 1.100438 | 2.205692 | TRUE | HP_31126 | TGCTGTTG | GTGAATTA | BC048817 | RIKEN cDNA A730098D12 gene | A730098D12Rik | 2.9 | SM2270 | d | 5 |
| 092308m3_V2MM_35142 | 1.100327 | 62.64426 | TRUE | HP_25794 | TGCTGTTG | CGACCTCC | NM_144902 | solute carrier family 35 (UDP-N-acetylglucosamine (UDP- | Slc35a3 | 2.11 | SM2393 | g | 9 |
| 092308m3_V2MM_44493 | 1.099446 | 75.83547 | TRUE | HP_28639 | TGCTGTTG | GAGATATT | NM_008978 | protein tyrosine phosphatase, non-receptor type 20 | Ptpn20 | 2.12 | SM2441 | e | 3 |
| mPool4_V2MM_17740 | 1.099198 | 61.29828 | TRUE | HP_24100 | TGCTGTTG | GCAACAGA | NM_145363 | oculospanin | | 2.15 | SM2587 | g | 7 |
| 092308m3_V2MM_23726 | 1.099118 | 75.83547 | TRUE | HP_24685 | TGCTGTTG | CTTGTTGT | NM_172272 | cD sequence BC027073 | BC027073 | 2.12 | SM2428 | f | 9 |
| 092308m3_V2MM_58066 | 1.0986 | 75.83547 | TRUE | HP_28030 | TGCTGTTG | GACTTTCT | NM_025391 | RIKEN cD 1110017C15 gene | 1110017C15Rik | 2.12 | SM2414 | a | 9 |
| 092308m3_V2MM_103631 | 1.098363 | 75.83547 | TRUE | NA | NA | NA | NA | NA | NA | NA | NA | NA | NA |
| 092308m3_V2MM_107543 | 1.09827 | 75.83547 | TRUE | HP_32862 | TGCTGTTG | CCGCAGAT | XM_284166 | RIKEN cD 5430427O21 gene | 5430427O21Rik | 2.14 | SM2503 | a | 6 |
| 092308m1_V2MM_120808 | 1.098122 | 3.004939 | TRUE | HP_34169 | TGCTGTTG | CTCAGAAA | XM_141251 | Mus musculus similar to cyclophilin A [Felis catus] (LOC241552), mR. | | 2.5 | SM2098 | b | 9 |
| 092308m1_V2MM_120808 | 1.098122 | 3.004939 | TRUE | HP_34169 | TGCTGTTG | CTCAGAAA | XM_141251 | Mus musculus similar to cyclophilin A [Felis catus] (LOC241552), mRNA. | | 2.5 | SM2098 | b | 9 |
| 092308m1_V2MM_165344 | 1.0978 | 1.959154 | TRUE | HP_38588 | TGCTGTTG | CGTCATTA | XM_356940 | similar to 60S ribosomal protein L9 | | 2.6 | SM2123 | f | 6 |
| mPool4_V2MM_262197 | 1.097653 | 75.83547 | TRUE | HP_52166 | TGCTGTTG | CTGATGCT | NM_199033 | expressed sequence AU067695 | AU067695 | 2.16 | SM2607 | e | 2 |
| 092308m1_V2MM_83706 | 1.097111 | 3.004939 | TRUE | HP_30523 | TGCTGTTG | CAGGAAAT | NM_023900 | pleckstrin homology domain containing, family J member | Plekhj1 | 2.6 | SM2113 | b | 2 |
| mPool2_V2MM_159987 | 1.097057 | 3.004939 | TRUE | HP_38058 | TGCTGTTG | GTCCCAAA | NM_198629 | hypothetical gene supported by AK045595 | | 2.6 | SM2131 | f | 3 |
| mPool2_V2MM_137311 | 1.096773 | 3.004939 | TRUE | HP_35803 | TGCTGTTG | CCAAGGAA | XM_153003 | Mus musculus LOC237998 (LOC237998), mRNA. | | 2.1 | SM2333 | g | 12 |
| 092308m1_V2MM_73791 | 1.096688 | 3.004939 | TRUE | HP_29561 | TGCTGTTG | GTCTCTGC | NM_009849 | ectonucleoside triphosphate diphosphohydrolase 2 | Entpd2 | 2.3 | SM2003 | g | 9 |
| 092308m1_V2MM_151927 | 1.095985 | 2.205692 | TRUE | HP_37259 | TGCTGTTG | CTGTTTAT | XM_195000 | expressed sequence AI413631 | AI413631 | 2.4 | SM2041 | e | 8 |
| 092308m3_V2MM_111750 | 1.095825 | 66.95036 | TRUE | NA | NA | NA | NA | NA | NA | NA | NA | NA | NA |
| mPool6_V2MM_68198 | 1.095784 | 75.83547 | TRUE | HP_29016 | TGCTGTTG | CTGTCTTT | NM_146930 | olfactory receptor 791 | Olfr791 | 2.8 | SM2216 | f | 7 |
| mPool2_V2MM_546 | 1.095609 | 3.004939 | TRUE | HP_22421 | TGCTGTTG | CACTGTGT | NM_007664 | cadherin 2 | Cdh2 | 2.7 | SM2165 | c | 10 |
| mPool2_V2MM_133000 | 1.095114 | 3.004939 | TRUE | HP_35374 | TGCTGTTG | GTTACATA | AK047658 | RIKEN cDNA C130080N23 gene | C130080N23Rik | 2.11 | SM2361 | g | 3 |
| mPool4_V2MM_43999 | 1.094979 | 75.83547 | TRUE | HP_26656 | TGCTGTTG | CTGTATTT | XM_139056 | gene model 290, (NCBI) | Gm290 | 2.16 | SM2617 | h | 6 |
| mPool2_V2MM_5165 | 1.094798 | 3.004939 | TRUE | HP_22872 | TGCTGTTG | CCAACCAA | NM_018768 | syntaxin 8 | Stx8 | 2.11 | SM2385 | f | 6 |
| mPool5_V2MM_8350 | 1.094178 | 59.99801 | TRUE | HP_23183 | TGCTGTTG | GTCAGTTA | NM_008223 | serine (or cysteine) proteinase inhibitor, clade D, membe | Serpind1 | NA | | | NA |
| mPool2_V2MM_121889 | 1.093951 | 2.205692 | TRUE | HP_34276 | TGCTGTTG | CTCTGAAC | XM_141920 | similar to hypothetical protein MGC37588 | | 2.11 | SM2363 | c | 5 |
| mPool2_V2MM_155145 | 1.093726 | 2.205692 | TRUE | HP_37578 | TGCTGTTG | CCTGTTAC | XM_197224 | Mus musculus LOC268804 (LOC268804), mRNA. | | 2.6 | SM2131 | h | 12 |

| | | | | | | | | | | | | | |
|---|---|---|---|---|---|---|---|---|---|---|---|---|---|
| 092308m1_V2MM_175932 | 1.093374 | 3.004939 | TRUE | HP_39641 | TGCTGTTG | CAGGAAT | XM_287564 | Mus musculus hypothetical gene supported by AK089795 (LOC330271), mRNA. | | 2.5 | SM2072 | b | 10 |
| 092308m1_V2MM_175932 | 1.093374 | 3.004939 | TRUE | HP_39641 | TGCTGTTG | CAGGAAT | XM_287564 | Mus musculus hypothetical gene supported by AK089795 (LOC330271), mR. | | 2.5 | SM2072 | b | 10 |
| 092308m3_V2MM_47249 | 1.093243 | 60.13437 | TRUE | HP_26973 | TGCTGTTG | CTCGGAAT | NM_177177 | RIKEN cD 5930403N24 gene | 5930403N24Rik | 2.12 | SM2441 | g | 8 |
| mPool4_V2MM_45829 | 1.092768 | 75.83547 | TRUE | HP_26834 | TGCTGTTG | CAGAAGT | XM_489874 | similar to TRAV9-1 | | NA | | | NA |
| 092308m1_V2MM_148520 | 1.09275 | 2.205692 | TRUE | HP_36923 | TGCTGTTG | CTGTTTAT | XM_163870 | Mus musculus LOC234017 (LOC234017), mR. | | 2.4 | SM2035 | f | 9 |
| 092308m3_V2MM_113083 | 1.092457 | 75.83547 | TRUE | HP_33407 | TGCTGTTG | GCCATAA | XM_136816 | similar to 60S ribosomal protein L35 | | 2.14 | SM2503 | f | 5 |
| mPool2_V2MM_154208 | 1.092442 | 3.004939 | TRUE | HP_37484 | TGCTGTTG | CTATGAAC | XM_196676 | Mus musculus LOC271710 (LOC271710), mRNA. | | 2.6 | SM2132 | b | 3 |
| mPool7_V2MM_178954 | 1.092104 | 3.004939 | TRUE | HP_39942 | TGCTGTTG | GGAAGCA | XM_288248 | Mus musculus LOC332522 (LOC332522), mR. | | 2.5 | SM2081 | b | 1 |
| mPool7_V2MM_178954 | 1.092104 | 3.004939 | TRUE | HP_39942 | TGCTGTTG | GGAAGCA | XM_288248 | Mus musculus LOC332522 (LOC332522), mRNA. | | 2.5 | SM2081 | b | 1 |
| 092308m3_V2MM_194272 | 1.091981 | 75.83547 | TRUE | HP_41244 | TGCTGTTG | CCAGGAT | NM_175350 | RIKEN cD 4933402B14 gene | 4933402B14Rik | 2.13 | SM2460 | h | 5 |
| mPool2_V2MM_143000 | 1.09158 | 3.004939 | TRUE | HP_36371 | TGCTGTTG | CTGTTAAC | XM_158204 | Mus musculus LOC241380 (LOC241380), mR. | | 2.4 | SM2050 | g | 5 |
| mPool2_V2MM_143000 | 1.09158 | 3.004939 | TRUE | HP_36371 | TGCTGTTG | CTGTTAAC | XM_158204 | Mus musculus LOC241380 (LOC241380), mRNA. | | 2.4 | SM2050 | g | 5 |
| 092308m1_V2MM_122300 | 1.091047 | 2.205692 | TRUE | HP_34316 | TGCTGTTG | GTAATAA | XM_142110 | gene model 381, (NCBI) | Gm381 | 2.6 | SM2103 | d | 10 |
| 092308m1_V2MM_122300 | 1.091047 | 2.205692 | TRUE | HP_34316 | TGCTGTTG | GTAATAA | XM_142110 | gene model 381, (NCBI) | Gm381 | 2.6 | SM2103 | d | 10 |
| mPool7_V2MM_142721 | 1.090622 | 3.004939 | TRUE | HP_36343 | TGCTGTTG | CAGATTCA | XM_157862 | Mus musculus LOC240631 (LOC240631), mR. | | 2.5 | SM2051 | h | 12 |
| mPool7_V2MM_142721 | 1.090622 | 3.004939 | TRUE | HP_36343 | TGCTGTTG | CAGATTCA | XM_157862 | Mus musculus LOC240631 (LOC240631), mRNA. | | 2.5 | SM2051 | h | 12 |
| 092308m3_V2MM_146518 | 1.090149 | 75.83547 | TRUE | HP_36722 | TGCTGTTG | GGCTGTG | XM_162119 | Mus musculus LOC231511 (LOC231511), mR. | | 2.14 | SM2510 | b | 7 |
| mPool4_V2MM_133211 | 1.090085 | 75.83547 | TRUE | NA | NA | NA | NA | NA | NA | NA | NA | NA | NA |
| 092308m1_V2MM_173837 | 1.08963 | 3.004939 | TRUE | HP_39432 | TGCTGTTG | CGTGTTCC | XM_287101 | Mus musculus hypothetical gene supported by AK030988 (LOC329361), mR. | | 2.5 | SM2084 | e | 3 |
| mPool4_V2MM_128749 | 1.089256 | 59.99801 | TRUE | HP_34956 | TGCTGTTG | CTATGAGT | XM_145899 | Mus musculus similar to putative pheromone receptor (Go-VN3) (LOC244115), m | | 2.14 | SM2550 | d | 6 |
| mPool2_V2MM_137685 | 1.08918 | 3.004939 | TRUE | HP_35840 | TGCTGTTG | CTCATATT | XM_153460 | Mus musculus LOC238186 (LOC238186), mRNA. | | 2.1 | SM2334 | d | 10 |
| 092308m1_V2MM_93507 | 1.089168 | 3.004939 | TRUE | HP_31482 | TGCTGTTG | GTAAAGC | NM_181413 | ankyrin repeat and SAM domain containing 1 | Anks1 | 2.6 | SM2106 | e | 11 |
| mPool4_V2MM_42825 | 1.089081 | 61.29828 | TRUE | HP_26542 | TGCTGTTG | CGCGGTTT | NM_028688 | RIKEN cDNA 1700029M20 gene | 1700029M20Rik | 2.15 | SM2583 | c | 8 |
| 092308m3_V2MM_98396 | 1.088204 | 75.83547 | TRUE | HP_31962 | TGCTGTTG | GATACTGC | XM_112451 | Mus musculus LOC195654 (LOC195654), mR. | | 2.13 | SM2477 | g | 11 |
| mPool2_V2MM_147640 | 1.087987 | 3.004939 | TRUE | HP_36835 | TGCTGTTG | CTGTCTAA | XM_163135 | Mus musculus LOC243740 (LOC243740), mRNA. | | 2.4 | SM2046 | b | 10 |
| mPool2_V2MM_147640 | 1.087987 | 3.004939 | TRUE | HP_36835 | TGCTGTTG | CTGTCTAA | XM_163135 | Mus musculus LOC243740 (LOC243740), mR. | | 2.4 | SM2046 | b | 10 |
| mPool4_V2MM_177587 | 1.087748 | 75.83547 | TRUE | HP_39806 | TGCTGTTG | GCTGTGAT | XM_287909 | Mus musculus LOC330964 (LOC330964), mRNA. | | NA | | | NA |
| mPool2_V2MM_97303 | 1.087746 | 2.205692 | TRUE | HP_38644 | TGCTGTTG | CAGATTTA | XM_146592 | Mus musculus similar to protease [Mus musculus] (LOC210103), mRNA. | | 2.6 | SM2118 | c | 9 |
| 092308m3_V2MM_233631 | 1.087213 | 75.83547 | TRUE | HP_48553 | TGCTGTTG | CGAAAGG | XM_136764 | similar to RNP particle component | | 2.14 | SM2509 | e | 3 |
| mPool2_V2MM_142284 | 1.087074 | 3.004939 | TRUE | HP_36299 | TGCTGTTG | GCAAATGT | XM_157529 | Mus musculus LOC240393 (LOC240393), mRNA. | | 2.1 | SM2338 | a | 11 |
| mPool6_V2MM_224135 | 1.086731 | 75.83547 | TRUE | HP_47621 | TGCTGTTG | CCCATACA | XM_195628 | vomeronasal 1 receptor, D11 | V1rd11 | 2.8 | SM2213 | f | 3 |
| 092308m1_V2MM_125714 | 1.086598 | 3.004939 | TRUE | HP_34655 | TGCTGTTG | CATAGAGC | XM_144191 | Mus musculus similar to Nedd4 binding protein 1 (LOC214795), mR. | | 2.5 | SM2099 | e | 6 |
| 092308m1_V2MM_151143 | 1.086586 | 3.004939 | TRUE | HP_37181 | TGCTGTTG | GTGTGATT | XM_194518 | Mus musculus similar to pol polyprotein [Trichosurus vulpecula] (LOC270309), m | | 2.4 | SM2031 | f | 3 |
| mPool2_V2MM_88780 | 1.086251 | 2.205692 | TRUE | HP_31021 | TGCTGTTG | GACCATAC | NM_175494 | zinc finger protein 367 | Zfp367 | 2.6 | SM2143 | f | 1 |
| mPool6_V2MM_155147 | 1.086239 | 75.83547 | TRUE | HP_37578 | TGCTGTTG | CTGTTAGC | XM_197224 | Mus musculus LOC268804 (LOC268804), mRNA. | | 2.8 | SM2212 | b | 5 |
| 092308m1_V2MM_91516 | 1.086147 | 3.004939 | TRUE | HP_31288 | TGCTGTTG | CAGGACAT | NM_178373 | cell death-inducing DFFA-like effector c | Cidec | 2.6 | SM2109 | d | 1 |
| 092308m1_V2MM_68795 | 1.085744 | 2.205692 | TRUE | HP_29074 | TGCTGTTG | GAGATAC | NM_053142 | protocadherin beta 17 | Pcdhb17 | 2.3 | SM2014 | c | 3 |
| mPool2_V2MM_110894 | 1.085267 | 2.205692 | TRUE | HP_33191 | TGCTGTTG | CAGGGCA | BC048918 | RIKEN cDNA 4931406O17 gene | 4931406O17Rik | 2.7 | SM2155 | e | 3 |
| mPool2_V2MM_208740 | 1.085141 | 3.004939 | TRUE | HP_42652 | TGCTGTTG | GAGTCAA | XM_157959 | Mus musculus LOC213485 (LOC213485), mRNA. | | 2.1 | SM2335 | c | 5 |
| 092308m3_V2MM_145760 | 1.085064 | 59.99801 | TRUE | HP_36647 | TGCTGTTG | CACTGTGC | XM_161356 | similar to 60S ribosomal protein L29 (P23) | | 2.14 | SM2523 | c | 7 |
| mPool7_V2MM_68821 | 1.084678 | 3.004939 | TRUE | HP_29076 | TGCTGTTG | GAGATTTA | NM_026036 | chemokine-like factor super family 6 | Cklfsf6 | 2.8 | SM2238 | e | 12 |
| mPool2_V2MM_106722 | 1.084084 | 2.205692 | TRUE | HP_32782 | TGCTGTTG | CTGGTGAC | NM_024433 | methylthioadenosine phosphorylase | Mtap | 2.6 | SM2149 | c | 5 |
| mPool2_V2MM_9744 | 1.083877 | 3.004939 | TRUE | HP_23319 | TGCTGTTG | CATTAAAC | NM_008219 | hemoglobin Z, beta-like embryonic chain | Hbb-bh1 | 2.11 | SM2378 | b | 5 |
| mPool5_V2MM_177410 | 1.083826 | 75.83547 | TRUE | HP_39789 | TGCTGTTG | GTTTCTTT | XM_489247 | hypothetical gene supported by AK080922 | | NA | | | NA |
| 092308m3_V2MM_87004 | 1.08359 | 75.83547 | TRUE | HP_30846 | TGCTGTTG | CACGTTTA | NM_174997 | cD sequence BC043098 | BC043098 | 2.9 | SM2266 | g | 11 |
| 092308m3_V2MM_87004 | 1.08359 | 75.83547 | TRUE | HP_30846 | TGCTGTTG | CACGTTTA | NM_174997 | cDNA sequence BC043098 | BC043098 | 2.9 | SM2266 | g | 11 |
| mPool4_V2MM_146577 | 1.083291 | 75.83547 | TRUE | HP_36728 | TGCTGTTG | CTGTGACT | XM_162158 | Mus musculus LOC243180 (LOC243180), mRNA. | | 2.16 | SM2644 | f | 6 |
| mPool4_V2MM_73597 | 1.083253 | 75.83547 | TRUE | HP_29542 | TGCTGTTG | GGGATTAT | NM_176935 | RIKEN cDNA F730015K02 gene | F730015K02Rik | 2.16 | SM2615 | c | 12 |
| mPool4_V2MM_108049 | 1.082411 | 75.83547 | TRUE | HP_32912 | TGCTGTTG | CTGTGTAA | AK078040 | cDNA sequence BC052360 | BC052360 | 2.16 | SM2624 | b | 5 |
| mPool7_V2MM_90810 | 1.08233 | 2.205692 | TRUE | HP_31222 | TGCTGTTG | CATAATCT | NM_177864 | RIKEN cDNA A030013N09 gene | A030013N09Rik | 2.9 | SM2261 | a | 1 |
| mPool7_V2MM_185500 | 1.082286 | 3.004939 | TRUE | HP_40597 | TGCTGTTG | CTTCTCAA | XM_289381 | Mus musculus LOC333054 (LOC333054), mRNA. | | 2.5 | SM2095 | c | 3 |
| mPool7_V2MM_185500 | 1.082286 | 3.004939 | TRUE | HP_40597 | TGCTGTTG | CTTCTCAA | XM_289381 | Mus musculus LOC333054 (LOC333054), mR. | | 2.5 | SM2095 | c | 3 |
| 092308m1_V2MM_146752 | 1.082242 | 2.205692 | TRUE | HP_36746 | TGCTGTTG | GCACAATT | XM_162261 | Mus musculus LOC208184 (LOC208184), mR. | | 2.4 | SM2046 | f | 6 |

| | | | | | | | | | | | | | | |
|---|---|---|---|---|---|---|---|---|---|---|---|---|---|---|
| 092308m1_V2MM_146752 | 1.082242 | 2.205692 | TRUE | HP_36746 | TGCTGTTG | GCACAATT | XM_162261 | Mus musculus LOC208184 (LOC208184), mRNA. | | 2.4 | SM2046 | f | | 6 |
| mPool7_V2MM_87298 | 1.0819 | 3.004939 | TRUE | HP_30875 | TGCTGTTG | CTCATACA | NM_175126 | zinc finger, CCHC domain containing 3 | Zcchc3 | 2.9 | SM2257 | h | | 5 |
| mPool7_V2MM_87298 | 1.0819 | 3.004939 | TRUE | HP_30875 | TGCTGTTG | CTCATACA | NM_175126 | zinc finger, CCHC domain containing 3 | Zcchc3 | 2.9 | SM2257 | h | | 5 |
| mPool4_V2MM_106653 | 1.081656 | 75.83547 | TRUE | HP_32775 | TGCTGTTG | GCACTTAT | XM_131400 | mitochondrial carrier triple repeat 1 | Mcart1 | 2.16 | SM2627 | b | | 10 |
| mPool2_V2MM_125934 | 1.081409 | 3.004939 | TRUE | HP_34677 | TGCTGTTG | CTAAGCAT | XM_144292 | gene model 446, (NCBI) | Gm446 | 2.11 | SM2360 | h | | 8 |
| mPool2_V2MM_128875 | 1.080771 | 3.004939 | TRUE | HP_34968 | TGCTGTTG | GCAATAAT | NM_206897 | olfactory receptor 6 | Olfr6 | 2.11 | SM2353 | f | | 9 |
| mPool4_V2MM_176399 | 1.080719 | 75.83547 | TRUE | HP_39688 | TGCTGTTG | GGAAGGA | XM_287660 | Mus musculus LOC330446 (LOC330446), mRNA. | | NA | | | | NA |
| mPool4_V2MM_38510 | 1.080539 | 75.83547 | TRUE | HP_26123 | TGCTGTTG | GCCAACG | NM_031384 | testis expressed gene 11 | Tex11 | 2.16 | SM2604 | f | | 9 |
| 092308m1_V2MM_170515 | 1.080342 | 2.205692 | TRUE | HP_39100 | TGCTGTTG | GCATATTA | AK087847 | RIKEN cDNA E330029E12 gene | E330029E12Rik | 2.5 | SM2068 | f | | 4 |
| 092308m1_V2MM_170515 | 1.080342 | 2.205692 | TRUE | HP_39100 | TGCTGTTG | GCATATTA | AK087847 | RIKEN cD E330029E12 gene | E330029E12Rik | 2.5 | SM2068 | f | | 4 |
| mPool7_V2MM_181518 | 1.079946 | 3.004939 | TRUE | HP_40199 | TGCTGTTG | CTCTAAAT | XM_288689 | Mus musculus LOC332089 (LOC332089), mRNA. | | 2.5 | SM2087 | c | | 9 |
| mPool7_V2MM_181518 | 1.079946 | 3.004939 | TRUE | HP_40199 | TGCTGTTG | CTCTAAAT | XM_288689 | Mus musculus LOC332089 (LOC332089), mR. | | 2.5 | SM2087 | c | | 9 |
| 092308m3_V2MM_208446 | 1.079644 | 61.29828 | TRUE | HP_42623 | TGCTGTTG | GCACGATT | XM_153091 | Mus musculus LOC238058 (LOC238058), mR. | | 2.14 | SM2527 | g | | 7 |
| 092308m1_V2MM_137884 | 1.079517 | 1.959154 | TRUE | HP_35860 | TGCTGTTG | CTCTGGTT | XM_153684 | Mus musculus LOC212115 (LOC212115), mR. | | 2.4 | SM2035 | g | | 2 |
| 092308m3_V2MM_89934 | 1.079505 | 75.83547 | TRUE | HP_31136 | TGCTGTTG | CCTTATAA | NM_177696 | hypothetical protein 4921513O04 | | 2.13 | SM2451 | d | | 7 |
| mPool4_V2MM_51895 | 1.079182 | 60.13437 | TRUE | HP_27428 | TGCTGTTG | CTGCTGCT | NM_020521 | vomeronasal 1 receptor, B5 | V1rb5 | 2.15 | SM2568 | b | | 7 |
| mPool2_V2MM_85234 | 1.079054 | 1.959154 | TRUE | HP_30671 | TGCTGTTG | GATACATC | NM_133740 | heterogeneous nuclear ribonucleoprotein methyltransfe | Hrmt1l3 | 2.6 | SM2136 | d | | 5 |
| mPool4_V2MM_211052 | 1.078541 | 75.83547 | TRUE | HP_42878 | TGCTGTTG | CAGAGTCT | NM_198624 | RIKEN cDNA 4922504M18 gene | 4922504M18Rik | 2.11 | SM2372 | a | | 1 |
| 092308m1_V2MM_131977 | 1.078112 | 3.004939 | TRUE | HP_35273 | TGCTGTTG | CAAAGGA | XM_147934 | ATP-binding cassette, sub-family D (ALD), member 2 | Abcd2 | 2.5 | SM2099 | c | | 12 |
| mPool2_V2MM_96911 | 1.078062 | 3.004939 | TRUE | HP_31815 | TGCTGTTG | GTCTCAAT | XM_111099 | Mus musculus LOC195045 (LOC195045), mRNA. | | 2.6 | SM2113 | e | | 10 |
| mPool2_V2MM_96911 | 1.078062 | 3.004939 | TRUE | HP_31815 | TGCTGTTG | GTCTCAAT | XM_111099 | Mus musculus LOC195045 (LOC195045), mR. | | 2.6 | SM2113 | e | | 10 |
| mPool6_V2MM_67650 | 1.077974 | 75.83547 | TRUE | HP_28963 | TGCTGTTG | CTAATAAT | AK080731 | RIKEN cDNA A930024F17 gene | A930024F17Rik | 2.8 | SM2224 | f | | 6 |
| mPool7_V2MM_200136 | 1.077826 | 2.205692 | TRUE | HP_41817 | TGCTGTTG | CTTGGAAT | XM_287596 | Mus musculus hypothetical gene supported by AK083439 (LOC330337), mRNA. | | 2.9 | SM2291 | c | | 6 |
| 092308m1_V2MM_118377 | 1.077701 | 2.205692 | TRUE | HP_33931 | TGCTGTTG | CCTTATAT | XM_484607 | similar to zinc finger protein 40 | | 2.5 | SM2100 | c | | 4 |
| mPool4_V2MM_42233 | 1.077665 | 75.83547 | TRUE | HP_26484 | TGCTGTTG | CATGATGA | NM_138590 | zinc finger, CCHC domain containing 7 | Zcchc7 | 2.15 | SM2576 | c | | 7 |
| mPool7_V2MM_193635 | 1.077653 | 2.205692 | TRUE | HP_28461 | TGCTGTTG | CAGGACA | NM_001004357 | contactin associated protein-like 2 | Cntnap2 | 2.8 | SM2247 | f | | 5 |
| mPool2_V2MM_116764 | 1.077433 | 3.004939 | TRUE | HP_33771 | TGCTGTTG | GTGCCCTC | XM_139080 | Mus musculus similar to 40S ribosomal protein S17 (LOC219110), mRNA. | | 2.7 | SM2153 | f | | 2 |
| mPool2_V2MM_84902 | 1.076825 | 3.004939 | TRUE | HP_30640 | TGCTGTTG | CCAGCTTC | NM_029792 | beta-1,3-glucuronyltransferase 1 (glucuronosyltransferas | B3gat1 | 2.6 | SM2133 | a | | 5 |
| mPool2_V2MM_211035 | 1.076824 | 3.004939 | TRUE | HP_42876 | TGCTGTTG | CCAGGAG | AK012792 | RIKEN cDNA 1700037C18 gene | 1700037C18Rik | 2.11 | SM2369 | d | | 8 |
| 092308m3_V2MM_146523 | 1.076642 | 61.29828 | TRUE | HP_36723 | TGCTGTTG | CAGCAGG | XM_162120 | Mus musculus LOC231512 (LOC231512), mR. | | 2.14 | SM2512 | h | | 2 |
| mPool2_V2MM_147347 | 1.075924 | 3.004939 | TRUE | HP_36805 | TGCTGTTG | GCTATTAT | XM_162888 | Mus musculus LOC208753 (LOC208753), mRNA. | | 2.1 | SM2334 | f | | 6 |
| mPool6_V2MM_165199 | 1.075698 | 75.83547 | TRUE | HP_38573 | TGCTGTTG | GCAATGAC | XM_285193 | Mus musculus similar to HDCMA18P protein; DKFZP564K112 protein [Homo sapi | | 2.7 | SM2199 | c | | 10 |
| 092308m3_V2MM_136883 | 1.075253 | 75.83547 | TRUE | HP_35760 | TGCTGTTG | GCAACAA | XM_152202 | Mus musculus LOC237477 (LOC237477), mRNA. | | 2.1 | SM2325 | g | | 10 |
| 092308m3_V2MM_136883 | 1.075253 | 75.83547 | TRUE | HP_35760 | TGCTGTTG | GCAACAA | XM_152202 | Mus musculus LOC237477 (LOC237477), mR. | | 2.1 | SM2325 | g | | 10 |
| mPool4_V2MM_203384 | 1.075085 | 61.29828 | TRUE | HP_42133 | TGCTGTTG | CCTCTGAG | BC042576 | expressed sequence AI480556 | AI480556 | 2.16 | SM2607 | b | | 2 |
| mPool4_V2MM_156717 | 1.074767 | 75.83547 | TRUE | HP_37734 | TGCTGTTG | GTGACCTT | XM_204182 | RIKEN cDNA 1520402A15 gene | 1520402A15Rik | NA | | | | NA |
| mPool2_V2MM_89734 | 1.074742 | 3.004939 | TRUE | HP_31116 | TGCTGTTG | CTCTGATT | AK083443 | zinc finger and BTB domain containing 10 | Zbtb10 | 2.6 | SM2145 | g | | 1 |
| 092308m1_V2MM_157751 | 1.074614 | 2.205692 | TRUE | HP_37836 | TGCTGTTG | CACAAGG | XM_205561 | Mus musculus similar to actin alpha 1 skeletal muscle protein (LOC277868), mR. | | 2.6 | SM2117 | b | | 12 |
| 092308m3_V2MM_88011 | 1.074361 | 75.83547 | TRUE | HP_30945 | TGCTGTTG | CAGAGAT | AK030734 | RIKEN cD 5730427M17 gene | 5730427M17Rik | 2.9 | SM2251 | h | | 3 |
| 092308m3_V2MM_88011 | 1.074361 | 75.83547 | TRUE | HP_30945 | TGCTGTTG | CAGAGAT | AK030734 | RIKEN cDNA 5730427M17 gene | 5730427M17Rik | 2.9 | SM2251 | h | | 3 |
| 092308m1_V2MM_166317 | 1.072477 | 2.205692 | TRUE | HP_38684 | TGCTGTTG | GAAAGAA | XM_285469 | Mus musculus LOC333526 (LOC333526), mR. | | 2.6 | SM2122 | a | | 6 |
| 092308m3_V2MM_226144 | 1.072446 | 62.64426 | TRUE | HP_47817 | TGCTGTTG | CACTGTGA | NM_144954 | peptidylprolyl isomerase (cyclophilin)-like 2 | Ppil2 | 2.13 | SM2460 | e | | 1 |
| mPool7_V2MM_100976 | 1.072249 | 2.205692 | TRUE | HP_32215 | TGCTGTTG | CCACATGT | AK021047 | RIKEN cDNA C030005H24 gene | C030005H24Rik | 2.7 | SM2162 | f | | 5 |
| 092308m1_V2MM_166964 | 1.071827 | 2.205692 | TRUE | HP_38748 | TGCTGTTG | CACACCTC | XM_285643 | Mus musculus similar to pol protein [Sus scrofa] (LOC331244), mR. | | 2.6 | SM2120 | e | | 4 |
| mPool4_V2MM_211189 | 1.071332 | 75.83547 | TRUE | HP_42890 | TGCTGTTG | GATGGAT | XM_140094 | Mus musculus similar to protein (peptidyl-prolyl cis/trans isomerase) NIMA-inter | | 2.14 | SM2543 | h | | 12 |
| 092308m1_V2MM_185026 | 1.070939 | 2.205692 | TRUE | HP_40549 | TGCTGTTG | CTCTCCTG | XM_289299 | Mus musculus LOC332946 (LOC332946), mR. | | 2.5 | SM2090 | c | | 2 |
| mPool4_V2MM_127095 | 1.070844 | 73.80895 | TRUE | HP_34792 | TGCTGTTG | CACAGAA | XM_144978 | Mus musculus similar to 60S RIBOSOMAL PROTEIN L17 (L23) (AMINO ACID STAR | | 2.11 | SM2363 | c | | 10 |
| 092308m1_V2MM_175193 | 1.070763 | 3.004939 | TRUE | HP_39567 | TGCTGTTG | CATACAGT | XM_287389 | Rap1, GTPase-activating protein 1 | Rap1ga1 | 2.5 | SM2085 | e | | 8 |
| mPool6_V2MM_47980 | 1.070741 | 69.72441 | TRUE | HP_27044 | TGCTGTTG | GACAGTTT | NM_008858 | protein kinase C, mu | Prkcm | 2.7 | SM2187 | f | | 7 |
| 092308m3_V2MM_90669 | 1.070686 | 75.83547 | TRUE | HP_31208 | TGCTGTTG | CTCCTGAA | NM_177837 | hypothetical protein 6530439I21 | | 2.13 | SM2469 | g | | 9 |
| 092308m1_V2MM_140396 | 1.0698 | 3.004939 | TRUE | HP_36111 | TGCTGTTG | GATAATGC | XM_155935 | Mus musculus LOC239788 (LOC239788), mR. | | 2.4 | SM2032 | b | | 3 |
| 092308m3_V2MM_98720 | 1.069336 | 75.83547 | TRUE | HP_31994 | TGCTGTTG | GAGACAG | XM_112920 | Mus musculus LOC210461 (LOC210461), mR. | | 2.13 | SM2455 | e | | 7 |
| 092308m1_V2MM_122382 | 1.068725 | 3.004939 | TRUE | HP_34325 | TGCTGTTG | CTAAGTCT | XM_142162 | Mus musculus similar to protein tyrosine phosphatase 4a1 [Rattus norvegicus] (L | | 2.6 | SM2111 | c | | 10 |

| | | | | | | | | | | | | | |
|---|---|---|---|---|---|---|---|---|---|---|---|---|---|
| mPool4_V2MM_60184 | 1.068674 | 75.83547 | TRUE | HP_28234 | TGCTGTTG | GGATTAA | NM_177120 | RIKEN cDNA D130060J02 gene | D130060J02Rik | 2.16 | SM2634 | c | 1 |
| mPool4_V2MM_133449 | 1.068415 | 75.83547 | TRUE | HP_35418 | TGCTGTTG | CCTGGAA | XM_149251 | RIKEN cDNA 9430008C03 gene | 9430008C03Rik | 2.14 | SM2550 | f | 2 |
| 092308m1_V2MM_129450 | 1.068051 | 2.205692 | TRUE | HP_35025 | TGCTGTTG | CAAATTAC | XM_146305 | Mus musculus similar to pol protein [Sus scrofa] (LOC215546), mRNA. | | 2.6 | SM2102 | f | 9 |
| 092308m1_V2MM_129450 | 1.068051 | 2.205692 | TRUE | HP_35025 | TGCTGTTG | CAAATTAC | XM_146305 | Mus musculus similar to pol protein [Sus scrofa] (LOC215546), mR. | | 2.6 | SM2102 | f | 9 |
| 092308m1_V2MM_89673 | 1.067954 | 2.205692 | TRUE | HP_31110 | TGCTGTTG | CCTACACT | NM_177645 | RIKEN cD C430010P07 gene | C430010P07Rik | 2.6 | SM2108 | e | 2 |
| mPool5_V2MM_15935 | 1.067465 | 75.83547 | TRUE | HP_23924 | TGCTGTTG | CTGTAAG | NM_153086 | RIKEN cDNA 9430022F06 gene | 9430022F06Rik | NA | | | NA |
| mPool2_V2MM_7485 | 1.067043 | 3.004939 | TRUE | HP_23099 | TGCTGTTG | GCAAAGT | NM_010864 | myosin Va | Myo5a | 2.7 | SM2164 | b | 12 |
| mPool4_V2MM_140479 | 1.066998 | 75.83547 | TRUE | HP_36119 | TGCTGTTG | GCAAGAA | XM_155983 | Mus musculus LOC239813 (LOC239813), mRNA. | | 2.16 | SM2638 | f | 6 |
| mPool5_V2MM_20282 | 1.066837 | 75.83547 | TRUE | HP_24348 | TGCTGTTG | GTGTAGC | NM_019512 | transcription elongation regulator 1 (CA150) | Tcerg1 | NA | | | NA |
| 092308m1_V2MM_137402 | 1.066813 | 2.205692 | TRUE | HP_35812 | TGCTGTTG | CAGTGAA | XM_153111 | Mus musculus LOC238071 (LOC238071), mR. | | 2.5 | SM2056 | a | 2 |
| mPool7_V2MM_98417 | 1.066493 | 3.004939 | TRUE | HP_31964 | TGCTGTTG | GTGACAT | NM_198860 | expressed sequence AI646023 | AI646023 | 2.9 | SM2264 | d | 2 |
| mPool7_V2MM_98417 | 1.066493 | 3.004939 | TRUE | HP_31964 | TGCTGTTG | GTGACAT | NM_198860 | expressed sequence AI646023 | AI646023 | 2.9 | SM2264 | d | 2 |
| mPool2_V2MM_203294 | 1.066437 | 2.205692 | TRUE | HP_42125 | TGCTGTTG | CACTTCGA | XM_138241 | Mus musculus similar to hypothetical protein A030003A19 (LOC217854), mRNA. | | 2.1 | SM2318 | a | 12 |
| 092308m1_V2MM_168602 | 1.06633 | 2.205692 | TRUE | HP_38910 | TGCTGTTG | GGGAACT | XM_286025 | Mus musculus similar to Glyceraldehyde 3-phosphate dehydrogese (GAPDH) (LO | | 2.6 | SM2114 | g | 12 |
| mPool2_V2MM_85330 | 1.066277 | 2.205692 | TRUE | HP_30681 | TGCTGTTG | CTTATTGA | NM_133992 | ubiquitin specific protease 52 | Usp52 | 2.6 | SM2147 | g | 4 |
| mPool7_V2MM_92544 | 1.066099 | 3.004939 | TRUE | HP_31389 | TGCTGTTG | CAGAGTA | NM_178730 | RIKEN cDNA 4732406D01 gene | 4732406D01Rik | 2.9 | SM2259 | c | 10 |
| 092308m3_V2MM_113493 | 1.066081 | 75.83547 | TRUE | HP_33448 | TGCTGTTG | GCTCCATC | XM_136990 | gene model 225, (NCBI) | Gm225 | 2.13 | SM2499 | h | 2 |
| mPool4_V2MM_201436 | 1.065901 | 75.83547 | TRUE | HP_41946 | TGCTGTTG | GCAAGAA | XM_288243 | Mus musculus LOC332514 (LOC332514), mRNA. | | 2.16 | SM2650 | e | 6 |
| 092308m1_V2MM_177744 | 1.065799 | 2.205692 | TRUE | HP_39822 | TGCTGTTG | GCTTCTGC | XM_287939 | Mus musculus hypothetical gene supported by AK040448 (LOC330996), mR. | | 2.5 | SM2067 | g | 5 |
| 092308m1_V2MM_177744 | 1.065799 | 2.205692 | TRUE | HP_39822 | TGCTGTTG | GCTTCTGC | XM_287939 | Mus musculus hypothetical gene supported by AK040448 (LOC330996), mRNA. | | 2.5 | SM2067 | g | 5 |
| 092308m1_V2MM_146947 | 1.065289 | 3.004939 | TRUE | HP_36765 | TGCTGTTG | CCTCCATT | XM_162472 | Mus musculus LOC243393 (LOC243393), mR. | | 2.4 | SM2034 | c | 7 |
| mPool7_V2MM_195615 | 1.06522 | 3.004939 | TRUE | HP_41374 | TGCTGTTG | CATAGGT | XM_112000 | Mus musculus similar to PROBABLE POL POLYPROTEIN (LOC194862), mRNA. | | 2.8 | SM2249 | a | 11 |
| 092308m3_V2MM_116930 | 1.065095 | 75.83547 | TRUE | HP_10912 | TGCTGTTG | CAGTTTGA | XM_139159 | gene model 295, (NCBI) | Gm295 | 2.1 | SM2319 | h | 10 |
| 092308m3_V2MM_116930 | 1.065095 | 75.83547 | TRUE | HP_10912 | TGCTGTTG | CAGTTTGA | XM_139159 | gene model 295, (NCBI) | Gm295 | 2.1 | SM2319 | h | 10 |
| 092308m3_V2MM_234378 | 1.065083 | 75.83547 | TRUE | HP_48625 | TGCTGTTG | CAGTGGT | XM_126867 | sorting nexin 13 | Snx13 | 2.14 | SM2501 | c | 4 |
| 092308m3_V2MM_86838 | 1.064907 | 75.83547 | TRUE | HP_30830 | TGCTGTTG | CTGCCAAC | NM_173784 | RIKEN cD 9630054F20 gene | 9630054F20Rik | 2.13 | SM2482 | h | 8 |
| mPool2_V2MM_2745 | 1.064804 | 2.205692 | TRUE | HP_22635 | TGCTGTTG | CTGGAACT | NM_026401 | mitochondrial ribosomal protein 63 | Mrp63 | 2.11 | SM2376 | a | 11 |
| mPool4_V2MM_242217 | 1.064744 | 75.83547 | TRUE | HP_49396 | TGCTGTTG | GGTCCAGT | XM_146000 | Mus musculus similar to voltage-dependent calcium channel gamma-3 subunit [F | | 2.14 | SM2550 | d | 5 |
| 092308m3_V2MM_197702 | 1.064463 | 75.83547 | TRUE | HP_41576 | TGCTGTTG | GCCAGAT | NM_181750 | R3H domain (binds single-stranded nucleic acids) | R3hdm | 2.13 | SM2471 | f | 3 |
| mPool2_V2MM_360 | 1.064181 | 3.004939 | TRUE | HP_22403 | TGCTGTTG | CACCACCT | NM_008262 | one cut domain, family member 1 | Onecut1 | 2.11 | SM2377 | d | 6 |
| mPool4_V2MM_228959 | 1.064096 | 75.83547 | TRUE | HP_48094 | TGCTGTTG | CTAAGGG | XM_290162 | Mus musculus LOC333780 (LOC333780), mRNA. | | NA | | | NA |
| 092308m1_V2MM_120081 | 1.063959 | 1.959154 | TRUE | HP_34098 | TGCTGTTG | CTGACAG | XM_140755 | gene model 1285, (NCBI) | Gm1285 | 2.5 | SM2098 | g | 9 |
| mPool2_V2MM_111339 | 1.063452 | 3.004939 | TRUE | HP_33235 | TGCTGTTG | CTACCATC | BC048657 | RIKEN cDNA 1700007N14 gene | 1700007N14Rik | 2.7 | SM2162 | f | 12 |
| 092308m3_V2MM_36601 | 1.06314 | 75.83547 | TRUE | HP_25937 | TGCTGTTG | CTGTTGCT | NM_008623 | myelin protein zero | Mpz | 2.12 | SM2435 | g | 7 |
| 092308m3_V2MM_91709 | 1.063074 | 75.83547 | TRUE | HP_31307 | TGCTGTTG | CACACACT | AK044024 | echinoderm microtubule associated protein like 5 | Eml5 | 2.13 | SM2466 | g | 4 |
| 092308m3_V2MM_209367 | 1.063001 | 75.83547 | TRUE | HP_42714 | TGCTGTTG | GCTGAAA | XM_487119 | similar to LRRGT00126 | | 2.14 | SM2511 | e | 5 |
| mPool7_V2MM_90534 | 1.062392 | 2.205692 | TRUE | HP_31195 | TGCTGTTG | CATTTAAA | NM_177812 | hypothetical protein 4932411K12 | | 2.9 | SM2260 | a | 7 |
| mPool7_V2MM_90534 | 1.062392 | 2.205692 | TRUE | HP_31195 | TGCTGTTG | CATTTAAA | NM_177812 | hypothetical protein 4932411K12 | | 2.9 | SM2260 | a | 7 |
| 092308m3_V2MM_42445 | 1.062023 | 75.83547 | TRUE | HP_26505 | TGCTGTTG | CCATGACC | NM_010269 | ganglioside-induced differentiation-associated-protein 2 | Gdap2 | 2.12 | SM2444 | a | 6 |
| mPool7_V2MM_61649 | 1.061566 | 3.004939 | TRUE | HP_28377 | TGCTGTTG | CCTTCACT | AK007918 | immunoglobulin heavy chain 1a (serum IgG2a) | Igh-1a | 2.8 | SM2229 | g | 2 |
| mPool4_V2MM_131402 | 1.061378 | 75.83547 | TRUE | HP_35216 | TGCTGTTG | GCAAGAA | XM_147640 | hypothetical LOC209550 | | 2.11 | SM2365 | b | 3 |
| mPool7_V2MM_208684 | 1.061349 | 2.205692 | TRUE | HP_42646 | TGCTGTTG | GAGACAA | XM_156894 | Mus musculus LOC213222 (LOC213222), mRNA. | | 2.1 | SM2325 | b | 2 |
| mPool7_V2MM_208684 | 1.061349 | 2.205692 | TRUE | HP_42646 | TGCTGTTG | GAGACAA | XM_156894 | Mus musculus LOC213222 (LOC213222), mR. | | 2.1 | SM2325 | b | 2 |
| mPool7_V2MM_199862 | 1.061291 | 2.205692 | TRUE | HP_41790 | TGCTGTTG | CAAGTGA | XM_289016 | Mus musculus LOC332676 (LOC332676), mRNA. | | 2.9 | SM2285 | b | 5 |
| 092308m1_V2MM_68043 | 1.061107 | 2.205692 | TRUE | HP_29001 | TGCTGTTG | CTGCATG | NM_022328 | myeloid/lymphoid or mixed lineage-leukemia translocati | Mllt1 | 2.4 | SM2024 | c | 8 |
| mPool6_V2MM_15144 | 1.061088 | 75.83547 | TRUE | HP_23847 | TGCTGTTG | CTATTCAG | NM_011631 | tumor rejection antigen gp96 | Tra1 | 2.7 | SM2182 | h | 9 |
| mPool4_V2MM_214399 | 1.06069 | 75.83547 | TRUE | HP_43199 | TGCTGTTG | GCACAAG | XM_143688 | Mus musculus similar to ribosomal protein S6 (LOC230046), mRNA. | | 2.16 | SM2648 | b | 2 |
| mPool2_V2MM_115925 | 1.060636 | 2.205692 | TRUE | HP_33688 | TGCTGTTG | CTGATTCA | BC027395 | RIKEN cDNA A530094I17 gene | A530094I17Rik | 2.7 | SM2163 | c | 4 |
| mPool5_V2MM_266 | 1.060593 | 75.83547 | TRUE | HP_22393 | TGCTGTTG | CACACCCT | NM_026418 | regulator of G-protein signalling 10 | Rgs10 | NA | | | NA |
| mPool4_V2MM_132613 | 1.060413 | 75.83547 | TRUE | HP_35336 | TGCTGTTG | CAGACCC | XM_128781 | lysocardiolipin acyltransferase | Lycat | 2.14 | SM2541 | c | 8 |
| 092308m1_V2MM_161203 | 1.06032 | 3.004939 | TRUE | HP_38177 | TGCTGTTG | CACACTAC | XM_283807 | Mus musculus hypothetical gene supported by AK032464 (LOC329551), mR. | | 2.6 | SM2122 | c | 5 |
| mPool6_V2MM_167800 | 1.059933 | 75.83547 | TRUE | HP_38831 | TGCTGTTG | GGAACTT | XM_285839 | Mus musculus similar to heterochromatin protein 1 alpha [Xenopus laevis] (LOC3 | | 2.7 | SM2189 | d | 5 |
| mPool4_V2MM_103876 | 1.059256 | 75.83547 | TRUE | HP_32501 | TGCTGTTG | CTGTGCTA | AK088543 | CWF19-like 1, cell cycle control (S. pombe) | Cwf19l1 | 2.16 | SM2611 | e | 11 |

| | | | | | | | | | | | | | | | |
|---|---|---|---|---|---|---|---|---|---|---|---|---|---|---|---|
| 092308m3_V2MM_103164 | 1.058881 | 61.29828 | TRUE | HP_32430 | TGCTGTTG | CAGGCATT | XM_128857 | RIKEN cD 4631422C13 gene | 4631422C13Rik | | 2.14 | SM2502 | d | | 1 |
| mPool2_V2MM_125463 | 1.058296 | 2.205692 | TRUE | HP_34630 | TGCTGTTG | GTCAGACT | XM_144020 | Mus musculus similar to urea transporter [Rattus norvegicus] (LOC230714), mRN | | | 2.5 | SM2098 | a | | 4 |
| mPool2_V2MM_125463 | 1.058296 | 2.205692 | TRUE | HP_34630 | TGCTGTTG | GTCAGACT | XM_144020 | Mus musculus similar to urea transporter [Rattus norvegicus] (LOC230714), mR. | | | 2.5 | SM2098 | a | | 4 |
| mPool6_V2MM_69804 | 1.058202 | 66.95036 | TRUE | HP_29172 | TGCTGTTG | GTACTTGT | NM_013761 | serine racemase | Srr | | 2.8 | SM2223 | f | | 6 |
| mPool7_V2MM_173410 | 1.058149 | 3.004939 | TRUE | HP_39389 | TGCTGTTG | CGTCCTAC | AK089549 | RIKEN cDNA 4930401B06 gene | 4930401B06Rik | | 2.9 | SM2281 | g | | 2 |
| mPool7_V2MM_147742 | 1.058049 | 3.004939 | TRUE | HP_36845 | TGCTGTTG | CAACCCA | XM_163232 | Mus musculus LOC243940 (LOC243940), mRNA. | | | 2.1 | SM2328 | b | | 5 |
| 092308m1_V2MM_74764 | 1.056543 | 3.004939 | TRUE | HP_29656 | TGCTGTTG | CAGGCTAA | NM_176934 | RIKEN cD D930046H04 gene | D930046H04Rik | | 2.4 | SM2029 | f | | 1 |
| mPool5_V2MM_15968 | 1.056227 | 75.83547 | TRUE | HP_23927 | TGCTGTTG | CTGTCCTT | NM_054055 | solute carrier family 13 (sodium-dependent dicarboxylat | Slc13a3 | NA | | | | NA | |
| 092308m1_V2MM_172502 | 1.056035 | 2.205692 | TRUE | HP_39298 | TGCTGTTG | CCTCACTT | AK035848 | expressed sequence AI449023 | AI449023 | | 2.5 | SM2063 | b | | 5 |
| mPool2_V2MM_20348 | 1.055883 | 3.004939 | TRUE | HP_24354 | TGCTGTTG | GTGTTATT | NM_007625 | chromobox homolog 4 (Drosophila Pc class) | Cbx4 | | 2.7 | SM2164 | e | | 12 |
| 092308m3_V2MM_47484 | 1.055828 | 62.64426 | TRUE | HP_26996 | TGCTGTTG | CTGCTCA | XM_484349 | similar to RIKEN cD 1700001E04 | | | 2.12 | SM2436 | a | | 12 |
| 092308m3_V2MM_57318 | 1.055725 | 75.83547 | TRUE | HP_27957 | TGCTGTTG | CTTGAAAT | NM_172592 | splicing factor, arginine/serine-rich 12 | Sfrs12 | | 2.12 | SM2402 | g | | 12 |
| mPool7_V2MM_174282 | 1.055218 | 3.004939 | TRUE | HP_39476 | TGCTGTTG | CACCAGG | XM_287196 | Mus musculus hypothetical gene supported by AK034924 (LOC329550), mR. | | | 2.5 | SM2084 | c | | 1 |
| mPool7_V2MM_174282 | 1.055218 | 3.004939 | TRUE | HP_39476 | TGCTGTTG | CACCAGG | XM_287196 | Mus musculus hypothetical gene supported by AK034924 (LOC329550), mRNA. | | | 2.5 | SM2084 | c | | 1 |
| 092308m3_V2MM_34276 | 1.055123 | 66.95036 | TRUE | HP_12579 | TGCTGTTG | CCATGAT | NM_139061 | vacuolar protein sorting 54 (yeast) | Vps54 | | 2.12 | SM2419 | h | | 12 |
| 092308m3_V2MM_207845 | 1.054475 | 75.83547 | TRUE | HP_42564 | TGCTGTTG | GACTGATT | XM_153454 | gene model 599, (NCBI) | Gm599 | | 2.14 | SM2509 | e | | 12 |
| mPool4_V2MM_88782 | 1.05437 | 75.83547 | TRUE | HP_31021 | TGCTGTTG | CACCTTAT | NM_175494 | zinc finger protein 367 | Zfp367 | | 2.16 | SM2616 | g | | 11 |
| mPool4_V2MM_31057 | 1.054283 | 61.29828 | TRUE | HP_25396 | TGCTGTTG | CTCTCTGA | NM_028354 | RIKEN cDNA 2810481F14 gene | 2810481F14Rik | | 2.15 | SM2595 | d | | 6 |
| mPool4_V2MM_51575 | 1.053992 | 75.83547 | TRUE | HP_27397 | TGCTGTTG | CTCTCGAT | XM_283179 | RIKEN cDNA 4930420O11 gene | 4930420O11Rik | NA | | | | NA | |
| mPool2_V2MM_6215 | 1.052072 | 3.004939 | TRUE | HP_22975 | TGCTGTTG | CTCTAGAT | NM_007790 | chondroitin sulfate proteoglycan 6 | Cspg6 | | 2.11 | SM2381 | d | | 1 |
| 092308m1_V2MM_136245 | 1.051001 | 3.004939 | TRUE | HP_35696 | TGCTGTTG | CCAAATTT | XM_151569 | Mus musculus LOC241206 (LOC241206), mRNA. | | | 2.5 | SM2053 | e | | 8 |
| 092308m1_V2MM_136245 | 1.051001 | 3.004939 | TRUE | HP_35696 | TGCTGTTG | CCAAATTT | XM_151569 | Mus musculus LOC241206 (LOC241206), mR. | | | 2.5 | SM2053 | e | | 8 |
| mPool2_V2MM_117873 | 1.050928 | 3.004939 | TRUE | HP_33881 | TGCTGTTG | GTGAAATA | XM_139566 | Mus musculus LOC239654 (LOC239654), mRNA. | | | 2.11 | SM2368 | a | | 9 |
| mPool2_V2MM_1862 | 1.050504 | 3.004939 | TRUE | HP_22549 | TGCTGTTG | CGGAGTTT | NM_054073 | testis specific gene A13 | Tsga13 | | 2.11 | SM2376 | d | | 10 |
| mPool2_V2MM_91444 | 1.050326 | 3.004939 | TRUE | HP_25251 | TGCTGTTG | CACCCATT | NM_178266 | membrane-bound transcription factor protease, site 2 | Mbtps2 | | 2.6 | SM2134 | b | | 10 |
| mPool4_V2MM_122378 | 1.050001 | 75.83547 | TRUE | HP_34324 | TGCTGTTG | CCTTGAGA | XM_142158 | Mus musculus similar to Transcription factor BTF3 (RNA polymerase B transcripti | | | 2.14 | SM2535 | a | | 10 |
| mPool6_V2MM_44055 | 1.049997 | 75.83547 | TRUE | HP_26662 | TGCTGTTG | CTGTGCT | NM_138650 | diacylglycerol kinase, gamma | Dgkg | | 2.7 | SM2186 | b | | 10 |
| 092308m3_V2MM_91042 | 1.049878 | 75.83547 | TRUE | HP_31245 | TGCTGTTG | CTCCGCAT | NM_177909 | solute carrier family 9 (sodium/hydrogen exchanger), iso | Slc9a9 | | 2.13 | SM2485 | c | | 3 |
| mPool7_V2MM_175917 | 1.049243 | 3.004939 | TRUE | HP_39639 | TGCTGTTG | CCATTCA | AK029922 | RIKEN cDNA 4930528J11 gene | 4930528J11Rik | | 2.5 | SM2083 | d | | 4 |
| mPool7_V2MM_175917 | 1.049243 | 3.004939 | TRUE | HP_39639 | TGCTGTTG | CCATTCA | AK029922 | RIKEN cD 4930528J11 gene | 4930528J11Rik | | 2.5 | SM2083 | d | | 4 |
| mPool4_V2MM_7997 | 1.049229 | 61.29828 | TRUE | HP_23148 | TGCTGTTG | GCTATAG | NM_175449 | RIKEN cDNA A630077B13 gene | A630077B13Rik | | 2.16 | SM2608 | a | | 10 |
| 092308m1_V2MM_139580 | 1.048557 | 3.004939 | TRUE | HP_36030 | TGCTGTTG | CCAACATA | XM_155213 | Mus musculus LOC210767 (LOC210767), mR. | | | 2.4 | SM2044 | d | | 2 |
| 092308m1_V2MM_139580 | 1.048557 | 3.004939 | TRUE | HP_36030 | TGCTGTTG | CCAACATA | XM_155213 | Mus musculus LOC210767 (LOC210767), mRNA. | | | 2.4 | SM2044 | d | | 2 |
| 092308m3_V2MM_232128 | 1.048307 | 75.83547 | TRUE | HP_48408 | TGCTGTTG | GAAATTTA | BC054732 | gametogenetin binding protein 1 | | | 2.13 | SM2490 | h | | 12 |
| mPool4_V2MM_118081 | 1.047956 | 60.13437 | TRUE | HP_33901 | TGCTGTTG | GCCAATA | XM_139678 | Mus musculus similar to G protein pathway suppressor 2 [Homo sapiens] (LOC23 | | | 2.16 | SM2641 | f | | 2 |
| 092308m1_V2MM_143461 | 1.047696 | 2.205692 | TRUE | HP_36417 | TGCTGTTG | GCAGAAA | XM_158745 | Mus musculus LOC228665 (LOC228665), mR. | | | 2.4 | SM2035 | f | | 12 |
| mPool4_V2MM_61399 | 1.047397 | 75.83547 | TRUE | HP_28353 | TGCTGTTG | GTGTGAA | NM_145978 | PDZ and LIM domain 2 | Pdlim2 | | 2.15 | SM2591 | f | | 11 |
| 092308m1_V2MM_118252 | 1.047317 | 3.004939 | TRUE | HP_33918 | TGCTGTTG | GTGGAAAT | XM_139741 | Fgfr1 oncogene partner | Fgfr1op | | 2.6 | SM2111 | a | | 7 |
| 092308m1_V2MM_118252 | 1.047317 | 3.004939 | TRUE | HP_33918 | TGCTGTTG | GTGGAAAT | XM_139741 | Fgfr1 oncogene partner | Fgfr1op | | 2.6 | SM2111 | a | | 7 |
| mPool6_V2MM_64823 | 1.047046 | 75.83547 | TRUE | HP_28687 | TGCTGTTG | CTTTGGTT | XM_484640 | similar to Ab2-450 | | | 2.7 | SM2176 | c | | 6 |
| mPool7_V2MM_103141 | 1.046707 | 3.004939 | TRUE | HP_32428 | TGCTGTTG | GCAAATAT | AK006077 | RIKEN cD 1700018A14 gene | 1700018A14Rik | | 2.9 | SM2297 | f | | 9 |
| mPool7_V2MM_103141 | 1.046707 | 3.004939 | TRUE | HP_32428 | TGCTGTTG | GCAAATAT | AK006077 | RIKEN cDNA 1700018A14 gene | 1700018A14Rik | | 2.9 | SM2297 | f | | 9 |
| mPool6_V2MM_39468 | 1.045732 | 35.99881 | TRUE | HP_26215 | TGCTGTTG | GGAATCAT | NM_008778 | p21 (CDKN1A)-activated kinase 3 | Pak3 | | 2.7 | SM2187 | h | | 9 |
| 092308m3_V2MM_55209 | 1.045139 | 75.83547 | TRUE | HP_27751 | TGCTGTTG | CCCTGTGT | NM_177121 | RIKEN cD B930095G15 gene | B930095G15Rik | | 2.12 | SM2416 | a | | 11 |
| mPool7_V2MM_197443 | 1.044533 | 3.004939 | TRUE | HP_14003 | TGCTGTTG | CCATTATA | NM_011669 | ubiquitin specific protease 12 | Usp12 | | 2.9 | SM2258 | c | | 9 |
| mPool6_V2MM_167646 | 1.044461 | 61.29828 | TRUE | HP_38815 | TGCTGTTG | GACATCAA | XM_285809 | Mus musculus similar to ORF1 (LOC332897), mRNA. | | | 2.8 | SM2201 | f | | 4 |
| 092308m3_V2MM_32954 | 1.044446 | 75.83547 | TRUE | HP_25582 | TGCTGTTG | CAAGGAA | NM_145519 | FERM, RhoGEF and pleckstrin domain protein 2 | Farp2 | | 2.12 | SM2433 | f | | 6 |
| mPool4_V2MM_136971 | 1.044195 | 75.83547 | TRUE | HP_35769 | TGCTGTTG | GCTTGGCT | XM_152300 | Mus musculus LOC237573 (LOC237573), mRNA. | | | 2.14 | SM2530 | b | | 2 |
| 092308m3_V2MM_142279 | 1.043338 | 75.83547 | TRUE | HP_36299 | TGCTGTTG | GTAAAGT | XM_157529 | Mus musculus LOC240393 (LOC240393), mR. | | | 2.1 | SM2339 | a | | 2 |
| 092308m3_V2MM_142279 | 1.043338 | 75.83547 | TRUE | HP_36299 | TGCTGTTG | GTAAAGT | XM_157529 | Mus musculus LOC240393 (LOC240393), mRNA. | | | 2.1 | SM2339 | a | | 2 |
| mPool5_V2MM_16573 | 1.043251 | 75.83547 | TRUE | HP_23986 | TGCTGTTG | GACACAAA | NM_053265 | RIKEN cDNA 4930433N12 gene | 4930433N12Rik | NA | | | | NA | |
| mPool2_V2MM_108119 | 1.043203 | 3.004939 | TRUE | HP_32919 | TGCTGTTG | CACGTGGA | AK005862 | RIKEN cDNA 1700011F03 gene | 1700011F03Rik | | 2.7 | SM2164 | e | | 1 |
| mPool7_V2MM_74533 | 1.04288 | 3.004939 | TRUE | HP_29633 | TGCTGTTG | CAGACTGT | NM_026041 | RIKEN cDNA 2810430M08 gene | 2810430M08Rik | | 2.8 | SM2244 | f | | 8 |

| | | | | | | | | | | | | | | | |
|---|---|---|---|---|---|---|---|---|---|---|---|---|---|---|---|
| mPool6_V2MM_73490 | 1.042751 | 75.83547 | TRUE | HP_29532 | TGCTGTTG | GGCCAAAT | NM_172809 | RIKEN cDNA E130115J16 gene | E130115J16Rik | | 2.8 | SM2217 | e | | 4 |
| 092308m3_V2MM_140153 | 1.042084 | 61.29828 | TRUE | HP_36087 | TGCTGTTG | GAGTATG( | XM_155758 | Mus musculus LOC239449 (LOC239449), mR. | | | 2.14 | SM2520 | c | | 8 |
| mPool5_V2MM_14799 | 1.042082 | 75.83547 | TRUE | HP_10626 | TGCTGTTG | CGCATTG( | XM_140330 | Mus musculus LOC240277 (LOC240277), mRNA. | | NA | | | | NA | |
| mPool2_V2MM_191993 | 1.042071 | 3.004939 | TRUE | HP_32602 | TGCTGTTG | CAGCAGA/ | NM_207298 | cerebral endothelial cell adhesion molecule 1 | Ceecam1 | | 2.1 | SM2319 | a | | 4 |
| mPool5_V2MM_165457 | 1.041182 | 75.83547 | TRUE | HP_38599 | TGCTGTTG | CTGCCGAT | XM_285253 | similar to ribosomal protein S18 | | NA | | | | NA | |
| 092308m1_V2MM_186362 | 1.04079 | 3.004939 | TRUE | HP_40683 | TGCTGTTG | CTGGCCTT | XM_289526 | Mus musculus LOC333250 (LOC333250), mR. | | | 2.5 | SM2094 | h | | 6 |
| mPool7_V2MM_70807 | 1.040609 | 3.004939 | TRUE | HP_29270 | TGCTGTTG | CAGAGTG( | NM_025466 | gastrokine 1 | Gkn1 | | 2.8 | SM2230 | b | | 8 |
| 092308m1_V2MM_135142 | 1.040175 | 3.004939 | TRUE | HP_35586 | TGCTGTTG | CTGAGTAT | XM_150350 | UDP-glucuronosyltransferase 1 family, member 1 | Ugt1a1 | | 2.4 | SM2049 | f | | 2 |
| 092308m3_V2MM_25922 | 1.040067 | 66.95036 | TRUE | HP_24897 | TGCTGTTG | CCCAGAG( | NM_008507 | linker of T-cell receptor pathways | Lnk | | 2.12 | SM2423 | b | | 10 |
| mPool6_V2MM_65976 | 1.039859 | 75.83547 | TRUE | HP_28800 | TGCTGTTG | GTCTGTCC | NM_033583 | protocadherin gamma subfamily C, 5 | Pcdhgc5 | | 2.8 | SM2224 | c | | 5 |
| mPool2_V2MM_113971 | 1.039781 | 3.004939 | TRUE | HP_33495 | TGCTGTTG | CAGAGTC/ | XM_137221 | RIKEN cDNA 4930503E15 gene | 4930503E15Rik | | 2.6 | SM2148 | c | | 4 |
| mPool7_V2MM_83943 | 1.03967 | 3.004939 | TRUE | HP_30546 | TGCTGTTG | CACAAGA/ | NM_026279 | RIKEN cDNA 2310026E23 gene | 2310026E23Rik | | 2.9 | SM2267 | e | | 11 |
| mPool7_V2MM_64334 | 1.03946 | 3.004939 | TRUE | HP_28639 | TGCTGTTG | CTGCTCCC | XM_139182 | Mus musculus similar to Cell division protein kinase 4 (Cyclin-dependent kinase 4 | | | 2.8 | SM2234 | e | | 7 |
| mPool4_V2MM_149309 | 1.039383 | 75.83547 | TRUE | HP_37001 | TGCTGTTG | CAAACCA( | XM_164753 | Mus musculus LOC244771 (LOC244771), mRNA. | | | 2.16 | SM2637 | b | | 9 |
| mPool4_V2MM_181709 | 1.039156 | 75.83547 | TRUE | HP_40218 | TGCTGTTG | CAATTACT | XM_288722 | Mus musculus LOC332139 (LOC332139), mRNA. | | | 2.16 | SM2649 | d | | 3 |
| 092308m3_V2MM_234790 | 1.038584 | 61.29828 | TRUE | HP_48666 | TGCTGTTG | CAGTGAG( | XM_135684 | RIKEN cD 1700022L20 gene | 1700022L20Rik | | 2.13 | SM2496 | d | | 4 |
| mPool2_V2MM_99576 | 1.038574 | 3.004939 | TRUE | HP_32077 | TGCTGTTG | CCTTTGTA | XM_125901 | DNA segment, Chr 10, ERATO Doi 516, expressed | D10Ertd516e | | 2.6 | SM2133 | c | | 2 |
| mPool2_V2MM_85489 | 1.038327 | 3.004939 | TRUE | HP_30697 | TGCTGTTG | CGTTTATT | NM_146394 | olfactory receptor 1278 | Olfr1278 | | 2.6 | SM2140 | f | | 10 |
| mPool7_V2MM_180102 | 1.038005 | 3.004939 | TRUE | HP_40057 | TGCTGTTG | GAGATAG( | XM_288450 | Mus musculus LOC331788 (LOC331788), mRNA. | | | 2.9 | SM2275 | c | | 6 |
| mPool7_V2MM_62812 | 1.03753 | 3.004939 | TRUE | HP_28490 | TGCTGTTG | CATGGCG( | NM_013845 | receptor tyrosine kinase-like orphan receptor 1 | Ror1 | | 2.8 | SM2228 | b | | 12 |
| mPool5_V2MM_10711 | 1.037528 | 75.83547 | TRUE | NA | NA | NA | NA | NA | NA | NA | | NA | NA | NA | |
| mPool4_V2MM_35547 | 1.037486 | 69.72441 | TRUE | HP_25834 | TGCTGTTG | CTACACTC | NM_027797 | RIKEN cDNA 5530601I19 gene | 5530601I19Rik | | 2.15 | SM2578 | a | | 9 |
| mPool2_V2MM_69797 | 1.037427 | 3.004939 | TRUE | HP_14267 | TGCTGTTG | GTACATAT | NM_146239 | PCTAIRE-motif protein kinase 2 | Pctk2 | | 2.11 | SM2392 | f | | 4 |
| mPool6_V2MM_163146 | 1.037413 | 75.83547 | TRUE | HP_38370 | TGCTGTTG | GTTCTTGA | AK054302 | expressed sequence C87414 | C87414 | | 2.8 | SM2210 | e | | 5 |
| 092308m1_V2MM_74079 | 1.037347 | 2.205692 | TRUE | HP_29589 | TGCTGTTG | CAACGGC/ | NM_011121 | polo-like kise 1 (Drosophila) | Plk1 | | 2.4 | SM2026 | c | | 9 |
| mPool5_V2MM_162011 | 1.037001 | 75.83547 | TRUE | HP_38257 | TGCTGTTG | GTTTGCA( | XM_284146 | Mus musculus similar to ubiquitin-conjugating enzyme MHR6BN [Mus musculus] | | NA | | | | NA | |
| mPool2_V2MM_165679 | 1.036851 | 3.004939 | TRUE | HP_38621 | TGCTGTTG | CAAGAGA( | XM_285302 | Mus musculus similar to 60S RIBOSOMAL PROTEIN L29 (P23) (LOC332391), mRN/ | | | 2.6 | SM2126 | f | | 3 |
| 092308m3_V2MM_42041 | 1.036744 | 75.83547 | TRUE | HP_26465 | TGCTGTTG | CAGTCAGT | X12807 | ubiquitin specific protease 53 | Usp53 | | 2.12 | SM2441 | h | | 7 |
| mPool2_V2MM_208035 | 1.036333 | 3.004939 | TRUE | HP_42583 | TGCTGTTG | GCAAAGAT | XM_154889 | Mus musculus LOC239181 (LOC239181), mRNA. | | | 2.1 | SM2347 | b | | 6 |
| mPool7_V2MM_174581 | 1.03601 | 3.004939 | TRUE | HP_39506 | TGCTGTTG | GTGGGTA/ | XM_287256 | Mus musculus hypothetical gene supported by AK035260 (LOC329691), mR. | | | 2.5 | SM2081 | h | | 2 |
| mPool7_V2MM_174581 | 1.03601 | 3.004939 | TRUE | HP_39506 | TGCTGTTG | GTGGGTA/ | XM_287256 | Mus musculus hypothetical gene supported by AK035260 (LOC329691), mRNA. | | | 2.5 | SM2081 | h | | 2 |
| 092308m3_V2MM_115612 | 1.035959 | 75.83547 | TRUE | HP_33657 | TGCTGTTG | GGATCTT( | XM_138573 | RIKEN cD 3110004L20 gene | 3110004L20Rik | | 2.13 | SM2493 | c | | 7 |
| mPool2_V2MM_101418 | 1.035595 | 3.004939 | TRUE | HP_32258 | TGCTGTTG | CAGTCTTG | AK029536 | tetratricopeptide repeat domain 18 | Ttc18 | | 2.7 | SM2155 | c | | 10 |
| mPool2_V2MM_109358 | 1.035054 | 3.004939 | TRUE | HP_33041 | TGCTGTTG | CCGACAAT | NM_207239 | general transcription factor III C 1 | Gtf3c1 | | 2.7 | SM2159 | g | | 5 |
| mPool4_V2MM_154320 | 1.034916 | 75.83547 | TRUE | HP_37496 | TGCTGTTG | GGCCGTC( | XM_196729 | Mus musculus LOC270718 (LOC270718), mRNA. | | NA | | | | NA | |
| mPool4_V2MM_73116 | 1.034783 | 75.83547 | TRUE | HP_29495 | TGCTGTTG | GCAACAT( | NM_029377 | RIKEN cDNA 1810054G18 gene | 1810054G18Rik | | 2.15 | SM2592 | b | | 6 |
| 092308m1_V2MM_147297 | 1.034673 | 2.205692 | TRUE | HP_36800 | TGCTGTTG | GCAGTTA/ | XM_162853 | Mus musculus LOC243690 (LOC243690), mR. | | | 2.4 | SM2044 | f | | 10 |
| mPool7_V2MM_90318 | 1.034474 | 2.205692 | TRUE | HP_31174 | TGCTGTTG | GTGACTAT | NM_177774 | serine-arginine repressor protein | | | 2.9 | SM2269 | a | | 11 |
| mPool4_V2MM_239676 | 1.034311 | 75.83547 | TRUE | HP_49147 | TGCTGTTG | CATAGTCA | XM_141952 | similar to P38IP protein | | | 2.16 | SM2641 | b | | 12 |
| mPool2_V2MM_211494 | 1.032269 | 3.004939 | TRUE | HP_42919 | TGCTGTTG | GACCTAGT | U46151 | t-complex-associated testis expressed 2 | Tcte2 | | 2.11 | SM2359 | h | | 8 |
| 092308m3_V2MM_51918 | 1.032177 | 75.83547 | TRUE | HP_27430 | TGCTGTTG | CTGGAAGT | AK122577 | rosbin, round spermatid basic protein 1 | Rsbn1 | | 2.12 | SM2443 | c | | 6 |
| mPool5_V2MM_2668 | 1.031032 | 59.99801 | TRUE | HP_22628 | TGCTGTTG | CTGCCATG | NM_023217 | pyroglutamyl-peptidase I | Pgpep1 | NA | | | | NA | |
| mPool7_V2MM_178962 | 1.030997 | 3.004939 | TRUE | HP_39943 | TGCTGTTG | CAGAAAGT | XM_288249 | Mus musculus LOC332523 (LOC332523), mRNA. | | | 2.9 | SM2289 | f | | 2 |
| 092308m3_V2MM_44089 | 1.030989 | 75.83547 | TRUE | HP_26665 | TGCTGTTG | CTGTGTTG | NM_027927 | RIKEN cD 1110020M19 gene | 1110020M19Rik | | 2.12 | SM2413 | a | | 4 |
| 092308m3_V2MM_97149 | 1.03066 | 75.83547 | TRUE | HP_31839 | TGCTGTTG | GTATCATG | XM_111258 | Mus musculus similar to high sulfur protein B2E [Rattus norvegicus] (LOC193248) | | | 2.13 | SM2459 | b | | 12 |
| 092308m3_V2MM_93056 | 1.030498 | 75.83547 | TRUE | HP_31439 | TGCTGTTG | GTCAGTAT | NM_178896 | expressed sequence AI836376 | AI836376 | | 2.9 | SM2254 | e | | 8 |
| 092308m3_V2MM_93056 | 1.030498 | 75.83547 | TRUE | HP_31439 | TGCTGTTG | GTCAGTAT | NM_178896 | expressed sequence AI836376 | AI836376 | | 2.9 | SM2254 | e | | 8 |
| 092308m3_V2MM_203341 | 1.030199 | 75.83547 | TRUE | HP_42129 | TGCTGTTG | CAGATTGT | XM_485514 | similar to U4/U6 small nuclear ribonucleoprotein Prp4 (U4/U6 snRNP 60 kDa pro | | | 2.9 | SM2298 | d | | 6 |
| 092308m3_V2MM_203341 | 1.030199 | 75.83547 | TRUE | HP_42129 | TGCTGTTG | CAGATTGT | XM_485514 | similar to U4/U6 small nuclear ribonucleoprotein Prp4 (U4/U6 snRNP 60 kDa pro | | | 2.9 | SM2298 | d | | 6 |
| 092308m3_V2MM_83844 | 1.029695 | 75.83547 | TRUE | HP_30536 | TGCTGTTG | GATTAAAT | NM_025700 | phosphoglucomutase 1 | Pgm1 | | 2.9 | SM2270 | c | | 9 |
| 092308m3_V2MM_83844 | 1.029695 | 75.83547 | TRUE | HP_30536 | TGCTGTTG | GATTAAAT | NM_025700 | phosphoglucomutase 1 | Pgm1 | | 2.9 | SM2270 | c | | 9 |
| 092308m1_V2MM_155626 | 1.029604 | 3.004939 | TRUE | HP_37626 | TGCTGTTG | GGATTCTT | XM_197656 | Mus musculus LOC271914 (LOC271914), mR. | | | 2.6 | SM2120 | b | | 5 |
| 092308m3_V2MM_97554 | 1.029561 | 75.83547 | TRUE | HP_31879 | TGCTGTTG | CAAACCA/ | XM_290002 | gene model 865, (NCBI) | Gm865 | | 2.9 | SM2269 | b | | 1 |

| | | | | | | | | | | | | | | | |
|---|---|---|---|---|---|---|---|---|---|---|---|---|---|---|---|
| 092308m3_V2MM_97554 | 1.029561 | 75.83547 | TRUE | HP_31879 | TGCTGTTG | CAAACCA/ | XM_290002 | gene model 865, (NCBI) | Gm865 | | 2.9 | SM2269 | b | | 1 |
| mPool7_V2MM_76042 | 1.029073 | 2.205692 | TRUE | HP_29781 | TGCTGTTG | CCTGCATG | NM_173434 | RIKEN cDNA 9930111J21 gene | 9930111J21Rik | | 2.8 | SM2241 | g | | 4 |
| 092308m1_V2MM_181518 | 1.028745 | 2.205692 | TRUE | HP_40199 | TGCTGTTG | CTCTAAAT | XM_288689 | Mus musculus LOC332089 (LOC332089), mRNA. | | | 2.5 | SM2087 | c | | 9 |
| 092308m1_V2MM_181518 | 1.028745 | 2.205692 | TRUE | HP_40199 | TGCTGTTG | CTCTAAAT | XM_288689 | Mus musculus LOC332089 (LOC332089), mR. | | | 2.5 | SM2087 | c | | 9 |
| mPool5_V2MM_201709 | 1.028687 | 75.83547 | TRUE | HP_41973 | TGCTGTTG | CCTTCATG | XM_289635 | Mus musculus LOC333397 (LOC333397), mRNA. | | NA | | | | NA | |
| mPool2_V2MM_131164 | 1.02775 | 3.004939 | TRUE | HP_35192 | TGCTGTTG | GGAGATA. | XM_483957 | RIKEN cDNA 1810038L18 gene | 1810038L18Rik | | 2.11 | SM2367 | h | | 6 |
| mPool2_V2MM_208122 | 1.027538 | 3.004939 | TRUE | HP_42591 | TGCTGTTG | GATCCGA/ | XM_193891 | similar to CTCL tumor antigen se57-1 | | | 2.1 | SM2346 | b | | 6 |
| mPool2_V2MM_22815 | 1.027533 | 3.004939 | TRUE | HP_24595 | TGCTGTTG | CTCCTAAG | NM_144814 | REST corepressor 3 | Rcor3 | | 2.11 | SM2389 | a | | 10 |
| mPool6_V2MM_194376 | 1.027353 | 75.83547 | TRUE | HP_41254 | TGCTGTTG | CACACTCT | NM_011215 | protein tyrosine phosphatase, receptor type, N polypepti | Ptprn2 | | 2.7 | SM2186 | c | | 11 |
| mPool6_V2MM_160997 | 1.027295 | 69.72441 | TRUE | HP_38157 | TGCTGTTG | GGAGTTC/ | AK077027 | RIKEN cDNA 1700123D08 gene | 1700123D08Rik | | 2.7 | SM2191 | a | | 7 |
| mPool6_V2MM_154139 | 1.026699 | 75.83547 | TRUE | HP_37478 | TGCTGTTG | CTGTGATC | XM_196600 | Mus musculus LOC271617 (LOC271617), mRNA. | | | 2.8 | SM2206 | b | | 4 |
| 092308m3_V2MM_52990 | 1.02624 | 75.83547 | TRUE | HP_27536 | TGCTGTTG | GGCAGAA | NM_010266 | guanine deamise | Gda | | 2.12 | SM2403 | c | | 5 |
| mPool2_V2MM_124387 | 1.026054 | 2.205692 | TRUE | HP_34523 | TGCTGTTG | CATAGTAC | XM_489896 | similar to TDPOZ2 | | | 2.1 | SM2349 | b | | 4 |
| mPool4_V2MM_230666 | 1.025793 | 75.83547 | TRUE | HP_48265 | TGCTGTTG | CGACATTC | XM_287672 | Mus musculus hypothetical gene supported by AK039174 (LOC330486), mRNA. | | NA | | | | NA | |
| mPool2_V2MM_2631 | 1.025366 | 3.004939 | TRUE | HP_22624 | TGCTGTTG | CTGATTCA | NM_134025 | peroxisomal biogenesis factor 12 | Pex12 | | 2.11 | SM2376 | h | | 5 |
| mPool6_V2MM_49051 | 1.024933 | 75.83547 | TRUE | HP_27149 | TGCTGTTG | GCTGGTTC | NM_021420 | serine/threonine kinase 4 | Stk4 | | 2.7 | SM2184 | f | | 12 |
| 092308m1_V2MM_177262 | 1.024861 | 3.004939 | TRUE | NA | NA | NA | NA | NA | NA | NA | | NA | NA | NA | |
| mPool4_V2MM_177464 | 1.024538 | 75.83547 | TRUE | HP_39794 | TGCTGTTG | CAGATGG/ | XM_489259 | hypothetical protein 4930565O14 | | NA | | | | NA | |
| 092308m3_V2MM_109489 | 1.02428 | 75.83547 | TRUE | HP_33054 | TGCTGTTG | CATCTATC | BC051545 | RIKEN cD 1190003J15 gene | 1190003J15Rik | | 2.14 | SM2505 | h | | 7 |
| 092308m3_V2MM_170614 | 1.024236 | 75.83547 | TRUE | HP_39110 | TGCTGTTG | CTGTGGTT | NM_177374 | RIKEN cD 6720458F09 gene | 6720458F09Rik | | 2.12 | SM2407 | e | | 1 |
| 092308m1_V2MM_181959 | 1.024213 | 3.004939 | TRUE | HP_40243 | TGCTGTTG | GCACTAAC | XM_288763 | Mus musculus LOC332210 (LOC332210), mRNA. | | | 2.5 | SM2085 | d | | 5 |
| 092308m1_V2MM_181959 | 1.024213 | 3.004939 | TRUE | HP_40243 | TGCTGTTG | GCACTAAC | XM_288763 | Mus musculus LOC332210 (LOC332210), mR. | | | 2.5 | SM2085 | d | | 5 |
| mPool7_V2MM_96983 | 1.023916 | 3.004939 | TRUE | HP_31822 | TGCTGTTG | CATCTAAT | XM_111147 | Mus musculus similar to pol polyprotein [Trichosurus vulpecula] (LOC211398), m | | | 2.9 | SM2254 | b | | 7 |
| mPool2_V2MM_121182 | 1.023698 | 2.205692 | TRUE | HP_34206 | TGCTGTTG | GCCACTA/ | XM_141519 | RIKEN cDNA 0610030I09 gene | 0610030I09Rik | | 2.5 | SM2098 | c | | 5 |
| mPool2_V2MM_121182 | 1.023698 | 2.205692 | TRUE | HP_34206 | TGCTGTTG | GCCACTA/ | XM_141519 | RIKEN cD 0610030I09 gene | 0610030I09Rik | | 2.5 | SM2098 | c | | 5 |
| mPool7_V2MM_98179 | 1.023653 | 3.004939 | TRUE | HP_31941 | TGCTGTTG | CAGCTGGT | XM_112240 | similar to 60S acidic ribosomal protein P1 | | | 2.9 | SM2267 | b | | 1 |
| mPool4_V2MM_192324 | 1.023398 | 75.83547 | TRUE | HP_41114 | TGCTGTTG | CTGAGAAC | NM_201367 | gene model 1012, (NCBI) | Gm1012 | | 2.16 | SM2606 | b | | 2 |
| mPool7_V2MM_183771 | 1.022688 | 3.004939 | TRUE | HP_40424 | TGCTGTTG | CCTATTAG | XM_289077 | Mus musculus LOC333530 (LOC333530), mR. | | | 2.5 | SM2084 | g | | 11 |
| mPool7_V2MM_183771 | 1.022688 | 3.004939 | TRUE | HP_40424 | TGCTGTTG | CCTATTAG | XM_289077 | Mus musculus LOC333530 (LOC333530), mRNA. | | | 2.5 | SM2084 | g | | 11 |
| 092308m3_V2MM_89502 | 1.022388 | 75.83547 | TRUE | HP_31093 | TGCTGTTG | CAAACACT | AK049924 | hypothetical protein C630011I23 | | | 2.13 | SM2462 | e | | 12 |
| mPool7_V2MM_197025 | 1.022161 | 3.004939 | TRUE | HP_41511 | TGCTGTTG | GTCTGGTT | NM_178643 | RIKEN cDNA C230052I12 gene | C230052I12Rik | | 2.9 | SM2255 | h | | 2 |
| mPool2_V2MM_14560 | 1.022023 | 3.004939 | TRUE | HP_23790 | TGCTGTTG | CCTGTAAA | NM_008314 | 5-hydroxytryptamine (serotonin) receptor 5A | Htr5a | | 2.11 | SM2376 | a | | 10 |
| mPool7_V2MM_203535 | 1.021606 | 3.004939 | TRUE | HP_20561 | TGCTGTTG | CAGTATA/ | XM_139351 | similar to cadherin 12, type 2 preproprotein; Br-cadherin; cadherin-12; N-cadheri | | | 2.1 | SM2302 | d | | 7 |
| 092308m1_V2MM_83736 | 1.021131 | 3.004939 | TRUE | HP_30526 | TGCTGTTG | GGGACTA. | NM_024286 | popeye domain containing 3 | Popdc3 | | 2.6 | SM2107 | c | | 5 |
| mPool7_V2MM_101488 | 1.020777 | 2.205692 | TRUE | HP_32265 | TGCTGTTG | GCTTTACC | AK016963 | RIKEN cDNA 4933428D01 gene | 4933428D01Rik | | 2.9 | SM2300 | d | | 9 |
| 092308m3_V2MM_141336 | 1.020763 | 75.83547 | TRUE | HP_36204 | TGCTGTTG | GTGTCAA/ | XM_156590 | Mus musculus LOC239860 (LOC239860), mR. | | | 2.14 | SM2515 | g | | 7 |
| 092308m3_V2MM_106905 | 1.020321 | 75.83547 | TRUE | HP_32799 | TGCTGTTG | CTGGCTGT | XM_131670 | Mus musculus similar to rhomboid-related protein [Homo sapiens] (LOC230727), | | | 2.1 | SM2314 | a | | 2 |
| 092308m3_V2MM_106905 | 1.020321 | 75.83547 | TRUE | HP_32799 | TGCTGTTG | CTGGCTGT | XM_131670 | Mus musculus similar to rhomboid-related protein [Homo sapiens] (LOC230727), | | | 2.1 | SM2314 | a | | 2 |
| 092308m3_V2MM_107803 | 1.020263 | 75.83547 | TRUE | NA | NA | NA | NA | NA | NA | NA | | NA | NA | NA | |
| mPool4_V2MM_122555 | 1.019963 | 75.83547 | TRUE | HP_34342 | TGCTGTTG | GCTATTCT | XM_142262 | patched domain containing 1 | Ptchd1 | | 2.14 | SM2540 | e | | 1 |
| 092308m3_V2MM_84105 | 1.019841 | 75.83547 | TRUE | HP_30562 | TGCTGTTG | CTGTTCTT. | BC072625 | RIKEN cD 1500010J02 gene | 1500010J02Rik | | 2.13 | SM2482 | c | | 1 |
| 092308m3_V2MM_54351 | 1.019756 | 75.83547 | TRUE | HP_27668 | TGCTGTTG | CAGTCTGT | NM_010293 | glucokise activity, related sequence 1 | Gk-rs1 | | 2.12 | SM2413 | g | | 10 |
| 092308m1_V2MM_178244 | 1.019755 | 3.004939 | TRUE | HP_39871 | TGCTGTTG | CAAGCATC | AK032250 | hypothetical protein 6430511F03 | | | 2.5 | SM2077 | c | | 10 |
| mPool4_V2MM_215074 | 1.019408 | 62.64426 | TRUE | HP_43264 | TGCTGTTG | CCTTCGCC | XM_146272 | Mus musculus similar to eukaryotic translation initiation factor 3, subunit 3 gamr | | | 2.16 | SM2647 | a | | 2 |
| mPool7_V2MM_205539 | 1.018911 | 2.205692 | TRUE | HP_42339 | TGCTGTTG | GAGCTTA/ | XM_127914 | Mus musculus RIKEN cDNA 4921505C17 gene (4921505C17Rik), mRNA. | | | 2.9 | SM2297 | h | | 1 |
| mPool4_V2MM_23788 | 1.018786 | 75.83547 | TRUE | HP_24690 | TGCTGTTG | GAAAGAA | NM_145509 | RIKEN cDNA 5430435G22 gene | 5430435G22Rik | NA | | | | NA | |
| 092308m1_V2MM_171419 | 1.018642 | 3.004939 | TRUE | HP_39190 | TGCTGTTG | CTTACATC | AK050207 | expressed sequence AI448102 | AI448102 | | 2.5 | SM2089 | b | | 2 |
| 092308m1_V2MM_171419 | 1.018642 | 3.004939 | TRUE | HP_39190 | TGCTGTTG | CTTACATC | AK050207 | expressed sequence AI448102 | AI448102 | | 2.5 | SM2089 | b | | 2 |
| mPool7_V2MM_175932 | 1.018612 | 3.004939 | TRUE | HP_39641 | TGCTGTTG | CAGGAAT/ | XM_287564 | Mus musculus hypothetical gene supported by AK089795 (LOC330271), mRNA. | | | 2.5 | SM2072 | b | | 10 |
| mPool7_V2MM_175932 | 1.018612 | 3.004939 | TRUE | HP_39641 | TGCTGTTG | CAGGAAT/ | XM_287564 | Mus musculus hypothetical gene supported by AK089795 (LOC330271), mR. | | | 2.5 | SM2072 | b | | 10 |
| 092308m1_V2MM_118975 | 1.018479 | 3.004939 | TRUE | HP_33990 | TGCTGTTG | CAGACATC | XM_140147 | Mus musculus similar to ribosomal protein L3 [Bos taurus] (LOC225107), mR. | | | 2.6 | SM2104 | a | | 8 |
| mPool4_V2MM_240424 | 1.018023 | 75.83547 | TRUE | HP_49220 | TGCTGTTG | GATATTCT | XM_146535 | ATP-binding cassette, sub-family E (OABP), member 1 | Abce1 | | 2.14 | SM2547 | g | | 9 |
| 092308m3_V2MM_29496 | 1.017978 | 75.83547 | TRUE | HP_25245 | TGCTGTTG | CACAGCC/ | NM_009154 | sema domain, seven thrombospondin repeats (type 1 an | Sema5a | | 2.12 | SM2419 | c | | 4 |

| | | | | | | | | | | | | | |
|---|---|---|---|---|---|---|---|---|---|---|---|---|---|
| mPool7_V2MM_175377 | 1.017508 | 3.004939 | TRUE | NA | NA | NA | NA | NA | NA | NA | NA | NA | NA |
| 092308m1_V2MM_117965 | 1.017506 | 3.004939 | TRUE | HP_33890 | TGCTGTTG | CTTAGATG | XM_139620 | Mus musculus similar to protein phosphatase 2 (formerly 2A), regulatory subunit | | 2.6 | SM2104 | e | 1 |
| 092308m1_V2MM_117965 | 1.017506 | 3.004939 | TRUE | HP_33890 | TGCTGTTG | CTTAGATG | XM_139620 | Mus musculus similar to protein phosphatase 2 (formerly 2A), regulatory subunit | | 2.6 | SM2104 | e | 1 |
| mPool2_V2MM_65699 | 1.017324 | 3.004939 | TRUE | HP_28773 | TGCTGTTG | GGGAAGA | XM_489719 | similar to Calcium-dependent phospholipase A2 precursor (Phosphatidylcholine 2 | | 2.7 | SM2166 | f | 12 |
| mPool2_V2MM_129888 | 1.017254 | 3.004939 | TRUE | HP_35068 | TGCTGTTG | GAGCTGA | BC041776 | RIKEN cDNA D630040G17 gene | D630040G17Rik | 2.11 | SM2353 | g | 6 |
| 092308m1_V2MM_132331 | 1.017208 | 3.004939 | TRUE | HP_35308 | TGCTGTTG | CCAGTTCT | XM_358823 | leucine rich repeat containing 15 | Lrrc15 | 2.5 | SM2099 | f | 9 |
| 092308m1_V2MM_132331 | 1.017208 | 3.004939 | TRUE | HP_35308 | TGCTGTTG | CCAGTTCT | XM_358823 | leucine rich repeat containing 15 | Lrrc15 | 2.5 | SM2099 | f | 9 |
| 092308m1_V2MM_177372 | 1.017051 | 2.205692 | TRUE | HP_39785 | TGCTGTTG | GGCTTAA | XM_287859 | EST AA546796 | AA546796 | 2.5 | SM2086 | h | 12 |
| 092308m3_V2MM_87647 | 1.016675 | 75.83547 | TRUE | HP_30909 | TGCTGTTG | CCACTATA | NM_175212 | RIKEN cD 4930438D12 gene | 4930438D12Rik | 2.9 | SM2263 | f | 5 |
| 092308m3_V2MM_87647 | 1.016675 | 75.83547 | TRUE | HP_30909 | TGCTGTTG | CCACTATA | NM_175212 | RIKEN cDNA 4930438D12 gene | 4930438D12Rik | 2.9 | SM2263 | f | 5 |
| 092308m3_V2MM_103151 | 1.016447 | 60.13437 | TRUE | HP_19897 | TGCTGTTG | GGAAGTT | AK129176 | RIKEN cD 4933432P15 gene | 4933432P15Rik | 2.13 | SM2490 | h | 10 |
| mPool7_V2MM_81933 | 1.01615 | 2.205692 | TRUE | HP_30352 | TGCTGTTG | GTGTCAA | NM_011228 | RAB33A, member of RAS oncogene family | Rab33a | 2.8 | SM2248 | d | 2 |
| mPool4_V2MM_198821 | 1.016099 | 75.83547 | TRUE | HP_41687 | TGCTGTTG | CTGTCTCT | XM_158119 | Mus musculus LOC241223 (LOC241223), mRNA. | | 2.16 | SM2642 | b | 6 |
| 092308m3_V2MM_195113 | 1.016007 | 75.83547 | TRUE | HP_41326 | TGCTGTTG | CCCTTTAG | NM_177770 | cD sequence BC043934 | BC043934 | 2.9 | SM2257 | g | 5 |
| 092308m3_V2MM_195113 | 1.016007 | 75.83547 | TRUE | HP_41326 | TGCTGTTG | CCCTTTAG | NM_177770 | cDNA sequence BC043934 | BC043934 | 2.9 | SM2257 | g | 5 |
| mPool4_V2MM_72595 | 1.015902 | 75.83547 | TRUE | HP_29444 | TGCTGTTG | CTTACTGG | NM_146768 | olfactory receptor 1099 | Olfr1099 | 2.15 | SM2596 | d | 5 |
| mPool7_V2MM_182098 | 1.015748 | 2.205692 | TRUE | HP_40257 | TGCTGTTG | CAATATAC | XM_288786 | Mus musculus LOC332247 (LOC332247), mRNA. | | 2.5 | SM2075 | c | 5 |
| 092308m1_V2MM_117945 | 1.01553 | 3.004939 | TRUE | HP_33888 | TGCTGTTG | GCAACTTC | XM_139608 | Mus musculus similar to glyceraldehyde 3-phosphate dehydrogese [Cavia porcell | | 2.6 | SM2101 | g | 5 |
| 092308m1_V2MM_166779 | 1.015057 | 2.205692 | TRUE | HP_38730 | TGCTGTTG | CGAAGTA | XM_285585 | similar to hypothetical protein MGC18736 | | 2.6 | SM2117 | b | 8 |
| mPool2_V2MM_87712 | 1.014996 | 3.004939 | TRUE | HP_30916 | TGCTGTTG | CAGAAGA | NM_175229 | serine/arginine repetitive matrix 2 | Srrm2 | 2.6 | SM2146 | d | 9 |
| mPool4_V2MM_34143 | 1.014927 | 75.83547 | TRUE | HP_25697 | TGCTGTTG | CCAGATTA | NM_133702 | RIKEN cDNA 1500002M01 gene | 1500002M01Rik | 2.15 | SM2599 | f | 6 |
| 092308m3_V2MM_38454 | 1.014729 | 75.83547 | TRUE | HP_26117 | TGCTGTTG | GCATCTAT | NM_030725 | syptotagmin 13 | Syt13 | 2.12 | SM2402 | b | 1 |
| mPool5_V2MM_16666 | 1.014572 | 75.83547 | TRUE | HP_23996 | TGCTGTTG | GACATGA | NM_008356 | interleukin 13 receptor, alpha 2 | Il13ra2 | NA | | | NA |
| 092308m3_V2MM_44042 | 1.014523 | 75.83547 | TRUE | HP_26660 | TGCTGTTG | CTGTGAG | NM_008992 | ATP-binding cassette, sub-family D (ALD), member 4 | Abcd4 | 2.12 | SM2405 | d | 3 |
| 092308m3_V2MM_105139 | 1.014326 | 75.83547 | TRUE | HP_32626 | TGCTGTTG | CAGTCGA | AK016085 | RIKEN cD 4930550G17 gene | 4930550G17Rik | 2.14 | SM2504 | h | 2 |
| mPool6_V2MM_160128 | 1.014306 | 66.95036 | TRUE | HP_38071 | TGCTGTTG | GTTGCTAT | XM_283382 | Mus musculus hypothetical gene supported by AK045597 (LOC328725), mRNA. | | 2.8 | SM2203 | e | 7 |
| mPool2_V2MM_146752 | 1.014109 | 3.004939 | TRUE | HP_36746 | TGCTGTTG | GCACAAT | XM_162261 | Mus musculus LOC208184 (LOC208184), mRNA. | | 2.4 | SM2046 | f | 6 |
| mPool2_V2MM_146752 | 1.014109 | 3.004939 | TRUE | HP_36746 | TGCTGTTG | GCACAAT | XM_162261 | Mus musculus LOC208184 (LOC208184), mR. | | 2.4 | SM2046 | f | 6 |
| 092308m3_V2MM_141285 | 1.013997 | 75.83547 | TRUE | HP_36199 | TGCTGTTG | CCTGCACA | XM_156562 | Mus musculus LOC224376 (LOC224376), mR. | | 2.4 | SM2031 | b | 7 |
| 092308m1_V2MM_139305 | 1.013963 | 3.004939 | TRUE | HP_36002 | TGCTGTTG | CGTATTTG | XM_154906 | Mus musculus LOC210795 (LOC210795), mR. | | 2.4 | SM2034 | b | 10 |
| mPool7_V2MM_94362 | 1.013487 | 3.004939 | TRUE | HP_31565 | TGCTGTTG | CACTTTAG | NM_183138 | cDNA sequence BC037432 | BC037432 | 2.9 | SM2267 | b | 3 |
| mPool2_V2MM_36920 | 1.013437 | 3.004939 | TRUE | HP_25969 | TGCTGTTG | GAACGTT | XM_487594 | similar to hypothetical protein 4930509O22 | | 2.11 | SM2392 | g | 4 |
| mPool6_V2MM_156731 | 1.013234 | 75.83547 | TRUE | HP_42605 | TGCTGTTG | GCCAGCA | AK007436 | a disintegrin-like and metalloprotease (reprolysin type) w | Adamts9 | 2.7 | SM2191 | h | 3 |
| 092308m3_V2MM_83918 | 1.011765 | 75.83547 | TRUE | HP_30544 | TGCTGTTG | CACAGTG | NM_026225 | component of oligomeric golgi complex 6 | Cog6 | 2.13 | SM2469 | c | 1 |
| 092308m3_V2MM_96517 | 1.011348 | 75.83547 | TRUE | HP_31776 | TGCTGTTG | CAGCCAA | XM_110787 | UPF3 regulator of nonsense transcripts homolog B (yeast | Upf3b | 2.13 | SM2459 | h | 5 |
| 092308m1_V2MM_88072 | 1.011282 | 2.205692 | TRUE | HP_30951 | TGCTGTTG | GACAGTT | AK030699 | a disintegrin-like and metalloprotease (reprolysin type) w | Adamts9 | 2.6 | SM2112 | f | 5 |
| 092308m3_V2MM_207279 | 1.010713 | 75.83547 | TRUE | HP_42509 | TGCTGTTG | GGAAACA | XM_157114 | Mus musculus LOC212321 (LOC212321), mRNA. | | 2.1 | SM2345 | d | 5 |
| 092308m3_V2MM_207279 | 1.010713 | 75.83547 | TRUE | HP_42509 | TGCTGTTG | GGAAACA | XM_157114 | Mus musculus LOC212321 (LOC212321), mR. | | 2.1 | SM2345 | d | 5 |
| mPool2_V2MM_12407 | 1.010561 | 3.004939 | TRUE | HP_23580 | TGCTGTTG | CAAAGCA | BC055079 | cytidine monophospho-N-acetylneuraminic acid hydroxy | Cmah | 2.11 | SM2384 | b | 2 |
| 092308m3_V2MM_142710 | 1.010315 | 75.83547 | TRUE | HP_36342 | TGCTGTTG | CTCAGCTT | XM_157854 | Mus musculus LOC240626 (LOC240626), mR. | | 2.14 | SM2514 | e | 6 |
| mPool2_V2MM_95486 | 1.010231 | 2.205692 | TRUE | HP_31676 | TGCTGTTG | GATATAA | NM_198014 | RIKEN cDNA 9630044O09 gene | 9630044O09Rik | 2.6 | SM2143 | d | 7 |
| mPool2_V2MM_214278 | 1.010164 | 2.205692 | TRUE | HP_43187 | TGCTGTTG | CCCTTCAA | XM_149291 | hypothetical gene supported by AK012899 | | 2.11 | SM2364 | b | 7 |
| mPool4_V2MM_241460 | 1.01016 | 69.72441 | TRUE | HP_49321 | TGCTGTTG | GACAGAC | NM_145475 | ceramide kinase | Cerk | 2.14 | SM2538 | a | 11 |
| 092308m3_V2MM_47098 | 1.009889 | 75.83547 | TRUE | HP_26958 | TGCTGTTG | CTATGCA | NM_138671 | cD sequence BC004012 | BC004012 | 2.12 | SM2406 | a | 4 |
| mPool7_V2MM_174426 | 1.009885 | 3.004939 | TRUE | HP_39491 | TGCTGTTG | CCGTGATT | XM_287229 | Mus musculus hypothetical gene supported by AK077932 (LOC329624), mRNA. | | 2.5 | SM2077 | d | 12 |
| mPool7_V2MM_174426 | 1.009885 | 3.004939 | TRUE | HP_39491 | TGCTGTTG | CCGTGATT | XM_287229 | Mus musculus hypothetical gene supported by AK077932 (LOC329624), mR. | | 2.5 | SM2077 | d | 12 |
| 092308m3_V2MM_101337 | 1.009748 | 75.83547 | TRUE | HP_32250 | TGCTGTTG | CACCCACT | AY262012 | RIKEN cD 1810058I14 gene | 1810058I14Rik | 2.13 | SM2499 | e | 11 |
| 092308m3_V2MM_232796 | 1.009462 | 59.99801 | TRUE | HP_48472 | TGCTGTTG | CCGTTCTG | XM_136073 | RIKEN cD 4930534P07 gene | 4930534P07Rik | 2.13 | SM2497 | e | 6 |
| mPool4_V2MM_160731 | 1.009436 | 75.83547 | TRUE | HP_38131 | TGCTGTTG | GTATGGT | XM_283619 | Mus musculus hypothetical gene supported by AK087209 (LOC329109), mRNA. | | NA | | | NA |
| 092308m3_V2MM_112952 | 1.009361 | 75.83547 | TRUE | HP_33394 | TGCTGTTG | CAGATTTG | NM_207228 | testis specific 10 | Tsga10 | 2.1 | SM2313 | e | 6 |
| 092308m3_V2MM_112952 | 1.009361 | 75.83547 | TRUE | HP_33394 | TGCTGTTG | CAGATTTG | NM_207228 | testis specific 10 | Tsga10 | 2.1 | SM2313 | e | 6 |
| mPool4_V2MM_53953 | 1.009128 | 60.13437 | TRUE | HP_27629 | TGCTGTTG | CACTTTCA | NM_146593 | olfactory receptor 1111 | Olfr1111 | 2.15 | SM2600 | h | 6 |
| mPool4_V2MM_52866 | 1.008972 | 70.87591 | TRUE | HP_27523 | TGCTGTTG | GCTGCCCT | NM_177268 | ankyrin repeat domain 16 | Ankrd16 | 2.16 | SM2607 | h | 10 |

| | | | | | | | | | | | | | |
|---|---|---|---|---|---|---|---|---|---|---|---|---|---|
| 092308m3_V2MM_97736 | 1.008525 | 75.83547 | TRUE | NA | NA | NA | NA | NA | NA | NA | NA | NA | NA |
| 092308m3_V2MM_113117 | 1.008227 | 75.83547 | TRUE | HP_33410 | TGCTGTTG | CACCCAAT | XM_136832 | Mus musculus similar to translation elongation factor eEF-1 alpha chain - rat (LO( | | 2.1 | SM2321 | f | 2 |
| 092308m3_V2MM_113117 | 1.008227 | 75.83547 | TRUE | HP_33410 | TGCTGTTG | CACCCAAT | XM_136832 | Mus musculus similar to translation elongation factor eEF-1 alpha chain - rat (LO( | | 2.1 | SM2321 | f | 2 |
| 092308m1_V2MM_78563 | 1.00817 | 2.205692 | TRUE | HP_91860 | TGCTGTTG | GAGAAGG | NM_010461 | homeo box B8 | Hoxb8 | 2.4 | SM2024 | c | 6 |
| 092308m3_V2MM_51341 | 1.008164 | 60.13437 | TRUE | HP_27374 | TGCTGTTG | CTCACTGC | NM_146591 | olfactory receptor 1101 | Olfr1101 | 2.12 | SM2405 | h | 10 |
| mPool4_V2MM_103762 | 1.007856 | 75.83547 | TRUE | HP_32490 | TGCTGTTG | GTCTTAAC | AK008974 | RIKEN cDNA 2210418G03 gene | 2210418G03Rik | 2.16 | SM2611 | d | 11 |
| mPool7_V2MM_174506 | 1.007285 | 2.205692 | TRUE | HP_39499 | TGCTGTTG | CGTGCATT | XM_287244 | Mus musculus hypothetical gene supported by AK051302 (LOC329648), mR. | | 2.5 | SM2071 | g | 2 |
| mPool7_V2MM_174506 | 1.007285 | 2.205692 | TRUE | HP_39499 | TGCTGTTG | CGTGCATT | XM_287244 | Mus musculus hypothetical gene supported by AK051302 (LOC329648), mRNA. | | 2.5 | SM2071 | g | 2 |
| mPool2_V2MM_118793 | 1.007218 | 3.004939 | TRUE | HP_33972 | TGCTGTTG | CCGTAACT | XM_140066 | Mus musculus similar to caspr5 protein isoform 1 (LOC215422), mRNA. | | 2.11 | SM2373 | h | 7 |
| mPool2_V2MM_177570 | 1.00699 | 3.004939 | TRUE | HP_39805 | TGCTGTTG | CGCATTGT | AF064874 | hyperpolarization-activated, cyclic nucleotide-gated K+ 4 | Hcn4 | 2.11 | SM2383 | h | 8 |
| mPool2_V2MM_26867 | 1.006194 | 3.004939 | TRUE | HP_24990 | TGCTGTTG | CTGACATT | NM_008577 | solute carrier family 3 (activators of dibasic and neutral a | Slc3a2 | 2.11 | SM2392 | h | 2 |
| mPool2_V2MM_95815 | 1.005847 | 2.205692 | TRUE | HP_31708 | TGCTGTTG | CACTGAG( | XM_147508 | similar to hypothetical protein MGC39650 | | 2.6 | SM2134 | f | 7 |
| mPool2_V2MM_125961 | 1.005739 | 3.004939 | TRUE | HP_34680 | TGCTGTTG | CTAACTGT | XM_144310 | RIKEN cDNA 6820424L24 gene | 6820424L24Rik | 2.11 | SM2356 | c | 3 |
| mPool7_V2MM_135599 | 1.005654 | 3.004939 | TRUE | HP_35632 | TGCTGTTG | CCTGTTGA | XM_150820 | Mus musculus LOC240684 (LOC240684), mRNA. | | 2.1 | SM2328 | h | 6 |
| 092308m3_V2MM_226962 | 1.005203 | 75.83547 | TRUE | HP_47897 | TGCTGTTG | GATTTCTT | NM_011879 | IK cytokine | Ik | 2.13 | SM2468 | d | 9 |
| mPool4_V2MM_117124 | 1.004907 | 75.83547 | TRUE | HP_33807 | TGCTGTTG | CAAGGACT | XM_139231 | Mus musculus similar to ferritin light chain 1 [Rattus norvegicus] (LOC223167), m | | 2.16 | SM2631 | c | 12 |
| mPool2_V2MM_100976 | 1.00486 | 2.205692 | TRUE | HP_32215 | TGCTGTTG | CCACATGT | AK021047 | RIKEN cDNA C030005H24 gene | C030005H24Rik | 2.7 | SM2162 | f | 5 |
| mPool4_V2MM_126152 | 1.004792 | 75.83547 | TRUE | NA | NA | NA | NA | NA | NA | NA | NA | NA | NA |
| 092308m3_V2MM_46080 | 1.004251 | 70.87591 | TRUE | HP_26859 | TGCTGTTG | CATACGAT | NM_009619 | a disintegrin and metalloprotease domain 3 (cyritestin) | Adam3 | 2.12 | SM2411 | h | 8 |
| mPool4_V2MM_193599 | 1.003953 | 75.83547 | TRUE | HP_41197 | TGCTGTTG | CCACCCTG | XM_196564 | gene model 1857, (NCBI) | Gm1857 | NA | | | NA |
| mPool7_V2MM_97565 | 1.003879 | 2.205692 | TRUE | HP_31880 | TGCTGTTG | CTCCTGTT | XM_111696 | Mus musculus similar to bM332P19.4 (novel 7 transmembrane receptor (rhodop | | 2.9 | SM2254 | a | 12 |
| mPool7_V2MM_83471 | 1.003796 | 1.959154 | TRUE | HP_30501 | TGCTGTTG | CACACATT | NM_016976 | glutamate receptor, metabotropic 1 | Grm1 | 2.9 | SM2267 | a | 2 |
| 092308m3_V2MM_236801 | 1.003457 | 70.87591 | TRUE | HP_48865 | TGCTGTTG | CTGAACCA | XM_156561 | Mus musculus LOC224374 (LOC224374), mR. | | 2.14 | SM2513 | a | 8 |
| 092308m3_V2MM_40451 | 1.003434 | 75.83547 | TRUE | HP_26311 | TGCTGTTG | GTCTCCCT | BC023187 | cell division cycle 27 homolog (S. cerevisiae) | Cdc27 | 2.12 | SM2423 | a | 8 |
| mPool4_V2MM_237471 | 1.003214 | 75.83547 | TRUE | HP_48931 | TGCTGTTG | CGGCCATT | XM_165211 | Mus musculus LOC244976 (LOC244976), mRNA. | | 2.14 | SM2530 | d | 11 |
| mPool7_V2MM_61822 | 1.00277 | 3.004939 | TRUE | HP_28394 | TGCTGTTG | CAATATTT | NM_029466 | ADP-ribosylation factor-like 8 | Arl8 | 2.8 | SM2236 | h | 6 |
| mPool2_V2MM_125790 | 1.002692 | 3.004939 | TRUE | HP_34663 | TGCTGTTG | CATTCAAC | XM_144231 | Mus musculus similar to cadherin EGF LAG seven-pass G-type receptor 2; EGF-lik | | 2.11 | SM2351 | f | 7 |
| mPool6_V2MM_162043 | 1.002089 | 75.83547 | TRUE | HP_38260 | TGCTGTTG | CAGCCCA( | XM_284161 | F-box and leucine-rich repeat protein 18 | Fbxl18 | 2.7 | SM2196 | d | 1 |
| mPool4_V2MM_60911 | 1.001915 | 75.83547 | TRUE | HP_28305 | TGCTGTTG | GTCATTGA | NM_016783 | progesterone receptor membrane component 1 | Pgrmc1 | 2.15 | SM2572 | h | 3 |
| mPool4_V2MM_155733 | 1.001903 | 75.83547 | TRUE | HP_37637 | TGCTGTTG | CTAGCAA( | XM_197725 | Mus musculus LOC271995 (LOC271995), mRNA. | | NA | | | NA |
| mPool4_V2MM_66644 | 1.001814 | 75.83547 | TRUE | HP_28865 | TGCTGTTG | CATAGGA( | NM_177386 | Scm-like with four mbt domains 2 | Sfmbt2 | 2.16 | SM2625 | g | 10 |
| 092308m3_V2MM_149151 | 1.001777 | 75.83547 | TRUE | HP_36986 | TGCTGTTG | CTCTTCAT | XM_164545 | Mus musculus LOC244536 (LOC244536), mRNA. | | 2.1 | SM2341 | g | 1 |
| 092308m3_V2MM_149151 | 1.001777 | 75.83547 | TRUE | HP_36986 | TGCTGTTG | CTCTTCAT | XM_164545 | Mus musculus LOC244536 (LOC244536), mR. | | 2.1 | SM2341 | g | 1 |
| mPool2_V2MM_137293 | 1.001524 | 3.004939 | TRUE | HP_35801 | TGCTGTTG | CCATTATA | XM_152993 | Mus musculus similar to hypothetical protein FLJ33318 [Homo sapiens] (LOC217: | | 2.1 | SM2338 | a | 7 |
| mPool7_V2MM_202634 | 1.001395 | 3.004939 | TRUE | HP_42063 | TGCTGTTG | GAACAGT( | XM_130353 | RIKEN cDNA 4833423E24 gene | 4833423E24Rik | 2.1 | SM2301 | a | 4 |
| mPool2_V2MM_146803 | 1.001393 | 3.004939 | TRUE | HP_36751 | TGCTGTTG | GACGAGT/ | XM_162338 | Mus musculus LOC243316 (LOC243316), mRNA. | | 2.1 | SM2336 | h | 11 |

**Supplementary Table 2: Positive regulators of CSR recovered by the shRNA screen**

| Gene.Name | log2.ratio | q.value... | Hit? | Codex_hai | Codex_97 | Codex_19 | Codex_Ac | Codex_gene.name | Codex_gene.symbol | OB_Releas | OB_Plate | OB_Well_I | OB_Well_( |
|---|---|---|---|---|---|---|---|---|---|---|---|---|---|
| mPool2_V2MM_97177 | -3.01397 | 3.004939 | TRUE | HP_31841 | TGCTGTTG | CACTGGA | XM_1113 | Mus musculus LOC193319 (LOC193319), mRNA. | | 2.6 | SM2143 | h | 2 |
| mPool2_V2MM_134134 | -2.76328 | 3.004939 | TRUE | HP_35486 | TGCTGTTG | GATGTCAT | XM_1496 | RIKEN cDNA 2510016D11 gene | 2510016D11Rik | 2.11 | SM2355 | f | 5 |
| 092308m3_V2MM_85278 | -2.60582 | 94.45693 | TRUE | HP_30676 | TGCTGTTG | CTACTCTA | AK036588 | RIKEN cD 4930504E06 gene | 4930504E06Rik | 2.13 | SM2463 | d | 8 |
| mPool2_V2MM_103707 | -2.59919 | 3.004939 | TRUE | HP_32484 | TGCTGTTG | CCAGCCG | D17583 | proprotein convertase subtilisin/kexin type 5 | Pcsk5 | 2.7 | SM2162 | a | 7 |
| mPool2_V2MM_97000 | -2.49072 | 3.004939 | TRUE | HP_31824 | TGCTGTTG | GCTACACA | XM_1111 | Mus musculus LOC193100 (LOC193100), mRNA. | | 2.6 | SM2144 | c | 3 |
| mPool4_V2MM_7180 | -2.46141 | 61.94057 | TRUE | HP_23069 | TGCTGTTG | GAGATTA | XM_2845 | similar to 14-3-3 protein tau (14-3-3 protein theta) | | 2.15 | SM2568 | h | 5 |
| mPool7_V2MM_97179 | -2.43531 | 3.004939 | TRUE | HP_31842 | TGCTGTTG | GCTGCAC | XM_1113 | Mus musculus LOC193319 (LOC193319), mRNA. | | 2.9 | SM2261 | a | 4 |
| mPool6_V2MM_165503 | -2.40109 | 75.98381 | TRUE | HP_38604 | TGCTGTTG | CTCCTAGT | XM_2852 | Mus musculus similar to p47 protein [Rattus norvegicus] (LOC328893), mRNA. | | 2.8 | SM2211 | d | 12 |
| 092308m3_V2MM_108572 | -2.39461 | 84.31789 | TRUE | HP_32963 | TGCTGTTG | GCAAGAT | NM_0010 | similar to hypothetical protein MGC33926 | | 2.14 | SM2509 | a | 8 |
| mPool2_V2MM_118099 | -2.39241 | 3.004939 | TRUE | HP_33903 | TGCTGTTG | CTGCCATA | XM_1396 | Mus musculus similar to cell division cycle 5-like; CDC5 (cell division cycle 5, S. | | 2.11 | SM2354 | f | 1 |
| 092308m1_V2MM_167492 | -2.36025 | 3.004939 | TRUE | HP_38800 | TGCTGTTG | GGCTATC | XM_2857 | Mus musculus similar to Involved in chitin synthase III activity, also required fo | | 2.6 | SM2118 | f | 1 |
| 092308m1_V2MM_75974 | -2.28985 | 3.004939 | TRUE | HP_29774 | TGCTGTTG | CCTCCAG | NM_1469 | olfactory receptor 1188 | Olfr1188 | 2.3 | SM2013 | f | 11 |
| mPool2_V2MM_120528 | -2.26478 | 3.004939 | TRUE | HP_34142 | TGCTGTTG | GCTTCCAA | XM_1409 | Mus musculus similar to pyrophosphatase (inorganic); inorganic pyrophosphat | | 2.11 | SM2358 | b | 5 |
| mPool2_V2MM_101727 | -2.22288 | 3.004939 | TRUE | HP_32289 | TGCTGTTG | GTGACCC | NM_2076 | fibronectin type III domain containing 3 | Fndc3 | 2.7 | SM2154 | b | 8 |
| mPool7_V2MM_140140 | -2.21765 | 2.577834 | TRUE | HP_36086 | TGCTGTTG | CGCCAAG | XM_1557 | Mus musculus LOC239680 (LOC239680), mRNA. | | 2.1 | SM2331 | c | 2 |
| 092308m3_V2MM_49812 | -2.21119 | 94.45693 | TRUE | HP_7094 | TGCTGTTG | CAAGAAA | NM_0100 | dipeptidylpeptidase 4 | Dpp4 | 2.12 | SM2405 | f | 6 |
| mPool7_V2MM_103335 | -2.18174 | 3.004939 | TRUE | HP_32447 | TGCTGTTG | CGGGTCC | XM_1289 | AFG3(ATPase family gene 3)-like 2 (yeast) | Afg3l2 | 2.7 | SM2163 | e | 12 |
| 092308m1_V2MM_181263 | -2.13466 | 3.004939 | TRUE | HP_40173 | TGCTGTTG | GAAGCAC | XM_2886 | Mus musculus LOC332051 (LOC332051), mR. | | 2.5 | SM2064 | g | 12 |
| 092308m1_V2MM_120524 | -2.13057 | 2.577834 | TRUE | HP_34141 | TGCTGTTG | GACGCTT | XM_1409 | Mus musculus similar to pyrophosphatase (inorganic); inorganic pyrophosphat | | 2.5 | SM2098 | c | 2 |
| mPool7_V2MM_186646 | -2.12306 | 2.577834 | TRUE | HP_40711 | TGCTGTTG | GCCCTGA | XM_2895 | Mus musculus LOC333325 (LOC333325), mRNA. | | 2.5 | SM2096 | a | 1 |
| mPool7_V2MM_186646 | -2.12306 | 2.577834 | TRUE | HP_40711 | TGCTGTTG | GCCCTGA | XM_2895 | Mus musculus LOC333325 (LOC333325), mR. | | 2.5 | SM2096 | a | 1 |
| 092308m1_V2MM_142193 | -2.11035 | 3.004939 | TRUE | HP_36290 | TGCTGTTG | GAGACAA | XM_1573 | Mus musculus LOC240317 (LOC240317), mR. | | 2.5 | SM2055 | d | 1 |
| 092308m1_V2MM_142193 | -2.11035 | 3.004939 | TRUE | HP_36290 | TGCTGTTG | GAGACAA | XM_1573 | Mus musculus LOC240317 (LOC240317), mRNA. | | 2.5 | SM2055 | d | 1 |
| mPool6_V2MM_161518 | -2.10709 | 94.45693 | TRUE | HP_38208 | TGCTGTTG | CCCAAGA | AK076999 | hypothetical protein 4931422A14 | | 2.7 | SM2193 | h | 12 |
| 092308m1_V2MM_172001 | -2.1042 | 2.577834 | TRUE | HP_39248 | TGCTGTTG | GGTGTGC | XM_2867 | Mus musculus hypothetical gene supported by AK084346 (LOC328677), mR. | | 2.5 | SM2066 | d | 3 |
| 092308m3_V2MM_194937 | -2.08482 | 94.45693 | TRUE | HP_41308 | TGCTGTTG | CATGACA | XM_1131 | similar to serine/threonine protein kise 6; aurora-A | | 2.13 | SM2464 | c | 12 |
| 092308m1_V2MM_186646 | -2.08014 | 3.004939 | TRUE | HP_40711 | TGCTGTTG | GCCCTGA | XM_2895 | Mus musculus LOC333325 (LOC333325), mRNA. | | 2.5 | SM2096 | a | 1 |
| 092308m1_V2MM_186646 | -2.08014 | 3.004939 | TRUE | HP_40711 | TGCTGTTG | GCCCTGA | XM_2895 | Mus musculus LOC333325 (LOC333325), mR. | | 2.5 | SM2096 | a | 1 |
| 092308m3_V2MM_22267 | -2.06953 | 94.45693 | TRUE | HP_24542 | TGCTGTTG | CGCTCAG | NM_0085 | lactotransferrin | Ltf | 2.12 | SM2428 | g | 8 |
| mPool2_V2MM_211021 | -2.06336 | 2.577834 | TRUE | HP_42875 | TGCTGTTG | CGCTTCCA | XM_1409 | Mus musculus similar to pyrophosphatase (inorganic); inorganic pyrophosphat | | 2.11 | SM2374 | f | 4 |
| 092308m1_V2MM_65132 | -2.05203 | 3.004939 | TRUE | HP_28717 | TGCTGTTG | GAGTTCA | NM_0262 | dual specificity phosphatase 6 | Dusp6 | 2.3 | SM2016 | c | 10 |
| mPool2_V2MM_140138 | -2.04771 | 2.577834 | TRUE | HP_36085 | TGCTGTTG | GCCAAGA | XM_1557 | Mus musculus LOC239680 (LOC239680), mRNA. | | 2.1 | SM2341 | a | 6 |
| mPool6_V2MM_168564 | -2.03668 | 61.94057 | TRUE | HP_38906 | TGCTGTTG | CAAACAA | XM_2860 | Mus musculus similar to hypothetical protein FLJ37794 [Homo sapiens] (LOC33 | | 2.7 | SM2194 | e | 11 |
| mPool2_V2MM_11840 | -2.01509 | 3.004939 | TRUE | HP_23524 | TGCTGTTG | GGCAAGC | XM_2836 | Cdc42 binding protein kinase alpha | Cdc42bpa | 2.11 | SM2376 | d | 7 |
| 092308m1_V2MM_187359 | -2.01502 | 3.004939 | TRUE | HP_40782 | TGCTGTTG | GACAGCA | XM_2897 | Mus musculus LOC333650 (LOC333650), mR. | | 2.5 | SM2091 | f | 1 |
| mPool2_V2MM_110610 | -2.01085 | 3.004939 | TRUE | HP_33164 | TGCTGTTG | CGTGCGG | XM_1348 | pygopus 1 | Pygo1 | 2.7 | SM2161 | h | 8 |
| mPool2_V2MM_94729 | -1.99219 | 3.004939 | TRUE | HP_31601 | TGCTGTTG | GACCAGG | NM_1832 | hypothetical protein D730018G16 | | 2.6 | SM2133 | b | 11 |
| 092308m1_V2MM_155300 | -1.9908 | 3.004939 | TRUE | HP_37593 | TGCTGTTG | GAACTTG | XM_1973 | Mus musculus LOC271478 (LOC271478), mR. | | 2.6 | SM2124 | f | 8 |
| 092308m3_V2MM_103329 | -1.98731 | 94.45693 | TRUE | HP_32447 | TGCTGTTG | CACTCCAT | XM_1289 | RIKEN cD 2310002L13 gene | 2310002L13Rik | 2.14 | SM2502 | b | 12 |
| mPool6_V2MM_71959 | -1.96709 | 94.45693 | TRUE | HP_29382 | TGCTGTTG | CTCCTCAT | NM_1773 | myosin ID | Myo1d | 2.8 | SM2220 | c | 11 |
| 092308m1_V2MM_128861 | -1.95678 | 3.004939 | TRUE | HP_34967 | TGCTGTTG | CAGTCACA | XM_1459 | Mus musculus LOC244149 (LOC244149), mR. | | NA | | | NA |
| mPool2_V2MM_132536 | -1.94503 | 2.577834 | TRUE | HP_35328 | TGCTGTTG | GTGAGCA | AF032130 | diabetic embryopathy 1 | Dep1 | 2.11 | SM2354 | c | 12 |
| mPool2_V2MM_210646 | -1.94248 | 3.004939 | TRUE | HP_42838 | TGCTGTTG | CACAAGC | XM_1453 | Mus musculus similar to hypothetical protein C230069C04 [Mus musculus] (LO | | 2.1 | SM2350 | h | 7 |
| mPool7_V2MM_70933 | -1.93896 | 3.004939 | TRUE | HP_29283 | TGCTGTTG | CAGGAAA | NM_0074 | astrotactin 1 | Astn1 | 2.8 | SM2235 | e | 12 |
| mPool7_V2MM_141291 | -1.93618 | 3.004939 | TRUE | HP_36200 | TGCTGTTG | CCAAAGG | XM_1565 | Mus musculus LOC239914 (LOC239914), mRNA. | | 2.5 | SM2051 | h | 6 |
| mPool7_V2MM_141291 | -1.93618 | 3.004939 | TRUE | HP_36200 | TGCTGTTG | CCAAAGG | XM_1565 | Mus musculus LOC239914 (LOC239914), mR. | | 2.5 | SM2051 | h | 6 |
| 092308m1_V2MM_136313 | -1.93453 | 2.577834 | TRUE | HP_35703 | TGCTGTTG | GACACAG | XM_1516 | Mus musculus LOC237290 (LOC237290), mRNA. | | 2.4 | SM2047 | b | 9 |
| 092308m1_V2MM_136313 | -1.93453 | 2.577834 | TRUE | HP_35703 | TGCTGTTG | GACACAG | XM_1516 | Mus musculus LOC237290 (LOC237290), mR. | | 2.4 | SM2047 | b | 9 |
| mPool2_V2MM_18822 | -1.92008 | 3.004939 | TRUE | HP_24206 | TGCTGTTG | GCTTGAG | NM_0109 | natural killer tumor recognition sequence | Nktr | 2.11 | SM2379 | a | 1 |

| | | | | | | | | | | | | | |
|---|---|---|---|---|---|---|---|---|---|---|---|---|---|
| 092308m1_V2MM_124412 | -1.91646 | 3.004939 | TRUE | HP_34526 | TGCTGTTG | GCAAGAG | XM_1434 | Mus musculus similar to envelope protein [Ovis aries] (LOC229636), mR. | | 2.5 | SM2099 | h | 3 |
| mPool7_V2MM_91890 | -1.90488 | 3.004939 | TRUE | HP_31325 | TGCTGTTG | GTCCCATA | NM_1786 | leukocyte-associated Ig-like receptor 1 | Lair1 | 2.9 | SM2258 | e | 2 |
| mPool7_V2MM_74594 | -1.89434 | 3.004939 | TRUE | HP_29639 | TGCTGTTG | CAGATCCT | NM_0260 | RIKEN cDNA 2700091N06 gene | 2700091N06Rik | 2.8 | SM2230 | c | 4 |
| mPool2_V2MM_138194 | -1.88746 | 3.004939 | TRUE | HP_35891 | TGCTGTTG | GCAAGGA | XM_1540 | Mus musculus LOC238593 (LOC238593), mRNA. | | 2.1 | SM2335 | e | 12 |
| mPool4_V2MM_122906 | -1.87838 | 94.45693 | TRUE | HP_34376 | TGCTGTTG | GATGTTA | XM_1424 | Mus musculus similar to serine (or cysteine) proteinase inhibitor, clade B, mem | | 2.16 | SM2634 | f | 4 |
| mPool7_V2MM_203354 | -1.878 | 3.004939 | TRUE | HP_42130 | TGCTGTTG | CCAGTGC | XM_1297 | RIKEN cDNA 9230112K08 gene | 9230112K08Rik | 2.1 | SM2302 | c | 3 |
| mPool6_V2MM_76167 | -1.86641 | 61.94057 | TRUE | HP_29793 | TGCTGTTG | CCTTGTCC | NM_0106 | killer cell lectin-like receptor subfamily C, member 2 | Klrc2 | 2.8 | SM2220 | c | 6 |
| mPool4_V2MM_123129 | -1.86044 | 69.72441 | TRUE | HP_34398 | TGCTGTTG | GTCATCTC | XM_2845 | Mus musculus similar to RIKEN cDNA D130040H23; hypothetical protein D130C | | 2.14 | SM2547 | h | 4 |
| 092308m1_V2MM_127303 | -1.84752 | 3.004939 | TRUE | HP_34813 | TGCTGTTG | CTGTGACT | AK032959 | RIKEN cD A430107O13 gene | A430107O13Rik | 2.5 | SM2100 | g | 9 |
| 092308m1_V2MM_127303 | -1.84752 | 3.004939 | TRUE | HP_34813 | TGCTGTTG | CTGTGACT | AK032959 | RIKEN cDNA A430107O13 gene | A430107O13Rik | 2.5 | SM2100 | g | 9 |
| mPool7_V2MM_102694 | -1.83953 | 2.577834 | TRUE | HP_32384 | TGCTGTTG | GTCCCAG | XM_1285 | RIKEN cDNA 1700065O13 gene | 1700065O13Rik | 2.1 | SM2317 | f | 8 |
| mPool6_V2MM_76415 | -1.83549 | 94.45693 | TRUE | HP_29817 | TGCTGTTG | CGGAAAC | NM_0074 | AT motif binding factor 1 | Atbf1 | 2.8 | SM2214 | g | 9 |
| 092308m1_V2MM_148226 | -1.83106 | 3.004939 | TRUE | HP_36893 | TGCTGTTG | GATAGCA | XM_16361 | Mus musculus LOC210679 (LOC210679), mRNA. | | 2.4 | SM2037 | h | 5 |
| 092308m1_V2MM_148226 | -1.83106 | 3.004939 | TRUE | HP_36893 | TGCTGTTG | GATAGCA | XM_16361 | Mus musculus LOC210679 (LOC210679), mR. | | 2.4 | SM2037 | h | 5 |
| mPool2_V2MM_104143 | -1.82611 | 3.004939 | TRUE | HP_32527 | TGCTGTTG | GACACAG | AK090278 | myelin protein zero-like 1 | Mpzl1 | 2.7 | SM2155 | d | 3 |
| mPool7_V2MM_114876 | -1.80953 | 3.004939 | TRUE | HP_33584 | TGCTGTTG | GAGAGGC | XM_1379 | inner mitochondrial membrane peptidase 2-like (S. c | Immp2l | 2.1 | SM2313 | c | 6 |
| mPool7_V2MM_95836 | -1.8095 | 3.004939 | TRUE | HP_31710 | TGCTGTTG | CAAGGAA | BC059847 | Rap guanine nucleotide exchange factor (GEF) 6 | Rapgef6 | 2.9 | SM2251 | e | 2 |
| mPool2_V2MM_103974 | -1.80287 | 3.004939 | TRUE | HP_32511 | TGCTGTTG | CTACTCAG | AK011565 | RIKEN cDNA 2610027F03 gene | 2610027F03Rik | 2.6 | SM2148 | d | 2 |
| 092308m1_V2MM_68829 | -1.80261 | 2.577834 | TRUE | HP_29077 | TGCTGTTG | GAGCCAA | NM_0287 | coagulation factor XIII, A1 subunit | F13a1 | 2.3 | SM2010 | g | 6 |
| mPool2_V2MM_136313 | -1.80045 | 3.004939 | TRUE | HP_35703 | TGCTGTTG | GACACAG | XM_1516 | Mus musculus LOC237290 (LOC237290), mR. | | 2.4 | SM2047 | b | 9 |
| mPool2_V2MM_136313 | -1.80045 | 3.004939 | TRUE | HP_35703 | TGCTGTTG | GACACAG | XM_1516 | Mus musculus LOC237290 (LOC237290), mRNA. | | 2.4 | SM2047 | b | 9 |
| mPool2_V2MM_104189 | -1.80013 | 3.004939 | TRUE | HP_32532 | TGCTGTTG | GCAAGAA | XM_1295 | activating transcription factor 6 | Atf6 | 2.9 | SM2299 | e | 2 |
| mPool4_V2MM_207646 | -1.79911 | 61.94057 | TRUE | HP_42544 | TGCTGTTG | CCTCCTCC | XM_1550 | Mus musculus LOC239237 (LOC239237), mRNA. | | 2.16 | SM2637 | e | 7 |
| mPool4_V2MM_28071 | -1.79667 | 94.45693 | TRUE | HP_25106 | TGCTGTTG | GCAATAG | NM_1455 | cytochrome P450, family 2, subfamily j, polypeptide | Cyp2j13 | 2.15 | SM2580 | c | 6 |
| mPool2_V2MM_18443 | -1.79286 | 3.004939 | TRUE | HP_24169 | TGCTGTTG | GCGATAC | NM_0081 | nuclear receptor subfamily 3, group C, member 1 | Nr3c1 | 2.7 | SM2165 | e | 8 |
| mPool7_V2MM_135220 | -1.79265 | 3.004939 | TRUE | HP_35594 | TGCTGTTG | GAAGGCA | AK005539 | RIKEN cDNA 1600025M17 gene | 1600025M17Rik | 2.4 | SM2046 | c | 6 |
| mPool7_V2MM_135220 | -1.79265 | 3.004939 | TRUE | HP_35594 | TGCTGTTG | GAAGGCA | AK005539 | RIKEN cD 1600025M17 gene | 1600025M17Rik | 2.4 | SM2046 | c | 6 |
| mPool2_V2MM_103335 | -1.79255 | 3.004939 | TRUE | HP_32447 | TGCTGTTG | CGGGTCC | XM_1289 | AFG3(ATPase family gene 3)-like 2 (yeast) | Afg3l2 | 2.7 | SM2163 | e | 12 |
| mPool4_V2MM_183220 | -1.78564 | 61.94057 | TRUE | HP_40369 | TGCTGTTG | GACATTTA | XM_2889 | Mus musculus LOC332636 (LOC332636), mRNA. | | NA | | | NA |
| mPool7_V2MM_184317 | -1.77685 | 3.004939 | TRUE | HP_40478 | TGCTGTTG | GCTAGGA | XM_2891 | Mus musculus LOC332792 (LOC332792), mRNA. | | 2.9 | SM2283 | e | 7 |
| 092308m3_V2MM_107135 | -1.77631 | 94.45693 | TRUE | HP_32822 | TGCTGTTG | CAGATCG | BC060239 | RIKEN cD 2610109H07 gene | 2610109H07Rik | 2.1 | SM2316 | e | 12 |
| 092308m3_V2MM_107135 | -1.77631 | 94.45693 | TRUE | HP_32822 | TGCTGTTG | CAGATCG | BC060239 | RIKEN cDNA 2610109H07 gene | 2610109H07Rik | 2.1 | SM2316 | e | 12 |
| mPool2_V2MM_105604 | -1.76756 | 3.004939 | TRUE | HP_32671 | TGCTGTTG | GCCTTAA | XM_13064 | RIKEN cDNA 5430405G05 gene | 5430405G05Rik | 2.7 | SM2159 | d | 4 |
| 092308m3_V2MM_60570 | -1.76738 | 91.10404 | TRUE | HP_28272 | TGCTGTTG | GGTCAGC | NM_1771 | RIKEN cD A730037L19 gene | A730037L19Rik | 2.12 | SM2445 | g | 10 |
| mPool2_V2MM_104028 | -1.76222 | 3.004939 | TRUE | HP_32516 | TGCTGTTG | CTGCACC | AK007727 | protein phosphatase 1, regulatory (inhibitor) subuni | Ppp1r12b | 2.6 | SM2150 | d | 8 |
| mPool4_V2MM_133732 | -1.76035 | 94.45693 | TRUE | HP_35446 | TGCTGTTG | CTCCATAG | XM_1494 | RIKEN cDNA 1110019J04 gene | 1110019J04Rik | 2.14 | SM2543 | h | 11 |
| mPool2_V2MM_6674 | -1.75621 | 3.004939 | TRUE | HP_23020 | TGCTGTTG | CTTCTGAG | NM_0234 | fumarylacetoacetate hydrolase domain containing 1 | Fahd1 | 2.11 | SM2374 | e | 11 |
| mPool2_V2MM_209710 | -1.7515 | 3.004939 | TRUE | HP_42747 | TGCTGTTG | CCAGCGC | XM_1649 | Mus musculus LOC244840 (LOC244840), mRNA. | | 2.1 | SM2338 | g | 3 |
| 092308m3_V2MM_26723 | -1.74485 | 94.45693 | TRUE | HP_24975 | TGCTGTTG | CTCCTACA | NM_1462 | olfactory receptor 125 | Olfr125 | 2.12 | SM2423 | b | 1 |
| 092308m1_V2MM_169934 | -1.74329 | 3.004939 | TRUE | HP_39042 | TGCTGTTG | GAATTGG | XM_4885 | hypothetical protein A730013G04 | | 2.6 | SM2116 | d | 1 |
| mPool4_V2MM_39314 | -1.74239 | 61.94057 | TRUE | HP_26200 | TGCTGTTG | GCTTCATA | NM_0197 | pseudouridine synthase 1 | Pus1 | 2.15 | SM2571 | a | 2 |
| 092308m1_V2MM_179056 | -1.7409 | 2.577834 | TRUE | HP_39953 | TGCTGTTG | GAGACAC | XM_2882 | Mus musculus LOC332540 (LOC332540), mR. | | 2.5 | SM2062 | c | 8 |
| mPool2_V2MM_83999 | -1.73108 | 3.004939 | TRUE | HP_30551 | TGCTGTTG | GTGCAGG | NM_0265 | reticulocalbin 3, EF-hand calcium binding domain | Rcn3 | 2.6 | SM2146 | g | 5 |
| mPool2_V2MM_131830 | -1.72847 | 2.577834 | TRUE | HP_35259 | TGCTGTTG | CTTCCCTT | XM_1478 | RIKEN cDNA 6330531I01 gene | 6330531I01Rik | 2.11 | SM2352 | b | 6 |
| mPool4_V2MM_134335 | -1.72829 | 81.56557 | TRUE | HP_35506 | TGCTGTTG | GCAATGT | XM_1497 | RIKEN cDNA 1700013D24 gene | 1700013D24Rik | 2.16 | SM2636 | b | 1 |
| 092308m3_V2MM_94017 | -1.72819 | 61.94057 | TRUE | HP_31531 | TGCTGTTG | GTCCATCT | NM_1830 | zinc finger protein 184 (Kruppel-like) | Zfp184 | 2.13 | SM2467 | g | 11 |
| mPool2_V2MM_214630 | -1.7239 | 3.004939 | TRUE | HP_43221 | TGCTGTTG | GGAGCAC | XM_1423 | Mus musculus similar to transitin - chicken (LOC235747), mRNA. | | 2.11 | SM2370 | c | 5 |
| mPool2_V2MM_82543 | -1.7208 | 3.004939 | TRUE | HP_30411 | TGCTGTTG | CTACGCA | NM_0097 | arylsulfatase B | Arsb | 2.6 | SM2135 | a | 10 |
| mPool7_V2MM_77611 | -1.71416 | 3.004939 | TRUE | HP_29932 | TGCTGTTG | CTGTGTG | NM_0138 | RAB GTPase activating protein 1-like | Rabgap1l | 2.8 | SM2244 | d | 12 |
| 092308m1_V2MM_128847 | -1.71414 | 2.577834 | TRUE | HP_34966 | TGCTGTTG | GGCTTCAT | XM_1459 | gene model 493, (NCBI) | Gm493 | 2.6 | SM2114 | a | 2 |
| 092308m1_V2MM_128453 | -1.70527 | 3.004939 | TRUE | HP_34926 | TGCTGTTG | GCATCAG | XM_1456 | similar to glyceraldehyde-3-phosphate dehydrogese | | 2.6 | SM2101 | b | 2 |

| | | | | | | | | | | | | | |
|---|---|---|---|---|---|---|---|---|---|---|---|---|---|
| 092308m1_V2MM_128453 | -1.70527 | 3.004939 | TRUE | HP_34926 | TGCTGTTG | GCATCAG/ | XM_14568 | similar to glyceraldehyde-3-phosphate dehydrogenase | | 2.6 | SM2101 | b | 2 |
| mPool2_V2MM_162734 | -1.70277 | 3.004939 | TRUE | HP_38329 | TGCTGTTG | CGGAGAA | NM_0010( | RIKEN cDNA C330011M18 gene | C330011M18Rik | 2.6 | SM2132 | a | 6 |
| mPool7_V2MM_205110 | -1.70119 | 3.004939 | TRUE | HP_42298 | TGCTGTTG | CAGCTTCC | XM_12924 | RIKEN cDNA E430027O22 gene | E430027O22Rik | 2.1 | SM2306 | f | 12 |
| mPool7_V2MM_148841 | -1.69981 | 2.577834 | TRUE | HP_36955 | TGCTGTTG | CAAGTCAT | XM_16419 | Mus musculus LOC214871 (LOC214871), mRNA. | | 2.1 | SM2331 | e | 9 |
| mPool2_V2MM_123260 | -1.69908 | 3.004939 | TRUE | HP_34411 | TGCTGTTG | GAGCACA( | XM_14284 | Mus musculus similar to glyceraldehyde-3-phosphate dehydrogenase [Mus mu | | 2.11 | SM2362 | h | 5 |
| mPool2_V2MM_104196 | -1.6928 | 3.004939 | TRUE | HP_32533 | TGCTGTTG | CTCTGCTC | XM_12959 | SLAM family member 8 | Slamf8 | 2.7 | SM2161 | c | 3 |
| mPool2_V2MM_157493 | -1.68681 | 3.004939 | TRUE | HP_37811 | TGCTGTTG | CCAAGGG. | XM_20529 | Mus musculus similar to Putative nuclear protein family member, nematode sp | | 2.6 | SM2129 | h | 10 |
| mPool6_V2MM_74478 | -1.68356 | 94.45693 | TRUE | HP_29628 | TGCTGTTG | CAGAAATT | NM_0335( | Mpv17 transgene, kidney disease mutant-like | Mpv17l | 2.7 | SM2173 | c | 2 |
| mPool6_V2MM_223220 | -1.68255 | 61.94057 | TRUE | HP_47539 | TGCTGTTG | CAGAAGT( | XM_2053( | RIKEN cDNA 6330583I20 gene | 6330583I20Rik | 2.8 | SM2204 | g | 11 |
| mPool2_V2MM_137651 | -1.67784 | 3.004939 | TRUE | HP_35837 | TGCTGTTG | GACCCAG( | XM_15343 | Mus musculus LOC238181 (LOC238181), mRNA. | | 2.1 | SM2341 | g | 6 |
| mPool7_V2MM_208359 | -1.67542 | 3.004939 | TRUE | HP_42615 | TGCTGTTG | CAGCAGA/ | XM_19387 | gene model 630, (NCBI) | Gm630 | 2.1 | SM2325 | a | 4 |
| 092308m3_V2MM_112509 | -1.67193 | 61.94057 | TRUE | HP_33351 | TGCTGTTG | CATTACAT | XM_13653 | Mus musculus similar to ribosomal protein S11 [Rattus norvegicus] (LOC24102! | | 2.13 | SM2495 | h | 2 |
| mPool6_V2MM_79246 | -1.67003 | 61.94057 | TRUE | HP_30091 | TGCTGTTG | GATGTAT( | NM_03019 | RIKEN cDNA 2610029D06 gene | 2610029D06Rik | 2.8 | SM2222 | d | 1 |
| 092308m1_V2MM_90225 | -1.66895 | 3.004939 | TRUE | HP_31164 | TGCTGTTG | CGAAAGC( | NM_17775 | hypothetical protein C130098D09 | | 2.6 | SM2107 | e | 6 |
| 092308m1_V2MM_90225 | -1.66895 | 3.004939 | TRUE | HP_31164 | TGCTGTTG | CGAAAGC( | NM_17775 | hypothetical protein C130098D09 | | 2.6 | SM2107 | e | 6 |
| 092308m1_V2MM_127671 | -1.66774 | 3.004939 | TRUE | HP_34849 | TGCTGTTG | CTCAGCAT | BC056221 | zinc finger protein 583 | Zfp583 | 2.5 | SM2097 | c | 5 |
| mPool2_V2MM_123359 | -1.66748 | 3.004939 | TRUE | HP_34421 | TGCTGTTG | GACCACA/ | XM_14291 | Mus musculus similar to ubiquitin-protein ligase (EC 6.3.2.19) - mouse (LOC207 | | 2.5 | SM2098 | f | 6 |
| mPool2_V2MM_123359 | -1.66748 | 3.004939 | TRUE | HP_34421 | TGCTGTTG | GACCACA/ | XM_14291 | Mus musculus similar to ubiquitin-protein ligase (EC 6.3.2.19) - mouse (LOC207 | | 2.5 | SM2098 | f | 6 |
| 092308m3_V2MM_116736 | -1.6642 | 66.95036 | TRUE | HP_33768 | TGCTGTTG | CTATCCAT | XM_48734 | similar to MGC64482 protein | | 2.13 | SM2492 | b | 7 |
| mPool4_V2MM_20522 | -1.65974 | 94.45693 | TRUE | HP_24371 | TGCTGTTG | GTTTGATC | XM_14042 | similar to Profilin I | | 2.16 | SM2636 | b | 11 |
| mPool6_V2MM_79410 | -1.65445 | 94.45693 | TRUE | HP_30107 | TGCTGTTG | GCAATATT | NM_01059 | potassium voltage-gated channel, shaker-related sul | Kcnab3 | 2.8 | SM2216 | f | 3 |
| mPool2_V2MM_6980 | -1.64164 | 3.004939 | TRUE | HP_23050 | TGCTGTTG | GACCGCA/ | NM_01949 | MAD2 (mitotic arrest deficient, homolog)-like 1 (yea | Mad2l1 | 2.11 | SM2375 | a | 9 |
| 092308m1_V2MM_183272 | -1.63987 | 3.004939 | TRUE | HP_40374 | TGCTGTTG | CAAAGGG. | XM_28899 | Mus musculus LOC332654 (LOC332654), mRNA. | | 2.5 | SM2092 | d | 8 |
| 092308m1_V2MM_183272 | -1.63987 | 3.004939 | TRUE | HP_40374 | TGCTGTTG | CAAAGGG. | XM_28899 | Mus musculus LOC332654 (LOC332654), mR. | | 2.5 | SM2092 | d | 8 |
| mPool2_V2MM_128453 | -1.63312 | 3.004939 | TRUE | HP_34926 | TGCTGTTG | GCATCAG/ | XM_14568 | similar to glyceraldehyde-3-phosphate dehydrogenase | | 2.6 | SM2101 | b | 2 |
| mPool2_V2MM_128453 | -1.63312 | 3.004939 | TRUE | HP_34926 | TGCTGTTG | GCATCAG/ | XM_14568 | similar to glyceraldehyde-3-phosphate dehydrogese | | 2.6 | SM2101 | b | 2 |
| mPool2_V2MM_119061 | -1.63155 | 3.004939 | TRUE | HP_33998 | TGCTGTTG | CCGTCCAT | XM_14019 | RIKEN cDNA 9430020K01 gene | 9430020K01Rik | 2.1 | SM2339 | g | 10 |
| 092308m1_V2MM_173738 | -1.62256 | 3.004939 | TRUE | HP_39422 | TGCTGTTG | CAACCCA/ | XM_28708 | Mus musculus LOC329351 (LOC329351), mR. | | 2.5 | SM2089 | g | 9 |
| mPool4_V2MM_162858 | -1.6199 | 61.94057 | TRUE | HP_38341 | TGCTGTTG | GACTCAG/ | XM_28452 | Mus musculus hypothetical gene supported by AK034278 (LOC330994), mRNA. | | NA | | | NA |
| mPool7_V2MM_198838 | -1.61612 | 3.004939 | TRUE | HP_41689 | TGCTGTTG | CACACCTT | XM_48896 | hypothetical gene supported by AK076684 | | 2.9 | SM2293 | h | 1 |
| 092308m3_V2MM_90076 | -1.61511 | 84.31789 | TRUE | HP_31150 | TGCTGTTG | GTCAGAA( | NM_17772 | hypothetical protein 9430004M15 | | 2.13 | SM2457 | d | 8 |
| mPool6_V2MM_222498 | -1.60069 | 94.45693 | TRUE | HP_47475 | TGCTGTTG | GAGAGAC | AK129480 | RIKEN cDNA 5330421F07 gene | 5330421F07Rik | 2.7 | SM2189 | g | 5 |
| 092308m1_V2MM_179796 | -1.59801 | 3.004939 | TRUE | HP_40027 | TGCTGTTG | GGCTTGG/ | XM_28839 | Mus musculus LOC331719 (LOC331719), mR. | | 2.5 | SM2080 | e | 2 |
| mPool2_V2MM_13615 | -1.59662 | 3.004939 | TRUE | HP_23698 | TGCTGTTG | CCAAGGA( | NM_03119 | renin 1 structural | Ren1 | 2.11 | SM2380 | b | 2 |
| mPool2_V2MM_120401 | -1.59644 | 3.004939 | TRUE | HP_34129 | TGCTGTTG | GAAAGAC. | XM_1409( | Mus musculus similar to hypothetical protein DKFZp761J139 [Homo sapiens] (L | | 2.11 | SM2355 | d | 4 |
| 092308m3_V2MM_22100 | -1.59545 | 75.98381 | TRUE | HP_24526 | TGCTGTTG | CGAAGAA( | NM_0314( | arsete resistance protein 2 | | 2.11 | SM2400 | c | 5 |
| mPool7_V2MM_180334 | -1.59438 | 3.004939 | TRUE | HP_40080 | TGCTGTTG | CCTCACAA | XM_28849 | Mus musculus LOC331843 (LOC331843), mRNA. | | 2.9 | SM2274 | e | 12 |
| 092308m3_V2MM_32592 | -1.58446 | 81.56557 | TRUE | HP_25547 | TGCTGTTG | GTATCTAT | AY266418 | expressed sequence AI553587 | AI553587 | 2.11 | SM2398 | f | 9 |
| mPool7_V2MM_104189 | -1.58242 | 3.004939 | TRUE | HP_32532 | TGCTGTTG | GCAAGAA( | XM_12957 | activating transcription factor 6 | Atf6 | 2.9 | SM2299 | e | 2 |
| 092308m1_V2MM_145253 | -1.58196 | 3.004939 | TRUE | HP_36596 | TGCTGTTG | CACTGAA/ | XM_1607( | Mus musculus LOC214721 (LOC214721), mR. | | 2.4 | SM2047 | h | 6 |
| 092308m1_V2MM_61821 | -1.58193 | 3.004939 | TRUE | HP_28394 | TGCTGTTG | CAATAGG/ | NM_0098( | catalase | Cat | 2.3 | SM2012 | e | 12 |
| mPool6_V2MM_63084 | -1.58116 | 94.45693 | TRUE | HP_28517 | TGCTGTTG | CCTCAGCT | NM_03018 | RIKEN cDNA 4930555I21 gene | 4930555I21Rik | 2.7 | SM2180 | g | 2 |
| mPool4_V2MM_22743 | -1.57862 | 79.16495 | TRUE | HP_24588 | TGCTGTTG | CTCATGG/ | NM_00869 | 4-nitrophenylphosphatase domain and non-neurona | Nipsnap1 | 2.15 | SM2568 | a | 1 |
| 092308m3_V2MM_51765 | -1.57826 | 84.31789 | TRUE | HP_27415 | TGCTGTTG | CTGAGCG | NM_01673 | GPI-anchored membrane protein 1 | Gpiap1 | 2.12 | SM2407 | f | 1 |
| mPool4_V2MM_202299 | -1.57639 | 70.87591 | TRUE | HP_42031 | TGCTGTTG | GCACTTG( | XM_12768 | RIKEN cDNA 4930474F22 gene | 4930474F22Rik | 2.16 | SM2617 | h | 7 |
| 092308m1_V2MM_148837 | -1.57423 | 3.004939 | TRUE | HP_36954 | TGCTGTTG | GAGCAAG | XM_16419 | Mus musculus LOC214871 (LOC214871), mR. | | 2.1 | SM2326 | e | 7 |
| 092308m1_V2MM_148837 | -1.57423 | 3.004939 | TRUE | HP_36954 | TGCTGTTG | GAGCAAG | XM_16419 | Mus musculus LOC214871 (LOC214871), mRNA. | | 2.1 | SM2326 | e | 7 |
| mPool2_V2MM_134882 | -1.57268 | 3.004939 | TRUE | HP_35560 | TGCTGTTG | GTCAGCA( | XM_15014 | guanylate cyclase 1, soluble, alpha 2 | Gucy1a2 | 2.11 | SM2360 | b | 7 |
| 092308m3_V2MM_103023 | -1.56319 | 94.45693 | TRUE | HP_32416 | TGCTGTTG | GCAAATG/ | AK010224 | protein phosphatase 1B, magnesium dependent, bet | Ppm1b | 2.1 | SM2321 | c | 12 |
| 092308m3_V2MM_103023 | -1.56319 | 94.45693 | TRUE | HP_32416 | TGCTGTTG | GCAAATG/ | AK010224 | protein phosphatase 1B, magnesium dependent, bet | Ppm1b | 2.1 | SM2321 | c | 12 |
| mPool4_V2MM_120517 | -1.56156 | 94.45693 | TRUE | HP_34141 | TGCTGTTG | CATATTAT | XM_14097 | similar to glyceraldehyde-3-phosphate dehydrogenase | | 2.16 | SM2632 | b | 6 |

| | | | | | | | | | | | | | |
|---|---|---|---|---|---|---|---|---|---|---|---|---|---|
| mPool2_V2MM_89190 | -1.55945 | 2.577834 | TRUE | HP_31062 | TGCTGTTG | GACTGTCC | NM_1773 | G protein-coupled receptor 149 | Gpr149 | 2.6 | SM2108 | e | 5 |
| mPool2_V2MM_89190 | -1.55945 | 2.577834 | TRUE | HP_31062 | TGCTGTTG | GACTGTCC | NM_1773 | G protein-coupled receptor 149 | Gpr149 | 2.6 | SM2108 | e | 5 |
| mPool6_V2MM_73215 | -1.55536 | 94.45693 | TRUE | HP_29505 | TGCTGTTG | GCCATGAC | NM_0136 | protein tyrosine phosphatase, non-receptor type 5 | Ptpn5 | 2.7 | SM2174 | f | 12 |
| 092308m1_V2MM_78488 | -1.55322 | 3.004939 | TRUE | HP_30017 | TGCTGTTG | GACTTCAT | NM_1467 | Mus musculus olfactory receptor MOR258-4P (MOR258-4P), mR. | | 2.3 | SM2011 | c | 12 |
| mPool7_V2MM_204887 | -1.55242 | 3.004939 | TRUE | HP_42277 | TGCTGTTG | CATCCTTT | XM_1363 | Mus musculus similar to ribosomal protein S24 [Rattus norvegicus] (LOC22664 | | 2.1 | SM2314 | h | 9 |
| mPool6_V2MM_93304 | -1.55031 | 94.45693 | TRUE | HP_29640 | TGCTGTTG | CAGATGCT | NM_1810 | SoxLZ/Sox6 leucine zipper binding protein in testis | Solt | 2.7 | SM2180 | c | 6 |
| mPool7_V2MM_137419 | -1.54713 | 2.577834 | TRUE | HP_35813 | TGCTGTTG | GCATCACA | XM_15313 | Mus musculus LOC238086 (LOC238086), mRNA. | | 2.1 | SM2327 | a | 12 |
| 092308m3_V2MM_85598 | -1.54514 | 83.62456 | TRUE | HP_30707 | TGCTGTTG | CTCCTTAA | NM_1464 | olfactory receptor 738 | Olfr738 | 2.13 | SM2473 | g | 5 |
| 092308m3_V2MM_42118 | -1.54333 | 70.87591 | TRUE | HP_26473 | TGCTGTTG | CAGTTAAA | NM_1770 | RIKEN cD 4932435O22 gene | 4932435O22Rik | 2.11 | SM2395 | h | 7 |
| mPool2_V2MM_131607 | -1.53988 | 3.004939 | TRUE | HP_35236 | TGCTGTTG | GCCACACT | NM_0010 | RIKEN cDNA A830005F24 gene | A830005F24Rik | 2.11 | SM2367 | f | 7 |
| mPool2_V2MM_86435 | -1.53751 | 2.577834 | TRUE | HP_30791 | TGCTGTTG | CGAGAGC | NM_1723 | solute carrier family 16 (monocarboxylic acid transp | Slc16a13 | 2.6 | SM2138 | d | 2 |
| mPool4_V2MM_241660 | -1.53351 | 73.80895 | TRUE | HP_49341 | TGCTGTTG | GCTTCCTT | XM_1460 | Mus musculus similar to pol protein [Homo sapiens] (LOC233858), mRNA. | | 2.14 | SM2532 | h | 1 |
| 092308m3_V2MM_227242 | -1.52956 | 94.45693 | TRUE | HP_47924 | TGCTGTTG | CTTCTCCC | XM_1112 | Mus musculus similar to Chain A, Horf6 A Novel Human Peroxidase Enzyme (LC | | 2.13 | SM2480 | d | 11 |
| 092308m3_V2MM_20884 | -1.52836 | 84.31789 | TRUE | HP_24406 | TGCTGTTG | CACCAGAC | NM_1756 | RIKEN cD A230050P20 gene | A230050P20Rik | 2.11 | SM2398 | e | 9 |
| mPool2_V2MM_100937 | -1.52681 | 3.004939 | TRUE | HP_32211 | TGCTGTTG | GACTGCTC | AK005570 | RIKEN cDNA 1600032L17 gene | 1600032L17Rik | 2.6 | SM2149 | f | 11 |
| mPool4_V2MM_261981 | -1.52658 | 91.10404 | TRUE | HP_52211 | TGCTGTTG | GATATCTC | NM_0272 | RIKEN cDNA 1810046I24 gene | 1810046I24Rik | 2.15 | SM2578 | e | 6 |
| 092308m3_V2MM_114121 | -1.52459 | 94.45693 | TRUE | HP_33510 | TGCTGTTG | CCTCAACT | XM_1372 | similar to E2a-Pbx1-associated protein; amyloid-beta precursor protein intracel | | 2.13 | SM2498 | c | 10 |
| 092308m1_V2MM_148683 | -1.52063 | 3.004939 | TRUE | HP_36939 | TGCTGTTG | CAGGTCA | XM_1640 | Mus musculus LOC244396 (LOC244396), mR. | | 2.4 | SM2033 | a | 5 |
| mPool2_V2MM_114687 | -1.52061 | 3.004939 | TRUE | HP_33566 | TGCTGTTG | GAAGGAC | XM_1378 | Mus musculus similar to hypothetical protein MGC15396 [Homo sapiens] (LOC | | 2.7 | SM2155 | b | 11 |
| 092308m1_V2MM_123359 | -1.5176 | 3.004939 | TRUE | HP_34421 | TGCTGTTG | GACCACA | XM_14291 | Mus musculus similar to ubiquitin-protein ligase (EC 6.3.2.19) - mouse (LOC207 | | 2.5 | SM2098 | f | 6 |
| 092308m1_V2MM_123359 | -1.5176 | 3.004939 | TRUE | HP_34421 | TGCTGTTG | GACCACA | XM_14291 | Mus musculus similar to ubiquitin-protein ligase (EC 6.3.2.19) - mouse (LOC207 | | 2.5 | SM2098 | f | 6 |
| mPool2_V2MM_110128 | -1.51571 | 3.004939 | TRUE | HP_33116 | TGCTGTTG | GCCACTTT | XM_13441 | expressed sequence AA589507 | AA589507 | 2.6 | SM2150 | a | 11 |
| mPool2_V2MM_212456 | -1.50644 | 3.004939 | TRUE | HP_43012 | TGCTGTTG | CAGCGGC | XM_1459 | Mus musculus similar to SKAP55 homologue [Mus musculus] (LOC233817), mR | | 2.11 | SM2355 | a | 6 |
| mPool4_V2MM_19230 | -1.50379 | 61.94057 | TRUE | HP_24245 | TGCTGTTG | GGCATTG | NM_0532 | RIKEN cDNA 4930433N12 gene | 4930433N12Rik | 2.15 | SM2586 | b | 10 |
| mPool4_V2MM_118091 | -1.50336 | 79.16495 | TRUE | HP_33902 | TGCTGTTG | GGTATCTT | XM_1396 | Mus musculus similar to helix-destabilizing protein - rat (LOC224379), mRNA. | | 2.11 | SM2352 | f | 10 |
| 092308m3_V2MM_43637 | -1.50296 | 84.31789 | TRUE | HP_26621 | TGCTGTTG | CTGACAGC | NM_1725 | expressed sequence AI661017 | AI661017 | 2.12 | SM2402 | a | 10 |
| mPool6_V2MM_11401 | -1.50209 | 61.94057 | TRUE | HP_23481 | TGCTGTTG | GAGATGA | NM_0077 | CDC like kinase 4 | Clk4 | 2.7 | SM2184 | e | 3 |
| 092308m1_V2MM_77037 | -1.50169 | 3.004939 | TRUE | HP_29877 | TGCTGTTG | CTCTGTGC | NM_0105 | Mus musculus internexin neurol intermediate filament protein, alpha (I), mR. | | 2.4 | SM2025 | e | 3 |
| mPool4_V2MM_181159 | -1.50165 | 91.71214 | TRUE | HP_40163 | TGCTGTTG | CACTACAA | XM_2886 | Mus musculus LOC328450 (LOC328450), mRNA. | | NA | | | NA |
| 092308m3_V2MM_225247 | -1.50087 | 84.31789 | TRUE | HP_47729 | TGCTGTTG | CAAATATT | XM_4875 | similar to peptidylprolyl isomerase A | | 2.13 | SM2456 | c | 6 |
| mPool4_V2MM_154815 | -1.49881 | 92.61223 | TRUE | HP_37545 | TGCTGTTG | CACACTCT | XM_1970 | Mus musculus LOC271091 (LOC271091), mRNA. | | NA | | | NA |
| 092308m3_V2MM_235032 | -1.49851 | 61.94057 | TRUE | HP_48690 | TGCTGTTG | TTCAACTC | XM_1953 | Mus musculus similar to hypothetical protein C230069C04 [Mus musculus] (LO | | 2.14 | SM2527 | b | 9 |
| mPool4_V2MM_170940 | -1.49811 | 84.31789 | TRUE | HP_39143 | TGCTGTTG | GTGCCTCT | XM_2865 | gene model 806, (NCBI) | Gm806 | NA | | | NA |
| 092308m3_V2MM_99591 | -1.49585 | 83.62456 | TRUE | HP_32078 | TGCTGTTG | CTAGTCTC | NM_0100 | cytochrome P450, family 27, subfamily b, polypeptic | Cyp27b1 | 2.13 | SM2467 | g | 5 |
| mPool6_V2MM_190416 | -1.49279 | 94.45693 | TRUE | HP_30118 | TGCTGTTG | GCATAATT | NM_1984 | creatine kinase, mitochondrial 2 | Ckmt2 | 2.7 | SM2186 | d | 2 |
| mPool2_V2MM_38692 | -1.48997 | 3.004939 | TRUE | HP_26140 | TGCTGTTG | GCCCGGA | NM_0117 | adenomatosis polyposis coli 2 | Apc2 | 2.11 | SM2389 | f | 6 |
| 092308m1_V2MM_123348 | -1.48953 | 3.004939 | TRUE | HP_34420 | TGCTGTTG | GCCTTGAC | XM_4882 | similar to Ig heavy chain precursor V region (108A) - mouse | | 2.6 | SM2103 | e | 12 |
| 092308m3_V2MM_236722 | -1.4882 | 70.87591 | TRUE | HP_48857 | TGCTGTTG | CTGGGAA | XM_1579 | Mus musculus LOC225982 (LOC225982), mR. | | 2.14 | SM2509 | e | 10 |
| 092308m3_V2MM_83145 | -1.48512 | 61.94057 | TRUE | HP_30469 | TGCTGTTG | CTTCGAGA | BC057080 | ATP-binding cassette, sub-family C (CFTR/MRP), mer | Abcc8 | 2.13 | SM2460 | e | 8 |
| 092308m1_V2MM_147672 | -1.48486 | 3.004939 | TRUE | HP_36838 | TGCTGTTG | CAAAGGG | XM_16317 | Mus musculus LOC243916 (LOC243916), mR. | | 2.4 | SM2034 | a | 6 |
| 092308m1_V2MM_147672 | -1.48486 | 3.004939 | TRUE | HP_36838 | TGCTGTTG | CAAAGGG | XM_16317 | Mus musculus LOC243916 (LOC243916), mRNA. | | 2.4 | SM2034 | a | 6 |
| mPool7_V2MM_179060 | -1.48276 | 2.577834 | TRUE | HP_39953 | TGCTGTTG | GCACAACT | XM_2882 | Mus musculus LOC332541 (LOC332541), mRNA. | | 2.9 | SM2278 | d | 2 |
| 092308m1_V2MM_131572 | -1.48124 | 3.004939 | TRUE | HP_35233 | TGCTGTTG | CTTTGAGT | XM_14771 | EDAR (ectodysplasin-A receptor)-associated death d | Edaradd | 2.5 | SM2099 | b | 5 |
| 092308m3_V2MM_53367 | -1.48018 | 83.62456 | TRUE | HP_27573 | TGCTGTTG | GTGGCGC | NM_0167 | ninjurin 2 | Ninj2 | 2.12 | SM2444 | e | 3 |
| 092308m1_V2MM_185950 | -1.47997 | 3.004939 | TRUE | HP_40642 | TGCTGTTG | CACCCAGA | XM_2894 | Mus musculus LOC333141 (LOC333141), mRNA. | | 2.5 | SM2091 | a | 11 |
| 092308m1_V2MM_185950 | -1.47997 | 3.004939 | TRUE | HP_40642 | TGCTGTTG | CACCCAGA | XM_2894 | Mus musculus LOC333141 (LOC333141), mR. | | 2.5 | SM2091 | a | 11 |
| mPool4_V2MM_51380 | -1.47394 | 84.31789 | TRUE | HP_27378 | TGCTGTTG | CTCAGTAA | NM_1471 | olfactory receptor 974 | Olfr974 | 2.15 | SM2590 | f | 1 |
| mPool7_V2MM_202787 | -1.47141 | 3.004939 | TRUE | HP_42077 | TGCTGTTG | CTGCTGTA | XM_1267 | keratin complex 1, acidic, gene 5 | Krt1-5 | 2.9 | SM2300 | c | 5 |
| 092308m1_V2MM_139512 | -1.47102 | 3.004939 | TRUE | HP_36023 | TGCTGTTG | GATACTCA | XM_15513 | Mus musculus LOC239257 (LOC239257), mRNA. | | 2.5 | SM2058 | b | 11 |
| 092308m1_V2MM_139512 | -1.47102 | 3.004939 | TRUE | HP_36023 | TGCTGTTG | GATACTCA | XM_15513 | Mus musculus LOC239257 (LOC239257), mR. | | 2.5 | SM2058 | b | 11 |
| mPool4_V2MM_240550 | -1.46993 | 61.94057 | TRUE | HP_49233 | TGCTGTTG | CATACCTC | XM_1489 | Mus musculus LOC212511 (LOC212511), mRNA. | | 2.14 | SM2532 | b | 8 |

| | | | | | | | | | | | | | | | |
|---|---|---|---|---|---|---|---|---|---|---|---|---|---|---|---|
| 092308m3_V2MM_234151 | -1.46913 | 87.58579 | TRUE | HP_48603 | TGCTGTTG | GTTATATC | XM_12887 | RIKEN cD 4833446K15 gene | 4833446K15Rik | | 2.13 | SM2486 | d | | 11 |
| 092308m1_V2MM_66996 | -1.46774 | 3.004939 | TRUE | HP_28899 | TGCTGTTG | CCCAGGT( | NM_0098: | calsequestrin 2 | Casq2 | | 2.4 | SM2028 | c | | 2 |
| 092308m1_V2MM_133568 | -1.4664 | 3.004939 | TRUE | HP_35430 | TGCTGTTG | CAGTTCAC | XM_14932 | RIKEN cD 4930565D16 gene | 4930565D16Rik | | 2.6 | SM2101 | a | | 8 |
| mPool4_V2MM_37223 | -1.46383 | 91.71214 | TRUE | HP_25998 | TGCTGTTG | GACATGG | NM_1447{ | jumonji domain containing 2C | Jmjd2c | | 2.15 | SM2593 | d | | 11 |
| 092308m3_V2MM_208722 | -1.46253 | 61.94057 | TRUE | NA | NA | NA | NA | NA | NA | NA | | NA | NA | NA | |
| 092308m3_V2MM_101581 | -1.46136 | 61.94057 | TRUE | HP_32274 | TGCTGTTG | GATGTAA( | XM_12772 | Mus musculus RIKEN cD 5830451P18 gene (5830451P18Rik), mR. | | | 2.13 | SM2497 | f | | 8 |
| mPool4_V2MM_175055 | -1.46101 | 66.95036 | TRUE | HP_39554 | TGCTGTTG | CCTCCATG | XM_2873( | Mus musculus hypothetical gene supported by AK034470 (LOC329914), mRNA. | | NA | | | | NA | |
| 092308m3_V2MM_94727 | -1.4605 | 79.16495 | TRUE | HP_31601 | TGCTGTTG | CACCTAG( | NM_1832: | hypothetical protein D730018G16 | | | 2.13 | SM2457 | b | | 8 |
| 092308m3_V2MM_107726 | -1.45865 | 81.56557 | TRUE | HP_32880 | TGCTGTTG | GCATGCTT | XM_1322{ | RIKEN cD 2310005C01 gene | 2310005C01Rik | | 2.13 | SM2487 | d | | 11 |
| 092308m3_V2MM_56000 | -1.45703 | 61.94057 | TRUE | HP_27829 | TGCTGTTG | CGGCTATT | NM_0318( | taste receptor, type 1, member 1 | Tas1r1 | | 2.12 | SM2438 | a | | 1 |
| mPool7_V2MM_107310 | -1.45554 | 3.004939 | TRUE | NA | NA | NA | NA | NA | NA | NA | | NA | NA | NA | |
| mPool2_V2MM_125122 | -1.45538 | 2.577834 | TRUE | HP_34596 | TGCTGTTG | CCAATCGA | XM_1438( | Mus musculus similar to alpha-2u-globulin IV precursor - mouse (LOC209155), ı | | | 2.11 | SM2370 | f | | 1 |
| 092308m3_V2MM_85733 | -1.4537 | 87.58579 | TRUE | HP_30721 | TGCTGTTG | CAGACAC( | NM_1464! | olfactory receptor 693 | Olfr693 | | 2.13 | SM2466 | c | | 1 |
| mPool4_V2MM_179954 | -1.45351 | 61.94057 | TRUE | HP_40042 | TGCTGTTG | CTTCCCTA | XM_28842 | Unknown (protein for MGC:58818) | | NA | | | | NA | |
| mPool4_V2MM_193883 | -1.45134 | 61.94057 | TRUE | HP_7949 | TGCTGTTG | CAGTTCTC | XM_19564 | Spi-B transcription factor (Spi-1/PU.1 related) | Spib | NA | | | | NA | |
| 092308m3_V2MM_39102 | -1.44456 | 70.87591 | TRUE | HP_26180 | TGCTGTTG | GCTCAACT | NM_0269: | CD209b antigen | Cd209b | | 2.12 | SM2424 | b | | 9 |
| 092308m1_V2MM_139336 | -1.44353 | 3.004939 | TRUE | HP_36005 | TGCTGTTG | GACACAC1 | XM_15494 | Mus musculus LOC211279 (LOC211279), mR. | | | 2.4 | SM2034 | e | | 2 |
| 092308m1_V2MM_139336 | -1.44353 | 3.004939 | TRUE | HP_36005 | TGCTGTTG | GACACAC1 | XM_15494 | Mus musculus LOC211279 (LOC211279), mRNA. | | | 2.4 | SM2034 | e | | 2 |
| mPool7_V2MM_173742 | -1.4395 | 3.004939 | TRUE | HP_39422 | TGCTGTTG | CCCAACCC | XM_2870{ | Mus musculus LOC329351 (LOC329351), mR. | | | 2.5 | SM2063 | b | | 7 |
| mPool7_V2MM_173742 | -1.4395 | 3.004939 | TRUE | HP_39422 | TGCTGTTG | CCCAACCC | XM_2870{ | Mus musculus LOC329351 (LOC329351), mRNA. | | | 2.5 | SM2063 | b | | 7 |
| mPool2_V2MM_131980 | -1.43675 | 3.004939 | TRUE | HP_35274 | TGCTGTTG | CTACCCAC | XM_1479: | ATP-binding cassette, sub-family D (ALD), member 2 | Abcd2 | | 2.11 | SM2369 | d | | 10 |
| mPool7_V2MM_145243 | -1.4367 | 3.004939 | TRUE | HP_36595 | TGCTGTTG | CAGGCTT( | XM_1607( | Mus musculus LOC214719 (LOC214719), mRNA. | | | 2.5 | SM2052 | c | | 11 |
| mPool7_V2MM_145243 | -1.4367 | 3.004939 | TRUE | HP_36595 | TGCTGTTG | CAGGCTT( | XM_1607( | Mus musculus LOC214719 (LOC214719), mR. | | | 2.5 | SM2052 | c | | 11 |
| mPool7_V2MM_83310 | -1.43126 | 3.004939 | TRUE | HP_30485 | TGCTGTTG | GACGCCA1 | NM_0120: | G protein-regulated inducer of neurite outgrowth 1 | Gprin1 | | 2.9 | SM2258 | b | | 7 |
| 092308m1_V2MM_153108 | -1.43075 | 3.004939 | TRUE | HP_37376 | TGCTGTTG | GTGTCAA1 | XM_19574 | Mus musculus similar to olfactory receptor MOR168-1 [Mus musculus] (LOC27( | | | 2.6 | SM2124 | b | | 9 |
| mPool2_V2MM_29618 | -1.42881 | 3.004939 | TRUE | HP_25257 | TGCTGTTG | CACGTTCC | NM_0086: | microtubule-associated protein 1 B | Mtap1b | | 2.11 | SM2389 | b | | 9 |
| mPool4_V2MM_94893 | -1.42359 | 61.94057 | TRUE | HP_31617 | TGCTGTTG | CAGTCTCT | NM_1832{ | RIKEN cDNA 2310005G13 gene | 2310005G13Rik | | 2.16 | SM2613 | c | | 12 |
| mPool2_V2MM_134470 | -1.42356 | 3.004939 | TRUE | HP_35519 | TGCTGTTG | CCACCTG( | XM_1498! | Mus musculus LOC232948 (LOC232948), mR. | | | 2.6 | SM2102 | d | | 5 |
| mPool2_V2MM_134470 | -1.42356 | 3.004939 | TRUE | HP_35519 | TGCTGTTG | CCACCTG( | XM_1498! | Mus musculus LOC232948 (LOC232948), mRNA. | | | 2.6 | SM2102 | d | | 5 |
| 092308m3_V2MM_21318 | -1.42149 | 61.94057 | TRUE | HP_24449 | TGCTGTTG | CAGCTAT( | NM_0282: | potassium large conductance calcium-activated char | Kcnmb2 | | 2.11 | SM2399 | a | | 6 |
| mPool2_V2MM_105479 | -1.42045 | 3.004939 | TRUE | HP_32659 | TGCTGTTG | CAGTGCA( | XM_1305: | ribosome binding protein 1 | Rrbp1 | | 2.7 | SM2163 | f | | 4 |
| mPool2_V2MM_128296 | -1.41817 | 3.004939 | TRUE | HP_34911 | TGCTGTTG | GCAAGGA | XM_14562 | Mus musculus similar to envelope protein [Ovis aries] (LOC233260), mRNA. | | | 2.11 | SM2358 | c | | 12 |
| 092308m1_V2MM_83373 | -1.41801 | 3.004939 | TRUE | HP_30491 | TGCTGTTG | CACAACA( | NM_0135{ | low density lipoprotein receptor-related protein ass | Lrpap1 | | 2.6 | SM2112 | e | | 6 |
| 092308m1_V2MM_83373 | -1.41801 | 3.004939 | TRUE | HP_30491 | TGCTGTTG | CACAACA( | NM_0135{ | low density lipoprotein receptor-related protein ass | Lrpap1 | | 2.6 | SM2112 | e | | 6 |
| mPool2_V2MM_133342 | -1.41632 | 3.004939 | TRUE | HP_35408 | TGCTGTTG | CAAGTGG( | NM_2072( | expressed sequence AI505034 | AI505034 | | 2.11 | SM2367 | h | | 5 |
| mPool2_V2MM_130089 | -1.41596 | 3.004939 | TRUE | HP_35088 | TGCTGTTG | CTGAAGG( | XM_1466( | DNA segment, Chr 8, ERATO Doi 738, expressed | D8Ertd738e | | 2.11 | SM2352 | g | | 11 |
| 092308m3_V2MM_44041 | -1.41573 | 94.45693 | TRUE | HP_26660 | TGCTGTTG | CTGTGACT | NM_0096 | activation-induced cytidine deamise | Aicda | | 2.12 | SM2412 | a | | 10 |
| 092308m1_V2MM_122265 | -1.40751 | 2.577834 | TRUE | HP_34313 | TGCTGTTG | GGAAGTA | XM_1421( | similar to RIKEN cD 9230105E10 gene | | | 2.6 | SM2105 | f | | 3 |
| mPool7_V2MM_94292 | -1.40665 | 3.004939 | TRUE | HP_31558 | TGCTGTTG | GTCCAGA( | NM_1831: | RIKEN cDNA 9530002K18 gene | 9530002K18Rik | | 2.6 | SM2139 | f | | 1 |
| 092308m3_V2MM_23589 | -1.4048 | 94.45693 | TRUE | HP_24671 | TGCTGTTG | CTTATGAC | NM_1724: | FERM domain containing 4A | Frmd4a | | 2.12 | SM2424 | g | | 11 |
| mPool4_V2MM_207015 | -1.40442 | 73.80895 | TRUE | HP_42483 | TGCTGTTG | GTCCTTCA | XM_19507 | Mus musculus LOC271513 (LOC271513), mRNA. | | NA | | | | NA | |
| mPool6_V2MM_169664 | -1.40426 | 91.71214 | TRUE | HP_39015 | TGCTGTTG | CTGAAGA1 | AK039689 | hypothetical protein A330087I24 | | | 2.7 | SM2197 | a | | 1 |
| mPool4_V2MM_167242 | -1.40374 | 94.45693 | TRUE | HP_38775 | TGCTGTTG | GAACTATT | XM_28572 | Mus musculus similar to Extracellular calcium-sensing receptor precursor (CaSF | | NA | | | | NA | |
| 092308m1_V2MM_179859 | -1.40243 | 3.004939 | TRUE | HP_40033 | TGCTGTTG | GACACTG( | XM_2884( | Mus musculus LOC331732 (LOC331732), mR. | | | 2.5 | SM2085 | g | | 3 |
| 092308m1_V2MM_179859 | -1.40243 | 3.004939 | TRUE | HP_40033 | TGCTGTTG | GACACTG( | XM_2884( | Mus musculus LOC331732 (LOC331732), mRNA. | | | 2.5 | SM2085 | g | | 3 |
| 092308m1_V2MM_154726 | -1.40196 | 3.004939 | TRUE | HP_37536 | TGCTGTTG | GTGAGAA | XM_1969{ | cD sequence BC065123 | BC065123 | | 2.6 | SM2114 | g | | 10 |
| mPool7_V2MM_81662 | -1.39931 | 3.004939 | TRUE | HP_30326 | TGCTGTTG | GTGACAC1 | NM_0256( | RIKEN cDNA 2610029K21 gene | 2610029K21Rik | | 2.8 | SM2244 | g | | 10 |
| mPool2_V2MM_155044 | -1.3989 | 3.004939 | TRUE | HP_37568 | TGCTGTTG | CATCTGG( | XM_1971( | Mus musculus LOC271249 (LOC271249), mRNA. | | | 2.6 | SM2126 | e | | 12 |
| 092308m3_V2MM_27067 | -1.39768 | 84.31789 | TRUE | HP_25009 | TGCTGTTG | CTGGTATT | NM_1338( | fibrinogen, gamma polypeptide | Fgg | | 2.11 | SM2393 | e | | 7 |
| mPool4_V2MM_129244 | -1.3963 | 69.72441 | TRUE | HP_35005 | TGCTGTTG | CTTACGTC | XM_1462( | RIKEN cDNA A130072A22 gene | A130072A22Rik | | 2.14 | SM2550 | b | | 11 |
| mPool7_V2MM_111439 | -1.39518 | 2.577834 | TRUE | HP_33245 | TGCTGTTG | CGAGATC( | XM_1470( | Mus musculus similar to glyceraldehyde-3-phosphate dehydrogenase [Mus mu | | | 2.1 | SM2304 | a | | 4 |

| | | | | | | | | | | | | | |
|---|---|---|---|---|---|---|---|---|---|---|---|---|---|
| 092308m3_V2MM_59827 | -1.39358 | 61.94057 | TRUE | HP_28200 | TGCTGTTG | GCTGGTTT | NM_1731 | amyotrophic lateral sclerosis 2 (juvenile) chromoson | Als2cr2 | 2.12 | SM2446 | b | 4 |
| 092308m1_V2MM_154672 | -1.39325 | 3.004939 | TRUE | HP_37531 | TGCTGTTG | GTTGACA | XM_1969 | Mus musculus LOC270974 (LOC270974), mR. | | 2.6 | SM2117 | c | 1 |
| mPool4_V2MM_124567 | -1.39318 | 75.98381 | TRUE | HP_34541 | TGCTGTTG | CTTTGCTC | XM_1435 | Mus musculus LOC242201 (LOC242201), mRNA. | | 2.14 | SM2537 | b | 1 |
| mPool2_V2MM_98122 | -1.39291 | 3.004939 | TRUE | HP_31935 | TGCTGTTG | GACATCA | XM_1121 | Mus musculus similar to RNP particle component [Mus musculus] (LOC195745) | | 2.6 | SM2108 | d | 7 |
| mPool2_V2MM_98122 | -1.39291 | 3.004939 | TRUE | HP_31935 | TGCTGTTG | GACATCA | XM_1121 | Mus musculus similar to RNP particle component [Mus musculus] (LOC195745) | | 2.6 | SM2108 | d | 7 |
| 092308m1_V2MM_156913 | -1.39274 | 3.004939 | TRUE | HP_37753 | TGCTGTTG | CATCGCAC | XM_2045 | Mus musculus similar to hypothetical protein FLJ35757 [Homo sapiens] (LOC27 | | 2.6 | SM2125 | f | 8 |
| mPool4_V2MM_32654 | -1.39156 | 83.62456 | TRUE | HP_25553 | TGCTGTTG | GTCTAAAT | NM_1448 | RIKEN cDNA 2810036L13 gene | 2810036L13Rik | 2.15 | SM2599 | g | 3 |
| 092308m3_V2MM_88803 | -1.38574 | 61.94057 | TRUE | HP_31023 | TGCTGTTG | CGTAGAA | NM_1754 | paraneoplastic antigen MA2 | Pnma2 | 2.13 | SM2461 | a | 5 |
| 092308m3_V2MM_225072 | -1.38506 | 92.61223 | TRUE | HP_47712 | TGCTGTTG | CTATAAGT | NM_0117 | WW domain binding protein 5 | Wbp5 | 2.13 | SM2485 | c | 8 |
| mPool2_V2MM_154724 | -1.38502 | 3.004939 | TRUE | HP_37536 | TGCTGTTG | GAGAGAC | XM_1969 | cDNA sequence BC065123 | BC065123 | 2.6 | SM2127 | d | 5 |
| mPool2_V2MM_118498 | -1.38059 | 3.004939 | TRUE | HP_33943 | TGCTGTTG | CTGCCAG | XM_13987 | similar to ZFP421 | | 2.11 | SM2374 | h | 1 |
| mPool6_V2MM_158432 | -1.37853 | 61.94057 | TRUE | HP_37904 | TGCTGTTG | CTTGCCAA | XM_2066 | Mus musculus LOC279600 (LOC279600), mRNA. | | 2.7 | SM2190 | e | 8 |
| 092308m3_V2MM_225897 | -1.37735 | 70.87591 | TRUE | HP_47793 | TGCTGTTG | CCGTTGTT | NM_1753 | RIKEN cD B630019K06 gene | B630019K06Rik | 2.13 | SM2483 | e | 7 |
| mPool7_V2MM_170934 | -1.3767 | 3.004939 | TRUE | HP_39142 | TGCTGTTG | CCAAGCA | AK079388 | RIKEN cDNA D130012G24 gene | D130012G24Rik | 2.9 | SM2284 | c | 2 |
| 092308m3_V2MM_101489 | -1.37251 | 94.45693 | TRUE | HP_32266 | TGCTGTTG | GGAAGCA | XM_1276 | RIKEN cDNA 9230112D13 gene | 9230112D13Rik | 2.1 | SM2305 | a | 9 |
| 092308m3_V2MM_101489 | -1.37251 | 94.45693 | TRUE | HP_32266 | TGCTGTTG | GGAAGCA | XM_1276 | RIKEN cD 9230112D13 gene | 9230112D13Rik | 2.1 | SM2305 | a | 9 |
| mPool2_V2MM_215149 | -1.3689 | 2.577834 | TRUE | HP_43271 | TGCTGTTG | GCCACATA | XM_14231 | Mus musculus similar to helix-destabilizing protein - rat (LOC237236), mRNA. | | 2.11 | SM2369 | d | 7 |
| mPool2_V2MM_204126 | -1.36833 | 3.004939 | TRUE | HP_42203 | TGCTGTTG | CAGCCGC | BC020182 | RIKEN cDNA 2700082O15 gene | 2700082O15Rik | 2.1 | SM2316 | b | 11 |
| 092308m3_V2MM_42370 | -1.36728 | 66.95036 | TRUE | HP_26497 | TGCTGTTG | CCAATAGT | NM_0168 | metaxin 2 | Mtx2 | 2.12 | SM2404 | a | 3 |
| mPool7_V2MM_101678 | -1.36617 | 3.004939 | TRUE | HP_32284 | TGCTGTTG | CAGAGAC | XM_1277 | RIKEN cDNA 1300010F03 gene | 1300010F03Rik | 2.9 | SM2299 | a | 11 |
| mPool7_V2MM_101678 | -1.36617 | 3.004939 | TRUE | HP_32284 | TGCTGTTG | CAGAGAC | XM_1277 | RIKEN cD 1300010F03 gene | 1300010F03Rik | 2.9 | SM2299 | a | 11 |
| mPool4_V2MM_82725 | -1.36544 | 83.62456 | TRUE | HP_30428 | TGCTGTTG | GGATTCG | NM_0102 | folylpolyglutamyl synthetase | Fpgs | 2.15 | SM2571 | c | 8 |
| mPool6_V2MM_159484 | -1.36379 | 61.94057 | TRUE | HP_38007 | TGCTGTTG | GTTCTAGC | XM_2831 | Mus musculus hypothetical gene supported by AK044328 (LOC328229), mRNA. | | 2.7 | SM2195 | c | 8 |
| 092308m1_V2MM_99611 | -1.36241 | 3.004939 | TRUE | HP_32080 | TGCTGTTG | GGCACAT | XM_12592 | glutamise 2 (liver, mitochondrial) | Gls2 | 2.6 | SM2107 | a | 1 |
| mPool4_V2MM_187384 | -1.36142 | 61.94057 | TRUE | HP_40785 | TGCTGTTG | CAGACAA | XM_28974 | Mus musculus LOC333655 (LOC333655), mRNA. | | NA | | | NA |
| mPool7_V2MM_173741 | -1.36138 | 3.004939 | TRUE | HP_39422 | TGCTGTTG | CCAACCCA | XM_2870 | Mus musculus LOC329351 (LOC329351), mR. | | 2.5 | SM2063 | g | 11 |
| mPool7_V2MM_173741 | -1.36138 | 3.004939 | TRUE | HP_39422 | TGCTGTTG | CCAACCCA | XM_2870 | Mus musculus LOC329351 (LOC329351), mRNA. | | 2.5 | SM2063 | g | 11 |
| 092308m3_V2MM_98954 | -1.36056 | 94.45693 | TRUE | HP_32017 | TGCTGTTG | CAGTGCTT | XM_1250 | similar to RIKEN cD 2610524H06 gene | | 2.13 | SM2454 | a | 7 |
| 092308m3_V2MM_42986 | -1.35869 | 92.61223 | TRUE | HP_26558 | TGCTGTTG | CGGTCGA | NM_1725 | RIKEN cD 5730420B22 gene | 5730420B22Rik | 2.12 | SM2437 | c | 1 |
| 092308m1_V2MM_185518 | -1.3564 | 3.004939 | TRUE | HP_40599 | TGCTGTTG | GACATCCT | XM_2893 | Mus musculus LOC333052 (LOC333052), mR. | | 2.5 | SM2064 | g | 8 |
| 092308m3_V2MM_146213 | -1.35602 | 84.31789 | TRUE | HP_36692 | TGCTGTTG | CCAAGAA | XM_1618 | Mus musculus LOC243054 (LOC243054), mR. | | 2.14 | SM2518 | c | 9 |
| mPool6_V2MM_157770 | -1.35372 | 61.94057 | TRUE | HP_37838 | TGCTGTTG | GAGGTAG | XM_2055 | gene model 725, (NCBI) | Gm725 | 2.7 | SM2200 | h | 9 |
| mPool4_V2MM_180763 | -1.35284 | 61.94057 | TRUE | HP_40123 | TGCTGTTG | CCAAGTC | XM_2885 | Mus musculus LOC331947 (LOC331947), mRNA. | | NA | | | NA |
| 092308m1_V2MM_142390 | -1.35274 | 3.004939 | TRUE | HP_36310 | TGCTGTTG | GCCAGAA | XM_15761 | Mus musculus LOC212132 (LOC212132), mRNA. | | 2.5 | SM2053 | c | 11 |
| 092308m1_V2MM_142390 | -1.35274 | 3.004939 | TRUE | HP_36310 | TGCTGTTG | GCCAGAA | XM_15761 | Mus musculus LOC212132 (LOC212132), mR. | | 2.5 | SM2053 | c | 11 |
| mPool4_V2MM_238628 | -1.35262 | 94.45693 | TRUE | HP_49045 | TGCTGTTG | CAAGAAT | XM_1442 | Mus musculus similar to RIKEN cDNA 4933431D05 [Mus musculus] (LOC21132 | | 2.14 | SM2533 | a | 4 |
| mPool4_V2MM_163605 | -1.34703 | 81.56557 | TRUE | HP_38416 | TGCTGTTG | CAATCACA | XM_28482 | Mus musculus similar to preferentially expressed antigen in melanoma; melan | | NA | | | NA |
| 092308m3_V2MM_44066 | -1.34618 | 79.16495 | TRUE | HP_26663 | TGCTGTTG | CTGTGGC | U83509 | angiopoietin 1 | Angpt1 | 2.12 | SM2445 | f | 9 |
| mPool4_V2MM_120385 | -1.34548 | 73.80895 | TRUE | HP_34128 | TGCTGTTG | CATCATCA | XM_1408 | Mus musculus similar to KIAA1422 protein [Homo sapiens] (LOC241281), mRN | | 2.14 | SM2531 | g | 4 |
| mPool7_V2MM_105908 | -1.34458 | 3.004939 | TRUE | HP_32701 | TGCTGTTG | GATGCAC | XM_1308 | progesterone receptor membrane component 2 | Pgrmc2 | 2.1 | SM2306 | e | 12 |
| mPool2_V2MM_13680 | -1.34365 | 3.004939 | TRUE | HP_23704 | TGCTGTTG | CCACCGA | NM_0264 | S100 calcium binding protein A16 | S100a16 | 2.11 | SM2374 | d | 9 |
| mPool2_V2MM_89835 | -1.34335 | 3.004939 | TRUE | HP_31126 | TGCTGTTG | CTCCTTAC | BC048817 | RIKEN cDNA A730098D12 gene | A730098D12Rik | 2.6 | SM2138 | c | 2 |
| mPool7_V2MM_208064 | -1.34097 | 3.004939 | TRUE | HP_42586 | TGCTGTTG | CCTGCTCT | XM_4840 | hypothetical protein B230105J10 | | 2.1 | SM2325 | h | 12 |
| mPool4_V2MM_112349 | -1.33971 | 94.45693 | TRUE | HP_33335 | TGCTGTTG | CTGCCCAT | XM_1364 | Mus musculus similar to pol protein (LOC226335), mRNA. | | 2.16 | SM2622 | d | 1 |
| 092308m1_V2MM_66675 | -1.33916 | 3.004939 | TRUE | HP_28868 | TGCTGTTG | CATCTGG | NM_1768 | RIKEN cD 1110055N21 gene | 1110055N21Rik | 2.4 | SM2025 | g | 1 |
| mPool4_V2MM_123074 | -1.33868 | 91.71214 | TRUE | HP_34393 | TGCTGTTG | GTGTTCTA | XM_4854 | similar to RIKEN cDNA 1700029I01 | | 2.14 | SM2540 | f | 7 |
| 092308m1_V2MM_173741 | -1.33731 | 3.004939 | TRUE | HP_39422 | TGCTGTTG | CCAACCCA | XM_2870 | Mus musculus LOC329351 (LOC329351), mR. | | 2.5 | SM2063 | g | 11 |
| 092308m1_V2MM_173741 | -1.33731 | 3.004939 | TRUE | HP_39422 | TGCTGTTG | CCAACCCA | XM_2870 | Mus musculus LOC329351 (LOC329351), mRNA. | | 2.5 | SM2063 | g | 11 |
| 092308m3_V2MM_38253 | -1.33723 | 91.71214 | TRUE | HP_26098 | TGCTGTTG | GATTTCAT | NM_1723 | RIKEN cD A530046H20 gene | A530046H20Rik | 2.11 | SM2400 | a | 9 |
| mPool4_V2MM_47319 | -1.33562 | 70.87591 | TRUE | HP_26980 | TGCTGTTG | CTCTTATT | AK083728 | RIKEN cDNA 2410004I17 gene | 2410004I17Rik | 2.16 | SM2609 | a | 5 |
| 092308m1_V2MM_66766 | -1.33537 | 3.004939 | TRUE | HP_28877 | TGCTGTTG | CCAAGAG | BC039216 | CLIP associating protein 1 | Clasp1 | 2.4 | SM2023 | g | 7 |

| | | | | | | | | | | | | | | | |
|---|---|---|---|---|---|---|---|---|---|---|---|---|---|---|---|
| mPool2_V2MM_117196 | -1.33516 | 3.004939 | TRUE | HP_33814 | TGCTGTTG | CAAGCACT | XM_13925 | Mus musculus similar to Eukaryotic translation initiation factor 4B (eIF-4B) (LO( | | | 2.7 | SM2156 | b | | 12 |
| 092308m3_V2MM_85604 | -1.32808 | 61.94057 | TRUE | HP_30708 | TGCTGTTG | CAATATTC | NM_1464 | olfactory receptor 887 | Olfr887 | | 2.13 | SM2457 | c | | 5 |
| 092308m1_V2MM_167114 | -1.32572 | 3.004939 | TRUE | HP_38762 | TGCTGTTG | CTAATGTT | XM_2856 | similar to meltrin alpha | | | 2.6 | SM2116 | b | | 9 |
| mPool6_V2MM_25178 | -1.32393 | 79.16495 | TRUE | HP_24825 | TGCTGTTG | CAATTGA | NM_1337 | RIKEN cDNA 2610018G03 gene | 2610018G03Rik | | 2.7 | SM2182 | a | | 4 |
| 092308m1_V2MM_175119 | -1.32324 | 3.004939 | TRUE | HP_39560 | TGCTGTTG | CAGTTCAC | AK045802 | CUB and Sushi multiple domains 2 | Csmd2 | | 2.5 | SM2082 | a | | 7 |
| 092308m1_V2MM_175119 | -1.32324 | 3.004939 | TRUE | HP_39560 | TGCTGTTG | CAGTTCAC | AK045802 | CUB and Sushi multiple domains 2 | Csmd2 | | 2.5 | SM2082 | a | | 7 |
| mPool6_V2MM_153077 | -1.32323 | 91.71214 | TRUE | HP_37373 | TGCTGTTG | GCCTCTAT | XM_19572 | Mus musculus similar to ribosomal protein S24 [Rattus norvegicus] (LOC27009 | | | 2.7 | SM2191 | a | | 1 |
| 092308m3_V2MM_205209 | -1.31904 | 83.62456 | TRUE | HP_42308 | TGCTGTTG | GGCAACG | XM_13784 | type I hair keratin KA36 | | | 2.13 | SM2486 | a | | 9 |
| mPool4_V2MM_160039 | -1.3186 | 91.10404 | TRUE | HP_38062 | TGCTGTTG | CACTCAAT | AK028773 | gene model 1600, (NCBI) | Gm1600 | NA | | | | NA | |
| mPool4_V2MM_5128 | -1.31635 | 84.31789 | TRUE | HP_22868 | TGCTGTTG | CATGTATC | NM_0194 | pannexin 1 | Panx1 | | 2.15 | SM2569 | b | | 4 |
| mPool4_V2MM_1623 | -1.31557 | 83.62456 | TRUE | HP_22526 | TGCTGTTG | CGAATTAT | NM_0258 | prolactin like protein I | | | 2.15 | SM2579 | a | | 9 |
| mPool4_V2MM_241330 | -1.31555 | 66.95036 | TRUE | HP_49309 | TGCTGTTG | GGTCAAA | NM_1990 | cDNA sequence BC057593 | BC057593 | | 2.14 | SM2543 | h | | 5 |
| mPool6_V2MM_81833 | -1.3138 | 94.45693 | TRUE | HP_30342 | TGCTGTTG | GTGGATA | NM_1729 | sorting nexin 14 | Snx14 | | 2.8 | SM2218 | f | | 12 |
| 092308m1_V2MM_134027 | -1.31181 | 2.577834 | TRUE | HP_35475 | TGCTGTTG | CAAGTAA | XM_48617 | LOC434363 | | | 2.6 | SM2104 | d | | 11 |
| 092308m3_V2MM_91675 | -1.31032 | 70.87591 | TRUE | HP_31304 | TGCTGTTG | CTGGGTA | NM_1784 | RIKEN cD 5330426L24 gene | 5330426L24Rik | | 2.9 | SM2262 | g | | 5 |
| 092308m3_V2MM_91675 | -1.31032 | 70.87591 | TRUE | HP_31304 | TGCTGTTG | CTGGGTA | NM_1784 | RIKEN cDNA 5330426L24 gene | 5330426L24Rik | | 2.9 | SM2262 | g | | 5 |
| mPool7_V2MM_184309 | -1.30813 | 2.577834 | TRUE | HP_40478 | TGCTGTTG | GCTATCAC | XM_28917 | Mus musculus LOC332799 (LOC332799), mRNA. | | | 2.5 | SM2077 | a | | 5 |
| mPool7_V2MM_184309 | -1.30813 | 2.577834 | TRUE | HP_40478 | TGCTGTTG | GCTATCAC | XM_28917 | Mus musculus LOC332799 (LOC332799), mR. | | | 2.5 | SM2077 | a | | 5 |
| 092308m3_V2MM_194627 | -1.30799 | 83.62456 | TRUE | HP_41278 | TGCTGTTG | CTCAGTGT | NM_1753 | RIKEN cD C630002B14 gene | C630002B14Rik | | 2.12 | SM2449 | c | | 9 |
| mPool4_V2MM_262180 | -1.30529 | 83.62456 | TRUE | HP_52396 | TGCTGTTG | CATACCAT | NM_1756 | RIKEN cDNA A730027B03 gene | A730027B03Rik | | 2.16 | SM2617 | d | | 5 |
| mPool4_V2MM_178783 | -1.30516 | 61.94057 | TRUE | HP_39925 | TGCTGTTG | GGCACTA | XM_28821 | Mus musculus LOC332488 (LOC332488), mRNA. | | NA | | | | NA | |
| mPool6_V2MM_162762 | -1.30423 | 61.94057 | TRUE | HP_38332 | TGCTGTTG | CAACATGA | XM_48621 | similar to PIP-1 protein precursor | | | 2.8 | SM2210 | a | | 6 |
| mPool4_V2MM_174556 | -1.30319 | 92.96229 | TRUE | HP_39504 | TGCTGTTG | CATATTCC | XM_28725 | Mus musculus hypothetical gene supported by AK084244 (LOC329673), mRNA. | | NA | | | | NA | |
| mPool4_V2MM_126785 | -1.30246 | 83.62456 | TRUE | HP_34762 | TGCTGTTG | GTCACTCC | XM_1448 | similar to immunoglobulin light chain variable region | | | 2.14 | SM2545 | f | | 1 |
| mPool4_V2MM_197560 | -1.30223 | 92.61223 | TRUE | HP_41562 | TGCTGTTG | CTCATTTA | NM_2071 | olfactory receptor 117 | Olfr117 | | 2.15 | SM2575 | a | | 12 |
| 092308m3_V2MM_83906 | -1.3021 | 61.94057 | TRUE | HP_30542 | TGCTGTTG | CACTATTT | NM_0261 | RIKEN cD 1300007F04 gene | 1300007F04Rik | | 2.13 | SM2464 | c | | 2 |
| 092308m3_V2MM_54623 | -1.3016 | 81.56557 | TRUE | HP_27694 | TGCTGTTG | CCAACATT | NM_1468 | olfactory receptor 147 | Olfr147 | | 2.12 | SM2446 | b | | 8 |
| 092308m1_V2MM_157571 | -1.29801 | 3.004939 | TRUE | HP_37818 | TGCTGTTG | GCCTTCGA | AY244804 | synovial sarcoma, X member A, breakpoint 1 | Ssxa1 | | 2.6 | SM2118 | f | | 8 |
| mPool4_V2MM_58227 | -1.29753 | 84.31789 | TRUE | HP_28045 | TGCTGTTG | GAGATAC | NM_1385 | G elongation factor | Gfm | | 2.16 | SM2603 | a | | 12 |
| mPool7_V2MM_185950 | -1.29746 | 2.577834 | TRUE | HP_40642 | TGCTGTTG | CACCCAG | XM_2894 | Mus musculus LOC333141 (LOC333141), mR. | | | 2.5 | SM2091 | a | | 11 |
| mPool7_V2MM_185950 | -1.29746 | 2.577834 | TRUE | HP_40642 | TGCTGTTG | CACCCAG | XM_2894 | Mus musculus LOC333141 (LOC333141), mRNA. | | | 2.5 | SM2091 | a | | 11 |
| mPool6_V2MM_161166 | -1.29506 | 61.94057 | TRUE | HP_38173 | TGCTGTTG | GACCTCC | AK047497 | transmembrane 9 superfamily protein member 4 | Tm9sf4 | | 2.7 | SM2197 | h | | 8 |
| mPool2_V2MM_124526 | -1.2948 | 3.004939 | TRUE | HP_34537 | TGCTGTTG | CAGTCAG | XM_1434 | Mus musculus similar to Ribonucleoside-diphosphate reductase M2 chain (Rib | | | 2.11 | SM2373 | b | | 3 |
| 092308m1_V2MM_157567 | -1.29479 | 3.004939 | TRUE | HP_37818 | TGCTGTTG | CCTTCGAC | AY244804 | synovial sarcoma, X member A, breakpoint 1 | Ssxa1 | | 2.6 | SM2121 | f | | 12 |
| mPool6_V2MM_81574 | -1.29099 | 61.94057 | TRUE | HP_30317 | TGCTGTTG | GTCTAGTA | XM_3557 | similar to Ig V-K167 precursor | | | 2.8 | SM2225 | g | | 1 |
| mPool2_V2MM_6071 | -1.29085 | 3.004939 | TRUE | HP_22961 | TGCTGTTG | CTATGAG | NM_0231 | guanine nucleotide binding protein (G protein), beta | Gnb1l | | 2.11 | SM2379 | g | | 1 |
| mPool4_V2MM_24963 | -1.28929 | 61.94057 | TRUE | HP_24804 | TGCTGTTG | GTGCATA | NM_1460 | hypothetical protein MGC7717 | | | 2.15 | SM2586 | c | | 9 |
| mPool4_V2MM_166206 | -1.28687 | 61.94057 | TRUE | HP_38673 | TGCTGTTG | GAAACAT | XM_4883 | similar to glyceraldehyde-3-phosphate dehydrogenase | | NA | | | | NA | |
| mPool2_V2MM_149737 | -1.28442 | 3.004939 | TRUE | HP_37044 | TGCTGTTG | CAGCACA | XM_1651 | similar to 60S ribosomal protein L5 | | | 2.4 | SM2039 | c | | 7 |
| mPool2_V2MM_149737 | -1.28442 | 3.004939 | TRUE | HP_37044 | TGCTGTTG | CAGCACA | XM_1651 | similar to 60S ribosomal protein L5 | | | 2.4 | SM2039 | c | | 7 |
| mPool4_V2MM_48544 | -1.28018 | 61.94057 | TRUE | HP_27100 | TGCTGTTG | GCACAGA | XM_4854 | RIKEN cDNA A330015D16 gene | A330015D16Rik | | 2.16 | SM2610 | g | | 4 |
| 092308m3_V2MM_207901 | -1.27975 | 92.61223 | TRUE | HP_42569 | TGCTGTTG | CGGGACT | BC075693 | RIKEN cD 1190002H09 gene | 1190002H09Rik | | 2.14 | SM2524 | c | | 1 |
| mPool4_V2MM_167810 | -1.27811 | 73.80895 | TRUE | HP_38832 | TGCTGTTG | CAAAGCA | XM_28584 | Mus musculus similar to MAP2 RNA trans-acting protein MARTA1 [Rattus norve | | NA | | | | NA | |
| mPool4_V2MM_129056 | -1.27586 | 70.87591 | TRUE | HP_34986 | TGCTGTTG | CTGTCCTC | XM_14611 | Mus musculus LOC211996 (LOC211996), mRNA. | | | 2.16 | SM2635 | f | | 1 |
| mPool4_V2MM_109499 | -1.27258 | 61.94057 | TRUE | HP_33055 | TGCTGTTG | CGCTAAA | XM_13391 | leucine rich repeat containing 27 | Lrrc27 | | 2.16 | SM2620 | a | | 5 |
| mPool4_V2MM_2614 | -1.27149 | 83.62456 | TRUE | HP_22622 | TGCTGTTG | CTGATCCT | AY462059 | kininogen 2 | Kng2 | | 2.15 | SM2564 | c | | 9 |
| 092308m3_V2MM_226824 | -1.26976 | 91.71214 | TRUE | HP_47883 | TGCTGTTG | CTGAAGA | NM_0082 | heat shock factor 1 | Hsf1 | | 2.13 | SM2469 | f | | 6 |
| 092308m1_V2MM_175115 | -1.26835 | 3.004939 | TRUE | HP_39560 | TGCTGTTG | CTTGGGC | XM_28737 | hypothetical protein 9630027E11 | | | 2.5 | SM2092 | a | | 10 |
| 092308m1_V2MM_184309 | -1.26779 | 2.577834 | TRUE | HP_40478 | TGCTGTTG | GCTATCAC | XM_28917 | Mus musculus LOC332799 (LOC332799), mRNA. | | | 2.5 | SM2077 | a | | 5 |
| 092308m1_V2MM_184309 | -1.26779 | 2.577834 | TRUE | HP_40478 | TGCTGTTG | GCTATCAC | XM_28917 | Mus musculus LOC332799 (LOC332799), mR. | | | 2.5 | SM2077 | a | | 5 |
| 092308m3_V2MM_142716 | -1.26747 | 70.87591 | TRUE | HP_36342 | TGCTGTTG | GTGTAGA | XM_1578 | Mus musculus LOC240627 (LOC240627), mR. | | | 2.14 | SM2528 | e | | 12 |

| | | | | | | | | | | | | | | | |
|---|---|---|---|---|---|---|---|---|---|---|---|---|---|---|---|
| mPool4_V2MM_35512 | -1.26449 | 70.87591 | TRUE | HP_25830 | TGCTGTTG | CGTTGTTC | NM_1460 | RIKEN cDNA 5730446C15 gene | 5730446C15Rik | | 2.15 | SM2593 | c | | 4 |
| 092308m1_V2MM_178676 | -1.26448 | 3.004939 | TRUE | HP_39915 | TGCTGTTG | GAGCACA | XM_2882 | Mus musculus LOC332471 (LOC332471), mR. | | | 2.5 | SM2084 | c | | 6 |
| mPool7_V2MM_68690 | -1.26221 | 3.004939 | TRUE | HP_29064 | TGCTGTTG | GACTGTA | NM_0531 | protocadherin beta 19 | Pcdhb19 | | 2.8 | SM2241 | a | | 8 |
| 092308m3_V2MM_235893 | -1.25921 | 61.94057 | TRUE | HP_48775 | TGCTGTTG | CGGCAAG | XM_15907 | Mus musculus LOC241849 (LOC241849), mR. | | | 2.14 | SM2522 | c | | 6 |
| mPool5_V2MM_161755 | -1.25886 | 61.94057 | TRUE | HP_38231 | TGCTGTTG | GGTACAG | XM_2840 | gene model 1369, (NCBI) | Gm1369 | NA | | | | NA | |
| mPool4_V2MM_4938 | -1.25854 | 79.16495 | TRUE | HP_22850 | TGCTGTTG | CAGCTCA | NM_0259 | RIKEN cDNA 1600013P15 gene | 1600013P15Rik | | 2.16 | SM2602 | f | | 2 |
| mPool2_V2MM_88981 | -1.25801 | 3.004939 | TRUE | HP_31041 | TGCTGTTG | GGCGAAA | AK048755 | zinc finger protein 291 | Zfp291 | | 2.6 | SM2109 | f | | 6 |
| mPool2_V2MM_88981 | -1.25801 | 3.004939 | TRUE | HP_31041 | TGCTGTTG | GGCGAAA | AK048755 | zinc finger protein 291 | Zfp291 | | 2.6 | SM2109 | f | | 6 |
| 092308m1_V2MM_134469 | -1.25749 | 3.004939 | TRUE | HP_35519 | TGCTGTTG | CACCTGG | XM_1498 | Mus musculus LOC232948 (LOC232948), mR. | | | 2.6 | SM2111 | c | | 4 |
| 092308m1_V2MM_134469 | -1.25749 | 3.004939 | TRUE | HP_35519 | TGCTGTTG | CACCTGG | XM_1498 | Mus musculus LOC232948 (LOC232948), mRNA. | | | 2.6 | SM2111 | c | | 4 |
| mPool4_V2MM_158562 | -1.25548 | 73.80895 | TRUE | HP_37917 | TGCTGTTG | CCCATGG | XM_2067 | Mus musculus LOC279190 (LOC279190), mRNA. | | NA | | | | NA | |
| 092308m3_V2MM_25777 | -1.25536 | 94.45693 | TRUE | HP_24883 | TGCTGTTG | CCACTTCT | NM_1454 | solute carrier family 10 (sodium/bile acid cotranspo | Slc10a3 | | 2.12 | SM2430 | a | | 11 |
| mPool4_V2MM_34194 | -1.25299 | 91.10404 | TRUE | HP_25702 | TGCTGTTG | CCAGTATA | XM_1381 | Mus musculus similar to phosphoglycerate kinase 1 [Mus musculus] (LOC2177 | | | 2.16 | SM2606 | g | | 6 |
| 092308m1_V2MM_159114 | -1.25213 | 3.004939 | TRUE | NA | NA | NA | NA | NA | NA | NA | | NA | NA | NA | |
| 092308m3_V2MM_238159 | -1.25136 | 61.94057 | TRUE | HP_48999 | TGCTGTTG | CAGACAG | XM_48891 | RIKEN cD 4930532M18 gene | 4930532M18Rik | | 2.14 | SM2528 | b | | 2 |
| mPool6_V2MM_222819 | -1.24963 | 61.94057 | TRUE | HP_47504 | TGCTGTTG | GAGATCA | XM_1955 | CDK5 regulatory subunit associated protein 2 | Cdk5rap2 | | 2.8 | SM2208 | d | | 6 |
| mPool4_V2MM_47189 | -1.24858 | 94.45693 | TRUE | HP_26967 | TGCTGTTG | CTCATTCA | NM_0323 | plasmalemma vesicle associated protein | Plvap | | 2.15 | SM2580 | f | | 10 |
| 092308m3_V2MM_225052 | -1.24683 | 94.45693 | TRUE | HP_47710 | TGCTGTTG | CACTCAAT | NM_1466 | olfactory receptor 305 | Olfr305 | | 2.13 | SM2470 | b | | 11 |
| mPool4_V2MM_141668 | -1.24671 | 94.45693 | TRUE | HP_36238 | TGCTGTTG | GTTTGGA | XM_1568 | Mus musculus similar to pol protein [Phascolarctos cinereus] (LOC215430), mR | | | 2.1 | SM2343 | g | | 1 |
| mPool6_V2MM_223433 | -1.24335 | 91.10404 | TRUE | HP_47559 | TGCTGTTG | CGCTTTCT | AK122512 | RIKEN cDNA D630023B12 gene | D630023B12Rik | | 2.8 | SM2211 | d | | 6 |
| 092308m3_V2MM_136886 | -1.23916 | 94.45693 | TRUE | HP_35760 | TGCTGTTG | CTCCTCTC | XM_1522 | Mus musculus LOC237477 (LOC237477), mR. | | | 2.14 | SM2524 | b | | 11 |
| mPool6_V2MM_31499 | -1.2387 | 92.96229 | TRUE | HP_25440 | TGCTGTTG | CTGTCTCC | NM_1455 | dual-specificity tyrosine-(Y)-phosphorylation regulat | Dyrk3 | | 2.7 | SM2184 | a | | 11 |
| mPool4_V2MM_127985 | -1.23817 | 84.31789 | TRUE | HP_34880 | TGCTGTTG | CTGATGTC | XM_14547 | Mus musculus similar to endogenous retroviral family W, env(C7), member 1 (s | | | 2.11 | SM2356 | d | | 12 |
| mPool2_V2MM_148226 | -1.23617 | 3.004939 | TRUE | HP_36893 | TGCTGTTG | GATAGCA | XM_16361 | Mus musculus LOC210679 (LOC210679), mRNA. | | | 2.4 | SM2037 | h | | 5 |
| mPool2_V2MM_148226 | -1.23617 | 3.004939 | TRUE | HP_36893 | TGCTGTTG | GATAGCA | XM_16361 | Mus musculus LOC210679 (LOC210679), mR. | | | 2.4 | SM2037 | h | | 5 |
| mPool6_V2MM_70875 | -1.23603 | 84.31789 | TRUE | HP_29277 | TGCTGTTG | CAGCCTCT | NM_0111 | protease, serine, 18 | Prss18 | | 2.8 | SM2220 | a | | 2 |
| mPool4_V2MM_262024 | -1.23592 | 61.94057 | TRUE | HP_52238 | TGCTGTTG | CTCTCTTC | NM_0531 | protocadherin beta 9 | Pcdhb9 | | 2.15 | SM2586 | g | | 7 |
| 092308m1_V2MM_172467 | -1.23288 | 3.004939 | TRUE | HP_39295 | TGCTGTTG | GCCTCTGT | XM_28682 | similar to hypothetical protein | | | 2.5 | SM2079 | h | | 5 |
| mPool4_V2MM_65023 | -1.23271 | 61.94057 | TRUE | HP_28707 | TGCTGTTG | GAGATGA | NM_0106 | keratin complex-1, acidic, gene C29 | Krt1-c29 | | 2.15 | SM2563 | a | | 10 |
| mPool2_V2MM_122154 | -1.23154 | 3.004939 | TRUE | HP_34302 | TGCTGTTG | GATGCCA | XM_1420 | Mus musculus similar to Ubiquitin-conjugating enzyme E2 C (Ubiquitin-protein | | | 2.11 | SM2365 | h | | 3 |
| mPool2_V2MM_20872 | -1.23072 | 3.004939 | TRUE | HP_24405 | TGCTGTTG | CACATTCC | NM_0087 | Notch gene homolog 1 (Drosophila) | Notch1 | | 2.7 | SM2165 | g | | 2 |
| mPool4_V2MM_240500 | -1.22852 | 73.80895 | TRUE | HP_49228 | TGCTGTTG | CATTGGC | XM_14591 | RIKEN cDNA 9230119C12 gene | 9230119C12Rik | | 2.14 | SM2542 | e | | 3 |
| mPool7_V2MM_98857 | -1.22618 | 3.004939 | TRUE | HP_32008 | TGCTGTTG | GAGAACA | XM_11317 | Mus musculus LOC195639 (LOC195639), mRNA. | | | 2.9 | SM2262 | g | | 11 |
| mPool6_V2MM_62727 | -1.22579 | 91.71214 | TRUE | HP_28482 | TGCTGTTG | CATATTAC | NM_1469 | olfactory receptor 1260 | Olfr1260 | | 2.8 | SM2222 | e | | 7 |
| mPool6_V2MM_161871 | -1.22533 | 69.72441 | TRUE | HP_38243 | TGCTGTTG | GAACTAA | AK005913 | expressed sequence AW538212 | AW538212 | | 2.7 | SM2200 | e | | 8 |
| mPool6_V2MM_79192 | -1.22349 | 61.94057 | TRUE | HP_30086 | TGCTGTTG | GATCTTCT | NM_0074 | a disintegrin and metalloprotease domain 7 | Adam7 | | 2.8 | SM2225 | g | | 7 |
| mPool6_V2MM_71365 | -1.22318 | 61.94057 | TRUE | HP_29325 | TGCTGTTG | CGATAGA | AK086463 | RIKEN cDNA D930030K17 gene | D930030K17Rik | | 2.8 | SM2222 | f | | 6 |
| mPool4_V2MM_196233 | -1.22266 | 83.62456 | TRUE | NA | NA | NA | NA | NA | NA | NA | | NA | NA | NA | |
| 092308m3_V2MM_87937 | -1.22179 | 81.56557 | TRUE | HP_30938 | TGCTGTTG | GTCTGAC | NM_1752 | formin-family protein FHOS2 | | | 2.13 | SM2468 | a | | 6 |
| 092308m3_V2MM_49774 | -1.22113 | 69.72441 | TRUE | HP_27221 | TGCTGTTG | CAAATATC | NM_0220 | PERP, TP53 apoptosis effector | Perp | | 2.12 | SM2407 | h | | 2 |
| mPool4_V2MM_261891 | -1.21976 | 73.80895 | TRUE | HP_52346 | TGCTGTTG | CTAACATG | NM_1726 | interferon activated gene 205 | Ifi205 | | 2.15 | SM2600 | f | | 12 |
| mPool6_V2MM_68880 | -1.21934 | 79.16495 | TRUE | HP_29082 | TGCTGTTG | GAGGATA | NM_1728 | hypothetical protein A030012E10 | | | 2.8 | SM2216 | h | | 6 |
| mPool6_V2MM_160030 | -1.2187 | 69.72441 | TRUE | HP_38062 | TGCTGTTG | CTCCGACT | NM_1770 | RIKEN cDNA C330018J07 gene | C330018J07Rik | | 2.8 | SM2204 | c | | 9 |
| mPool4_V2MM_199187 | -1.21864 | 83.62456 | TRUE | HP_41723 | TGCTGTTG | CCTTAAG | XM_2891 | Mus musculus LOC332829 (LOC332829), mRNA. | | NA | | | | NA | |
| mPool4_V2MM_221128 | -1.21826 | 66.95036 | TRUE | HP_45406 | TGCTGTTG | CAGTGGT | NM_1467 | olfactory receptor 1271 | Olfr1271 | | 2.16 | SM2603 | e | | 4 |
| 092308m1_V2MM_141850 | -1.2165 | 3.004939 | TRUE | HP_36256 | TGCTGTTG | GGTCTAG | XM_1570 | Mus musculus LOC209953 (LOC209953), mRNA. | | | 2.4 | SM2048 | d | | 11 |
| 092308m1_V2MM_141850 | -1.2165 | 3.004939 | TRUE | HP_36256 | TGCTGTTG | GGTCTAG | XM_1570 | Mus musculus LOC209953 (LOC209953), mR. | | | 2.4 | SM2048 | d | | 11 |
| 092308m1_V2MM_159933 | -1.21588 | 3.004939 | TRUE | HP_38052 | TGCTGTTG | CACTGAG | NM_0010 | RIKEN cD F830031D20 gene | F830031D20Rik | | 2.6 | SM2125 | d | | 10 |
| mPool4_V2MM_166290 | -1.21491 | 91.71214 | TRUE | HP_38682 | TGCTGTTG | CTGCTATT | XM_2854 | Mus musculus similar to 40S ribosomal protein SA (P40) (34/67 kDa laminin rec | | | 2.16 | SM2649 | b | | 1 |
| mPool5_V2MM_11368 | -1.21475 | 73.80895 | TRUE | HP_23478 | TGCTGTTG | GAGAAGC | NM_1451 | defensin beta 19 | Defb19 | NA | | | | NA | |
| mPool4_V2MM_164633 | -1.21413 | 70.87591 | TRUE | HP_38517 | TGCTGTTG | GATCTAA | XM_2850 | Mus musculus similar to ribosomal protein L7a, cytosolic [validated] - rat (LOC3 | | NA | | | | NA | |

| | | | | | | | | | | | | | | | |
|---|---|---|---|---|---|---|---|---|---|---|---|---|---|---|---|
| mPool7_V2MM_107175 | -1.21399 | 3.004939 | TRUE | HP_26257 | TGCTGTTG | GGCTCTCA | XM_35556 | similar to Speer1-ps1 protein | | | 2.1 | SM2317 | h | | 5 |
| mPool4_V2MM_210878 | -1.21085 | 75.98381 | TRUE | HP_42861 | TGCTGTTG | CCGGCTCA | XM_35771 | gene model 1424, (NCBI) | Gm1424 | | 2.14 | SM2540 | e | | 12 |
| mPool2_V2MM_83373 | -1.20955 | 3.004939 | TRUE | HP_30491 | TGCTGTTG | CACAACAC | NM_0135 | low density lipoprotein receptor-related protein ass | Lrpap1 | | 2.6 | SM2112 | e | | 6 |
| mPool2_V2MM_83373 | -1.20955 | 3.004939 | TRUE | HP_30491 | TGCTGTTG | CACAACAC | NM_0135 | low density lipoprotein receptor-related protein ass | Lrpap1 | | 2.6 | SM2112 | e | | 6 |
| 092308m3_V2MM_32588 | -1.2091 | 61.94057 | TRUE | HP_25546 | TGCTGTTG | GTAGTTGA | NM_0274 | pregncy-specific glycoprotein 21 | Psg21 | | 2.12 | SM2419 | f | | 5 |
| 092308m1_V2MM_179082 | -1.2067 | 3.004939 | TRUE | HP_39955 | TGCTGTTG | CGTAGGC | XM_28827 | Mus musculus LOC332544 (LOC332544), mR. | | | 2.5 | SM2079 | c | | 6 |
| mPool4_V2MM_5226 | -1.20568 | 61.94057 | TRUE | HP_22878 | TGCTGTTG | CCACTCAC | NM_1752 | methionyl aminopeptidase 1 | Metap1 | | 2.16 | SM2618 | d | | 11 |
| mPool4_V2MM_28887 | -1.20394 | 84.31789 | TRUE | HP_25186 | TGCTGTTG | GTCATGGA | NM_1448 | WD repeat domain 36 | Wdr36 | | 2.15 | SM2587 | h | | 8 |
| mPool6_V2MM_168164 | -1.20275 | 70.87591 | TRUE | HP_38867 | TGCTGTTG | CATCTAGT | XM_28592 | Mus musculus similar to hypothetical protein FLJ11159 [Homo sapiens] (LOC33 | | | 2.7 | SM2198 | a | | 11 |
| mPool4_V2MM_152299 | -1.20233 | 87.58579 | TRUE | HP_37296 | TGCTGTTG | CAGCTAG | XM_19525 | gene model 650, (NCBI) | Gm650 | NA | | | | NA | |
| 092308m3_V2MM_91443 | -1.20181 | 81.56557 | TRUE | HP_24786 | TGCTGTTG | GGTATAA | NM_1782 | membrane-bound transcription factor protease, site | Mbtps2 | | 2.13 | SM2483 | f | | 3 |
| 092308m3_V2MM_49690 | -1.20067 | 61.94057 | TRUE | HP_27212 | TGCTGTTG | GAGATAT | NM_0205 | beta-microseminoprotein | Msmb | | 2.12 | SM2407 | f | | 8 |
| 092308m1_V2MM_134470 | -1.2004 | 2.577834 | TRUE | HP_35519 | TGCTGTTG | CCACCTGC | XM_1498 | Mus musculus LOC232948 (LOC232948), mRNA. | | | 2.6 | SM2102 | d | | 5 |
| 092308m1_V2MM_134470 | -1.2004 | 2.577834 | TRUE | HP_35519 | TGCTGTTG | CCACCTGC | XM_1498 | Mus musculus LOC232948 (LOC232948), mR. | | | 2.6 | SM2102 | d | | 5 |
| 092308m3_V2MM_100520 | -1.20036 | 83.62456 | TRUE | HP_32170 | TGCTGTTG | CTGGGAT | AK014558 | RIKEN cD 4632408A20 gene | 4632408A20Rik | | 2.13 | SM2499 | a | | 2 |
| mPool4_V2MM_32586 | -1.19886 | 69.72441 | TRUE | HP_25546 | TGCTGTTG | GTACTCCT | NM_1336 | RIKEN cDNA 4930469P12 gene | 4930469P12Rik | | 2.15 | SM2591 | h | | 3 |
| mPool2_V2MM_2359 | -1.19852 | 3.004939 | TRUE | HP_22598 | TGCTGTTG | CTCGGTCA | NM_0194 | solute carrier family 2, (facilitated glucose transport | Slc2a8 | | 2.11 | SM2379 | h | | 12 |
| 092308m3_V2MM_91061 | -1.19706 | 79.16495 | TRUE | HP_31247 | TGCTGTTG | GAAAGGA | XM_4857 | similar to CD sequence BC061212 | | | 2.13 | SM2470 | a | | 1 |
| 092308m1_V2MM_144752 | -1.19677 | 2.577834 | TRUE | HP_36546 | TGCTGTTG | GGTATGA | XM_16025 | Mus musculus LOC213626 (LOC213626), mR. | | | 2.4 | SM2031 | b | | 1 |
| 092308m3_V2MM_226545 | -1.19291 | 73.80895 | TRUE | HP_47856 | TGCTGTTG | CAAGCTC | NM_1776 | expressed sequence C80587 | C80587 | | 2.13 | SM2461 | b | | 1 |
| mPool6_V2MM_223607 | -1.19238 | 73.80895 | TRUE | HP_47574 | TGCTGTTG | GCTGCAG | XM_28522 | Mus musculus similar to KRAB-zinc finger protein KZF-1 [Rattus norvegicus] (LO | | | 2.8 | SM2210 | h | | 10 |
| mPool4_V2MM_28545 | -1.19183 | 61.94057 | TRUE | HP_25153 | TGCTGTTG | GCTCAACA | NM_1447 | jumonji domain containing 2C | Jmjd2c | | 2.15 | SM2600 | h | | 7 |
| mPool2_V2MM_130611 | -1.19111 | 3.004939 | TRUE | HP_35139 | TGCTGTTG | GACATCCC | XM_14695 | Mus musculus similar to groucho-related gene 1 protein (LOC235489), mRNA. | | | 2.11 | SM2362 | b | | 7 |
| mPool4_V2MM_162153 | -1.19069 | 94.45693 | TRUE | HP_38271 | TGCTGTTG | GCCATGGA | XM_28422 | gene model 765, (NCBI) | Gm765 | NA | | | | NA | |
| mPool4_V2MM_134010 | -1.1903 | 61.94057 | TRUE | HP_35474 | TGCTGTTG | CCTCTTCT | AK017143 | RIKEN cDNA 5031425E22 gene | 5031425E22Rik | | 2.14 | SM2531 | c | | 2 |
| mPool4_V2MM_31930 | -1.19029 | 61.94057 | TRUE | HP_25481 | TGCTGTTG | GAGGAGC | NM_1724 | RIKEN cDNA B230206N24 gene | B230206N24Rik | | 2.15 | SM2590 | h | | 9 |
| 092308m1_V2MM_136268 | -1.19008 | 3.004939 | TRUE | HP_35698 | TGCTGTTG | CAAAGAG | XM_1515 | Mus musculus LOC213852 (LOC213852), mRNA. | | | 2.4 | SM2030 | g | | 11 |
| 092308m1_V2MM_136268 | -1.19008 | 3.004939 | TRUE | HP_35698 | TGCTGTTG | CAAAGAG | XM_1515 | Mus musculus LOC213852 (LOC213852), mR. | | | 2.4 | SM2030 | g | | 11 |
| mPool4_V2MM_28035 | -1.1896 | 61.94057 | TRUE | HP_25103 | TGCTGTTG | GCAAAGA | NM_0079 | esterase 1 | Es1 | | 2.15 | SM2569 | a | | 8 |
| 092308m3_V2MM_137426 | -1.18904 | 61.94057 | TRUE | HP_35814 | TGCTGTTG | GTCCATAT | XM_1531 | Mus musculus LOC238089 (LOC238089), mR. | | | 2.14 | SM2520 | h | | 8 |
| 092308m3_V2MM_136806 | -1.18873 | 83.62456 | TRUE | HP_35752 | TGCTGTTG | GCTACTCA | XM_1521 | Mus musculus similar to small nuclear ribonucleoprotein-associated protein N | | | 2.14 | SM2528 | d | | 7 |
| 092308m3_V2MM_88671 | -1.18823 | 92.61223 | TRUE | HP_31010 | TGCTGTTG | CTCGTACA | NM_1754 | RIKEN cD A930012O16 gene | A930012O16Rik | | 2.13 | SM2470 | e | | 10 |
| mPool4_V2MM_71213 | -1.18799 | 94.45693 | TRUE | HP_29310 | TGCTGTTG | CATGCAA | NM_0097 | bombesin-like receptor 3 | Brs3 | | 2.15 | SM2569 | a | | 10 |
| mPool4_V2MM_202241 | -1.18786 | 73.80895 | TRUE | HP_42025 | TGCTGTTG | CAGCTAG | XM_1362 | gene model 1302, (NCBI) | Gm1302 | | 2.16 | SM2620 | f | | 5 |
| 092308m1_V2MM_95833 | -1.18777 | 3.004939 | TRUE | HP_31710 | TGCTGTTG | CAAATGG | BC059847 | Rap guanine nucleotide exchange factor (GEF) 6 | Rapgef6 | | 2.6 | SM2112 | f | | 10 |
| mPool4_V2MM_170243 | -1.18719 | 73.80895 | TRUE | HP_39073 | TGCTGTTG | CTCAGCTT | XM_28641 | Mus musculus hypothetical gene supported by AK044531 (LOC328072), mRNA. | | NA | | | | NA | |
| 092308m1_V2MM_71080 | -1.18607 | 3.004939 | TRUE | HP_29297 | TGCTGTTG | CAGTGTC | NM_1734 | nuclear receptor interacting protein 1 | Nrip1 | | 2.4 | SM2026 | h | | 2 |
| mPool4_V2MM_240135 | -1.18574 | 91.10404 | TRUE | HP_49192 | TGCTGTTG | CTACCGA | XM_1426 | similar to hypothetical protein FLJ21628 | | | 2.14 | SM2533 | f | | 10 |
| mPool4_V2MM_15649 | -1.18533 | 91.71214 | TRUE | HP_23897 | TGCTGTTG | CTGATCTA | NM_0312 | chordin-like 1 | Chrdl1 | | 2.15 | SM2590 | e | | 8 |
| mPool4_V2MM_184423 | -1.18441 | 81.56557 | TRUE | HP_40489 | TGCTGTTG | GCCTTAAC | XM_2891 | Mus musculus LOC332829 (LOC332829), mRNA. | | NA | | | | NA | |
| 092308m1_V2MM_176042 | -1.18391 | 3.004939 | TRUE | HP_39652 | TGCTGTTG | CCCTGAG | AK032533 | hypothetical protein 6430584L05 | | | 2.5 | SM2074 | b | | 10 |
| 092308m3_V2MM_84819 | -1.18334 | 84.31789 | TRUE | HP_30631 | TGCTGTTG | CAGGTAG | NM_0295 | RIKEN cD 5830472M02 gene | 5830472M02Rik | | 2.13 | SM2476 | d | | 9 |
| mPool2_V2MM_88600 | -1.18329 | 3.004939 | TRUE | HP_31003 | TGCTGTTG | CTCAGATC | NM_1754 | ethanolamine kinase 2 | Etnk2 | | 2.6 | SM2137 | d | | 3 |
| mPool4_V2MM_101531 | -1.18307 | 81.56557 | TRUE | HP_90480 | TGCTGTTG | CTTCTAGA | AK034323 | sterile alpha motif domain containing 4 | Samd4 | | 2.16 | SM2624 | e | | 5 |
| mPool2_V2MM_85682 | -1.18182 | 3.004939 | TRUE | HP_30716 | TGCTGTTG | CAGCTGT | NM_1464 | olfactory receptor 934 | Olfr934 | | 2.6 | SM2143 | c | | 12 |
| mPool4_V2MM_33730 | -1.18132 | 92.96229 | TRUE | HP_25657 | TGCTGTTG | CAGTCCTA | NM_0292 | Rho GTPase activating protein 24 | Arhgap24 | | 2.15 | SM2598 | f | | 6 |
| mPool2_V2MM_93786 | -1.18098 | 3.004939 | TRUE | HP_31508 | TGCTGTTG | CCTCAACC | NM_1818 | G protein-coupled receptor 100 | Gpr100 | | 2.6 | SM2133 | f | | 6 |
| 092308m3_V2MM_37046 | -1.17923 | 91.10404 | TRUE | HP_25981 | TGCTGTTG | GAATGAT | NM_1817 | RIKEN cD 2610208M17 gene | 2610208M17Rik | | 2.12 | SM2427 | c | | 9 |
| 092308m3_V2MM_94887 | -1.17907 | 83.62456 | TRUE | HP_31617 | TGCTGTTG | CCTCCTTC | AK011447 | RIKEN cD 2610018I05 gene | 2610018I05Rik | | 2.9 | SM2268 | e | | 3 |
| 092308m3_V2MM_94887 | -1.17907 | 83.62456 | TRUE | HP_31617 | TGCTGTTG | CCTCCTTC | AK011447 | RIKEN cDNA 2610018I05 gene | 2610018I05Rik | | 2.9 | SM2268 | e | | 3 |
| mPool5_V2MM_164042 | -1.17874 | 94.45693 | TRUE | HP_38459 | TGCTGTTG | CATCCTCC | XM_2849 | Mus musculus similar to env protein [Homo sapiens] (LOC327892), mRNA. | | NA | | | | NA | |

| | | | | | | | | | | | | | | | |
|---|---|---|---|---|---|---|---|---|---|---|---|---|---|---|---|
| mPool6_V2MM_64870 | -1.17774 | 66.95036 | TRUE | HP_28691 | TGCTGTTG | GAAGTAT | NM_0113 | nuclear receptor co-repressor 1 | Ncor1 | | 2.8 | SM2222 | a | | 8 |
| 092308m3_V2MM_202196 | -1.177 | 84.31789 | TRUE | HP_42021 | TGCTGTTG | CTGGGAA | AK011120 | RIKEN cD 2510049J12 gene | 2510049J12Rik | | 2.13 | SM2491 | h | | 9 |
| 092308m3_V2MM_46348 | -1.17698 | 94.45693 | TRUE | HP_26885 | TGCTGTTG | CCATTGCT | NM_0094 | SH2 domain binding protein 1 (tetratricopeptide rep | Sh2bp1 | | 2.12 | SM2438 | f | | 12 |
| 092308m3_V2MM_204092 | -1.17682 | 61.94057 | TRUE | HP_42200 | TGCTGTTG | CCATCCAT | XM_1318 | cD sequence BC025458 | BC025458 | | 2.13 | SM2489 | g | | 11 |
| mPool6_V2MM_156258 | -1.17626 | 94.45693 | TRUE | HP_37689 | TGCTGTTG | CGCATCTC | XM_1982 | Mus musculus LOC272635 (LOC272635), mRNA. | | | 2.8 | SM2209 | f | | 10 |
| mPool6_V2MM_167461 | -1.17543 | 91.71214 | TRUE | HP_38797 | TGCTGTTG | GTACCAA | XM_2857 | Mus musculus similar to sterol carrier protein 2, liver [Mus musculus] (LOC329 | | | 2.8 | SM2211 | e | | 8 |
| mPool4_V2MM_227942 | -1.17502 | 70.87591 | TRUE | HP_47993 | TGCTGTTG | CTATTCCT | XM_2870 | Mus musculus hypothetical gene supported by BC039973 (LOC329350), mRNA. | | | 2.16 | SM2649 | b | | 11 |
| 092308m3_V2MM_110848 | -1.17278 | 94.45693 | TRUE | HP_33187 | TGCTGTTG | GAGATGC | XM_1350 | expressed sequence AV228068 | AV228068 | | 2.9 | SM2300 | e | | 2 |
| 092308m3_V2MM_110848 | -1.17278 | 94.45693 | TRUE | HP_33187 | TGCTGTTG | GAGATGC | XM_1350 | expressed sequence AV228068 | AV228068 | | 2.9 | SM2300 | e | | 2 |
| mPool4_V2MM_163041 | -1.17229 | 84.31789 | TRUE | HP_38359 | TGCTGTTG | CAAACTCT | XM_2846 | calcium channel, voltage-dependent, gamma subuni | Cacng7 | NA | | | | NA | |
| 092308m3_V2MM_195907 | -1.17176 | 94.45693 | TRUE | HP_41402 | TGCTGTTG | GCCATATC | NM_1464 | olfactory receptor 998 | Olfr998 | | 2.9 | SM2268 | g | | 5 |
| 092308m3_V2MM_195907 | -1.17176 | 94.45693 | TRUE | HP_41402 | TGCTGTTG | GCCATATC | NM_1464 | olfactory receptor 998 | Olfr998 | | 2.9 | SM2268 | g | | 5 |
| 092308m3_V2MM_141759 | -1.17157 | 91.71214 | TRUE | HP_36247 | TGCTGTTG | GTCAGCTA | XM_1569 | Mus musculus LOC225066 (LOC225066), mR. | | | 2.14 | SM2514 | a | | 9 |
| mPool4_V2MM_15548 | -1.17108 | 84.31789 | TRUE | HP_23886 | TGCTGTTG | CTGAATAT | NM_0532 | vomeronasal 1 receptor, B7 | V1rb7 | | 2.15 | SM2598 | e | | 1 |
| mPool4_V2MM_194280 | -1.17107 | 94.45693 | TRUE | HP_41245 | TGCTGTTG | GGAATTG | NM_1778 | cDNA sequence BC022651 | BC022651 | | 2.16 | SM2620 | d | | 5 |
| mPool2_V2MM_109324 | -1.17066 | 3.004939 | TRUE | HP_33038 | TGCTGTTG | GGAACTG | XM_3559 | RIKEN cDNA 1110033K02 gene | 1110033K02Rik | | 2.1 | SM2308 | g | | 10 |
| 092308m3_V2MM_203551 | -1.17024 | 83.62456 | TRUE | HP_42149 | TGCTGTTG | CAGTGTAT | AK010407 | zinc finger protein 511 | Zfp511 | | 2.13 | SM2496 | h | | 1 |
| 092308m3_V2MM_50809 | -1.17022 | 92.61223 | TRUE | HP_27321 | TGCTGTTG | CGACCCAT | NM_0010 | olfactory receptor 194 | Olfr194 | | 2.12 | SM2438 | h | | 9 |
| mPool4_V2MM_160529 | -1.16995 | 70.87591 | TRUE | HP_38110 | TGCTGTTG | GCACTTAT | AK076800 | hypothetical gene supported by AK076800; BC048609 | | NA | | | | NA | |
| mPool2_V2MM_141850 | -1.16763 | 3.004939 | TRUE | HP_36256 | TGCTGTTG | GGTCTAGC | XM_1570 | Mus musculus LOC209953 (LOC209953), mRNA. | | | 2.4 | SM2048 | d | | 11 |
| mPool2_V2MM_141850 | -1.16763 | 3.004939 | TRUE | HP_36256 | TGCTGTTG | GGTCTAGC | XM_1570 | Mus musculus LOC209953 (LOC209953), mR. | | | 2.4 | SM2048 | d | | 11 |
| mPool6_V2MM_223173 | -1.16676 | 75.98381 | TRUE | HP_47535 | TGCTGTTG | CTAACATC | XM_2846 | RIKEN cDNA 1700023I07 gene | 1700023I07Rik | | 2.7 | SM2197 | d | | 2 |
| mPool6_V2MM_63704 | -1.16646 | 92.61223 | TRUE | HP_28577 | TGCTGTTG | CTATTTCC | NM_1729 | COP9 (constitutive photomorphogenic) homolog, su | Cops7b | | 2.8 | SM2217 | d | | 10 |
| mPool6_V2MM_78301 | -1.16612 | 94.45693 | TRUE | HP_29999 | TGCTGTTG | GACATTTC | NM_0256 | ubiquitin-activating enzyme E1-domain containing 1 | Ube1dc1 | | 2.8 | SM2220 | h | | 2 |
| 092308m3_V2MM_50010 | -1.16555 | 61.94057 | TRUE | HP_27244 | TGCTGTTG | CACCATGA | NM_0292 | RIKEN cD 1700001L23 gene | 1700001L23Rik | | 2.12 | SM2411 | c | | 6 |
| mPool4_V2MM_183952 | -1.16547 | 79.16495 | TRUE | HP_40442 | TGCTGTTG | GAGTATCC | XM_2891 | Mus musculus LOC333602 (LOC333602), mRNA. | | NA | | | | NA | |
| 092308m3_V2MM_31772 | -1.16539 | 92.61223 | TRUE | HP_25466 | TGCTGTTG | GAAGTAA | NM_0274 | RIKEN cD 2610312E17 gene | 2610312E17Rik | | 2.11 | SM2400 | g | | 9 |
| mPool6_V2MM_67828 | -1.16535 | 61.94057 | TRUE | HP_28980 | TGCTGTTG | CTCCGAGT | NM_0110 | olfactory marker protein | Omp | | 2.8 | SM2225 | c | | 6 |
| 092308m3_V2MM_43199 | -1.16487 | 91.71214 | TRUE | HP_26579 | TGCTGTTG | CTATTGTT | NM_1773 | RIKEN cD A530053G22 gene | A530053G22Rik | | 2.12 | SM2442 | g | | 9 |
| mPool4_V2MM_186155 | -1.16405 | 91.10404 | TRUE | HP_40662 | TGCTGTTG | CTCATCAA | XM_2894 | Mus musculus LOC333197 (LOC333197), mRNA. | | NA | | | | NA | |
| 092308m3_V2MM_56273 | -1.16338 | 94.45693 | TRUE | HP_27855 | TGCTGTTG | CTCACCTG | NM_1725 | expressed sequence AU067744 | AU067744 | | 2.12 | SM2441 | a | | 6 |
| mPool6_V2MM_62092 | -1.1631 | 84.31789 | TRUE | HP_28420 | TGCTGTTG | CACCTTGA | NM_0255 | RIKEN cDNA 2310005N03 gene | 2310005N03Rik | | 2.8 | SM2220 | a | | 1 |
| 092308m3_V2MM_30944 | -1.16256 | 83.62456 | TRUE | HP_25385 | TGCTGTTG | CTCCATTA | BC038377 | RIKEN cD 4933426L22 gene | 4933426L22Rik | | 2.12 | SM2422 | d | | 6 |
| mPool4_V2MM_206630 | -1.16233 | 61.94057 | TRUE | HP_42445 | TGCTGTTG | CATTTCGC | XM_1551 | Mus musculus LOC239254 (LOC239254), mRNA. | | | 2.16 | SM2645 | h | | 5 |
| mPool4_V2MM_47886 | -1.15966 | 83.62456 | TRUE | HP_27035 | TGCTGTTG | GAAGTAA | NM_1385 | DNA segment, Chr 6, Wayne State University 176, ex | D6Wsu176e | | 2.15 | SM2588 | b | | 11 |
| mPool4_V2MM_65957 | -1.15959 | 61.94057 | TRUE | HP_28798 | TGCTGTTG | GTCTCAGT | NM_0262 | solute carrier family 25 (mitochondrial carrier, phos | Slc25a26 | | 2.15 | SM2594 | g | | 8 |
| mPool4_V2MM_210432 | -1.15913 | 92.61223 | TRUE | HP_42818 | TGCTGTTG | CACCCTGA | XM_4878 | similar to Heat shock cognate 71 kDa protein | | | 2.14 | SM2538 | a | | 7 |
| mPool6_V2MM_163262 | -1.15683 | 61.94057 | TRUE | HP_38381 | TGCTGTTG | GAGACCA | AK016974 | RIKEN cDNA 4933428M09 gene | 4933428M09Rik | | 2.7 | SM2190 | h | | 2 |
| 092308m3_V2MM_42626 | -1.15605 | 75.98381 | TRUE | HP_26523 | TGCTGTTG | CCTTCTAT | NM_1726 | F11 receptor | F11r | | 2.12 | SM2407 | f | | 6 |
| mPool6_V2MM_161264 | -1.15437 | 92.61223 | TRUE | HP_38183 | TGCTGTTG | CAGTTATA | XM_2838 | TRAF2 and NCK interacting kinase | Tnik | | 2.7 | SM2193 | b | | 1 |
| mPool4_V2MM_32090 | -1.15413 | 92.61223 | TRUE | HP_25497 | TGCTGTTG | GCCACTGC | NM_1724 | ornithine aminotransferase-like 1 | Oatl1 | NA | | | | NA | |
| 092308m1_V2MM_137990 | -1.15407 | 3.004939 | TRUE | HP_35871 | TGCTGTTG | CTGTGTGC | XM_1537 | Mus musculus LOC238452 (LOC238452), mR. | | | 2.4 | SM2049 | e | | 5 |
| 092308m1_V2MM_137990 | -1.15407 | 3.004939 | TRUE | HP_35871 | TGCTGTTG | CTGTGTGC | XM_1537 | Mus musculus LOC238452 (LOC238452), mRNA. | | | 2.4 | SM2049 | e | | 5 |
| mPool4_V2MM_10538 | -1.15341 | 61.94057 | TRUE | HP_23396 | TGCTGTTG | CTCTGGAT | NM_0272 | mitochondrial ribosomal protein L12 | Mrpl12 | | 2.15 | SM2592 | c | | 10 |
| mPool2_V2MM_124343 | -1.15137 | 3.004939 | TRUE | HP_34519 | TGCTGTTG | CATCTGG | XM_1433 | Mus musculus similar to ribosomal protein L13; 60S ribosomal protein L13; bre | | | 2.11 | SM2372 | g | | 2 |
| mPool4_V2MM_61465 | -1.15076 | 84.31789 | TRUE | HP_28359 | TGCTGTTG | GTGTTTAA | NM_1726 | cDNA sequence BC021523 | BC021523 | | 2.15 | SM2587 | h | | 10 |
| mPool4_V2MM_202335 | -1.15048 | 91.71214 | TRUE | HP_42034 | TGCTGTTG | GGCTTCTA | XM_1389 | potassium channel, subfamily K, member 16 | Kcnk16 | | 2.16 | SM2614 | b | | 6 |
| mPool6_V2MM_67581 | -1.14969 | 70.87591 | TRUE | HP_28956 | TGCTGTTG | CGGCCTTT | NM_0530 | CD163 antigen | Cd163 | | 2.8 | SM2223 | b | | 3 |
| mPool4_V2MM_167754 | -1.14947 | 70.87591 | TRUE | HP_38826 | TGCTGTTG | CCTATATG | XM_2858 | Mus musculus similar to cyclase-associated protein homologue [Rattus norvegi | | NA | | | | NA | |
| mPool6_V2MM_65889 | -1.14862 | 94.45693 | TRUE | HP_28791 | TGCTGTTG | GTCACATC | NM_0139 | ubiquitin specific protease 25 | Usp25 | | 2.7 | SM2176 | b | | 7 |
| mPool7_V2MM_194256 | -1.14707 | 3.004939 | TRUE | HP_41243 | TGCTGTTG | CGTTACAC | NM_1786 | F-box and leucine-rich repeat protein 21 | Fbxl21 | | 2.9 | SM2253 | g | | 10 |

| | | | | | | | | | | | | | |
|---|---|---|---|---|---|---|---|---|---|---|---|---|---|
| mPool4_V2MM_109076 | -1.14579 | 61.94057 | TRUE | HP_33014 | TGCTGTTG | CAGATAG | NM_0074 | amyloid beta (A4) precursor protein-binding, family | Apba2 | 2.16 | SM2617 | a | 11 |
| mPool4_V2MM_216973 | -1.14533 | 84.31789 | TRUE | HP_45007 | TGCTGTTG | CTATCTAC | NM_0101 | exostoses (multiple) 1 | Ext1 | 2.15 | SM2575 | a | 1 |
| 092308m1_V2MM_149737 | -1.14337 | 3.004939 | TRUE | HP_37044 | TGCTGTTG | CAGCACA | XM_1651 | similar to 60S ribosomal protein L5 | | 2.4 | SM2039 | c | 7 |
| 092308m1_V2MM_149737 | -1.14337 | 3.004939 | TRUE | HP_37044 | TGCTGTTG | CAGCACA | XM_1651 | similar to 60S ribosomal protein L5 | | 2.4 | SM2039 | c | 7 |
| 092308m1_V2MM_145550 | -1.14249 | 2.577834 | TRUE | HP_36626 | TGCTGTTG | CAATGTA | XM_1610 | Mus musculus LOC230170 (LOC230170), mRNA. | | 2.4 | SM2047 | a | 8 |
| 092308m1_V2MM_145550 | -1.14249 | 2.577834 | TRUE | HP_36626 | TGCTGTTG | CAATGTA | XM_1610 | Mus musculus LOC230170 (LOC230170), mR. | | 2.4 | SM2047 | a | 8 |
| mPool7_V2MM_112924 | -1.14114 | 3.004939 | TRUE | HP_33391 | TGCTGTTG | GTGACAG | XM_13671 | Mus musculus LOC213721 (LOC213721), mRNA. | | 2.1 | SM2321 | a | 8 |
| 092308m3_V2MM_82257 | -1.14099 | 94.45693 | TRUE | HP_30383 | TGCTGTTG | CGGCCAG | NM_0077 | procollagen, type IV, alpha 5 | Col4a5 | 2.7 | SM2168 | c | 12 |
| 092308m3_V2MM_82257 | -1.14099 | 94.45693 | TRUE | HP_30383 | TGCTGTTG | CGGCCAG | NM_0077 | procollagen, type IV, alpha 5 | Col4a5 | 2.7 | SM2168 | c | 12 |
| 092308m3_V2MM_147982 | -1.14091 | 84.31789 | TRUE | HP_36869 | TGCTGTTG | CCCAAAG | XM_1633 | Mus musculus LOC244043 (LOC244043), mRNA. | | 2.1 | SM2336 | b | 9 |
| 092308m3_V2MM_147982 | -1.14091 | 84.31789 | TRUE | HP_36869 | TGCTGTTG | CCCAAAG | XM_1633 | Mus musculus LOC244043 (LOC244043), mR. | | 2.1 | SM2336 | b | 9 |
| 092308m3_V2MM_48412 | -1.14055 | 83.62456 | TRUE | HP_27087 | TGCTGTTG | GATCACA | NM_0290 | RIKEN cD 4933428M03 gene | 4933428M03Rik | 2.12 | SM2445 | f | 4 |
| mPool2_V2MM_212554 | -1.13935 | 3.004939 | TRUE | HP_43022 | TGCTGTTG | GGTCCCA | XM_1463 | similar to 40S ribosomal protein S7 (S8) | | 2.1 | SM2349 | e | 6 |
| mPool6_V2MM_169113 | -1.13831 | 83.62456 | TRUE | HP_38961 | TGCTGTTG | CTCAGTTA | XM_2861 | similar to High mobility group protein 2 (HMG-2) | | 2.7 | SM2192 | d | 4 |
| mPool2_V2MM_120454 | -1.13675 | 3.004939 | TRUE | HP_34135 | TGCTGTTG | GTAAACA | XM_3571 | similar to basic transcription factor 3 | | 2.11 | SM2353 | h | 1 |
| 092308m3_V2MM_45381 | -1.13625 | 94.45693 | TRUE | HP_26791 | TGCTGTTG | GTCGCAT | NM_0206 | TATA box binding protein (Tbp)-associated factor, R | Taf1b | 2.12 | SM2410 | f | 6 |
| mPool4_V2MM_2704 | -1.13459 | 91.71214 | TRUE | HP_22631 | TGCTGTTG | CTGCTATT | NM_1530 | zinc finger protein 472 | Zfp472 | 2.15 | SM2598 | g | 12 |
| mPool6_V2MM_168290 | -1.13311 | 83.62456 | TRUE | HP_38879 | TGCTGTTG | GAGAGAT | XM_2859 | Mus musculus similar to interferon gamma inducible protein 30; lysosomal thic | | 2.7 | SM2199 | e | 8 |
| mPool4_V2MM_242264 | -1.13198 | 61.94057 | TRUE | HP_49401 | TGCTGTTG | CCATTCTC | XM_1409 | Mus musculus similar to constitutive photomorphogenic protein 1 [Mus muscu | | 2.14 | SM2549 | f | 7 |
| mPool4_V2MM_134038 | -1.13116 | 92.96229 | TRUE | HP_35476 | TGCTGTTG | GAACTAA | NM_0010 | RIKEN cDNA E130018O15 gene | E130018O15Rik | 2.14 | SM2543 | c | 2 |
| mPool6_V2MM_223466 | -1.13099 | 91.71214 | TRUE | HP_47561 | TGCTGTTG | CTGGAGA | XM_2849 | Mus musculus similar to single-strand DNA binding protein [Rattus norvegicus] | | 2.7 | SM2197 | f | 7 |
| mPool6_V2MM_62966 | -1.13072 | 69.72441 | TRUE | HP_28505 | TGCTGTTG | CCAGTTCT | NM_0080 | UDP-N-acetyl-alpha-D-galactosamine:(N-acetylneura | Galgt1 | 2.8 | SM2214 | g | 7 |
| 092308m3_V2MM_205067 | -1.12995 | 94.45693 | TRUE | HP_42294 | TGCTGTTG | CTCTCCTC | XM_1373 | Mus musculus similar to cytochrome c oxidase subunit VIIc [Mus musculus] (LO | | 2.1 | SM2306 | h | 9 |
| 092308m3_V2MM_205067 | -1.12995 | 94.45693 | TRUE | HP_42294 | TGCTGTTG | CTCTCCTC | XM_1373 | Mus musculus similar to cytochrome c oxidase subunit VIIc [Mus musculus] (LO | | 2.1 | SM2306 | h | 9 |
| 092308m3_V2MM_225216 | -1.12851 | 61.94057 | TRUE | HP_47727 | TGCTGTTG | GAAGATA | XM_10991 | similar to hepatitis A virus cellular receptor 1; T-cell immunoglobulin and mucir | | 2.13 | SM2467 | b | 6 |
| mPool6_V2MM_157354 | -1.12674 | 73.80895 | TRUE | HP_37797 | TGCTGTTG | CTTCTACT | XM_35871 | similar to hypothetical protein FLJ36601 | | 2.7 | SM2192 | h | 3 |
| mPool4_V2MM_90738 | -1.12612 | 61.94057 | TRUE | HP_31215 | TGCTGTTG | CGGACCC | NM_1778 | hypothetical protein A630008I04 | | 2.16 | SM2610 | e | 11 |
| mPool2_V2MM_142390 | -1.12499 | 3.004939 | TRUE | HP_36310 | TGCTGTTG | GCCAGAA | XM_15761 | Mus musculus LOC212132 (LOC212132), mRNA. | | 2.5 | SM2053 | c | 11 |
| mPool2_V2MM_142390 | -1.12499 | 3.004939 | TRUE | HP_36310 | TGCTGTTG | GCCAGAA | XM_15761 | Mus musculus LOC212132 (LOC212132), mR. | | 2.5 | SM2053 | c | 11 |
| 092308m1_V2MM_98998 | -1.12496 | 3.004939 | TRUE | HP_32021 | TGCTGTTG | CTAACTGA | AK122468 | pleckstrin homology domain containing, family G (w | Plekhg1 | 2.6 | SM2112 | a | 11 |
| 092308m3_V2MM_88104 | -1.12454 | 94.45693 | TRUE | HP_30955 | TGCTGTTG | CCCTCTAC | NM_1753 | RIKEN cD 4931414L13 gene | 4931414L13Rik | 2.13 | SM2474 | g | 1 |
| mPool4_V2MM_211824 | -1.1241 | 61.94057 | TRUE | HP_42950 | TGCTGTTG | GACTCCAT | AK038838 | RIKEN cDNA A230067G21 gene | A230067G21Rik | 2.16 | SM2640 | d | 2 |
| mPool4_V2MM_8790 | -1.12372 | 94.45693 | TRUE | HP_23227 | TGCTGTTG | CAAAGAA | XM_1421 | Mus musculus similar to argonaute 5 protein [Mus musculus] (LOC245644), mR | | 2.15 | SM2586 | c | 4 |
| mPool6_V2MM_64724 | -1.12339 | 66.95036 | TRUE | HP_28677 | TGCTGTTG | CTTGACCT | NM_1734 | regulator of G-protein signaling 12 | Rgs12 | 2.8 | SM2223 | c | 1 |
| mPool5_V2MM_180307 | -1.12306 | 92.61223 | TRUE | HP_40078 | TGCTGTTG | CACTCGA | XM_2884 | Mus musculus LOC331838 (LOC331838), mRNA. | | NA | | | NA |
| mPool4_V2MM_125223 | -1.12185 | 79.16495 | TRUE | HP_34606 | TGCTGTTG | CTAATTGT | XM_1438 | similar to Elongation factor 1-alpha 1 (EF-1-alpha-1) (Elongation factor 1 A-1) (e | | 2.14 | SM2549 | g | 1 |
| 092308m1_V2MM_169056 | -1.12162 | 3.004939 | TRUE | HP_38955 | TGCTGTTG | CCTTCCTG | XM_2861 | RIKEN cD 8430439C15 gene | 8430439C15Rik | 2.6 | SM2119 | a | 1 |
| mPool4_V2MM_193877 | -1.12159 | 61.94057 | TRUE | HP_41213 | TGCTGTTG | CTCATGTG | XM_3553 | gene model 998, (NCBI) | Gm998 | NA | | | NA |
| mPool4_V2MM_238613 | -1.12121 | 84.31789 | TRUE | HP_49044 | TGCTGTTG | CATATACA | NM_0010 | solute carrier family 5 (sodium/glucose cotransporte | Slc5a12 | 2.14 | SM2538 | f | 5 |
| 092308m3_V2MM_47832 | -1.12101 | 61.94057 | TRUE | HP_27030 | TGCTGTTG | GAAATTG | NM_0166 | programmed cell death protein 7 | Pdcd7 | 2.12 | SM2446 | f | 6 |
| mPool2_V2MM_93782 | -1.11898 | 3.004939 | TRUE | HP_31508 | TGCTGTTG | GACAGCC | NM_1818 | RIKEN cDNA 4933401K09 gene | 4933401K09Rik | 2.6 | SM2140 | c | 1 |
| mPool2_V2MM_210543 | -1.11884 | 3.004939 | TRUE | HP_42829 | TGCTGTTG | CCCAGATT | AK017962 | RIKEN cDNA 5830432E09 gene | 5830432E09Rik | 2.11 | SM2358 | d | 2 |
| mPool4_V2MM_121327 | -1.11739 | 91.10404 | TRUE | HP_34220 | TGCTGTTG | GAAGAAA | XM_1416 | Mus musculus similar to glyceraldehyde-3-phosphate dehydrogenase [Merione | | 2.16 | SM2648 | c | 12 |
| mPool4_V2MM_125806 | -1.11644 | 91.10404 | TRUE | HP_34664 | TGCTGTTG | CTAAGAC | XM_1442 | Mus musculus similar to protease [Mus musculus] (LOC211321), mRNA. | | 2.14 | SM2543 | f | 3 |
| 092308m3_V2MM_84004 | -1.11643 | 94.45693 | TRUE | HP_30552 | TGCTGTTG | CACGCCT | AK031571 | cell division cycle associated 8 | Cdca8 | 2.13 | SM2471 | h | 12 |
| 092308m3_V2MM_54737 | -1.11625 | 91.71214 | TRUE | HP_27705 | TGCTGTTG | CCACTTAT | XM_4857 | similar to immunoglobulin light chain variable region | | 2.12 | SM2413 | b | 3 |
| mPool4_V2MM_122450 | -1.11608 | 84.31789 | TRUE | HP_34331 | TGCTGTTG | GGTATAG | XM_1421 | Mus musculus similar to actin (LOC209541), mRNA. | | 2.11 | SM2373 | g | 6 |
| mPool6_V2MM_166975 | -1.11602 | 91.10404 | TRUE | HP_38749 | TGCTGTTG | GTGATAC | XM_2856 | Mus musculus similar to hypothetical protein FLJ10891 [Homo sapiens] (LOC33 | | 2.7 | SM2190 | f | 12 |
| 092308m3_V2MM_106827 | -1.11556 | 61.94057 | TRUE | HP_32792 | TGCTGTTG | GAGGAAA | XM_1315 | hypothetical protein LOC230595 | | 2.14 | SM2503 | g | 7 |
| mPool4_V2MM_206616 | -1.11493 | 75.98381 | TRUE | HP_42443 | TGCTGTTG | GAGCTGC | XM_1636 | Mus musculus LOC209415 (LOC209415), mRNA. | | 2.16 | SM2647 | a | 5 |
| mPool4_V2MM_123202 | -1.11482 | 91.10404 | TRUE | HP_34405 | TGCTGTTG | CTGTTGG | XM_4882 | similar to hypothetical protein FLJ38281 | | 2.15 | SM2551 | b | 5 |

| | | | | | | | | | | | | | | | |
|---|---|---|---|---|---|---|---|---|---|---|---|---|---|---|---|
| mPool6_V2MM_160295 | -1.11426 | 91.10404 | TRUE | HP_38087 | TGCTGTTG | GATCTAAA | XM_48463 | hypothetical gene supported by AK089217 | | | 2.7 | SM2188 | b | | 12 |
| mPool4_V2MM_55261 | -1.1132 | 66.95036 | TRUE | HP_27756 | TGCTGTTG | CCCTTTCCT | NM_14630 | olfactory receptor 420 | Olfr420 | | 2.15 | SM2600 | h | | 12 |
| mPool4_V2MM_26347 | -1.11252 | 94.45693 | TRUE | HP_24939 | TGCTGTTG | CCTTTGCT | NM_14620 | two pore segment channel 2 | Tpcn2 | NA | | | | NA | |
| mPool4_V2MM_167738 | -1.11179 | 91.10404 | TRUE | HP_38825 | TGCTGTTG | CTTTCCAG | XM_28582 | similar to RIKEN cDNA 4732496O08 | | NA | | | | NA | |
| mPool4_V2MM_57154 | -1.11173 | 94.45693 | TRUE | HP_27941 | TGCTGTTG | CTGTGTTT | BC056964 | RIKEN cDNA 4930553M18 gene | 4930553M18Rik | | 2.15 | SM2598 | a | | 1 |
| mPool4_V2MM_133428 | -1.1113 | 61.94057 | TRUE | HP_35416 | TGCTGTTG | GAATATTT | BC072638 | RIKEN cDNA E130013N09 gene | E130013N09Rik | | 2.16 | SM2637 | f | | 5 |
| mPool4_V2MM_142311 | -1.11109 | 61.94057 | TRUE | HP_36302 | TGCTGTTG | GACATGTT | XM_15756 | Mus musculus LOC211517 (LOC211517), mRNA. | | | 2.16 | SM2641 | g | | 5 |
| mPool6_V2MM_163740 | -1.11104 | 70.87591 | TRUE | HP_38429 | TGCTGTTG | CCAGGACC | XM_28484 | RIKEN cDNA 4930513F16 gene | 4930513F16Rik | | 2.7 | SM2195 | c | | 2 |
| mPool6_V2MM_161433 | -1.11076 | 92.61223 | TRUE | HP_38200 | TGCTGTTG | GAGAGATA | AK036551 | hypothetical protein 9830132P13 | | | 2.7 | SM2188 | c | | 8 |
| mPool4_V2MM_130738 | -1.11039 | 84.31789 | TRUE | HP_35151 | TGCTGTTG | CGCTCAGC | XM_14700 | Mus musculus similar to extracellular matrix protein papilin [Drosophila melan | | | 2.14 | SM2549 | h | | 9 |
| mPool4_V2MM_161281 | -1.10928 | 92.61223 | TRUE | HP_38185 | TGCTGTTG | CTGCAAAT | XM_28383 | Mus musculus hypothetical gene supported by AK043341 (LOC329611), mRNA. | | NA | | | | NA | |
| mPool4_V2MM_11751 | -1.10859 | 94.45693 | TRUE | HP_23516 | TGCTGTTG | GCTTTATA | NM_14530 | thioredoxin domain containing 5 | Txndc5 | | 2.15 | SM2581 | a | | 9 |
| mPool4_V2MM_242002 | -1.10677 | 84.31789 | TRUE | HP_49375 | TGCTGTTG | GGAATCAT | AK006834 | RIKEN cDNA 1700058M13 gene | 1700058M13Rik | | 2.14 | SM2539 | c | | 8 |
| mPool4_V2MM_60660 | -1.10668 | 91.71214 | TRUE | HP_28281 | TGCTGTTG | GGTTCATC | NM_14634 | olfactory receptor 1259 | Olfr1259 | | 2.15 | SM2588 | g | | 1 |
| mPool4_V2MM_331 | -1.10664 | 92.61223 | TRUE | HP_22400 | TGCTGTTG | CACATCAG | NM_02649 | mitochondrial ribosomal protein S11 | Mrps11 | | 2.15 | SM2592 | f | | 3 |
| mPool4_V2MM_57357 | -1.10662 | 87.58579 | TRUE | HP_27961 | TGCTGTTG | CTTGTTCC | NM_14650 | olfactory receptor 843 | Olfr843 | | 2.15 | SM2587 | b | | 11 |
| mPool4_V2MM_19223 | -1.10651 | 84.31789 | TRUE | HP_24245 | TGCTGTTG | GGCATTAT | NM_01201 | exonuclease 1 | Exo1 | | 2.15 | SM2569 | f | | 3 |
| mPool4_V2MM_165318 | -1.10604 | 61.94057 | TRUE | HP_38585 | TGCTGTTG | GAAAGAAT | XM_28522 | zinc finger protein 206 | Zfp206 | | 2.16 | SM2649 | d | | 6 |
| mPool2_V2MM_152825 | -1.10451 | 3.004939 | TRUE | HP_37348 | TGCTGTTG | GGGACCAC | XM_19557 | Mus musculus similar to large subunit ribosomal protein L36a [Rattus norvegicu | | | 2.6 | SM2126 | b | | 11 |
| 092308m3_V2MM_60764 | -1.10438 | 91.71214 | TRUE | HP_28291 | TGCTGTTG | GTAGAGAT | BC023914 | expressed sequence AI315068 | AI315068 | | 2.12 | SM2402 | d | | 10 |
| 092308m3_V2MM_106505 | -1.10434 | 94.45693 | TRUE | HP_32760 | TGCTGTTG | CTACTGAA | AK011611 | RIKEN cD 2610029I01 gene | 2610029I01Rik | | 2.13 | SM2487 | b | | 3 |
| mPool4_V2MM_124287 | -1.10327 | 61.94057 | TRUE | HP_34513 | TGCTGTTG | GCATTGTC | NM_20727 | tigger transposable element derived 4 | Tigd4 | | 2.14 | SM2539 | a | | 3 |
| mPool4_V2MM_222434 | -1.10301 | 70.87591 | TRUE | HP_47469 | TGCTGTTG | CCTGCTCC | XM_19688 | Mus musculus LOC270881 (LOC270881), mRNA. | | NA | | | | NA | |
| mPool4_V2MM_128833 | -1.10227 | 94.45693 | TRUE | HP_34964 | TGCTGTTG | CTTATGAT | XM_14593 | Mus musculus similar to Glyceraldehyde 3-phosphate dehydrogenase (GAPDH) | | | 2.16 | SM2633 | f | | 7 |
| mPool4_V2MM_230502 | -1.10114 | 61.94057 | TRUE | HP_48248 | TGCTGTTG | GGTGTACC | AK052790 | hypothetical protein D730003B17 | | | 2.16 | SM2650 | c | | 2 |
| mPool4_V2MM_262501 | -1.10086 | 91.10404 | TRUE | HP_52303 | TGCTGTTG | CCTACATC | NM_18310 | hypothetical protein 4930549O06 | | | 2.16 | SM2620 | b | | 8 |
| mPool6_V2MM_163301 | -1.10041 | 62.64426 | TRUE | HP_38385 | TGCTGTTG | CAAATTAA | AK038641 | RIKEN cDNA A230052C13 gene | A230052C13Rik | | 2.8 | SM2208 | f | | 9 |
| 092308m3_V2MM_90177 | -1.09978 | 79.16495 | TRUE | HP_31160 | TGCTGTTG | GTTATCCT | NM_17774 | killer immunoglobulin-like receptor-like 2 | | | 2.9 | SM2259 | d | | 7 |
| 092308m3_V2MM_90177 | -1.09978 | 79.16495 | TRUE | HP_31160 | TGCTGTTG | GTTATCCT | NM_17774 | killer immunoglobulin-like receptor-like 2 | | | 2.9 | SM2259 | d | | 7 |
| mPool4_V2MM_197044 | -1.09976 | 92.61223 | TRUE | HP_41513 | TGCTGTTG | GGAACATC | NM_02685 | mitochondrial ribosomal protein L52 | Mrpl52 | | 2.15 | SM2589 | b | | 10 |
| mPool2_V2MM_161333 | -1.09943 | 3.004939 | TRUE | HP_38190 | TGCTGTTG | CAAAGAAC | AK053020 | hypothetical protein D930017F01 | | | 2.6 | SM2128 | g | | 7 |
| 092308m3_V2MM_39079 | -1.09907 | 61.94057 | TRUE | HP_26177 | TGCTGTTG | GCTATCAA | NM_14602 | cD sequence BC034054 | BC034054 | | 2.11 | SM2398 | f | | 3 |
| mPool2_V2MM_147547 | -1.09889 | 3.004939 | TRUE | HP_36825 | TGCTGTTG | GAATGTTA | XM_16307 | Mus musculus LOC243797 (LOC243797), mRNA. | | | 2.1 | SM2333 | e | | 5 |
| mPool4_V2MM_48431 | -1.09881 | 75.98381 | TRUE | HP_27089 | TGCTGTTG | GATCTATA | NM_03187 | SWI/SNF related, matrix associated, actin dependen | Smarcd2 | | 2.15 | SM2588 | e | | 9 |
| mPool7_V2MM_209029 | -1.09832 | 2.577834 | TRUE | HP_42680 | TGCTGTTG | GCTCTCCA | XM_19489 | Mus musculus similar to vomeronasal 1 receptor, I3 [Mus musculus] (LOC2686 | | | 2.1 | SM2329 | f | | 10 |
| mPool7_V2MM_197460 | -1.09798 | 3.004939 | TRUE | HP_41553 | TGCTGTTG | GCCTTTCT | NM_17552 | RIKEN cDNA A930026L03 gene | A930026L03Rik | | 2.9 | SM2266 | a | | 9 |
| mPool7_V2MM_80810 | -1.0964 | 3.004939 | TRUE | HP_30243 | TGCTGTTG | GGCAAGA | NM_17530 | MOB1, Mps One Binder kinase activator-like 2C (yea | Mobkl2c | | 2.8 | SM2233 | a | | 3 |
| mPool4_V2MM_9776 | -1.09626 | 83.62456 | TRUE | HP_23322 | TGCTGTTG | CATTTATA | NM_02727 | RIKEN cDNA 2810422O20 gene | 2810422O20Rik | | 2.15 | SM2583 | f | | 11 |
| mPool7_V2MM_204674 | -1.09548 | 3.004939 | TRUE | HP_42256 | TGCTGTTG | GAGACTCA | XM_13558 | hypothetical LOC236262 | | | 2.1 | SM2317 | e | | 7 |
| mPool4_V2MM_132161 | -1.09499 | 83.62456 | TRUE | HP_35291 | TGCTGTTG | CCTCTACT | AK048159 | mitogen activated protein kinase 1 | Mapk1 | | 2.14 | SM2537 | d | | 11 |
| mPool7_V2MM_174445 | -1.09406 | 3.004939 | TRUE | HP_39493 | TGCTGTTG | GGCTAAGA | XM_48899 | hypothetical gene supported by AK040598 | | | 2.9 | SM2282 | d | | 3 |
| mPool4_V2MM_230920 | -1.09252 | 91.10404 | TRUE | HP_48290 | TGCTGTTG | CCCACTTC | XM_28738 | gene model 831, (NCBI) | Gm831 | NA | | | | NA | |
| 092308m3_V2MM_99559 | -1.09111 | 79.16495 | TRUE | HP_32075 | TGCTGTTG | CACCTCAT | NM_17877 | transmembrane protein 16D (eight membrane-span | Tmem16d | | 2.9 | SM2262 | h | | 10 |
| 092308m3_V2MM_99559 | -1.09111 | 79.16495 | TRUE | HP_32075 | TGCTGTTG | CACCTCAT | NM_17877 | transmembrane protein 16D (eight membrane-span | Tmem16d | | 2.9 | SM2262 | h | | 10 |
| 092308m3_V2MM_47138 | -1.0902 | 75.98381 | TRUE | HP_26962 | TGCTGTTG | CTCAGAAC | NM_17714 | RIKEN cD B230317F23 gene | B230317F23Rik | | 2.12 | SM2447 | a | | 1 |
| 092308m3_V2MM_88260 | -1.08921 | 94.45693 | TRUE | HP_30970 | TGCTGTTG | CTGGGTCT | NM_17535 | RIKEN cD E330013P04 gene | E330013P04Rik | | 2.13 | SM2460 | f | | 4 |
| 092308m3_V2MM_88260 | -1.08921 | 94.45693 | TRUE | HP_30970 | TGCTGTTG | CTGGGTCT | NM_17535 | RIKEN cDNA E330013P04 gene | E330013P04Rik | | 2.13 | SM2460 | f | | 4 |
| 092308m3_V2MM_224981 | -1.08908 | 62.64426 | TRUE | HP_47703 | TGCTGTTG | CCCTTCCC | NM_17860 | D segment, Chr 10, University of California at Los An | D10Ucla1 | | 2.13 | SM2477 | h | | 6 |
| mPool7_V2MM_137990 | -1.08886 | 2.577834 | TRUE | HP_35871 | TGCTGTTG | CTGTGTGC | XM_15377 | Mus musculus LOC238452 (LOC238452), mRNA. | | | 2.4 | SM2049 | e | | 5 |
| mPool7_V2MM_137990 | -1.08886 | 2.577834 | TRUE | HP_35871 | TGCTGTTG | CTGTGTGC | XM_15377 | Mus musculus LOC238452 (LOC238452), mR. | | | 2.4 | SM2049 | e | | 5 |
| 092308m3_V2MM_191422 | -1.08654 | 91.71214 | TRUE | HP_26388 | TGCTGTTG | CAAGGAGT | NM_18330 | a disintegrin and metalloprotease domain 28 | Adam28 | | 2.12 | SM2411 | a | | 11 |

| | | | | | | | | | | | | | | | |
|---|---|---|---|---|---|---|---|---|---|---|---|---|---|---|---|
| mPool2_V2MM_168105 | -1.08627 | 2.577834 | TRUE | HP_38861 | TGCTGTTG | CAAGGTC | XM_2859 | Mus musculus similar to Eukaryotic translation initiation factor 3 subunit 6 (eIF | | | 2.6 | SM2129 | c | | 3 |
| mPool7_V2MM_171565 | -1.08609 | 3.004939 | TRUE | HP_39205 | TGCTGTTG | GCATCTAC | AK041537 | hypothetical protein A630020A06 | | | 2.5 | SM2077 | e | | 4 |
| mPool7_V2MM_171565 | -1.08609 | 3.004939 | TRUE | HP_39205 | TGCTGTTG | GCATCTAC | AK041537 | hypothetical protein A630020A06 | | | 2.5 | SM2077 | e | | 4 |
| mPool6_V2MM_20758 | -1.08542 | 94.45693 | TRUE | HP_24394 | TGCTGTTG | CAATGTG | NM_0197 | expressed in non-metastatic cells 4, protein | Nme4 | | 2.7 | SM2187 | c | | 10 |
| mPool4_V2MM_241615 | -1.08507 | 92.61223 | TRUE | HP_49337 | TGCTGTTG | GAGTACC | BC052040 | cDNA sequence BC052040 | BC052040 | | 2.14 | SM2542 | e | | 1 |
| 092308m3_V2MM_82769 | -1.0848 | 94.45693 | TRUE | HP_30433 | TGCTGTTG | CTGTTCTC | NM_0103 | guanine nucleotide binding protein (G protein), gam | | 10090 | 2.13 | SM2485 | b | | 5 |
| mPool2_V2MM_82828 | -1.08445 | 3.004939 | TRUE | HP_30438 | TGCTGTTG | CAGCCCA | NM_0105 | insulin receptor substrate 3 | Irs3 | | 2.6 | SM2139 | d | | 5 |
| mPool2_V2MM_149240 | -1.08441 | 3.004939 | TRUE | HP_36995 | TGCTGTTG | GTCATCAC | XM_16467 | Mus musculus LOC209941 (LOC209941), mRNA. | | | 2.1 | SM2338 | g | | 2 |
| mPool4_V2MM_21116 | -1.08416 | 92.61223 | TRUE | HP_24429 | TGCTGTTG | CAGACAA | NM_1462 | cDNA sequence BC026374 | BC026374 | | 2.15 | SM2583 | d | | 12 |
| mPool2_V2MM_119182 | -1.08262 | 3.004939 | TRUE | HP_34010 | TGCTGTTG | CTATGTGT | XM_1402 | Mus musculus similar to 40S RIBOSOMAL PROTEIN S21 (LOC225141), mRNA. | | | 2.11 | SM2357 | a | | 8 |
| mPool4_V2MM_173106 | -1.08197 | 62.64426 | TRUE | HP_39359 | TGCTGTTG | GCATTCAC | XM_2869 | hypothetical LOC329092 | | NA | | | | NA | |
| 092308m3_V2MM_43319 | -1.08171 | 91.10404 | TRUE | HP_26590 | TGCTGTTG | CTCATCAT | NM_1463 | olfactory receptor 62 | Olfr62 | | 2.12 | SM2417 | e | | 1 |
| 092308m3_V2MM_27431 | -1.08137 | 94.45693 | TRUE | HP_25045 | TGCTGTTG | GACAAGA | NM_0092 | sigl recognition particle 14 | Srp14 | | 2.11 | SM2398 | h | | 4 |
| mPool4_V2MM_182128 | -1.08102 | 73.80895 | TRUE | HP_40260 | TGCTGTTG | GAGCTTC | XM_2887 | Mus musculus LOC332252 (LOC332252), mRNA. | | NA | | | | NA | |
| mPool6_V2MM_63825 | -1.0808 | 94.45693 | TRUE | HP_28589 | TGCTGTTG | CTCATCTA | NM_1468 | olfactory receptor 992 | Olfr992 | | 2.8 | SM2216 | f | | 2 |
| mPool7_V2MM_173489 | -1.07809 | 3.004939 | TRUE | HP_39397 | TGCTGTTG | GAGCACA | AK040646 | RIKEN cDNA E130008O17 gene | E130008O17Rik | | 2.5 | SM2072 | e | | 1 |
| mPool7_V2MM_173489 | -1.07809 | 3.004939 | TRUE | HP_39397 | TGCTGTTG | GAGCACA | AK040646 | RIKEN cD E130008O17 gene | E130008O17Rik | | 2.5 | SM2072 | e | | 1 |
| mPool4_V2MM_202311 | -1.0774 | 61.94057 | TRUE | NA | NA | NA | NA | NA | NA | NA | | NA | NA | NA | |
| mPool4_V2MM_392 | -1.07692 | 91.71214 | TRUE | HP_22406 | TGCTGTTG | CACCGAA | NM_0305 | cDNA sequence BC004044 | BC004044 | | 2.15 | SM2582 | b | | 10 |
| 092308m3_V2MM_59485 | -1.07677 | 91.71214 | TRUE | HP_28166 | TGCTGTTG | GCCTTTAG | NM_0285 | actin related protein M2 | | | 2.12 | SM2402 | c | | 8 |
| mPool6_V2MM_71135 | -1.0765 | 69.72441 | TRUE | HP_29302 | TGCTGTTG | CATATACC | NM_0113 | SEC61, gamma subunit | Sec61g | | 2.8 | SM2217 | e | | 2 |
| mPool4_V2MM_241316 | -1.07624 | 61.94057 | TRUE | HP_49307 | TGCTGTTG | CCTGAAG | NM_1420 | Mus musculus similar to baculoviral IAP repeat-containing 6 [Homo sapiens] (L | | | 2.14 | SM2544 | c | | 4 |
| mPool4_V2MM_13345 | -1.07513 | 73.80895 | TRUE | HP_23671 | TGCTGTTG | CAGTTACT | AK172981 | cDNA sequence BC004022 | BC004022 | | 2.15 | SM2588 | h | | 6 |
| mPool4_V2MM_241882 | -1.07461 | 87.58579 | TRUE | HP_49363 | TGCTGTTG | GAGTGTT | XM_1431 | LOC434684 | | | 2.14 | SM2533 | e | | 7 |
| mPool4_V2MM_183062 | -1.07252 | 92.61223 | TRUE | HP_40353 | TGCTGTTG | CAGTTATA | XM_2889 | Mus musculus LOC332601 (LOC332601), mRNA. | | NA | | | | NA | |
| mPool4_V2MM_230934 | -1.07114 | 83.62456 | TRUE | HP_48291 | TGCTGTTG | CAGAGAG | AK035916 | hypothetical protein 9630017O17 | | NA | | | | NA | |
| 092308m3_V2MM_207884 | -1.06961 | 91.71214 | TRUE | HP_42568 | TGCTGTTG | GTTAATAT | XM_48765 | similar to ribosomal protein L6 | | | 2.14 | SM2509 | b | | 11 |
| mPool4_V2MM_195522 | -1.06825 | 83.62456 | TRUE | HP_41366 | TGCTGTTG | GCCATTG | NM_0272 | minichromosome maintenance deficient 10 (S. cerev | Mcm10 | | 2.15 | SM2579 | e | | 2 |
| mPool4_V2MM_80207 | -1.06784 | 94.45693 | TRUE | HP_30184 | TGCTGTTG | GCGTTTAT | NM_0254 | asparagine-linked glycosylation 5 homolog (yeast, d | Alg5 | | 2.15 | SM2584 | g | | 4 |
| mPool4_V2MM_148697 | -1.06743 | 69.72441 | TRUE | HP_36940 | TGCTGTTG | GTTTCTCT | XM_1640 | Mus musculus LOC244398 (LOC244398), mRNA. | | | 2.16 | SM2636 | b | | 3 |
| mPool4_V2MM_154219 | -1.06719 | 61.94057 | TRUE | HP_37486 | TGCTGTTG | GAGATAC | XM_1966 | LOC433292 | | | 2.16 | SM2648 | f | | 12 |
| mPool4_V2MM_125995 | -1.06658 | 91.71214 | TRUE | HP_34683 | TGCTGTTG | CTATCTCT | XM_4878 | similar to apurinic/apyrimidinic endonuclease 2 | | | 2.16 | SM2628 | h | | 7 |
| mPool4_V2MM_26609 | -1.06632 | 84.31789 | TRUE | HP_24964 | TGCTGTTG | CTATTAAC | AF371320 | brain and acute leukemia, cytoplasmic | Baalc | | 2.15 | SM2598 | h | | 8 |
| mPool4_V2MM_120762 | -1.06494 | 61.94057 | TRUE | HP_34165 | TGCTGTTG | CCTGTTCC | XM_1411 | Mus musculus similar to olfactory receptor MOR225-2 [Mus musculus] (LOC22 | | | 2.16 | SM2640 | c | | 7 |
| mPool7_V2MM_192349 | -1.06386 | 2.577834 | TRUE | HP_32504 | TGCTGTTG | GATGCCA | NM_1994 | expressed sequence AA408556 | AA408556 | | 2.1 | SM2315 | g | | 6 |
| mPool4_V2MM_25595 | -1.06384 | 73.80895 | TRUE | HP_24865 | TGCTGTTG | CAGTTGG | XM_1492 | zinc finger protein 217 | Zfp217 | | 2.16 | SM2643 | b | | 7 |
| 092308m3_V2MM_147348 | -1.06319 | 61.94057 | TRUE | HP_36805 | TGCTGTTG | GACTCAC | XM_1628 | Mus musculus LOC208761 (LOC208761), mR. | | | 2.14 | SM2526 | c | | 1 |
| 092308m3_V2MM_84648 | -1.06261 | 91.71214 | TRUE | HP_30615 | TGCTGTTG | CTGACTAT | NM_0288 | elongation protein 3 homolog (S. cerevisiae) | Elp3 | | 2.13 | SM2475 | g | | 9 |
| mPool6_V2MM_63375 | -1.06196 | 94.45693 | TRUE | HP_28545 | TGCTGTTG | CGCTTCCA | AF074881 | histone deacetylase 3 | Hdac3 | | 2.8 | SM2215 | b | | 10 |
| mPool4_V2MM_221317 | -1.06163 | 69.72441 | TRUE | HP_45425 | TGCTGTTG | CTCATCAT | NM_1469 | olfactory receptor 353 | Olfr353 | | 2.15 | SM2578 | a | | 1 |
| mPool4_V2MM_59942 | -1.05986 | 66.95036 | TRUE | HP_28211 | TGCTGTTG | GCTTTCCT | NM_0292 | ring finger protein 121 | Rnf121 | | 2.15 | SM2599 | g | | 9 |
| mPool2_V2MM_7196 | -1.05964 | 3.004939 | TRUE | HP_23071 | TGCTGTTG | GAGCCAA | NM_0298 | B-cell CLL/lymphoma 7A | Bcl7a | | 2.11 | SM2386 | f | | 3 |
| mPool4_V2MM_197644 | -1.05946 | 94.45693 | TRUE | HP_41571 | TGCTGTTG | GACTTACT | XM_1244 | ATP synthase, H+ transporting, mitochondrial F0 cor | Atp5l | | 2.16 | SM2611 | b | | 9 |
| 092308m3_V2MM_49632 | -1.05942 | 94.45693 | TRUE | HP_27207 | TGCTGTTG | GTTACTTC | NM_0169 | toll-like receptor 5 | Tlr5 | | 2.12 | SM2407 | h | | 4 |
| mPool4_V2MM_220109 | -1.05815 | 92.61223 | TRUE | HP_45308 | TGCTGTTG | GGAAATA | NM_1467 | olfactory receptor 491 | Olfr491 | | 2.15 | SM2596 | d | | 2 |
| mPool4_V2MM_168161 | -1.05749 | 79.16495 | TRUE | HP_38866 | TGCTGTTG | CTAGTAG | XM_2859 | Mus musculus similar to hypothetical protein FLJ11159 [Homo sapiens] (LOC33 | | NA | | | | NA | |
| mPool4_V2MM_181834 | -1.0568 | 94.45693 | TRUE | HP_40230 | TGCTGTTG | CCTGTGG | XM_2887 | Mus musculus LOC332186 (LOC332186), mRNA. | | NA | | | | NA | |
| mPool4_V2MM_102126 | -1.05669 | 92.61223 | TRUE | HP_32328 | TGCTGTTG | CACACCAT | XM_1281 | RIKEN cDNA 4930415O20 gene | 4930415O20Rik | | 2.16 | SM2609 | a | | 2 |
| 092308m3_V2MM_29468 | -1.05622 | 61.94057 | TRUE | HP_25243 | TGCTGTTG | CACACCCA | NM_0093 | testicular serine protease 2 | Tesp2 | | 2.12 | SM2433 | a | | 4 |
| mPool4_V2MM_43108 | -1.05483 | 92.96229 | TRUE | HP_26570 | TGCTGTTG | CTACAAA | NM_0292 | RIKEN cDNA 1700001E04 gene | 1700001E04Rik | | 2.16 | SM2604 | h | | 11 |
| 092308m1_V2MM_154088 | -1.05384 | 3.004939 | TRUE | HP_37472 | TGCTGTTG | CGAGTAC | XM_1964 | myosin VI | Myo6 | | 2.6 | SM2123 | c | | 2 |

| | | | | | | | | | | | | | | | |
|---|---|---|---|---|---|---|---|---|---|---|---|---|---|---|---|
| mPool6_V2MM_154811 | -1.05328 | 94.45693 | TRUE | HP_37545 | TGCTGTTG | GACTCCGA | XM_19702 | Mus musculus LOC271072 (LOC271072), mRNA. | | | 2.8 | SM2204 | c | | 12 |
| mPool4_V2MM_31675 | -1.05276 | 69.72441 | TRUE | HP_25457 | TGCTGTTG | CTTGCCAA | NM_0093 | tissue factor pathway inhibitor 2 | Tfpi2 | | 2.15 | SM2572 | g | | 1 |
| mPool4_V2MM_13901 | -1.05234 | 91.10404 | TRUE | HP_23726 | TGCTGTTG | CCCAAGTT | NM_0270 | RIKEN cDNA 1700013G24 gene | 1700013G24Rik | | 2.15 | SM2594 | b | | 6 |
| mPool4_V2MM_136695 | -1.05203 | 79.16495 | TRUE | HP_35741 | TGCTGTTG | GGATCATT | XM_15204 | Mus musculus LOC237418 (LOC237418), mRNA. | | | 2.16 | SM2648 | c | | 10 |
| mPool4_V2MM_122525 | -1.05062 | 94.45693 | TRUE | HP_34339 | TGCTGTTG | CTCCAAAT | XM_48839 | similar to forkhead box R2 | | | 2.11 | SM2358 | f | | 2 |
| mPool6_V2MM_192732 | -1.0502 | 81.56557 | TRUE | HP_27534 | TGCTGTTG | GGATTAAC | BC052474 | mitogen-activated protein kinase kinase kinase kinase 6 | | | 2.7 | SM2184 | g | | 9 |
| mPool4_V2MM_111247 | -1.04982 | 94.45693 | TRUE | HP_33226 | TGCTGTTG | GGCTGAAA | AK015052 | RIKEN cDNA 4930402K13 gene | 4930402K13Rik | | 2.16 | SM2615 | c | | 1 |
| 092308m3_V2MM_197152 | -1.0492 | 92.96229 | TRUE | HP_41523 | TGCTGTTG | CTCAATAC | NM_0289 | zinc finger protein 336 | Zfp336 | | 2.13 | SM2484 | d | | 9 |
| mPool2_V2MM_119312 | -1.04901 | 3.004939 | TRUE | HP_34023 | TGCTGTTG | CATTTACA | XM_14034 | similar to RIKEN cDNA 5430401O09 gene | | | 2.11 | SM2352 | e | | 12 |
| mPool4_V2MM_128042 | -1.04832 | 87.58579 | TRUE | HP_34886 | TGCTGTTG | GGGACATC | XM_1455 | Mus musculus similar to hypothetical protein MGC45586 [Homo sapiens] (LOC | | | 2.14 | SM2538 | e | | 4 |
| 092308m3_V2MM_50233 | -1.04822 | 83.62456 | TRUE | HP_27266 | TGCTGTTG | CAGACGGT | NM_0290 | serine caroboxypeptidase 1 | Scpep1 | | 2.12 | SM2408 | d | | 5 |
| mPool4_V2MM_262409 | -1.04715 | 92.61223 | TRUE | HP_52461 | TGCTGTTG | CTGGGTAC | NM_1984 | RIKEN cDNA C030039L03 gene | C030039L03Rik | | 2.16 | SM2605 | a | | 11 |
| mPool6_V2MM_67834 | -1.04654 | 92.61223 | TRUE | HP_28981 | TGCTGTTG | CTCCTATA | NM_0294 | zinc finger protein 393 | Zfp393 | | 2.8 | SM2225 | h | | 6 |
| mPool6_V2MM_68015 | -1.04641 | 84.31789 | TRUE | HP_28998 | TGCTGTTG | CTGATCAA | NM_0074 | adenylate cyclase 7 | Adcy7 | | 2.8 | SM2220 | g | | 11 |
| mPool7_V2MM_206354 | -1.04543 | 2.577834 | TRUE | HP_42418 | TGCTGTTG | CCTGGATC | XM_1365 | Mus musculus similar to Glyceraldehyde 3-phosphate dehydrogenase (GAPDH) | | | 2.1 | SM2303 | h | | 12 |
| 092308m1_V2MM_79639 | -1.04447 | 3.004939 | TRUE | HP_30129 | TGCTGTTG | GCCAATCA | AK085850 | RIKEN cD D830016O14 gene | D830016O14Rik | | 2.3 | SM2001 | b | | 11 |
| mPool4_V2MM_112142 | -1.04307 | 91.10404 | TRUE | HP_33314 | TGCTGTTG | CTCTTGAA | XM_13634 | Mus musculus similar to Exocyst complex component Sec8 (LOC240873), mRN | | | 2.16 | SM2624 | b | | 7 |
| mPool4_V2MM_160582 | -1.04191 | 61.94057 | TRUE | HP_38116 | TGCTGTTG | CACCTTCT | XM_28357 | Mus musculus hypothetical gene supported by AK083061 (LOC329079), mRNA. | | | 2.16 | SM2650 | f | | 12 |
| mPool6_V2MM_44133 | -1.04115 | 92.61223 | TRUE | HP_26669 | TGCTGTTG | CTTACATC | NM_1773 | calcium/calmodulin-dependent protein kinase ID | Camk1d | | 2.7 | SM2185 | c | | 4 |
| mPool4_V2MM_13935 | -1.04105 | 83.62456 | TRUE | HP_23729 | TGCTGTTG | CCCACTCT | BC034893 | RIKEN cDNA 3110037K17 gene | 3110037K17Rik | | 2.15 | SM2586 | a | | 2 |
| mPool4_V2MM_41094 | -1.0408 | 92.61223 | TRUE | HP_26373 | TGCTGTTG | CAAAGATC | NM_1392 | defensin beta 7 | Defb7 | | 2.15 | SM2596 | b | | 3 |
| 092308m3_V2MM_195103 | -1.03972 | 75.98381 | TRUE | HP_41325 | TGCTGTTG | GAAAGGA | NM_1531 | supervillin | Svil | | 2.12 | SM2450 | e | | 6 |
| mPool4_V2MM_238407 | -1.03944 | 94.45693 | TRUE | HP_49024 | TGCTGTTG | CCTCCTTA | XM_1466 | Mus musculus similar to protease [Mus musculus] (LOC244731), mRNA. | | | 2.14 | SM2537 | e | | 12 |
| 092308m3_V2MM_195127 | -1.03931 | 61.94057 | TRUE | HP_41327 | TGCTGTTG | CCACAATC | NM_1776 | hypothetical protein A030013D21 | | | 2.13 | SM2451 | b | | 6 |
| mPool4_V2MM_38269 | -1.0392 | 62.64426 | TRUE | HP_26099 | TGCTGTTG | GCAAATAT | NM_0276 | spermatogenesis associated 1 | Spata1 | | 2.16 | SM2603 | f | | 1 |
| mPool4_V2MM_169478 | -1.03746 | 92.61223 | TRUE | HP_38997 | TGCTGTTG | CTCTTCGA | XM_2862 | Mus musculus hypothetical gene supported by AK038490 (LOC327786), mRNA. | | NA | | | | NA | |
| mPool4_V2MM_126910 | -1.03708 | 94.45693 | TRUE | HP_34774 | TGCTGTTG | GACCTCCT | XM_14487 | similar to Glyceraldehyde 3-phosphate dehydrogenase, liver (GAPDH) | | | 2.1 | SM2329 | c | | 1 |
| mPool4_V2MM_131566 | -1.03681 | 94.45693 | TRUE | HP_35232 | TGCTGTTG | CAGACCTT | XM_14771 | gene model 528, (NCBI) | Gm528 | | 2.16 | SM2632 | d | | 10 |
| mPool4_V2MM_170226 | -1.03646 | 84.31789 | TRUE | HP_39072 | TGCTGTTG | GTCATCTG | XM_28641 | hypothetical LOC328082 | | NA | | | | NA | |
| mPool5_V2MM_6667 | -1.03628 | 92.61223 | TRUE | NA | NA | NA | NA | NA | NA | NA | | NA | NA | NA | |
| 092308m3_V2MM_60584 | -1.03603 | 79.16495 | TRUE | HP_28273 | TGCTGTTG | GGTCCTCT | XM_48489 | similar to Diacylglycerol kise, delta (Diglyceride kise) (DGK-delta) (DAG kise delt | | | 2.12 | SM2446 | f | | 12 |
| mPool4_V2MM_195190 | -1.03532 | 84.31789 | TRUE | HP_41333 | TGCTGTTG | CGACCAAC | NM_1464 | olfactory receptor 1288 | Olfr1288 | | 2.15 | SM2582 | f | | 6 |
| mPool6_V2MM_9789 | -1.03397 | 92.61223 | TRUE | HP_23323 | TGCTGTTG | CATTTGCA | NM_0077 | conserved helix-loop-helix ubiquitous kinase | Chuk | | 2.7 | SM2185 | f | | 11 |
| mPool4_V2MM_192555 | -1.03354 | 83.62456 | TRUE | HP_24925 | TGCTGTTG | CCTATATA | NM_1449 | SWI/SNF related, matrix associated, actin dependen | Smarca3 | | 2.16 | SM2602 | h | | 11 |
| 092308m1_V2MM_70649 | -1.03042 | 3.004939 | TRUE | HP_29255 | TGCTGTTG | CACTCCAC | AF152344 | prostaglandin F2 receptor negative regulator | Ptgfrn | | 2.3 | SM2010 | e | | 7 |
| mPool4_V2MM_61507 | -1.03027 | 70.87591 | TRUE | HP_28364 | TGCTGTTG | GTTATCCT | NM_0238 | RAB3C, member RAS oncogene family | Rab3c | | 2.15 | SM2571 | h | | 3 |
| 092308m3_V2MM_231358 | -1.02993 | 94.45693 | TRUE | HP_48333 | TGCTGTTG | CGCAAAGC | XM_12844 | RIKEN cD 4930432O21 gene | 4930432O21Rik | | 2.13 | SM2487 | e | | 3 |
| mPool4_V2MM_124135 | -1.02986 | 94.45693 | TRUE | HP_34498 | TGCTGTTG | CCAAATAT | XM_14331 | Mus musculus similar to peptidoglycan recognition protein-I-beta precursor (L | | | 2.1 | SM2347 | b | | 11 |
| 092308m3_V2MM_194412 | -1.02963 | 94.45693 | TRUE | HP_41257 | TGCTGTTG | CGGGTAAT | NM_1777 | zinc finger protein 31 | Zfp31 | | 2.13 | SM2451 | g | | 12 |
| mPool6_V2MM_36475 | -1.02961 | 92.96229 | TRUE | HP_25925 | TGCTGTTG | CTGTATTC | NM_0087 | p21 (CDKN1A)-activated kinase 3 | Pak3 | | 2.7 | SM2186 | d | | 9 |
| mPool4_V2MM_153946 | -1.02938 | 83.62456 | TRUE | HP_37458 | TGCTGTTG | CCTCCTGG | XM_19641 | gene model 682, (NCBI) | Gm682 | NA | | | | NA | |
| mPool6_V2MM_161946 | -1.02931 | 91.10404 | TRUE | HP_38251 | TGCTGTTG | GAATTTGA | XM_28412 | Mus musculus hypothetical gene supported by AK036030 (LOC330162), mRNA. | | | 2.7 | SM2192 | c | | 6 |
| 092308m1_V2MM_66341 | -1.02841 | 3.004939 | TRUE | HP_28835 | TGCTGTTG | CACGCCAA | NM_0263 | processing of precursor 1, ribonuclease P/MRP famil | Pop1 | | 2.4 | SM2024 | g | | 1 |
| mPool6_V2MM_53756 | -1.02767 | 83.62456 | TRUE | HP_27610 | TGCTGTTG | CACATCCG | NM_1772 | cyclin-dependent kinase-like 2 (CDC2-related kinase) | Cdkl2 | | 2.7 | SM2184 | h | | 1 |
| mPool4_V2MM_109969 | -1.02765 | 92.61223 | TRUE | HP_33101 | TGCTGTTG | GGCTCCTC | XM_13425 | RIKEN cDNA 9430098E02 gene | 9430098E02Rik | | 2.16 | SM2625 | h | | 7 |
| mPool4_V2MM_5400 | -1.0271 | 61.94057 | TRUE | HP_22895 | TGCTGTTG | CCCATCAA | NM_1535 | Rho-related BTB domain containing 2 | Rhobtb2 | | 2.15 | SM2591 | d | | 2 |
| 092308m3_V2MM_225134 | -1.02703 | 83.62456 | TRUE | HP_47718 | TGCTGTTG | GATTGGGA | XM_10934 | similar to ribosomal protein | | | 2.13 | SM2458 | c | | 8 |
| 092308m3_V2MM_31917 | -1.02448 | 61.94057 | TRUE | HP_25480 | TGCTGTTG | GAGCTAAC | NM_0197 | SEC23B (S. cerevisiae) | Sec23b | | 2.12 | SM2426 | a | | 12 |
| mPool4_V2MM_211180 | -1.02407 | 92.96229 | TRUE | HP_33224 | TGCTGTTG | CTCAAGAA | XM_14292 | hypothetical LOC333463 | | | 2.14 | SM2531 | b | | 2 |
| mPool4_V2MM_125567 | -1.02375 | 81.56557 | TRUE | HP_34641 | TGCTGTTG | GGACAGCA | XM_14409 | Mus musculus similar to CG17097-PB [Drosophila melanogaster] (LOC230826), | | | 2.16 | SM2641 | c | | 3 |
| mPool4_V2MM_177396 | -1.02346 | 73.80895 | TRUE | HP_39787 | TGCTGTTG | GCCATAGC | AK054405 | hypothetical protein E330022O07 | | NA | | | | NA | |

| | | | | | | | | | | | | | |
|---|---|---|---|---|---|---|---|---|---|---|---|---|---|
| 092308m3_V2MM_29857 | -1.02344 | 91.71214 | TRUE | HP_17809 | TGCTGTTG | CAGCAATT | NM_1448 | COX15 homolog, cytochrome c oxidase assembly pro | Cox15 | 2.11 | SM2399 | h | 8 |
| mPool6_V2MM_40359 | -1.02207 | 61.94057 | TRUE | HP_26302 | TGCTGTTG | GTCCCACA | XM_13292 | guanylate cyclase 2c | Gucy2c | 2.7 | SM2182 | a | 5 |
| 092308m3_V2MM_206944 | -1.02203 | 70.87591 | TRUE | HP_42476 | TGCTGTTG | CATGTGG | XM_1538 | Mus musculus LOC238472 (LOC238472), mR. | | 2.14 | SM2526 | d | 6 |
| 092308m3_V2MM_232487 | -1.02047 | 94.45693 | TRUE | HP_48443 | TGCTGTTG | GCAATTCT | XM_1371 | similar to hypothetical protein MGC37588 | | 2.14 | SM2508 | g | 2 |
| mPool4_V2MM_128653 | -1.019 | 94.45693 | TRUE | HP_34946 | TGCTGTTG | GTCTTTAC | XM_1458 | similar to tripartite motif protein TRIM30 isoform alpha | | 2.14 | SM2545 | d | 8 |
| mPool4_V2MM_118802 | -1.0182 | 91.71214 | TRUE | HP_33973 | TGCTGTTG | GCCTGACT | XM_14007 | Mus musculus similar to transcription factor IIIA [Mus musculus] (LOC240124), | | 2.16 | SM2643 | b | 6 |
| mPool6_V2MM_153186 | -1.01776 | 94.45693 | TRUE | HP_37384 | TGCTGTTG | GACTGAGT | AK029407 | RIKEN cDNA 4833431D13 gene | 4833431D13Rik | 2.7 | SM2196 | a | 12 |
| mPool4_V2MM_15891 | -1.01651 | 83.62456 | TRUE | HP_23920 | TGCTGTTG | CTGGGTCA | NM_0188 | mesothelin | Msln | 2.15 | SM2573 | h | 9 |
| 092308m3_V2MM_53619 | -1.01644 | 61.94057 | TRUE | HP_27597 | TGCTGTTG | CAAGGGCA | NM_1725 | transcriptiol adaptor 2 (ADA2 homolog, yeast)-like | Tada2l | 2.12 | SM2437 | a | 5 |
| 092308m3_V2MM_116730 | -1.01572 | 92.61223 | TRUE | HP_33768 | TGCTGTTG | CATGCTAT | XM_48733 | similar to granzyme H splice variant 2 | | 2.14 | SM2501 | h | 5 |
| mPool6_V2MM_66315 | -1.01454 | 84.31789 | TRUE | HP_28833 | TGCTGTTG | CACCCTTC | NM_0080 | FYVE, RhoGEF and PH domain containing 1 | Fgd1 | 2.7 | SM2176 | h | 8 |
| 092308m3_V2MM_111537 | -1.01444 | 94.45693 | TRUE | HP_33255 | TGCTGTTG | CAGTTCCA | XM_13591 | cD sequence BC023829 | BC023829 | 2.14 | SM2503 | b | 1 |
| mPool4_V2MM_97336 | -1.01443 | 84.31789 | TRUE | HP_31857 | TGCTGTTG | CCTTACAT | XM_1114 | Mus musculus similar to vomeronasal receptor V1RH19 [Mus musculus] (LOC1 | | 2.16 | SM2617 | h | 11 |
| 092308m1_V2MM_151088 | -1.0129 | 3.004939 | TRUE | HP_37176 | TGCTGTTG | CCCACAG | NM_0274 | cD sequence BC033915 | BC033915 | 2.4 | SM2030 | e | 3 |
| mPool6_V2MM_69142 | -1.0106 | 66.95036 | TRUE | HP_29108 | TGCTGTTG | GCAATTAT | NM_1728 | beta galactoside alpha 2,6 sialyltransferase 2 | St6gal2 | 2.8 | SM2221 | a | 12 |
| mPool6_V2MM_67102 | -1.01006 | 75.98381 | TRUE | HP_12245 | TGCTGTTG | CCCTGGA | NM_0080 | gamma-aminobutyric acid (GABA-A) receptor, subur | Gabra3 | 2.8 | SM2220 | a | 3 |
| mPool4_V2MM_107883 | -1.00821 | 84.31789 | TRUE | HP_32896 | TGCTGTTG | CCACTCAG | XM_1323 | RIKEN cDNA 1700016K13 gene | 1700016K13Rik | 2.16 | SM2624 | g | 12 |
| mPool4_V2MM_39325 | -1.00645 | 69.72441 | TRUE | HP_26201 | TGCTGTTG | GCTTCTAC | NM_1342 | vomeronasal 1 receptor, G11 | V1rg11 | 2.15 | SM2579 | c | 7 |
| 092308m3_V2MM_140236 | -1.006 | 61.94057 | TRUE | HP_36095 | TGCTGTTG | GTGTTGCT | XM_1558 | Mus musculus LOC239484 (LOC239484), mR. | | 2.14 | SM2520 | a | 2 |
| 092308m3_V2MM_36248 | -1.00558 | 73.80895 | TRUE | HP_25902 | TGCTGTTG | CTGCCTAA | NM_0241 | YY1 associated factor 2 | Yaf2 | 2.11 | SM2396 | g | 8 |
| mPool4_V2MM_122598 | -1.00516 | 94.45693 | TRUE | HP_34346 | TGCTGTTG | GTCCTTAG | XM_1422 | Mus musculus similar to differentiation inhibitor protein Id2A - human (LOC23 | | 2.14 | SM2548 | g | 2 |
| mPool6_V2MM_222672 | -1.00483 | 94.45693 | TRUE | HP_47491 | TGCTGTTG | GCCTGTAC | XM_2858 | similar to zinc finger, MYND domain containing 12 | | 2.8 | SM2209 | c | 10 |
| 092308m3_V2MM_97963 | -1.00454 | 73.80895 | TRUE | HP_31919 | TGCTGTTG | CCAATTTC | XM_1119 | Mus musculus similar to intracisterl A particles [Mus musculus] (LOC194861), n | | 2.13 | SM2484 | c | 8 |
| mPool4_V2MM_13117 | -1.00387 | 94.45693 | TRUE | HP_23649 | TGCTGTTG | CAGCTCTC | NM_1531 | acetyl-Coenzyme A acetyltransferase 3 | Acat3 | 2.15 | SM2600 | g | 7 |
| mPool6_V2MM_163964 | -1.00381 | 79.16495 | TRUE | HP_38451 | TGCTGTTG | CAGGTAG | XM_2848 | Mus musculus similar to eukaryotic translation initiation factor 3, subunit 6 48k | | 2.7 | SM2191 | g | 11 |
| mPool4_V2MM_14535 | -1.00371 | 94.45693 | TRUE | HP_23788 | TGCTGTTG | CCTGGAG | NM_0084 | kinesin family member 3A | Kif3a | 2.15 | SM2574 | g | 4 |
| 092308m3_V2MM_106066 | -1.00351 | 92.96229 | TRUE | HP_32717 | TGCTGTTG | GTCATCTC | XM_35541 | gene model 1015, (NCBI) | Gm1015 | 2.14 | SM2507 | a | 10 |
| 092308m3_V2MM_227516 | -1.00222 | 92.61223 | TRUE | HP_47951 | TGCTGTTG | CTGGGAA | NM_1819 | RAS-like, estrogen-regulated, growth-inhibitor | Rerg | 2.13 | SM2482 | d | 12 |
| mPool4_V2MM_165848 | -1.00215 | 91.10404 | TRUE | HP_38638 | TGCTGTTG | CACAGGA | XM_2853 | plakophilin 4 | Pkp4 | 2.16 | SM2649 | a | 3 |
| mPool4_V2MM_212070 | -1.00156 | 94.45693 | TRUE | HP_42974 | TGCTGTTG | GAGACAG | XM_1488 | armadillo repeat containing 4 | Armc4 | 2.16 | SM2647 | c | 4 |
| mPool6_V2MM_160135 | -1.00145 | 94.45693 | TRUE | HP_38072 | TGCTGTTG | GCAACTTA | XM_2833 | Mus musculus hypothetical gene supported by AK029312 (LOC328736), mRNA. | | 2.7 | SM2200 | d | 7 |
| mPool4_V2MM_199657 | -1.00126 | 61.94057 | TRUE | HP_41769 | TGCTGTTG | GGCCCAA | XM_2885 | Mus musculus LOC331931 (LOC331931), mRNA. | | NA | | | NA |
| mPool2_V2MM_98049 | -1.00081 | 2.577834 | TRUE | HP_31928 | TGCTGTTG | CAGAATGA | XM_1120 | Mus musculus similar to RIKEN cDNA 2610028L19 (LOC195712), mRNA. | | 2.6 | SM2141 | h | 7 |
